# Development of Potent G Protein Pathway-Biased GPR183 Agonists

**DOI:** 10.64898/2026.07.27.740726

**Authors:** Kaustubh R. Bhuskute, Asmita Manandhar, Viktoria M. S. Kjær, Francesco Casartelli, Gertrud M. Hjortø, Maria Ioanna Koutsaki, Udhayabhaskar Sathyanarayanan, Emilia Hjortkilde, Rita Turcio, Mette M. Rosenkilde, Trond Ulven, Elisabeth Rexen Ulven

## Abstract

GPR183 is an oxysterol-sensing GPCR predominantly expressed in lymphoid organs and tissues. Activation of the receptor by oxysterol 7α,25-OHC leads to Gα_i_ protein-mediated signaling as well as β-arrestin2 recruitment. GPR183/oxysterol signaling modulates localization of lymphoid cells, consequently the receptor is associated with several inflammation-associated diseases and is an interesting potential drug target. Previously, we reported the discovery of moderately potent G protein-biased partial agonists for GPR183 from a virtual screening based on the scaffold of the antagonist NIBR189. Herein, we present the detailed structure-activity investigations and optimizations, which led to the identification of full agonists for GPR183 with complete bias for Gα_i_ protein signaling and low nanomolar potency, including **63** (TUG-2604) with potency and efficacy similar to 7α,25-OHC. Notably, 63 was unable to induce migration of human dendritic cells but inhibited migration induced by 7α,25-OHC. This compound will be valuable for further explorations of the signaling-specific function and drug target potential of GPR183.

## Introduction

GPR183, also known as Epstein-Barr Virus induced gene 2 (EBI2), is a chemotactic G protein-coupled receptor.^1,2^ Oxysterols, formed endogenously by oxidation of cholesterols, act as agonists for the receptor, with 7α,25-dihydroxycholesterol (7α,25-OHC) being the most potent of these.^3,4^ GPR183 signals via Gα_i_ protein and recruits β-arrestin2, and 7α,25-OHC acts as a balanced agonist of both pathways.^3,5–7^ The receptor is primarily expressed in lymphoid tissue and immune cells, including peripheral blood mononuclear cells (PBMCs), macrophages, and dendritic cells, and is central in immunometabolism, and the gut immune system.^7–10^ The receptor is also expressed in the CNS in astrocytes, oligodendrocytes, and microglia.^11–13^ GPR183 signaling induced by 7α,25-OHC plays an important role in modulating innate and adaptive immunity by guiding the localization of lymphoid cells *in vivo*.^14–17^ Both GPR183 agonists and antagonists are of therapeutic interest. Recently, GPR183 antagonists were shown to be a potential therapeutic approach for the treatment of rheumatoid arthritis,^18^ neuropathic pain,^19,20^ viral infections,^21^ and inflammatory bowel disease,^22–25^ whereas GPR183 agonists have been suggested as therapeutic strategies for diseases such as tuberculosis,^26,27^ multiple sclerosis,^11,28,29^ and systemic lupus erythematosus.^30,31^

Both agonists and antagonists of GPR183 have been reported. The inverse agonist and antagonist GSK682753A, discovered through screening independently of the oxysterol agonists, was the first ligand reported for GPR183 (Chart 1).^25,32^ Gessier *et al.* later identified a low potency agonist NIBR51 and the antagonist NIBR127.^33^ Subsequent optimization of NIBR127 led to the potent and selective competitive GPR183 antagonist NIBR189.^33^ Recently, further optimization of NIBR189 has led to the discovery of additional GPR183 antagonists, including Cmpd 32 and Cmpd 33, which have displayed efficacy in murine models of collagen-induced arthritis and dextran sodium sulfate (DSS)-induced experimental colitis, respectively.^18,24^ The conserved GPCR sodium binding site has also been identified in GPR183 and shown to exert negative allosteric modulation of 7α,25-OHC-mediated activation.^34^

We previously conducted a virtual screening based on the piperazine diamide scaffold of NIBR189 that led to the discovery of Cmpd 15 and Cmpd 16, and subsequent exploration gave TUG-2201 and TUG-2202, which were found to be G protein-biased partial agonists for GPR183.^35^ This study provided preliminary SAR insights, indicating that the eastern part of the molecule was responsible for the switch from antagonism to partial agonism, however the structural information was limited due to a small number of compounds. Herein, we report further SAR explorations and optimizations leading to potent GPR183 full agonists with complete Gα_i_-biased signaling.

**Chart 1.**
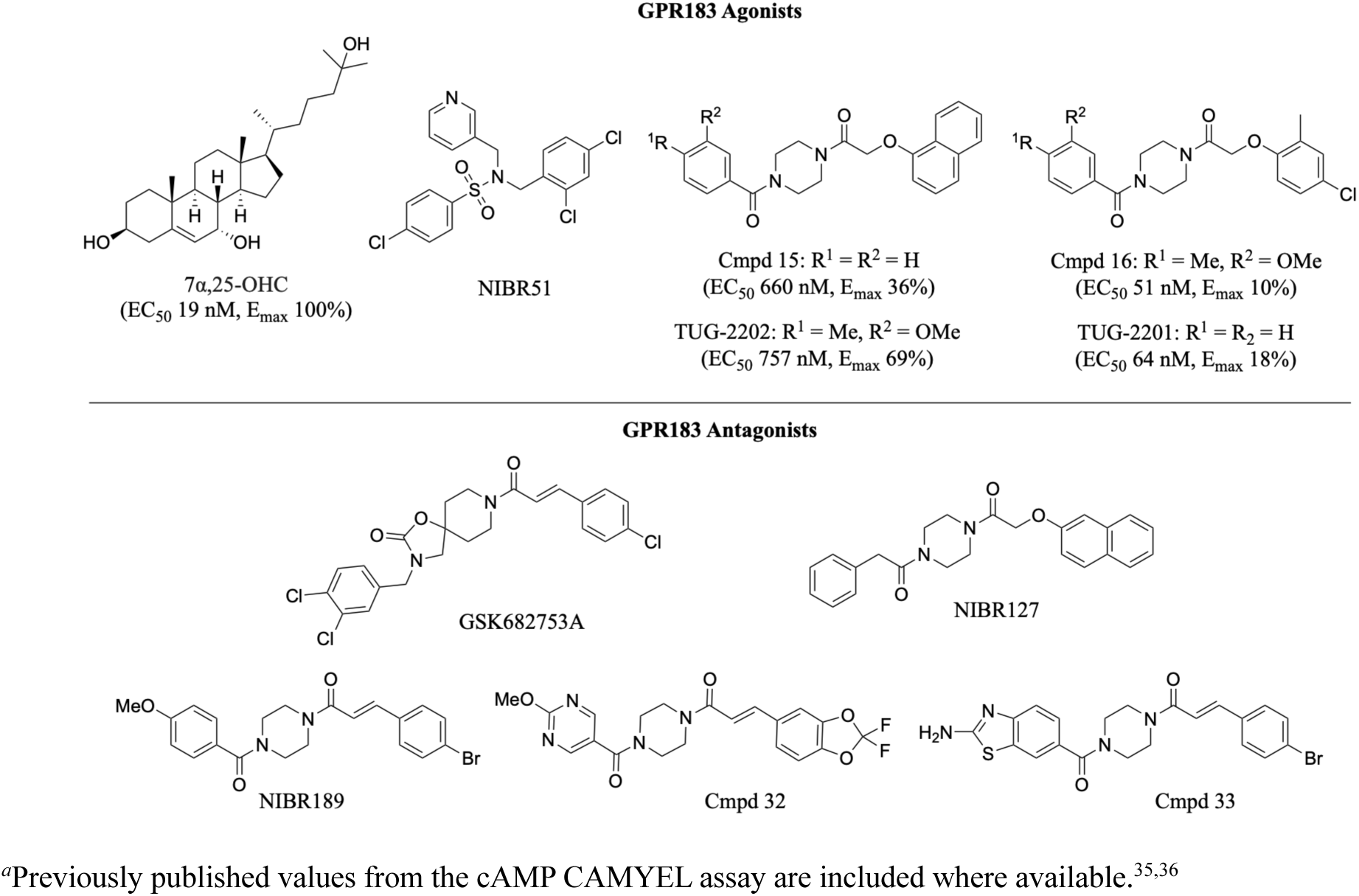
Structures of GPR183 relevant ligands*^a^*.

## Results and discussion

### SAR exploration

GPR183 is predominantly Gα_i_-coupled, leading to inhibition of adenylyl cyclase and reduced cAMP accumulation.^7^ The activity of all compounds were tested with the cAMP sensor using YFP-EPAC-Rluc (CAMYEL) BRET biosensor in CHO-K1 cells transiently transfected with the human GPR183, as described previously.^35^ In this assay, the most potent endogenous agonist 7α,25-OHC displayed pEC_50_ 8.39 ± 0.05. CHO cells have shown a clean response with low constitutive activity compared e.g. to HEK293 cells, potentially caused by the presence of oxysterols in the medium of the latter cell type.^37^

Previously, we reported the observation that the substitution pattern on the eastern phenoxyacetamide part of ligands such as NIBR127 appeared to direct functional activity so that turning to 1-naphthyl (Cmpd 15, TUG-2202) or replacing with an *ortho/para*-disubstituted phenyl (Cmpd 16, TUG-2201) produced partial GPR183 agonists (Chart 1), whereas the corresponding unsubstituted phenoxy analogues were inactive.^35^ To further explore the dependence of functional activity on the structure of the eastern part, we synthesized mono-substituted analogues of TUG-2201. The *ortho*-methylated **1** was found to retain partial agonism albeit with lower potency, whereas **2** was inactive (Table 1). Furthermore, the 2-naphthyl analogue **3** was devoid of agonistic activity, in contrast to the 1-naphthyl analogue Cmpd 15. These results supported our previous conclusions and pointed at the *ortho-*substitution as the hot spot for control of functional activity.

**Table 1.**
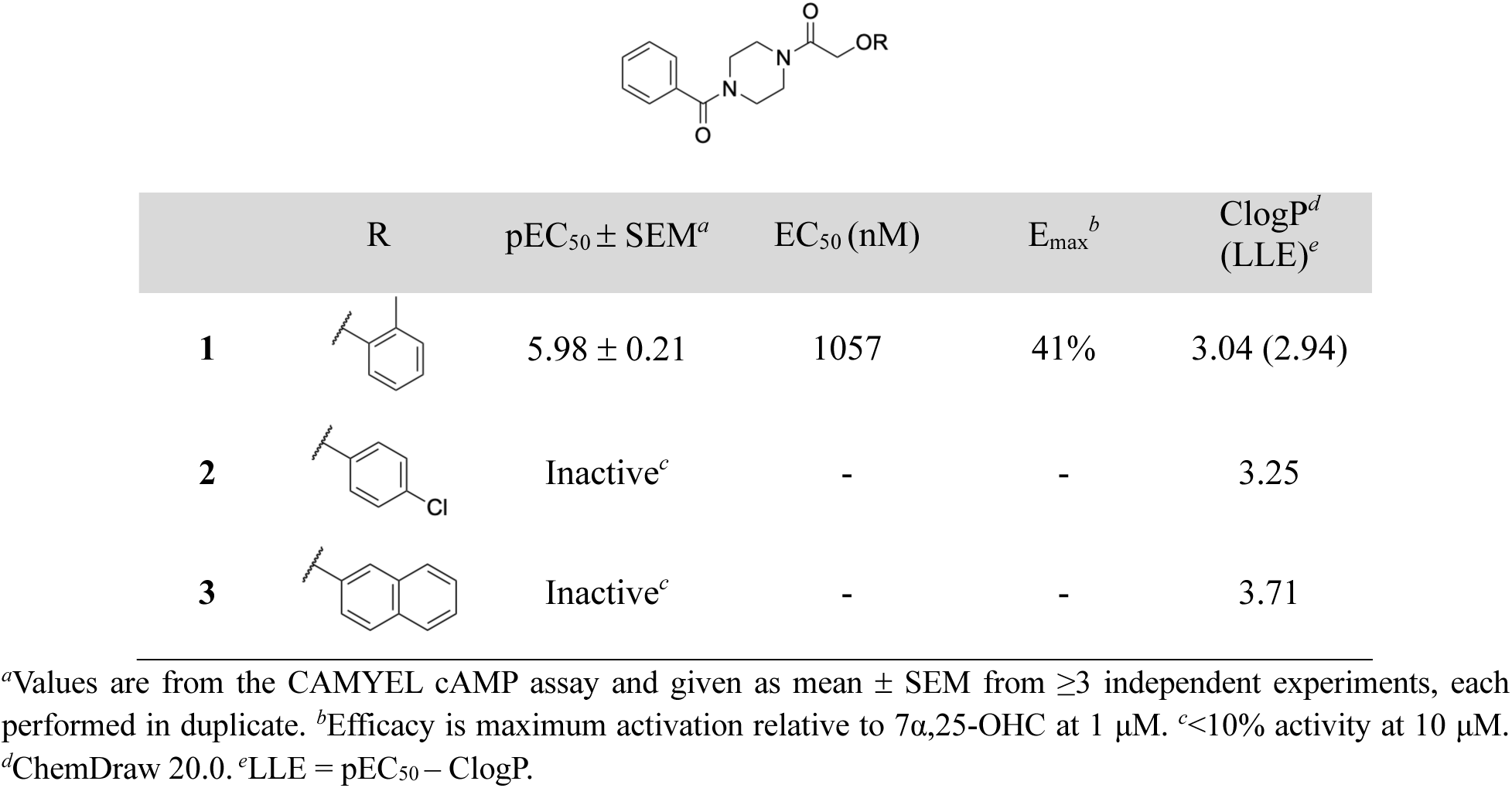
Exploration of the Eastern Part.

|  | R | pEC<sub>50</sub> ± SEM<sup>a</sup> | EC<sub>50</sub> (nM) | E<sub>max</sub><sup>b</sup> | ClogP<sup>d</sup><br>(LLE)<sup>e</sup> |
| --- | --- | --- | --- | --- | --- |
| **1** |  | 5.98 ± 0.21 | 1057 | 41% | 3.04 (2.94) |
| **2** |  | Inactive<sup>c</sup> | - | - | 3.25 |
| **3** |  | Inactive<sup>c</sup> | - | - | 3.71 |

<sup>a</sup>Values are from the CAMYEL cAMP assay and given as mean ± SEM from ≥3 independent experiments, each performed in duplicate. <sup>b</sup>Efficacy is maximum activation relative to 7α,25-OHC at 1 μM. <sup>c</sup><10% activity at 10 μM.
<sup>d</sup>ChemDraw 20.0. <sup>e</sup>LLE = pEC<sub>50</sub> – ClogP.

The 1-naphthoxyacetamide eastern part gave the most interesting combination of high partial agonism and good potency. We therefore retained this part and turned attention to the western part, initially exploring substituted benzamides. *para-*Methoxybenzoyl **4** displayed slightly improved potency (pEC_50_ = 6.43) and efficacy of 68% compared to Cmpd 15 (Table 2). In contrast, the more hydrophilic *ortho,meta-*dimethoxybenzoyl **5** lost almost 10-fold potency compared to **4**. Interestingly, replacing the phenyl ring with benzodioxane **6** was detrimental to agonistic activity whereas benzodioxole **7** retained the potency and efficacy, indicating sensitivity to steric hindrance in this part. Insertion of various linear and cyclic spacers in between the piperazine diamide core and the phenyl ring (**8-16**) led to partial agonism with pEC_50_ 6.1-7.2, with the *gem*-cyclopentane spacer (**10**) as the most potent with pEC_50_ 7.23 and an efficacy of 42%. The carbamide analogue **17** was inactive, which could be due to increased polarity in combination with rotational constraints, whereas ketones **18** and **19** were well-tolerated. Overall, the binding pocket occupied by the western benzene ring seems to be flexible in accommodating different substitution patterns and spacers of various chain length and bulkiness.

**Table 2.**
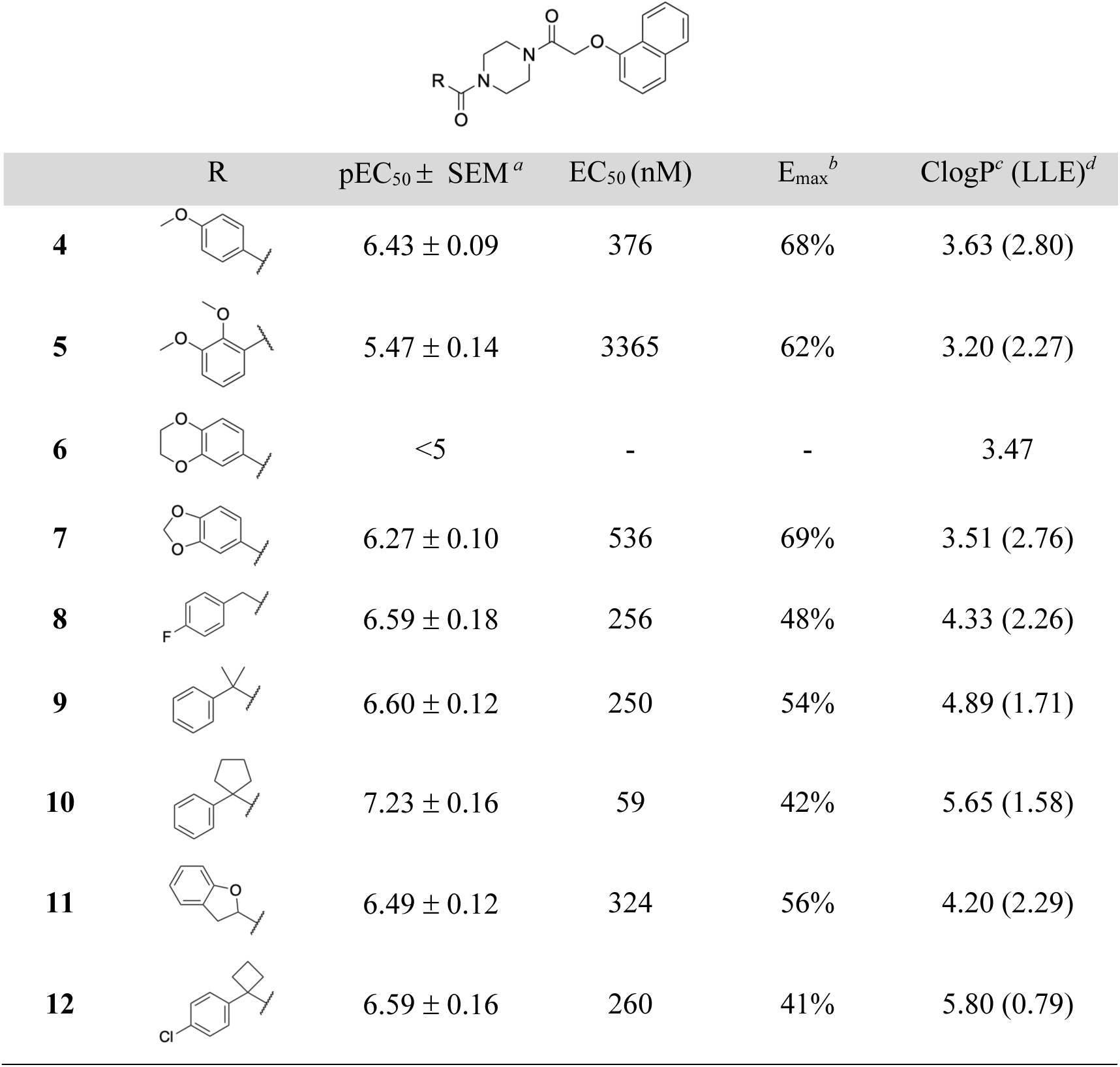
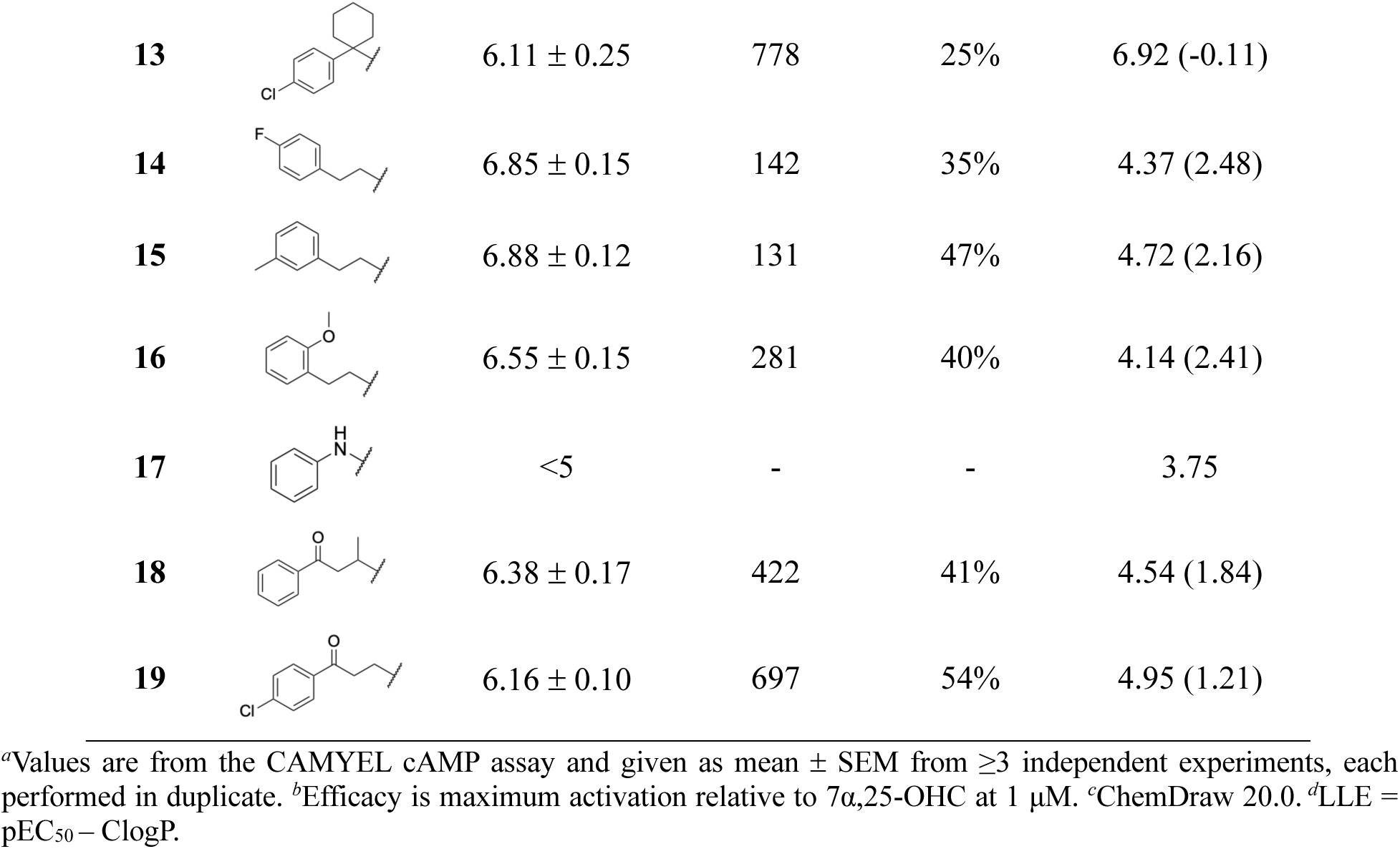
Variations of the Western Part.

|  | R | pEC<sub>50</sub> ± SEM<sup>a</sup> | EC<sub>50</sub> (nM) | E<sub>max</sub><sup>b</sup> | ClogP<sup>c</sup> (LLE)<sup>d</sup> |
| --- | --- | --- | --- | --- | --- |
| **4** |  | 6.43 ± 0.09 | 376 | 68% | 3.63 (2.80) |
| **5** |  | 5.47 ± 0.14 | 3365 | 62% | 3.20 (2.27) |
| **6** |  | <5 | - | - | 3.47 |
| **7** |  | 6.27 ± 0.10 | 536 | 69% | 3.51 (2.76) |
| **8** |  | 6.59 ± 0.18 | 256 | 48% | 4.33 (2.26) |
| **9** |  | 6.60 ± 0.12 | 250 | 54% | 4.89 (1.71) |
| **10** |  | 7.23 ± 0.16 | 59 | 42% | 5.65 (1.58) |
| **11** |  | 6.49 ± 0.12 | 324 | 56% | 4.20 (2.29) |
| **12** |  | 6.59 ± 0.16 | 260 | 41% | 5.80 (0.79) |
| 13 | | $6.11 \pm 0.25$ | 778 | 25% | 6.92 (-0.11) |
| 14 | | $6.85 \pm 0.15$ | 142 | 35% | 4.37 (2.48) |
| 15 | | $6.88 \pm 0.12$ | 131 | 47% | 4.72 (2.16) |
| 16 | | $6.55 \pm 0.15$ | 281 | 40% | 4.14 (2.41) |
| 17 |  | <5 | - | - | 3.75 |
| 18 | | $6.38 \pm 0.17$ | 422 | 41% | 4.54 (1.84) |
| 19 | | $6.16 \pm 0.10$ | 697 | 54% | 4.95 (1.21) |

<sup>a</sup>Values are from the CAMYEL cAMP assay and given as mean ± SEM from $\geq 3$ independent experiments, each performed in duplicate. <sup>b</sup>Efficacy is maximum activation relative to 7α ,25-OHC at 1 μM. <sup>c</sup>ChemDraw 20.0. <sup>d</sup>LLE = pEC<sub>50</sub> – ClogP.

We then went on to explore non-aromatic western parts of various lengths, bulkiness, and polarity. The short acetamide **20** was inactive and elongation to butyramide **21** showed poor activity (Table 3). Unsaturation of the butyl chain (**22**) led to a small increase in potency, whereas insertion of an oxygen ether of the same length (**23**) abolished agonistic activity. Compounds bearing bulkier, branched alkyl groups (**24-27**), terminal cyclopentyl **28** and hexanoyl **29** displayed partial agonistic activity with micromolar potency. Addition of steric bulk improved the potency without significantly altering the efficacy as both **27** and **28** were more potent compared to **24**, and steric bulk in the alpha-position (**26** (TUG-2292)) led to a further increase in potency. Introduction of the very bulky adamantyl **31** led to a super agonist with an efficacy of 123% but with a reduced EC_50_ of 2 μM, whereas the even more bulky dimethyl-adamantyl **32** was 10-fold more potent but only a partial agonist. These results indicate that the western part also has the potential to regulate efficacy in a complex fashion, perhaps by affecting the positioning of the eastern part in the binding.

**Table 3.**
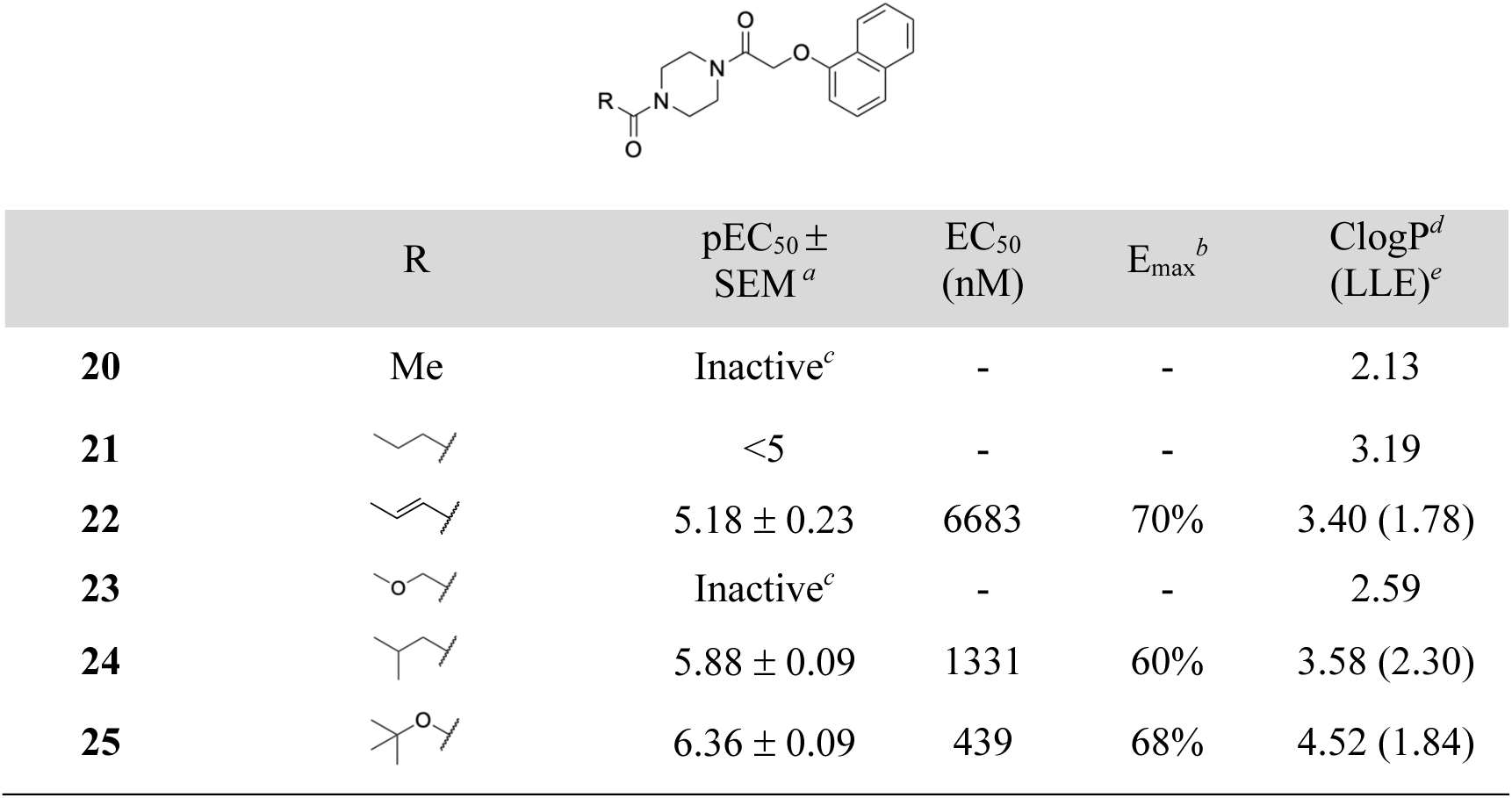
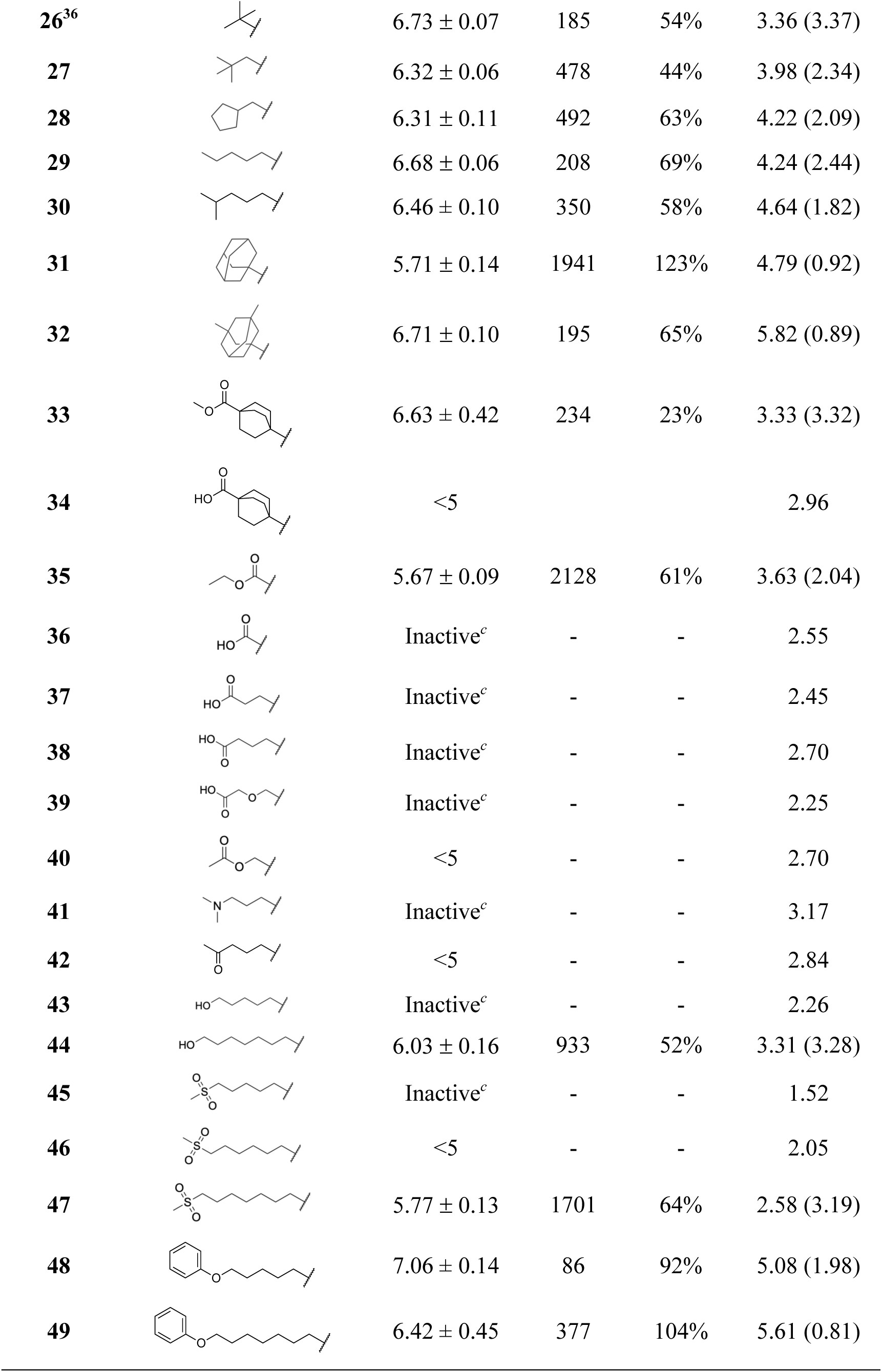
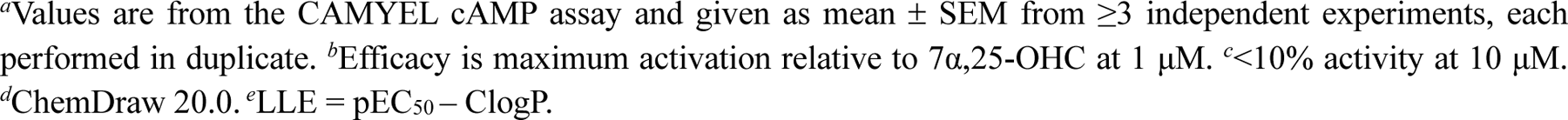
Explorations of the Western Part.

|  | R | pEC<sub>50</sub> ± SEM<sup>a</sup> | EC<sub>50</sub> (nM) | E<sub>max</sub><sup>b</sup> | ClogP<sup>d</sup> (LLE)<sup>e</sup> |
| --- | --- | --- | --- | --- | --- |
| **20** | Me | Inactive<sup>c</sup> | - | - | 2.13 |
| **21** |  | <5 | - | - | 3.19 |
| **22** |  | 5.18 ± 0.23 | 6683 | 70% | 3.40 (1.78) |
| **23** |  | Inactive<sup>c</sup> | - | - | 2.59 |
| **24** |  | 5.88 ± 0.09 | 1331 | 60% | 3.58 (2.30) |
| **25** |  | 6.36 ± 0.09 | 439 | 68% | 4.52 (1.84) |
| 26<sup>36</sup> | | $6.73 \pm 0.07$ | 185 | 54% | 3.36 (3.37) |
| 27 | | $6.32 \pm 0.06$ | 478 | 44% | 3.98 (2.34) |
| 28 | | $6.31 \pm 0.11$ | 492 | 63% | 4.22 (2.09) |
| 29 | | $6.68 \pm 0.06$ | 208 | 69% | 4.24 (2.44) |
| 30 | | $6.46 \pm 0.10$ | 350 | 58% | 4.64 (1.82) |
| 31 | | $5.71 \pm 0.14$ | 1941 | 123% | 4.79 (0.92) |
| 32 | | $6.71 \pm 0.10$ | 195 | 65% | 5.82 (0.89) |
| 33 | | $6.63 \pm 0.42$ | 234 | 23% | 3.33 (3.32) |
| 34 |  | <5 |  |  | 2.96 |
| 35 | | $5.67 \pm 0.09$ | 2128 | 61% | 3.63 (2.04) |
| 36 |  | Inactive<sup>c</sup> | - | - | 2.55 |
| 37 |  | Inactive<sup>c</sup> | - | - | 2.45 |
| 38 |  | Inactive<sup>c</sup> | - | - | 2.70 |
| 39 |  | Inactive<sup>c</sup> | - | - | 2.25 |
| 40 |  | <5 | - | - | 2.70 |
| 41 |  | Inactive<sup>c</sup> | - | - | 3.17 |
| 42 |  | <5 | - | - | 2.84 |
| 43 |  | Inactive<sup>c</sup> | - | - | 2.26 |
| 44 | | $6.03 \pm 0.16$ | 933 | 52% | 3.31 (3.28) |
| 45 |  | Inactive<sup>c</sup> | - | - | 1.52 |
| 46 |  | <5 | - | - | 2.05 |
| 47 | | $5.77 \pm 0.13$ | 1701 | 64% | 2.58 (3.19) |
| 48 | | $7.06 \pm 0.14$ | 86 | 92% | 5.08 (1.98) |
| 49 | | $6.42 \pm 0.45$ | 377 | 104% | 5.61 (0.81) |

<sup>a</sup>Values are from the CAMYEL cAMP assay and given as mean ± SEM from $\geq 3$ independent experiments, each performed in duplicate. <sup>b</sup>Efficacy is maximum activation relative to 7α ,25-OHC at 1 μM. <sup>c</sup><10% activity at 10 μM. <sup>d</sup>ChemDraw 20.0. <sup>e</sup>LLE = pEC<sub>50</sub> – ClogP.

Lipophilicity is a key parameter for successful drug discovery.^38^ The preference of GPR183 for lipids and lipophilic compounds represents a challenge for the identification of high-quality ligands. For this reason, we closely monitored the lipophilicity and ligand lipophilicity efficiently (LLE),^39^ even if the current focus was pEC_50_ and E_max_ in the cAMP assay. To lower lipophilicity of the compounds, we then incorporated various polar terminal moieties. Introduction of an ester-functionalized bicyclo[2.2.2]octane (**33**) led to sustained potency compared to **32** but with reduced efficacy. The corresponding carboxylic acid (**34**) abolished agonistic activity. Similarly, the oxalate derivative **35** showed agonistic activity similarly to the shorter aliphatic analogues, whereas the corresponding acid (**36**) was completely inactive, as were the extended carboxylic acids **37-39**, indicating that a terminal negative charge is not well tolerated, at least not in the vicinity of the piperazine. Flipping the ester (**40**) led to a weak but not completely inactive compound. Exchanging the terminal group to dimethyl amine (**41**) to introduce a positive charge also led to an inactive compound.

Since both positive and negative charges seemed to be unfavored, we moved on to explore neutral polar groups such as terminal keto (**42**), hydroxy (**43-44**) and sulfonyl groups (**45-47**). Interestingly, only **44** and **47**, both of which contain C8 linker in between the polar group and the piperazine core, retained partial agonistic activity whereas the shorter analogues **42**, **43**, **45** and **46** were inactive or displayed poor potency. Converting the hydroxy analogues to the corresponding phenoxy analogues **48** and **49**, increased the potency and efficacy to full agonists, with the shorter analogue being 4-fold more potent. Overall, it seems like polar moieties can be incorporated in the western region of the compounds, but the distance to the piperazine core is important and masking of the polarity seems to be an advantage.

We then directed attention back to the eastern part while keeping the C6 aliphatic tail of **29** based on its balance between higher efficacy and good potency. First, the oxymethyl spacer was replaced with the corresponding (*E*)-alkene (**50**) and ethylene (**51**) linker, resulting in inactive compounds and supporting the importance of oxymethyl linker for agonistic activity (Table 4).^35^ Keeping the oxymethyl linker, the 1-naphthyl ring was replaced with various aromatic/heteroaromatic moieties. Replacement to the partly reduced tetrahydronaphthyl (**52**) led to 5-fold reduction in potency compared to **29**. Next, in order to reduce lipophilicity and pick up potential hydrogen bond interactions in the eastern part, we performed a partial nitrogen walk around the naphthalene ring. Of the five successfully synthesized compounds, **53** and **54** were micromolar potency partial agonists with preserved LLE, indicating that the nitrogen atoms were tolerated but not contributing to activity. The remaining quinoline analogues were inactive or close to inactive.

**Table 4.**
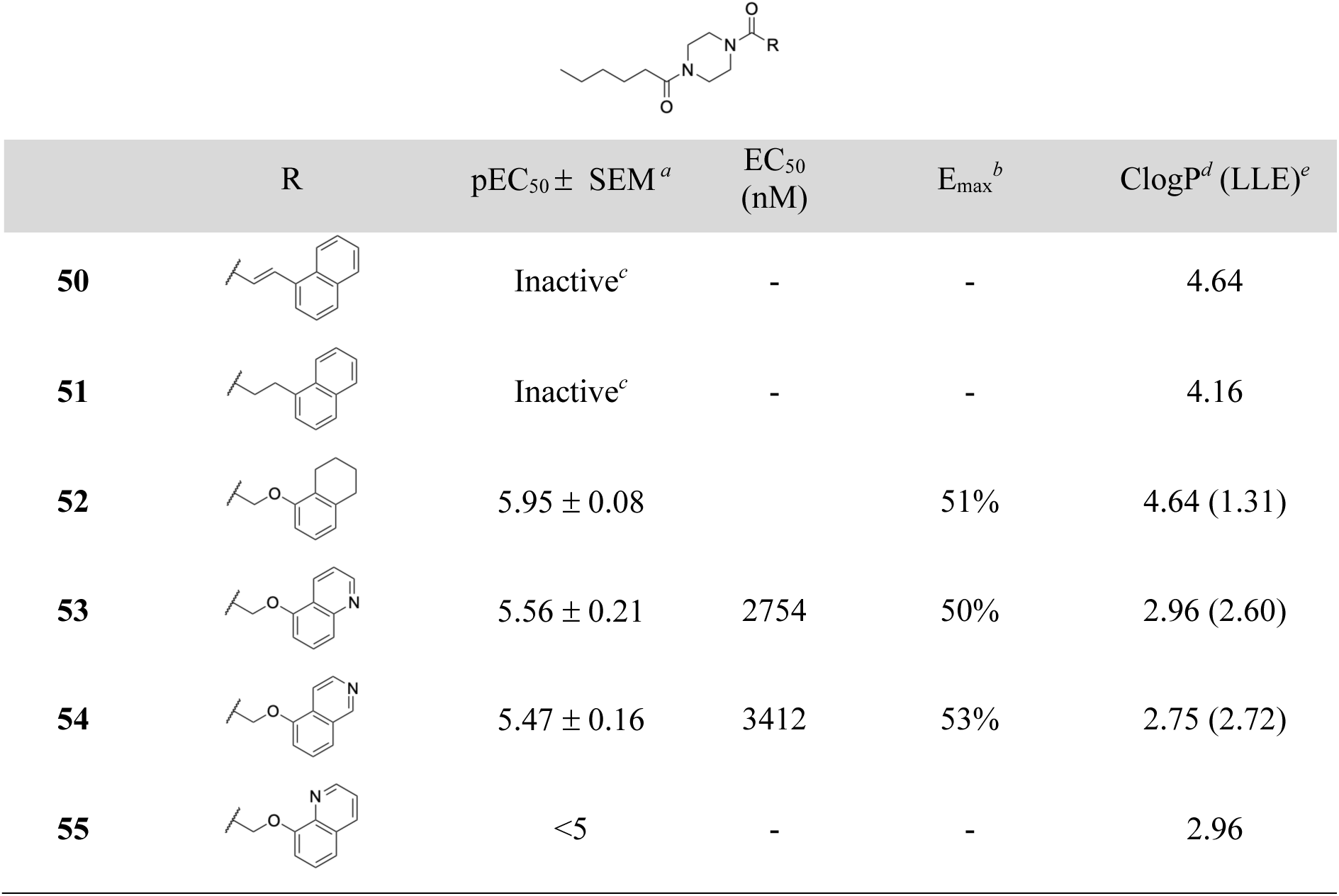
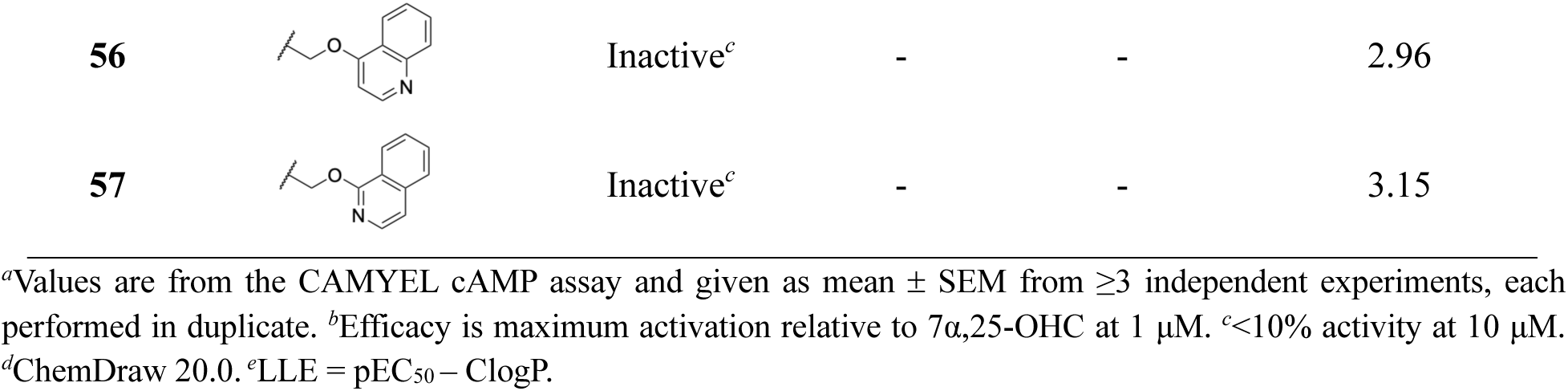
Exploration of Aromatic and Heteroaromatic Variations of the Naphthalene.

| | R | pEC<sub>50</sub> ± SEM <sup>a</sup> | EC<sub>50</sub> (nM) | E<sub>max</sub> <sup>b</sup> | ClogP<sup>d</sup> (LLE)<sup>e</sup> |
| --- | --- | --- | --- | --- | --- |
| **50** |  | Inactive<sup>c</sup> | - | - | 4.64 |
| **51** |  | Inactive<sup>c</sup> | - | - | 4.16 |
| **52** | | 5.95 ± 0.08 | | 51% | 4.64 (1.31) |
| **53** | | 5.56 ± 0.21 | 2754 | 50% | 2.96 (2.60) |
| **54** | | 5.47 ± 0.16 | 3412 | 53% | 2.75 (2.72) |
| **55** |  | <5 | - | - | 2.96 |
| **56** |  | Inactive<sup>c</sup> | - | - | 2.96 |
| **57** |  | Inactive<sup>c</sup> | - | - | 3.15 |

<sup>a</sup>Values are from the CAMYEL cAMP assay and given as mean ± SEM from $\geq 3$ independent experiments, each performed in duplicate. <sup>b</sup>Efficacy is maximum activation relative to 7α ,25-OHC at 1 μM. <sup>c</sup><10% activity at 10 μM.
<sup>d</sup>ChemDraw 20.0. <sup>e</sup>LLE = pEC<sub>50</sub> – ClogP.

Since the exploration of the eastern part did not result in improved compounds and the phenoxy analogues **48** and **49** indicated that there was additional space to explore in the western binding region, we went back to explore longer aliphatic substituent. The bromo intermediates **58-60** to the earlier sulfones (**45-47**) displayed full agonism in the GPR183 cAMP accumulation assay, with the longest analogue **60** being 60-fold more potent that the shorter analogues and showing nanomolar potency (Table 5). To investigate the depths of the western binding pocket, analogues with alkyl chain lengths of C8-C16 were tested and were all found to be active with potency in the order C16 < C6 < C12 < C8 < C9 < C10. Except for the less potent analogues, **65** and **29** (Table 3), all were full agonists. The C10 analogue **63** was the most potent GPR183 agonist with single digit nanomolar potency (EC_50_ = 5.4 nM, 90% activation relative to 7α,25-OHC). Insertion of an amide functionality in the C10 chain (**66**) led to a drastic decrease in potency, while introduction of terminal ester (**67**) was better tolerated but still led to a 20-fold decrease in potency. Extension of the ester by two carbons (**68**) further decreased the potency, and hydrolysis to give carboxylic acid **69** led to a complete loss of potency, in line with all previously explored carboxylic acids (Table 3). Finally, the extended C12 methyl ester (**70**) and corresponding acid (**71**) both showed decent agonist potency with partial efficacy, supporting the notion that polar heteroatoms were only well-tolerated at least eight carbon atoms further away from the piperazine core, but that the incorporation seems to be associated with a considerable decrease in both potency and efficacy. The outstanding potency and efficacy of **63** directed our attention to this compound even if it was not among the compounds with the highest LLE. Compounds **26**, **33**, **44** and **67** demonstrate that higher LLE is attainable, but their moderate potency made them less attractive for further studies than **63**.

**Table 5.** Alkyl Chain-Containing Western Substituents.

|  | R | pEC<sub>50</sub> ± SEM<sup>a</sup> | EC<sub>50</sub><br>(nM) | E<sub>max</sub><sup>b</sup> | ClogP<sup>c</sup><br>(LLE)<sup>d</sup> |
| --- | --- | --- | --- | --- | --- |
| **58** |  | 5.70 ± 0.13 | 2008 | 109% | 4.10 (1.60) |
| **59** |  | 5.86 ± 0.17 | 1393 | 140% | 4.63 (1.23) |
| **60** |  | 7.68 ± 0.09 | 21 | 91% | 5.15 (2.53) |
| **61** |  | 7.50 ± 0.15 | 32 | 88% | 5.30 (2.20) |
| **62** |  | 7.97 ± 0.09 | 11 | 91% | 5.83 (2.14) |
| **63** |  | 8.27 ± 0.07 | 5.4 | 90% | 6.35 (1.92) |
| **64** |  | 7.32 ± 0.06 | 48 | 91% | 7.42 (-0.10) |
| **65** |  | 5.85 ± 0.15 | 1415 | 61% | 9.53 (-3.68) |
| **66** |  | <5 | - | - | 2.47 |
| **67** |  | 6.97 ± 0.11 | 108 | 76% | 3.73 (3.24) |
| **68** |  | 5.92 ± 0.15 | 1202 | 65% | 4.79 (1.21) |
| **69** |  | Inactive<sup>e</sup> | - | - | 4.41 |
| **70** |  | 6.24 ± 0.11 | 581 | 78% | 5.85 (0.39) |
| **71** |  | 6.64 ± 0.24 | 229 | 50% | 5.47 (1.17) |

<sup>a</sup>Values are from the CAMYEL cAMP assay and given as mean ± SEM from ≥3 independent experiments, each performed in duplicate. <sup>b</sup>Efficacy is maximum activation relative to 7α,25-OHC at 1 μM. <sup>c</sup>ChemDraw 20.0. <sup>d</sup>LLE = pEC<sub>50</sub> – ClogP. <sup>e</sup><10% activity at 10 μM.

To evaluate their general potential, we investigated the kinetic aqueous solubility of a selection of the most potent and efficacious compounds (**48**, **59**, **60**, **63-65**) in a previously established assay.^40^ As expected in light of their high lipophilicity, all compounds showed poor (<5 μM) aqueous solubility. None of the tested compounds generated PAINS^41^ hits by in silico screening.^42^ Testing in the standard MTT (3-(4,5-dimethyl-2-thiazolyl)-2,5-diphenyl-2*H*-tetrazolium bromide) assay^43^ did not indicate reduced cell viability of **63** at 3 or 10 μM concentration (see the SI).

The agonist series were derived from the antagonists NIBR127 an NIBR189 (Chart 1),^35^ thus, we wished to investigate if the inactive compounds acted as antagonists. However, none of the compounds with pEC_50_ <5 showed significant antagonism of 7α,25-OHC, except **2** and **3** that inhibited the 7α,25-OHC to 38% and 29%, respectively (see the SI). This is notable, since **2** and **3** are the only analogues that lack the phenoxyacetamide *ortho*-substituent. This demonstrate that the *ortho*-substituent is a reliable switch for G protein activation, albeit with varying efficacy, and that inactive compounds lose affinity rather than become antagonists. We furthermore confirmed that the activity of **63** can be fully inhibited by NIBR189, supporting that agonists and antagonists of the series compete for the same binding site (see the SI).

While the SAR for agonist activity is clearly associated with the *ortho*-substituent at the eastern phenoxyacetamide part and the potency is associated with the length of the western acylamide chain, the structural basis for high efficacy is less clear, with the various lengths of the eastern alkyl chains consistently being close to full agonists while bromo-terminal octanoyl **59** jumps to 140%. Likewise, the bulky adamantly **31** jumps to 123% while the smaller *tert*-butyl (**26**), the larger dimethyladamantyl (**32**) and other analogs of similar size all are partial agonists.

Selected compounds (**48**, **59**, **60**, **62-64**) were tested on the free fatty acid sensing receptors FFA1, FFA4 and GPR84, the cannabinoid CB_1_ receptor, the lysophosphatidic acid LPA_1_ receptor, and the chemokine CCR7 receptor (see below). None of the compounds caused substantial activation of any of the receptors (see the SI). Furthermore, **29** and **64** did not give any response up to 10 μM in cells expressing CAMYEL but not GPR183 (not shown), suggesting that the compounds do not activate constitutively expressed receptors that signal by cAMP.

### Biased Signaling

We previously found that the partial GPR183 G_i_ agonists TUG-2201, TUG-2202, and **26** (TUG-2292) had no effect on β-arrestin2.^35,36^ Therefore, we next tested selected compounds, including **1, 24, 27**–**29, 35**, and **62**–**64** in a BRET-based β-arrestin2 recruitment assay (Figure 1). None of the compounds showed any activity except **64**, which displayed 25% β-arrestin2 recruitment relative to 7α,25-OHC at 10 μM concentration. This is consistent with the previous results and confirms that members of this compound series is not only highly G protein-biased, but indeed G protein specific for most members, and furthermore demonstrates that this also holds true for compounds that are highly potent full agonists in the cAMP assay, such as **63** and **62**. To confirm that the strong G protein bias is consistent throughout the series, compounds **48, 59, 60**, with 91-140% efficacy in the cAMP assay, were tested in the β-arrestin2 assay at 1 and 10 μM concentration and did not display significant activity. Although calculation of bias factor is recommended for comparison of signaling bias,^44^ this concept is unapplicable if a signal is not detected in both pathways.

**Figure 1.**
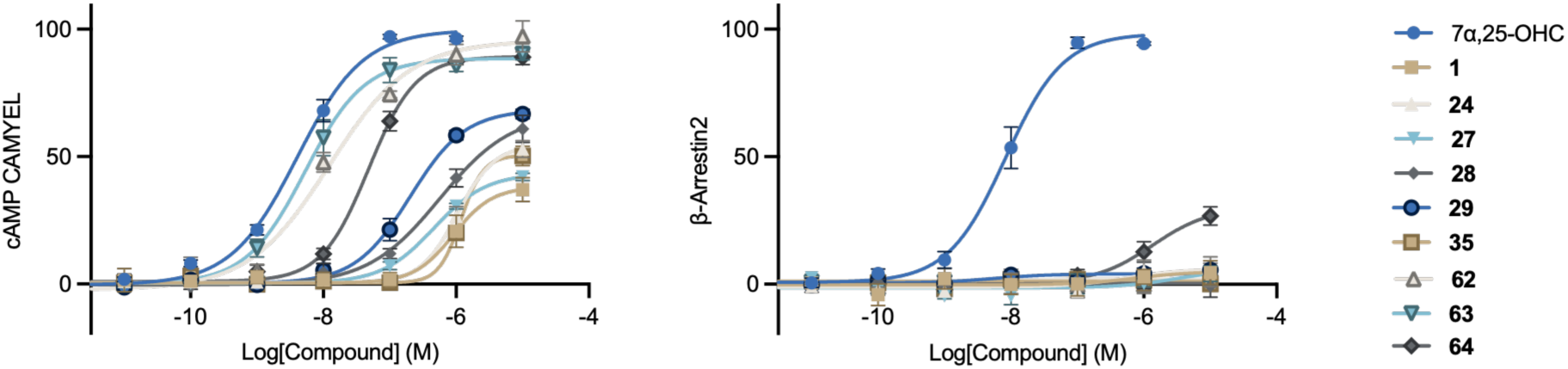
G protein signaling (cAMP) elicited by selected compounds is shown on the left whereas the analysis of β-arrestin2 recruitment is shown on the right. BRET ratio (525 nm/475 nm), normalized to the max of 7α,25-OHC (%). Data from ≥3 independent experiments, each performed in duplicate and given as mean ± SEM.

Receptor internalization is linked to β-arrestin2 recruitment. We investigated the ability of 7α,25-OHC and selected compound to promote internalization GPR183 labeled with Lumi4-Tb.^36,45^ Consistent with the absence of β-arrestin2 signaling, 7α,25-OHC effectively induced internalization but none of the G protein specific agonists **27, 28, 29, 63** or **64** increased intracellular fluorescence (Figure 2).

**Figure 2.**
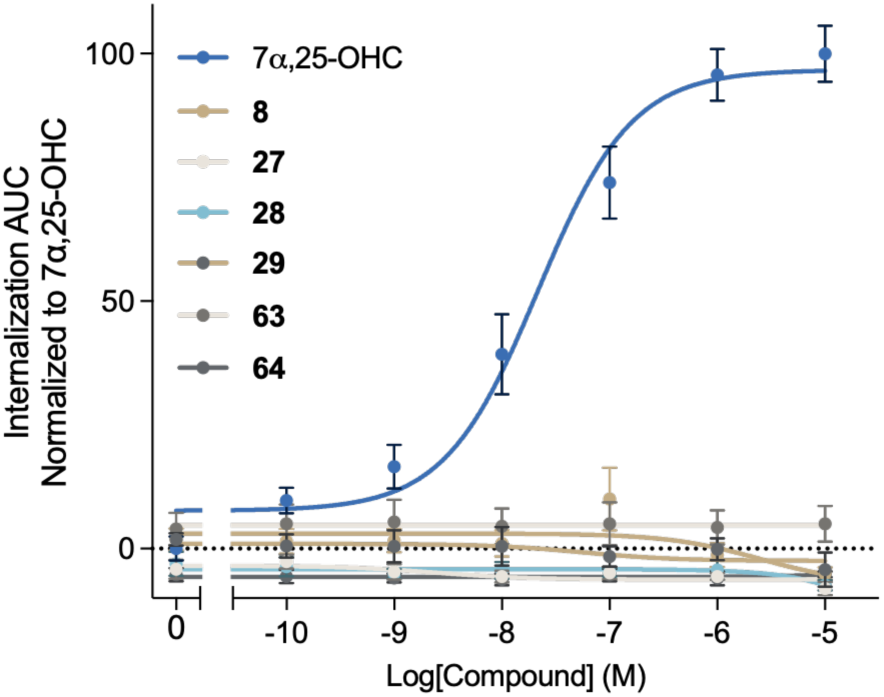
Ligand-induced internalization of GPR183 in HEK293A cells. Data from 3 independent experiments, each performed in duplicate and given as mean ± SEM.

GPR183-promoted chemotaxis is found to be mediated by β-arrestin2 signaling.^36^ Consistent with this, **29, 64** and **63** were unable to induce *in vitro* migration of human stimulated dendritic cells, as also previously observed for **26** (Figure 3).^36^ Thus, our results confirm that GPR183-promoted G protein activation do not result in receptor internalization or migration of dendritic cells. These properties are of high interest for conditions where GPR183 agonists is a relevant treatment modality, since our results suggest that the compounds would not cause receptor desensitization and internalization, and undesired agonist-induced immune cell mobilization and infiltration could be avoided.

**Figure 3.**
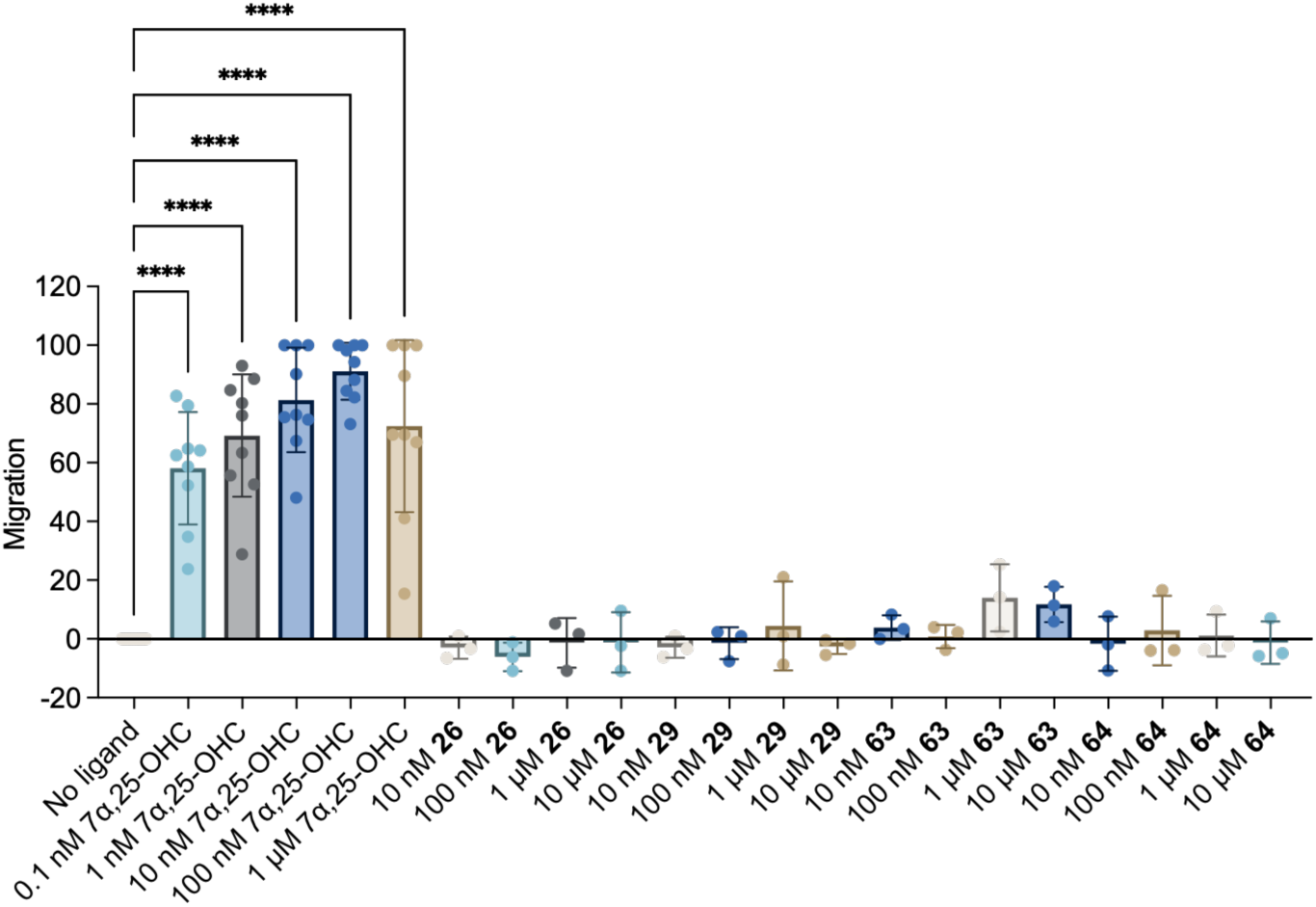
Ligand-induced migration of human stimulated dendritic cells. Normalized to maximum 7α,25-OHC Data from ≥3 independent experiments, each performed in duplicate and given as mean ± SEM. One-way ANOVA with Dunnett’s test for multiple comparisons. ****, p<0.0001.

Given that **63** potently stimulates Gi signaling of GPR183 without β-arrestin2 recruitment required for chemotaxis, we hypothesized that the compound would compete with 7α,25-OHC and antagonize its chemoattractant effects. To test the hypothesis, we investigated the effect of the compound on 7α,25-OHC-induced dendritic cell migration. Indeed, **63** dose-dependently inhibited 7α,25-OHC-induced migration with complete inhibition obtained around 1 μM (Figure 4A). As a control experiment, we tested the effect of **63** on CCL19-induced dendritic cell migration, which acts through the chemokine CCR7 receptor.^46^ In this system, **63** showed no effect (Figure 4B), demonstrating that the inhibition of 7α,25-OHC-induced migration is mediated by GPR183 and also that **63** has no effect on CCR7.

**Figure 4.**
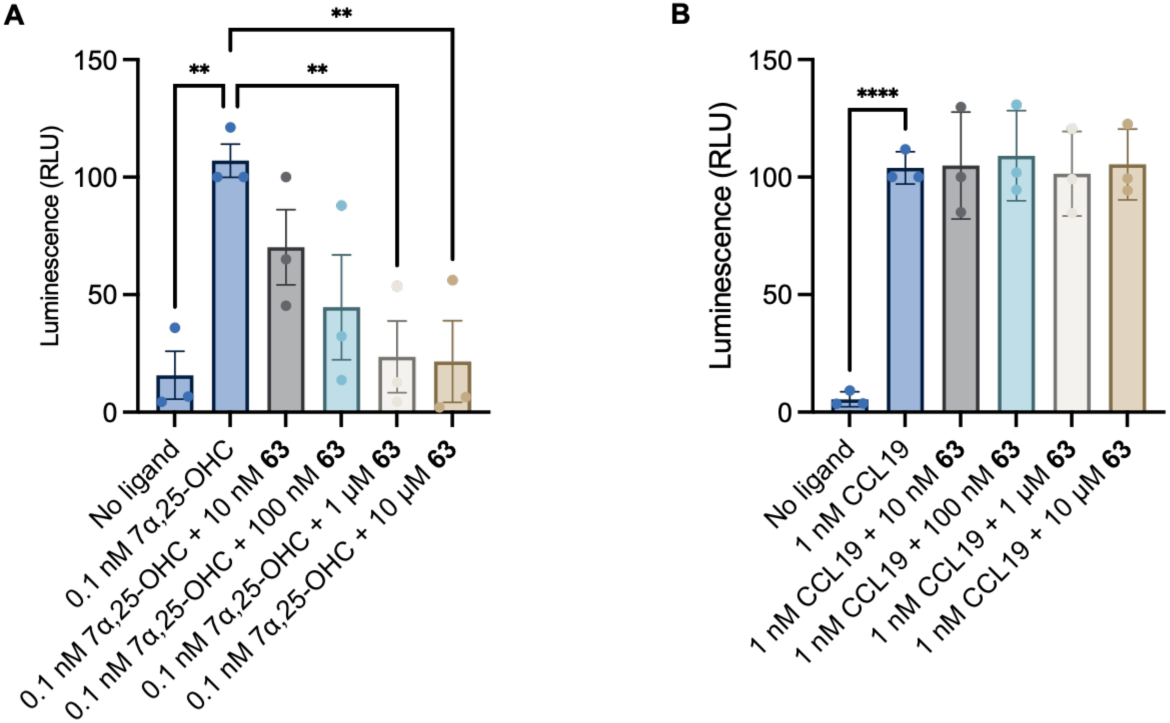
Inhibition of dendritic cell migration by **63** (TUG-2604). Migration of human dendritic cells was indued by 7α,25-OHC (panel A) or CCL19 (panel B), each treated with increasing concentration of **63**. A: Inhibition of 7α,25-OHC-induced human dendritic cell migration. B: Inhibition of CCL19-induced human dendritic cell migration. **, p<0.01; ****, p<0.0001.

### Synthesis

Compounds were synthesized by coupling of 1-Boc-piperazine (**A**) with 2-chloroacetyl chloride followed by nucleophilic substitution with the relevant phenols, Boc-deprotection to **D**, and coupling with carboxylic acids or acyl chlorides, as described previously (Scheme 1).^35^ Final coupling of the intermediate **D** with carboxylic acids, phenyl isocyanate and anhydrides afforded **1-7, 9-33, 35, 37-44, 58-65, 67-68**, and **70** respectively. Carboxylic acids **36, 34, 69** and **71** were prepared from the ester hydrolysis of the respective esters, compound **66** was synthesized from the corresponding amine by coupling with propionic acid, and sulfones **45-47** were prepared by reaction of the corresponding bromo compounds **58-60** with sodium methanesulfinate.

**Scheme 1a.**
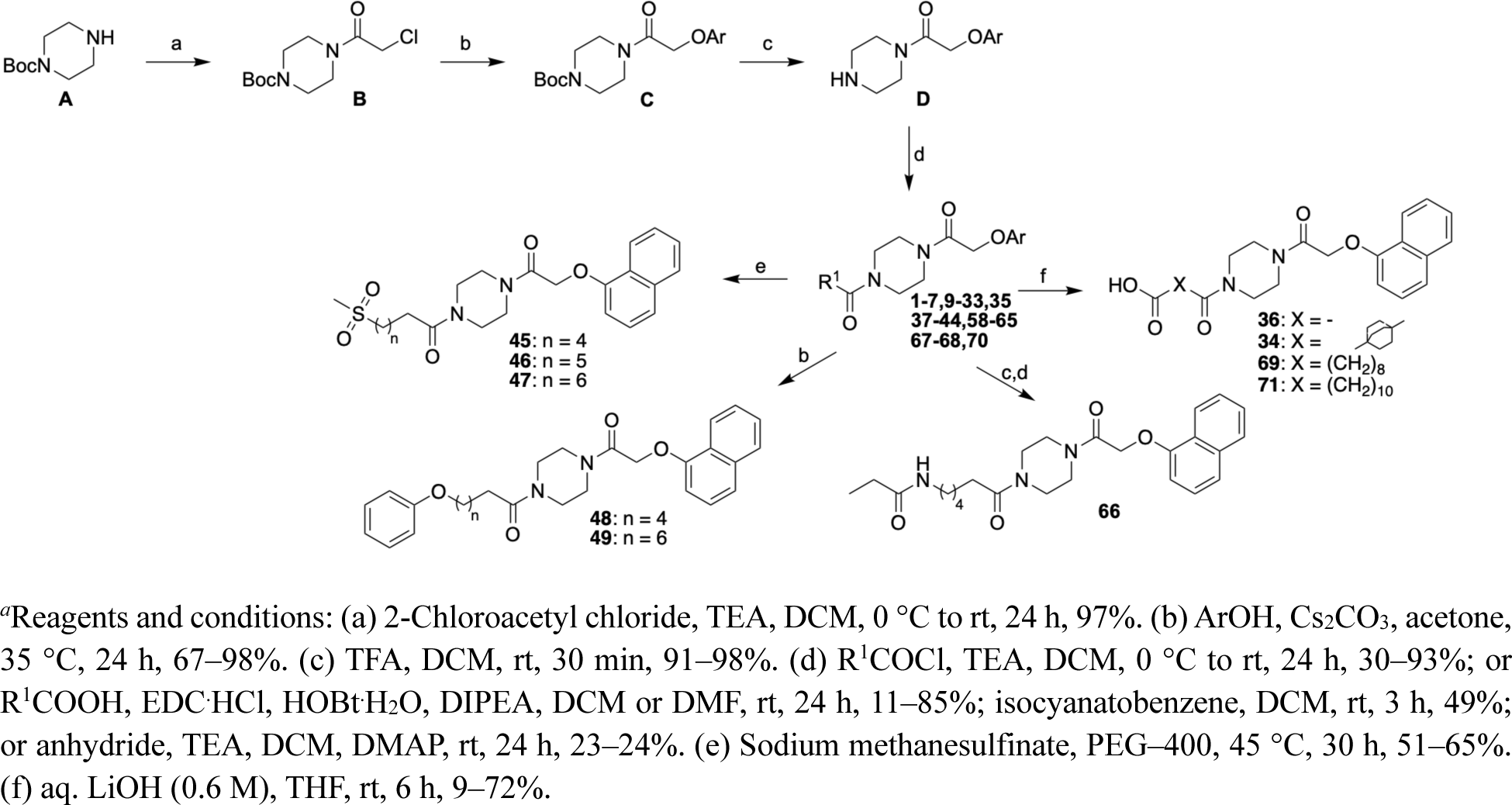
Synthesis of compounds 1-7, 9-49, 58-71.

For compounds **8, 52-57**, the reactions order was changed to facilitate variation of the eastern substituent to produce various aryloxyacetamides via chloroacetamide **G** as well as alkene **50** and ethylene analogue **51** (Scheme 2).

**Scheme 2a.**
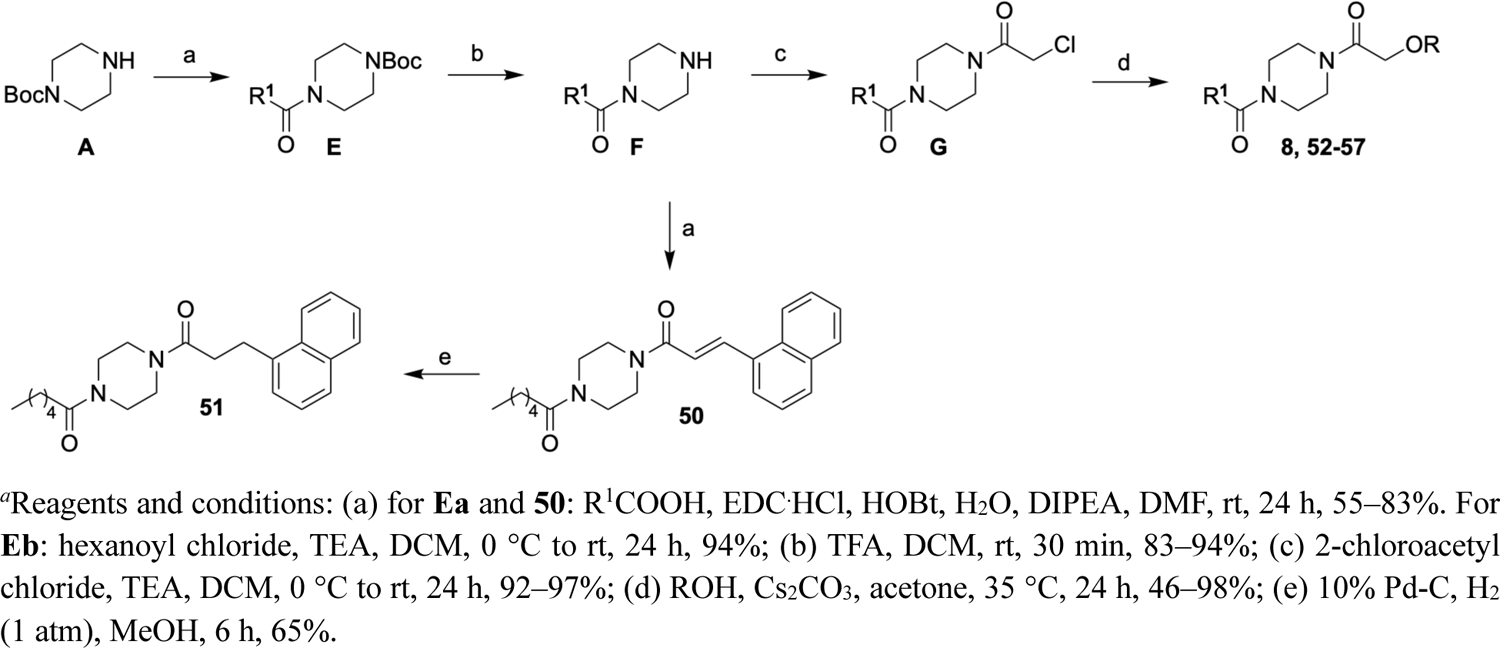
Synthesis of compounds 8, 50-57.

## Conclusion

We here report a systematic SAR investigation of piperazine aryloxyacetamide GPR183 ligands with focus on optimizing agonistic activity. Our findings confirm that *ortho*-substituted phenoxyacetamides behave as agonists in the G protein dependent cAMP accumulation assay and that the 1-naphthyloxyacetamide eastern part is favored for obtaining high efficacy and potency. The methyleneoxy linker is also crucial for agonistic activity. On the other side, the western carboxamide part is more accommodating to variations in chain length and bulkiness, whereas no polar moieties in vicinity to the piperazine diamide core were tolerated. Modifications of the western side affected both the potency and efficacy of the agonists and allowed discovery of full agonists with efficacy and potency similar to 7α,25-OHC. Importantly, compounds of this series were generally found unable to induce β-arrestin2 recruitment. Even for the full agonists **62** and **63** with EC_50_ down to 5 nM, no β-arrestin2 recruitment was detected up to 10 μM, implying G protein-specific agonism with >1000-fold signaling pathway selectivity by direct assay readouts. However, the 25% β-arrestin2 activation of **64** at 10 μM, even if the bias still is high, illustrate that some compounds of the series may activate β-arrestin2. Furthermore, consistent with the general absence of β-arrestin2 recruitment, the compounds did not promote receptor internalization. The compounds were also largely unable to induce chemotaxis of human stimulated dendritic cells in vitro and **63** was demonstrated to inhibit the chemoattractant effect of 7α,25-OHC. This profile is of interest for therapeutic strategies that involve GPR183 activation. Biased GPCR agonism has attracted much attention and still holds potential to revolutionize GPCR-targeted drug discovery even if not all initial attempts at this strategy have been successful.^47^ Compounds such as **63** (TUG-2604) will be valuable for further exploration of specific GPR183 G protein signaling, that can contribute to define and validate such therapeutic strategies. As the most potent full agonists have high lipophilicity, we expect that these compounds will be most useful for in vitro studies. Optimizations to identify biased GPR183 agonists with lower lipophilicity that will be better suited for in vivo studies are currently in progress in our laboratories.

## Experimental Section

### General

All reagents were of commercial grade and were used without further purification. Anhydrous reactions were carried out under argon atmosphere in flame-dried glassware. Anhydrous chromatographic grade DCM and DMF were obtained from a Waters SG solvent purification system. Silica gel 60 F254 precoated plates (Merck) were used for TLC. Flash chromatography of compounds was performed either manually using silica gel 60 (40–64 μm) or automated using Reveleris®X2 Flash Chromatography System, Buchi. ^1^H NMR spectra were obtained at 400 or 600 MHz, and ^13^C NMR spectra were obtained at 100 or 151 MHz and calibrated relative to residual solvent peaks. Analytical HPLC was performed on a Dionex UltiMate system using a Gemini-NX C18 column (4.6 mm × 250 mm, 3 μm, 110 Å); flow 1 mL/min; UV detection at 254 nm; mobile phase A: water:TFA, 100:0.1, v/v; and mobile phase B: MeCN:water:TFA, 90:10:0.1, v/v/v. Data were acquired and processed using the Chromeleon Software v. 6.80. The analysis was performed by (a) Method A: 40-100% mobile phase B in mobile phase A over 10 min; (b) Method B: 50-100% mobile phase B in mobile phase A over 10 min; (c) Method C: 50-100% mobile phase B in mobile phase A over 20 min; (d) Method D: 0-100% mobile phase B in mobile phase A over 15 min; (e) Method E: 20-100% mobile phase B in mobile phase A over 15 min; (f) Method F: 30-100% mobile phase B in mobile phase A over 15 min; (g) Method G: 0-100% mobile phase B in mobile phase A over 20 min; (h) Method H: 0-100% mobile phase B in mobile phase A over 10 min; (i) Method I: 50-100% mobile phase B in mobile phase A over 15 min. All test compounds except **36** (94.0% pure), **58** (94.0% pure), **60** (94.4% pure), **71** (94.5% pure) were >95% pure by HPLC. Preparative HPLC was performed on a Dionex UltiMate system using a Gemini-NX C18 column (21.2 mm × 250 mm, 5 μm, 110 Å) with mobile phase A, water:TFA, 100:0.1, v/v; mobile phase B, MeCN:water:TFA, 90:10:0.1, v/v/v. HRMS (MALDI) was performed on a QExactive Orbitrap mass spectrometer equipped with a SMALDI5 ion source using 2,5-dihydroxybenzoic acid as a matrix. Rotamer peaks are indicated by asterisk (*). Integrals reported as H represent sum of rotamers, whereas integrals reported as H′ and H″ represent specific rotamers where H′ + H″ = H for the relevant signal.

### 1-(4-Benzoylpiperazin-1-yl)-2-(4-chlorophenoxy)ethan-1-one (2)

Step 1: *tert*-Butyl 4-(2-chloroacetyl)piperazine-1-carboxylate (**B**): A flame dried round bottom flask was sequentially charged with 1-bocpiperazine **A** (2.2 g, 11.8 mmol) DCM (200 mL), and TEA (2.5 mL, 17.9 mmol). Then, 2-chloroacetyl chloride (1.9 mL, 23.6 mmol) was added dropwise at 0 °C and the mixture was allowed to stir at rt. After 24 h, the reaction mixture was washed with aq. NaOH solution (1 M, 200 mL) and brine (200 mL). The organic layer was dried over anhydrous Na_2_SO_4_, filtered and the solvent was evaporated under reduced pressure to afford the target compound (3.01 g, 97%) as a white solid. The product was used in the next reaction without further purification. R_f_ = 0.63 (MeOH:DCM; 2:8); ^1^H NMR (400 MHz, CDCl_3_) δ 4.06 (s, 2H), 3.59 (t, *J* = 5.3 Hz, 2H), 3.49 (s, 4H), 3.43 (t, *J* = 5.4 Hz, 2H), 1.46 (s, 9H); ^13^C NMR (101 MHz, CDCl_3_) δ 165.4, 154.6, 80.6, 46.3, 43.5, 42.1, 40.9, 28.5. ^1^H NMR spectra are in accordance with the literature.

Step 2: *tert*-Butyl 4-(2-(4-chlorophenoxy)acetyl)piperazine-1-carboxylate (**Ca**): A round bottom flask was charged with **B** (400 mg, 1.52 mmol), 4-chlorophenol (293 mg, 2.3 mmol), Cs_2_CO_3_ (992 mg, 3.04 mmol), KI (13 mg, 76 µmol), and acetone (12 mL). The reaction mixture was stirred at 35 °C for 24 h. The solvent was evaporated, and the residue was suspended in EtOAc (20 mL). The organic solution was washed with aq. NaOH solution (1 M, 2 x 10 mL) and brine (10 mL). The organic phase was dried over Na_2_SO_4_, filtered and the solvent was evaporated under reduced pressure. The residue was purified by flash chromatography (60% EtOAc in heptane) to afford the target compound (444 mg, 82%) as a white solid. Melting Point Range: 95.7-99.3 °C; R_f_ = 0.34 (EtOAc:Heptane; 1:1); ^1^H NMR (400 MHz, CDCl_3_) δ 7.28 – 7.20 (m, 2H), 6.92 – 6.83 (m, 2H), 4.68 (s, 2H), 3.62 – 3.50 (m, 4H), 3.46 – 3.36 (m, 4H), 1.46 (s, 9H); ^13^C NMR (101 MHz, CDCl_3_) δ 166.5, 156.4, 154.6, 129.7, 126.9, 116.0, 80.6, 68.1, 45.4, 42.1, 28.5; HPLC: t*_R_* = 7.62 min, purity 99.4%, Method A; HRMS (MALDI) calcd for C_17_H_24_ClN_2_O_4_^+^ (M+H^+^) 355.1419, found 355.1415.

Step 3: 2-(4-Chlorophenoxy)-1-(piperazin-1-yl)ethan-1-one (**Da**): To a solution of **Ca** (200 mg, 0.56 mmol) in DCM (3 mL) was added TFA (0.86 mL, 11.3 mmol). The mixture was stirred at rt for 30 min and evaporated to dryness. The residue was dissolved in DCM (10 mL), and the solution was washed with aq. Na_2_CO_3_ (2M, 2 x10 mL). The organic layer was dried over Na_2_SO_4_, filtered, and evaporated to dryness to afford the target compound (140 mg, 98%) as a colorless oil. The product was used in the next reaction without further purification. R_f_ = 0.72 (MeOH:DCM; 2:8); ^1^H NMR (400 MHz, CDCl_3_) δ 7.25 – 7.20 (m, 2H), 6.92 – 6.84 (m, 2H), 4.66 (s, 2H), 3.63 – 3.49 (m, 4H), 2.88 – 2.79 (m, 4H), 1.84 (s, 1H); ^13^C NMR (101 MHz, CDCl_3_) δ 166.1, 156.5, 129.5, 126.6, 116.0, 67.9, 46.6, 46.3, 45.8, 43.2.

Step 4: To a solution of **Da** (120 mg, 0.47 mmol) in DCM (4 mL) was added TEA (0.13 mL, 0.93 mmol) and cooled to 0 °C. Benzoyl chloride (80 µL, 0.69 mmol) was added dropwise over 15 min. The reaction mixture was allowed to stir at rt. After 24 h, the reaction mixture was washed with 0.1 N HCl_aq_ (x2) and brine. The organic layer was dried with anhydrous MgSO_4_, filtered, and the solvent was evaporated under reduced pressure. The crude was purified by flash chromatography (70% EtOAc in heptane) to afford the title compound (157 mg, 93%) as a white solid. Melting Point Range: 113.3-117.9 °C; R_f_ = 0.33 (EtOAc:Heptane; 7:3); ^1^H NMR (600 MHz, CDCl_3_) δ 7.47 – 7.37 (m, 5H), 7.27 – 7.22 (m, 2H), 6.88 (d, *J* = 8.4 Hz, 2H), 4.70 (s, 2H), 3.63* (s, 6H’), 3.44* (s, 2H”); ^13^C NMR (151 MHz, CDCl_3_) δ 170.8, 166.6, 156.3, 135.1, 130.3, 129.8, 128.8, 127.2, 127.1, 116.0, 68.2, 45.6, 42.4; HPLC: t*_R_* = 6.62 min, purity 99.5%, Method A; HRMS (MALDI) m/z calcd for C_19_H_20_ClN_2_O_3_^+^ (M+H^+^) 359.1157, found 359.1152.

### 1-(4-Benzoylpiperazin-1-yl)-2-(o-tolyloxy)ethan-1-one (1)

Step 1: *tert*-Butyl 4-(2-(*o*-tolyloxy)acetyl)piperazine-1-carboxylate (**Cb**): The target compound was synthesized according to the procedure described for **Ca** using *o*-cresol (247 mg, 2.28 mmol) to afford 420 mg (82%) of a white solid after purification by flash chromatography (60% EtOAc in heptane). Melting Point Range: 89.4-93.3 °C; R_f_ = 0.42 (EtOAc:Heptane; 1:1); ^1^H NMR (400 MHz, CDCl_3_) δ 7.19 – 7.10 (m, 2H), 6.90 (t, *J* = 1.1 Hz, 1H), 6.84 (d, *J* = 8.9 Hz, 1H), 4.71 (s, 2H), 3.63 – 3.56 (m, 4H), 3.46 – 3.37 (m, 4H), 2.24 (s, 3H), 1.46 (s, 9H); ^13^C NMR (101 MHz, CDCl_3_) δ 167.0, 155.9, 154.6, 131.2, 127.2, 126.6, 121.6, 111.0, 80.5, 68.2, 45.5, 42.1, 28.5, 16.4; HRMS (MALDI) m/z calcd for C_18_H_27_N_2_O_4_^+^ (M+H^+^) 335.1965, found 335.1962.

Step 2: 1-(Piperazin-1-yl)-2-(*o*-tolyloxy)ethan-1-one (**Db**): The target compound was synthesized according to the procedure described for **Da** using **Cb** (200 mg, 0.60 mmol) to yield 130 mg (93%) of a colorless oil. R_f_ = 0.76 (MeOH:DCM; 2:8); ^1^H NMR (400 MHz, CDCl_3_) δ 7.19 – 7.10 (m, 2H), 6.89 (t, *J* = 1.1 Hz, 1H), 6.86 – 6.83 (m, 1H), 4.69 (s, 2H), 3.65 – 3.55 (m, 4H), 2.85 (t, *J* = 4.8 Hz, 4H), 2.25 (s, 3H), 1.89 (s, 1H); ^13^C NMR (101 MHz, CDCl_3_) δ 166.8, 156.1, 131.1, 127.1, 126.7, 121.4, 111.1, 68.1, 46.8, 46.5, 46.0, 43.3, 16.4.

Step 3: The title compound was synthesized according to the procedure described for **2** using **Db** (120 mg, 0.51 mmol) to afford 122 mg (70%) of a white solid after purification by flash chromatography (70% EtOAc in heptane). Melting Point Range: 132.6-134.6 °C; R_f_ = 0.32 (EtOAc:Heptane; 7:3); ^1^H NMR (400 MHz, CDCl_3_) δ 7.49 – 7.35 (m, 5H), 7.15 (t, *J* = 7.1 Hz, 2H), 6.91 (t, *J* = 7.4 Hz, 1H), 6.86 (d, *J* = 8.3 Hz, 1H), 4.73 (s, 2H), 3.68* (s, 6H’), 3.47* (s, 2H”), 2.24 (s, 3H); ^13^C NMR (101 MHz, CDCl_3_) δ 170.8, 167.2, 155.8, 135.2, 131.2, 130.3, 128.8, 127.2, 126.6, 121.7, 111.0, 68.2, 45.7, 42.4, 16.4; HPLC: t*_R_* = 6.49 min, purity 99.5%, Method A; HRMS (MALDI) m/z calcd for C_20_H_23_N_2_O_3_^+^ (M+H^+^) 339.1703, found 339.1699.

### 1-(4-Benzoylpiperazin-1-yl)-2-(naphthalen-2-yloxy)ethan-1-one (3)

Step 1: *tert*-Butyl 4-(2-(naphthalen-2-yloxy)acetyl)piperazine-1-carboxylate (**Cc**): The target compound was synthesized according to the procedure described for **Ca** using naphthalen-2-ol (288 mg, 2 mmol) to afford 368 mg (75%) of a white solid after purification by flash chromatography (60% EtOAc in heptane). R_f_ = 0.42 (EtOAc:Heptane; 1:1); ^1^H NMR (400 MHz, CDCl_3_) δ 7.81 – 7.71 (m, 3H), 7.48 – 7.42 (m, 1H), 7.39 – 7.33 (m, 1H), 7.24 – 7.14 (m, 2H), 4.82 (s, 2H), 3.64 – 3.57 (m, 4H), 3.48 – 3.36 (m, 4H), 1.46 (s, 9H); ^13^C NMR (101 MHz, CDCl_3_) δ 166.8, 155.7, 154.6, 134.5, 130.0, 129.5, 127.8, 127.1, 126.8, 124.3, 118.4, 107.4, 80.5, 68.1, 45.6, 42.2, 28.5.

Step 2: 2-(Naphthalen-2-yloxy)-1-(piperazin-1-yl)ethan-1-one (**Dc**): The target compound was synthesized according to the procedure described for **Da** using **Cc** (300 mg, 0.81 mmol) to yield 214 mg (98%) of a yellow oil. R_f_ = 0.26 (EtOAc:Heptane; 7:3); ^1^H NMR (400 MHz, CDCl_3_) δ 7.80 – 7.71 (m, 3H), 7.44 (t, *J* = 1.3 Hz, 1H), 7.35 (t, *J* = 1.3 Hz, 1H), 7.23 – 7.15 (m, 2H), 4.80 (s, 2H), 3.65 – 3.55 (m, 4H), 2.89 – 2.79 (m, 4H), 1.78 (s, 1H); ^13^C NMR (101 MHz, CDCl_3_) δ 166.5, 155.9, 134.5, 129.8, 129.5, 127.8, 127.1, 126.7, 124.2, 118.5, 107.5, 67.9, 46.9, 46.5, 46.0, 43.4.

Step 3: The title compound was synthesized according to the procedure described for **2** using **Dc** (50 mg, 0.18 mmol) to afford 57 mg (82%) of a white solid after purification by flash chromatography (70% EtOAc in heptane). Melting Point Range: 154.1-156.8 °C; R_f_ = 0.56 (MeOH:DCM; 5:95); ^1^H NMR (400 MHz, CDCl_3_) δ 7.81 – 7.72 (m, 3H), 7.50 – 7.33 (m, 7H), 7.24 – 7.13 (m, 2H), 4.84 (s, 2H), 3.68* (s, 6H’), 3.46* (s, 2H”); ^13^C NMR (151 MHz, CDCl_3_) δ 170.7, 166.8, 155.6, 135.2, 134.4, 130.2, 130.0, 129.5, 128.8, 127.8, 127.2, 127.1, 126.8, 124.4, 118.2, 107.4, 68.0, 45.6, 42.4; HPLC: t*_R_* = 7.06 min, purity 99.2%, Method A; HRMS (MALDI) m/z calcd for C_23_H_23_N_2_O_3_^+^ (M+H^+^) 375.1703, found 375.1697.

### 1-(4-(4-Methoxybenzoyl)piperazin-1-yl)-2-(naphthalen-1-yloxy)ethan-1-one (4)

A flame-dried round bottom flask was charged with 4-methoxybenzoic acid (56 mg, 0.37 mmol), HOBt.H_2_O (55 mg, 0.41 mmol), EDC·HCl (78 mg, 0.41 mmol), and **Dd** (100 mg, 0.37 mmol). The flask was evacuated and backfilled with argon three times. Then, anhydrous DCM (2.2 mL) was added. The mixture was cooled down to 0 °C and DIPEA (130 μL, 0.75 μmol) was added. The ice bath was removed, and the mixture was stirred at rt. After 24 h, DCM (20 mL) was added and the organic phase was washed with aq. HCl (1.0 M, 2 x 20 mL), followed by aq. NaOH (1.0 M, 2 x 20 mL) and brine (30 mL). The organic layer was dried over anhydrous MgSO_4_, filtered, and the filtrate was evaporated onto Celite and purified by flash column chromatography (65-90% EtOAc in heptane) to afford the title compound (84 mg, 56%) as a faint orange solid. R_f_ = 0.24 (EtOAc:Heptane; 3:1); ^1^H NMR (600 MHz, CDCl_3_) δ 8.21 (d, *J* = 8.2 Hz, 1H), 7.82 (d, *J* = 7.6 Hz, 1H), 7.55 – 7.46 (m, 3H), 7.40 – 7.32 (m, 3H), 6.93 – 6.87 (m, 3H), 4.92 (s, 2H), 3.82 (s, 3H), 3.80 – 3.43 (m, 8H); ^13^C NMR (600 MHz, CDCl_3_) δ 170.7, 166.9, 161.3, 153.3, 134.7, 129.4, 127.8, 127.1, 126.8, 125.88, 125.87, 125.4, 121.7, 121.5, 114.0, 105.3, 68.4, 55.5, 45.8, 42.5; HPLC: t*_R_* = 13.01 min, purity 99.7%, Method B; HRMS (MALDI) m/z calcd for C_24_H_24_N_2_O_4_Na^+^ (M+Na^+^) 427.1628, found 427.1637.

### 1-(4-(2,3-Dimethoxybenzoyl)piperazin-1-yl)-2-(naphthalen-1-yloxy)ethan-1-one (5)

The title compound was synthesized according to the procedure described for **4** using 2,3-dimethoxybenzoic acid (20 mg, 0.110 mmol) and DMF (0.8 mL) as the solvent to afford 35 mg (73%) of a white solid after purification by flash chromatography (0-60% EtOAc in heptane). R_f_ = 0.25 (EtOAc:Heptane; 6:1); ^1^H NMR (600 MHz, CDCl_3_) δ 8.24 and 8.15 (2d, *J* = 7.6 Hz and *J* = 8.3 Hz, 1H), 7.86 – 7.76 (m, 1H), 7.57 – 7.41 (m, 3H), 7.36 (q, *J* = 8.1 Hz, 1H), 7.12 – 7.04 (m, 1H), 6.97 – 6.86 (m, 2H), 6.79 (d, *J* = 7.6 Hz, 1H), 4.97 – 4.81 (m, 2H), 4.11 – 3.92 (m, 2H), 3.91 – 3.76 (m, 6H), 3.73 – 3.46 (m, 4H), 3.35 – 3.30 (m, 1H), 3.15 – 3.12 (m, 1H); ^13^C NMR (151 MHz, CDCl_3_) δ 167.8, 167.7*, 166.92*, 166.87, 153.4, 152.8, 152.7, 145.1, 134.8, 130.8, 130.7, 127.8, 126.9, 126.8*, 125.94, 125.91, 125.87, 125.8, 125.4, 125.1, 121.7, 121.6, 119.3*, 119.2, 113.5, 113.4*, 105.3, 68.4*, 68.3, 61.7, 56.0, 47.4, 46.9, 46.2, 45.7, 42.7, 42.3, 42.2, 41.7; HPLC: t*_R_* = 12.89 min, purity > 99.99%, Method D; HRMS (MALDI) m/z calcd for C_25_H_26_N_2_O_5_Na^+^ (M+Na^+^) 457.1734, found 457.1731.

### 1-(4-(2,3-Dihydrobenzo[b][1,4]dioxine-6-carbonyl)piperazin-1-yl)-2-(naphthalen-1-yloxy)ethan-1-one (6)

The title compound was synthesized according to the procedure described for **4** using 2,3-dihydrobenzo[*b*][1,4]dioxine-6-carboxylic acid (17 mg, 91.6 μmol) and DMF (0.6 mL) as the solvent for the reaction to afford 13 mg (33%) of a white solid after purification by flash chromatography (0-10% MeOH in DCM). R_f_ = 0.43 (MeOH:DCM; 1:9); ^1^H NMR (600 MHz, DMSO-*d_6_*) δ 8.22 (d, *J* = 7.6 Hz, 1H), 7.87 (d, *J* = 7.6 Hz, 1H), 7.56 – 7.47 (m, 3H), 7.40 (t, *J* = 8.0 Hz, 1H), 6.97 – 6.89 (m, 4H), 5.05 (s, 2H), 4.27 (d, *J* = 1.8 Hz, 4H), 3.69 – 3.41 (m, 8H); ^13^C NMR (151 MHz, DMSO-*d_6_*) δ 168.7, 165.9, 153.4, 144.7, 143.0, 134.0, 128.4, 127.4, 126.5, 125.4, 124.8, 121.6, 120.5, 120.3, 116.9, 116.3, 105.6, 66.3, 64.0; HPLC: t*_R_* = 12.82 min, purity 95.03%, Method D; HRMS (MALDI) m/z calcd for C_25_H_24_N_2_O_5_Na^+^ (M+Na^+^) 455.1577, found 455.1573.

### 1-(4-(Benzo[d][1,3]dioxole-5-carbonyl)piperazin-1-yl)-2-(naphthalen-1-yloxy)ethan-1-one (7)

The title compound was synthesized according to the procedure described for **4** using benzo[*d*][1,3]dioxole-5-carboxylic acid (20 mg, 0.120 mmol) and DMF (0.8 mL) as the solvent for the reaction to afford 35 mg (70%) of a white solid after purification by flash chromatography (0-10% MeOH in DCM). R_f_ = 0.73 (MeOH:DCM; 1:9); ^1^H NMR (600 MHz, CDCl_3_) δ 8.20 (d, *J* = 8.6 Hz, 1H), 7.84 – 7.80 (m, 1H), 7.54 – 7.46 (m, 3H), 7.37 (t, *J* = 7.9 Hz, 1H), 6.93 – 6.84 (m, 3H), 6.80 (d, *J* = 7.9 Hz, 1H), 5.99 (s, 2H), 4.92 (s, 2H), 3.81 – 3.36 (m, 8H); ^13^C NMR (151 MHz, CDCl_3_) δ 170.2, 166.9, 153.2, 149.3, 147.9, 134.7, 127.8, 126.8, 125.9, 121.9, 121.7, 121.5, 108.4, 108.2, 105.3, 101.7, 68.3; HPLC: t*_R_* = 12.43 min, purity 98.2%, Method B; HRMS (MALDI) m/z calcd for C_24_H_22_N_2_O_5_Na^+^ (M+Na^+^) 441.1421, found 441.1424.

### 2-(4-Fluorophenyl)-1-(4-(2-(naphthalen-1-yloxy)acetyl)piperazin-1-yl)ethan-1-one (8)

Step 1: *tert*-Butyl 4-(2-(4-fluorophenyl)acetyl)piperazine-1-carboxylate (**Ea**): The target compound was synthesized according to the procedure described for **4** using 4-fluorophenylacetic acid (415 mg, 2.69 mmol), **A** (500 mg, 2.68 mmol), and DMF (4.0 mL) as the solvent for the reaction to afford 715 mg (83%) of a white solid without purification. R_f_ = 0.70 (MeOH:DCM; 2:8); ^1^H NMR (400 MHz, CDCl_3_) δ 7.29 – 7.16 (m, 2H), 7.05 – 6.96 (m, 2H), 3.70 (s, 2H), 3.63 – 3.56 (m, 2H), 3.46 – 3.34 (m, 4H), 3.29 – 3.21 (m, 2H), 1.44 (s, 9H); ^13^C NMR (101 MHz, CDCl_3_) δ 169.6, 163.2, 160.8, 154.6, 130.63*, 130.59*, 130.4*, 130.3*, 115.7 (d, *J* = 21.6 Hz), 80.5, 46.0, 41.8, 40.2, 28.5.

Step 2: 2-(4-Fluorophenyl)-1-(piperazin-1-yl)ethan-1-one (**Fa**): The target compound was synthesized according to the procedure described for **Da** using **Ea** (700 mg, 2.17 mmol) to yield 400 mg (83%) of a white solid. R_f_ = 0.77 (MeOH:DCM; 2:8); ^1^H NMR (400 MHz, CDCl_3_) δ 7.28 – 7.16 (m, 2H), 7.02 (t, *J* = 8.9 Hz, 1H), 3.69 (s, 2H), 3.64 (t, *J* = 5.2 Hz, 2H), 3.45 (t, *J* = 5.0 Hz, 2H), 2.84 (t, *J* = 5.2 Hz, 2H), 2.72 (t, *J* = 5.0 Hz, 2H), 2.42 (s, 1H); ^13^C NMR (101 MHz, CDCl_3_) δ 169.5, 160.7, 149.5, 130.4, 130.3, 115.7 (d, *J* = 21.6 Hz), 47.3, 46.2, 45.9, 43.0, 40.1.

Step 3: 2-Chloro-1-(4-(2-(4-fluorophenyl)acetyl)piperazin-1-yl)ethan-1-one (**Ga**): The target compound was synthesized according to the procedure described for **B** using **Fa** (390 mg, 1.75 mmol) to yield 485 mg (92%) of a yellow oil. The product was used in the next step without further purification. R_f_ = 0.70 (MeOH:DCM; 1:9); ^1^H NMR (400 MHz, CDCl_3_) δ 7.25 – 7.16 (m, 2H), 7.02 (t, *J* = 8.6 Hz, 2H), 4.06* (s, 1.3H’), 4.03* (s, 0.7H”), 3.72* (s, 3H’), 3.66* (s, 1H”), 3.58* (s, 1H’), 3.50* (s, 3H”), 3.47* (s, 1.3H’), 3.34* (s, 0.7H”); ^13^C NMR (101 MHz, CDCl_3_) δ 169.8*, 169.6*,165.6*, 165.4*, 163.3, 160.8, 130.3, 115.9 (d, *J* = 21.6 Hz), 46.2*, 46.1*, 45.6*, 42.1*, 41.8*, 41.4*, 40.8, 40.3*, 40.1*.

Step 4: The title compound was synthesized according to the procedure described for **Ca** using **Ga** (150 mg, 0.50 mmol) to afford 190 mg (93%) of a brown solid after purification by preparative HPLC (30-100% mobile phase B in mobile phase A in 10 min). R_f_ = 0.40 (EtOAc:Heptane; 9:1); ^1^H NMR (400 MHz, CDCl_3_) δ 8.22 – 8.10 (m, 1H), 7.86 – 7.79 (m, 1H), 7.57 – 7.44 (m, 3H), 7.36 (t, *J* = 8.0 Hz, 1H), 7.21 – 7.09 (m, 2H), 7.06 – 6.95 (m, 2H), 6.86 (d, *J* = 7.6 Hz, 1H), 4.92* (s, 1.2H’), 4.88* (s, 0.8H”), 3.73* (s, 1H’), 3.73* (s, 1H”), 3.70 – 3.59* (m, 3.5H’), 3.59 – 3.41* (m, 3.5H”), 3.38* (s, 0.5H’), 3.36* (s, 0.5H”); ^19^F NMR (376 MHz, CDCl_3_) δ −115.47*, −115.53*; ^13^C NMR (151 MHz, CDCl_3_) δ 170.7*, 170.6*, 167.6*, 167.4*, 163.0, 161.3, 153.1*, 152.9*, 134.7, 130.3*, 130.25*, 130.18*, 130.14*, 129.75*, 129.73*, 128.0*, 127.8*, 127.0, 126.0*, 125.9*, 125.8*, 125.3*, 125.2*, 121.9, 121.4*, 121.3*, 115.8 (d, *J* = 21.6 Hz), 105.3*, 105.2*, 68.2*, 67.9*, 46.4*, 45.9*, 45.5*, 45.4*, 42.4*, 42.3*, 42.2*, 41.9*, 40.0*, 39.9*; HPLC: t*_R_* = 7.06 min, purity 96.1%, Method F; HRMS (MALDI) m/z calcd for C_24_H_24_FN_2_O_3_^+^ (M+H^+^) 407.1765, found 407.1757.

### 2-Methyl-1-(4-(2-(naphthalen-1-yloxy)acetyl)piperazin-1-yl)-2-phenylpropan-1-one (9)

The title compound was synthesized according to the procedure described for **4** using 2-methyl-2-phenylpropanoic acid (24 mg, 0.15 mmol) and DMF (2.0 mL) as the solvent for the reaction to afford 47 mg (76%) of a white foam after purification by flash chromatography (50% EtOAc in heptane). R_f_ = 0.38 (EtOAc:Heptane; 7:3); ^1^H NMR (600 MHz, CDCl_3_) δ 8.14 (s, 1H), 7.83 – 7.79 (m, 1H), 7.54 – 7.44 (m, 3H), 7.36 – 7.29 (m, 3H), 7.24 – 7.18 (m, 3H), 6.82 (d, *J* = 7.6 Hz, 1H), 4.80 (s, 2H), 3.58* (s, 4.3H’), 3.07* (s, 3.7H”), 1.52 (s, 6H); ^13^C NMR (151 MHz, CDCl_3_) δ 175.2, 166.7, 153.3, 146.0, 134.7, 129.2, 127.7, 126.83, 126.76, 125.81, 125.76, 125.4, 124.8, 121.6, 121.5, 105.2, 68.1, 47.1, 45.2, 41.6, 28.3; HPLC: t*_R_* = 8.47 min, purity 99.7%, Method A; HRMS (MALDI) m/z calcd for C_26_H_29_N_2_O_3_^+^ (M+H^+^) 417.2172, found 417.2166.

### 2-(Naphthalen-1-yloxy)-1-(4-(1-phenylcyclopentane-1-carbonyl)piperazin-1-yl)ethan-1-one (10)

The title compound was synthesized according to the procedure described for **4** using 1-phenylcyclopentane-1-carboxylic acid (28 mg, 0.15 mmol) and DMF (2.0 mL) as the solvent for the reaction to afford 48 mg (73%) of a yellow solid after purification by flash chromatography (70% EtOAc in heptane). R_f_ = 0.31 (EtOAc:Heptane; 9:1); ^1^H NMR (400 MHz, CDCl_3_) δ 8.14 (s, 1H), 7.84 – 7.77 (m, 1H), 7.55 – 7.43 (m, 3H), 7.37 – 7.27 (m, 3H), 7.25 – 7.15 (m, 3H), 6.82 (d, *J* = 7.6 Hz, 1H), 4.79 (s, 2H), 3.58* (s, 4.3H’), 3.06* (s, 3.7H”), 2.43 – 2.32 (m, 2H), 2.05 – 1.92 (m, 2H), 1.79 – 1.63 (m, 4H); ^13^C NMR (151 MHz, CDCl_3_) δ 174.9, 166.7, 145.2, 134.7, 129.0, 127.7, 126.8, 126.6, 125.81, 125.76, 125.4, 125.1, 121.6, 105.2, 68.1, 58.5, 45.9*, 45.3*, 43.1*, 41.4*, 38.3, 25.3; HPLC: t*_R_* = 9.28 min, purity 97.2%, Method A; HRMS (MALDI) m/z calcd for C_28_H_31_N_2_O_3_^+^ (M+H^+^) 443.2329, found 443.2322.

### 1-(4-(2,3-Dihydrobenzofuran-2-carbonyl)piperazin-1-yl)-2-(naphthalen-1-yloxy)ethan-1-one (11)

The title compound was synthesized according to the procedure described for **4** using 2,3-dihydrobenzofuran-2-carboxylic acid (25 mg, 0.15 mmol) and DMF (2.0 mL) as the solvent for the reaction to afford 37 mg (60%) of a clear oil after purification by flash chromatography (70% EtOAc in heptane). R_f_ = 0.32 (EtOAc:Heptane; 7:3); ^1^H NMR (400 MHz, CDCl_3_) δ 8.28 – 8.18 (m, 1H), 7.87 – 7.78 (m, 1H), 7.56 – 7.46 (m, 3H), 7.38 (t, *J* = 8.0 Hz, 1H), 7.19 (d, *J* = 7.4 Hz, 1H), 7.15 – 7.07 (m, 1H), 6.95 – 6.84 (m, 2H), 6.81 – 6.66 (m, 1H), 5.38 – 5.23 (m, 1H), 4.93 (s, 2H), 3.98 – 3.86 (m, 2H), 3.85 – 3.75* (m, 3H’), 3.75 – 3.18 (m, 5H”); ^13^C NMR (101 MHz, CDCl_3_) δ 167.6, 167.0, 158.3, 153.3, 134.7, 128.2, 128.2, 126.8, 125.8, 125.4, 125.1, 121.7, 121.52, 121.47, 109.5, 105.3, 79.2*, 79.0*, 68.5*, 68.2*, 46.1*, 45.7*, 45.5*, 43.0*, 42.7*, 42.5*, 42.1*, 31.8; HPLC: t*_R_* = 7.71 min, purity 98.1%, Method A; HRMS (MALDI) m/z calcd for C_25_H_25_N_2_O_4_^+^ (M+H^+^) 417.1809, found 417.1801.

### 1-(4-(1-(4-Chlorophenyl)cyclobutane-1-carbonyl)piperazin-1-yl)-2-(naphthalen-1-yloxy)ethan-1-one (12)

The title compound was synthesized according to the procedure described for **4** using 1-(4-chlorophenyl)cyclobutane-1-carboxylic acid (32 mg, 0.15 mmol) and DMF (2.0 mL) as the solvent for the reaction to afford 49 mg (72%) of a white foam after purification by flash chromatography (70% EtOAc in heptane). R_f_ = 0.29 (EtOAc:Heptane; 7:3); ^1^H NMR (600 MHz, CDCl_3_) δ 8.27 – 8.01 (m, 1H), 7.83 – 7.79 (m, 1H), 7.57 – 7.43 (m, 3H), 7.37 – 7.31 (m, 3H), 7.29 – 7.22 (m, 2H), 6.86 – 6.81 (m, 1H), 4.84* (s, 1.3H’), 4.77* (s, 0.7H”), 3.57* (s, 4H’), 3.19 – 3.05* (m, 2H”), 2.98* (s, 1.3H’), 2.91* (s, 0.7H”), 2.83 – 2.75 (m, 2H), 2.37 – 2.25 (m, 2H), 2.01 – 1.92* (m, 1H’), 1.92 – 1.82* (m, 1H”); ^13^C NMR (151 MHz, CDCl_3_) δ 173.9, 166.8*, 166.6*, 153.2, 141.7, 134.7, 132.7, 129.4, 127.8, 126.8, 126.5, 125.8, 125.3, 121.7, 121.5, 105.2, 68.2, 51.9, 45.7*, 45.4*, 45.2*, 45.0*, 42.7*, 42.1*, 42.0*, 41.6*, 32.3, 15.5; HPLC: t*_R_* = 9.37 min, purity 99.1%, Method A; HRMS (MALDI) m/z calcd for C_27_H_28_ClN_2_O_3_^+^ (M+H^+^) 463.1783, found 463.1775.

### 1-(4-(1-(4-Chlorophenyl)cyclohexane-1-carbonyl)piperazin-1-yl)-2-(naphthalen-1-yloxy)ethan-1-one (13)

The title compound was synthesized according to the procedure described for **4** using 1-(4-chlorophenyl)cyclohexane-1-carboxylic acid (36 mg, 0.15 mmol) and DMF (2.0 mL) as the solvent for the reaction to afford 43 mg (59%) of a white foam after purification by flash chromatography (70% EtOAc in heptane). R_f_ = 0.69 (EtOAc:Heptane; 8:2); ^1^H NMR (600 MHz, CDCl_3_) δ 8.14 (d, *J* = 8.3 Hz, 1H), 7.83 – 7.78 (m, 1H), 7.54 – 7.44 (m, 3H), 7.34 (t, *J* = 8.0 Hz, 1H), 7.31 – 7.27 (m, 2H), 7.20 – 7.14 (m, 2H), 6.83 (d, *J* = 7.6 Hz, 1H), 4.80 (s, 2H), 3.80 – 2.84* (m, 8H), 2.22* (s, 1H’), 2.21* (s, 1H”), 1.71 – 1.59 (m, 7H), 1.30 – 1.21 (m, 1H); ^13^C NMR (151 MHz, CDCl_3_) δ 173.6, 166.7, 153.3, 144.4, 134.7, 132.6, 129.3, 127.8, 126.80, 126.76, 125.83, 125.82, 125.3, 121.6, 121.5, 105.3, 68.2, 50.9, 45.2, 41.7, 36.9, 25.8, 23.5; HPLC: t*_R_* = 10.62 min, purity 99.2%, Method A; HRMS (MALDI) m/z calcd for C_29_H_32_ClN_2_O_3_^+^ (M+H^+^) 491.2096, found 491.2089.

### 3-(4-Fluorophenyl)-1-(4-(2-(naphthalen-1-yloxy)acetyl)piperazin-1-yl)propan-1-one (14)

The title compound was synthesized according to the procedure described for **4** using 3-(4-fluorophenyl)propanoic acid (25 mg, 0.15 mmol) and DMF (2.0 mL) as the solvent for the reaction to afford 40 mg (69%) of a white foam after purification by preparative HPLC (40-100% mobile phase B in mobile phase A in 10 min). R_f_ = 0.39 (EtOAc:Heptane; 7:3); ^1^H NMR (400 MHz, CDCl_3_) δ 8.23 – 8.14 (m, 1H), 7.85 – 7.79 (m, 1H), 7.56 – 7.46 (m, 3H), 7.36 (t, *J* = 8.0 Hz, 1H), 7.17 – 7.09 (m, 2H), 6.95 (t, *J* = 8.5 Hz, 2H), 6.87 (d, *J* = 7.7 Hz, 1H), 4.90 (s, 2H), 3.70 – 3.55 (m, 4H’), 3.55 – 3.49* (m, 2H”), 3.35* (s, 1.3H’), 3.28* (s, 0.7H”), 2.96 – 2.88 (m, 2H), 2.65 – 2.51 (m, 2H); ^19^F NMR (376 MHz, CDCl_3_) δ −116.62*, −116.64*; ^13^C NMR (101 MHz, CDCl_3_) δ 171.5*, 171.4*, 167.3*, 167.1*, 162.9, 160.4, 153.1, 136.3, 134.7, 130.0, 129.9, 127.8, 126.9, 125.9*, 125.8*, 125.3, 121.8, 121.4, 115.6, 115.4, 105.2, 68.2*, 68.0*, 45.9*, 45.5*, 42.2*, 42.1*, 34.9, 30.7; HPLC: t*_R_* = 7.92 min, purity 95.4%, Method A; HRMS (MALDI) m/z calcd for C_25_H_26_FN_2_O_3_^+^ (M+H^+^) 421.1922, found 421.1915.

### 1-(4-(2-(Naphthalen-1-yloxy)acetyl)piperazin-1-yl)-3-(m-tolyl)propan-1-one (15)

The title compound was synthesized according to the procedure described for **4** using 3-(*m*-tolyl)propanoic acid (25 mg, 0.15 mmol) and DMF (2.0 mL) as the solvent for the reaction to afford 44 mg (71%) of a clear oil after purification by flash chromatography (70% EtOAc in heptane). R_f_ = 0.34 (EtOAc:Heptane; 7:3); ^1^H NMR (600 MHz, CDCl_3_) δ 8.23 – 8.16 (m, 1H), 7.84 – 7.80 (m, 1H), 7.54 – 7.46 (m, 3H), 7.36 (t, *J* = 8.0 Hz, 1H), 7.18 – 7.10 (m, 1H), 7.02 – 6.96 (m, 3H), 6.88 (d, *J* = 7.7 Hz, 1H), 4.90* (s, 1.3H’), 4.88* (s, 0.7H”), 3.69 – 3.51* (m, 4H’), 3.46* (s, 2H”), 3.32* (s, 1.3H’), 3.26* (s, 0.7H”), 2.94 – 2.89 (m, 2H), 2.64 – 2.57* (m, 1.3H’), 2.57 – 2.52* (m, 0.7H”), 2.31 – 2.26 (m, 3H); ^13^C NMR (151 MHz, CDCl_3_) δ 171.1*, 171.0*, 166.9*, 166.8*, 153.2, 140.9, 138.3, 134.7, 129.4, 128.6, 127.8, 127.2, 126.8, 125.9, 125.8, 125.5, 125.3, 121.7, 121.5, 105.2, 68.4*, 68.1*, 45.8*, 45.5*, 45.3*, 42.1*, 41.9*, 41.4*, 35.0, 31.5, 21.4; HPLC: t*_R_* = 8.21 min, purity 99.1%, Method A; HRMS (MALDI) m/z calcd for C_26_H_29_N_2_O_3_^+^ (M+H^+^) 417.2172, found 417.2167.

### 3-(2-Methoxyphenyl)-1-(4-(2-(naphthalen-1-yloxy)acetyl)piperazin-1-yl)propan-1-one (16)

The title compound was synthesized according to the procedure described for **4** using 3-(2-methoxyphenyl)propanoic acid (27 mg, 0.15 mmol) and DMF (2.0 mL) as the solvent for the reaction to afford 47 mg (73%) of a white foam after purification by flash chromatography (70% EtOAc in heptane). R_f_ = 0.21 (EtOAc:Heptane; 7:3); ^1^H NMR (400 MHz, CDCl_3_) δ 8.20 (d, *J* = 7.7 Hz, 1H), 7.82 (d, *J* = 7.4 Hz, 1H), 7.56 – 7.45 (m, 3H), 7.36 (t, *J* = 8.0 Hz, 1H), 7.20 – 7.09 (m, 2H), 6.93 – 6.73 (m, 3H), 4.90 (s, 2H), 3.81* (s, 1.8H’), 3.76* (s, 1.2H), 3.64 – 3.52* (m, 4H’), 3.47* (s, 2H”), 3.37* (s, 1.3H’), 3.30* (s, 0.7H”), 2.92 (t, *J* = 7.8 Hz, 2H), 2.64 – 2.52 (m, 2H); ^13^C NMR (151 MHz, CDCl_3_) δ 171.8*, 171.6*, 166*.9, 166.7*, 157.5, 153.3, 134.7, 130.5, 129.0, 127.9, 127.8, 126.8, 125.9, 125.8, 125.4, 121.7, 121.5, 120.7, 110.4, 105.3, 68.5*, 68.2*, 55.4, 45.9*, 45.6*, 45.4*, 42.3*, 41.9*, 41.4*, 33.4, 27.2; HPLC: t*_R_* = 8.13 min, purity 97.8%, Method A; HRMS (MALDI) m/z calcd for C_26_H_29_N_2_O_4_^+^ (M+H^+^) 433.2122, found 433.2115.

### 4-(2-(Naphthalen-1-yloxy)acetyl)-N-phenylpiperazine-1-carboxamide (17)

A solution of isocyanatobenzene (0.025 mL, 0.20 mmol) and **Dd** (50 mg, 0.18 mmol) in DCM (2 mL) was stirred at rt for 3 h and concentrated under reduced pressure. The residue was purified by flash chromatography (SiO_2_, 80% EtOAc in heptane) to afford the title compound (35 mg, 49%) as a pink oil. R_f_ = 0.57 (EtOAc:Heptane; 8:2); ^1^H NMR (600 MHz, CDCl_3_) δ 8.25 – 8.19 (m, 1H), 7.85 – 7.79 (m, 1H), 7.55 – 7.47 (m, 2H), 7.37 (t, *J* = 7.9 Hz, 1H), 7.33 – 7.25 (m, 5H), 7.08 – 7.02 (m, 1H), 6.90 (d, *J* = 7.6 Hz, 1H), 6.31 (s, 1H), 4.93 (s, 2H), 3.78 – 3.71* (m, 4H’), 3.50 – 3.45* (m, 4H”); ^13^C NMR (151 MHz, CDCl_3_) δ 167.0, 154.9, 153.3, 138.6, 134.8, 129.1, 127.9, 126.9, 125.9, 125.4, 123.7, 121.7, 121.6, 121.1, 120.2, 105.3, 68.3, 45.4, 44.2, 42.0; HPLC: t*_R_* = 7.23 min, purity 95.0%, Method A; HRMS (MALDI) m/z calcd for C_23_H_24_N_3_O_3_^+^ (M+H^+^) 390.1812, found 390.1807.

### 2-Methyl-1-(4-(2-(naphthalen-1-yloxy)acetyl)piperazin-1-yl)-4-phenylbutane-1,4-dione (18)

The title compound was synthesized according to the procedure described for **4** using 2-methyl-4-oxo-4-phenylbutanoic acid (29 mg, 0.15 mmol) to afford 37 mg (57%) of a faint orange solid after purification by flash chromatography (20-75% EtOAc in heptane). R_f_ = 0.25 (EtOAc:Heptane; 4:1); ^1^H NMR (600 MHz, CDCl_3_) δ 8.27 – 8.20 (m, 1H), 7.95 (d, *J* = 8.1 Hz, 2H), 7.85 – 7.79 (m, 1H), 7.57 – 7.47 (m, 4H), 7.44 (t, *J* = 7.7 Hz, 2H), 7.41 – 7.34 (m, 1H), 6.95 – 6.89 (m, 1H), 4.93 (s, 2H), 3.91 – 3.31 (m, 10H), 2.96 – 2.88 (m, 1H), 1.22 – 1.14 (m, 3H); ^13^C NMR (151 MHz, CDCl_3_) δ 199.0, 166.9*, 166.8*, 153.4, 136.7, 134.7, 128.7, 128.2, 127.9*, 127.8*, 126.8, 125.9, 125.4, 121.7*, 121.6*, 105.4*, 105.3*, 68.5*, 68.2*, 46.2*, 45.9*, 45.7*, 43.1*, 42.5*, 42.4*, 41.9*, 31.1, 18.0; HPLC: t*_R_* = 13.53 min, purity 98.9%, Method D; HRMS (MALDI) m/z calcd for C_27_H_28_N_2_O_4_Na^+^ (M+Na^+^) 467.1941, found 467.1940.

### 1-(4-Chlorophenyl)-4-(4-(2-(naphthalen-1-yloxy)acetyl)piperazin-1-yl)butane-1,4-dione (19)

The title compound was synthesized according to the procedure described for **4** using 4-(4-chlorophenyl)-4-oxobutanoic acid (32 mg, 0.15 mmol) and DMF (2.0 mL) as the solvent for the reaction to afford 51 mg (74%) of a yellow oil after purification by flash chromatography (70% EtOAc in heptane). R_f_ = 0.32 (EtOAc:Heptane; 8:2); ^1^H NMR (600 MHz, CDCl_3_) δ 8.23 – 8.19 (m, 1H), 7.92 (d, *J* = 8.4 Hz, 2H), 7.84 – 7.79 (m, 1H), 7.53 – 7.46 (m, 3H), 7.45 – 7.40 (m, 2H), 7.40 – 7.33 (m, 1H), 6.93 – 6.87 (m, 1H), 4.92 (s, 2H), 3.79 – 3.61* (m, 4H’), 3.61 – 3.47* (m, 4H”), 3.28 (t, *J* = 6.3 Hz, 2H), 2.79 – 2.73* (m, 1.3H’), 2.70* (s, 0.7H”); ^13^C NMR (151 MHz, CDCl_3_) δ 197.9, 170.4*, 170.3*, 166.9*, 166.8*, 153.3, 139.7, 135.1, 134.7, 129.6, 129.0, 127.8, 126.8, 125.9, 125.4, 121.7, 121.5, 105.2, 68.4*, 68.2*, 45.7*, 45.6*, 45.5*, 45.2*, 42.2*, 42.1*, 41.6*, 33.5, 27.1; HPLC: t*_R_* = 8.21 min, purity 95.6%, Method A; HRMS (MALDI) m/z calcd for C_26_H_26_ClN_2_O_4_^+^ (M+H^+^) 465.1575, found 465.1569.

### 1-(4-Acetylpiperazin-1-yl)-2-(naphthalen-1-yloxy)ethan-1-one (20)

Step 1: *tert*-Butyl 4-(2-(naphthalen-1-yloxy)acetyl)piperazine-1-carboxylate (**25**): A round bottom flask was charged with **B** (850 mg, 3.24 mmol), naphthalen-1-ol (700 mg, 4.86 mmol), Cs_2_CO_3_ (2.11 g, 6.5 mmol), KI (27 mg, 0.16 mmol), and acetone (25 mL). The reaction mixture was stirred at 35 °C for 24 h. The solvent was evaporated, and the residue was suspended in EtOAc (20 mL). The organic solution was washed with aq. NaOH solution (1 M, 2 x 10 mL) and brine (10 mL). The organic phase was dried over Na_2_SO_4_, filtered and the solvent was evaporated under reduced pressure. The residue was purified by flash chromatography (60% EtOAc in heptane) to afford the target compound (0.86 g, 72%) as a red oil. R_f_ = 0.43 (EtOAc:Heptane; 1:1); ^1^H NMR (400 MHz, CDCl_3_) δ 8.26 – 8.18 (m, 1H), 7.86 – 7.77 (m, 1H), 7.57 – 7.45 (m, 3H), 7.36 (t, *J* = 8.0 Hz, 1H), 6.89 (d, *J* = 0.9 Hz, 1H), 4.90 (s, 2H), 3.68 – 3.59 (m, 4H), 3.45 – 3.35 (m, 4H), 1.44 (s, 9H); ^13^C NMR (101 MHz, CDCl_3_) δ 166.8, 154.5, 153.4, 134.7, 127.8, 126.8, 125.9, 125.8, 125.4, 121.62, 121.59, 105.3, 80.5, 68.3, 45.6, 42.2, 28.5; HPLC: t*_R_* = 8.4 min, purity 95%, Method B; HRMS calcd for C_21_H_27_N_2_O_4_^+^ (M+H^+^) 371.1965, found 371.1959.

Step 2: 2-(Naphthalen-1-yloxy)-1-(piperazin-1-yl)ethan-1-one (**Dd**): To a solution of **25** (800 mg, 2.16 mmol) in DCM (5 mL) was added TFA (3.5 mL, 43.2 mmol). The mixture was stirred at rt for 30 min and evaporated to dryness. The residue was dissolved in DCM (10 mL), and the solution was washed with aq. Na_2_CO_3_ (2 M, 2 x10 mL). The organic layer was dried over Na_2_SO_4_, filtered, and evaporated to dryness to afford the target compound (568 mg, 97%) as a black oil. The product was used in the next reaction without further purification. R_f_ = 0.66 (MeOH:DCM; 2:8); ^1^H NMR (600 MHz, CDCl_3_) δ 8.25 (d, *J* = 1.8 Hz, 1H), 7.81 (d, *J* = 1.8 Hz, 1H), 7.53 – 7.45 (m, 3H), 7.37 (t, *J* = 7.9 Hz, 1H), 6.89 (d, *J* = 7.6 Hz, 1H), 4.88 (s, 2H), 3.64 (t, *J* = 5.3 Hz, 4H), 2.86 – 2.79 (m, 4H), 1.89 (s, 1H); ^13^C NMR (151 MHz, CDCl_3_) δ 166.6, 153.5, 134.7, 127.7, 126.7, 125.9, 125.7, 125.5, 121.8, 121.4, 105.3, 68.3, 46.9, 46.5, 46.0, 43.4; HRMS calcd for C_16_H_19_N_2_O_2_^+^ (M+H^+^) 271.1441, found 271.1437.

Step 3: The title compound was synthesized according to the procedure described for **2** using **Dd** (75 mg, 0.28 mmol) and acetyl chloride (40 µL, 0.56 mmol) to afford 26 mg (30%) of a clear oil after purification by preparative HPLC (40-100% mobile phase B in mobile phase A in 10 min). R_f_ = 0.64 (MeOH:DCM; 1:9); ^1^H NMR (400 MHz, CDCl_3_) δ 8.25 – 8.16 (m, 1H), 7.85 – 7.78 (m, 1H), 7.55 – 7.45 (m, 3H), 7.36 (t, *J* = 8.0 Hz, 1H), 6.89 (d, *J* = 7.6 Hz, 1H), 4.91 (s, 2H), 3.73 – 3.61* (m, 4H’), 3.61 – 3.50* (m, 2H”), 3.45 – 3.39* (m, 1H’), 3.39 – 3.30* (m, 1H”), 2.09* (s, 1.8H’), 2.04* (s, 1.2H”); ^13^C NMR (101 MHz, CDCl_3_) δ 169.3*, 169.2*, 167.0*, 166.9*, 134.7, 127.8, 126.8, 125.9, 125.4, 121.7, 121.5, 105.3, 68.4*, 68.2*, 46.5*, 46.1*, 45.7*, 45.5*, 42.3*, 42.1*, 41.7*, 41.2*, 32.0*, 29.8*, 22.8*, 21.4*; HPLC: t*_R_* = 5.66 min, purity 99.4%, Method A; HRMS (MALDI) m/z calcd for C_18_H_21_N_2_O_3_^+^ (M+H^+^) 313.1546, found 313.1543.

### 1-(4-(2-(Naphthalen-1-yloxy)acetyl)piperazin-1-yl)butan-1-one (21)

The title compound was synthesized according to the procedure described for **2** using **Dd** (50 mg, 0.18 mmol) and butyryl chloride (30 µL, 0.29 mmol) to afford 52 mg (83%) of a clear oil after purification by flash chromatography (70% EtOAc in heptane). R_f_ = 0.21 (EtOAc:Heptane; 7:3); ^1^H NMR (600 MHz, CDCl_3_) δ 8.24 – 8.18 (m, 1H), 7.83 – 7.78 (m, 1H), 7.53 – 7.45 (m, 3H), 7.36 (t, *J* = 8.0 Hz, 1H), 6.89 (d, *J* = 7.6 Hz, 1H), 4.90 (s, 2H), 3.65* (s, 4H’), 3.57* (s, 2H”), 3.43* (s, 1.3H’), 3.37* (s, 0.7H”) 2.31 – 2.20 (m, 2H), 1.74 – 1.53 (m, 2H), 1.03 – 0.81 (m, 3H); ^13^C NMR (151 MHz, CDCl_3_) δ 171.8*, 171.7*, 166.9*, 166.8*, 153.3, 134.7, 127.8, 126.8, 125.9, 125.8, 125.3, 121.6, 121.5, 105.2, 68.4*, 68.1*, 45.7*, 45.5*, 45.3*, 42.3*, 41.8*, 41.2*, 35.2, 18.7, 14.0; HPLC: t*_R_* = 6.77 min, purity 99.4%, Method A; HRMS (MALDI) m/z calcd for C_20_H_25_N_2_O_3_^+^ (M+H^+^) 341.1859, found 341.1855.

### (*E*)-1-(4-(2-(Naphthalen-1-yloxy)acetyl)piperazin-1-yl)but-2-en-1-one (22)

To a solution of (*E*)-but-2-enoic acid (16 mg, 0.19 mmol) and 2-(naphthalen-1-yloxy)-1-(piperazin-1-yl)ethan-1-one (50 mg, 0.18 mmol) in DMF (2 mL) were added EDC·HCl (39 mg, 0.20 mmol), HOBt (31 mg, 0.20 mmol), and DIPEA (0.07 mL, 0.40 mmol). The reaction mixture was stirred overnight at rt and then partitioned between EtOAc and brine. The organic layer was separated, dried over anhydrous MgSO_4_, filtered, and concentrated *in vacuo*. The residue was purified by flash column chromatography (EtOAc:Heptane; 8:2) to give the title compound (47 mg, 75%) as a white solid. R_f_ = 0.52 (EtOAc:Heptane; 8:2); ^1^H NMR (400 MHz, CDCl_3_) δ 8.25 – 8.17 (m, 1H), 7.86 – 7.77 (m, 1H), 7.56 – 7.45 (m, 3H), 7.37 (t, *J* = 8.0 Hz, 1H), 6.95 – 6.82 (m, 2H), 6.20 (d, *J* = 14.7 Hz, 1H), 4.92 (s, 2H), 3.73 – 3.63* (m, 4.3H’), 3.57* (s, 3.7H”), 1.90 – 1.84 (m, 3H); ^13^C NMR (101 MHz, CDCl_3_) δ 166.9, 165.9, 153.3, 142.9, 134.7, 127.8, 126.8, 125.9, 125.4, 121.7, 121.5, 121.0, 105.3, 68.3, 45.7, 42.4, 18.4; HPLC: t*_R_* = 6.58 min, purity 99.7%, Method A; HRMS calcd for C_20_H_23_N_2_O_3_ (M+H^+^) 339.1703, found 339.1700.

### 2-Methoxy-1-(4-(2-(naphthalen-1-yloxy)acetyl)piperazin-1-yl)ethan-1-one (23)

The title compound was synthesized according to the procedure described for **2** using **Dd** (50 mg, 0.18 mmol) and 2-methoxyacetyl chloride (25 µL, 0.27 mmol) to afford 39 mg (61%) of a yellow oil after purification by flash chromatography (5% MeOH in DCM). R_f_ = 0.56 (MeOH:DCM; 5:95); ^1^H NMR (400 MHz, CDCl_3_) δ 8.24 – 8.16 (m, 1H), 7.85 – 7.78 (m, 1H), 7.56 – 7.45 (m, 3H), 7.36 (t, *J* = 8.0 Hz, 1H), 6.90 (d, *J* = 7.6 Hz, 1H), 4.91 (s, 2H), 4.07* (s, 1.3H’), 4.04* (s, 0.7H”), 3.72 – 3.63* (m, 4H’), 3.61 – 3.51* (m, 2H”), 3.51 – 3.41* (m, 2H’), 3.39* (s, 1.8H’), 3.37* (s, 1.2H”); ^13^C NMR (101 MHz, CDCl_3_) δ 167.8, 166.9, 153.2, 134.7, 127.8, 126.8, 125.9, 125.4, 121.7, 121.5, 105.3, 72.2*, 72.1*, 68.4*, 68.2*, 59.2*, 45.9*, 45.5*, 44.9*, 42.5*, 42.2*, 41.7*; HPLC: t*_R_* = 5.67 min, purity 99.0%, Method A; HRMS (MALDI) m/z calcd for C_19_H_23_N_2_O_4_^+^ (M+H^+^) 343.1652, found 343.1647.

### 3-Methyl-1-(4-(2-(naphthalen-1-yloxy)acetyl)piperazin-1-yl)butan-1-one (24)

The title compound was synthesized according to the procedure described for **2** using **Dd** (50 mg, 0.18 mmol) and 3-methylbutanoyl chloride (35 µL, 0.28 mmol) to afford 48 mg (72%) of a clear oil after purification by flash chromatography (30% acetone in heptane). R_f_ = 0.48 (Acetone:Heptane; 1:1); ^1^H NMR (400 MHz, CDCl_3_) δ 8.25 – 8.18 (m, 1H), 7.85 – 7.78 (m, 1H), 7.55 – 7.45 (m, 3H), 7.36 (t, *J* = 8.0 Hz, 1H), 6.89 (d, *J* = 7.6 Hz, 1H), 4.91 (s, 2H), 3.72 – 3.62* (m, 4H’), 3.59* (s, 2H”), 3.45* (s, 1.3H’), 3.39* (s, 0.7H”), 2.23 – 2.05 (m, 3H), 1.00 – 0.91 (m, 6H); ^13^C NMR (101 MHz, CDCl_3_) δ 171.3, 166.9, 153.3, 134.7, 127.8, 126.8, 125.8, 125.4, 121.7, 121.5, 105.3, 68.3, 45.7, 42.4, 42.1, 25.8, 22.8; HPLC: t*_R_* = 7.23 min, purity 99.1%, Method A; HRMS (MALDI) m/z calcd for C_21_H_27_N_2_O_3_^+^ (M+H^+^) 355.2016, found 355.2011.

### 3,3-Dimethyl-1-(4-(2-(naphthalen-1-yloxy)acetyl)piperazin-1-yl)butan-1-one (27)

The title compound was synthesized according to the procedure described for **2** using **Dd** (50 mg, 0.18 mmol) and *tert*-butylacetyl chloride (40 µL, 0.29 mmol) to afford 42 mg (62%) of a clear oil after purification by flash chromatography (70% EtOAc in heptane). R_f_ = 0.38 (EtOAc:Heptane; 7:3); ^1^H NMR (400 MHz, CDCl_3_) δ 8.26 – 8.17 (m, 1H), 7.86 – 7.78 (m, 1H), 7.56 – 7.45 (m, 3H), 7.36 (t, *J* = 8.0 Hz, 1H), 6.89 (d, *J* = 7.6 Hz, 1H), 4.91 (s, 2H), 3.70 – 3.63* (m, 4H’), 3.63 – 3.56* (m, 2H”), 3.49* (s, 1.3H’), 3.42* (s, 0.7H”), 2.24* (s, 1.3H’), 2.20* (s, 0.7H”), 1.03 (s, 9H); ^13^C NMR (151 MHz, CDCl_3_) δ 170.7, 166.9, 153.3, 134.7, 127.8, 126.8, 125.8, 125.4, 121.7, 121.5, 105.3, 68.3, 45.7, 44.8, 42.3, 31.6, 30.1; HPLC: t*_R_* = 7.72 min, purity 97.6%, Method A; HRMS (MALDI) m/z calcd for C_22_H_29_N_2_O_3_^+^ (M+H^+^) 369.2172, found 369.2167.

### 2-Cyclopentyl-1-(4-(2-(naphthalen-1-yloxy)acetyl)piperazin-1-yl)ethan-1-one (28)

The title compound was synthesized according to the procedure described for **2** using **Dd** (50 mg, 0.18 mmol) and 2-cyclopentylacetyl chloride (40 µL, 0.30 mmol) to afford 52 mg (72%) of a clear oil after purification by flash chromatography (70% EtOAc in heptane). R_f_ = 0.42 (EtOAc:Heptane; 7:3); ^1^H NMR (600 MHz, CDCl_3_) δ 8.24 – 8.18 (m, 1H), 7.83 – 7.79 (m, 1H), 7.54 – 7.45 (m, 3H), 7.36 (t, *J* = 7.9 Hz, 1H), 6.89 (d, *J* = 7.6 Hz, 1H), 4.91 (s, 2H), 3.65* (s, 4H’), 3.58* (s, 2H”), 3.45* (s, 1.3H’), 3.39* (s, 0.7H”), 2.38 – 2.31* (m, 1.3H’), 2.31 – 2.25* (m, 0.7H”), 2.25 – 2.13 (m, 1H), 1.88 – 1.77 (m, 2H), 1.66 – 1.57* (m, 2H’), 1.57 – 1.49* (m, 2H”), 1.19 – 1.05 (m, 2H); ^13^C NMR (151 MHz, CDCl_3_) δ 171.6*, 171.5*, 167.0*, 166.8*, 153.3, 134.7, 127.9*, 127.8*, 126.8, 125.9, 125.8, 125.4, 121.64, 121.55, 105.3*, 105.2*, 68.5*, 68.1*, 45.9*, 45.8*, 45.6*, 45.5*, 42.4*, 42.3*, 41.8*, 41.3*, 39.3, 36.7, 32.8, 25.0; HPLC: t*_R_* = 8.12 min, purity 99.9%, Method A; HRMS (MALDI) m/z calcd for C_23_H_29_N_2_O_3_^+^ (M+H^+^) 381.2172, found 381.2166.

### 1-(4-(2-(Naphthalen-1-yloxy)acetyl)piperazin-1-yl)hexan-1-one (29)

The title compound was synthesized according to the procedure described for **2** using **Dd** (75 mg, 0.28 mmol) and hexanoyl chloride (80 µL, 0.57 mmol) to afford 60 mg (59%) of a clear oil after purification by flash chromatography (80% EtOAc in heptane). R_f_ = 0.64 (EtOAc:Heptane; 8:2); ^1^H NMR (400 MHz, CDCl_3_) δ 8.25 – 8.17 (m, 1H), 7.85 – 7.78 (m, 1H), 7.55 – 7.45 (m, 3H), 7.36 (t, *J* = 8.0 Hz, 1H), 6.89 (d, *J* = 7.6 Hz, 1H), 4.91 (s, 2H), 3.72 – 3.61* (m, 4H’), 3.61 – 3.54* (m, 2H”), 3.48 – 3.41* (m, 1.3H’), 3.41 – 3.35* (m, 0.7H”), 2.34 – 2.19 (m, 2H), 1.65 – 1.55 (m, 2H), 1.37 – 1.21 (m, 4H), 0.94 – 0.85 (m, 3H); ^13^C NMR (151 MHz, CDCl_3_) δ 172.0*, 171.9*, 167.0*, 166.8*, 153.3, 134.7, 127.8, 126.8, 125.9, 125.8, 125.4, 121.7, 121.5, 105.3, 68.5*, 68.2*, 45.8*, 45.6*, 45.4*, 42.4*, 41.8*, 41.3*, 33.3, 31.7, 25.0, 22.4, 14.2; HPLC: t*_R_* = 8.05 min, purity 99.9%, Method A; HRMS (MALDI) m/z calcd for C_22_H_29_N_2_O_3_^+^ (M+H^+^) 369.2172, found 369.2167.

### 5-Methyl-1-(4-(2-(naphthalen-1-yloxy)acetyl)piperazin-1-yl)hexan-1-one (30)

To a flame-dried round bottom flask was added 5-methylhexanoic acid (25 mg, 0.19 mmol), 2-(naphthalen-1-yloxy)-1-(piperazin-1-yl)ethan-1-one (52 mg, 0.19 mmol), EDC · HCl (39 mg, 0.21 mmol) and HOBt hydrate (31 mg, 0.23 mmol) to anhydrous CH_2_Cl_2_ (4 mL). The mixture was put under an atmosphere of argon. After 10 min, DIPEA (70 μL, 0.38 mmol) was added at 0°C. The ice bath was removed. The resulting mixture was stirred at rt for 24 hrs. The mixture was then diluted with CH_2_Cl_2_ (3 mL) and the organic phase was washed with 1 M HCl (2 x 10 mL), aq. NaOH (10 mL) and brine (10 mL), dried over anhydrous MgSO_4_, filtered and evaporated *in vacuo*. The crude residue was purified by flash column chromatography (DCM:MeOH, 49:1), affording the title compound (49 mg, 67%) as an orange oil. ^1^H NMR (600 MHz, CDCl_3_) δ 8.15 – 8.13 (m, 1 H), 7.75 – 7.74 (m, 1H), 7.46 – 7.40 (m, 3H), 7.30 (t, *J =* 8.0, 1H), 6.83 (d, *J* = *7*.6, 1H), 4.85 (s, 2H), 3.61 – 3.52 (m, 6H), 3.37 – 3.32 (m, 2H), 1.54 – 1.50 (m, 2H), 1.24 – 1.12 (m, 3H), 0.88 – 0.86 (m, 6H); ^13^C NMR (151 MHz, CDCl_3_) δ 171.8, 171.6*, 166.8, 166.6*, 153.1, 153.0*, 134.5, 127.7*, 127.6, 126.6, 125.8*, 125.7*, 125.6, 125.2, 121.5, 121.4, 121.2*, 105.0, 68.3*, 68.0, 45.7*, 45.6*, 45.4*, 45.2*, 42.2*, 42.1*, 41.6*, 41.1*, 38.6, 33.4, 27.8, 23.0, 22.4; HPLC: t*_R_* = 14.01 min, purity 98.31%, Method D; HRMS (MALDI) m/z calcd for C_23_H_31_N_2_O_3_ (M+H^+^) 383.2329, found 383.2339.

### 1-(4-(Adamantane-1-carbonyl)piperazin-1-yl)-2-(naphthalen-1-yloxy)ethan-1-one (31)

The title compound was synthesized according to the procedure described for **4** using adamantane-1-carboxylic acid (27 mg, 0.15 mmol) and DMF (2.0 mL) as the solvent for the reaction to afford 20 mg (31%) of a clear oil after purification by flash chromatography (70% EtOAc in heptane). R_f_ = 0.45 (EtOAc:Heptane; 8:2); ^1^H NMR (400 MHz, CDCl_3_) δ 8.27 – 8.19 (m, 1H), 7.85 – 7.78 (m, 1H), 7.56 – 7.45 (m, 3H), 7.36 (t, *J* = 8.0 Hz, 1H), 6.89 (d, *J* = 7.6 Hz, 1H), 4.90 (s, 2H), 3.72 – 3.58* (m, 8H), 2.06 – 2.00 (m, 3H), 1.98 – 1.92 (m, 6H), 1.78 – 1.64 (m, 6H); ^13^C NMR (101 MHz, CDCl_3_) δ 176.1, 166.8, 153.3, 134.7, 127.8, 126.8, 125.9, 125.8, 125.4, 121.62, 121.61, 105.3, 68.2, 45.9*, 45.6*, 45.2*, 42.6*, 41.9, 39.2, 36.7, 28.5; HPLC: t*_R_* = 9.65 min, purity 98.3%, Method A; HRMS (MALDI) m/z calcd for C_27_H_33_N_2_O_3_^+^ (M+H^+^) 433.2485, found 433.2477.

### 1-(4-(3,5-Dimethyladamantane-1-carbonyl)piperazin-1-yl)-2-(naphthalen-1-yloxy)ethan-1-one (32)

The title compound was synthesized according to the procedure described for **4** using 3,5-dimethyladamantane-1-carboxylic acid (29 mg, 0.13 mmol) to afford 28 mg (46%) of a white solid after purification by flash chromatography (20-80% EtOAc in heptane). R_f_ = 0.55 (EtOAc:Heptane; 7:2); ^1^H NMR (600 MHz, CDCl_3_) δ 8.23 (d, *J* = 8.8 Hz, 1H), 7.82 (d, *J* = 7.6 Hz, 1H), 7.54 – 7.46 (m, 3H), 7.37 (t, *J* = 7.9 Hz, 1H), 6.89 (d, *J* = 7.6 Hz, 1H), 4.91 (s, 2H), 3.78 – 3.45 (m, 8H), 2.12 (s, 1H), 1.77 (s, 2H), 1.61 – 1.56 (m, 2H), 1.55 – 1.50 (m, 2H), 1.38 – 1.30 (m, 4H), 1.12 – 1.00 (m, 2H), 0.84 (s, 6H), ^13^C NMR (151 MHz, CDCl_3_) δ 175.9, 166.9, 153.4, 134.7, 127.8, 126.8, 125.9, 125.8, 125.4, 121.7*, 121.6*, 105.3, 68.3, 50.8, 45.9*, 45.6*, 45.4*, 45.3*, 43.8*, 42.9*, 42.6*, 37.8, 31.4, 30.8, 29.6; HPLC: t*_R_* = 12.07 min, purity 96.7%, Method C; HRMS (MALDI) m/z calcd for C_29_H_36_N_2_O_3_Na^+^ (M+Na^+^) 483.2618, found 483.2616.

### Methyl 4-(4-(2-(naphthalen-1-yloxy)acetyl)piperazine-1-carbonyl)bicyclo[2.2.2]octane-1-carboxylate (33)

A flame-dried round bottom flask was charged with 2-(naphthalen-1-yloxy)-1-(piperazin-1-yl)ethan-1-one (60 mg, 0.22 mmol), 4-(methoxycarbonyl)bicyclo[2.2.2]octane-1-carboxylic acid (47 mg, 0.22 mmol), EDC · HCl (47 mg, 0.24 mmol) and HOBt hydrate (37 mg, 0.24 mmol). Then, anhydrous DCM (1 mL) and DIPEA (77 μL, 0.44 mmol) were added. The resulting mixture was stirred at rt for overnight. The mixture was then diluted with EtOAc (2 mL) and water (2 mL). The organic phase was separated, and the aqueous phase was further extracted with EtOAc (2 x 2 mL). The combined organic phase was concentrated, and the crude was purified by flash chromatography (60% EtOAc in heptane) to afford the title compound (58 mg, 56%) as a colourless oil. R_f_ = 0.28 (EtOAc:Heptane; 4:1); ^1^H NMR (600 MHz, CDCl_3_) δ 8.25 – 8.14 (m, 1H), 7.86 – 7.77 (m, 1H), 7.56 – 7.46 (m, 3H), 7.37 (t, *J* = 8.0 Hz, 1H), 6.89 (d, *J* = 7.7 Hz, 1H), 4.91 (s, 2H), 3.69 – 3.61 (m, 9H), 3.60 – 3.56 (m, 2H), 1.88 – 1.79 (m, 12H); ^13^C NMR (151 MHz, CDCl_3_) δ 177.8, 175.5, 166.9, 153.3, 134.7, 127.8, 126.8, 125.9, 125.9, 125.4, 121.7, 121.6, 105.3, 68.3, 51.9, 45.8, 45.6, 45.2, 42.4, 39.8, 38.9, 28.1, 27.9; HPLC: t*_R_* = 6.67 min, purity 97.50%, Method I; HRMS (MALDI): m/z calcd for C_27_H_33_N_2_O_5_ (M+H^+^) 465.2384, found 465.2394.

### 4-(4-(2-(Naphthalen-1-yloxy)acetyl)piperazine-1-carbonyl)bicyclo[2.2.2]octane-1-carboxylic acid (34)

To a solution of **33** (19.7 mg, 42.2 μmol) in THF (212 μL) was added aq. LiOH (0.6 M, 212 μL, 0.13 mmol) and stirred at rt for 6 h. Then, the reaction mixture was acidified with aq. HCl (1.0 M) and extracted with EtOAc (3 x 2 mL). The combined organic phase was dried over anhydrous MgSO_4_, filtered, concentrated and the crude was purified by preparative HPLC to afford the title compound (8.5 mg, 44%) as a white solid. ^1^H NMR (600 MHz, CDCl_3_) δ 8.30 – 8.17 (m, 1H), 7.90 – 7.77 (m, 1H), 7.58 – 7.43 (m, 3H), 7.37 (t, *J* = 8.0 Hz, 1H), 6.93 – 6.80 (m, 1H), 4.91 (s, 2H), 3.71 – 3.54 (m, 8H), 1.90 – 1.81 (m, 12H); ^13^C NMR (151 MHz, CDCl_3_) δ 181.7, 175.4, 167.0, 153.3, 134.7, 127.8, 126.8, 125.9, 125.9, 125.4, 121.7, 121.6, 105.3, 68.3, 45.8, 45.6, 45.3, 42.4, 39.8, 38.6, 28.0, 27.8; HPLC: t*_R_* = 12.07 min, purity 97.98%, Method D; HRMS (MALDI) m/z calcd for C_26_H_31_N_2_O_5_ (M+H^+^) 451.2227, found 451.2240.

### Ethyl 2-(4-(2-(naphthalen-1-yloxy)acetyl)piperazin-1-yl)-2-oxoacetate (35)

The title compound was synthesized according to the procedure described for **2** using **Dd** (50 mg, 0.18 mmol) and ethyl 2-chloro-2-oxoacetate (30 µL, 0.27 mmol) to afford 48 mg (70%) of a yellow oil after purification by flash chromatography (40% acetone in heptane). R_f_ = 0.66 (Acetone:Heptane; 1:1); ^1^H NMR (400 MHz, CDCl_3_) δ 8.23 – 8.16 (m, 1H), 7.86 – 7.79 (m, 1H), 7.57 – 7.46 (m, 3H), 7.37 (t, *J* = 8.0 Hz, 1H), 6.90 (d, *J* = 7.6 Hz, 1H), 4.93* (s, 1H’), 4.91* (s, 1H”), 4.38 – 4.23 (m, 2H), 3.79 – 3.68* (m, 4H’), 3.63 – 3.59* (m, 1H’), 3.59 – 3.52* (m, 1H’), 3.44 – 3.35* (m, 2H”), 1.42 – 1.29 (m, 3H); ^13^C NMR (101 MHz, CDCl_3_) δ 167.0*, 166.9*, 162.3, 160.3*, 160.1*, 153.1, 134.8, 127.9, 126.9, 126.0, 125.9, 125.8, 125.3, 121.8, 121.4, 105.3, 68.4*, 68.3*, 62.6, 46.4*, 46.0*, 45.8*, 45.2*, 42.3*, 41.8*, 41.5*, 14.1; HPLC: t*_R_* = 6.84 min, purity 95.5%, Method A; HRMS (MALDI) m/z calcd for C_20_H_23_N_2_O_5_^+^ (M+H^+^) 371.1601, found 371.1596.

### 2-(4-(2-(Naphthalen-1-yloxy)acetyl)piperazin-1-yl)-2-oxoacetic acid (36)

To a solution of **35** (20 mg, 54 µmol) in THF (0.5 mL) was added aq. LiOH (0.6 M, 0.3 mL) and stirred vigorously. After reaction completion, the mixture was acidified with aq. HCl (1.0 M) and extracted with EtOAc. The organic phase was washed with brine, dried over MgSO_4_, filtered, concentrated and the crude was purified by preparative HPLC (0-100% mobile phase B in mobile phase A in 10 min) to afford the title compound (5 mg, 27%) as a clear oil. R_f_ = 0.16 (MeOH:DCM; 2:8); ^1^H NMR (400 MHz, CDCl_3_) δ 8.21 – 8.14 (m, 1H), 7.86 – 7.78 (m, 1H), 7.56 – 7.46 (m, 3H), 7.36 (t, *J* = 8.0 Hz, 1H), 6.88 (d, *J* = 7.7 Hz, 1H), 4.94 (s, 2H), 4.01* (s, 2H’), 3.81 – 3.71* (m, 4H”), 3.66* (s, 1H’), 3.60* (s, 1H”); ^13^C NMR (101 MHz, CDCl_3_) δ 167.8, 153.0, 134.8, 127.9, 127.0, 126.0, 125.8, 125.3, 122.0, 121.3, 105.3, 68.0, 46.9*, 46.5*, 45.9*, 45.3*, 43.8*, 43.7*, 42.7*, 42.0*; HPLC: t*_R_* = 5.34 min, purity 94.02%, Method H; HRMS (MALDI) m/z calcd for C_18_H_19_N_2_O_5_^+^ (M+H^+^) 343.1288, found 343.1284.

### 4-(4-(2-(Naphthalen-1-yloxy)acetyl)piperazin-1-yl)-4-oxobutanoic acid (37)

To a solution of **Dd** (65 mg, 0.24 mmol) in DCM (1.5 mL) were added TEA (50 µL, 0.36 mmol), succinic anhydride (32 mg, 0.32 mmol), and DMAP (40 mg, 0.33 mol). The reaction mixture was stirred at rt for 24 h. Water (5 mL) was added and extracted with EtOAc (3 x 5 mL). The combined organic layer was washed with aq. HCl (1.0 M, 10 mL), dried over MgSO_4_, filtered, concentrated and the crude was purified by flash chromatography (10% MeOH in DCM) to afford the title compound (20 mg, 23%) as a clear oil. R_f_ = 0.67 (MeOH:DCM; 1:9); ^1^H NMR (600 MHz, CDCl_3_) δ 8.19 (d, *J* = 7.6 Hz, 1H), 7.84 – 7.76 (m, 1H), 7.54 – 7.43 (m, 3H), 7.39 – 7.31 (m, 1H), 6.90 – 6.81 (m, 1H), 4.90 (s, 2H), 3.72 – 3.60* (m, 4H’), 3.60 – 3.54* (m, 2H”), 3.45* (s, 1.3H’), 3.39* (s, 0.7H”), 2.68 – 2.63 (m, 2H), 2.62 – 2.56* (m, 1.3H’), 2.56 – 2.51* (m, 0.7H”); ^13^C NMR (151 MHz, CDCl_3_) δ 176.7, 170.6*, 170.5*, 167.3*, 167.2*, 153.24*, 153.16*, 134.7*, 127.9*, 127.8*, 126.9*, 125.9*, 125.8*, 125.3*, 125.2*, 122.0*, 121.9*, 121.8*, 121.7*, 121.5*, 121.4*, 121.33*, 121.29*, 105.34*, 105.28*, 68.3*, 68.0*, 45.6*, 45.5*, 45.4*, 45.2*, 42.2*, 41.7*, 29.3, 27.9; HPLC: t*_R_* = 5.20 min, purity 97.7%, Method A; HRMS (MALDI) m/z calcd for C_20_H_23_N_2_O_5_^+^ (M+H^+^) 371.1601, found 371.1594.

### 5-(4-(2-(Naphthalen-1-yloxy)acetyl)piperazin-1-yl)-5-oxopentanoic acid (38)

The title compound was synthesized according to the procedure described for **37** (excluding DMAP) using glutaric anhydride (18 mg, 0.15 mmol) to afford 13 mg (24%) of a faint orange solid after purification by flash chromatography (0-17% MeOH in DCM). R_f_ = 0.68 (MeOH:DCM; 1:9); ^1^H NMR (600 MHz, CDCl_3_) δ 8.20 (d, *J* = 7.4 Hz, 1H), 7.84 – 7.78 (m, 1H), 7.53 – 7.43 (m, 3H), 7.36 (t, *J* = 8.0 Hz, 1H), 6.91 – 6.86 (m, 1H), 4.91 (s, 2H), 3.75 – 3.51 (m, 6H), 3.49 – 3.35 (m, 2H), 2.45 – 2.30 (m, 4H), 1.93 (s, 2H); ^13^C NMR (151 MHz, CDCl_3_) δ 177.4, 171.3*, 171.1*, 167.1*, 167.0*, 153.3, 134.7, 127.9*, 127.8*, 126.8, 126.0*, 125.9*, 125.84*, 125.81*, 125.4, 121.7, 121.54*, 121.47*, 105.4*, 105.3*, 68.4*, 68.1*, 45.73*, 45.68*, 45.5*, 45.3*, 42.4*, 42.2*, 41.9, 41.4, 33.2, 32.1, 20.2; HPLC: t*_R_* = 13.03 min, purity >99.9%, Method G; HRMS (MALDI) m/z calcd for C_21_H_24_N_2_O_5_Na^+^ (M+Na^+^) 407.1577, found 407.1578.

### 2-(2-(4-(2-(Naphthalen-1-yloxy)acetyl)piperazin-1-yl)-2-oxoethoxy)acetic acid (39)

The title compound was synthesized according to the procedure described for **37** (excluding DMAP) using 1,4-dioxane-2,6-dione (21 mg, 0.17 mmol) and the reaction was carried out at 50 ^°^C to afford 11 mg (24%) of a white solid after purification by flash chromatography (0-20% MeOH in DCM). R_f_ = 0.32 (MeOH:DCM; 3:7); ^1^H NMR (600 MHz, CDCl_3_) δ 8.21 – 8.16 (m, 1H), 7.81 (d, *J* = 7.2 Hz, 1H), 7.54 – 7.46 (m, 3H), 7.36 (t, *J* = 8.0 Hz, 1H), 6.88 (d, *J* = 7.2 Hz, 1H), 4.92 (s, 2H), 4.30 (d, *J* = 38.7 Hz, 2H), 4.16 (d, *J* = 10.7 Hz, 2H), 3.76 – 3.52 (m, 6H), 3.30 (d, *J* = 35.4 Hz, 2H); ^13^C NMR (151 MHz, CDCl_3_) δ 171.6, 169.4*, 169.3*, 167.2*, 167.0*, 153.1, 134.7, 128.0*, 127.9*, 126.9*, 126.9*, 126.0*, 125.9*, 125.8*, 125.3, 121.8, 121.41*, 121.36*, 105.4*, 105.3*, 71.5, 71.0, 68.3*, 68.1*, 45.3*, 45.2*, 44.8*, 44.3*, 42.5*, 42.1*, 42.0*, 41.9*; HPLC: t*_R_* = 10.80 min, purity >99.9%, Method B; HRMS (MALDI) m/z calcd for C_20_H_22_N_2_O_6_Na^+^ (M+Na^+^) 409.1370, found 409.1367.

### 2-(4-(2-(Naphthalen-1-yloxy)acetyl)piperazin-1-yl)-2-oxoethyl acetate (40)

The title compound was synthesized according to the procedure described for **4** using 2-acetoxyacetic acid (11 mg, 92.5 µmol) and DMF (2.0 mL) as the solvent for the reaction to afford 28 mg (82%) of a white foam after purification by flash chromatography (20-90% EtOAc in heptane). R_f_ = 0.31 (EtOAc:Heptane; 9:1); ^1^H NMR (400 MHz, CDCl_3_) δ 8.23 – 8.16 (m, 1H), 7.86 – 7.79 (m, 1H), 7.56 – 7.46 (m, 3H), 7.37 (t, *J* = 8.0 Hz, 1H), 6.90 (d, *J* = 7.7 Hz, 1H), 4.92 (s, 2H), 4.69* (s, 1H’), 4.65* (s, 1H”), 3.71* (s, 4H’), 3.57* (s, 2H”), 3.37* (s, 1.3H’), 3.30* (s, 0.7H”), 2.16 (s, 3H); ^13^C NMR (101 MHz, CDCl_3_) δ 170.6, 165.3, 153.2, 134.8, 127.9, 126.9, 125.9, 125.3, 121.8, 121.4, 105.3, 68.2, 61.3, 45.5, 44.5, 42.1, 20.7; HPLC: t*_R_* = 5.80 min, purity 97.8%, Method A; HRMS (MALDI) m/z calcd for C_20_H_23_N_2_O_5_^+^ (M+H^+^) 371.1601, found 371.1595.

### 4-(Dimethylamino)-1-(4-(2-(naphthalen-1-yloxy)acetyl)piperazin-1-yl)butan-1-one (41)

The title compound was synthesized according to the procedure described for **4** using 4-(dimethylamino)butanoic acid hydrochloride (31 mg, 0.18 mmol) and DMF (2.0 mL) as the solvent for the reaction to afford 18 mg (25%) of a clear oil after purification by flash chromatography (30% MeOH in DCM). R_f_ = 0.23 (MeOH:DCM; 3:7); ^1^H NMR (600 MHz, CDCl_3_) δ 8.20 (d, *J* = 7.7 Hz, 1H), 7.84 – 7.78 (m, 1H), 7.54 – 7.45 (m, 3H), 7.39 – 7.33 (m, 1H), 6.92 – 6.86 (m, 1H), 4.91 (s, 2H), 3.82 – 3.34 (m, 8H), 2.38 – 2.23 (m, 4H), 2.23 – 2.14 (m, 6H), 1.83 – 1.73 (m, 2H); ^13^C NMR (101 MHz, CDCl_3_) δ 171.7*, 171.6*, 167.0*, 166.8*, 153.3, 134.7, 127.8, 126.8, 125.9, 125.4, 121.7*, 121.5*, 105.3, 68.5*, 68.2*, 59.0, 45.8*, 45.5, 45.3*, 42.4*, 41.9*, 41.4*, 30.8, 23.1; HPLC: t*_R_* = 7.78 min, purity = 98.0%, Method H; HRMS (MALDI) m/z calcd for C_22_H_30_N_3_O_3_^+^ (M+H^+^) 384.2281, found 384.2276.

### 1-(4-(2-(Naphthalen-1-yloxy)acetyl)piperazin-1-yl)hexane-1,5-dione (42)

The title compound was synthesized according to the procedure described for **4** using 5-oxohexanoic acid (25 mg, 0.19 mmol) to afford 45 mg (65%) of a faint yellow oil after purification by flash chromatography (2% MeOH in DCM). ^1^H NMR (600 MHz, CD_3_OD) δ 8.33 – 8.27 (m, 1H), 7.84 – 7.79 (m, 1H), 7.53 – 7.45 (m, 3H), 7.37 (t, *J* = 8.0 Hz, 1H), 6.93 (d, *J* = 7.7 Hz, 1H), 5.03 (s, 2H), 3.76 – 3.51 (m, 8H), 2.59 – 2.52 (m, 1H), 2.44 – 2.35 (m, 2H), 2.12 (s, 2H), 1.85 – 1.78 (m, 1H), 1.65 – 1.56 (m, 2H), 1.27 – 1.22 (m, 1H); ^13^C NMR (151 MHz, CD_3_OD) δ 211.1, 174.1*, 174.0*, 173.9*, 173.8*, 169.2*, 169.1*, 154.9, 136.1, 128.6, 127.6, 126.8, 126.5, 122.8, 122.2, 106.5, 102.8, 68.3*, 68.1*, 46.7*, 46.2*, 45.94*, 45.89*, 43.2*, 42.9*, 42.8*, 42.3*, 37.0, 33.8*, 33.7*, 33.2*, 33.1*, 29.8, 21.2, 21.1, 20.3; HPLC: t*_R_* = 11.71 min, purity 95.7%, Method D; HRMS (MALDI) m/z calcd for C_22_H_27_N_2_O_4_^+^ (M+H^+^) 383.1965, found 383.1977.

### 6-Hydroxy-1-(4-(2-(naphthalen-1-yloxy)acetyl)piperazin-1-yl)hexan-1-one (43)

The title compound was synthesized according to the procedure described for **4** using 6-hydroxyhexanoic acid (25 mg, 0.19 mmol) to afford 30 mg (52%) of an orange oil after purification by flash chromatography (2% MeOH in DCM). ^1^H NMR (600 MHz, CD_3_OD) δ 8.35 – 8.29 (m, 1H), 7.86 – 7.81 (m, 1H), 7.54 – 7.47 (m, 3H), 7.39 (t, *J* = 8.0 Hz, 1H), 6.94 (d, *J* = 7.7 Hz, 1H), 5.07 – 5.03 (m, 2H), 4.40 (s, 1H), 3.79 – 3.52 (m, 9H), 2.48 – 2.38 (m, 2H), 1.83 – 1.76 (m, 1H), 1.71 – 1.61 (m, 2H), 1.61 – 1.54 (m, 1H), 1.51 – 1.39 (m, 2H); ^13^C NMR (151 MHz, CD_3_OD) δ 173.0*, 172.9*, 172.7*, 172.6*, 167.8*, 167.7*, 153.5, 134.7, 127.2, 126.2, 125.4, 125.1, 121.4, 120.9*, 120.8*, 105.1, 67.9, 66.9*, 66.7*, 61.3, 45.3*, 45.3*, 44.9*, 44.5*, 41.8*, 41.5*, 41.4*, 40.9*, 32.5*, 32.2*, 32.0*, 27.6, 25.3*, 24.9*, 24.8*, 24.3*; HPLC: t*_R_* = 11.60 min, purity 95.4%, Method D; HRMS (MALDI) m/z calcd for C_22_H_29_N_2_O_4_^+^ (M+H^+^) 385.2122, found 385.2131.

### 8-Hydroxy-1-(4-(2-(naphthalen-1-yloxy)acetyl)piperazin-1-yl)octan-1-one (44)

The title compound was synthesized according to the procedure described for **4** using 8-hydroxyoctanoic acid (18 mg, 0.11 mmol) to afford 5 mg (12%) of a colorless oil after purification by preparative HPLC. ^1^H NMR (600 MHz, CDCl_3_) δ 8.21 (d, *J* = 7.3 Hz, 1H), 7.86 – 7.79 (m, 1H), 7.56 – 7.46 (m, 3H), 7.41 – 7.34 (m, 1H), 6.93 – 6.88 (m, 1H), 4.93 (s, 2H), 3.76 – 3.53 (m, 8H), 3.49 – 3.36 (m, 2H), 2.36 – 2.23 (m, 2H), 1.78 – 1.48 (m, 4H), 1.42 – 1.20 (m, 6H); ^13^C NMR (151 MHz, CDCl_3_) δ 172.3*, 172,1*, 167.1*, 166.9*, 153.3*, 153.2*, 134.7, 127.9*, 127.8*, 126.8, 125.9*, 125.8*, 125.4, 121.7, 121.5*, 121.4*, 105.4*, 105.2*, 68.5*, 68.3*, 68.2*, 63.1, 45.9*, 45.8*, 45.6*, 45.4*, 42.4*, 42.3*, 41.9*, 41.4*, 33.3, 32.7, 29.4*, 29.3*, 29.2*, 29.0*, 28.2, 25.6*, 25.5*, 25.2*, 25.1*; HPLC: t*_R_* = 9.88 min, purity 95.8%, Method H; HRMS (MALDI) m/z calcd for C_24_H_33_N_2_O_4_^+^ (M+H^+^) 413.2435, found 413.2450.

### 6-(Methylsulfonyl)-1-(4-(2-(naphthalen-1-yloxy)acetyl)piperazin-1-yl)hexan-1-one (45)

A mixture of **58** (24 mg, 0.053 mmol) and sodium methanesulfinate (11 mg, 0.11 mmol) in PEG-400 (640 μL) was stirred at 45 °C for 30 h. The reaction mixture was cooled to rt, water was added, and the mixture was extracted with EtOAc (x3). The combined extracts were washed with brine, dried over MgSO_4_, and concentrated under reduced pressure. The crude residue was purified by preparative HPLC to afford the title compound as a white solid (14 mg, 60%). ^1^H NMR (600 MHz, CDCl_3_) δ 8.21 (d, *J* = 7.4 Hz, 1H), 7.85 – 7.80 (m, 1H), 7.55 – 7.46 (m, 3H), 7.41 – 7.33 (m, 1H), 6.94 – 6.87 (m, 1H), 4.93 (s, 2H), 3.75 – 3.62 (m, 4H), 3.62 – 3.56 (m, 2H), 3.49 – 3.34 (m, 2H), 3.04 – 2.98 (m, 2H), 2.89 (s, 3H), 2.38 – 2.26 (m, 2H), 1.91 – 1.83 (m, 2H), 1.71 – 1.62 (m, 2H), 1.55 – 1.46 (m, 2H); ^13^C NMR (151 MHz, CDCl_3_) δ 171.4*, 171.3*, 167.1*, 167.0*, 153.3*, 153.2*, 134.7, 127.9*, 127.8*, 126.8, 125.9*, 125.8*, 125.4, 121.7, 121.5*, 121.4*, 105.4*, 105.3*, 68.5*, 68.3*, 54.6, 45.8*, 45.6*, 45.3*, 42.4*, 42.3*, 42.0*, 41.5*, 40.8, 32.6, 28.1, 24.4, 22.3*, 22.2*; HPLC: t*_R_* = 11.62 min, purity 99.0%, Method D; HRMS (MALDI) m/z calcd for C_23_H_31_N_2_O_5_S^+^ (M+H^+^) 447.1948, found 447.1966.

### 7-(Methylsulfonyl)-1-(4-(2-(naphthalen-1-yloxy)acetyl)piperazin-1-yl)heptan-1-one (46)

The title compound was synthesized according to the procedure described for **45** using **59** (11.6 mg, 0.025 mmol) to afford 5.8 mg (51%) of a yellow oil after purification by preparative HPLC. ^1^H NMR (600 MHz, CDCl_3_) δ 8.25 – 8.17 (m, 1H), 7.88 – 7.79 (m, 1H), 7.55 – 7.45 (m, 3H), 7.41 – 7.34 (m, 1H), 6.94 – 6.87 (m, 1H), 4.92 (s, 2H), 3.77 – 3.53 (m, 6H), 3.46 – 3.32 (m, 2H), 2.99 (t, *J* = 8.1 Hz, 2H), 2.88 (s, 3H), 2.39 – 2.22 (m, 2H), 1.92 – 1.80 (m, 2H), 1.67 – 1.60 (m, 2H), 1.51 – 1.44 (m, 2H), 1.38 (s, 2H); ^13^C NMR (151 MHz, CDCl_3_) δ 171.5*, 171.4*, 167.1*, 166.9*, 153.3, 134.8, 127.9*, 127.8*, 126.8, 126.0*, 125.9*, 125.8*, 125.4, 121.7, 121.6*, 121.5*, 105.4*, 105.3*, 68.5*, 68.2*, 54.8, 45.8*, 45.6*, 45.3*, 42.3*, 42.4*, 41.9*, 41.4*, 40.7, 33.0, 28.9, 28.3, 24.7, 22.4*; HPLC: t*_R_* = 11.94 min, purity 99.3%, Method D; HRMS (MALDI) m/z calcd for C_24_H_33_N_2_O_5_S^+^ (M+H^+^) 461.2104, found 461.2121.

### 8-(Methylsulfonyl)-1-(4-(2-(naphthalen-1-yloxy)acetyl)piperazin-1-yl)octan-1-one (47)

The title compound was synthesized according to the procedure described for **45** using **60** (17 mg, 0.036 mmol) to afford 11 mg (65%) of a colorless oil after purification by preparative HPLC (50% mobile phase B in mobile phase A). R_f_ = 0.38 (MeOH: DCM; 1:19); ^1^H NMR (400 MHz, CDCl_3_) δ 8.20 (d, *J* = 9.3 Hz, 1H), 7.85 – 7.79 (m, 1H), 7.54 – 7.47 (m, 3H), 7.37 (t, *J* = 7.8 Hz, 1H), 6.92 – 6.88 (m, 1H), 4.93 (s, 2H), 3.75 – 3.57 (m, 6H), 3.48 – 3.38 (m, 2H), 2.99 (t, *J* = 8.1 Hz, 2H), 2.88 (s, 3H), 2.55 – 2.24 (m, 2H), 1.89 – 1.78 (m, 2H), 1.65 – 1.53 (m, 2H), 1.48 – 1.28 (m, 6H);^13^C NMR (101 MHz, CDCl_3_) δ 172.3, 167.3, 153.2, 134.8, 127.8, 126.9, 126.0, 125.8, 125.4, 121.8, 121.5, 105.3, 68.1, 54.9, 45.6, 45.5, 42.5, 42.0, 40.7, 33.2, 29.0, 28.9, 28.3, 25.1, 22.4; HPLC: t*_R_* = 5.37 min, purity 97.0%, Method I; HRMS (MALDI) m/z calcd for C_25_H_34_N_2_O_5_SNa^+^ (M+Na^+^) 497.2080, found 497.2086.

### 1-(4-(2-(Naphthalen-1-yloxy)acetyl)piperazin-1-yl)-7-phenoxyheptan-1-one (48)

To a round bottom flask was added **58** (18.2 mg, 0.041 mmol), phenol (5.7 mg, 0.061 mmol), cesium carbonate (26.5 mg, 0.081 mmol) and potassium iodide (1 mg, 0.006 mmol). Then acetone (350 μL) was added, and the reaction was stirred at 35°C. After 24 h, the solvent was evaporated, and the residue was dissolved in EtOAc. The organic solution was washed with aq. NaOH (1M) and brine. The organic phase was dried over anhydrous MgSO_4_, filtered, and the filtrate evaporated. The crude residue was purified by preparative HPLC to give the title compound (15 mg, 79%) as a clear oil. ^1^H NMR (600 MHz, CD_3_OD): δ 8.33 – 8.27 (m, 1H), 7.84 – 7.79 (m, 1H), 7.52 – 7.45 (m, 3H), 7.37 (t, *J* = 8.0 Hz, 1H), 7.23 (t, *J* = 7.8 Hz, 2H), 6.95 – 6.90 (m, 1H), 6.90 – 6.85 (m, 3H), 5.02 (d, *J* = 14.2 Hz, 2H), 4.00 – 3.93 (m, 2H), 3.71 – 3.54 (m, 8H), 2.43 (dt, *J* = 19.4, 7.5 Hz, 2H), 1.79 (q, *J* = 7.1 Hz, 2H), 1.68 (d, *J* = 14.7 Hz, 2H), 1.53 (t, *J* = 8.0 Hz, 2H); ^13^C NMR (151 MHz, CDCl_3_) δ 171.7*, 171.5*, 166.9*, 166.7*, 158.9, 153.1*, 153.0*, 134.5, 129.4, 127.7*, 127.6*, 126.6, 125.7*, 125.6*, 125.2*, 121.5*, 121.3*, 121.2*, 120.5, 114.4, 105.1*, 105.0*, 68.3*, 68.0*, 67.4, 45.63*, 45.59*, 45.4*, 45.2*, 42.2*, 42.1*, 41.7*, 41.2*, 33.0, 29.0, 25.8, 24.8; HPLC: t*_R_* = 8.31 min, purity 95.25%, Method I; HRMS (MALDI): m/z calcd for C_28_H_33_N_2_O_4_ (M+H^+^) 461.2435, found 461.2449.

### 1-(4-(2-(Naphthalen-1-yloxy)acetyl)piperazin-1-yl)-7-phenoxyheptan-1-one (49)

To a round bottom flask was added **59** (21 mg, 0.046 mmol), phenol (4.3 mg, 0.045 mmol), cesium carbonate (29.7 mg, 0.091 mmol) and potassium iodide (1 mg, 0.006 mmol). Then acetone (350 μL) was added, and the reaction was stirred at 35°C. After 24 h, the solvent was evaporated, and the residue was dissolved in EtOAc. The organic solution was washed with aq. NaOH (1M) and brine. The organic phase was dried over anhydrous MgSO_4_, filtered, and the filtrate evaporated. The crude was purified by preparative HPLC to give the title compound (15 mg, 67%) of a clear oil. ^1^H NMR (600 MHz, CD_3_OD) δ 8.37 – 8.31 (m, 1H), 7.89 – 7.82 (m, 1H), 7.56 – 7.48 (m, 3H), 7.40 (t, *J* = 7.9 Hz, 1H), 7.30 – 7.23 (m, 2H), 6.96 (d, *J* = 7.7 Hz, 1H), 6.94 – 6.88 (m, 3H), 5.06 (s, 2H), 4.04 – 3.96 (m, 2H), 3.79 – 3.57 (m, 8H), 2.51 – 2.40 (m, 2H), 1.85 – 1.75 (m, 2H), 1.73 – 1.62 (m, 2H), 1.62 – 1.51 (m, 2H), 1.51 – 1.42 (m, 2H); ^13^C NMR (151 MHz, CDCl_3_) δ 172.3, 167.1, 166.9*, 159.0, 153.0, 134.5, 129.4, 127.7*, 127.6, 126.7*, 125.7, 125.6*, 125.2, 121.6, 121.3, 121.2*, 120.5, 114.4, 105.1*, 105.0, 67.9, 67.5, 45.7*, 45.6*, 45.4*, 15.3*, 42.2*, 42.1*, 41.8*, 41.3*, 33.0, 29.03, 29.0, 25.8, 25.1; HPLC: t*_R_* = 8.99 min, purity 95.99%, Method I; HRMS (MALDI): m/z calcd for C_29_H_35_N_2_O_4_ (M+H^+^) 475.2591, found 475.2608.

### (*E*)-1-(4-(3-(Naphthalen-1-yl)acryloyl)piperazin-1-yl)hexan-1-one (50)

The title compound was synthesized according to the procedure described for **4** using TFA salt of **Gb** (108 mg, 0.53 mmol) and 3-(naphthalen-1-yl)acrylic acid (107 mg, 0.53 mmol) to afford 106 mg (55%) of a white solid after purification by flash chromatography (0-60% EtOAc in heptane). R_f_ = 0.54 (MeOH:DCM; 1:9); ^1^H NMR (600 MHz, CDCl_3_) δ 8.54 (d, *J* = 15.1 Hz, 1H), 8.21 (d, *J* = 8.3 Hz, 1H), 7.91 – 7.85 (m, 2H), 7.72 (d, *J* = 7.4 Hz, 1H), 7.60 – 7.51 (m, 2H), 7.48 (t, *J* = 7.7 Hz, 1H), 6.93 (d, *J* = 15.1 Hz, 1H), 3.86 – 3.51 (m, 6H), 2.36 (t, *J* = 7.6 Hz, 3H), 1.66 (t, *J* = 7.7 Hz, 1H), 1.61 (d, *J* = 1.9 Hz, 5H), 1.35 (s, 1H), 0.94 – 0.89 (m, 2H); ^13^C NMR (151 MHz, CDCl_3_) δ 172.2, 165.7, 141.1, 133.8, 132.9, 131.6, 130.3, 128.8, 126.9, 126.4, 125.5, 124.8, 123.8, 119.6, 45.8, 45.5, 42.4, 41.7, 33.5, 31.8, 25.1, 22.6, 14.1; HPLC: t*_R_* = 13.99 min, purity 98.1%, Method D; HRMS (MALDI) m/z calcd for C_23_H_28_N_2_O_2_Na^+^ (M+Na^+^) 387.2043, found 387.2041.

### 1-(4-(3-(Naphthalen-1-yl)propanoyl)piperazin-1-yl)hexan-1-one (51)

To a solution of **50** (40 mg, 0.11 mmol) in MeOH (2 mL) was added 10% Pd-C (3.6 mg). The mixture was evacuated and backfilled with argon (3x) followed with hydrogen (3x) and stirred rt under the atmosphere of hydrogen. After 6 h, the reaction mixture was filtered through a Celite pad. The filtrate was concentrated and purified by preparative HPLC to afford the title compound (24 mg, 65%) as a colorless oil. R_f_ = 0.54 (MeOH:DCM; 1:9); ^1^H NMR (600 MHz, CDCl_3_) δ 8.03 (d, *J* = 8.2 Hz, 1H), 7.86 (d, *J* = 8.0 Hz, 1H), 7.73 (d, *J* = 7.9 Hz, 1H), 7.55 – 7.45 (m, 2H), 7.43 – 7.33 (m, 2H), 3.67 – 2.97 (m, 10H), 2.77 (t, *J* = 7.8 Hz, 2H), 2.33 – 2.17 (m, 2H), 1.64 – 1.53 (m, 2H), 1.37 – 1.24 (m, 4H), 0.89 (t, *J* = 7.0 Hz, 3H); ^13^C NMR (151 MHz, CDCl_3_) δ 172.0*, 171.8*, 171.3*, 171.1*, 137.2*, 137.1*, 134.0, 131.8, 129.1*, 129.0*, 127.34*, 127.28*, 126.6*, 126.5*, 126.3, 125.8*, 125.7*, 123.6*, 123.5*, 45.6*, 45.4*, 45.3*, 45.2*, 41.8*, 41.5*, 41.3*, 34.2, 33.3, 31.7, 28.9, 28.8, 25.0, 22.6; HPLC: t*_R_* = 12.17 min, purity >99.8%, Method E; HRMS (MALDI) m/z calcd for C_23_H_31_N_2_O_2_ (M+H^+^) 367.2380, found 367.2390.

### 1-(4-(2-((5,6,7,8-Tetrahydronaphthalen-1-yl)oxy)acetyl)piperazin-1-yl)hexan-1-one (52)

Step 1: *tert*-Butyl 4-hexanoylpiperazine-1-carboxylate (**Eb**): The title compound was synthesized according to the procedure described for **2** using **A** (1.00 g, 5.37 mmol) and hexanoyl chloride (1.1 mL, 7.87 mmol) to afford 1.43 g (94%) of a white solid after purification by flash chromatography (50% EtOAc in heptane). R_f_ = 0.44 (EtOAc:Heptane; 7:3); ^1^H NMR (400 MHz, CDCl_3_) δ 3.65 – 3.32 (m, 8H), 2.31 (t, *J* = 7.4 Hz, 2H), 1.67 – 1.55 (m, 2H), 1.46 (s, 9H), 1.37 – 1.28 (m, 4H), 0.88 (t, *J* = 6.5 Hz, 3H); ^13^C NMR (101 MHz, CDCl_3_) δ 172.0, 154.7, 80.4, 45.5, 41.4, 33.5, 31.7, 28.5, 25.1, 22.6, 14.1.

Step 2: 1-(Piperazin-1-yl)hexan-1-one (**Fb**): The target compound was synthesized according to the procedure described for **Da** using **Eb** (1.0 g, 3.52 mmol) to yield 602 mg (94%) of a clear oil. R_f_ = 0.26 (EtOAc:Heptane; 9:1); ^1^H NMR (400 MHz, CDCl_3_) δ 3.90 (s, 2H), 3.80 (s, 2H), 3.24 (s, 4H), 2.40 – 2.32 (m, 2H), 1.61 (p, *J* = 7.4 Hz, 2H), 1.40 – 1.24 (m, 4H), 0.90 (t, *J* = 6.6 Hz, 3H); ^13^C NMR (101 MHz, CDCl_3_) δ 173.0, 43.8, 42.6, 38.5, 33.0, 31.5, 24.9, 22.5, 13.9.

Step 3: 1-(4-(2-Chloroacetyl)piperazin-1-yl)hexan-1-one (**Gb**): The target compound was synthesized according to the procedure described for **B** using **Fb** (250 mg, 1.36 mmol) to yield 342 mg (97%) of a yellow solid. R_f_ = 0.21 (EtOAc:Heptane; 8:2); ^1^H NMR (400 MHz, CDCl_3_) δ 4.08 (s, 2H), 3.70* (s, 1H’), 3.63* (s, 3H’), 3.55* (s, 1H”), 3.54 – 3.44* (m, 3H”), 2.33 (t, *J* = 7.7 Hz, 2H), 1.63 (p, *J* = 7.3 Hz, 2H), 1.39 – 1.27 (m, 4H), 0.90 (t, *J* = 6.5 Hz, 3H); ^13^C NMR (101 MHz, CDCl_3_) δ 172.2*, 172.0*, 165.6*, 165.3*, 46.5*, 46.2*, 45.6*, 45.2*, 42.3*, 42.2*, 41.5*, 41.1*, 40.8, 33.3, 31.7, 25.0, 22.5, 14.0.

Step 4: The title compound was synthesized according to the procedure described for **Ca** using **Gb** (40 mg, 0.15 mmol) and 5,6,7,8-tetrahydronaphthalen-1-ol (35 mg, 0.24 mmol) to afford 56 mg (98%) of a clear oil after purification by flash chromatography (90% EtOAc in heptane). R_f_ = 0.25 (EtOAc:Heptane; 7:3); ^1^H NMR (600 MHz, CDCl_3_) δ 7.04 (t, *J* = 7.9 Hz, 1H), 6.74 (d, *J* = 7.6 Hz, 1H), 6.66 (d, *J* = 8.1 Hz, 1H), 4.69 (s, 2H), 3.62* (s, 6H’), 3.45* (s, 2H”) 2.74 (t, *J* = 6.1 Hz, 2H), 2.65 (t, *J* = 6.2 Hz, 2H), 2.32 (t, *J* = 7.6 Hz, 2H), 1.82 – 1.71 (m, 4H), 1.62 (p, *J* = 7.4 Hz, 2H), 1.38 – 1.27 (m, 4H), 0.90 (t, *J* = 6.8 Hz, 3H); ^13^C NMR (151 MHz, CDCl_3_) δ 172.0, 167.2, 155.4, 139.1, 126.0, 122.7, 107.8, 68.0, 45.6*, 45.4*, 42.3*, 41.9*, 41.3*, 33.3, 31.7, 29.7, 25.0, 23.3, 22.8, 22.5, 14.0; HPLC: t*_R_* = 8.68 min, purity 98.1%, Method A; HRMS (MALDI) m/z calcd for C_22_H_33_N_2_O_3_^+^ (M+H^+^) 373.2485, found 373.2480.

### 1-(4-(2-(Quinolin-5-yloxy)acetyl)piperazin-1-yl)hexan-1-one (53)

The title compound was synthesized according to the procedure described for **Ca** using **Gb** (40 mg, 0.15 mmol) and quinolin-5-ol (34 mg, 0.23 mmol) to afford 44 mg (78%) of a yellow oil after purification by flash chromatography (90% EtOAc in heptane). R_f_ = 0.21 (Acetone:Heptane; 7:3); ^1^H NMR (600 MHz, CDCl_3_) δ 8.94 – 8.90 (m, 1H), 8.61 (d, *J* = 8.2 Hz, 1H), 7.79 (d, *J* = 8.5 Hz, 1H), 7.61 (t, *J* = 8.1 Hz, 1H), 7.43 (dd, *J* = 8.4, 4.2 Hz, 1H), 6.95 (d, *J* = 7.7 Hz, 1H), 4.93 (s, 2H), 3.65* (s, 2.5H’), 3.61* (s, 3.5H”), 3.45* (s, 2H’), 2.33 – 2.24 (m, 2H), 1.64 – 1.56 (m, 2H), 1.40 – 1.26 (m, 4H), 0.92 – 0.85 (m, 3H); ^13^C NMR (151 MHz, CDCl_3_) δ 172.1*, 171.8*, 166.3*, 166.2*, 153.1, 150.3, 148.3, 131.4, 131.2, 129.9, 122.3, 120.9, 120.7, 106.1, 68.3*, 68.0*, 45.8*, 45.6*, 45.4*, 45.3*, 42.3*, 42.2*, 41.7*, 41.3*, 33.3, 31.7, 25.0, 22.6, 14.0; HPLC: t*_R_* = 7.50 min, purity 99.7%, Method H; HRMS (MALDI) m/z calcd for C_21_H_28_N_3_O_3_^+^ (M+H^+^) 370.2125, found 370.2120.

### 1-(4-(2-(Isoquinolin-5-yloxy)acetyl)piperazin-1-yl)hexan-1-one (54)

The title compound was synthesized according to the procedure described for **Ca** using **Gb** (40 mg, 0.15 mmol) and isoquinolin-5-ol (34 mg, 0.23 mmol) to afford 44 mg (78%) of a yellow oil after purification by flash chromatography (90% EtOAc in heptane). R_f_ = 0.56 (Acetone:Heptane; 7:3); ^1^H NMR (600 MHz, CDCl_3_) δ 9.24 – 9.21 (m, 1H), 8.54 (d, *J* = 5.9 Hz, 1H), 8.02 – 7.96 (m, 1H), 7.60 (d, *J* = 8.2 Hz, 1H), 7.50 (t, *J* = 8.0 Hz, 1H), 7.15 – 7.07 (m, 1H), 4.93 (s, 2H), 3.68 – 3.64* (m, 2H’), 3.64 – 3.56* (m, 4H”), 3.49 – 3.43* (m, 1.3H’), 3.43 – 3.39* (m, 0.7H”), 2.33 – 2.23 (m, 2H), 1.62 – 1.56 (m, 2H), 1.35 – 1.26 (m, 4H), 0.90 – 0.86 (m, 3H); ^13^C NMR (101 MHz, CDCl_3_) δ 172.1*, 171.8*, 166.3, 162.8, 152.5, 151.7, 142.6, 129.5, 128.4, 127.6, 120.9, 114.9, 109.5, 68.3*, 67.9*, 45.7*, 45.4*, 45.3*, 42.3*, 41.7*, 41.2*, 33.3, 31.7, 25.0, 22.5, 14.0; HPLC: t*_R_* = 7.50 min, purity 99.0%, Method H; HRMS (MALDI) m/z calcd for C_21_H_28_N_3_O_3_^+^ (M+H^+^) 370.2125, found 370.2120.

### 1-(4-(2-(Quinolin-8-yloxy)acetyl)piperazin-1-yl)hexan-1-one (55)

The title compound was synthesized according to the procedure described for **Ca** using **Gb** (40 mg, 0.15 mmol) and quinolin-8-ol (34 mg, 0.23 mmol) to afford 26 mg (46%) of a yellow oil after purification by flash chromatography (3% MeOH in DCM). R_f_ = 0.52 (MeOH:DCM; 5:95); ^1^H NMR (600 MHz, CDCl_3_) δ 8.96 – 8.89 (m, 1H), 8.26 – 8.09 (m, 1H), 7.51 – 7.41 (m, 3H), 7.31 – 7.15 (m, 1H), 5.04 (s, 2H), 3.83* (s, 1H’), 3.74* (s, 1H”), 3.59* (s, 1H’), 3.55* (s, 1H”), 3.52 – 3.46* (m, 2H’), 3.38 – 3.34* (m, 2H”), 2.27 – 2.18 (m, 2H), 1.57 – 1.50 (m, 2H), 1.33 – 1.20 (m, 4H), 0.84 (t, *J* = 6.8 Hz, 3H); ^13^C NMR (151 MHz, CDCl_3_) δ 171.9, 167.0*, 166.7*, 153.2*, 152.8*, 149.5*, 148.9*, 139.8*, 139.0*, 137.2*, 136.3*, 129.70*, 129.66*, 127.4*, 126.8*, 122.0*, 121.9*, 121.1*, 110.8*, 110.2*, 69.4*, 69.1*, 45.9*, 45.5*, 45.3*, 42.4*, 42.3*, 41.8*, 41.2*, 33.3, 31.6, 24.9, 22.5, 14.0; HPLC: t*_R_* = 4.25 min, purity 97.3%, Method F; HRMS (MALDI) m/z calcd for C_21_H_28_N_3_O_3_^+^ (M+H^+^) 370.2125, found 370.2129.

### 1-(4-(2-(Quinolin-4-yloxy)acetyl)piperazin-1-yl)hexan-1-one (56)

The title compound was synthesized according to the procedure described for **Ca** using **Gb** (40 mg, 0.15 mmol) and quinolin-4-ol (34 mg, 0.23 mmol) to afford 32 mg (56%) of a clear oil after purification by preparative HPLC (0-100% mobile phase B in mobile phase A in 10 min). R_f_ = 0.11 (Acetone:Heptane; 9:1); ^1^H NMR (400 MHz, CDCl_3_) δ 8.40 (d, *J* = 7.9 Hz, 1H), 7.66 – 7.46 (m, 2H), 7.35 (t, *J* = 7.6 Hz, 1H), 7.11 (d, *J* = 8.8 Hz, 1H), 6.31 (d, *J* = 7.6 Hz, 1H), 5.05* (s, 0.7H’), 4.97* (s, 1.3H”), 3.77* (s, 1H’), 3.69 – 3.55* (m, 6H”), 3.52* (s, 1H’), 2.34 (t, *J* = 7.6 Hz, 2H), 1.65 (p, *J* = 5.0 Hz, 2H), 1.41 – 1.28 (m, 4H), 0.91 (t, *J* = 6.7 Hz, 3H); ^13^C NMR (101 MHz, CDCl_3_) δ 178.1, 172.2, 164.6, 144.4, 140.6, 132.6, 127.1, 124.2, 115.1, 110.4, 53.6, 45.2*, 44.9*, 42.5*, 41.4*, 33.4, 31.7, 25.0, 22.6, 14.1; HPLC: t*_R_* = 8.47 min, purity 96.1%, Method H; HRMS (MALDI) m/z calcd for C_21_H_28_N_3_O_3_^+^ (M+H^+^) 370.2125, found 370.2120.

### 1-(4-(2-(Isoquinolin-1-yloxy)acetyl)piperazin-1-yl)hexan-1-one (57)

The title compound was synthesized according to the procedure described for **Ca** using **Gb** (40 mg, 0.15 mmol) and isoquinolin-1-ol (34 mg, 0.23 mmol) to afford 48 mg (85%) of a clear oil after purification by flash chromatography (90% EtOAc in heptane). R_f_ = 0.23 (Acetone:Heptane; 7:3); ^1^H NMR (400 MHz, CDCl_3_) δ 8.35 (d, *J* = 8.0 Hz, 1H), 7.61 (t, *J* = 7.6 Hz, 1H), 7.55 – 7.41 (m, 2H), 7.16 – 6.99 (m, 1H), 6.51 (d, *J* = 7.3 Hz, 1H), 4.79 (s, 2H), 3.78 – 3.54* (m, 6H’), 3.54 – 3.40* (m, 2H”), 2.29 (t, *J* = 7.7 Hz, 2H), 1.60 (p, *J* = 7.2 Hz, 2H), 1.38 – 1.22 (m, 4H), 0.92 – 0.84 (m, 3H); ^13^C NMR (101 MHz, CDCl_3_) δ 172.0, 166.0, 162.2, 137.3, 132.5, 132.1, 127.9, 127.0, 126.1, 125.8, 106.5, 49.0, 45.2, 42.2, 41.3, 33.3, 31.6, 24.9, 22.5, 14.0; HPLC: t*_R_* = 7.50 min, purity 99.9%, Method A; HRMS (MALDI) m/z calcd for C_21_H_28_N_3_O_3_^+^ (M+H^+^) 370.2125, found 370.2119.

### 6-Bromo-1-(4-(2-(naphthalen-1-yloxy)acetyl)piperazin-1-yl)hexan-1-one (58)

The title compound was synthesized according to the procedure described for **4** using 6-bromohexanoic acid (170 mg, 0.87 mmol) to afford 118 mg (31%) of an orange solid after purification by flash chromatography (EtOAc). ^1^H NMR (400 MHz, CDCl_3_) δ 8.22 – 8.19 (m, 1H), 7.83 – 7.80 (m, 1H), 7.53 – 7.47 (m, 3H), 7.37 (t, *J* = 8.01 Hz, 1H), 6.90 (d, *J* = 7.48 Hz, 1H), 4.92 (s, 2H), 3.70 – 3.38 (m, 10H), 2.34 – 2.25 (m, 2H), 1.89 – 1.84 (m, 2H), 1.68 – 1.59 (m, 2H), 1.51 – 1.43 (m, 2H); ^13^C NMR (151 MHz, CDCl_3_) δ 171.5*, 171.3*, 167.0*, 166.8*, 153.3*, 153.2*, 134.7, 127.9*, 127.8*, 126.8, 126.0*, 125.9*, 125.8, 125.4, 121.72*, 121.68*, 121.5*, 121.4*, 105.3*, 105.2*, 68.5*, 68.2*, 45.8*, 45.6*, 45.3*, 44.9*, 42.4*, 42.3*, 41.9*, 41.3*, 33.7, 33.0, 32.6, 28.0, 24.3; HPLC: t*_R_* = 13.52 min, purity 94.0%, Method I; HRMS (MALDI) m/z calcd for C_22_H_28_N_2_O_3_^+^ (M+H^+^) 447.1278, found 447.1292.

### 7-Bromo-1-(4-(2-(naphthalen-1-yloxy)acetyl)piperazin-1-yl)heptan-1-one (59)

The title compound was synthesized according to the procedure described for **4** using 7-bromoheptanoic acid (183 mg, 0.87 mmol) to afford 45 mg (11%) of an orange oil after purification by preparative HPLC. ^1^H NMR (600 MHz, CDCl_3_) δ 8.21 (d, *J* = 7.2 Hz, 1H), 7.86 – 7.78 (m, 1H), 7.55 – 7.46 (m, 3H), 7.41 – 7.34 (m, 1H), 6.94 – 6.87 (m, 1H), 4.92 (s, 2H), 3.73 – 3.55 (m, 6H), 3.47 – 3.36 (m, 4H), 2.36 – 2.23 (m, 2H), 1.89 – 1.81 (m, 2H), 1.67 – 1.57 (m, 2H), 1.49 – 1.40 (m, 2H), 1.40 – 1.29 (m, 2H); ^13^C NMR (151 MHz, CDCl_3_) δ 171.7*, 171.5*, 167.0*, 166.8*, 153.2*, 153.1*, 134.6, 127.8*, 127.6*, 126.7, 125.84*, 125.78*, 125.7, 125.3, 121.6, 121.4*, 121.3*, 105.2*, 105.1*, 68.4*, 68.0*, 45.7*, 45.5*, 45.2*, 42.3*, 42.2*, 41.8*, 41.2*, 33.8, 33.0, 32.5, 28.4, 27.9, 24.9; HPLC: t*_R_* = 14.02 min, purity 96.7%, Method D; HRMS (MALDI) m/z calcd for C_23_H_30_BrN_2_O_3_^+^ (M+H^+^) 461.1434, found 461.1447.

### 8-Bromo-1-(4-(2-(naphthalen-1-yloxy)acetyl)piperazin-1-yl)heptan-1-one (60)

The title compound was synthesized according to the procedure described for **4** using 7-bromooctanoic acid (41 mg, 0.19 mmol) to afford 38 mg (43%) of a white solid after purification by flash chromatography (75-85% EtOAc in heptane). R_f_ = 0.32 (EtOAc:Heptane; 9:1); ^1^H NMR (600 MHz, CDCl_3_) δ 8.23 – 8.18 (m, 1H), 7.81 (d, *J* = 7.3 Hz, 1H), 7.55 – 7.44 (m, 3H), 7.36 (t, *J* = 8.0 Hz, 1H), 6.89 (d, *J* = 7.7 Hz, 1H), 4.91 (s, 2H), 3.74 – 3.54 (m, 6H), 3.48 – 3.41 (m, 2H), 3.38 (t, *J* = 7.3 Hz, 2H), 2.36 – 2.19 (m, 2H), 1.89 – 1.78 (m, 2H), 1.65 – 1.53 (m, 2H), 1.48 – 1.22 (m, 6H); ^13^C NMR (101 MHz, CDCl_3_) δ 171.8**\***, 171.7**\***, 167.0**\***, 166.8**\***, 153.3, 134.7, 127.9**\***, 127.8**\***, 126.8, 125.81**\***, 125.80**\***, 125.4, 121.4, 121.7, 121.5, 105.2, 68.5**\***, 68.2**\***, 45.8**\***, 45.6**\***, 45.3**\***, 42.4**\***, 41.8**\***, 41.3**\***, 34.0, 33.2, 32.8, 29.3, 28.6, 28.1, 25.1; HPLC: t*_R_* = 8.58 min, purity 94.4%, Method I; HRMS (MALDI) m/z calcd for C_24_H_31_BrN_2_O_3_Na^+^ (M+Na^+^) 497.1410, found 497.1418.

### 1-(4-(2-(Naphthalen-1-yloxy)acetyl)piperazin-1-yl)octan-1-one (61)

The title compound was synthesized according to the procedure described for **4** using octanoic acid (20 μL, 0.12 mmol) and DMF (1 mL) as the solvent to afford 39 mg (76%) of a clear oil after purification by flash chromatography (70% EtOAc in heptane). R_f_ = 0.36 (EtOAc:Heptane; 6:1); ^1^H NMR (600 MHz, CDCl_3_) δ 8.21 (d, *J* = 7.9 Hz, 1H), 7.85 – 7.79 (m, 1H), 7.53 – 7.45 (m, 3H), 7.40 – 7.33 (m, 1H), 6.92 – 6.87 (m, 1H), 4.91 (s, 2H), 3.74 – 3.31 (m, 8H), 2.43 – 2.13 (m, 2H), 1.33 – 1.24 (m, 10H), 0.87 (t, *J* = 6.7 Hz, 3H); ^13^C NMR (151 MHz, CDCl_3_) δ 167.0*, 166.8*, 153.3*, 153.2*, 134.7, 127.9*, 127.8*, 126.8, 125.95*, 125.87*, 125.8*, 125.4, 121.64*, 121.56*, 121.5*, 105.3*, 105.2*, 68.5*, 68.2*, 45.9*, 45.6*, 45.4*, 42.5*, 42.3*, 41.8*, 41.3*, 33.5, 31.8, 29.8, 29.5, 29.2, 25.3, 22.7, 14.2; HPLC: t*_R_* = 14.76 min, purity 99.6%, Method D; HRMS (MALDI) m/z calcd for C_24_H_32_N_2_O_3_Na^+^ (M+Na^+^) 419.2305, found 419.2308.

### 1-(4-(2-(Naphthalen-1-yloxy)acetyl)piperazin-1-yl)nonan-1-one (62)

The title compound was synthesized according to the procedure described for **4** using nonanoic acid (20 μL, 0.12 mmol) and DMF (0.8 mL) as the solvent to afford 18 mg (40%) of a clear oil after purification by flash chromatography (70% EtOAc in heptane). R_f_ = 0.37 (EtOAc:Heptane; 6:1); ^1^H NMR (600 MHz, CDCl_3_) δ 8.21 (d, *J* = 7.9 Hz, 1H), 7.84 – 7.78 (m, 1H), 7.53 – 7.45 (m, 3H), 7.40 – 7.33 (m, 1H), 6.92 – 6.87 (m, 1H), 4.91 (s, 2H), 3.71 – 3.54 (m, 6H), 3.46 – 3.33 (m, 2H), 2.34 – 2.21 (m, 2H), 1.64 – 1.53 (m, 2H), 1.30 – 1.20 (m, 10H), 0.87 (t, *J* = 7.0 Hz, 3H); ^13^C NMR (151 MHz, CDCl_3_) δ 172.0*, 171.9*, 167.0*, 166.8*, 153.3*, 153.2*, 134.7, 127.9*, 127.8*, 126.8, 125.94*, 125.87*, 125.8*, 125.4, 121.7*, 121.6*, 121.5*, 105.3* 105.2*, 68.5*, 68.1*, 45.6*, 45.6*, 45.4*, 42.4*, 42.3*, 41.8*, 41.3*, 33.4, 31.9, 29.53, 29.46, 29.2, 25.3, 22.7, 14.2; HPLC: t*_R_* = 15.29 min, purity 95.8%, Method D; HRMS (MALDI) m/z calcd for C_25_H_34_N_2_O_3_Na^+^ (M+Na^+^) 433.2461, found 433.2458.

### 1-(4-(2-(Naphthalen-1-yloxy)acetyl)piperazin-1-yl)decan-1-one (63)

The title compound was synthesized according to the procedure described for **4** using decanoic acid (68 mg, 0.39 mmol) and DMF (0.8 mL) as the solvent to afford 121 mg (73%) of a faint orange oil after purification by flash chromatography (60-75% EtOAc in heptane). R_f_ = 0.86 (MeOH:DCM; 1:9); ^1^H NMR (600 MHz, CDCl_3_) δ 8.21 (d, *J* = 7.6 Hz, 1H), 7.85 – 7.79 (m, 1H), 7.54 – 7.46 (m, 3H), 7.40 – 7.33 (m, 1H), 6.93 – 6.87 (m, 1H), 4.92 (s, 2H), 3.72 – 3.55 (m, 6H), 3.48 – 3.32 (m, 1H), 2.38 – 2.17 (m, 2H), 1.63 – 1.55 (m, 2H), 1.34 – 1.20 (m, 12H), 0.87 (t, *J* = 7.1 Hz, 3H); ^13^C NMR (151 MHz, CDCl_3_) δ 172.1*, 171.9*, 167.0*, 166.8*, 153.3*, 153.2*, 134.7, 127.9*, 127.8*, 126.8, 126.0*, 125.9*, 125.8*, 125.4, 121.70*, 121.68*, 121.6*, 121.5*, 105.3*, 105.2*, 68.5*, 68.2*, 45.9*, 45.6*, 45.4*, 42.4*, 42.3*, 41.8*, 41.3*, 33.4, 32.0, 29.57. 29.56, 29.53, 29.4, 25.3, 22.8, 14.2; HPLC: t*_R_* = 12.46 min, purity 97.29%, Method C; HRMS (MALDI) m/z calcd for C_26_H_36_N_2_O_3_Na^+^ (M+Na^+^) 447.2618, found 447.2624.

### 1-(4-(2-(Naphthalen-1-yloxy)acetyl)piperazin-1-yl)dodecan-1-one (64)

The title compound was synthesized according to the procedure described for **4** using lauric acid (16 mg, 0.18 mmol) and DMF (0.8 mL) as the solvent to afford 56 mg (67%) of a white solid after purification by flash chromatography (70% EtOAc in heptane). Melting Point Range: 84.0-87.5 °C; R_f_ = 0.32 (EtOAc:Heptane; 7:3); ^1^H NMR (400 MHz, CDCl_3_) δ 8.24 – 8.18 (m, 1H), 7.84 – 7.77 (m, 1H), 7.55 – 7.44 (m, 3H), 7.36 (t, *J* = 7.9 Hz, 1H), 6.89 (d, *J* = 7.6 Hz, 1H), 4.91 (s, 2H), 3.65* (s, 4H’), 3.58* (s, 2H”), 3.43* (s, 1.3H’), 3.38* (s, 0.7H”), 2.34 – 2.20 (m, 2H), 1.64 – 1.53 (m, 2H), 1.38 – 1.16 (m, 16H), 0.87 (t, *J* = 6.8 Hz, 3H); ^13^C NMR (101 MHz, CDCl_3_) δ 172.0, 166.9, 153.3, 134.7, 127.8, 126.8, 125.8, 125.4, 121.7, 121.5, 105.3, 68.4*, 68.2*, 45.7*, 45.6*, 45.5*, 42.3*, 41.8*, 41.3*, 33.4, 32.0, 29.7, 29.6, 29.54, 29.51, 29.4, 25.3, 22.8, 14.2; HPLC: t*_R_* = 11.96 min, purity 99.3%, Method A; HRMS (MALDI) m/z calcd for C_28_H_41_N_2_O_3_^+^ (M+H^+^) 453.3111, found 453.3104.

### 1-(4-(2-(Naphthalen-1-yloxy)acetyl)piperazin-1-yl)hexadecan-1-one (65)

The title compound was synthesized according to the procedure described for **4** using palmitic acid (23 mg, 90 µmol) and DMF (2 mL) as the solvent to afford 48 mg (85%) of a clear oil after purification by flash chromatography (70% EtOAc in heptane). R_f_ = 0.62 (EtOAc:Heptane; 7:3); ^1^H NMR (400 MHz, CDCl_3_) δ 8.25 – 8.17 (m, 1H), 7.85 – 7.78 (m, 1H), 7.56 – 7.45 (m, 3H), 7.36 (t, *J* = 8.0 Hz, 1H), 6.90 (d, *J* = 7.6 Hz, 1H), 4.91 (s, 2H), 3.66* (s, 4H’), 3.58* (s, 2H”), 3.44* (s, 1.3H’), 3.38* (s, 0.7H”), 2.37 – 2.20 (m, 2H), 1.63 – 1.55 (m, 2H), 1.36 – 1.22 (m, 24H), 0.87 (t, *J* = 6.5 Hz, 3H); ^13^C NMR (101 MHz, CDCl_3_) δ 172.0, 166.9, 153.3, 134.7, 127.8, 126.8, 125.8, 125.4, 121.7, 121.5, 105.3, 68.2, 45.6*, 42.4*, 41.8*, 41.3*, 33.4, 32.0, 29.81, 29.77, 29.73, 29.62, 29.56, 29.53, 29.48, 25.3, 22.8, 14.2; HPLC: t*_R_* = 14.85 min, purity 99.4%, Method A; HRMS (MALDI) m/z calcd for C_32_H_49_N_2_O_3_^+^ (M+H^+^) 509.3737, found 509.3733.

### *N*-(6-(4-(2-(Naphthalen-1-yloxy)acetyl)piperazin-1-yl)-6-oxohexyl)propionamide (66)

Step 1: *tert*-Butyl (6-(4-(2-(naphthalen-1-yloxy)acetyl)piperazin-1-yl)-6-oxohexyl)carbamate: The title compound was synthesized according to the procedure described for **4** using 6-((*tert*-butoxycarbonyl)amino)hexanoic acid (43 mg, 0.18 mmol) and DMF (0.9 mL) as the solvent for the reaction to afford 57 mg (64%) of a faint orange solid after purification by flash chromatography (70% EtOAc in heptane). R_f_ = 0.40 (MeOH:DCM; 1:9); ^1^H NMR (600 MHz, DMSO-*d_6_*) δ 8.22 (d, *J* = 8.2 Hz, 1H), 7.88 (d, *J* = 7.5 Hz, 1H), 7.56 – 7.47 (m, 3H), 7.40 (t, *J* = 7.9 Hz, 1H), 6.94 (d, *J* = 7.7 Hz, 1H), 5.05 (s, 2H), 3.62 – 3.43 (m, 8H), 2.89 (t, *J* = 7.0 Hz, 2H), 2.31 (t, *J* = 7.5 Hz, 2H), 1.48 (t, *J* = 7.6 Hz, 2H), 1.36 (s, 9H), 1.27 – 1.21 (m, 4H); ^13^C NMR (151 MHz, DMSO-*d_6_*) δ 170.8, 165.9, 155.5, 153.3, 134.0, 127.4, 126.4, 126.0, 125.4, 124.8, 121.5, 120.3, 105.6, 77.3, 66.4, 66.3*, 44.9, 44.6, 44.5*, 44.1, 41.5, 41.2, 41.1*, 40.7, 32.2, 29.3, 28.3, 26.0, 24.4; HPLC: t*_R_* = 13.05 min, purity 98.5%, Method D; HRMS (MALDI) m/z calcd for C_27_H_37_N_3_O_5_Na^+^ (M+Na^+^) 506.2625, found 506.2631.

Step 2: 6-(4-(2-(Naphthalen-1-yloxy)acetyl)piperazin-1-yl)-6-oxohexan-1-aminium 2,2,2-trifluoroacetate: The title compound was synthesized according to the procedure described for **Da** using *tert*-butyl (6-(4-(2-(naphthalen-1-yloxy)acetyl)piperazin-1-yl)-6-oxohexyl)carbamate (56 mg, 0.12 mmol) to yield 56 mg (97%) of a yellow solid. R_f_ = 0.30 (DCM:MeOH; 9:1); ^1^H NMR (600 MHz, DMSO-*d_6_*) δ 8.22 (d, *J* = 7.6 Hz, 1H), 7.88 (d, *J* = 7.7 Hz, 1H), 7.66 (s, 3H), 7.57 – 7.47 (m, 3H), 7.40 (t, *J* = 7.9 Hz, 1H), 6.94 (d, *J* = 7.7 Hz, 1H), 5.05 (s, 2H), 3.67 – 3.47 (m, 8H), 2.84 – 2.73 (m, 2H), 2.33 (t, *J* = 7.3 Hz, 2H), 1.59 – 1.44 (m, 4H), 1.35 – 1.27 (m, 2H); ^13^C NMR (151 MHz, DMSO-*d_6_*) δ 171.2, 166.5, 153.8, 134.5, 127.9, 127.0, 126.5, 125.9, 125.3, 122.0, 120.8, 106.1, 79.6, 66.9*, 66.8, 45.3, 45.0, 44.9, 44.6, 42.0, 41.7, 41.6, 41.2, 39.2, 32.4, 27.3, 26.0, 24.6; HPLC: t*_R_* = 9.33 min, purity 98.0%, Method D; HRMS (MALDI): m/z calcd for C_22_H_29_N_3_O_3_Na^+^ (M+Na^+^) 406.2101, found 406.2103.

Step 3: The title compound was synthesized according to the procedure described for **4** using 6-(4-(2-(naphthalen-1-yloxy)acetyl)piperazin-1-yl)-6-oxohexan-1-aminium 2,2,2-trifluoroacetate (26 mg, 53 μmol), propionic acid (4.0 μL, 52.9 μmol) and DMF (0.4 mL) as the solvent for the reaction to afford 11 mg (48%) of a white solid after purification by flash chromatography (0-10% MeOH in DCM). R_f_ = 0.48 (MeOH:DCM; 1:9); ^1^H NMR (600 MHz, CDCl_3_) δ 8.21 (d, *J* = 7.7 Hz, 1H), 7.84 – 7.79 (m, 1H), 7.54 – 7.46 (m, 3H), 7.40 – 7.33 (m, 1H), 6.93 – 6.87 (m, 1H), 5.61 (s, 1H), 4.92 (s, 2H), 3.71 – 3.53 (m, 6H), 3.48 – 3.33 (m, 2H), 3.27 – 3.21 (m, 2H), 2.34 – 2.24 (m, 2H), 2.17 (q, *J* = 7.6 Hz, 2H), 1.67 – 1.57 (m, 2H), 1.55 – 1.47 (m, 2H), 1.40 – 1.29 (m, 2H), 1.13 (t, *J* = 7.6 Hz, 3H); ^13^C NMR (151 MHz, CDCl_3_) δ 173.9, 171.7*, 171.5*, 167.0*, 166.9*, 153.3, 134.7, 127.9*, 127.8*, 126.8, 126.0*, 125.9*, 125.8*, 125.4, 121.7, 121.54*, 121.46*, 105.3*, 105.2*, 68.5*, 68.2*, 45.8*, 45.6*, 45.3*, 42.4*, 42.2*, 41.9*, 41.3*, 39.2, 33.0, 29.9, 29.4, 26.6, 24.5, 10.1; HPLC: t*_R_* = 12.06 min, purity 99.5%, Method D; HRMS (MALDI) m/z calcd for C_25_H_33_N_3_O_4_Na^+^ (M+Na^+^) 462.2363, found 462.2346.

### Methyl 8-(4-(2-(naphthalen-1-yloxy)acetyl)piperazin-1-yl)-8-oxooctanoate (67)

The title compound was synthesized according to the procedure described for **4** using 8-methoxy-8-oxooctanoic acid (65 mg, 0.34 mmol) to afford 65 mg (42%) of a yellow oil after purification by flash chromatography (80% EtOAc in heptane). ^1^H NMR (600 MHz, CDCl_3_) δ 8.24 – 8.18 (m, 1H), 7.85 – 7.79 (m, 1H), 7.55 – 7.46 (m, 3H), 7.40 – 7.33 (m, 1H), 6.93 – 6.87 (m, 1H), 4.92 (s, 2H), 3.76 – 3.52 (m, 9H), 3.51 – 3.35 (m, 2H), 2.40 – 2.23 (m, 4H), 1.71 – 1.52 (m, 4H), 1.41 – 1.25 (m, 4H); ^13^C NMR (151 MHz, CDCl_3_) δ 176.0, 174.3, 172.2*, 172.0*, 167.2*, 167.0*, 153.3, 134.7, 127.9*, 127.8*, 126.9, 125.9*, 125.8*, 125.4, 121.7, 121.5*, 121.4*, 105.4*, 105.3*, 68.5*, 68.1*, 51.6, 45.9*, 45.8*, 45.6*, 45.4*, 42.4*, 42.3*, 42.0*, 41.4*, 34.1, 33.4*, 33.2*, 29.1*, 29.01*, 28.95*, 28.8*, 25.1*, 25.0*, 24.8*, 24.6*; HPLC: t*_R_* = 12.90 min, purity 95.9%, Method D; HRMS (MALDI) m/z calcd for C_25_H_33_N_2_O_5_^+^ (M+H^+^) 441.2384, found 441.2397.

### Methyl 10-(4-(2-(naphthalen-1-yloxy)acetyl)piperazin-1-yl)-10-oxodecanoate (68)

A flame-dried round bottom flask was charged with 2-(naphthalen-1-yloxy)-1-(piperazin-1-yl)ethan-1-one (60 mg, 0.22 mmol), 10-methoxy-10-oxodecanoic acid (48 mg, 0.22 mmol), EDC · HCl (47 mg, 0.24 mmol) and HOBt hydrate (37 mg, 0.24 mmol). The flask was evacuated and backfilled with argon three times. Then, anhydrous DCM (1 mL) and DIPEA (77 μL, 0.44 mmol) were added. The resulting mixture was stirred at rt for overnight. The mixture was then diluted with EtOAc (2 mL) and water (2 mL). The organic phase was separated, and the aqueous phase was further extracted with EtOAc (2 x 2 mL). The combined organic phase was concentrated, and the crude was purified by flash column chromatography (60% EtOAc in heptane) to afford the title compound (37 mg, 34%) as a colorless oil. R_f_ = 0.26 (EtOAc:Heptane; 4:1); ^1^H NMR (600 MHz, CDCl_3_) δ 8.21 (d, *J* = 7.3 Hz, 1H), 7.81 (d, *J* = 7.6 Hz, 1H), 7.55 – 7.45 (m, 3H), 7.36 (t, *J* = 8.0 Hz, 1H), 6.90 (d, *J* = 7.6 Hz, 1H), 4.92 (s, 2H), 3.74 – 3.52 (m, 9H), 3.52 – 3.34 (m, 2H), 2.36 – 2.21 (m, 4H), 1.64 – 1.53 (m, 4H), 1.35 – 1.22 (m, 8H); ^13^C NMR (151 MHz, CDCl_3_) δ 174.4, 172.0, 167.0, 153.3, 134.7, 127.8, 126.8, 125.9, 125.4, 121.7, 121.5, 105.3, 68.5*, 68.2*, 51.6, 45.8*, 45.6*, 45.4*, 42.4*, 41.9*, 41.3*, 34.2, 33.4, 29.5, 29.3, 29.2, 29.2, 25.3, 25.0; HPLC: t*_R_* = 7.64 min, purity 95.0%, Method B; HRMS (MALDI) m/z calcd for C_27_H_37_N_2_O_5_ (M+H^+^) 469.2697, found 469.2708.

### 10-(4-(2-(Naphthalen-1-yloxy)acetyl)piperazin-1-yl)-10-oxodecanoic acid (69)

To a solution of **68** (22 mg, 47 μmol) in THF (232 μL) was added aq. LiOH (0.6 M, 232 μL, 0.14 mmol) and stirred at rt for 6 h. Then, the reaction mixture was acidified with aq. HCl (1.0 M) and extracted with EtOAc (3 x 2 mL). The combined organic phase was dried over anhydrous MgSO_4_, filtered, concentrated and the crude was purified by preparative HPLC to afford the title compound (2.0 mg, 9%) as a colorless oil. ^1^H NMR (600 MHz, CDCl_3_) δ 8.25 – 8.18 (m, 1H), 7.86 – 7.77 (m, 1H), 7.56 – 7.45 (m, 3H), 7.40 – 7.32 (m, 1H), 6.94 – 6.86 (m, 1H), 4.92 (s, 2H), 3.76 – 3.32 (m, 8H), 2.35 – 2.23 (m, 4H), 1.64 – 1.55 (m, 4H), 1.34 – 1.26 (m, 8H); ^13^C NMR (151 MHz, CDCl_3_) δ 176.3, 172.1, 167.3, 153.3, 134.8, 127.8, 126.9, 125.9, 125.8, 125.4, 121.7, 121.6, 105.3, 68.5*, 68.2*, 45.9*, 45.6*, 45.4*, 42.5*, 42.3*, 41.9*, 33.6, 33.3, 29.3, 29.1, 29.0, 25.2, 24.7; HPLC: t*_R_* = 12.74 min, purity 95.49%, Method B; HRMS (MALDI) m/z calcd for C_26_H_35_N_2_O_5_ (M+H^+^) 455.2540, found 455.2552.

### 1-(4-(2-(Naphthalen-1-yloxy)acetyl)piperazin-1-yl)dodecan-1-one (70)

The title compound was synthesized according to the procedure described for **4** using 12-methoxy-12-oxododecanoic acid (37 mg, 0.15 mmol) and DMF (2 mL) as the solvent to afford 72 mg (78%) of a brown oil after purification by flash chromatography (80% EtOAc in heptane). R_f_ = 0.26 (EtOAc:Heptane; 8:2); ^1^H NMR (600 MHz, CDCl_3_) δ 8.23 – 8.18 (m, 1H), 7.81 (d, *J* = 7.5 Hz, 1H), 7.53 – 7.45 (m, 3H), 7.36 (t, *J* = 8.0 Hz, 1H), 6.89 (d, *J* = 7.6 Hz, 1H), 4.91 (s, 2H), 3.65* (s, 7H’), 3.57* (s, 2H”), 3.43* (s, 1.3H’), 3.38* (s, 0.7H”), 2.29 (t, *J* = 7.5 Hz, 4H), 1.64 – 1.52 (m, 4H), 1.28 – 1.22 (m, 12H); ^13^C NMR (151 MHz, CDCl_3_) δ 174.4, 172.0*, 171.8*, 167.0*, 166.8*, 153.3, 134.7, 127.8, 126.8, 125.8, 125.4, 121.6, 121.5, 105.2, 68.5*, 68.2*, 51.5, 45.8*, 45.6*, 45.4*, 42.4*, 42.3*, 41.8*, 41.3*, 34.2, 33.4, 29.50, 29.48, 29.45, 29.3, 29.2, 25.3, 25.0; HPLC: t*_R_* = 9.59 min, purity 99.1%, Method A; HRMS (MALDI) m/z calcd for C_29_H_41_N_2_O_5_^+^ (M+H^+^) 497.3010, found 497.3006.

### 12-(4-(2-(Naphthalen-1-yloxy)acetyl)piperazin-1-yl)-12-oxododecanoic acid (71)

The title compound was synthesized according to the procedure described for **36** from **70** (20 mg, 40 μmol) to afford 14 mg (72%) of a clear oil after purification by flash chromatography (5% MeOH in DCM). R_f_ = 0.36 (EtOAc:Heptane; 9:1); ^1^H NMR (400 MHz, CDCl_3_) δ 8.25 – 8.16 (m, 1H), 7.85 – 7.78 (m, 1H), 7.56 – 7.43 (m, 3H), 7.36 (t, *J* = 8.0 Hz, 1H), 6.90 (d, *J* = 7.7 Hz, 1H), 4.92 (s, 2H), 3.67* (s, 4H’), 3.59* (s, 2H”), 3.45* (s, 1.3H’), 3.40* (s, 0.7H”), 2.39 – 2.22 (m, 4H), 1.67 – 1.52 (m, 4H), 1.36 – 1.22 (m, 12H); ^13^C NMR (101 MHz, CDCl_3_) δ 178.1, 177.3, 167.1, 153.3, 134.7, 127.8, 126.8, 125.9, 125.4, 121.7, 121.5, 105.3, 68.2, 45.7, 42.4, 34.0, 33.4, 29.4, 29.27, 29.26, 29.2, 29.1, 29.0, 25.3, 24.8; HPLC: t*_R_* = 7.31 min, purity 94.5%, Method H; HRMS (MALDI) m/z calcd for C_28_H_39_N_2_O_5_^+^ (M+H^+^) 483.2853, found 483.2849.

### Biological Assays

#### Cell culture

CHO-K1 cells (ATTC CCL61) were cultured in RPMI with 10% FBS, 180 μg/mL penicillin and 45 μg/ml streptomycin and incubated with 5% CO_2_ at 37°C. HEK293A cells were cultured in DMEM with 10% FBS, 180 μg/mL penicillin and 45 μg/mL streptomycin and incubated with 5% CO_2_ at 37°C. For both cell lines, cells were loosened by trypsin addition, split and transferred to new growth bottles (T75 from ThermoFisher) and supplied with new media once confluent (around every 3^rd^ day).

#### DNA constructs

For BRET assays human GPR183 CDS (corresponding to GenBank accession number L08177) was inserted in the pcDNA3.1+ plasmid with an N-terminal FLAG (M1)-tag and an upstream hemagglutinin signal peptide. The CAMYEL biosensor DNA and the BRET β-arrestin2 constructs were as described previously.^48,49^ For internalization assays, the human GPR183 CDS was inserted in the pcDNA5/FRT plasmid with an N-terminal SNAP tag downstream of a FLAG (M1)-tag and the hemagglutinin signal peptide.

#### CAMYEL cAMP Assay

CHO-K1 cells were seeded at 500,000 or 250,000 per well in a 6-well plate in growth medium. The following day or two days after, respectively, cells were transfected using Lipofectamine 2000 (Invitrogen) according to the manufacturers’ protocol. Growth medium was exchanged with 1 mL Opti-MEM and a mixture of 160 ng M1-GPR183 receptor DNA, and 840 ng CAMYEL sensor DNA was transfected into the cells of one well. Cells were incubated for approximately 24 h before assay was performed. On assay day cells were suspended in PBS + 5 mM glucose (3ml per well). 84 μL cell suspension was added to each well in an opaque flat bottomed 96-well plate. Subsequently, 10 μL 50 μM coelenterazine luciferase substrate in PBS was added to each well (5 μM in well) followed by 1 μL 100X ligand in DMSO. Five min post ligand addition, 5 μL 200 μM forskolin in PBS was added to each well (10 μM in well). At 40 min post ligand addition the BRET emissions at 525 nm (acceptor) and 485 nm (donor) were measured on a PerkinElmer EnVision plate reader and the BRET ratio was determined as acceptor counts/donor counts.

#### BRET β-arrestin2 Recruitment Assay

Cells were seeded and transfected like for the CAMYEL cAMP assay, but in this case with 330 ng M1-GPR183 receptor, 800 ng of mem-citrine-SH3 and 42 ng of Rluc-Arr3 DNA. The assay was performed as described for the cAMP BRET assay, except that 89 μL of cell suspension was added to each well, and forskolin was not added.

#### Real Time Internalization Assay

24 h before assay day HEK293A cells were seeded in white 384-well tissue-culture plates while simultaneously reverse transfected with 5 ng M1-SNAP-GPR183 DNA per well using Lipofectamine 2000 (Invitrogen). 20,000 cells in 30 μl DMEM growth medium were added to each poly-D-lysine coated well. The plate was incubated at 37°C with 5% CO_2_ until assay day. On assay day media was removed and 10 μL of 0.1 pmol/μL SNAP-Lumi4-Tb (Cisbio) in prewarmed Opti-MEM was added to each well followed by incubation at 37°C with 5% CO_2_ for 1 h. After incubation the wells were washed with 30 μL warm internalization buffer (HBSS (Gibco) + 1mM CaCl_2_ + 1mM MgCl_2_ + 20 mM HEPES + 0.1% BSA, pH = 7.4) 4 times to wash away unbound lumi-tag. 10 μL 50 μM Fluorescein-O′-acetic acid (Sigma-Aldrich) in prewarmed internalization buffer was added to each well, except for wells measuring donor signal only. 10 μL warm 2X ligand in internalization buffer was added using VIAFLO 384 channel electronic pipette (Integra) and the plate was immediately after inserted into Envision plate reader (Perkin Elmer) with a chamber temperature of 37 °C and measured. Signals emitted at 620 nm (Lumi4-Tb) and 520 nm (Fluorescein) were measured every 4 min for 92 min.

#### sDC Migration ChemoTX assay

sDCs were prepared as described previously and stored at −80°C.^50^ On assay day, cells were quickly thawn by the addition of warm X-vivo media (Lonza). They were subsequently spun down, resuspended, and left for acclimatization for 20-30 minutes. The system used for migration assay was the ChemoTX disposable chemotaxis system (Neuroprobe) with 5 μm pore size and 3.2 mm cell site diameter (101-5 series number). The ligands used for this experiment were diluted in media and placed in the bottom chamber (pipetting 31 μL) at varying concentrations and the filter containing the membranes was placed on top. 20 μL cell suspension was added on top of each chamber. The plate was incubated for 2.5 h for sDCs. After incubation the cells were carefully removed from the top of the filter using tissue paper. To dislodge the adhering sDCs, 15 μL 2 mM EDTA in PBS was added on top of each chamber and put in the fridge for 30 min. Filters were then carefully washed by holding it in a 45° angle and dripping PBS on top. The plate with the filter on top was spun down @ 500 x g for 10 min to make sure no cells were stuck to the bottom of the filter. The filter was then carefully removed. The contents of the bottom chamber were transferred to an opaque 96-well flat-bottomed plate with the use of a funnel plate (FP-1 Neuroprobe) and centrifugation @ 500 xg for 10 min. The number of cells migrated was quantified by the addition of 30 μL (1:1volume) Cell Titer Glo (Promega) to each well according to the manufacturers protocol. Luminescence was measured at 560 nm. Because the cell count added to each chamber was not always completely the same (not very variable however, ranging from 60-80.000 cells/chamber), the data was analyzed by normalizing to the background and maximum migration for each experiment. For the antagonist experiment testing the effect of **63** on 7α,25-OHC-induced dendritic cell chemotaxis, Corning Transwell 24 well plates with 5 μm pore size were used. Media with agonist at a fixed concentration known to induce chemotaxis, 7α,25-OHC (0.1 nM) or CCL19 (1 nM) and various concentrations of **63** (10 nM, 100 nM, 1 μM or 10 μM) was added to the bottom chamber (source) in a total volume of 600 μL, for the no ligand condition DMSO was added instead of **63**. sDCs added to the top chamber in the presence of **63** at a concentration matching the source or DMSO for the no ligand condition. The chambers were incubated for 2 hours in a humidified incubator at 37 degrees and 5% CO_2_. During the last 30 min. EDTA (final conc. of 0.5 mM) was added to the source chamber to detach transmigrated DCs from the filter. The filters were removed and the source suspension transferred to Eppendorf tubes and spun at 300 RCF for 5 min. The pellet was resuspended in 100 μL medium and transferred to a white 96 well plate with addition of 100 μL Cell Titer Glo. The signal was read as described above. CCL19 stimulates chemotaxis of sDCs through CCR7 expressed by these cells and was used as a counter screen for the specificity of **63**.

## Supporting Information

Cell viability, antagonists assays, counterscreens, NMR spectra and HPLC chromatograms (pdf)

## Acknowledgements

We thank Adrian Dragan, Katrine Schultz-Knudsen, and Maria Anastacia Mela for technical help with the in vitro cell-based experiments.

We thank the Lundbeck Foundation (grant # R307-2018-2950), the Novo Nordisk Foundation (grant # NNF21OC0069019) and the H2020 European Research Council (grant # 101055152) for generous financial support.

## Abbreviations

BRET: bioluminescence resonance energy transfer
cAMP: cyclic adenosine monophosphate
CHO-K1: Chinese hamster ovary-K1
DIPEA: *N*,*N*-diisopropylethylamine
DMEM: Dulbecco’s modified Eagle medium
EDC: 1-ethyl-3-(3-dimethylaminopropyl)carbodiimide
HEPES: 4-(2-hydroxyethyl)piperazine-1-ethane-sulfonic acid
HOBt: hydroxybenzotriazole
LLE: ligand lipophilicity efficiency
Rluc: Renilla Luciferase
sDCs: standard matured dendritic cells
TEA: triethylamine
YFP: yellow fluorescent protein
XPhos Pd G2: chloro(2-dicyclohexylphosphino-2′,4′,6′-triisopropyl-1,1′-biphenyl)[2-(2′-amino-1,1′-biphenyl)]palladium(II).

