## Supporting Information for "Development of Potent G Protein Pathway-Biased GPR183 Agonists"

### Cell viability

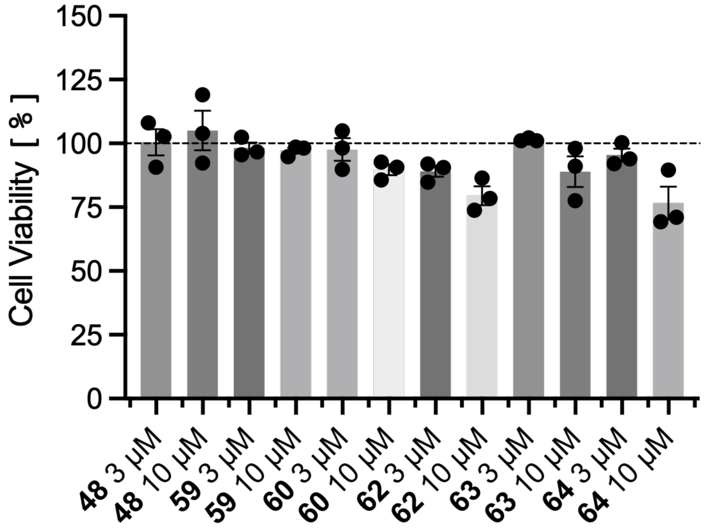

**Figure S1.** Cell viability as evaluated in the MTT assay.[1] Bars represent SEM.

### Antagonist screen of inactive compounds

Antagonism was evaluated in the CAMYEL assay as described in the Experimental section with the following modifications:83 µL of cell suspension were transferred to each well of an opaque, white flat-bottom 96-well plate. 1 µL of 100× ligand solution prepared as seven-point, 10-fold concentration range from 10 µM to 10 pM in DMSO was added to each well and were incubated for 30 minutes. Coelenterazine H was then added, corresponding to a final assay concentration of 5 µM. Subsequently, 1 µL of 100× agonist solution corresponding to the $EC_{80}$ concentration of the agonist prepared in DMSO was added to each well. Five minutes after agonist addition, 5 µL of 200 µM forskolin in assay buffer was added, resulting in a final forskolin concentration of 10 µM and a total assay volume of 100 µL per well. After 40 min after agonist addition, donor and acceptor emissions were measured at 485 and 525 nm, respectively, using a EnVision multimode plate reader (PerkinElmer). The BRET ratio was calculated as the acceptor emission divided by the donor emission.

**Table S2.** Antagonist screen of compounds with $pEC_{50}$ (mean ± SEM, n=2-3)

| | | Mean | SEM |
|---|---|---|---|
| **2** | TUG-2296 | 28.11 | 9.98 |
| **3** | TUG-2300 | 20.88 | 8.33 |
| **6** | TUG-2260 | 97.11 | 4.21 |
| **17** | TUG-2256 | 97.93 | 1.44 |
| **20** | TUG-2293 | 73.04 | 8.11 |
| **21** | TUG-2299 | 86.07 | 1.98 |
| **23** | TUG-2842 | 89.74 | 2.78 |
| **34** | TUG-2328 | 98.85 | 8.02 |
| **36** | TUG-2257 | 96.25 | 5.68 |
| **37** | TUG-2606 | 91.85 | 2.38 |
| **38** | TUG-2607 | 85.10 | 1.11 |
| **39** | TUG-2329 | 78.51 | 1.08 |
| **40** | TUG-2261 | 102.08 | 12.74 |
| **41** | TUG-2711 | 79.54 | 3.35 |
| **42** | TUG-2710 | 90.19 | 4.24 |
| **43** | TUG-2760 | 95.15 | 9.32 |
| **45** | TUG-2761 | 81.10 | 0.12 |
| **46** | TUG-2616 | 73.62 | 16.07 |
| **50** | TUG-2778 | 104.94 | 9.16 |
| **51** | TUG-2369 | 90.39 | 0.48 |
| **55** | TUG-2368 | 87.77 | 1.46 |
| **56** | TUG-2366 | 95.51 | 4.11 |
| **57** | TUG-2650 | 91.50 | 7.52 |
| **69** | TUG-2843 | 84.40 | 2.28 |

Inhibition of **63** by GPR183 antagonist

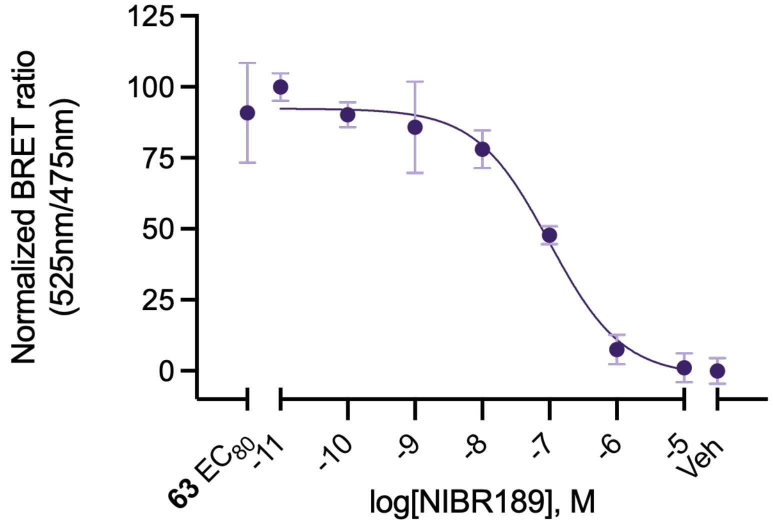

**Figure S2.** Inhibition of **63** by GPR183 antagonist NIBR189.

#### Counterscreen

##### IP-One Gq HTRF assay for screen on FFA1

Intracellular IP1 accumulation was measured using the homogeneous time-resolved fluorescence competitive immunoassay IP-One Gq HTRF kit (Revvity) according to the manufacturer's instructions. SNAP-human FFA1 expressing HEK cells were seeded at 1 × $10^4$ cells per well in poly-D-lysine coated 96-well plates and cultured overnight. Before the assay, the cells were washed with 100 µl of dPBS twice and 90 µl of assay buffer containing $LiCl_2$ was added and 10 µl of indicated agonist containing solution. The cells were incubated for 30 minutes for stimulation. Following the 30-minute stimulation period, cells were lysed and IP1 levels were determined in 384-well polystyrene microplates (Greiner Bio-One, 781074) using the IP-One HTRF detection reagents. HTRF signals were measured using a EnVision multimode plate reader (PerkinElmer) at emission wavelengths of 620 and 665 nm. The HTRF ratio was calculated from the fluorescence signals measured at 665 and 620 nm. IP1 concentrations were subsequently determined by correlating the 665/620 nm ratio with a previously generated IP1 standard curve.

#### Calcium mobilization assay for screen on FFA4

The calcium mobilization assay was performed using the ratiometric calcium indicator Fura-2 AM (Invitrogen, F1221). Cells stably expressing human FFA4-eYFP were seeded at $5 \times 10^5$ cells per well in black, clear flat-bottom 96-well polystyrene plates (Corning Falcon, 734-2480). For each experiment, technical replicates were seeded in triplicate. On the day of the assay, assay buffer was freshly prepared by supplementing HBSS with 20 mM HEPES, 1 mM $MgCl_2$, and 1 mM $CaCl_2$. Before the assay, the cells were washed twice with dPBS (Merck, D8537). After washing, 100 µL of assay buffer containing 3 µM Fura-2 AM was added, and the cells were incubated at 37 °C for 1 hour. Following incubation, the cells were washed once with 100 µL of dPBS. Subsequently, 50 µL of assay buffer was added, and the cells were incubated for an additional 15 minutes. Measurements were performed using a FlexStation 3 multimode plate reader (Molecular Devices). Basal fluorescence values were recorded first, followed by the addition of 50 µL of the indicated agonist diluted in assay buffer. Fluorescence was measured for approximately 2 minutes at intervals of approximately 4 seconds for each well. The Fura-2 ratio was calculated as the ratio of the emission signals obtained following excitation at 340 and 380 nm.

#### GRK activation assay for screen on CB1, GPR84 and LPA1

The intermolecular BRET measurements were performed to investigate GRK activation by Gβγ. 4 $\times 10^6$ cells were seeded into 10 cm dishes and transfected the following day using Lipofectamine 2000 (Invitrogen) according to the manufacturer's instructions. Transfection mixture included 2 µg of the indicated GPCR (CB1, GPR84, LPAR1), 1.2 µg of GRK-Nluc, 2.4 µg of Gαi, 1.6 µg of Gβ and 1.6 µg of Gγ plasmid DNA prepared in Opti-MEM with a DNA:Lipofectamine ratio of 1:2.5. The following day, 50,000 cells per well were seeded into a poly-D-lysine-coated opaque, white flat-bottom 96-well plate (Nunc, 136102).

For each experiment, technical replicates were seeded in triplicate. On the day of the assay, assay buffer was freshly prepared by supplementing HBSS with 20 mM HEPES, 1 mM $MgCl_2$ and 1 mM $CaCl_2$. Before the assay, the cells were washed twice with 100 µl of dPBS (Merck, D8537). After washing, 80 µl of assay buffer was added and 10 µl of the NLuc-substrate furimazine (Promega, N157B) diluted in assay buffer (1:100) was added. The cells were incubated for 5 minutes. The measurements were performed in a LUMIStar Omega plate reader (BMG LabTech). First, basal values were measured for 4 min every 30 seconds, followed by addition 10 µl of the indicated agonist diluted in the assay buffer and measurement of the stimulated values for 10 min. The BRET ratio was calculated as the acceptor emission divided by the donor emission.

**Table S1.** Screening of selected compounds on related receptors. Activation relative to 7α,25-OHC as mean ± SEM (n = 2-3)

| | Concentration | Mean (%) | SEM (%) |
|---|---|---|---|
| **FFA1** | | | |
| **48** | 3 µM | -0.12 | 2.84 |
| | 10 µM | 9.41 | 10.94 |
| **59** | 3 µM | -1.53 | 2.83 |
| | 10 µM | 12.13 | 9.10 |
| **60** | 3 µM | 0.37 | 4.84 |
| | 10 µM | -5.88 | 1.39 |
| **62** | 3 µM | -0.32 | 3.55 |
| | 10 µM | -2.20 | 1.82 |
| **63** | 3 µM | 4.09 | 4.90 |
| | 10 µM | 10.02 | 10.02 |
| **64** | 3 µM | -1.88 | 1.65 |
| | 10 µM | 1.42 | 2.70 |
| **FFA4** | | | |
| **48** | 3 µM | 1.87 | 1.65 |
| | 10 µM | -6.35 | 2.50 |
| **59** | 3 µM | 3.92 | 2.17 |
| | 10 µM | 1.53 | 6.07 |
| **60** | 3 µM | 1.17 | 5.87 |
| | 10 µM | -12.08 | 1.32 |
| **62** | 3 µM | 4.16 | 2.58 |
| | 10 µM | -3.89 | 1.22 |
| **63** | 3 µM | 0.24 | 4.76 |
| | 10 µM | 2.59 | 4.15 |
| **64** | 3 µM | -3.93 | 2.14 |
| | 10 µM | 0.16 | 0.47 |
| **LPA1** | | | |
| **48** | 3 µM | 4.53 | 1.45 |
| | 10 µM | 18.34 | 3.35 |
| **59** | 3 µM | 4.18 | 11.25 |
| | 10 µM | 13.38 | 13.53 |
| **60** | 3 µM | 13.57 | 0.18 |
| | 10 µM | 25.81 | 2.59 |
| **62** | 3 µM | 13.62 | 9.34 |
| | 10 µM | 33.45 | 8.15 |
| **63** | 3 µM | 8.36 | 4.64 |
| | 10 µM | 19.83 | 0.37 |
| **64** | 3 µM | 7.22 | 0.39 |
| | 10 µM | 11.29 | 0.47 |
| **GPR84** | | | |
| **48** | 3 µM | 12.10 | 7.70 |
| | 10 µM | 20.04 | 12.62 |
| **59** | 3 µM | -0.44 | 3.63 |
| | 10 µM | 9.20 | 4.74 |
| **60** | 3 µM | 16.83 | 11.69 |
| | 10 µM | 26.62 | 15.62 |
| **62** | 3 µM | 11.02 | 2.68 |
| | 10 µM | 28.61 | 12.24 |
| **63** | 3 µM | 14.68 | 8.45 |
| | 10 µM | 20.84 | 9.50 |

| | | | |
|---|---|---|---|
| **64** | 3 µM | 6.24 | 3.09 |
| | 10 µM | 11.91 | 4.95 |
| **CB1** | | | |
| **48** | 3 µM | 0.60 | 3.79 |
| | 10 µM | 1.61 | 10.31 |
| **59** | 3 µM | -0.96 | 8.94 |
| | 10 µM | -2.68 | 5.89 |
| **60** | 3 µM | -3.56 | 2.95 |
| | 10 µM | -6.19 | 8.32 |
| **62** | 3 µM | 1.79 | 4.48 |
| | 10 µM | 3.30 | 11.29 |
| **63** | 3 µM | 8.48 | 2.02 |
| | 10 µM | 4.57 | 2.58 |
| **64** | 3 µM | 13.19 | 1.27 |
| | 10 µM | 27.84 | 3.22 |

### NMR spectra and HPLC chromatograms

#### Compound 1 - TUG-2295

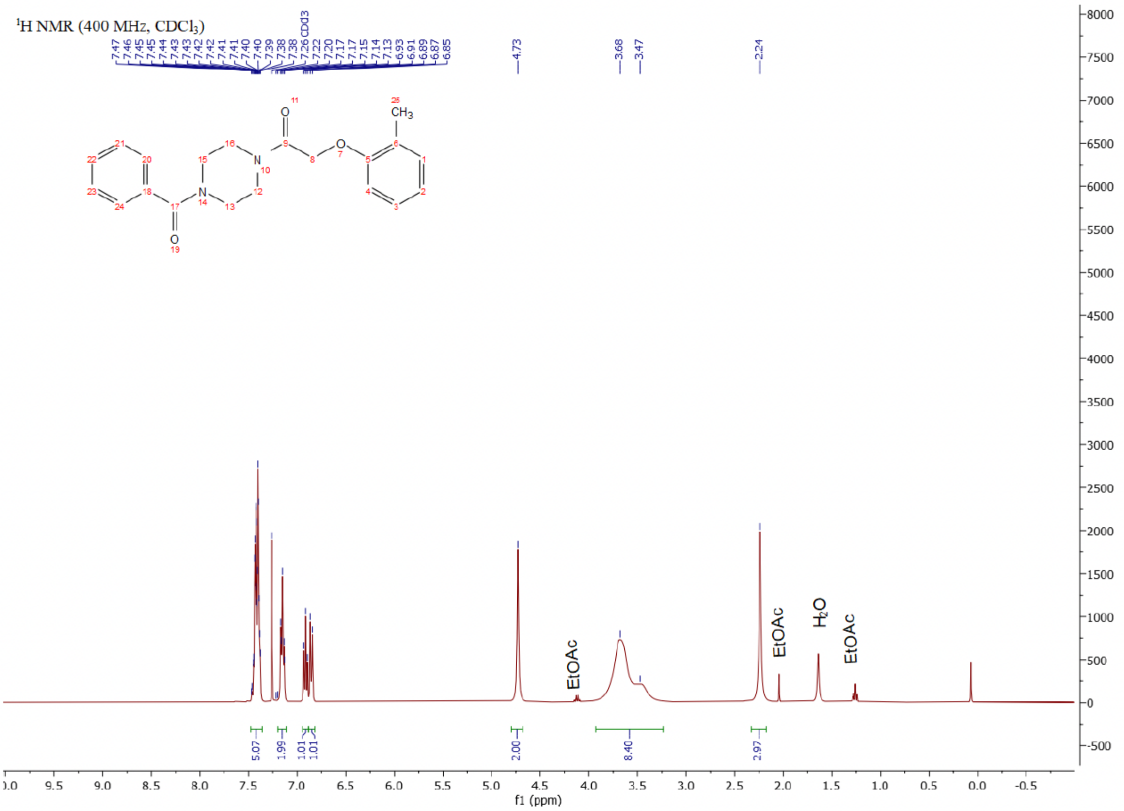

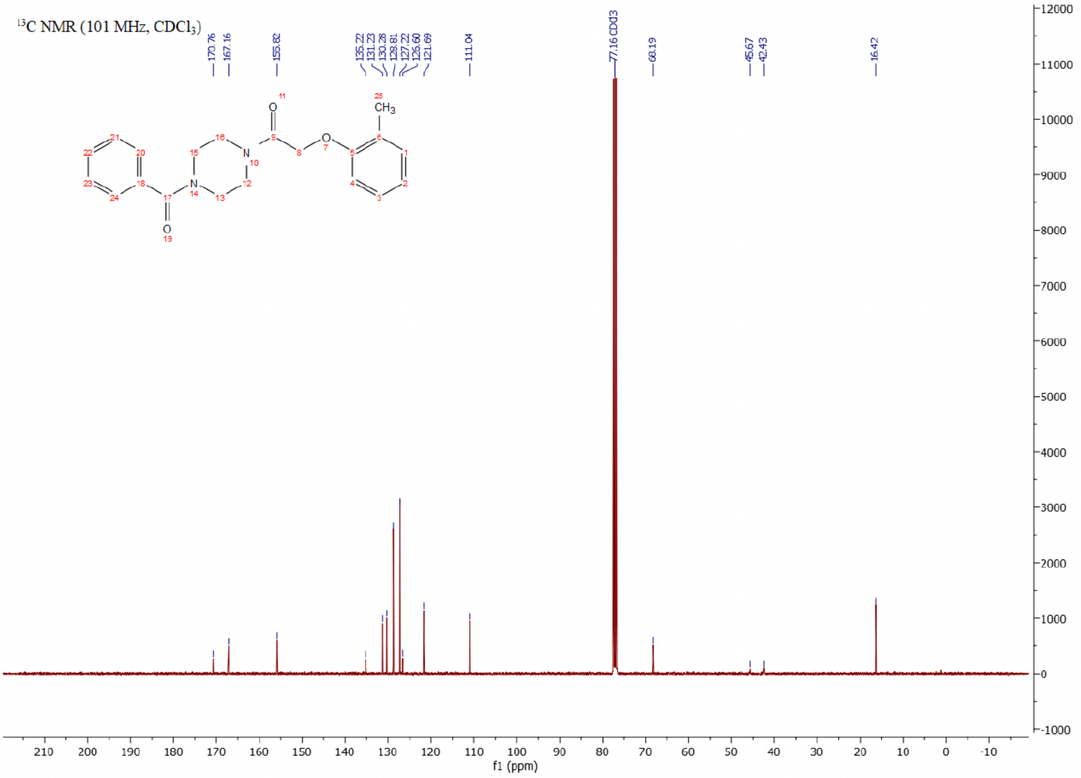

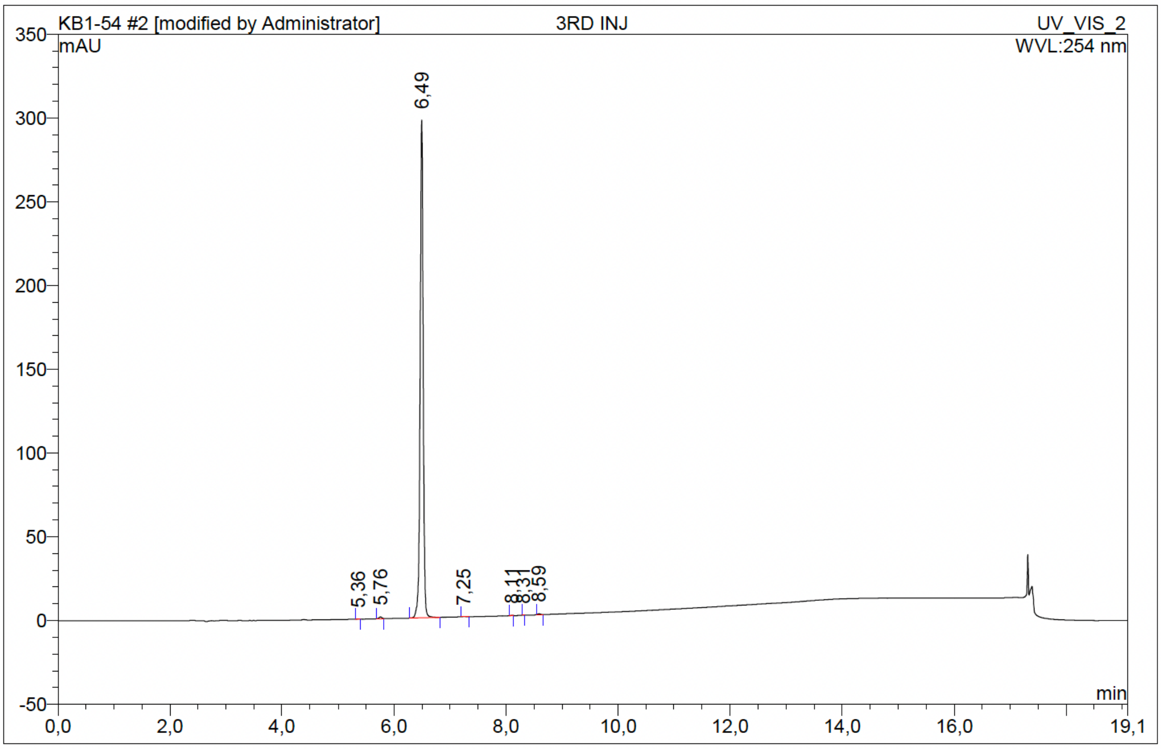

| No. | Ret.Time min | Peak Name | Height mAU | Area mAU*min | Rel.Area % | Amount | Resolution(EP) |
|---|---|---|---|---|---|---|---|
| 1 | 5,36 | n.a. | 0,066 | 0,003 | 0,02 | n.a. | 4,36 |
| 2 | 5,76 | n.a. | 0,972 | 0,056 | 0,29 | n.a. | 7,75 |
| 3 | 6,49 | n.a. | 297,183 | 19,106 | 99,46 | n.a. | 6,97 |
| 4 | 7,25 | n.a. | 0,128 | 0,009 | 0,05 | n.a. | 9,27 |
| 5 | 8,11 | n.a. | 0,128 | 0,005 | 0,03 | n.a. | 3,26 |
| 6 | 8,31 | n.a. | 0,081 | 0,003 | 0,01 | n.a. | 3,97 |
| 7 | 8,59 | n.a. | 0,523 | 0,027 | 0,14 | n.a. | n.a. |
| Total: | | | 299,081 | 19,209 | 100,00 | 0,000 | |

Compound 2 - TUG-2296

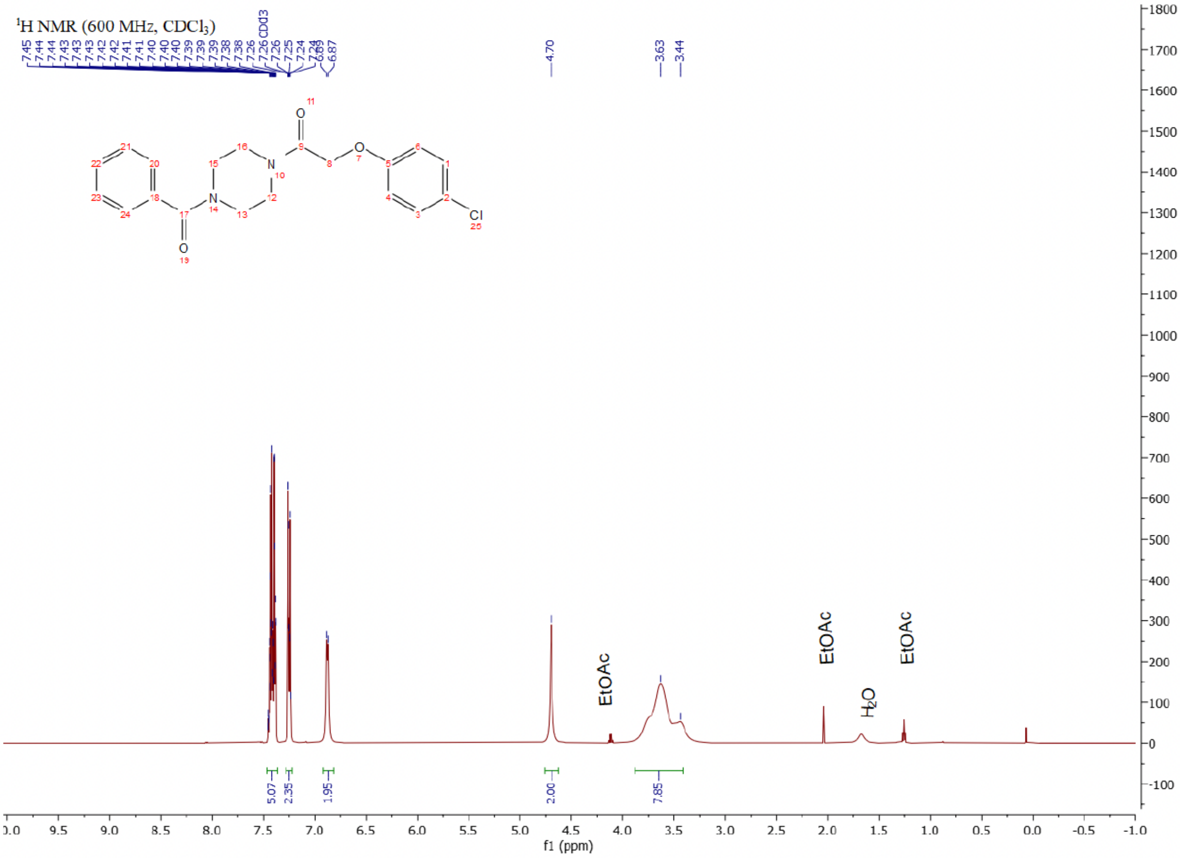

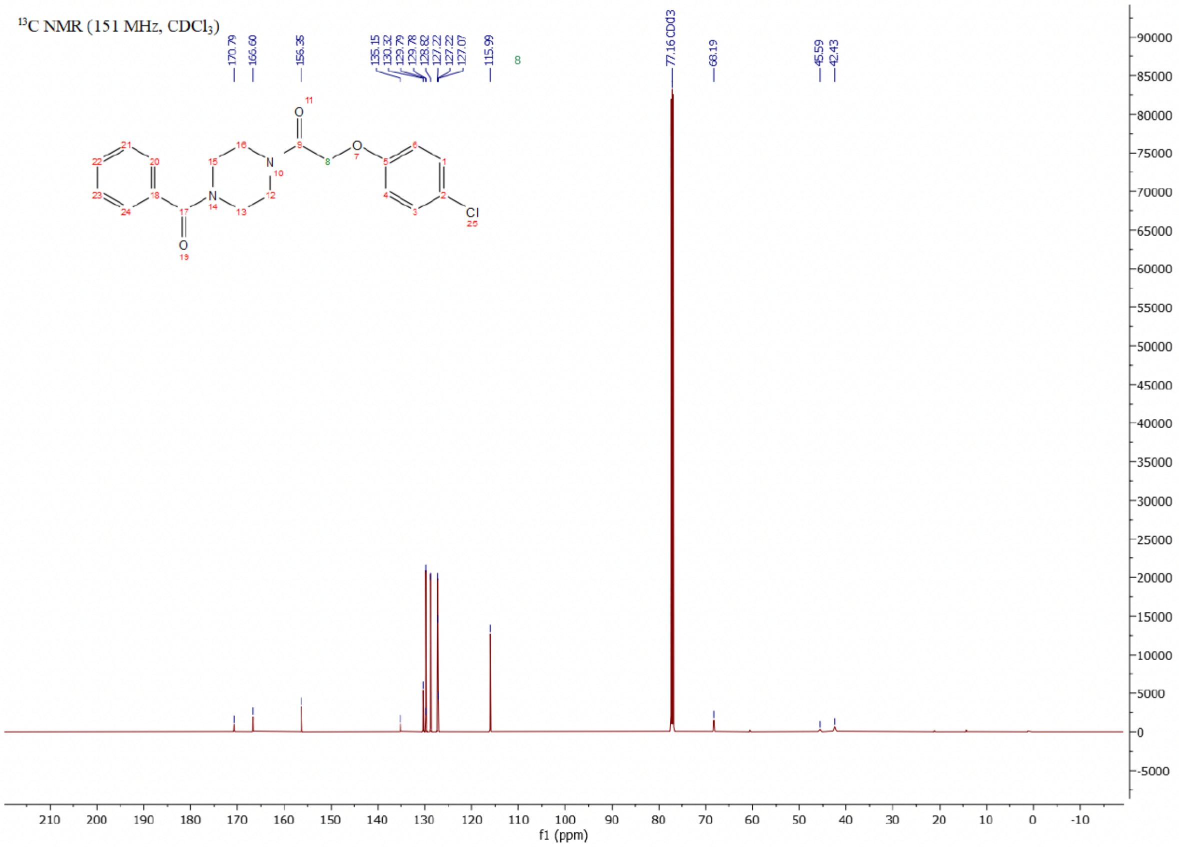

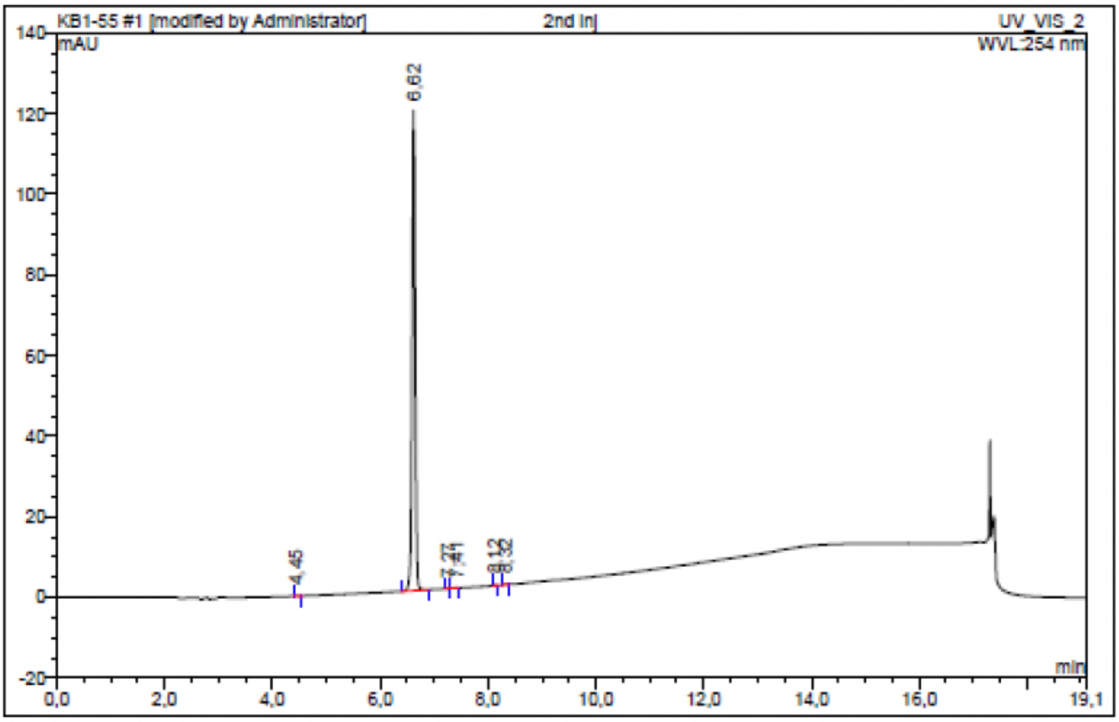

| No. | Ret.Time min | Peak Name | Height mAU | Area mAU*min | Rel.Area % | Amount | Resolution(EP) |
|---|---|---|---|---|---|---|---|
| 1 | 4,45 | n.a. | 0,137 | 0,008 | 0,10 | n.a. | 23,14 |
| 2 | 6,62 | n.a. | 119,128 | 7,610 | 99,54 | n.a. | 7,68 |
| 3 | 7,27 | n.a. | 0,063 | 0,002 | 0,03 | n.a. | 1,92 |
| 4 | 7,41 | n.a. | 0,111 | 0,006 | 0,08 | n.a. | 9,17 |
| 5 | 8,12 | n.a. | 0,169 | 0,008 | 0,10 | n.a. | 2,43 |
| 6 | 8,32 | n.a. | 0,200 | 0,011 | 0,15 | n.a. | n.a. |
| Total: | | | 119,808 | 7,645 | 100,00 | 0,000 | |

Compound 3 - TUG-2300

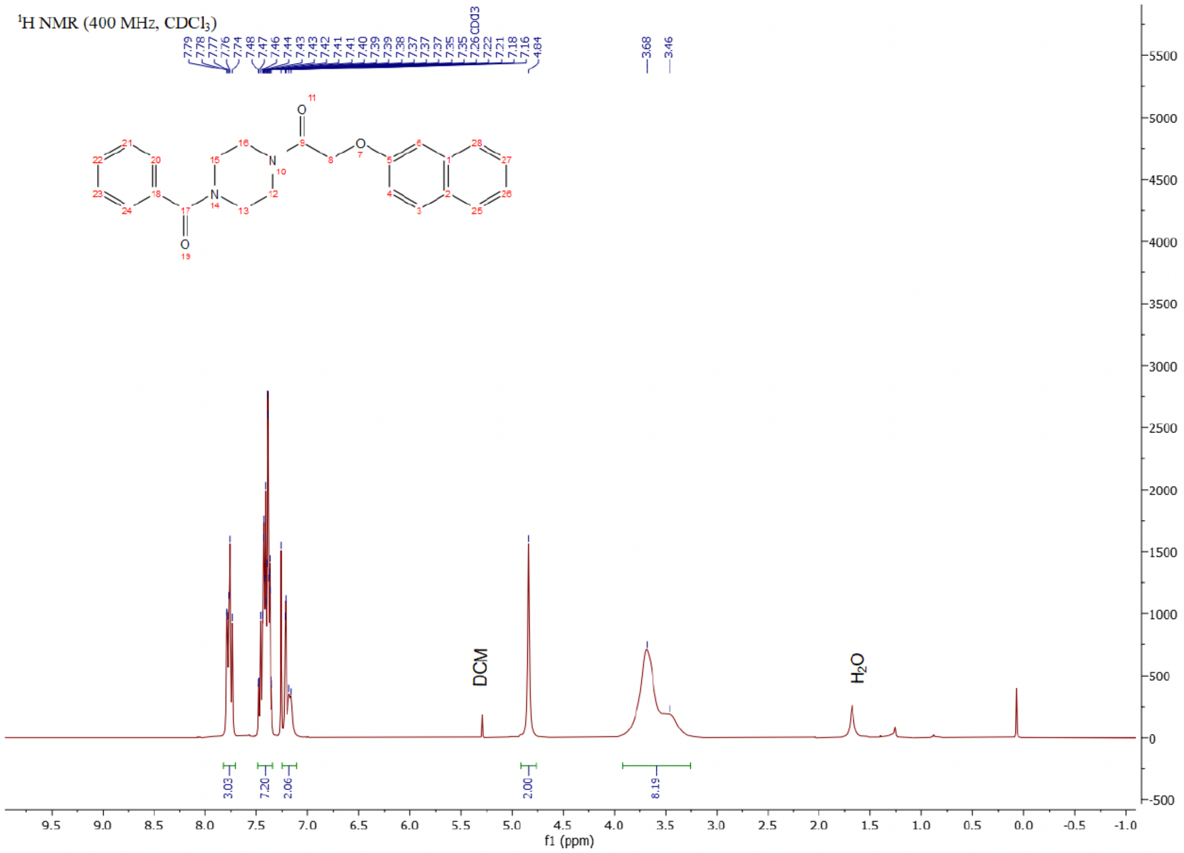

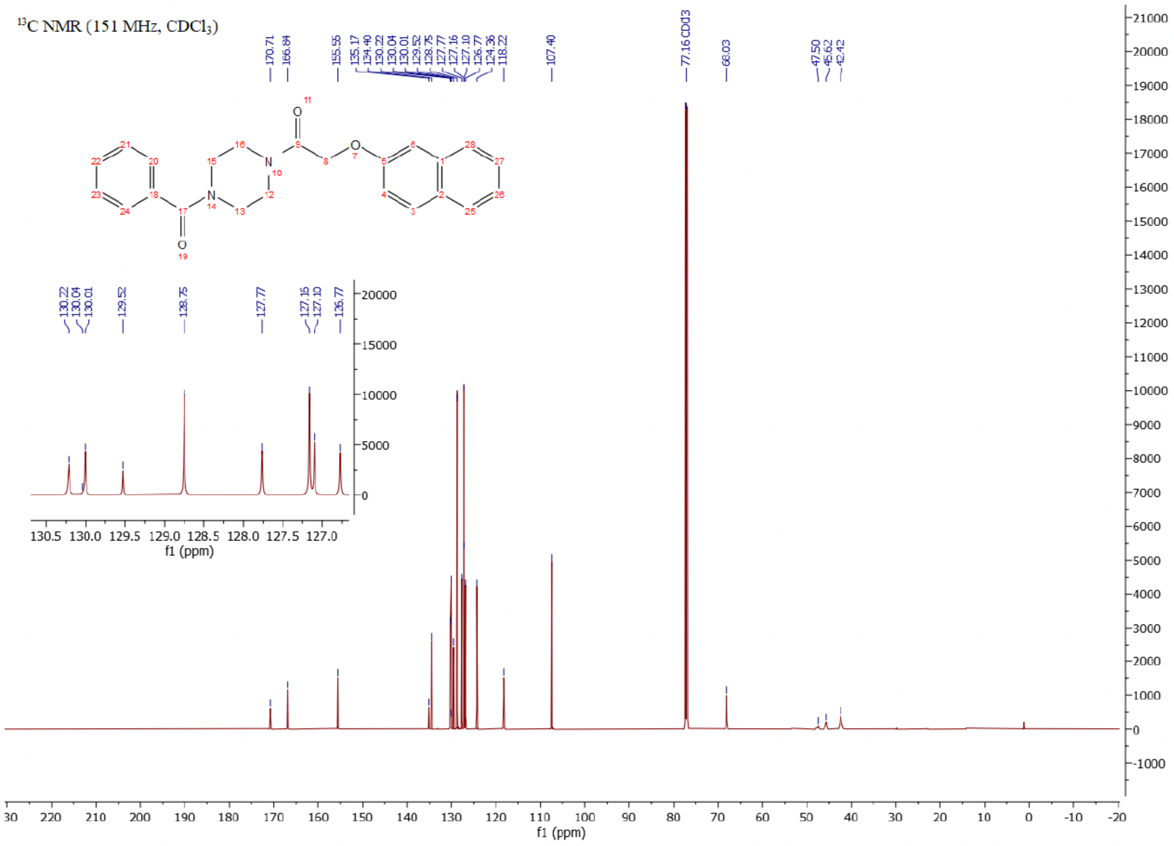

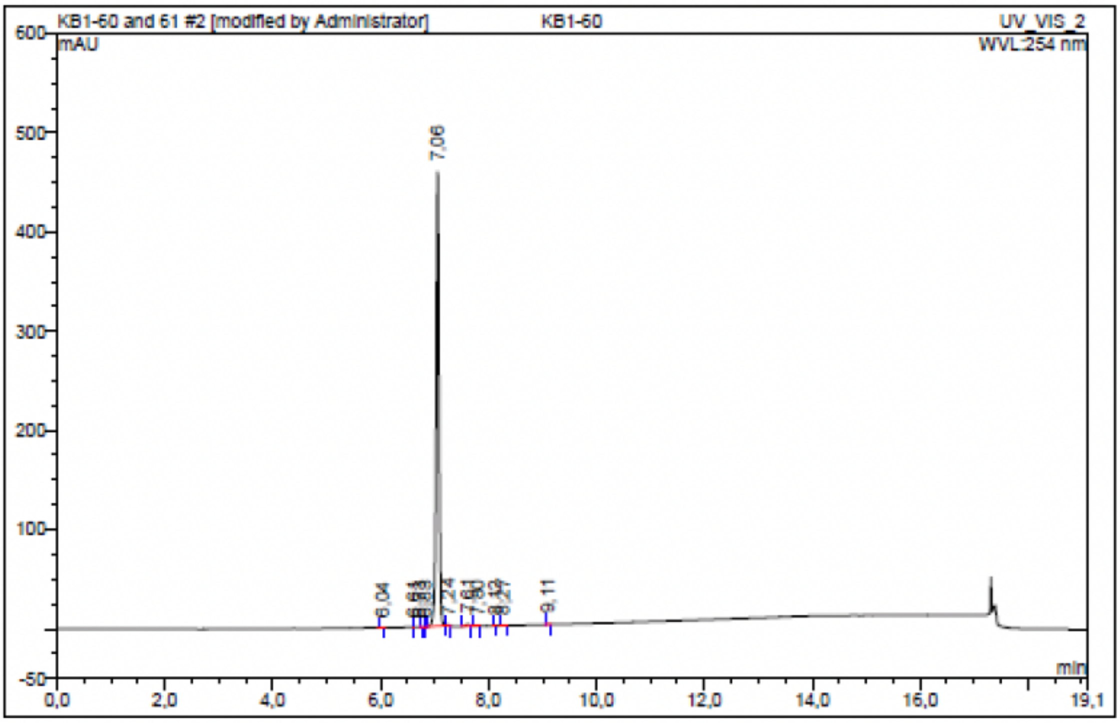

| No. | Ret.Time min | Peak Name | Height mAU | Area mAU*min | Rel.Area % | Amount | Resolution(EP) |
|---|---|---|---|---|---|---|---|
| 1 | 6,04 | n.a. | 0,102 | 0,009 | 0,03 | n.a. | 6,42 |
| 2 | 6,61 | n.a. | 0,007 | 0,000 | 0,00 | n.a. | 1,75 |
| 3 | 6,73 | n.a. | 0,024 | 0,002 | 0,01 | n.a. | 1,52 |
| 4 | 6,83 | n.a. | 0,010 | 0,000 | 0,00 | n.a. | 3,67 |
| 5 | 7,06 | n.a. | 457,429 | 29,025 | 99,19 | n.a. | 2,16 |
| 6 | 7,24 | n.a. | 0,867 | 0,041 | 0,14 | n.a. | 4,28 |
| 7 | 7,61 | n.a. | 1,875 | 0,114 | 0,39 | n.a. | 1,92 |
| 8 | 7,80 | n.a. | 0,237 | 0,015 | 0,05 | n.a. | 3,93 |
| 9 | 8,12 | n.a. | 0,226 | 0,009 | 0,03 | n.a. | 1,56 |
| 10 | 8,27 | n.a. | 0,381 | 0,027 | 0,09 | n.a. | 7,85 |
| 11 | 9,11 | n.a. | 0,400 | 0,022 | 0,08 | n.a. | n.a. |
| Total: | | | 461,558 | 29,263 | 100,00 | 0,000 | |

Compound 4 - TUG-2605

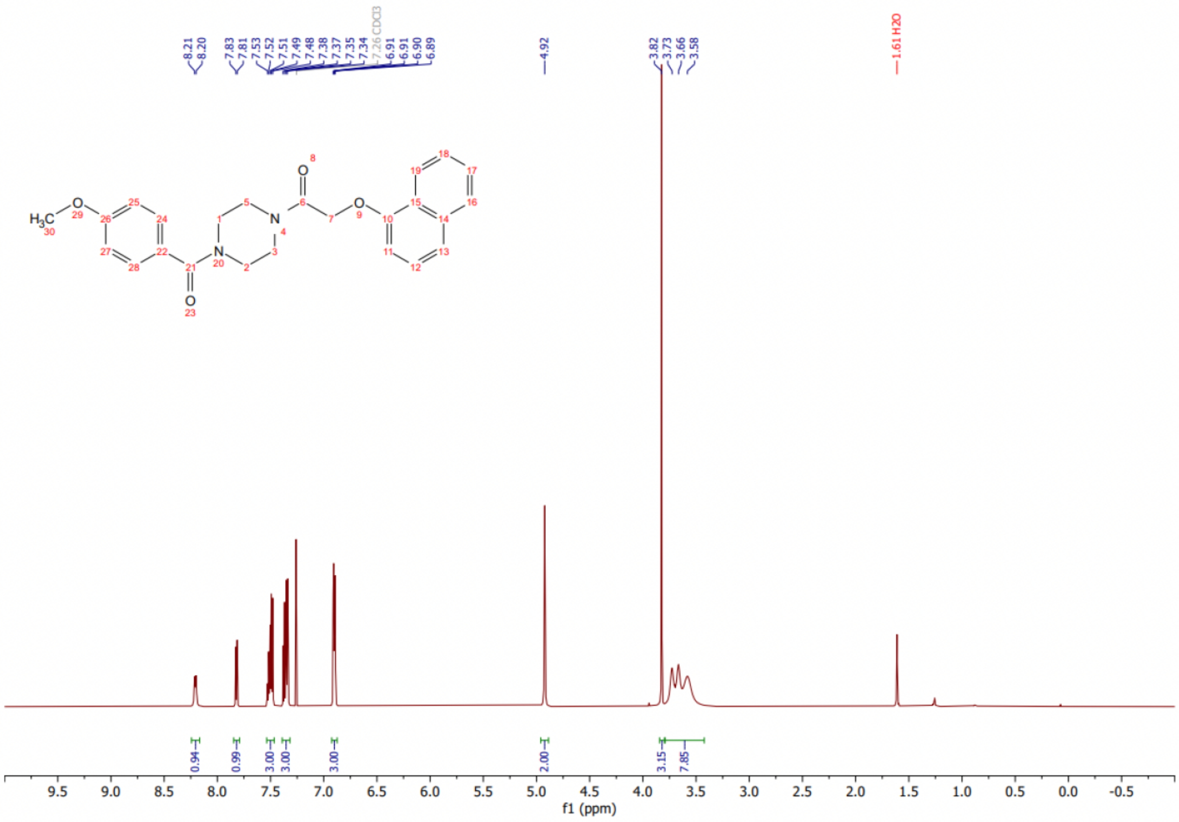

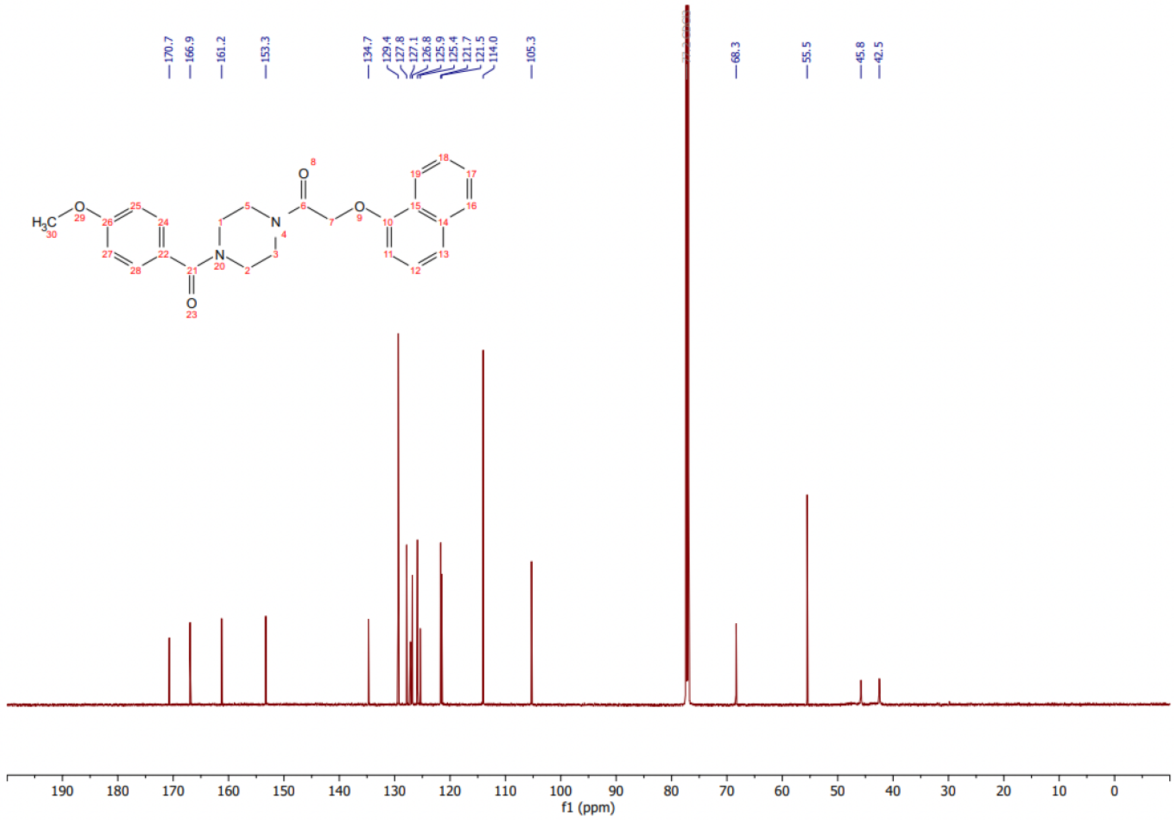

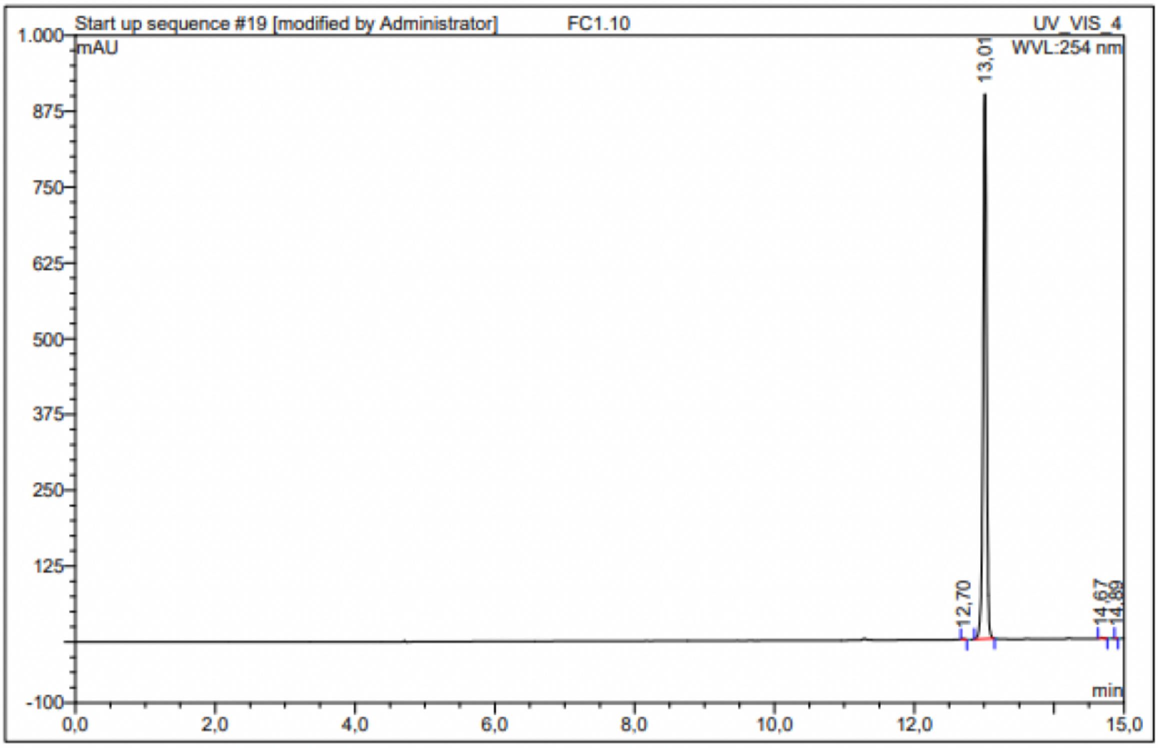

| No. | Ret.Time min | Peak Name | Height mAU | Area mAU*min | Rel.Area % | Amount | Resolution(EP) |
|---|---|---|---|---|---|---|---|
| 1 | 12,70 | n.a. | 1,049 | 0,049 | 0,09 | n.a. | 3,64 |
| 2 | 13,01 | n.a. | 898,213 | 52,559 | 99,74 | n.a. | 16,87 |
| 3 | 14,67 | n.a. | 1,293 | 0,086 | 0,16 | n.a. | 2,68 |
| 4 | 14,89 | n.a. | 0,074 | 0,002 | 0,00 | n.a. | n.a. |
| Total: | | | 900,629 | 52,697 | 100,00 | 0,000 | |

Compound 5 - TUG-2613

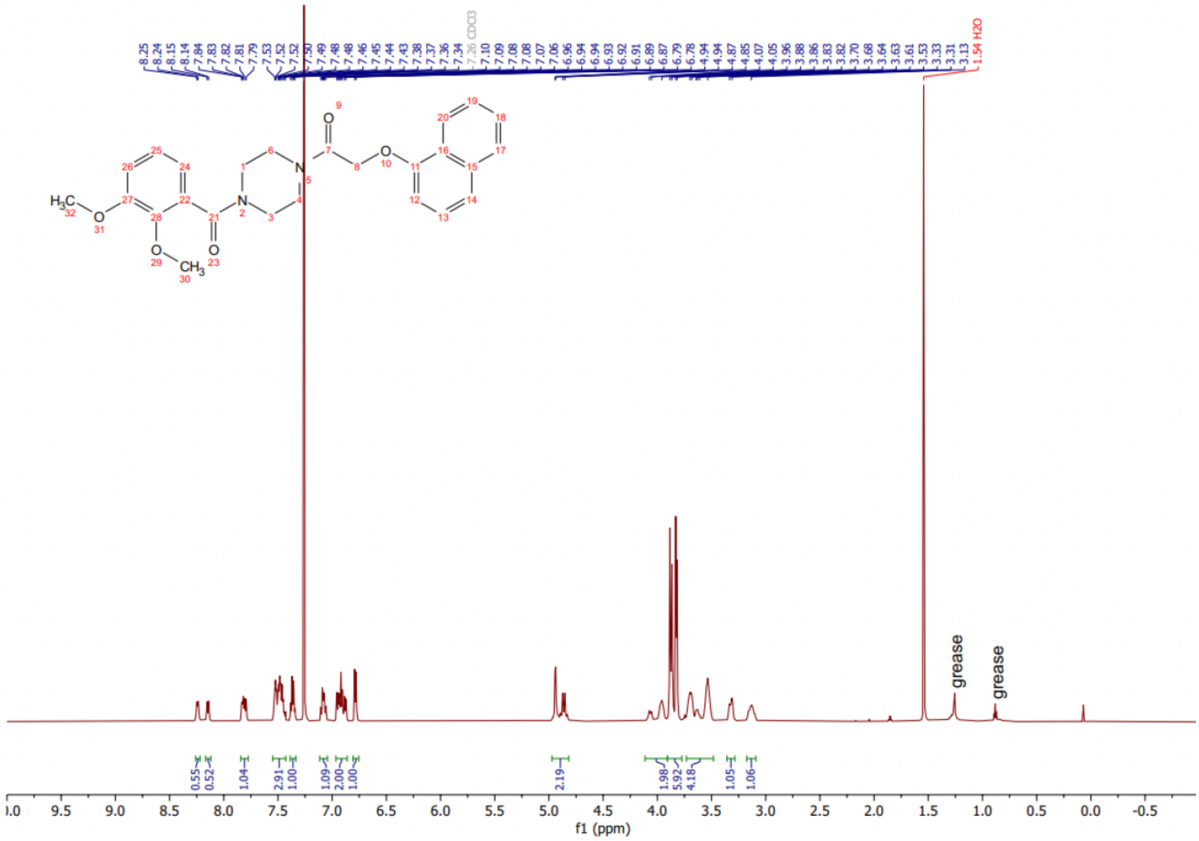

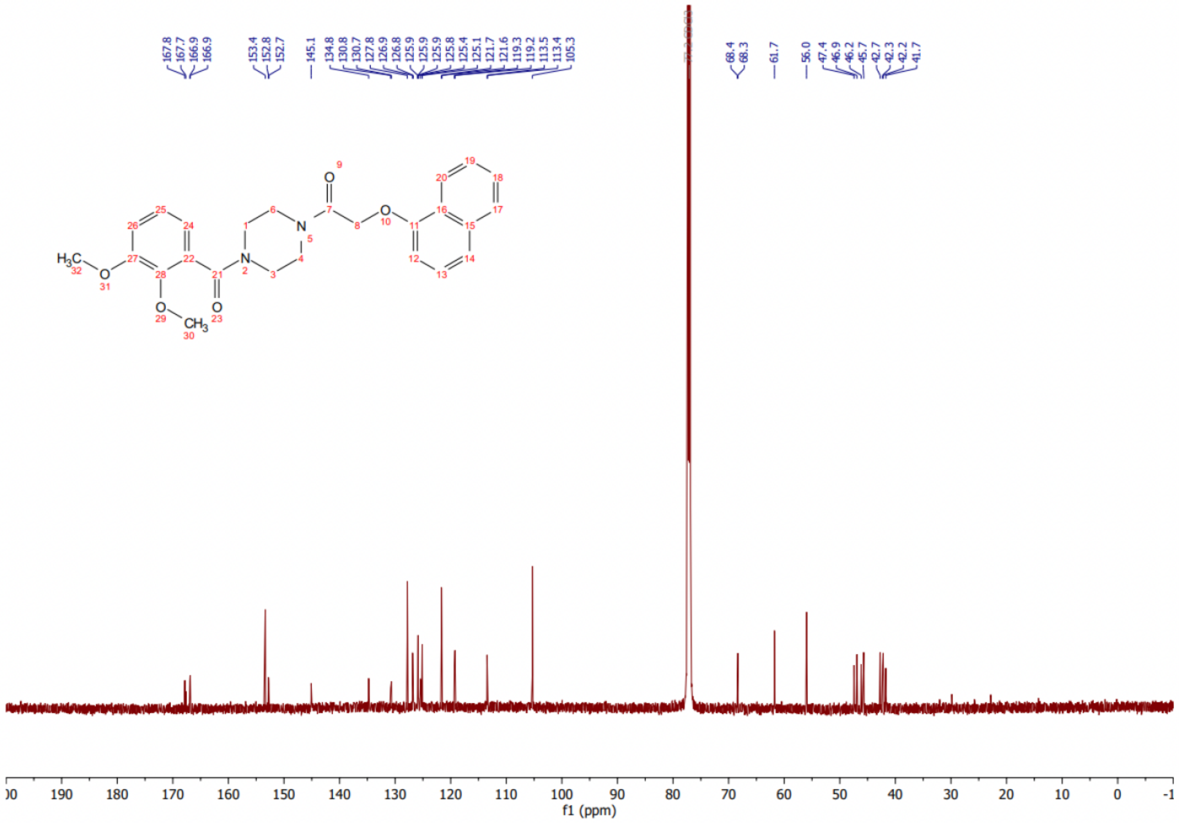

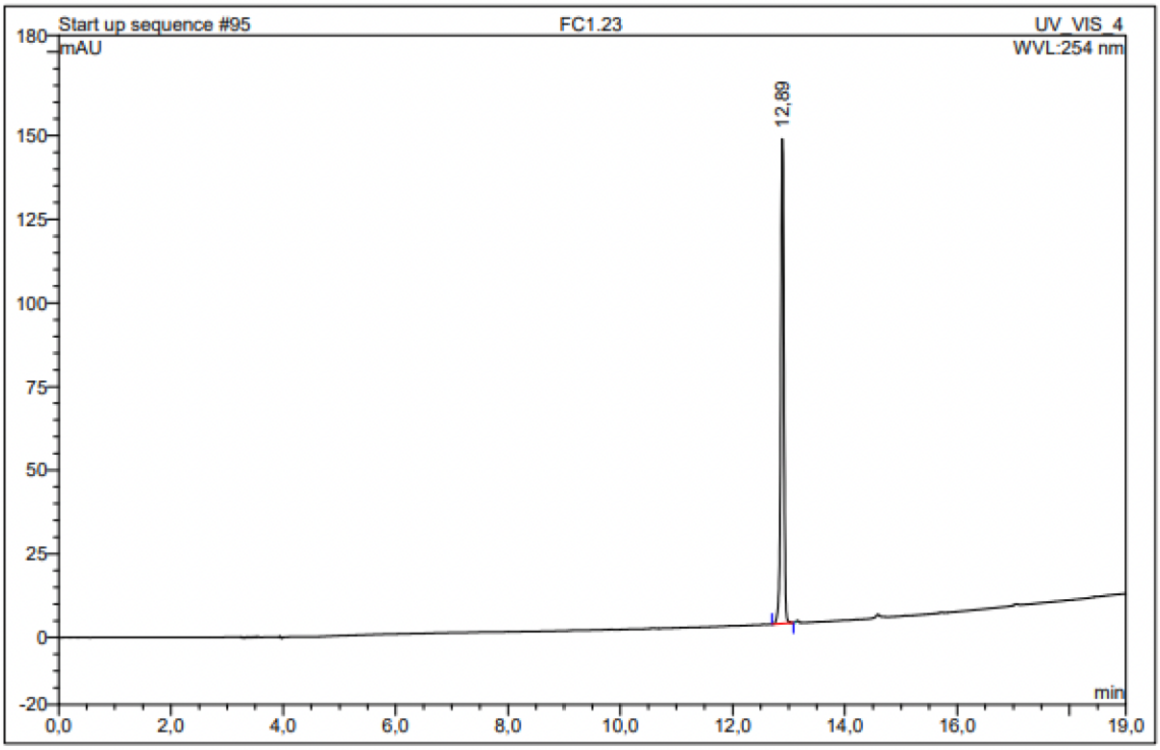

| No. | Ret.Time min | Peak Name | Height mAU | Area mAU*min | Rel.Area % | Amount | Resolution(EP) |
|---|---|---|---|---|---|---|---|
| 1 | 12,89 | n.a. | 144,877 | 8,645 | 100,00 | n.a. | n.a. |
| Total: | | | 144,877 | 8,645 | 100,00 | 0,000 | |

Compound 6 - TUG-2614

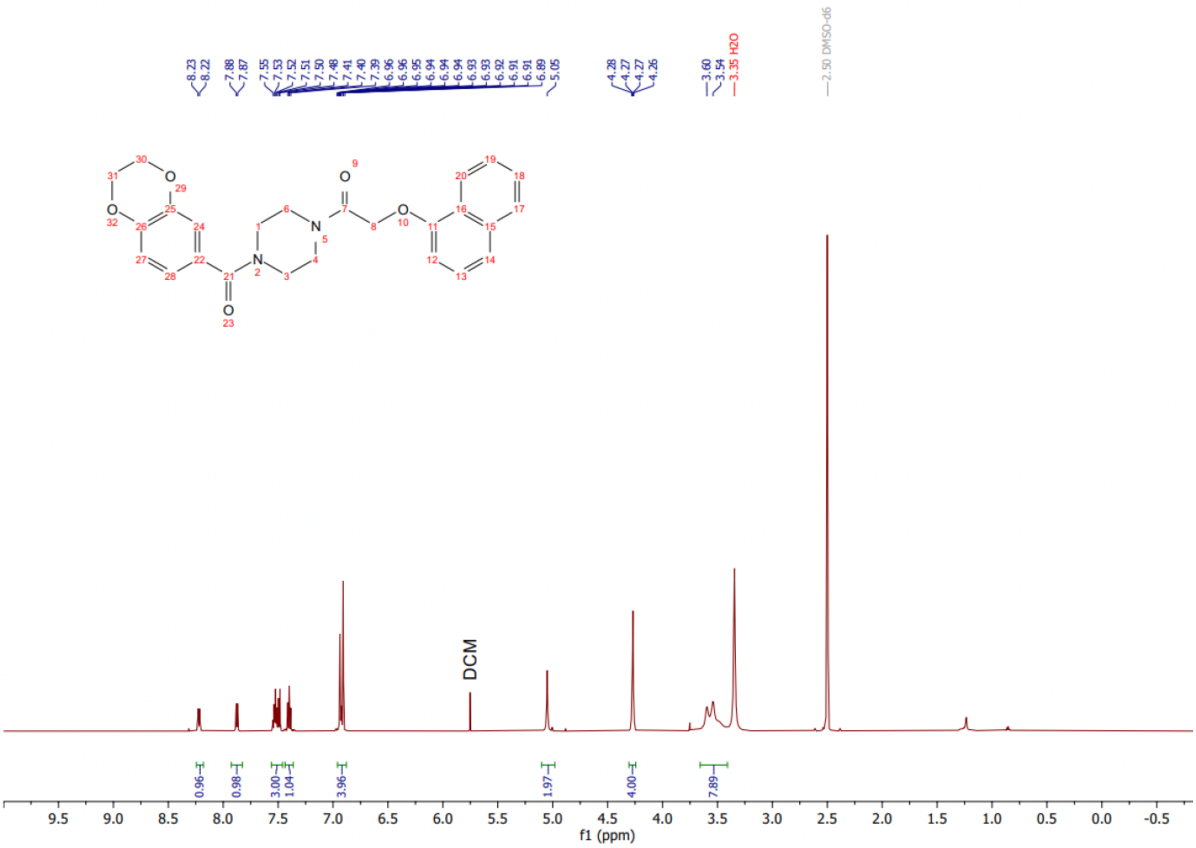

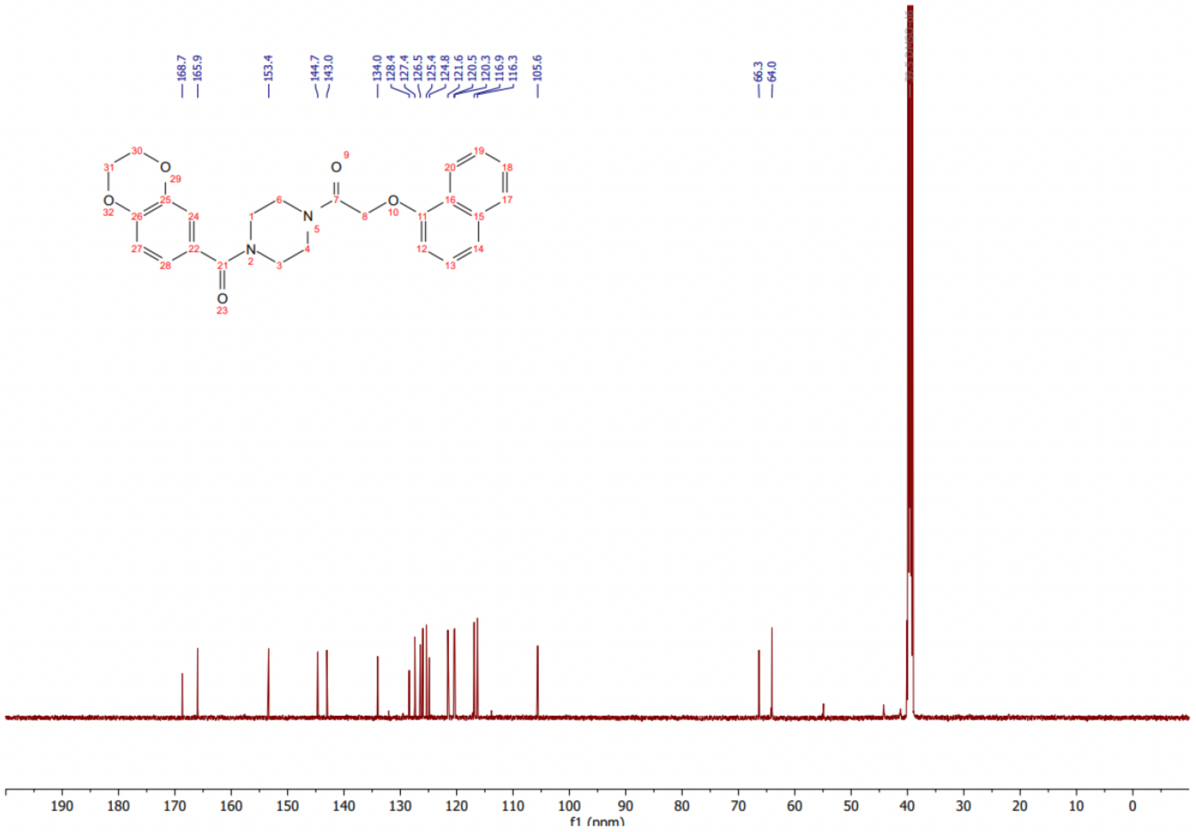

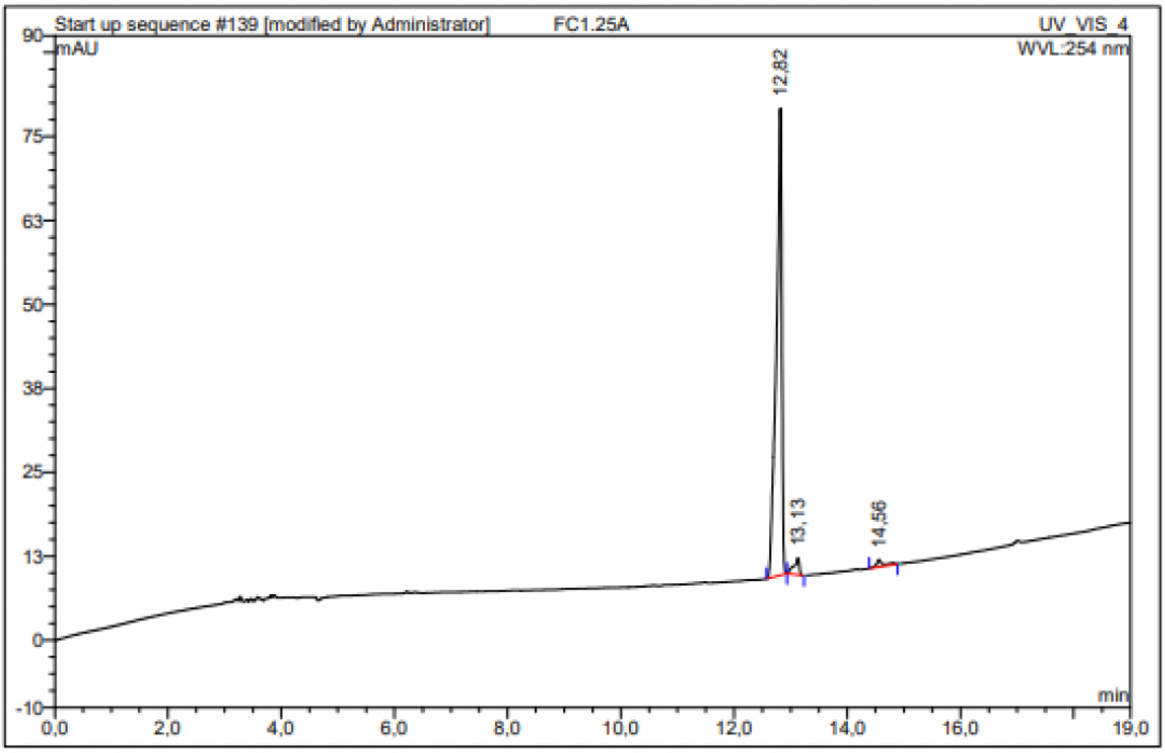

| No. | Ret.Time min | Peak Name | Height mAU | Area mAU*min | Rel.Area % | Amount | Resolution(EP) |
|---|---|---|---|---|---|---|---|
| 1 | 12,82 | n.a. | 69,518 | 7,837 | 95,03 | n.a. | 1,91 |
| 2 | 13,13 | n.a. | 2,513 | 0,269 | 3,26 | n.a. | 10,25 |
| 3 | 14,56 | n.a. | 1,054 | 0,142 | 1,72 | n.a. | n.a. |
| Total: | | | 73,085 | 8,247 | 100,00 | 0,000 | |

Compound 7 - TUG-2612

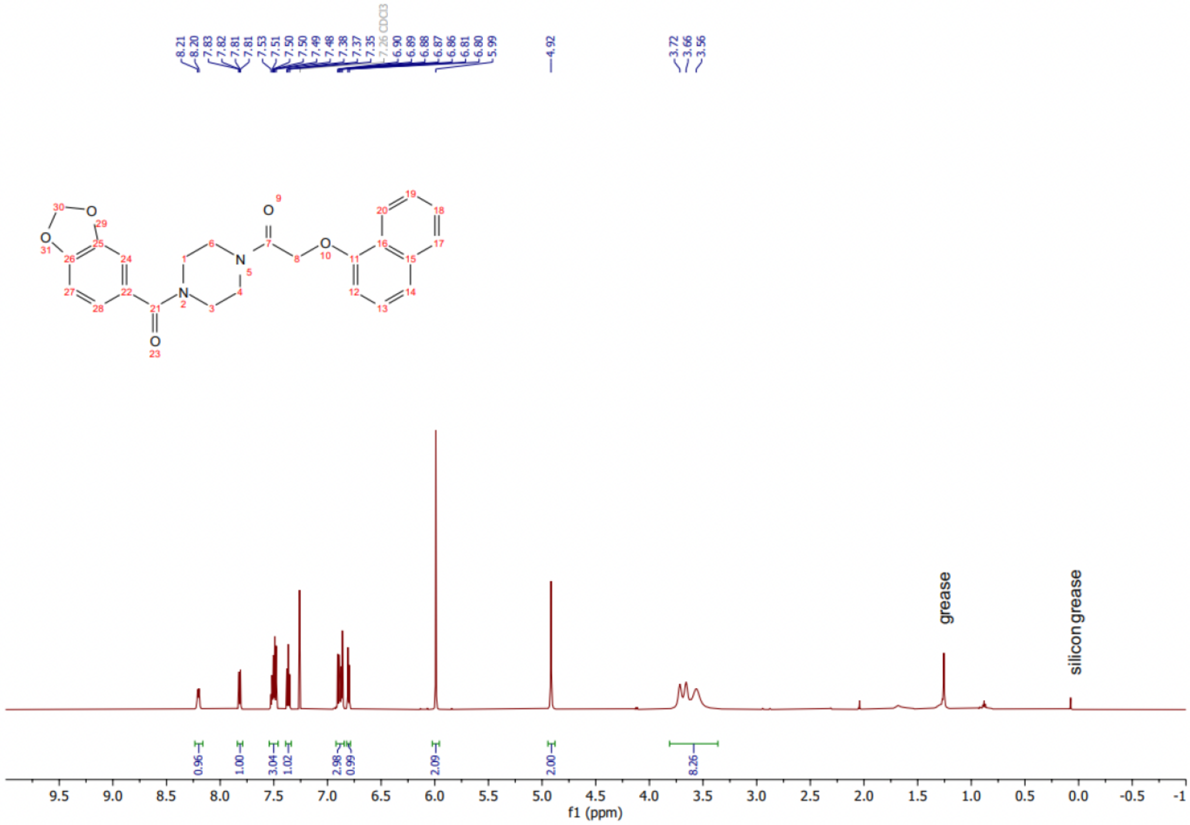

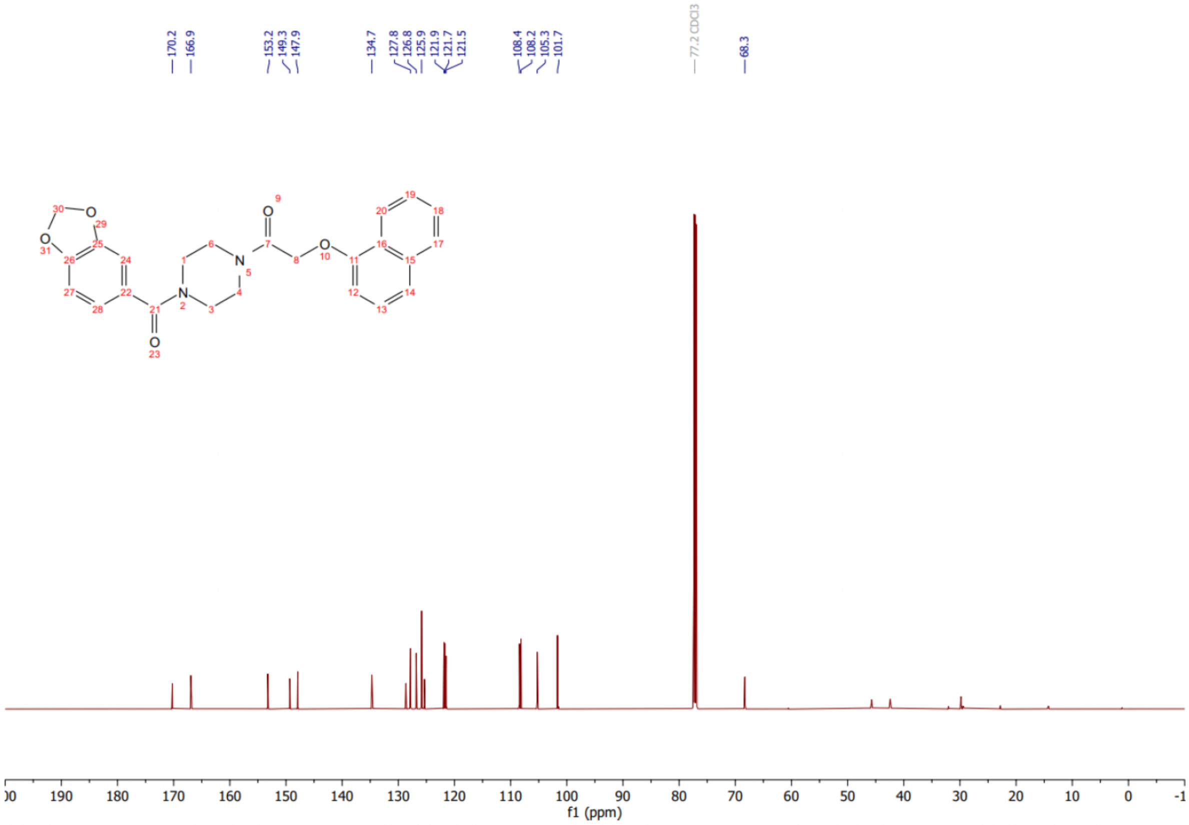

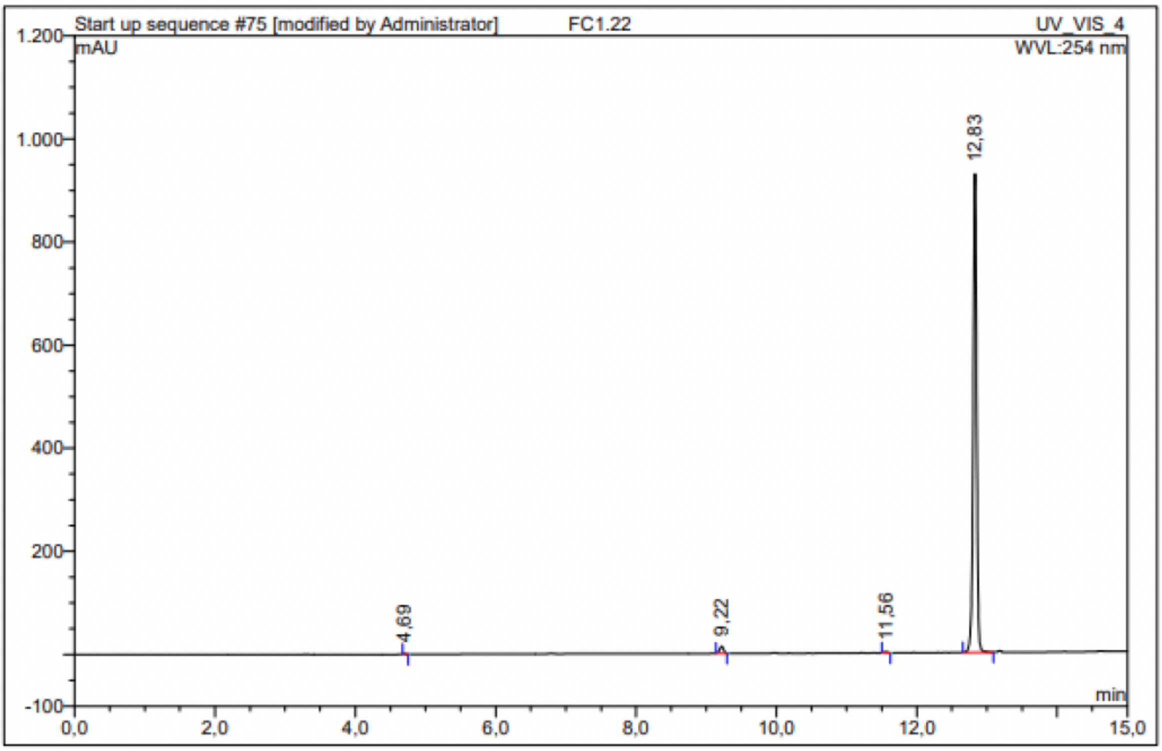

| No. | Ret.Time min | Peak Name | Height mAU | Area mAU*min | Rel.Area % | Amount | Resolution(EP) |
|---|---|---|---|---|---|---|---|
| 1 | 4,69 | n.a. | 2,491 | 0,038 | 0,07 | n.a. | 64,75 |
| 2 | 9,22 | n.a. | 13,417 | 0,870 | 1,57 | n.a. | 24,40 |
| 3 | 11,56 | n.a. | 2,159 | 0,117 | 0,21 | n.a. | 14,39 |
| 4 | 12,83 | n.a. | 928,178 | 54,225 | 98,15 | n.a. | n.a. |
| Total: | | | 946,245 | 55,250 | 100,00 | 0,000 | |

Compound 8 - TUG-2250

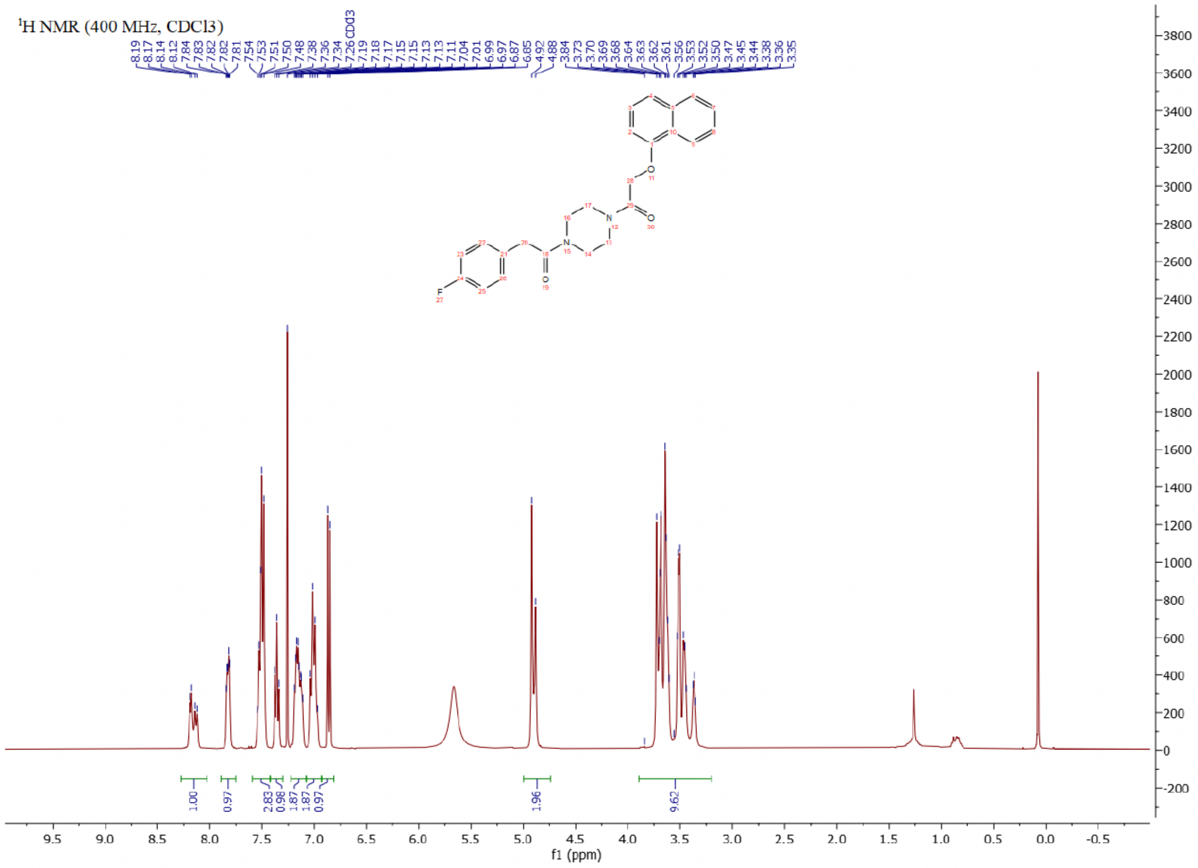

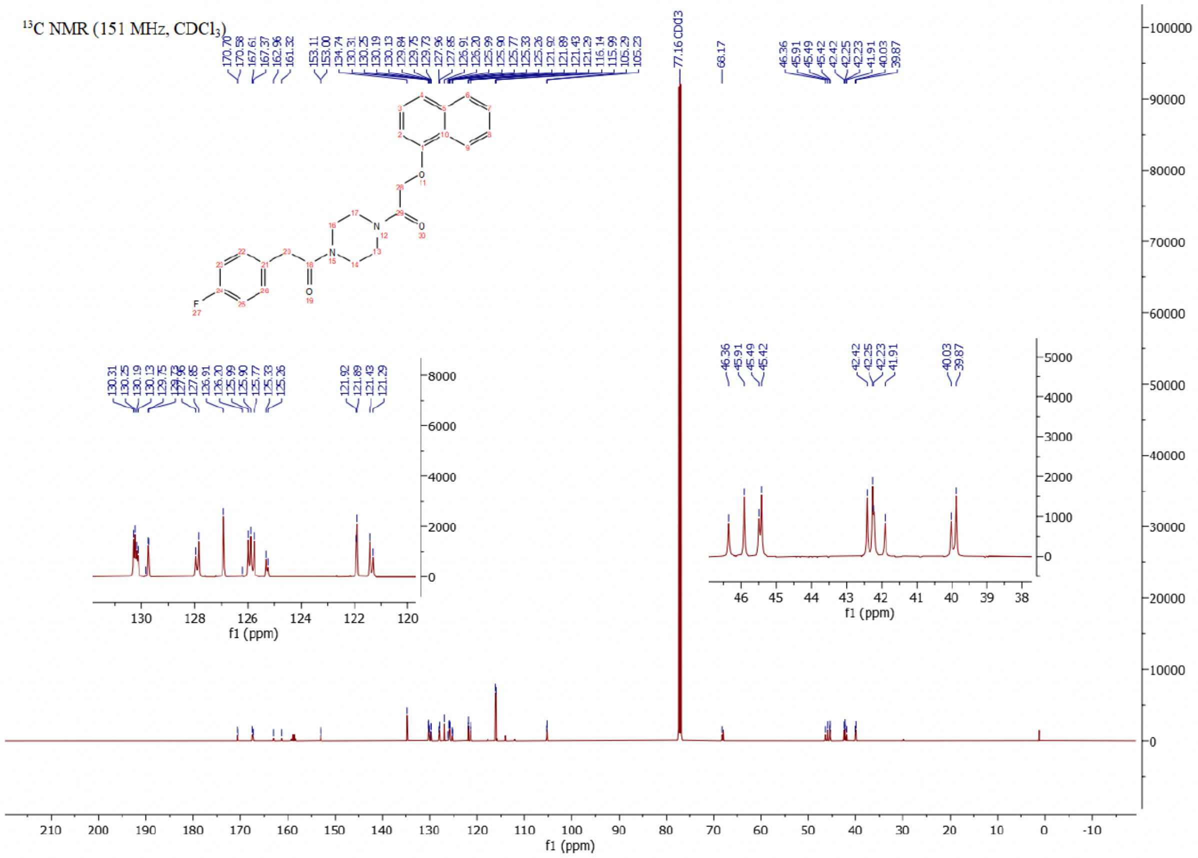

$^{19}$F NMR (376 MHz, $CDCl_3$)

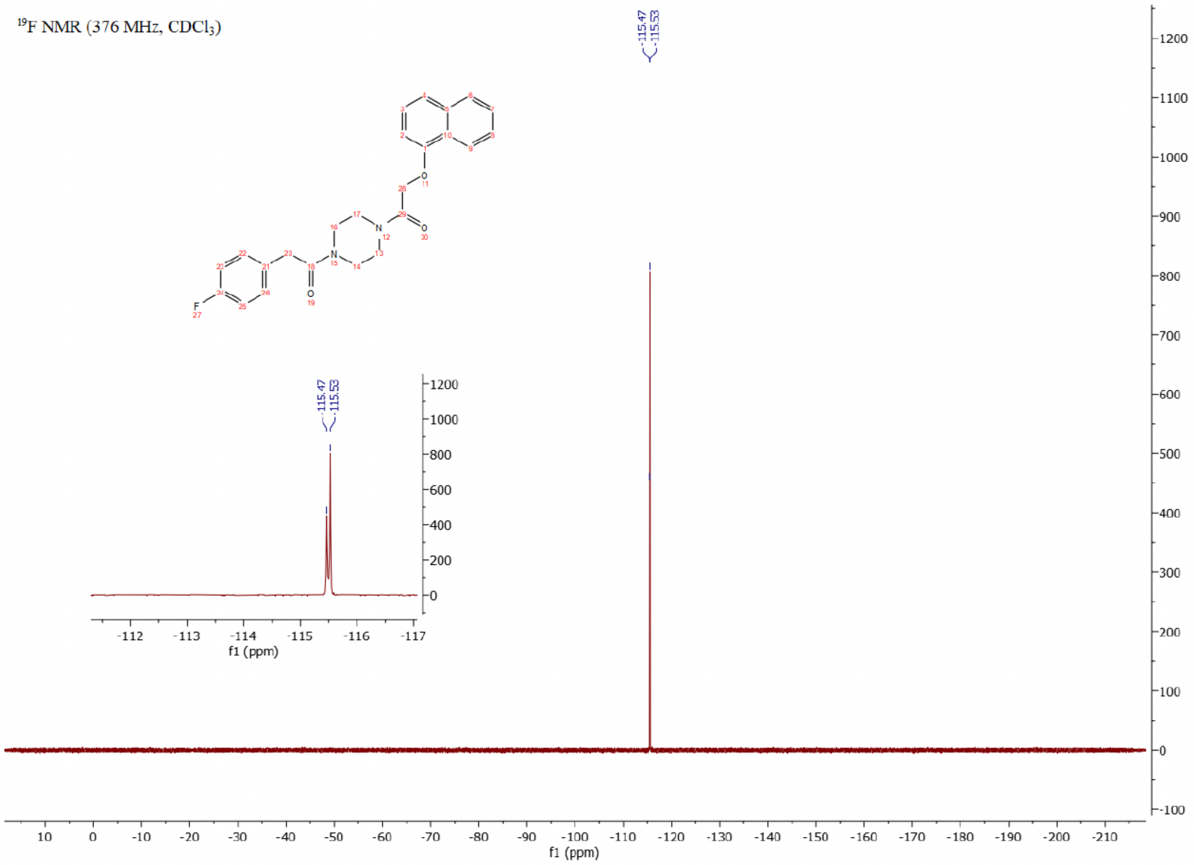

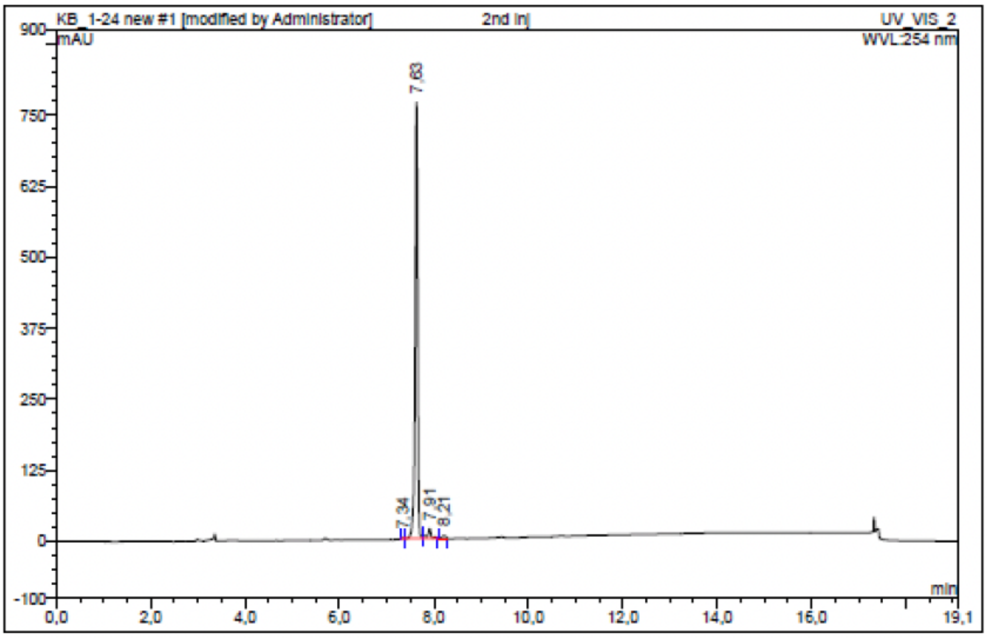

| No. | Ret.Time min | Peak Name | Height mAU | Area mAU*min | Rel.Area % | Amount | Resolution(EP) |
|---|---|---|---|---|---|---|---|
| 1 | 7,34 | n.a. | 1,041 | 0,042 | 0,08 | n.a. | 3,61 |
| 2 | 7,63 | n.a. | 766,996 | 48,046 | 96,11 | n.a. | 3,00 |
| 3 | 7,91 | n.a. | 16,953 | 1,488 | 2,98 | n.a. | 2,62 |
| 4 | 8,21 | n.a. | 4,981 | 0,415 | 0,83 | n.a. | n.a. |
| Total: | | | 789,971 | 49,991 | 100,00 | 0,000 | |

Compound 9 - TUG-2361

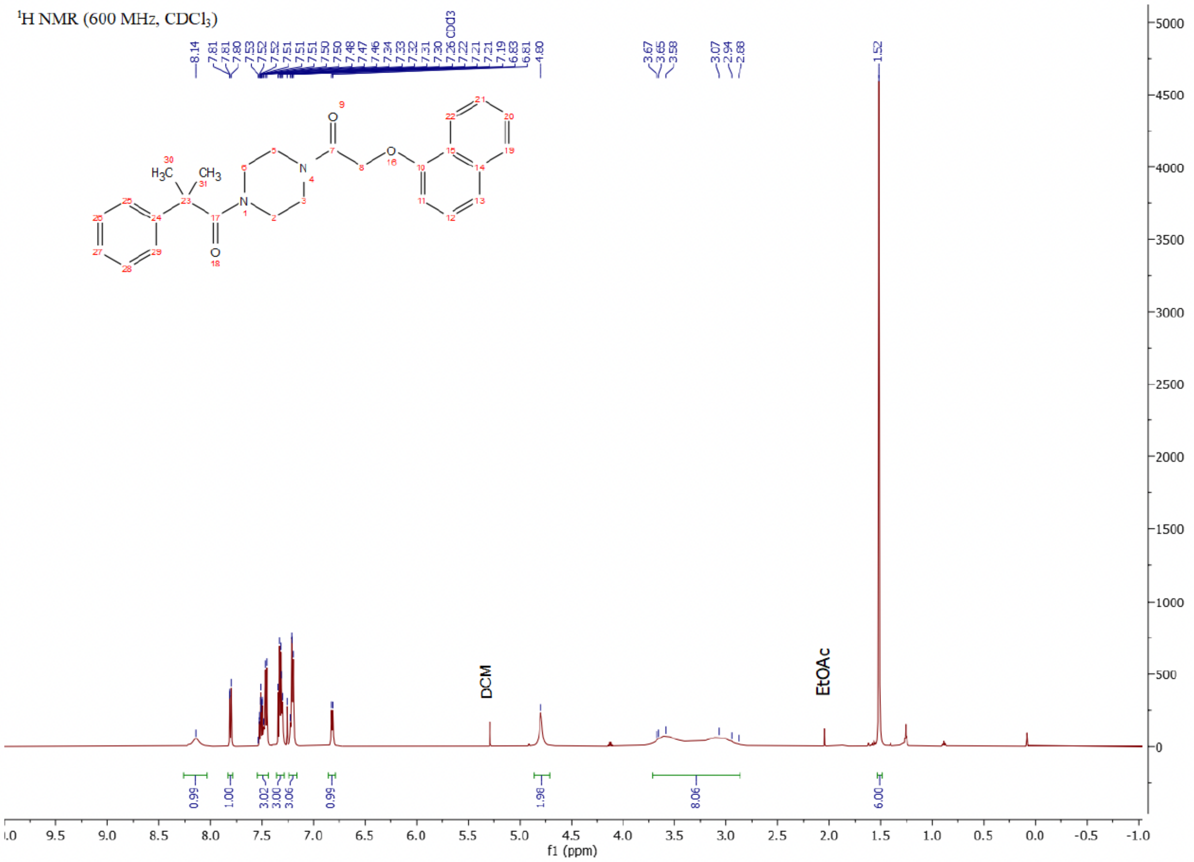

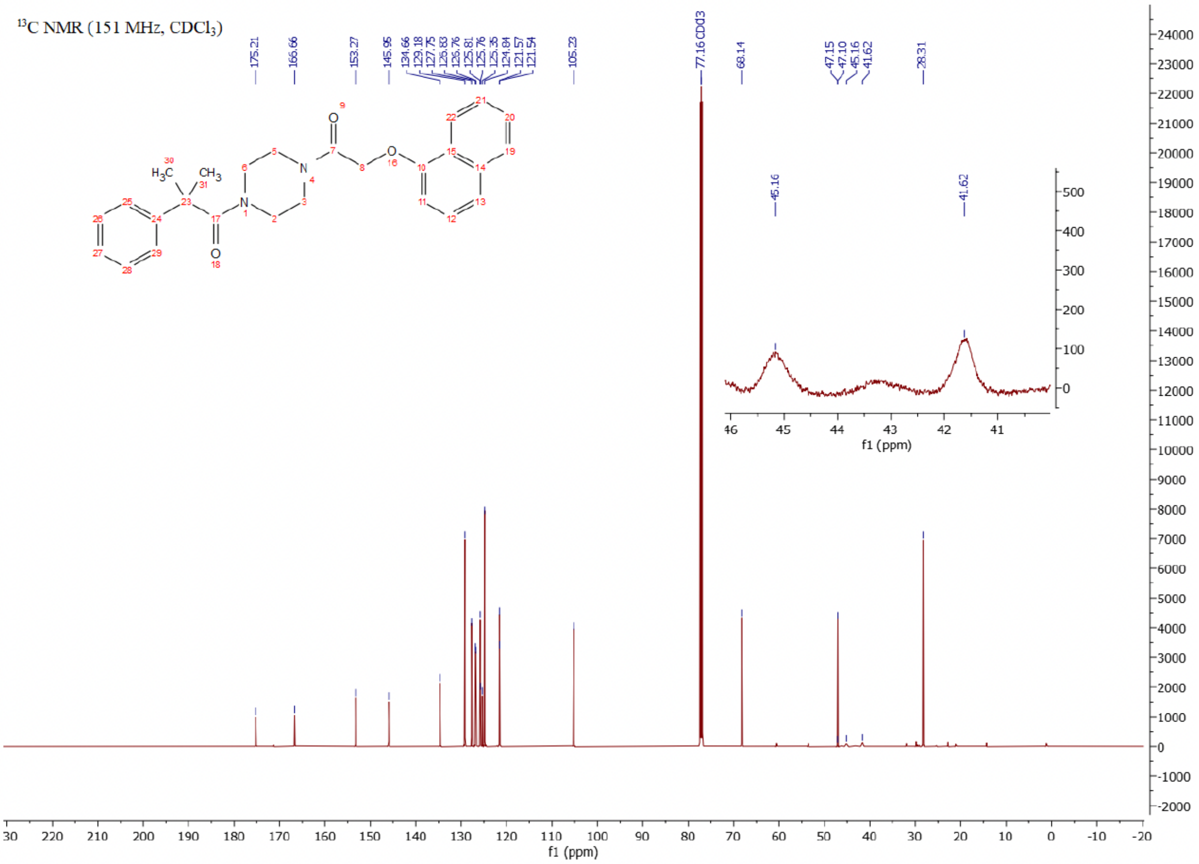

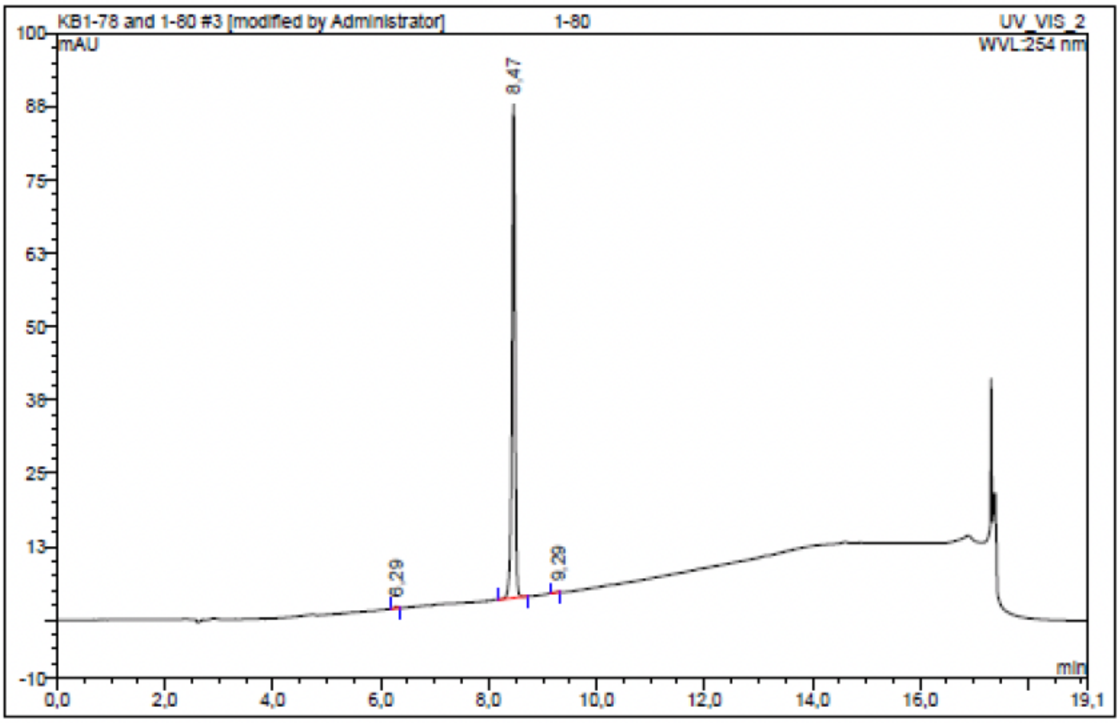

| No. | Ret.Time min | Peak Name | Height mAU | Area mAU*min | Rel.Area % | Amount | Resolution(EP) |
|---|---|---|---|---|---|---|---|
| 1 | 6,29 | n.a. | 0,289 | 0,016 | 0,29 | n.a. | 23,39 |
| 2 | 8,47 | n.a. | 84,187 | 5,600 | 99,68 | n.a. | 9,12 |
| 3 | 9,29 | n.a. | 0,075 | 0,002 | 0,03 | n.a. | n.a. |
| Total: | | | 84,551 | 5,618 | 100,00 | 0,000 | |

Compound 10 - TUG-2359

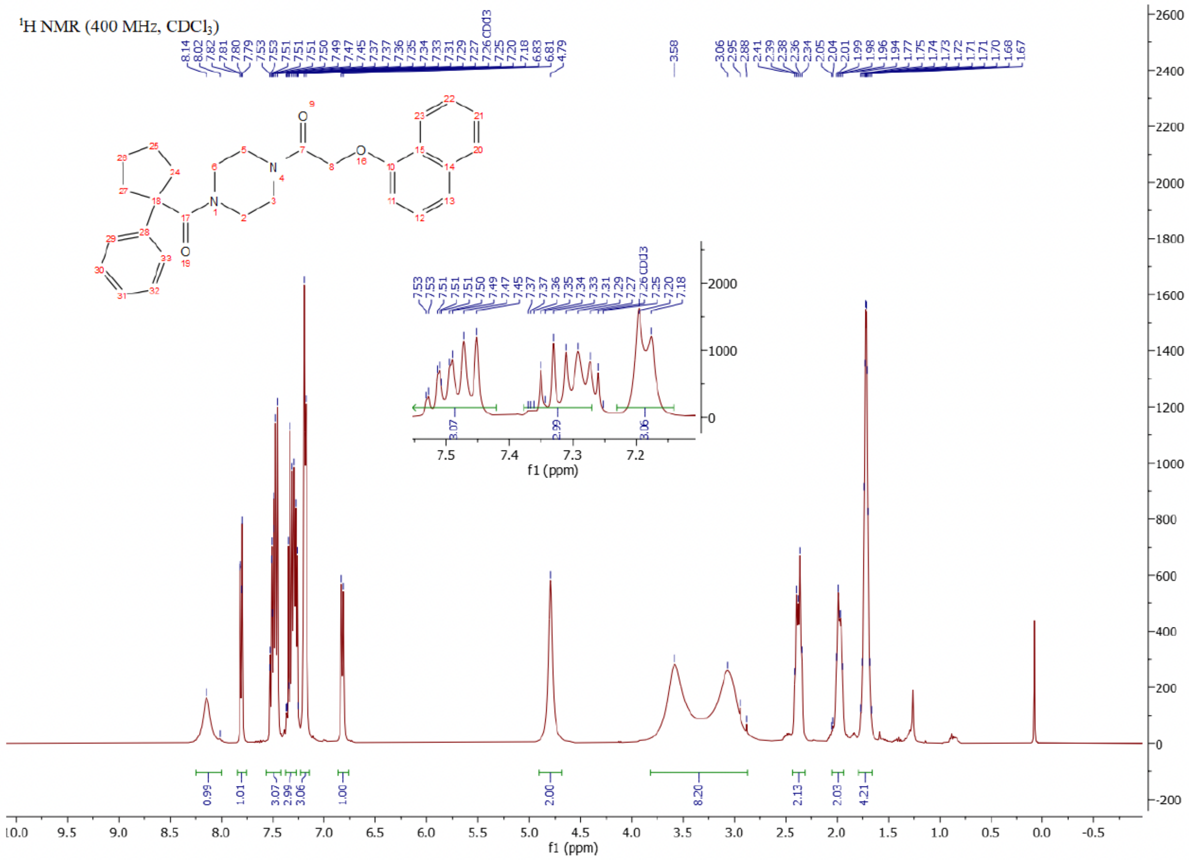

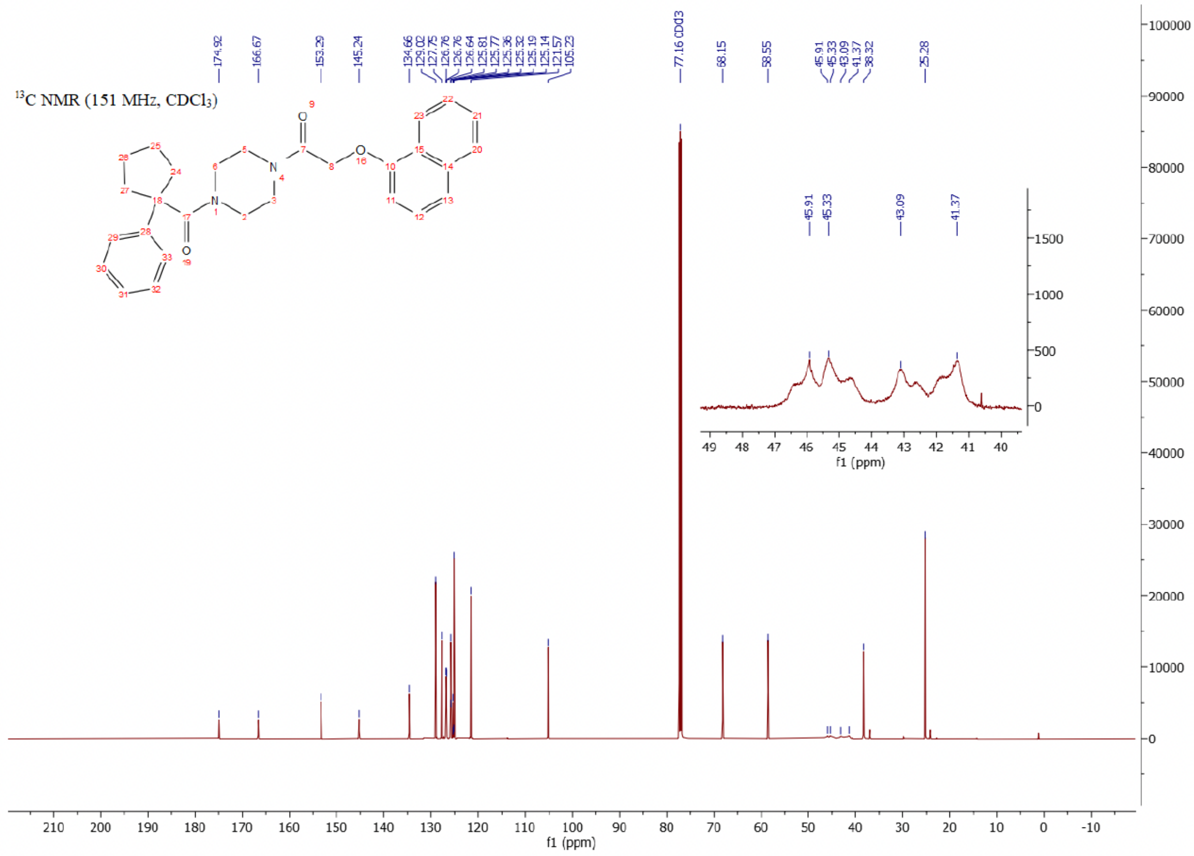

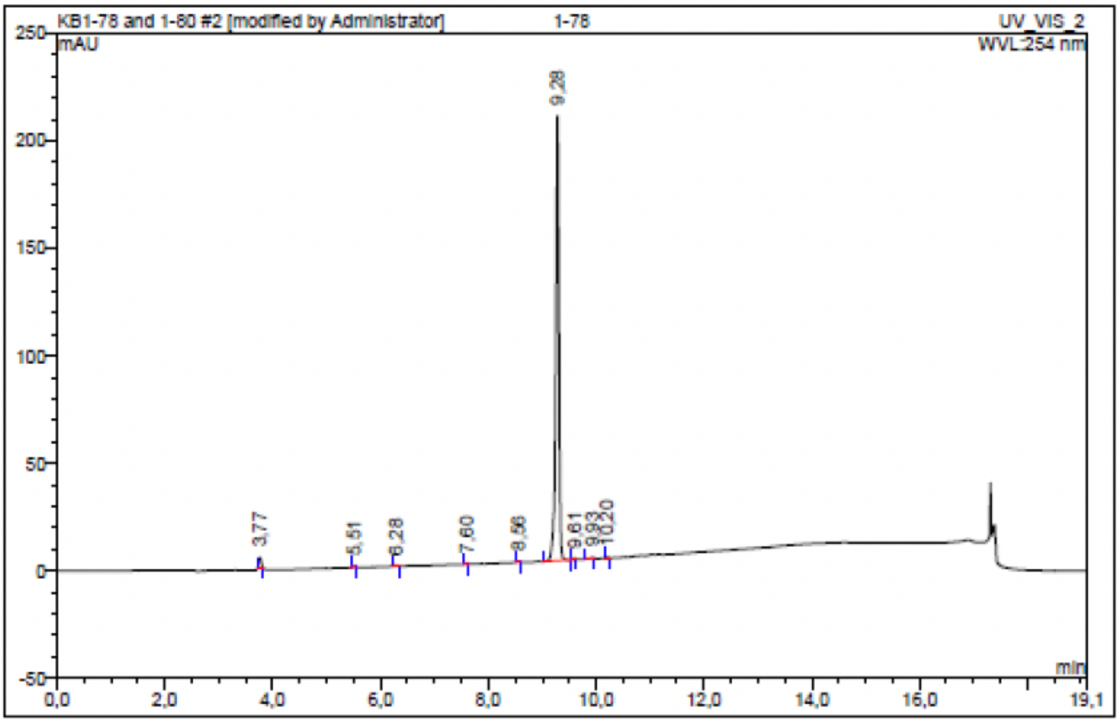

| No. | Ret.Time min | Peak Name | Height mAU | Area mAU*min | Rel.Area % | Amount | Resolution(EP) |
|---|---|---|---|---|---|---|---|
| 1 | 3,77 | n.a. | 4,880 | 0,278 | 1,89 | n.a. | 20,88 |
| 2 | 5,51 | n.a. | 0,424 | 0,018 | 0,12 | n.a. | 9,78 |
| 3 | 6,28 | n.a. | 0,543 | 0,030 | 0,20 | n.a. | 16,35 |
| 4 | 7,60 | n.a. | 0,495 | 0,022 | 0,15 | n.a. | 14,02 |
| 5 | 8,56 | n.a. | 0,348 | 0,013 | 0,09 | n.a. | 8,81 |
| 6 | 9,28 | n.a. | 206,814 | 14,324 | 97,20 | n.a. | 3,80 |
| 7 | 9,61 | n.a. | 0,375 | 0,015 | 0,10 | n.a. | 4,29 |
| 8 | 9,93 | n.a. | 0,353 | 0,019 | 0,13 | n.a. | 3,48 |
| 9 | 10,20 | n.a. | 0,401 | 0,018 | 0,12 | n.a. | n.a. |
| Total: | | | 214,633 | 14,737 | 100,00 | 0,000 | |

Compound 11 - TUG-2373

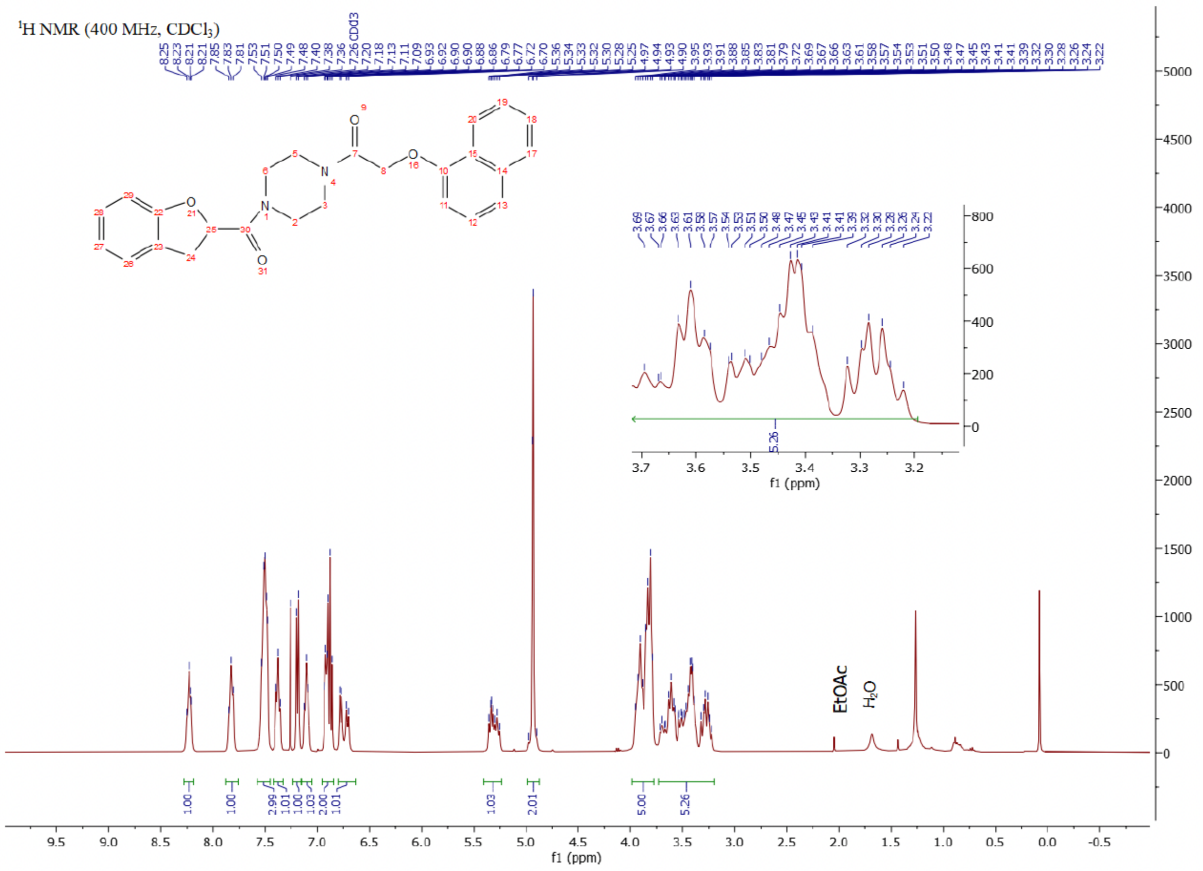

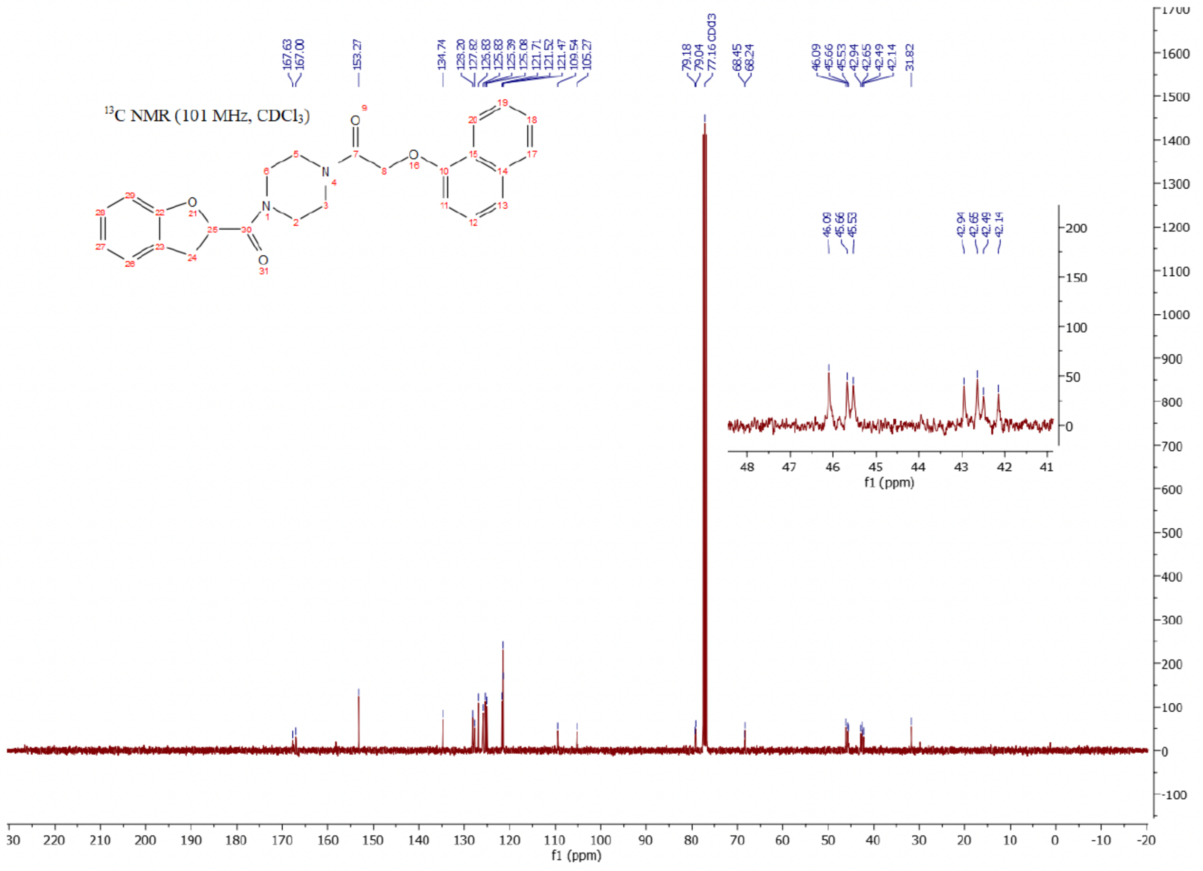

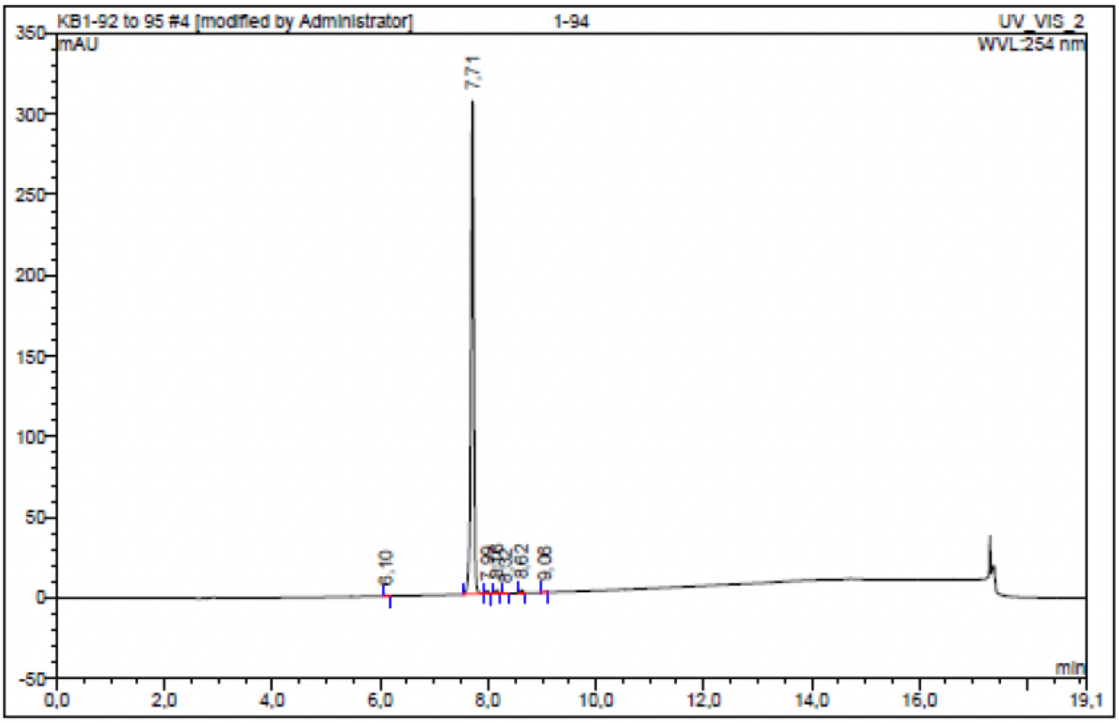

| No. | Ret.Time min | Peak Name | Height mAU | Area mAU*min | Rel.Area % | Amount | Resolution(EP) |
|---|---|---|---|---|---|---|---|
| 1 | 6,10 | n.a. | 0,532 | 0,030 | 0,15 | n.a. | 17,32 |
| 2 | 7,71 | n.a. | 305,527 | 19,360 | 98,13 | n.a. | 3,05 |
| 3 | 7,99 | n.a. | 1,581 | 0,088 | 0,44 | n.a. | 1,89 |
| 4 | 8,16 | n.a. | 2,184 | 0,124 | 0,63 | n.a. | 1,81 |
| 5 | 8,32 | n.a. | 0,214 | 0,011 | 0,06 | n.a. | 3,43 |
| 6 | 8,62 | n.a. | 1,674 | 0,098 | 0,49 | n.a. | 4,76 |
| 7 | 9,06 | n.a. | 0,320 | 0,018 | 0,09 | n.a. | n.a. |
| Total: | | | 312,032 | 19,728 | 100,00 | 0,000 | |

Compound 12 - TUG-2364

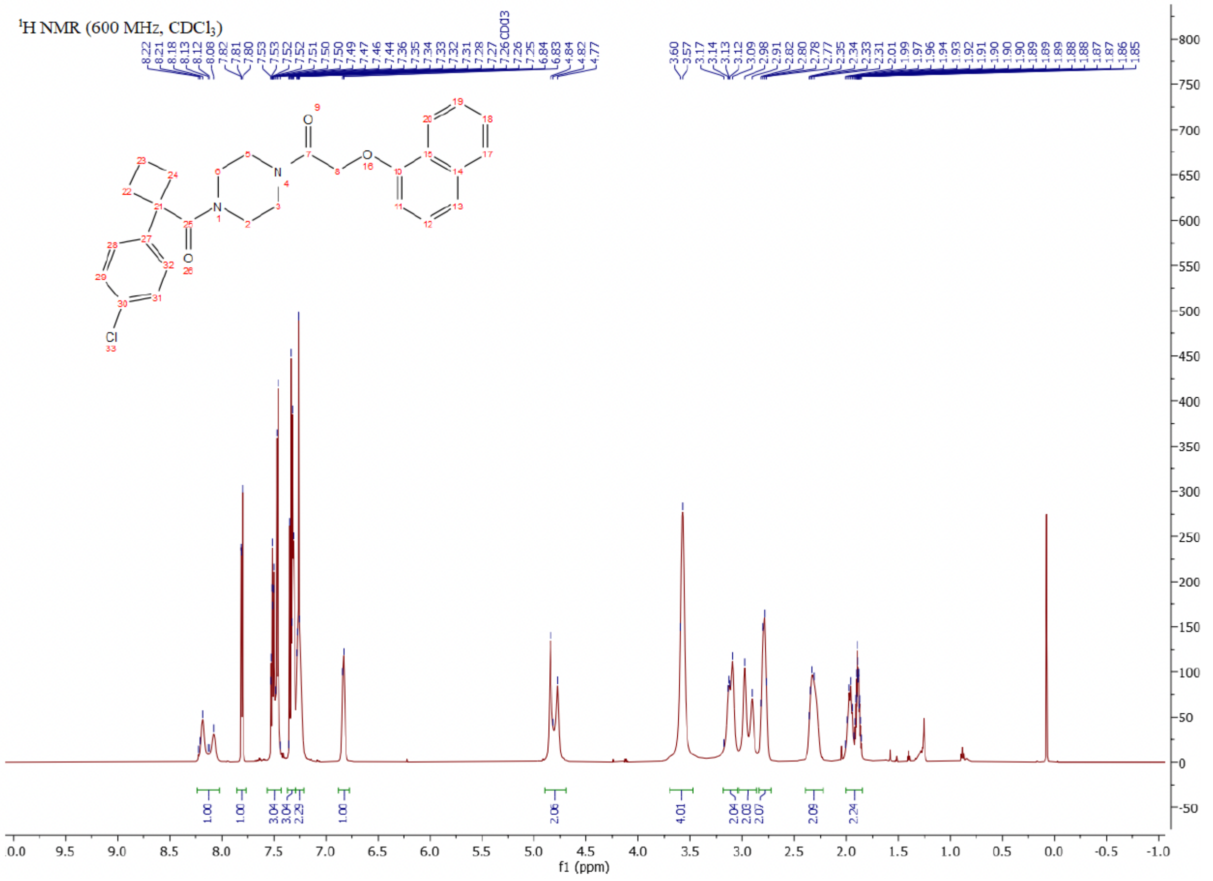

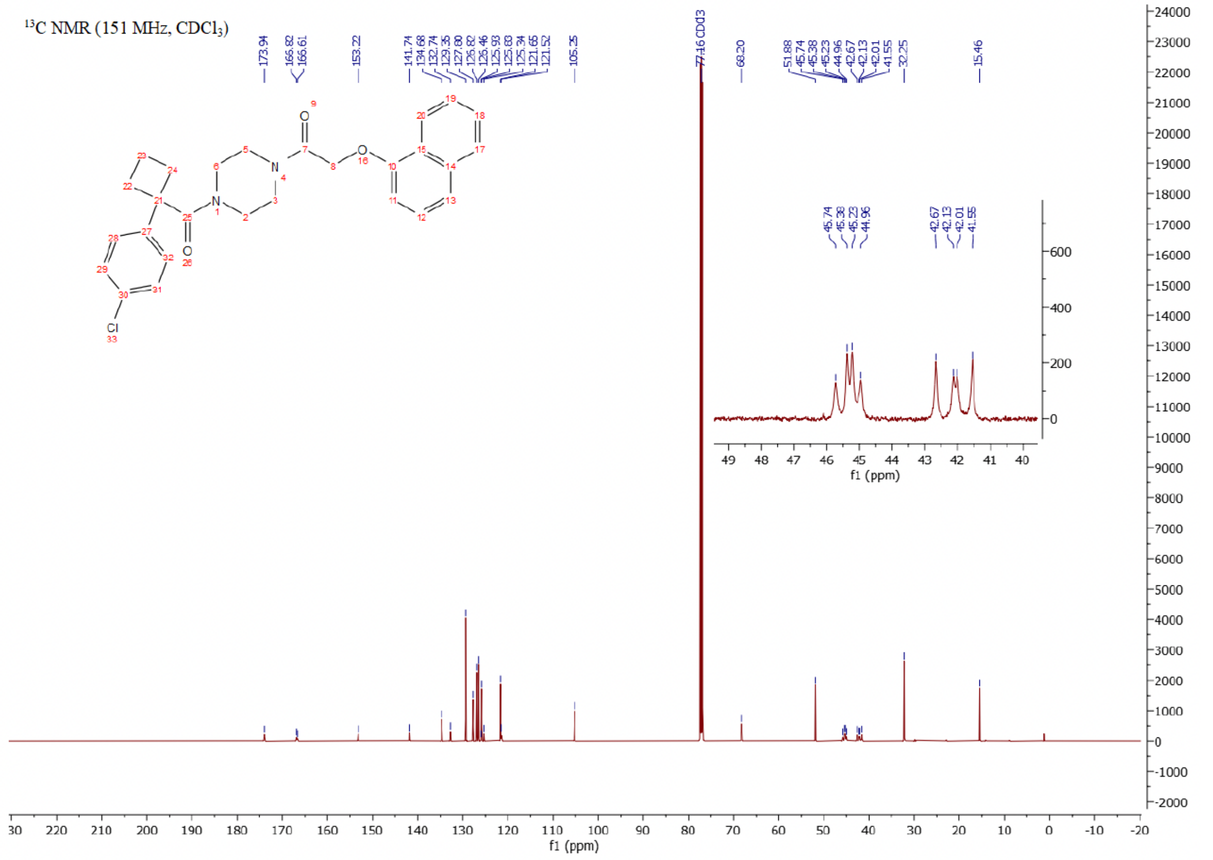

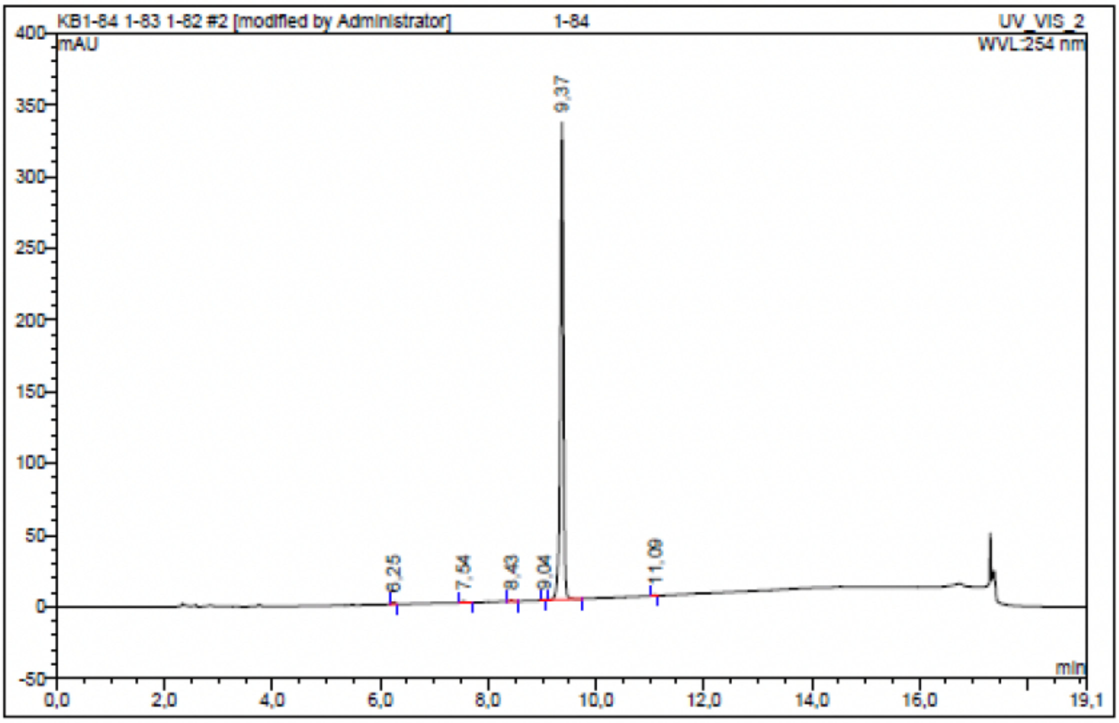

| No. | Ret.Time min | Peak Name | Height mAU | Area mAU*min | Rel.Area % | Amount | Resolution(EP) |
|---|---|---|---|---|---|---|---|
| 1 | 6,25 | n.a. | 1,388 | 0,078 | 0,34 | n.a. | 14,24 |
| 2 | 7,54 | n.a. | 1,038 | 0,058 | 0,25 | n.a. | 9,61 |
| 3 | 8,43 | n.a. | 0,664 | 0,039 | 0,17 | n.a. | 7,01 |
| 4 | 9,04 | n.a. | 0,223 | 0,011 | 0,05 | n.a. | 3,66 |
| 5 | 9,37 | n.a. | 332,766 | 22,869 | 99,07 | n.a. | 16,96 |
| 6 | 11,09 | n.a. | 0,457 | 0,028 | 0,12 | n.a. | n.a. |
| Total: | | | 336,535 | 23,083 | 100,00 | 0,000 | |

### Compound 13 - TUG-2371

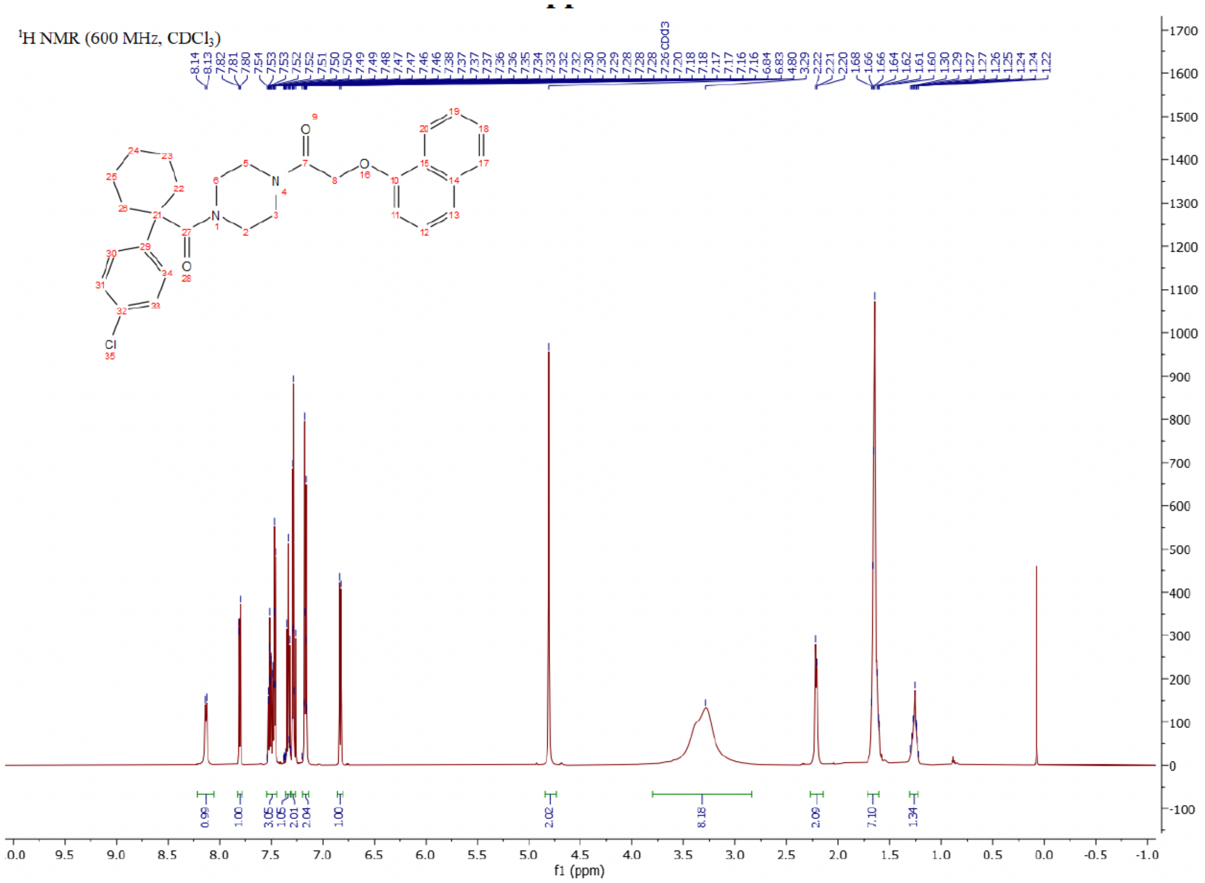

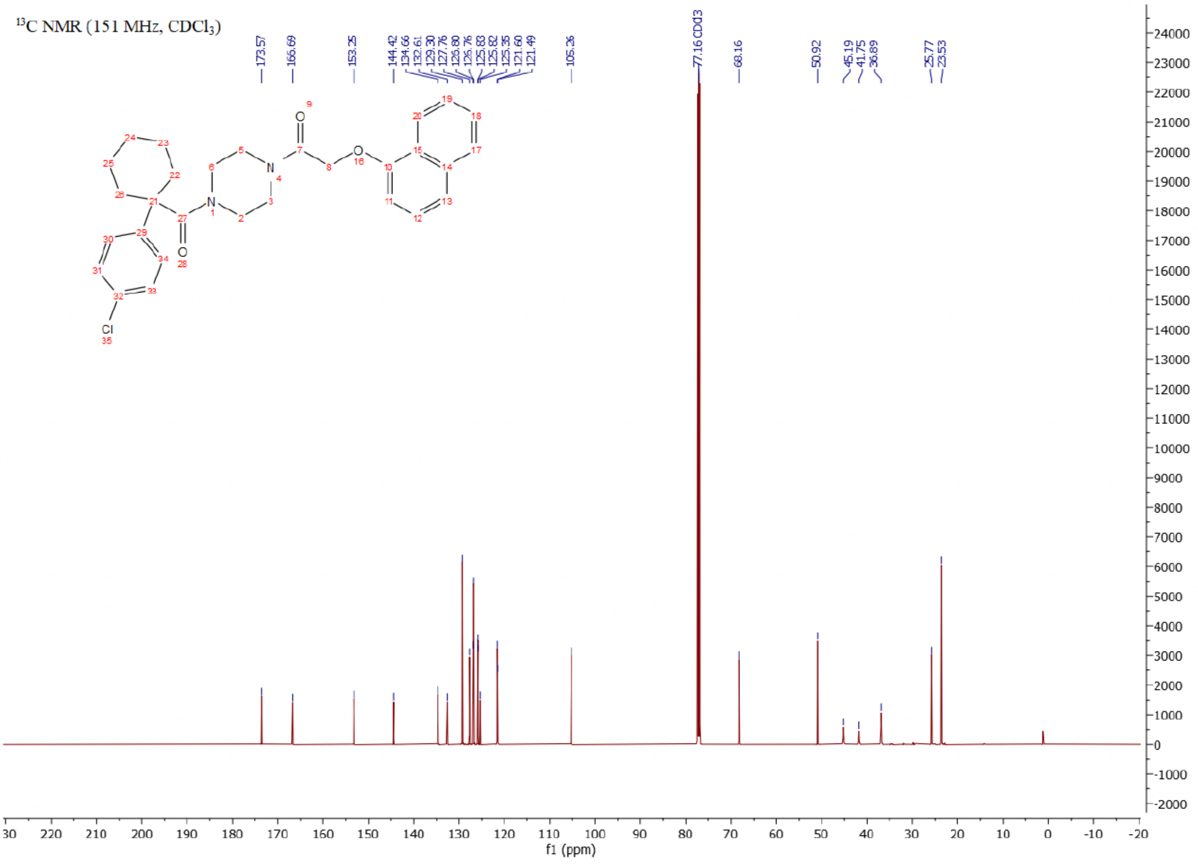

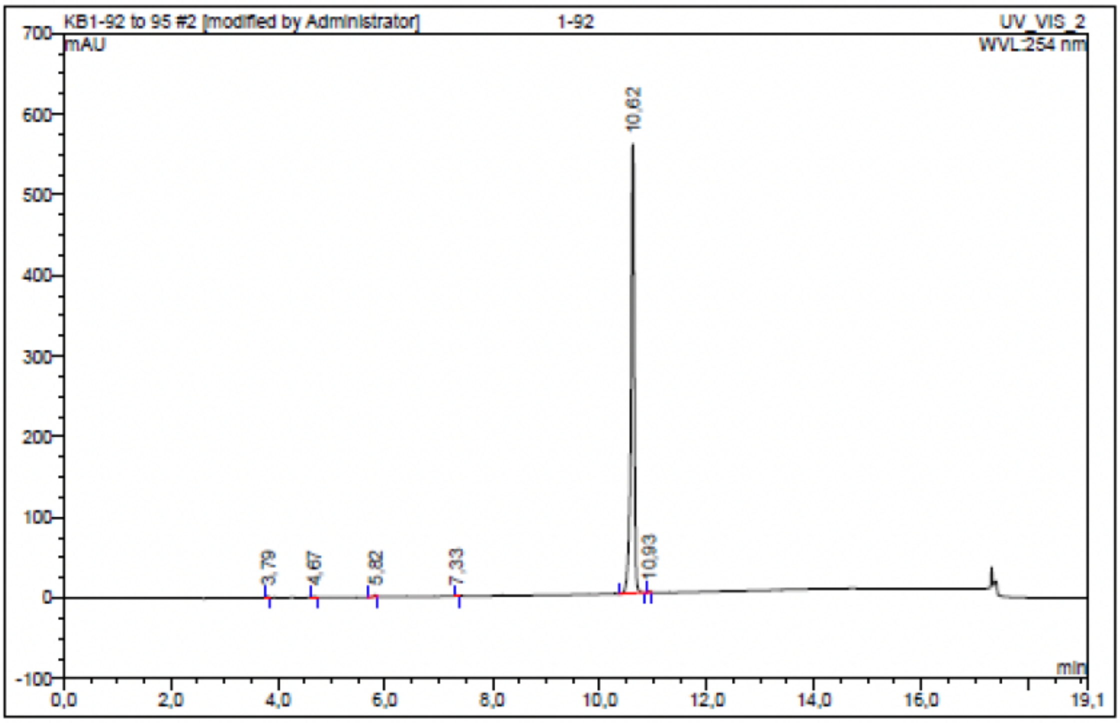

| No. | Ret.Time min | Peak Name | Height mAU | Area mAU*min | Rel.Area % | Amount | Resolution(EP) |
|---|---|---|---|---|---|---|---|
| 1 | 3,79 | n.a. | 1,452 | 0,071 | 0,17 | n.a. | 9,41 |
| 2 | 4,67 | n.a. | 1,149 | 0,072 | 0,17 | n.a. | 12,52 |
| 3 | 5,82 | n.a. | 1,568 | 0,064 | 0,15 | n.a. | 18,79 |
| 4 | 7,33 | n.a. | 1,332 | 0,064 | 0,15 | n.a. | 35,24 |
| 5 | 10,62 | n.a. | 556,939 | 41,261 | 99,17 | n.a. | 3,22 |
| 6 | 10,93 | n.a. | 1,432 | 0,075 | 0,18 | n.a. | n.a. |
| Total: | | | 563,872 | 41,606 | 100,00 | 0,000 | |

Compound 14 - TUG-2360

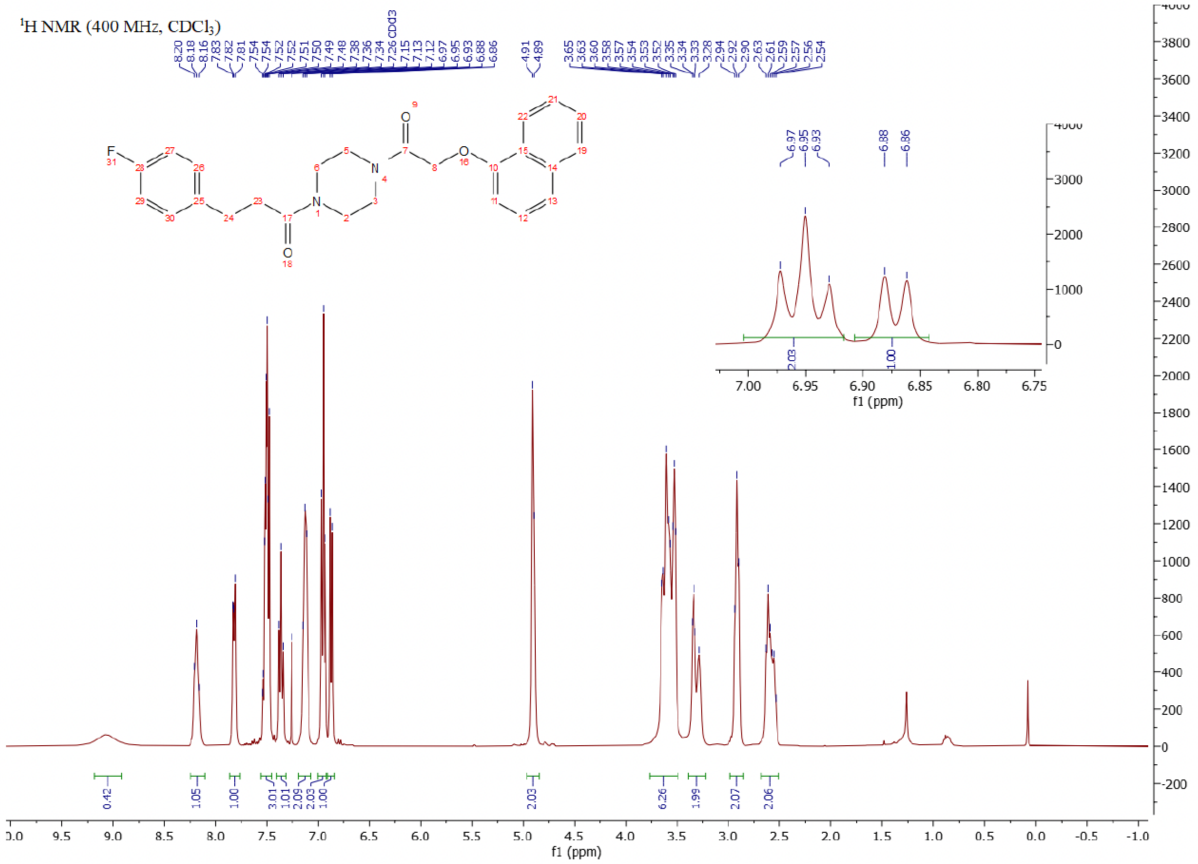

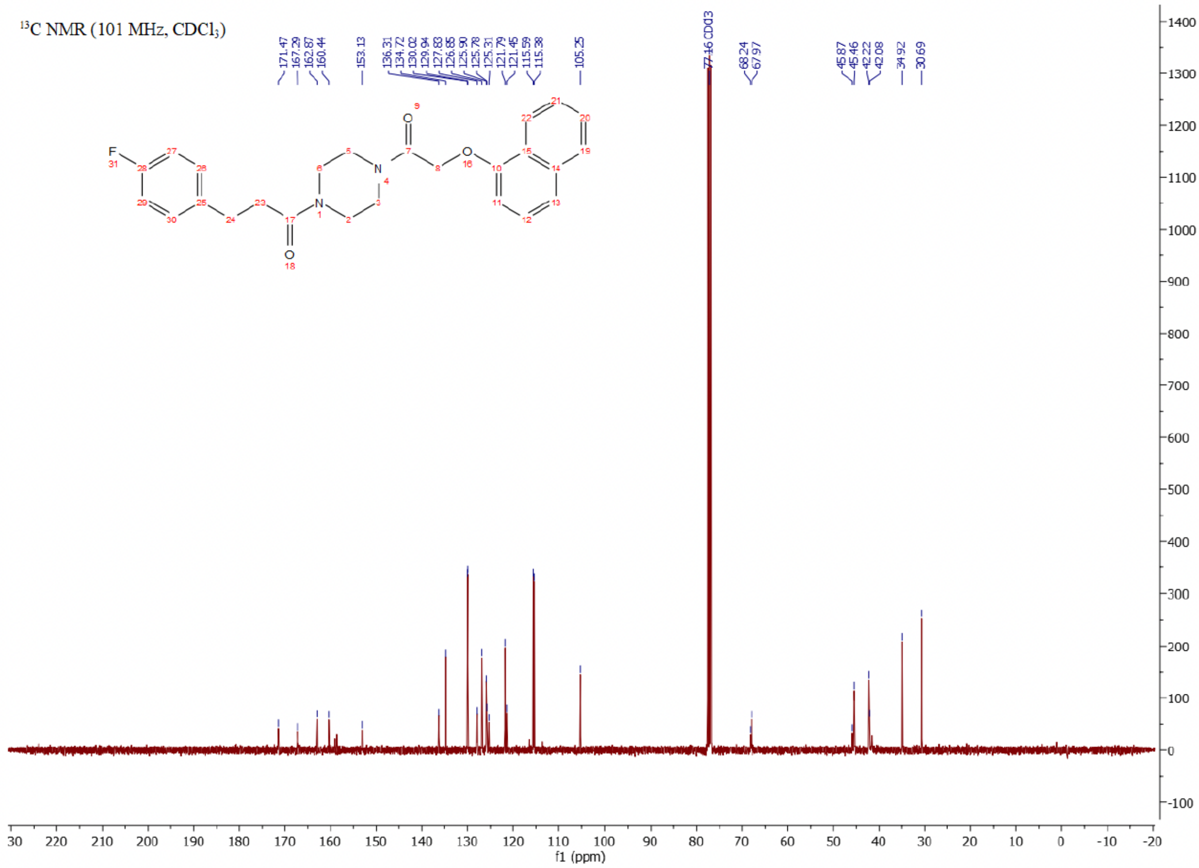

$^{19}F$ NMR (376 MHz, $CDCl_3$)

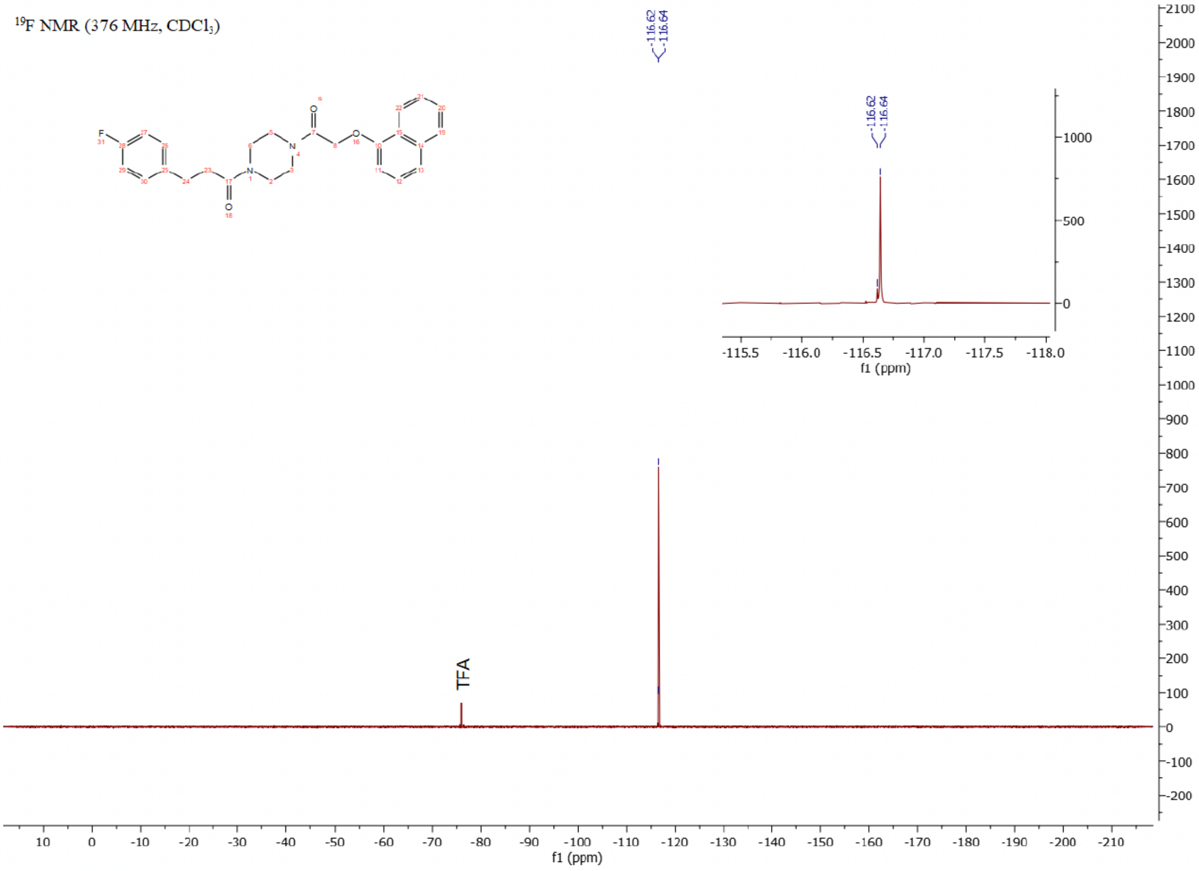

Compound 15 - TUG-2362

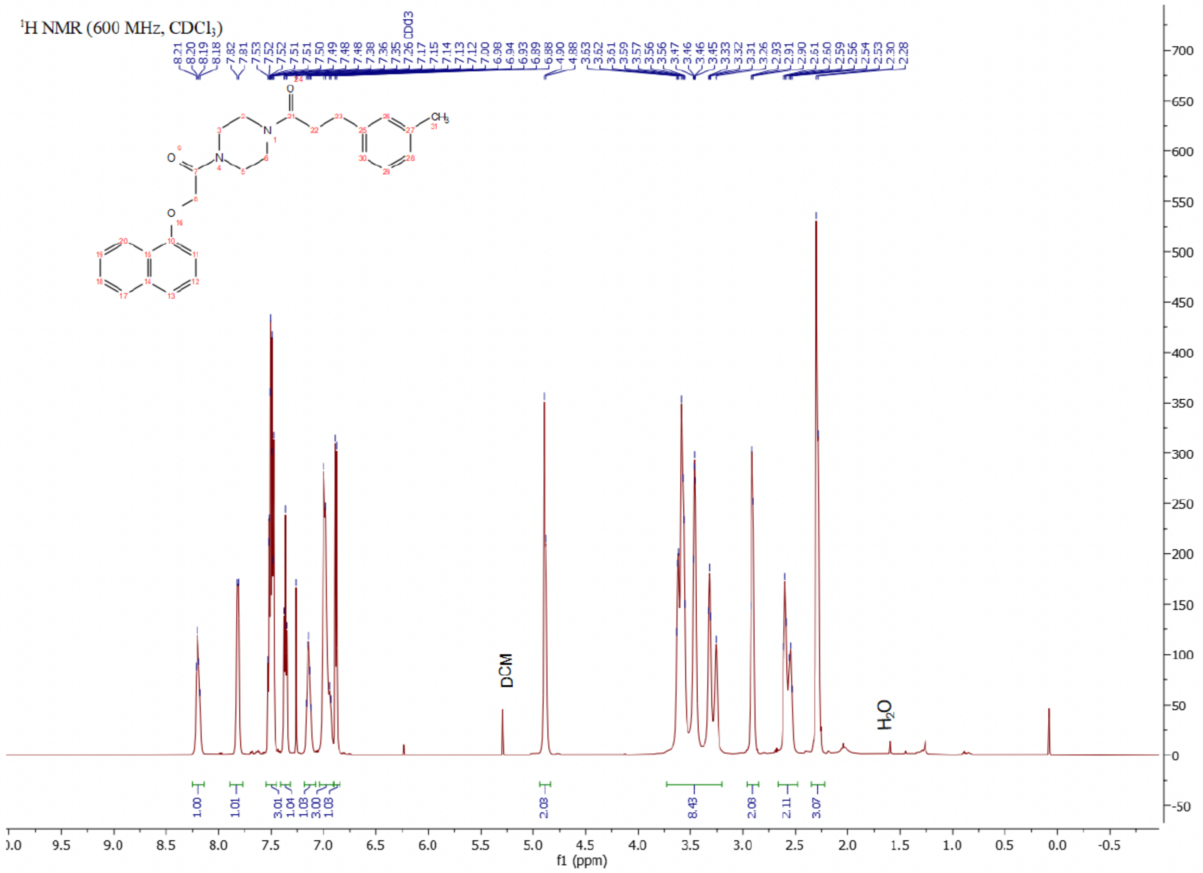

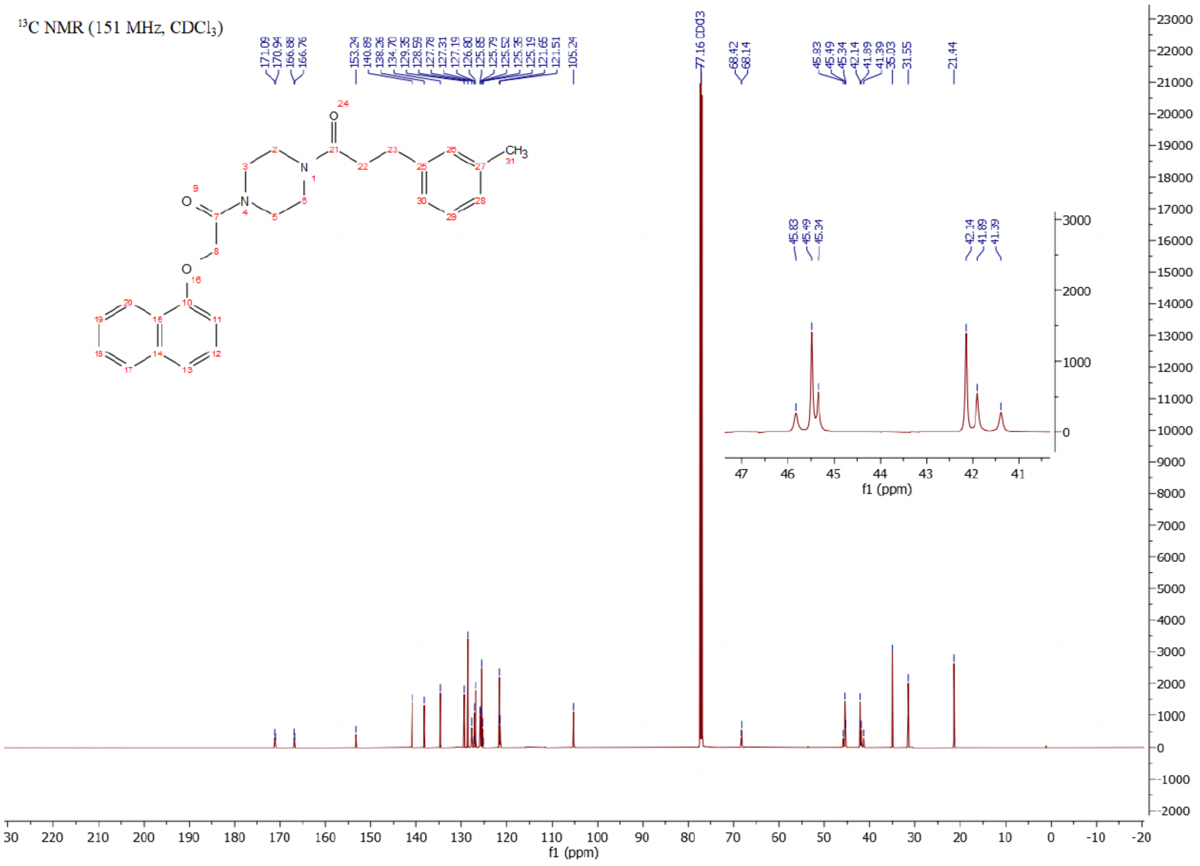

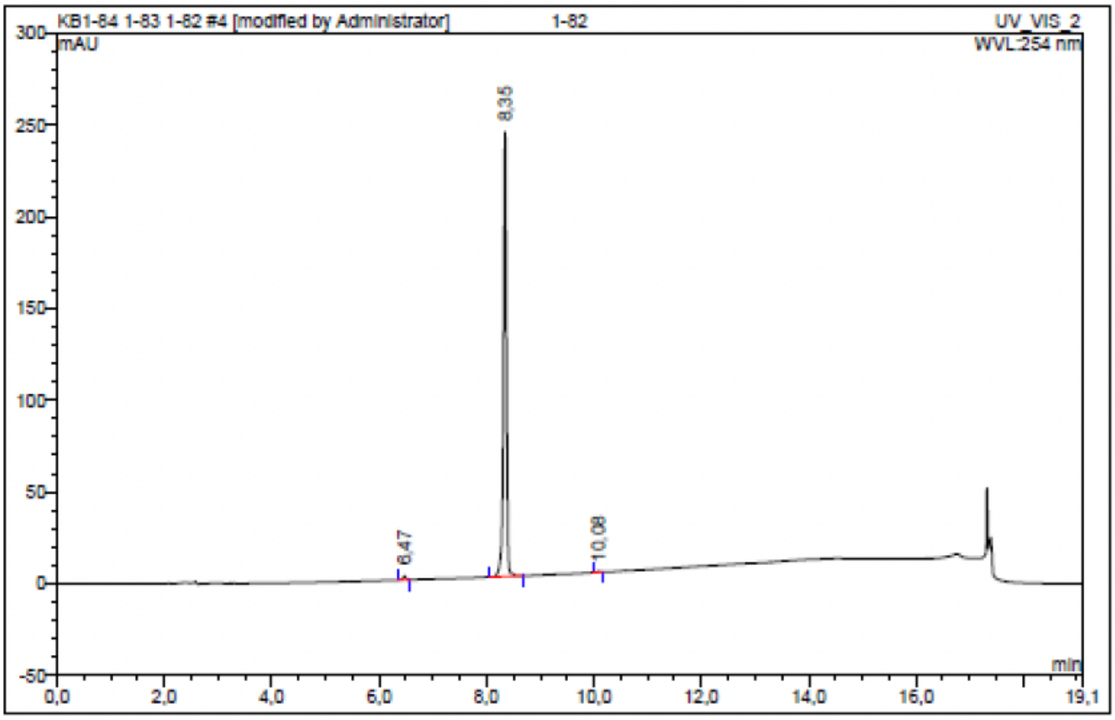

| No. | Ret.Time min | Peak Name | Height mAU | Area mAU*min | Rel.Area % | Amount | Resolution(EP) |
|---|---|---|---|---|---|---|---|
| 1 | 6,47 | n.a. | 2,064 | 0,128 | 0,79 | n.a. | 19,50 |
| 2 | 8,35 | n.a. | 242,192 | 16,037 | 99,07 | n.a. | 17,62 |
| 3 | 10,08 | n.a. | 0,370 | 0,023 | 0,14 | n.a. | n.a. |
| Total: | | | 244,626 | 16,188 | 100,00 | 0,000 | |

Compound 16 - TUG-2372

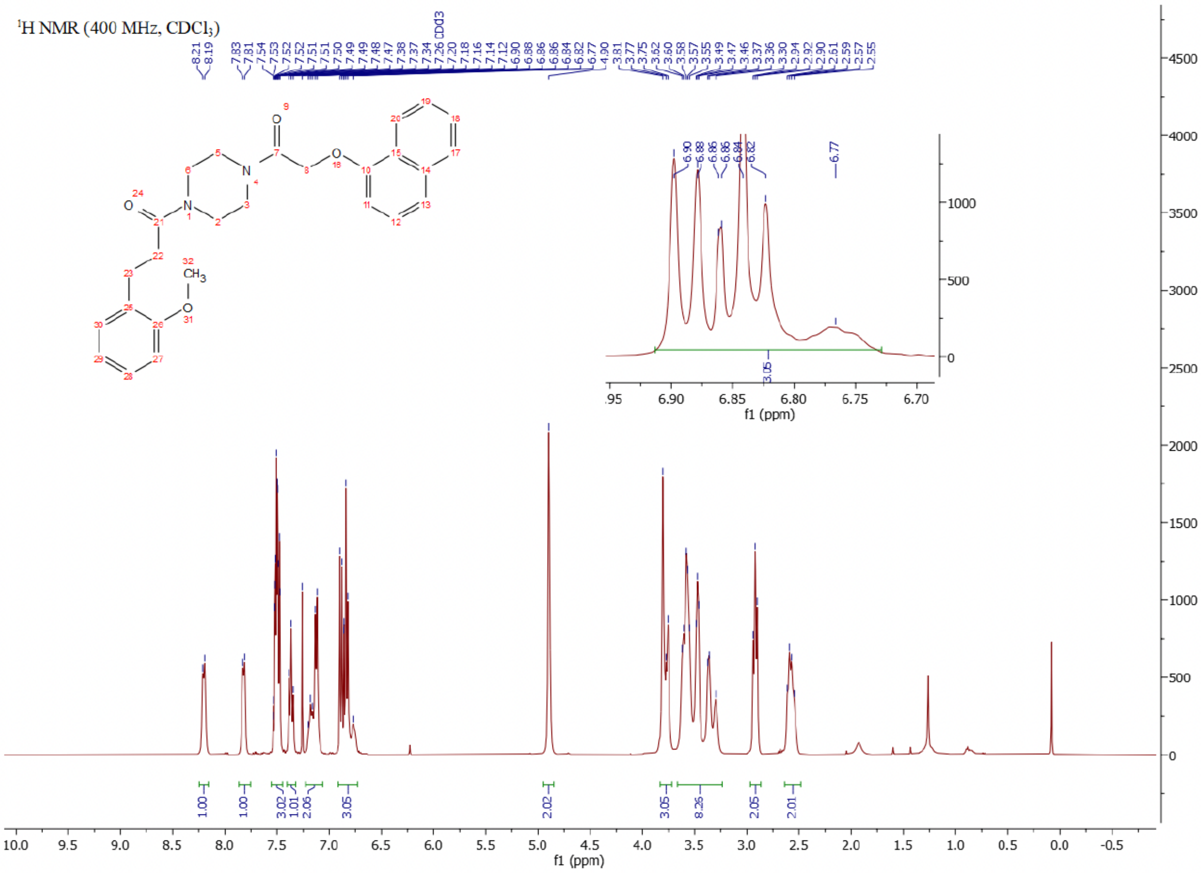

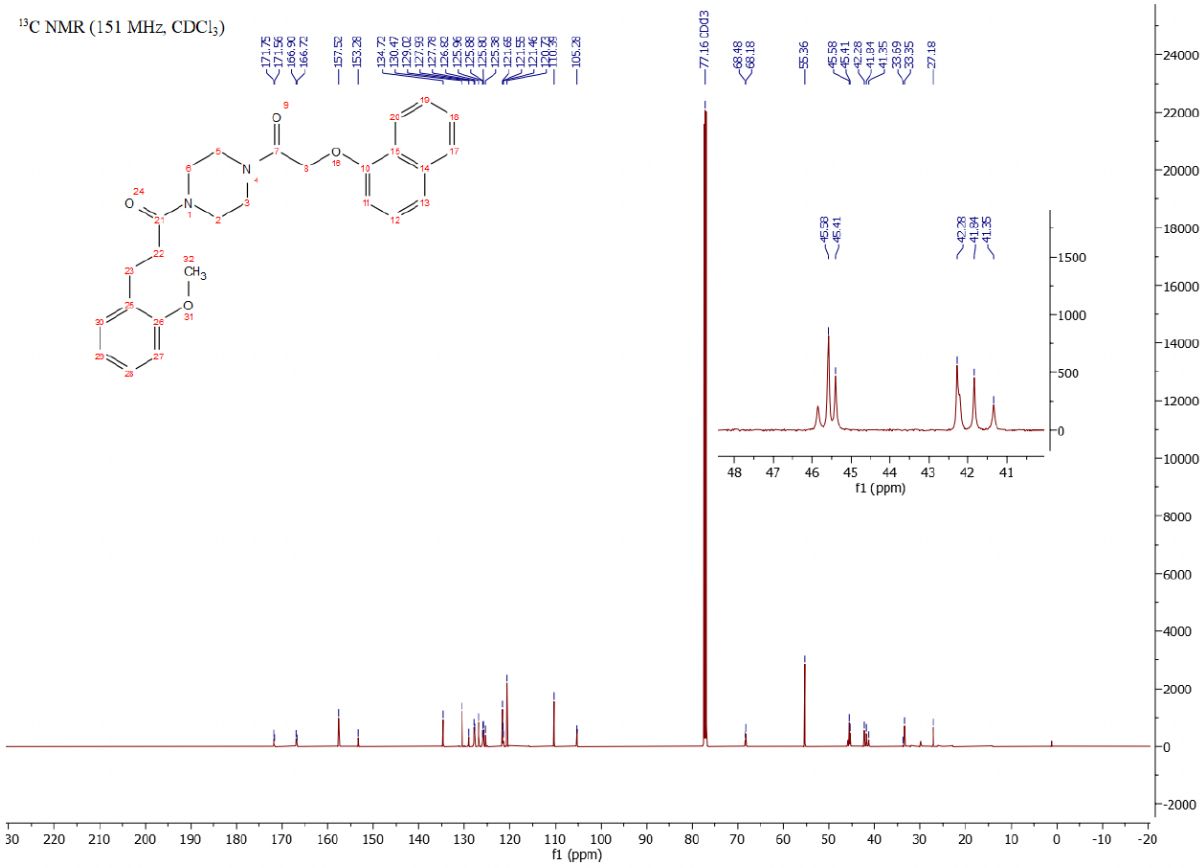

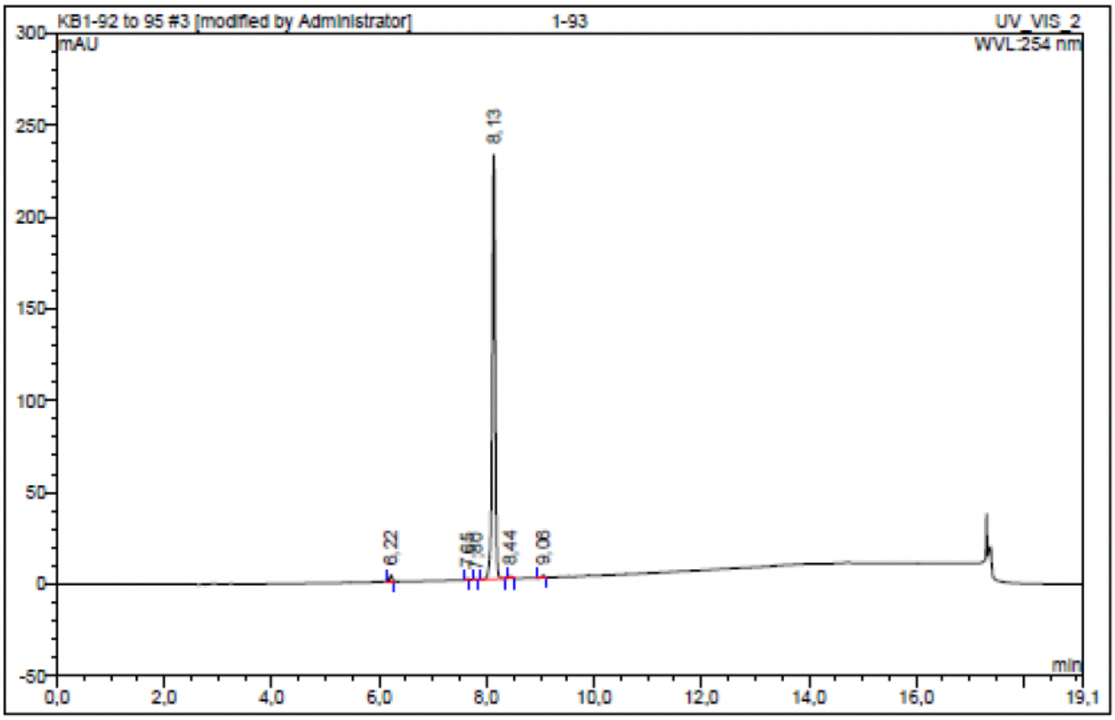

| No. | Ret.Time min | Peak Name | Height mAU | Area mAU*min | Rel.Area % | Amount | Resolution(EP) |
|---|---|---|---|---|---|---|---|
| 1 | 6,22 | n.a. | 3,257 | 0,182 | 1,18 | n.a. | 17,31 |
| 2 | 7,65 | n.a. | 0,623 | 0,028 | 0,18 | n.a. | 2,28 |
| 3 | 7,80 | n.a. | 0,171 | 0,006 | 0,04 | n.a. | 4,35 |
| 4 | 8,13 | n.a. | 231,150 | 15,114 | 97,76 | n.a. | 3,29 |
| 5 | 8,44 | n.a. | 1,239 | 0,070 | 0,45 | n.a. | 6,75 |
| 6 | 9,06 | n.a. | 1,100 | 0,061 | 0,40 | n.a. | n.a. |
| Total: | | | 237,540 | 15,461 | 100,00 | 0,000 | |

### Compound 17 - TUG-2260

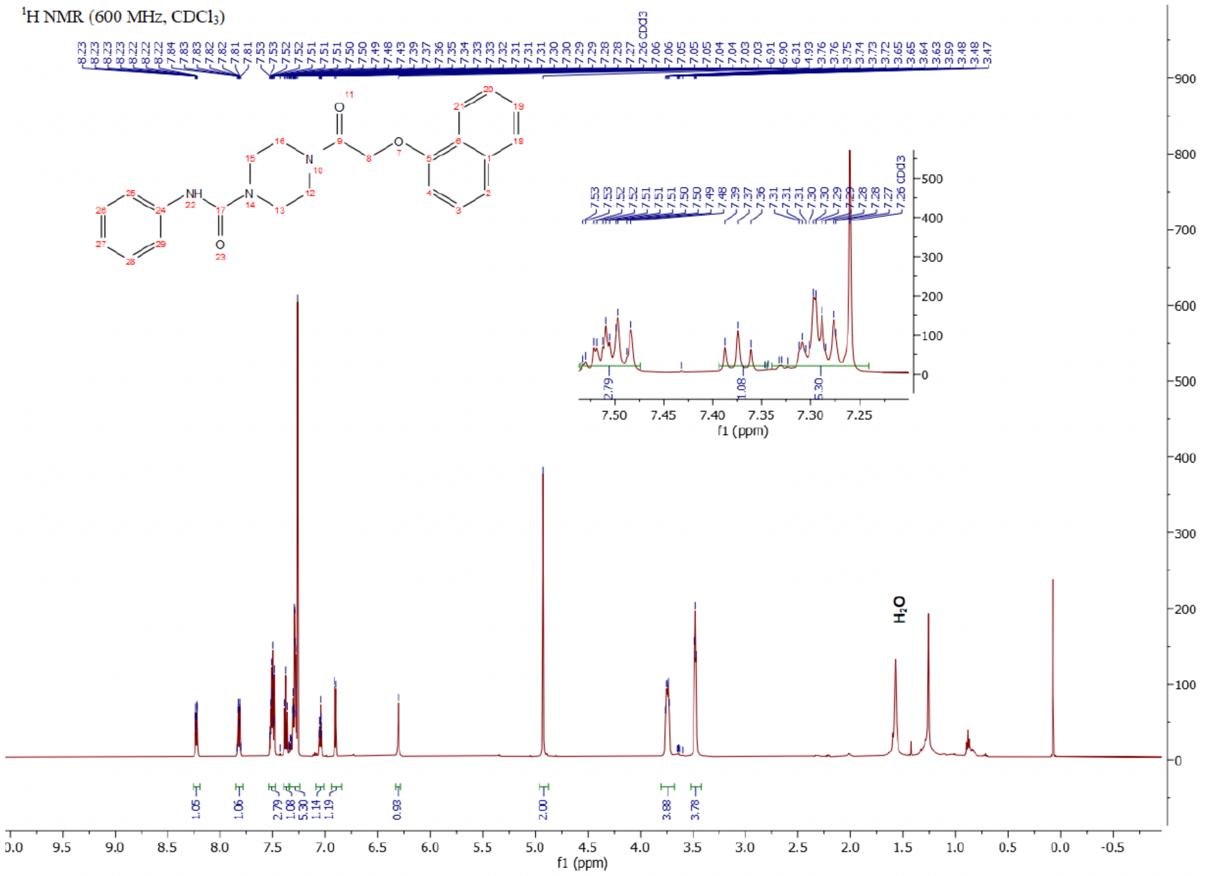

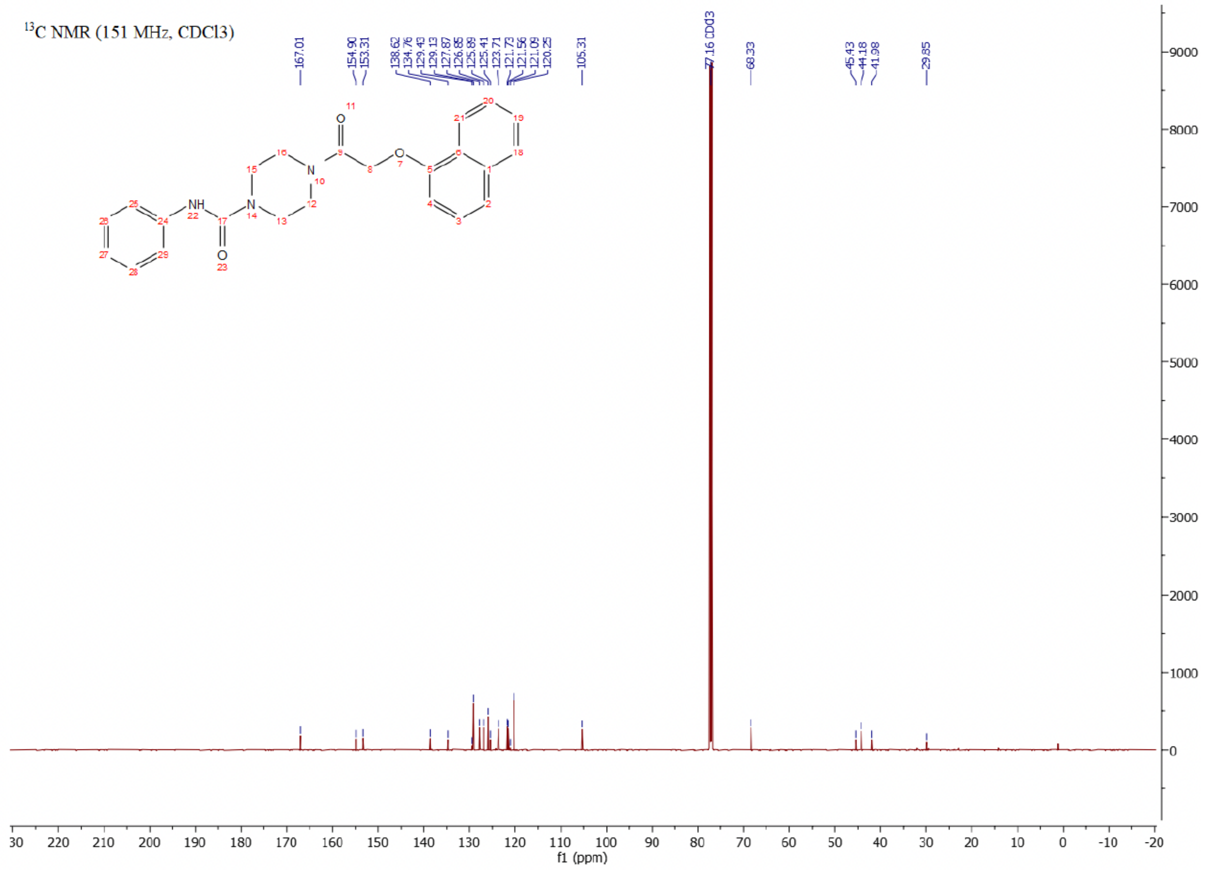

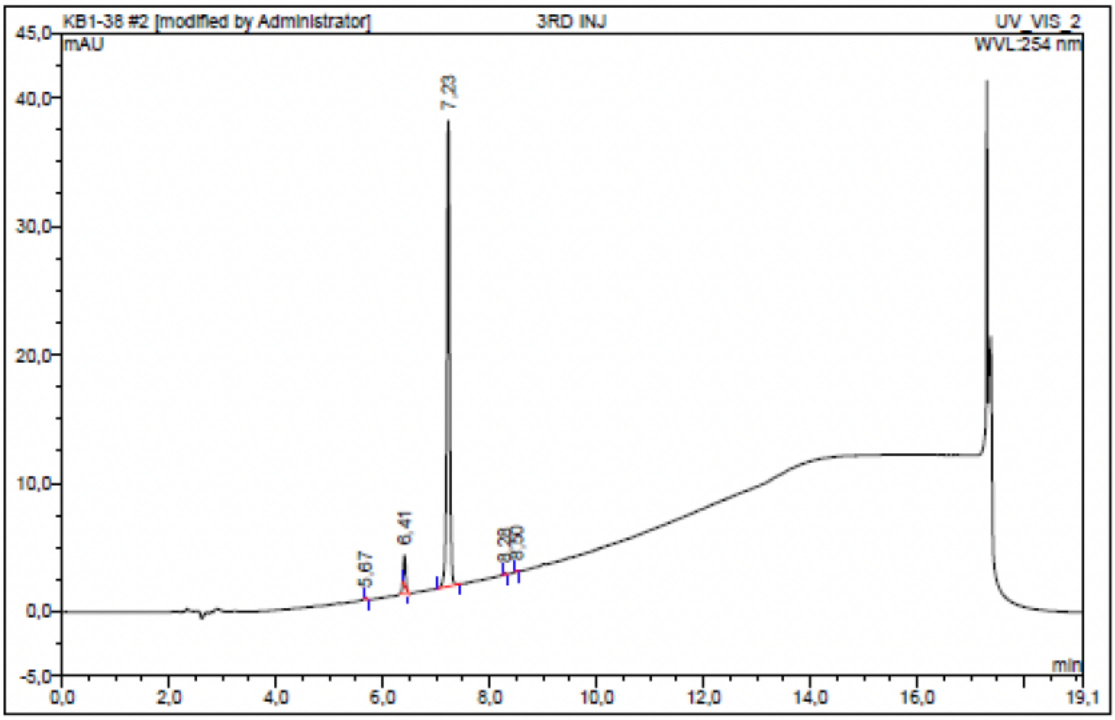

| No. | Ret.Time min | Peak Name | Height mAU | Area mAU*min | Rel.Area % | Amount | Resolution(EP) |
|---|---|---|---|---|---|---|---|
| 1 | 5,67 | n.a. | 0,113 | 0,005 | 0,19 | n.a. | 10,04 |
| 2 | 6,41 | n.a. | 2,353 | 0,110 | 4,52 | n.a. | 9,51 |
| 3 | 7,23 | n.a. | 36,261 | 2,304 | 95,01 | n.a. | 12,25 |
| 4 | 8,28 | n.a. | 0,122 | 0,005 | 0,22 | n.a. | 3,37 |
| 5 | 8,50 | n.a. | 0,046 | 0,002 | 0,06 | n.a. | n.a. |
| Total: | | | 38,895 | 2,425 | 100,00 | 0,000 | |

Compound 18 - TUG-2609

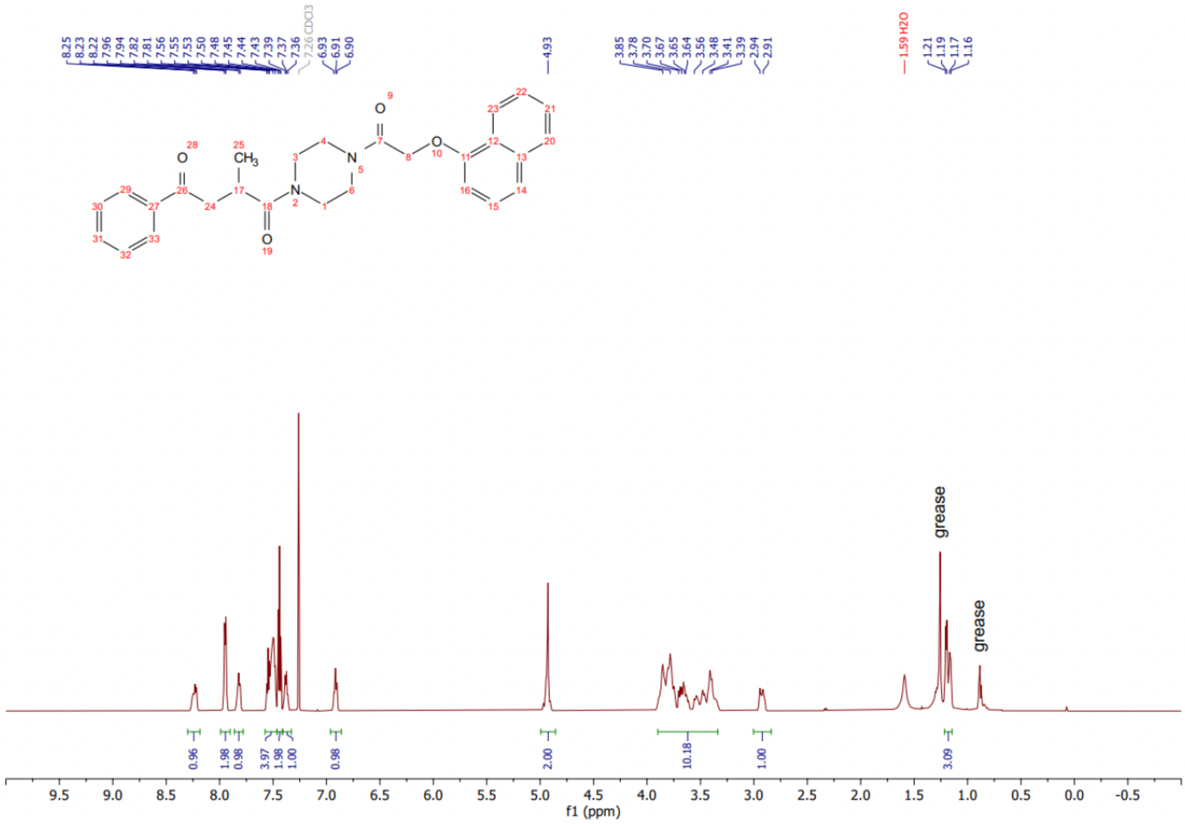

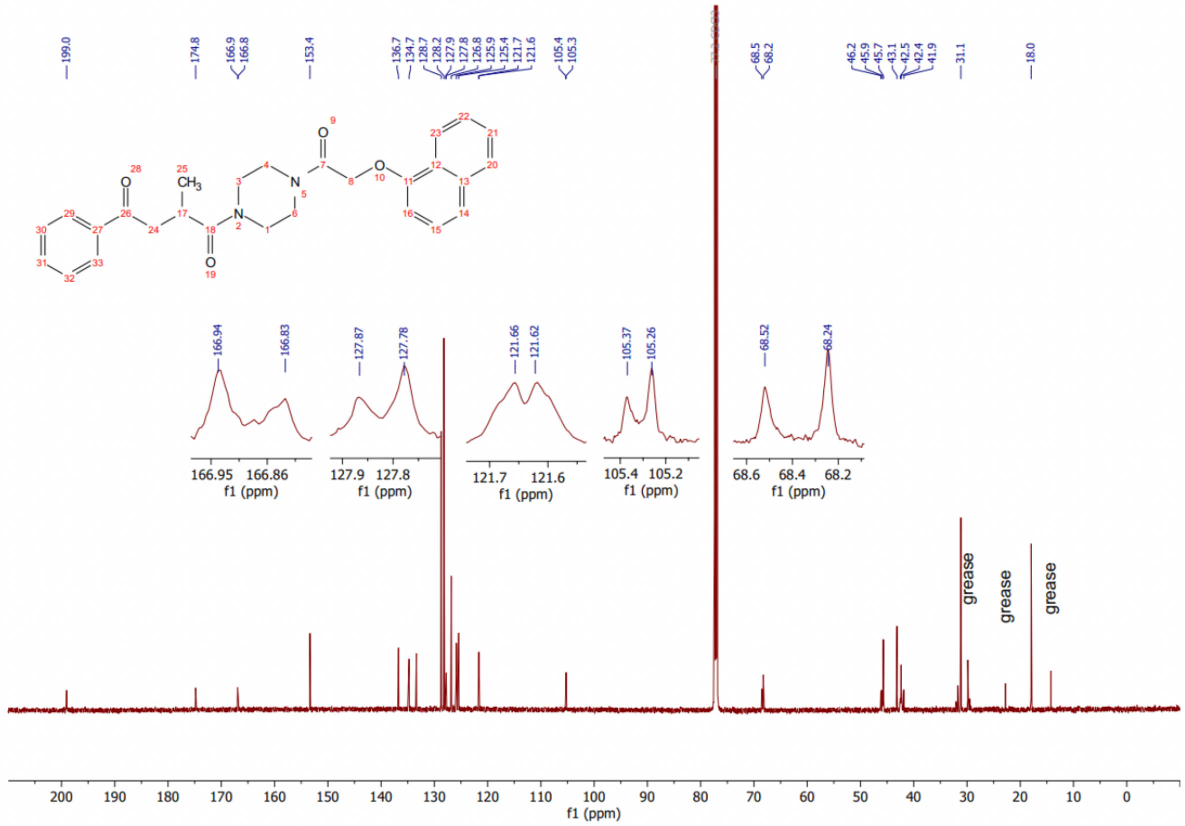

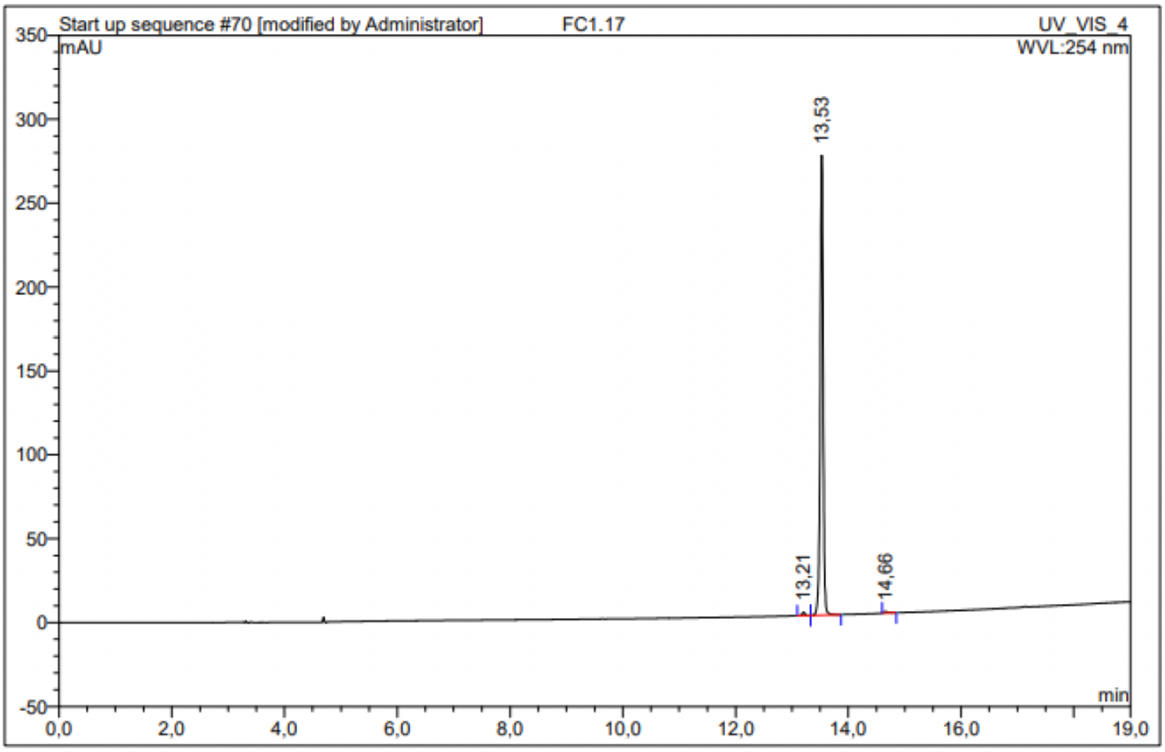

| No. | Ret.Time min | Peak Name | Height mAU | Area mAU*min | Rel.Area % | Amount | Resolution(EP) |
|---|---|---|---|---|---|---|---|
| 1 | 13,21 | n.a. | 2,089 | 0,119 | 0,70 | n.a. | 3,63 |
| 2 | 13,53 | n.a. | 274,074 | 16,735 | 98,86 | n.a. | 11,13 |
| 3 | 14,66 | n.a. | 0,862 | 0,075 | 0,44 | n.a. | n.a. |
| Total: | | | 277,025 | 16,929 | 100,00 | 0,000 | |

Compound 19 - TUG-2363

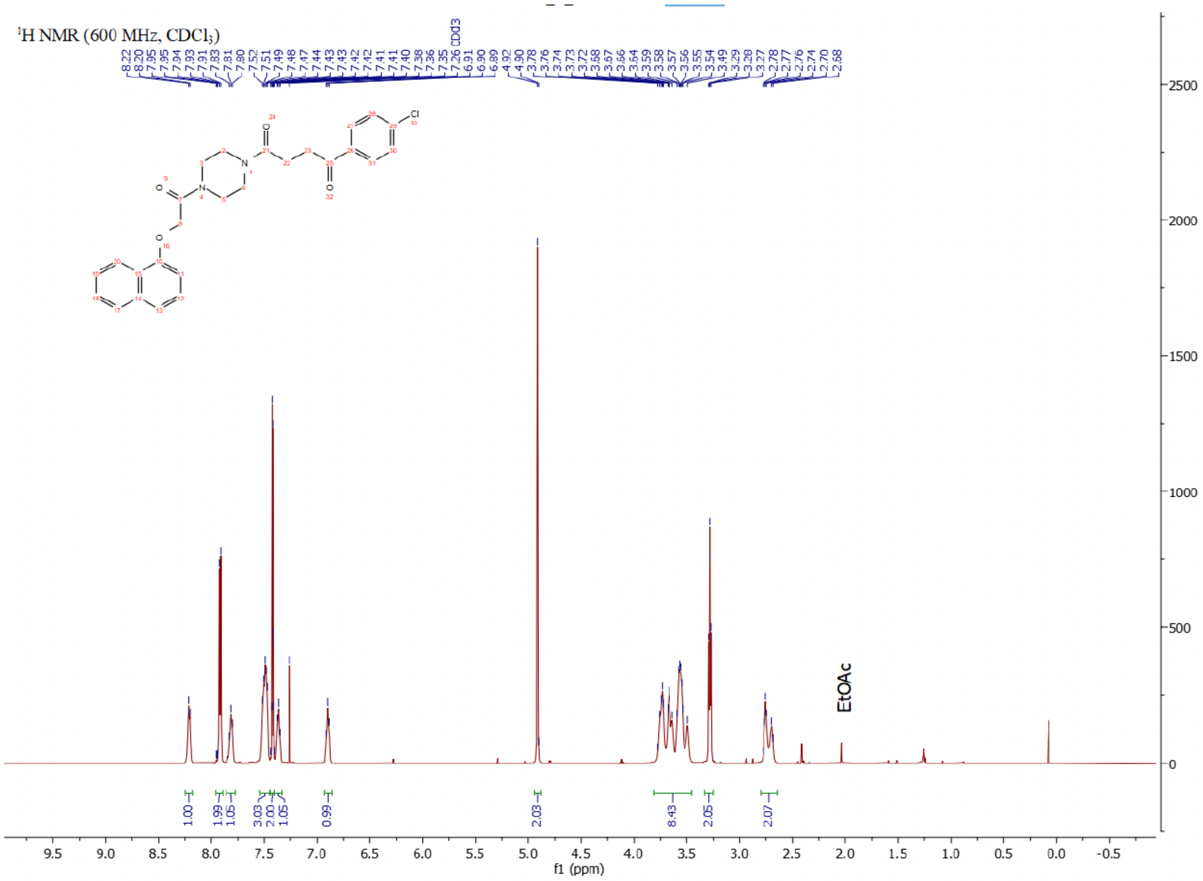

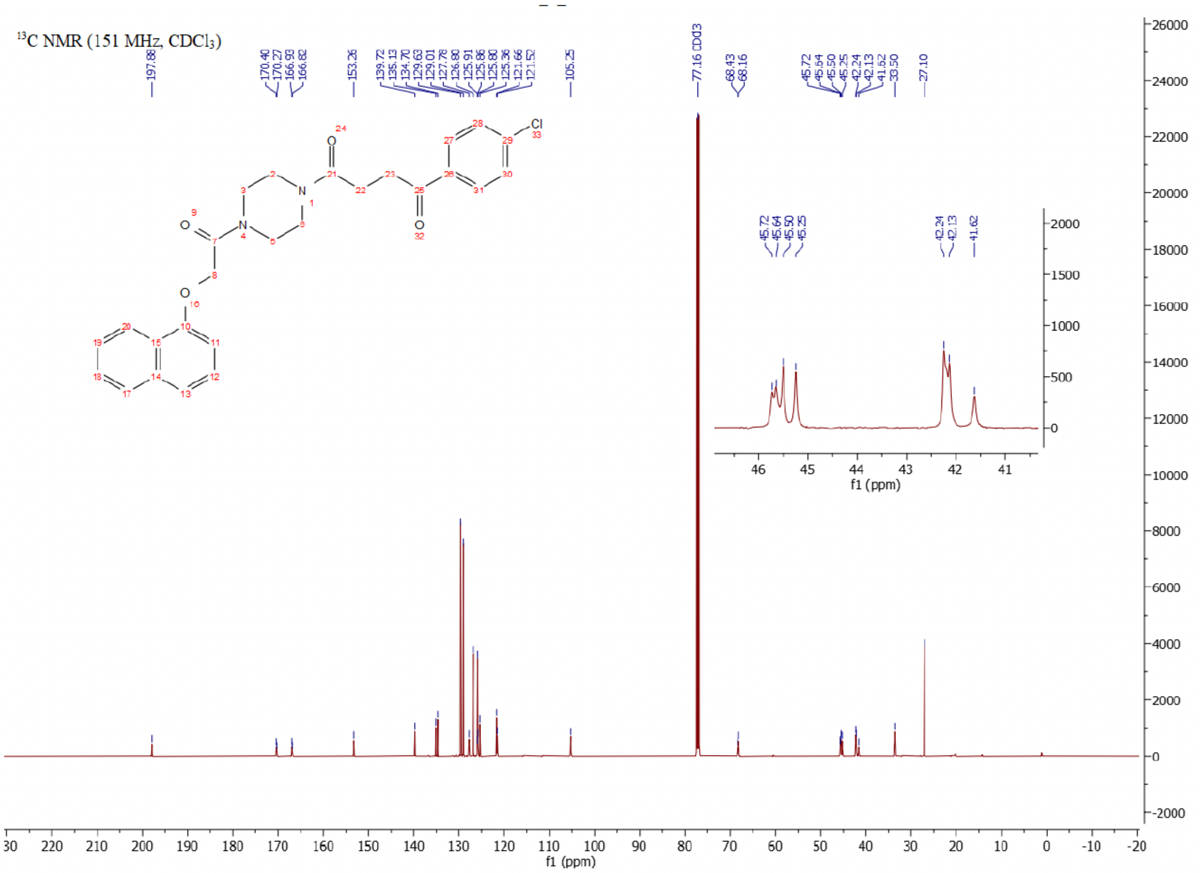

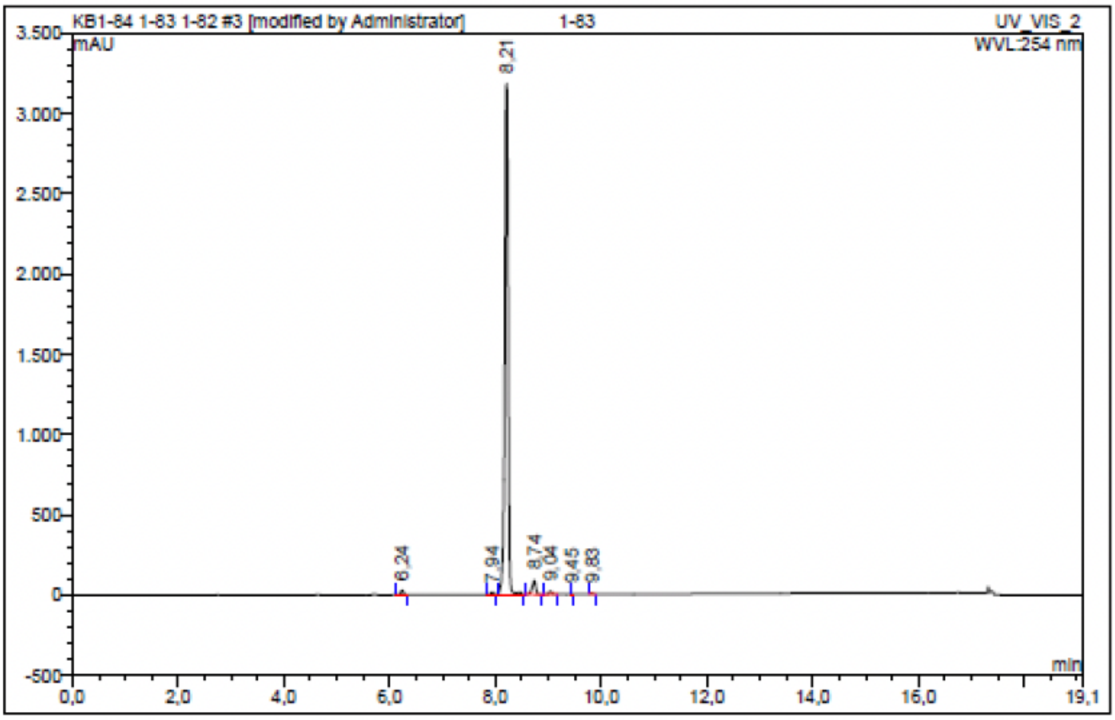

| No. | Ret.Time min | | Peak Name | Height mAU | Area mAU*min | Rel.Area % | Amount | Resolution(EP) |
|---|---|---|---|---|---|---|---|---|
| 1 | 6,24 | n.a. | | 27,948 | 1,716 | 0,68 | n.a. | 17,89 |
| 2 | 7,94 | n.a. | | 13,509 | 0,820 | 0,32 | n.a. | 2,55 |
| 3 | 8,21 | n.a. | | 3179,244 | 243,068 | 95,63 | n.a. | 4,44 |
| 4 | 8,74 | n.a. | | 86,624 | 6,878 | 2,71 | n.a. | 2,81 |
| 5 | 9,04 | n.a. | | 18,972 | 1,255 | 0,49 | n.a. | 4,70 |
| 6 | 9,45 | n.a. | | 1,390 | 0,061 | 0,02 | n.a. | 4,44 |
| 7 | 9,83 | n.a. | | 6,409 | 0,369 | 0,15 | n.a. | n.a. |
| Total: | | | | 3334,096 | 254,167 | 100,00 | 0,000 | |

Compound 20 - TUG-2256

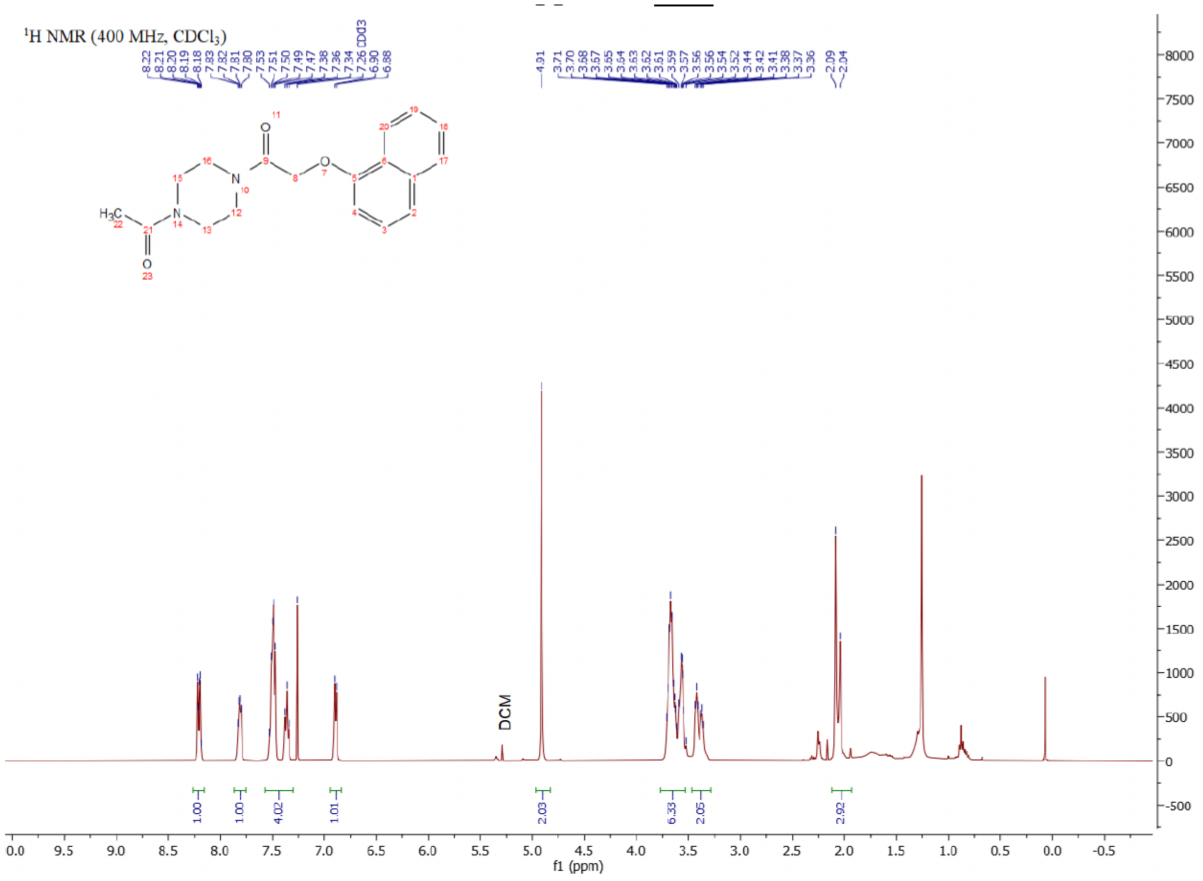

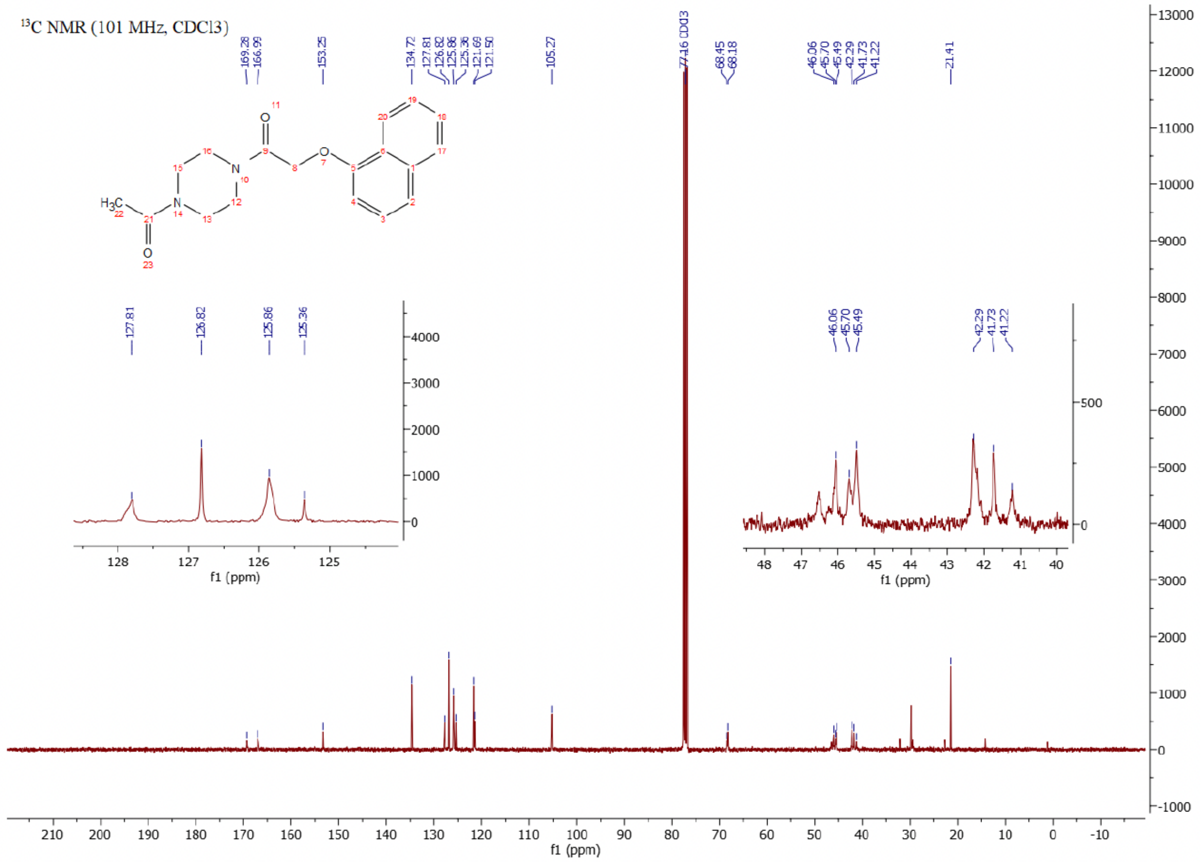

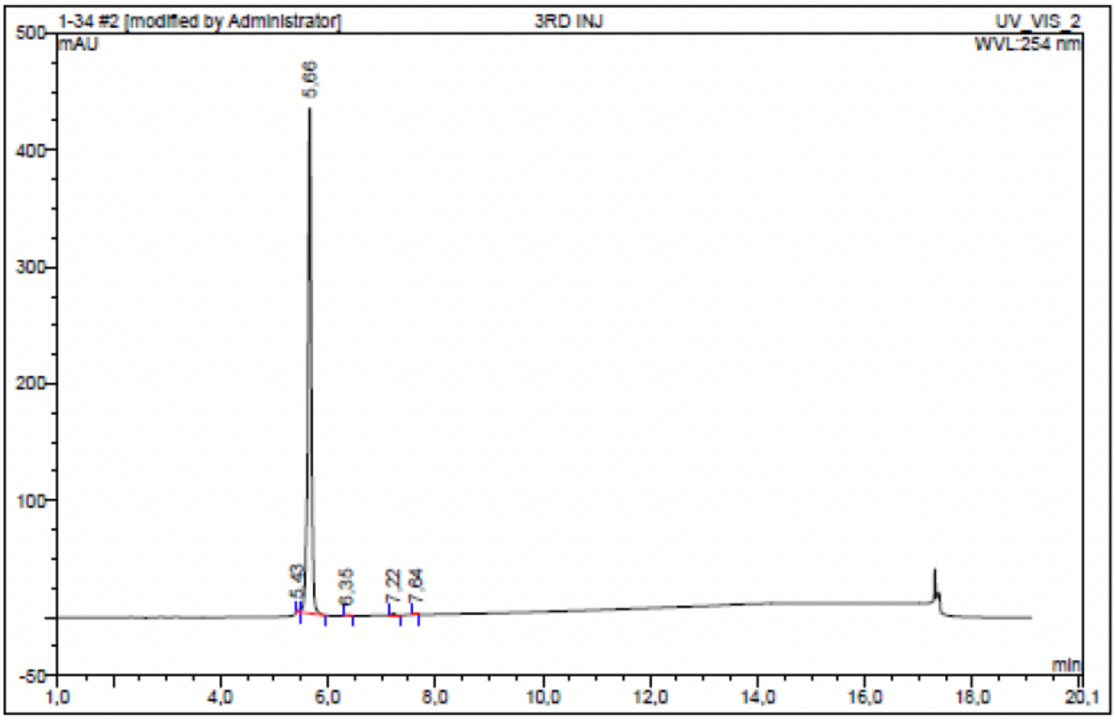

| No. | Ret.Time min | Peak Name | Height mAU | Area mAU*min | Rel.Area % | Amount | Resolution(EP) |
|---|---|---|---|---|---|---|---|
| 1 | 5,43 | n.a. | 1,674 | 0,057 | 0,18 | n.a. | 2,82 |
| 2 | 5,66 | n.a. | 432,465 | 32,280 | 99,40 | n.a. | 5,30 |
| 3 | 6,35 | n.a. | 0,130 | 0,012 | 0,04 | n.a. | 6,92 |
| 4 | 7,22 | n.a. | 1,262 | 0,080 | 0,25 | n.a. | 4,60 |
| 5 | 7,64 | n.a. | 0,913 | 0,047 | 0,14 | n.a. | n.a. |
| Total: | | | 436,444 | 32,475 | 100,00 | 0,000 | |

Compound 21 - TUG-2293

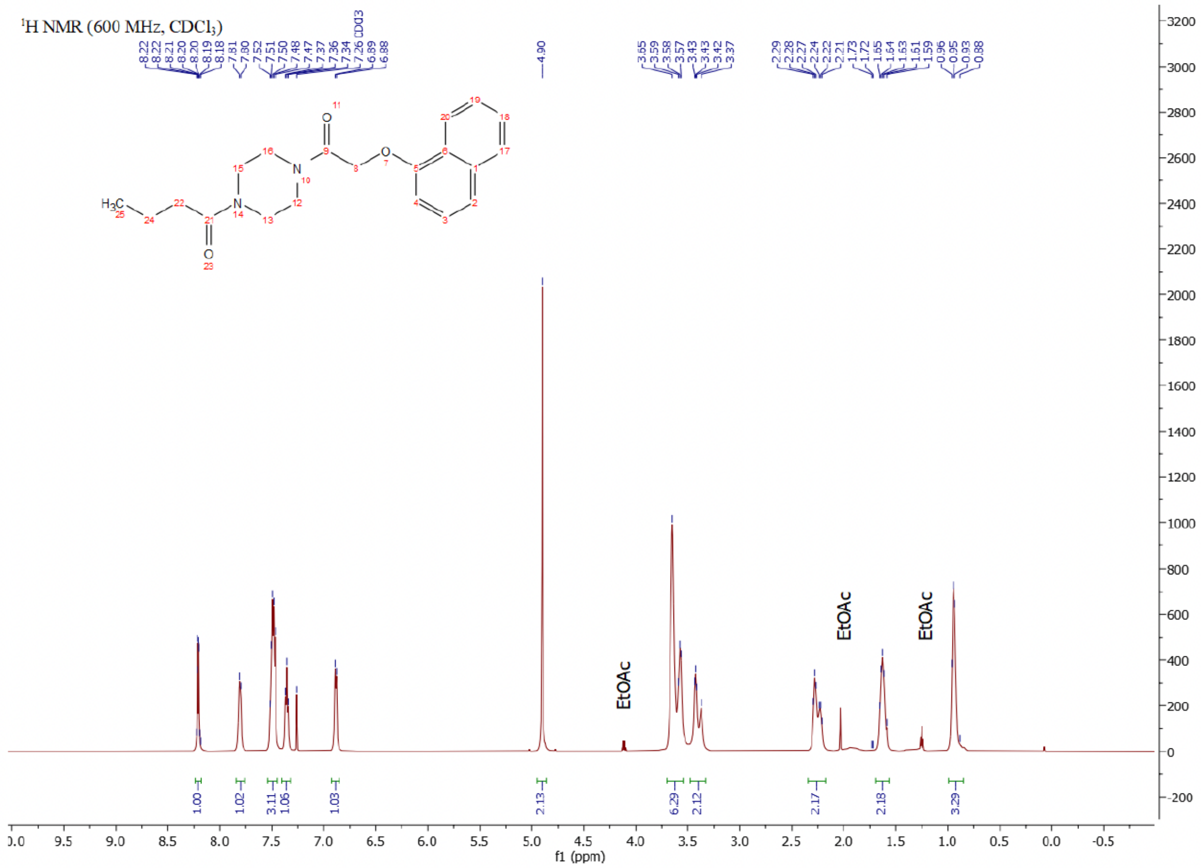

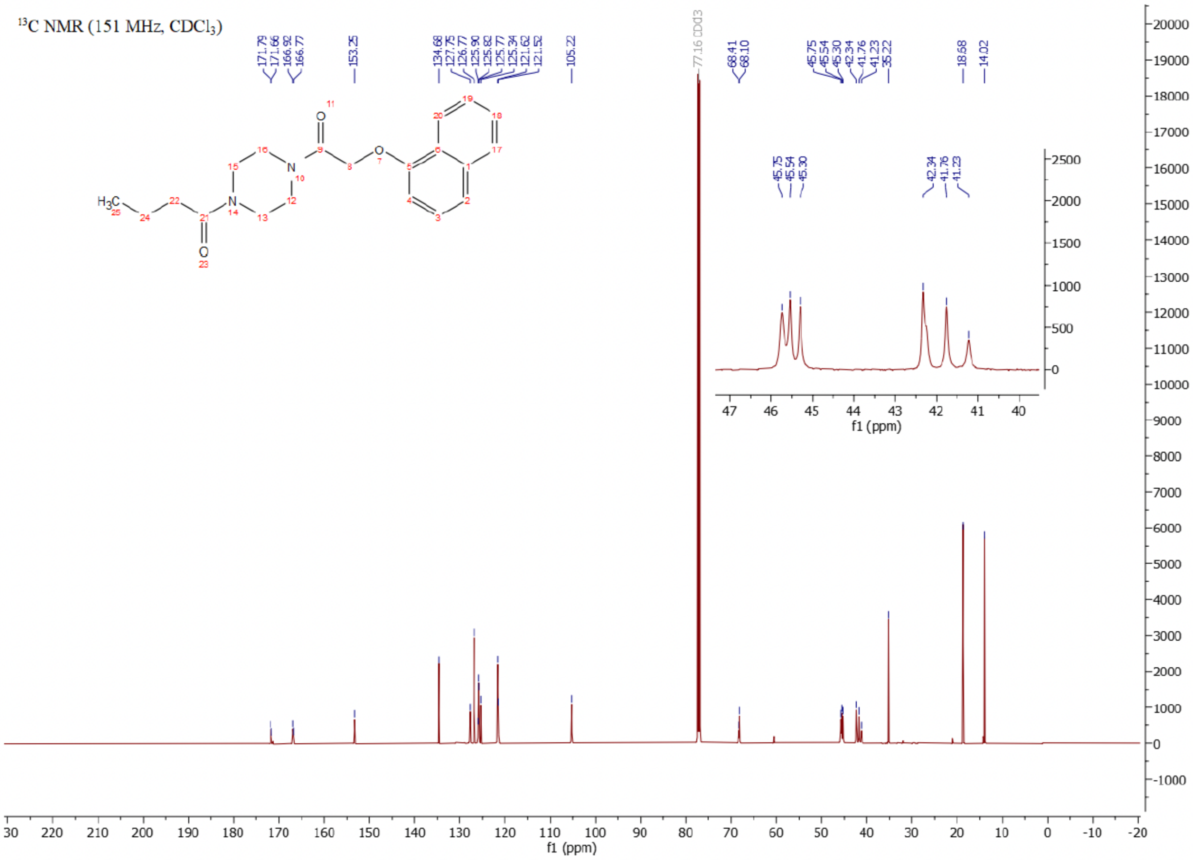

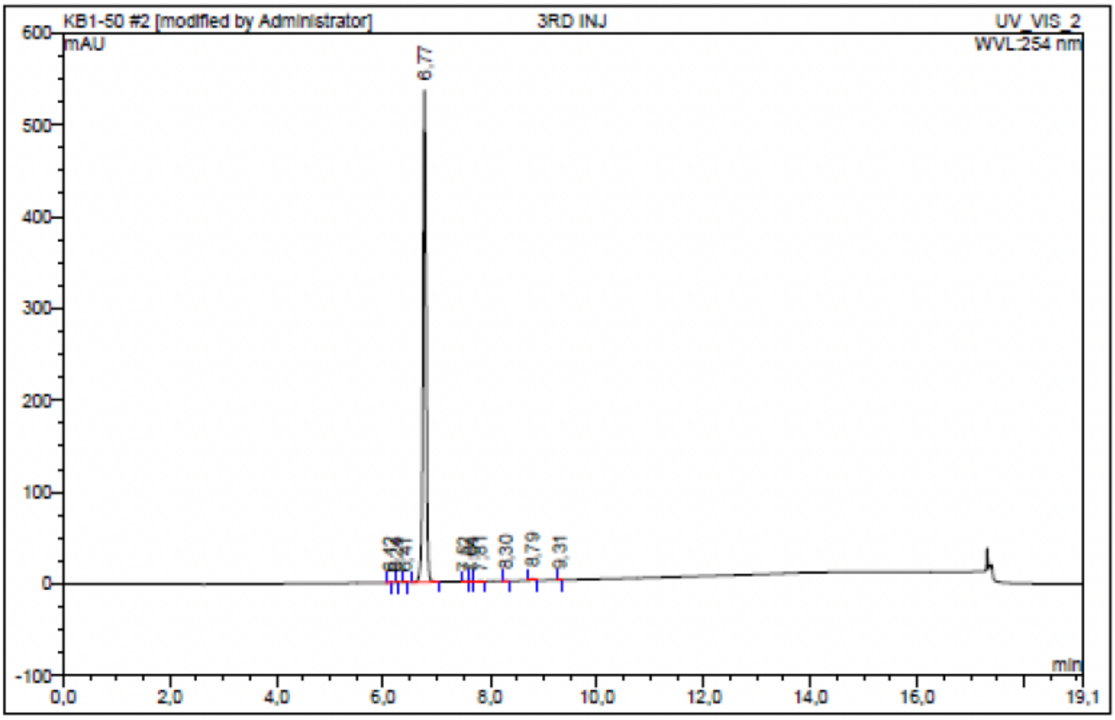

| No. | Ret.Time min | Peak Name | Height mAU | Area mAU*min | Rel.Area % | Amount | Resolution(EP) |
|---|---|---|---|---|---|---|---|
| 1 | 6,12 | n.a. | 0,255 | 0,010 | 0,03 | n.a. | 2,16 |
| 2 | 6,24 | n.a. | 0,009 | 0,000 | 0,00 | n.a. | 2,80 |
| 3 | 6,41 | n.a. | 0,406 | 0,020 | 0,06 | n.a. | 4,05 |
| 4 | 6,77 | n.a. | 535,620 | 35,130 | 99,36 | n.a. | 6,84 |
| 5 | 7,52 | n.a. | 0,253 | 0,017 | 0,05 | n.a. | 1,24 |
| 6 | 7,64 | n.a. | 0,171 | 0,007 | 0,02 | n.a. | 2,10 |
| 7 | 7,81 | n.a. | 0,272 | 0,016 | 0,05 | n.a. | 5,20 |
| 8 | 8,30 | n.a. | 0,240 | 0,014 | 0,04 | n.a. | 5,12 |
| 9 | 8,79 | n.a. | 1,981 | 0,127 | 0,36 | n.a. | 5,50 |
| 10 | 9,31 | n.a. | 0,297 | 0,016 | 0,04 | n.a. | n.a. |
| Total: | | | 539,503 | 35,356 | 100,00 | 0,000 | |

Compound 22 - TUG-2330

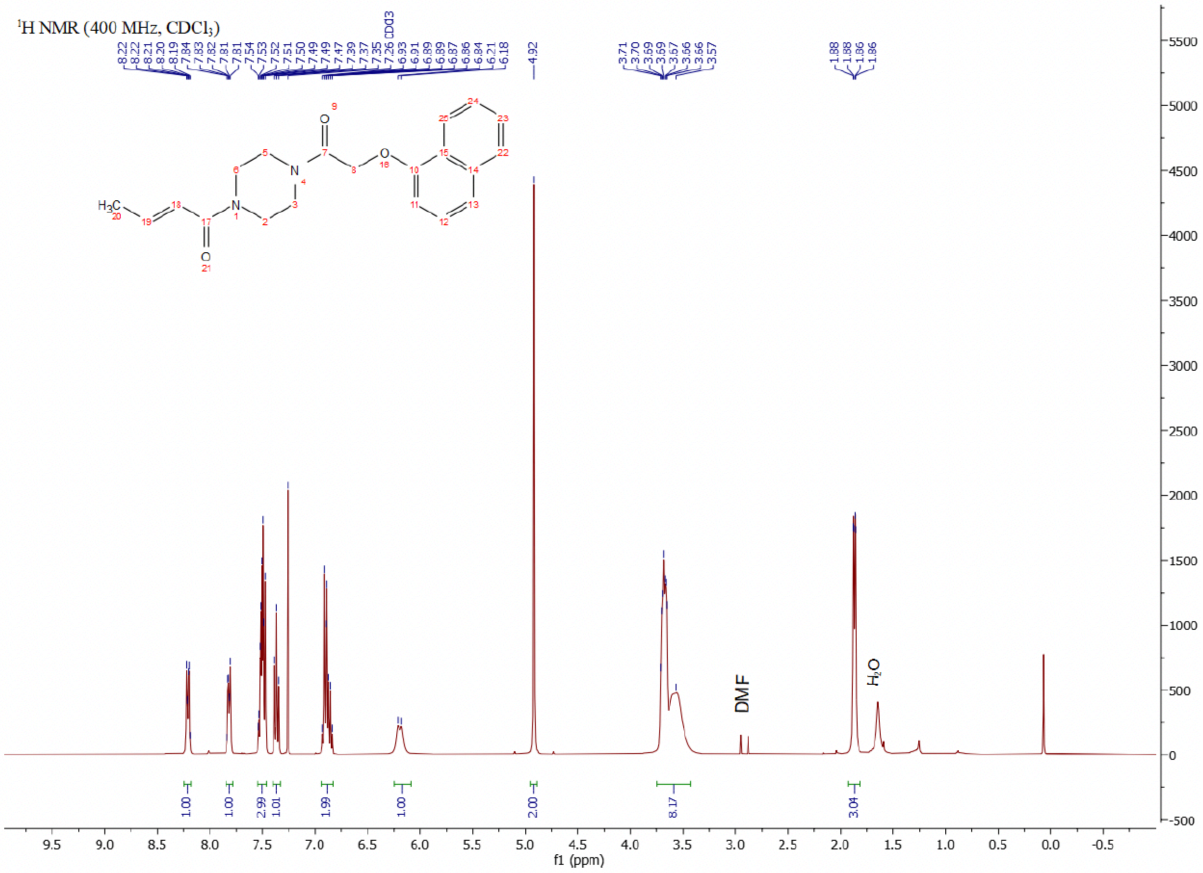

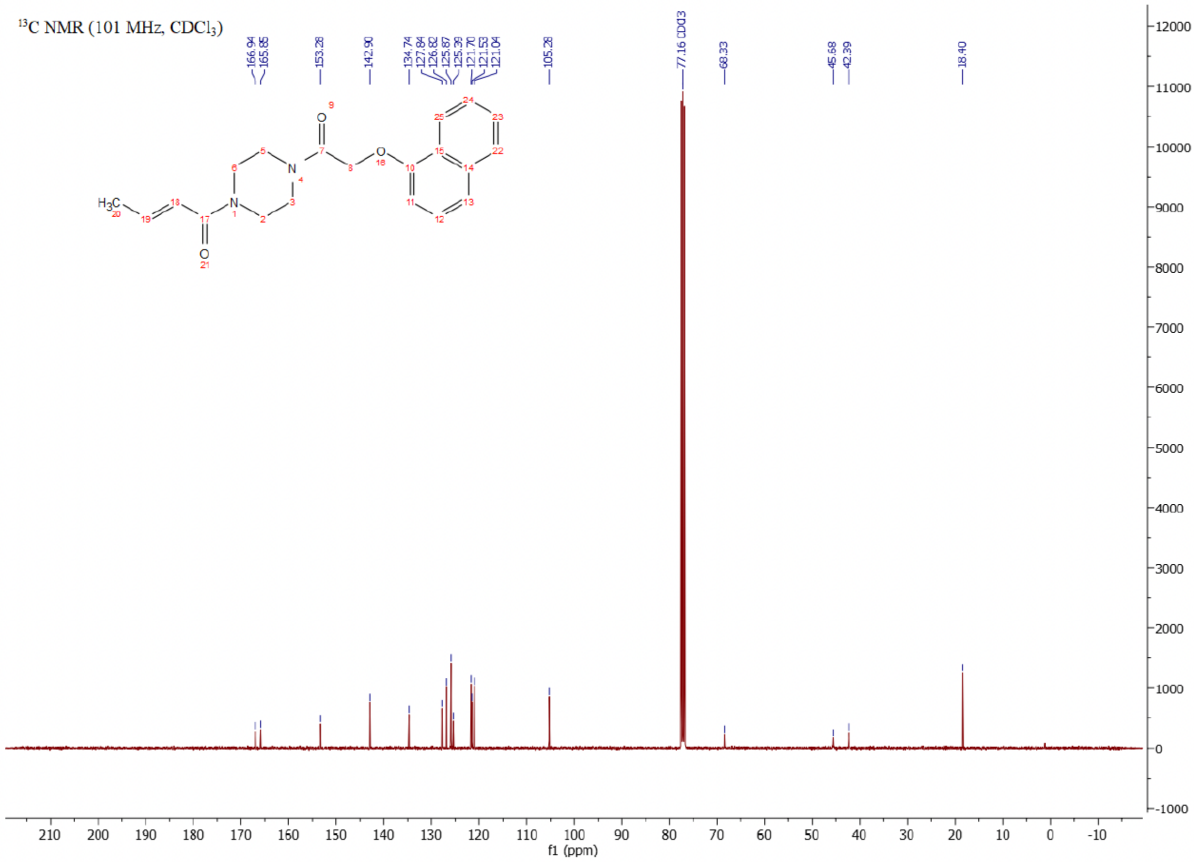

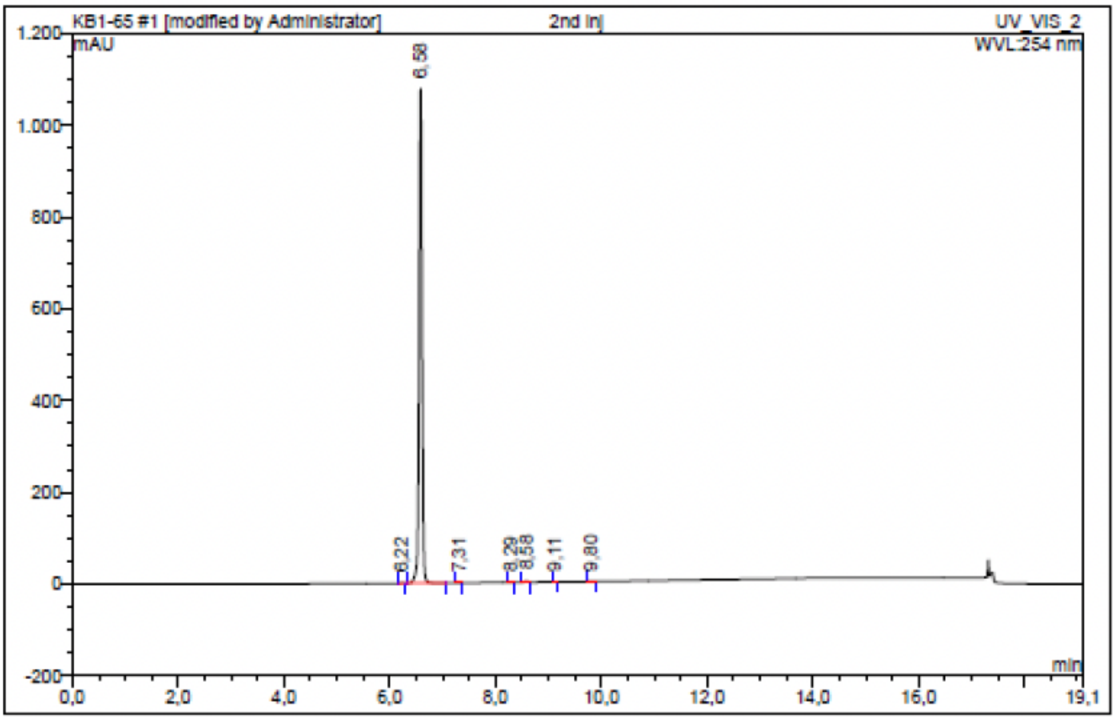

| No. | Ret.Time min | Peak Name | Height mAU | Area mAU*min | Rel.Area % | Amount | Resolution(EP) |
|---|---|---|---|---|---|---|---|
| 1 | 6,22 | n.a. | 0,321 | 0,015 | 0,02 | n.a. | 4,02 |
| 2 | 6,58 | n.a. | 1077,497 | 71,181 | 99,68 | n.a. | 7,27 |
| 3 | 7,31 | n.a. | 0,279 | 0,017 | 0,02 | n.a. | 11,02 |
| 4 | 8,29 | n.a. | 0,152 | 0,007 | 0,01 | n.a. | 3,34 |
| 5 | 8,58 | n.a. | 2,616 | 0,162 | 0,23 | n.a. | 5,90 |
| 6 | 9,11 | n.a. | 0,269 | 0,013 | 0,02 | n.a. | 7,54 |
| 7 | 9,80 | n.a. | 0,243 | 0,016 | 0,02 | n.a. | n.a. |
| Total: | | | 1081,376 | 71,412 | 100,00 | 0,000 | |

Compound 23 - TUG-2299

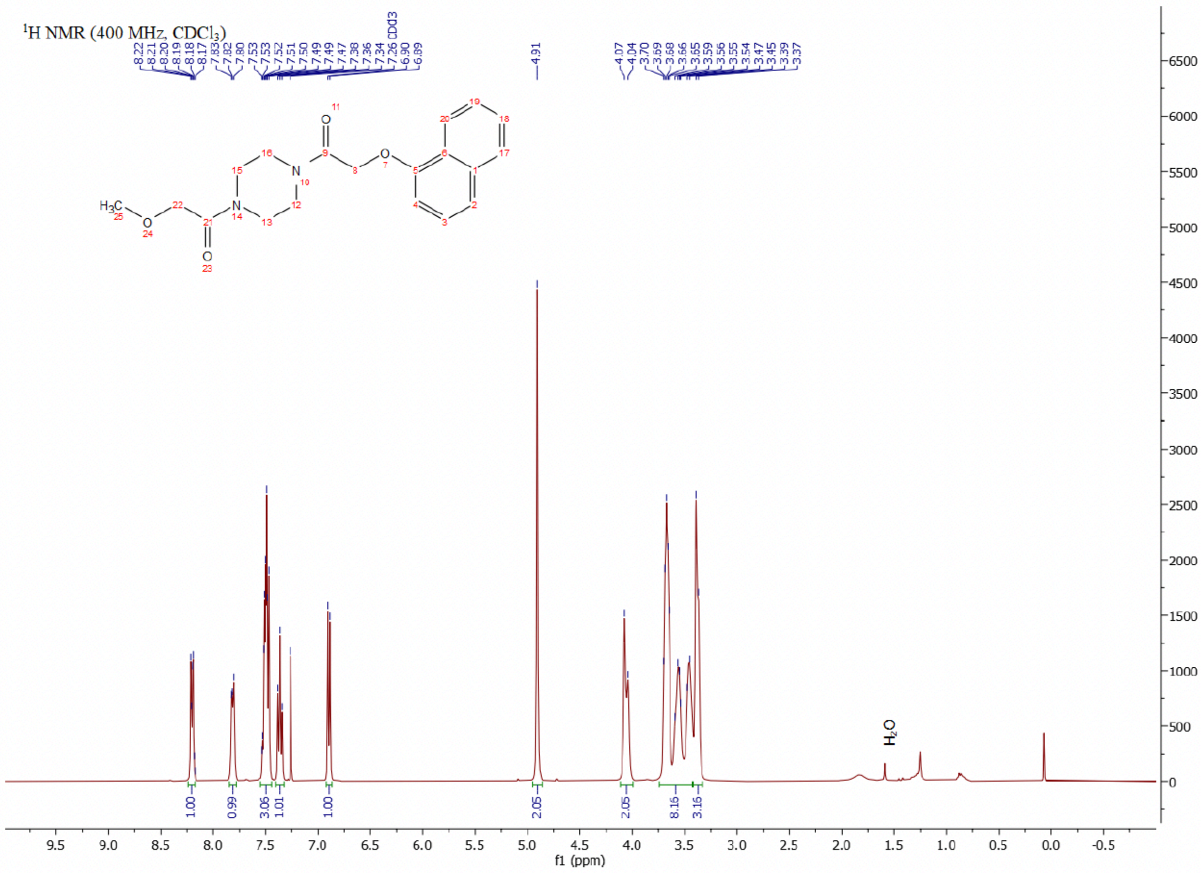

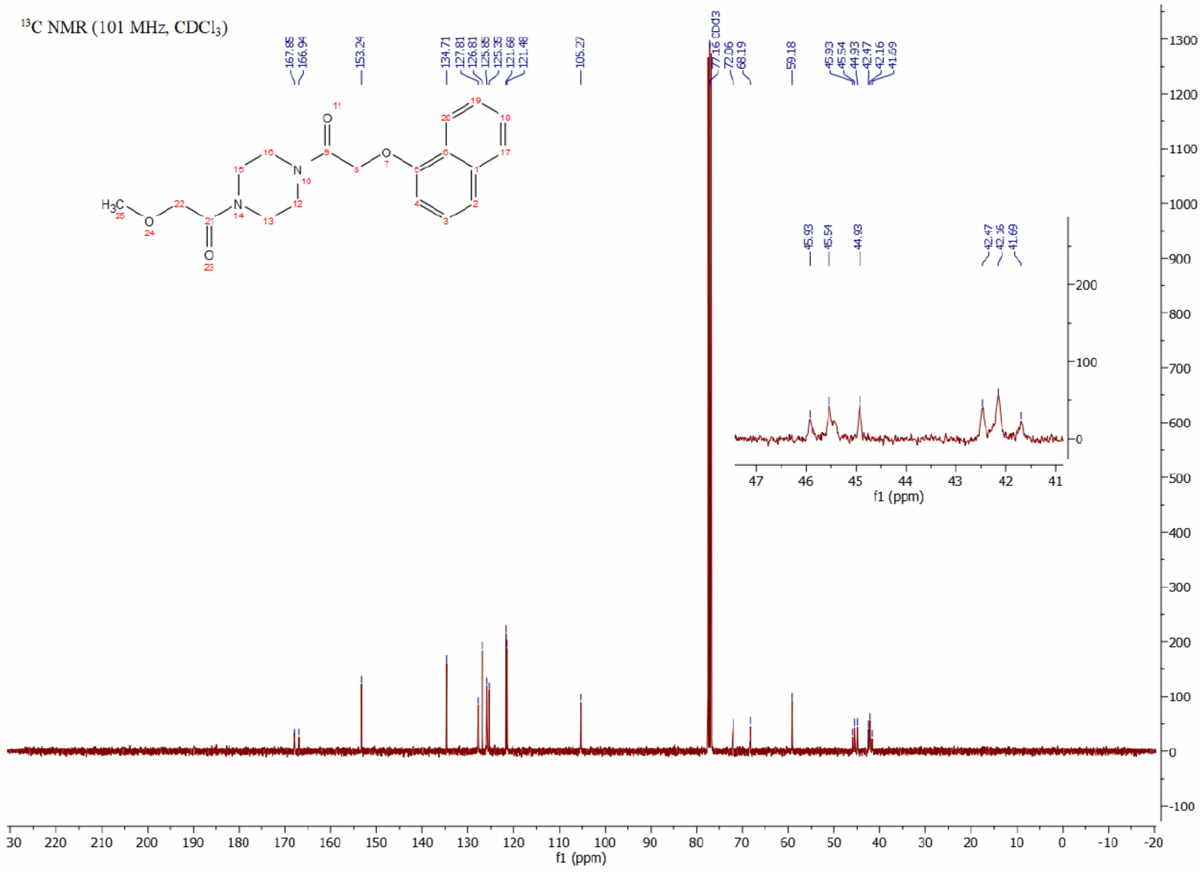

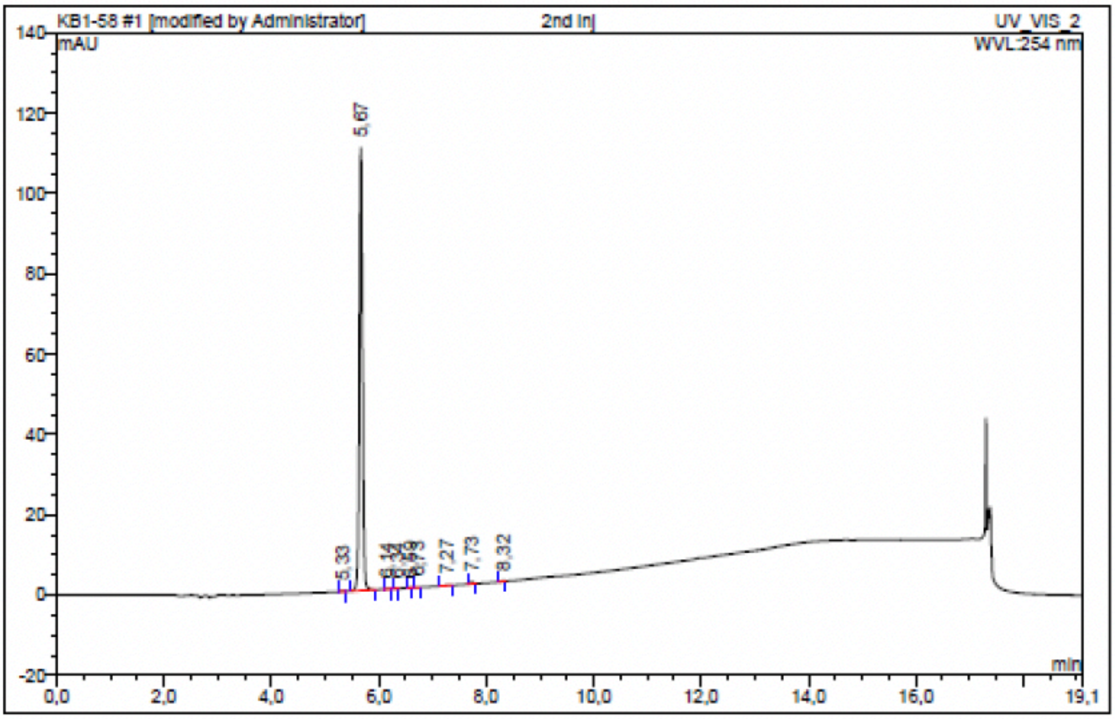

| No. | Ret.Time min | Peak Name | Height mAU | Area mAU*min | Rel.Area % | Amount | Resolution(EP) |
|---|---|---|---|---|---|---|---|
| 1 | 5,33 | n.a. | 0,100 | 0,006 | 0,08 | n.a. | 3,32 |
| 2 | 5,67 | n.a. | 110,302 | 7,483 | 99,03 | n.a. | 4,59 |
| 3 | 6,14 | n.a. | 0,071 | 0,005 | 0,06 | n.a. | 2,34 |
| 4 | 6,34 | n.a. | 0,080 | 0,004 | 0,05 | n.a. | 3,02 |
| 5 | 6,59 | n.a. | 0,067 | 0,003 | 0,05 | n.a. | 1,56 |
| 6 | 6,73 | n.a. | 0,038 | 0,002 | 0,03 | n.a. | 4,70 |
| 7 | 7,27 | n.a. | 0,104 | 0,011 | 0,14 | n.a. | 3,94 |
| 8 | 7,73 | n.a. | 0,510 | 0,032 | 0,43 | n.a. | 5,90 |
| 9 | 8,32 | n.a. | 0,166 | 0,010 | 0,13 | n.a. | n.a. |
| Total: | | | 111,436 | 7,557 | 100,00 | 0,000 | |

Compound 24 - TUG-2298

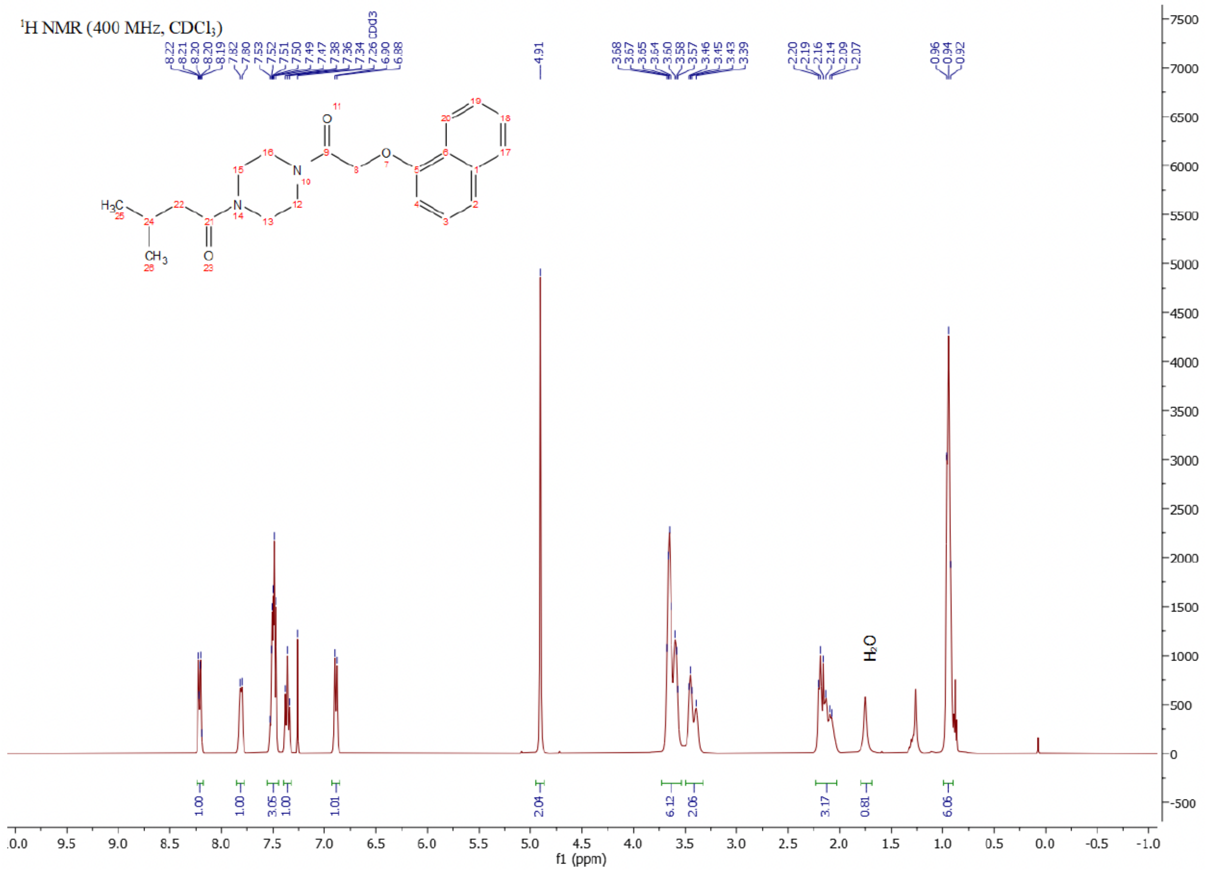

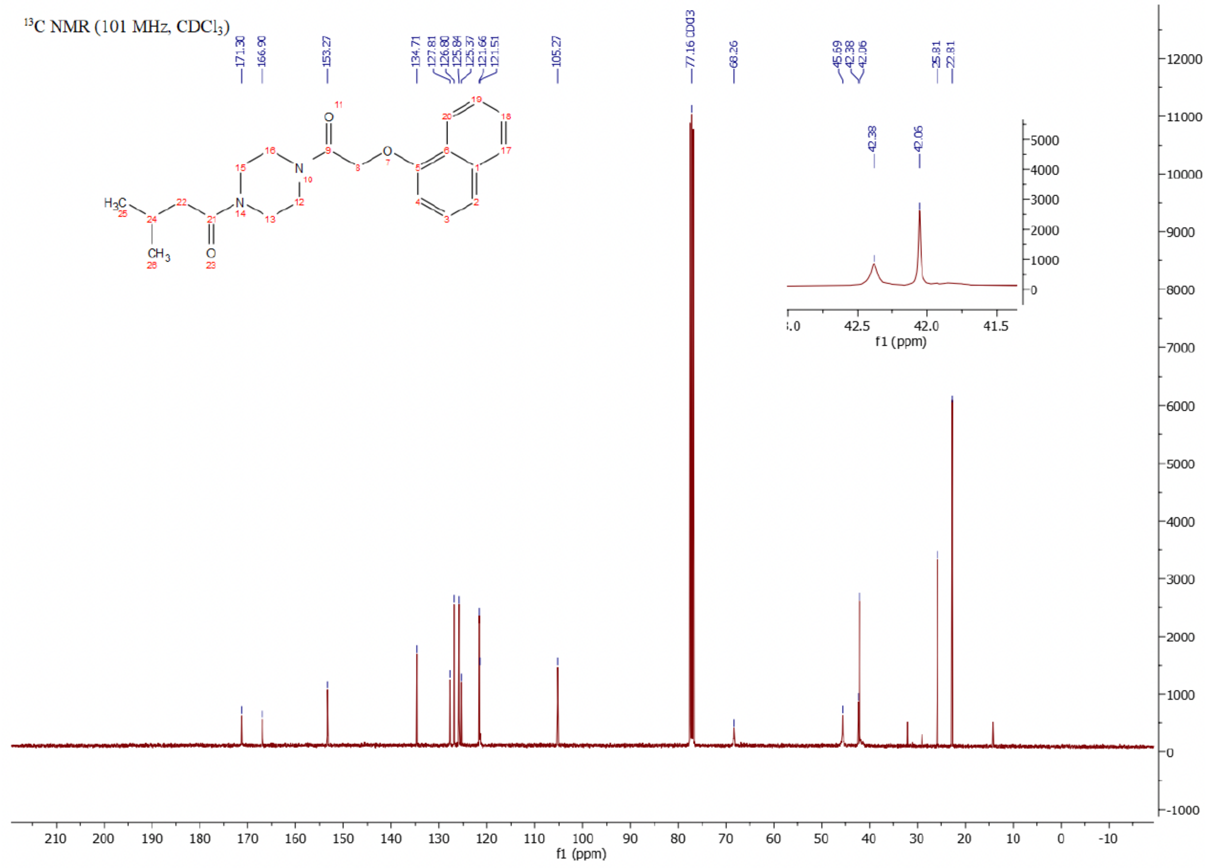

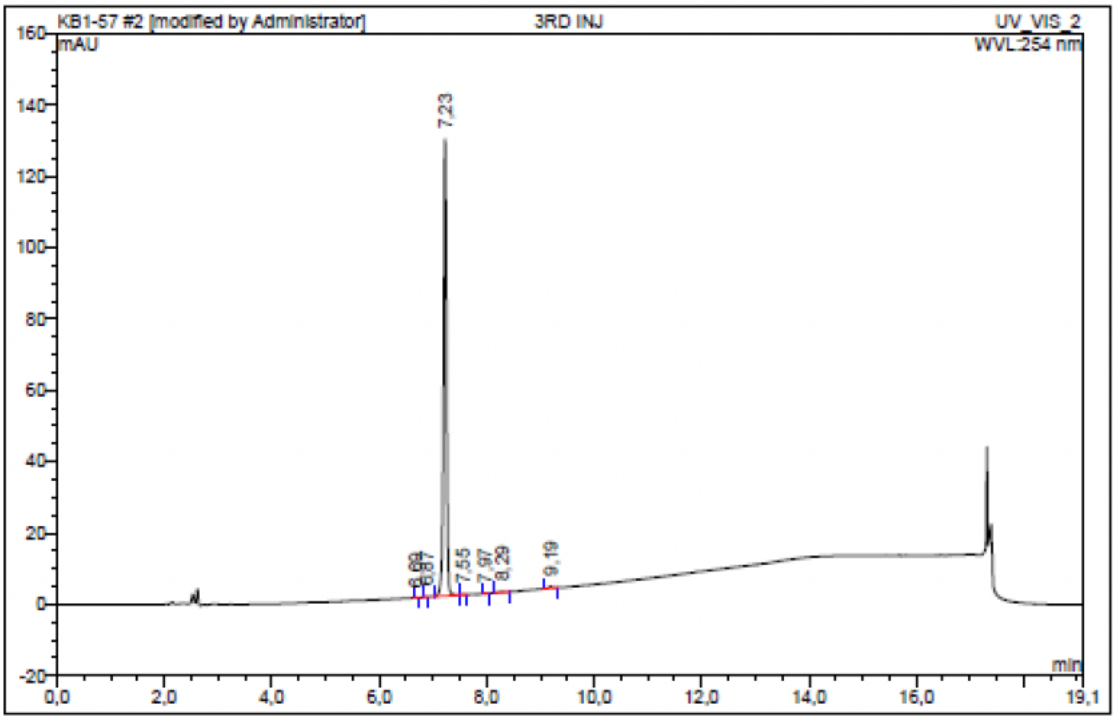

| No. | Ret.Time min | Peak Name | Height mAU | Area mAU*min | Rel.Area % | Amount | Resolution(EP) |
|---|---|---|---|---|---|---|---|
| 1 | 6,69 | n.a. | 0,096 | 0,005 | 0,06 | n.a. | 2,30 |
| 2 | 6,87 | n.a. | 0,100 | 0,005 | 0,06 | n.a. | 3,97 |
| 3 | 7,23 | n.a. | 128,003 | 8,426 | 99,12 | n.a. | 3,70 |
| 4 | 7,55 | n.a. | 0,096 | 0,005 | 0,05 | n.a. | 5,60 |
| 5 | 7,97 | n.a. | 0,054 | 0,002 | 0,03 | n.a. | 3,19 |
| 6 | 8,29 | n.a. | 0,216 | 0,019 | 0,22 | n.a. | 7,31 |
| 7 | 9,19 | n.a. | 0,487 | 0,039 | 0,46 | n.a. | n.a. |
| Total: | | | 129,052 | 8,501 | 100,00 | 0,000 | |

Compound 25 - TUG-2254

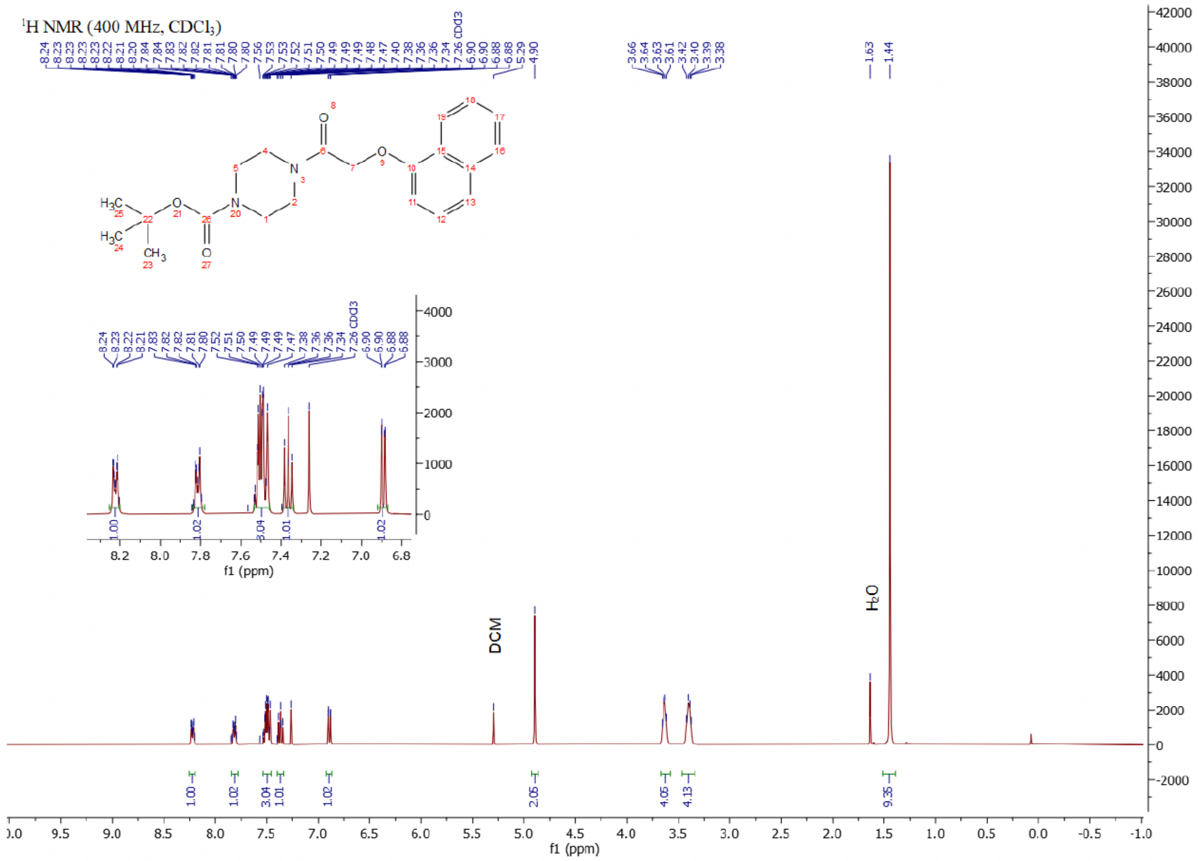

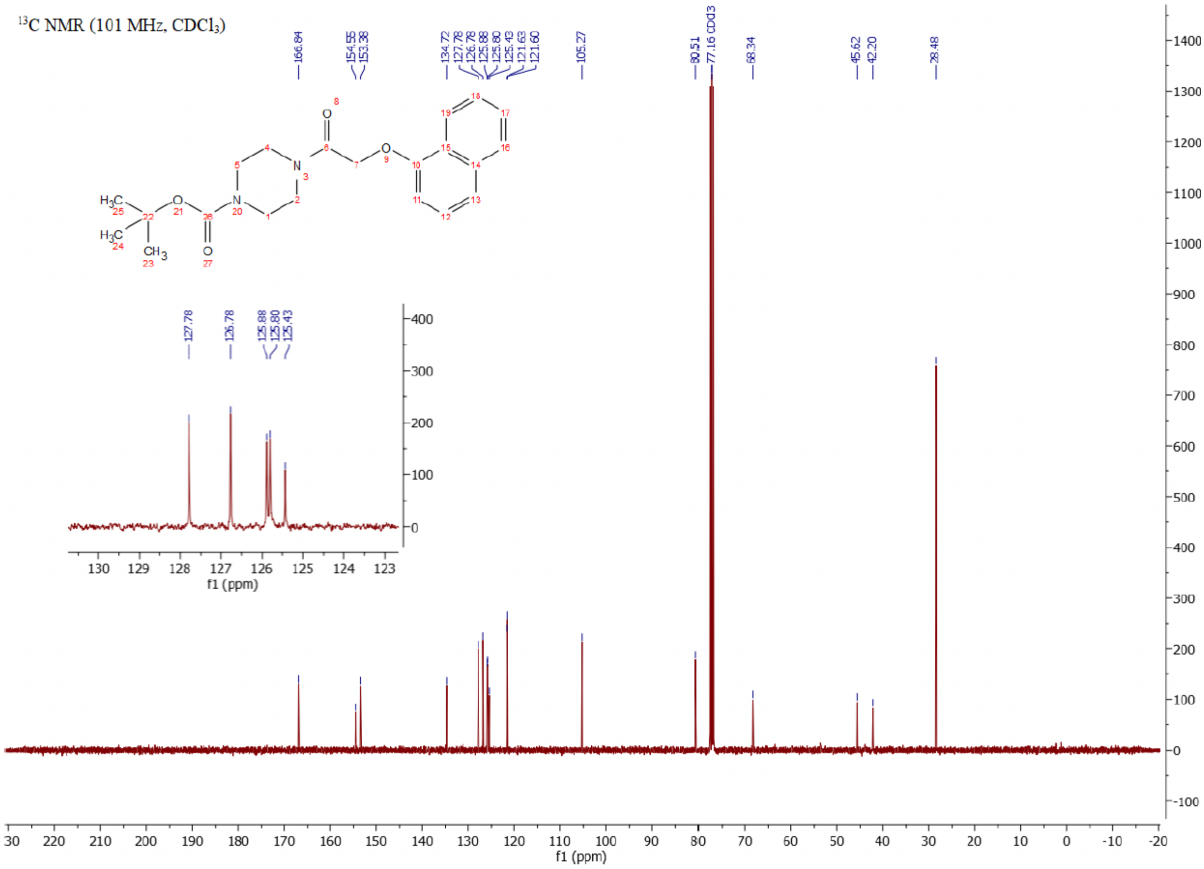

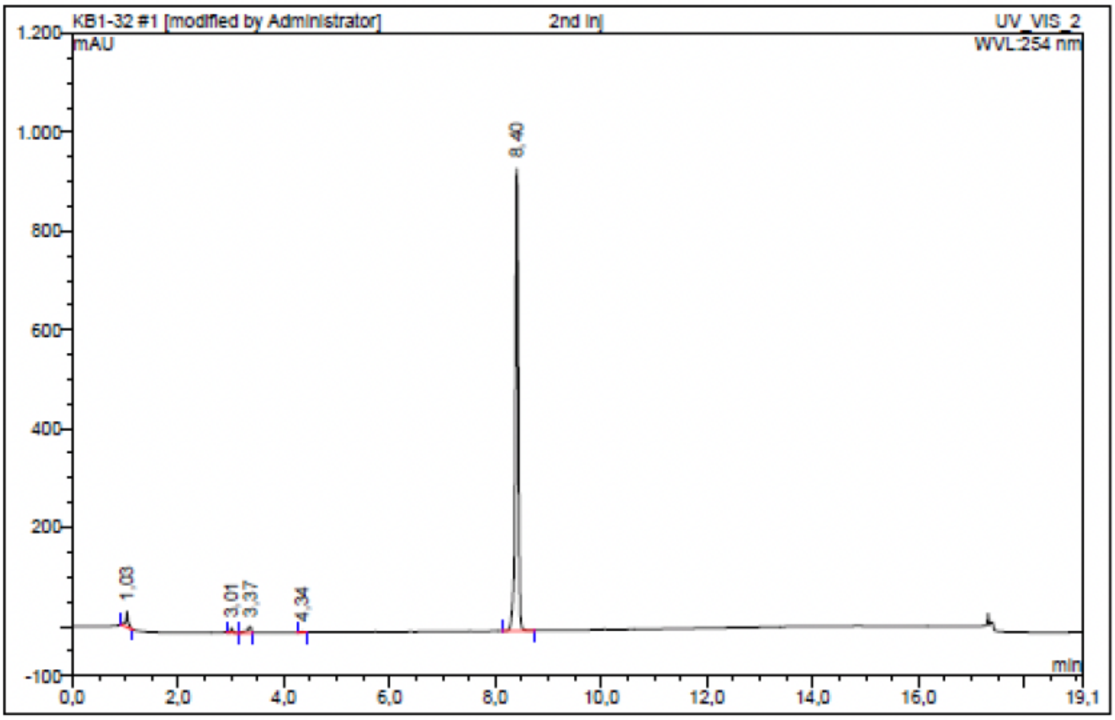

| No. | Ret.Time min | Peak Name | Height mAU | Area mAU*min | Rel.Area % | Amount | Resolution(EP) |
|---|---|---|---|---|---|---|---|
| 1 | 1,03 | n.a. | 31,209 | 1,631 | 2,50 | n.a. | 36,38 |
| 2 | 3,01 | n.a. | 9,529 | 0,378 | 0,58 | n.a. | 4,49 |
| 3 | 3,37 | n.a. | 12,575 | 1,181 | 1,81 | n.a. | 9,79 |
| 4 | 4,34 | n.a. | 1,202 | 0,073 | 0,11 | n.a. | 41,36 |
| 5 | 8,40 | n.a. | 935,184 | 61,993 | 95,00 | n.a. | n.a. |
| Total: | | | 989,700 | 65,256 | 100,00 | 0,000 | |

Compound 26 - TUG-2292

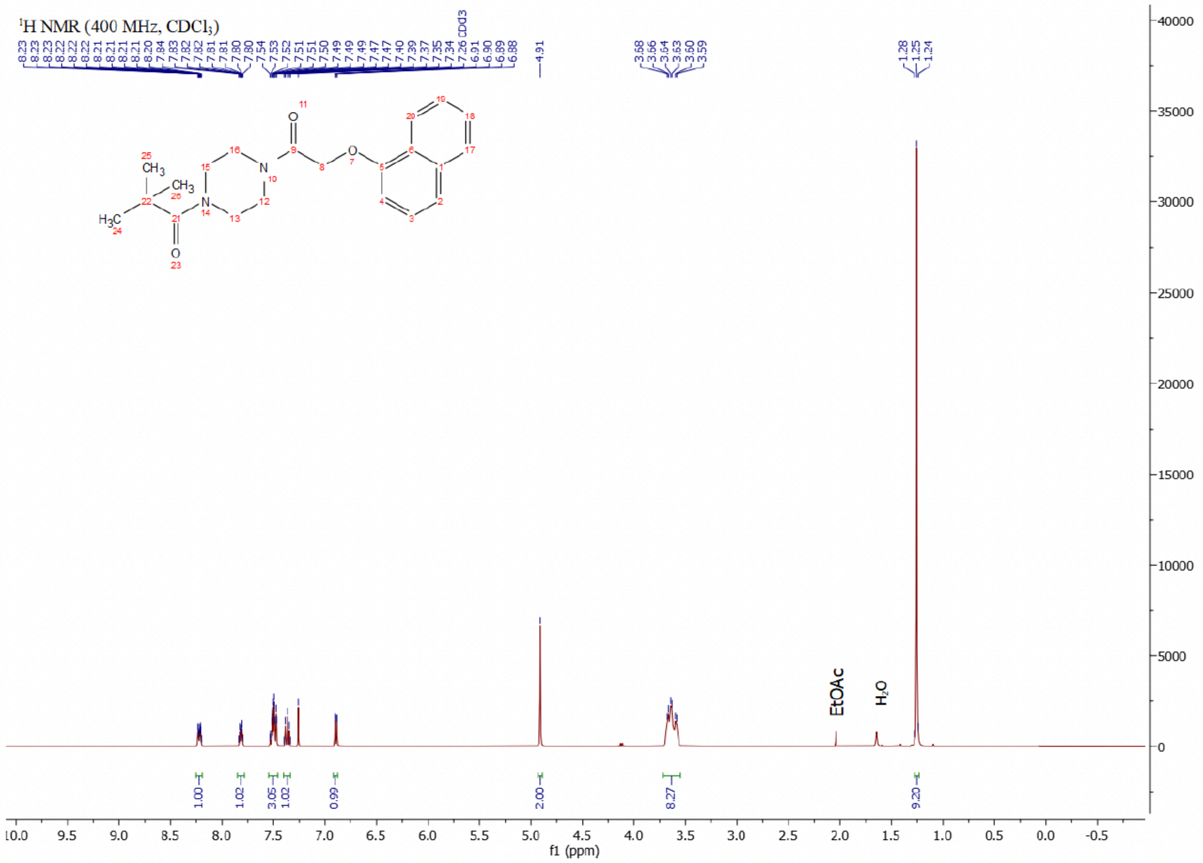

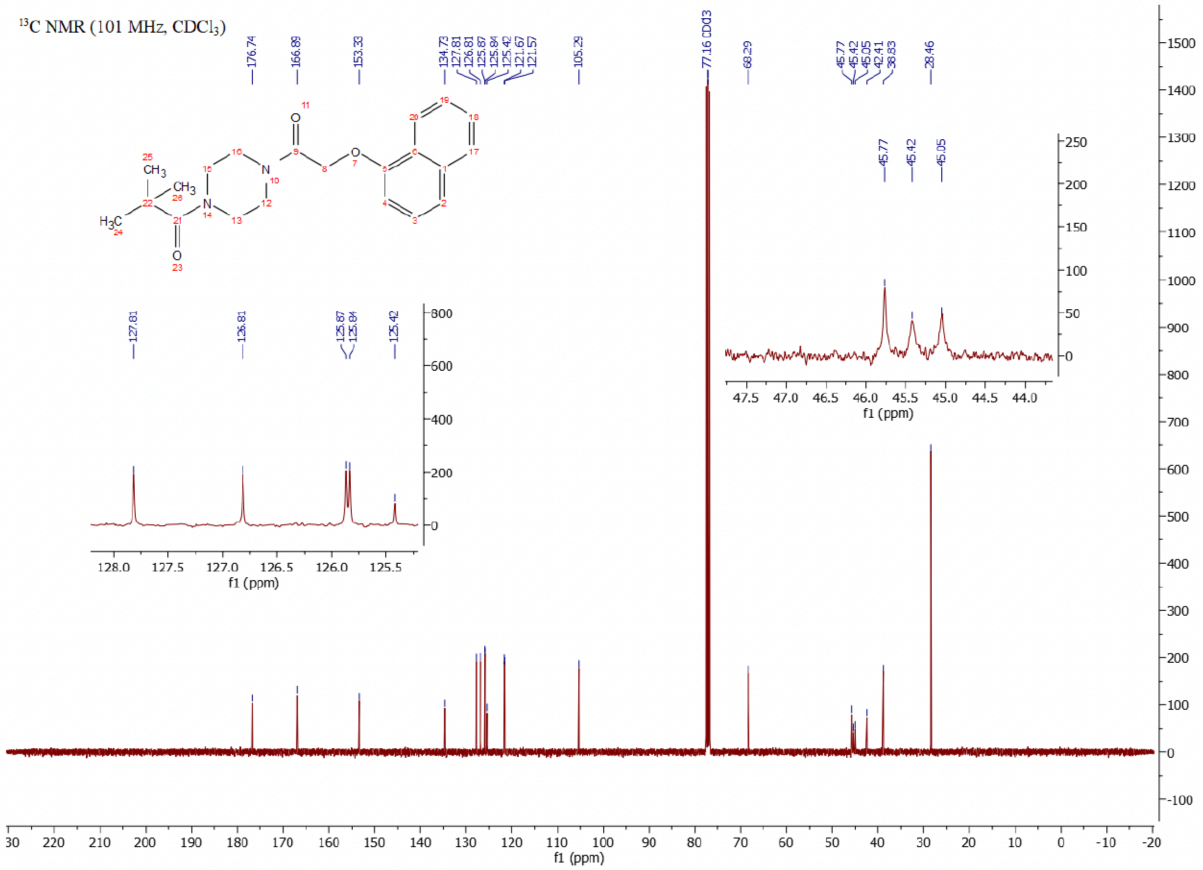

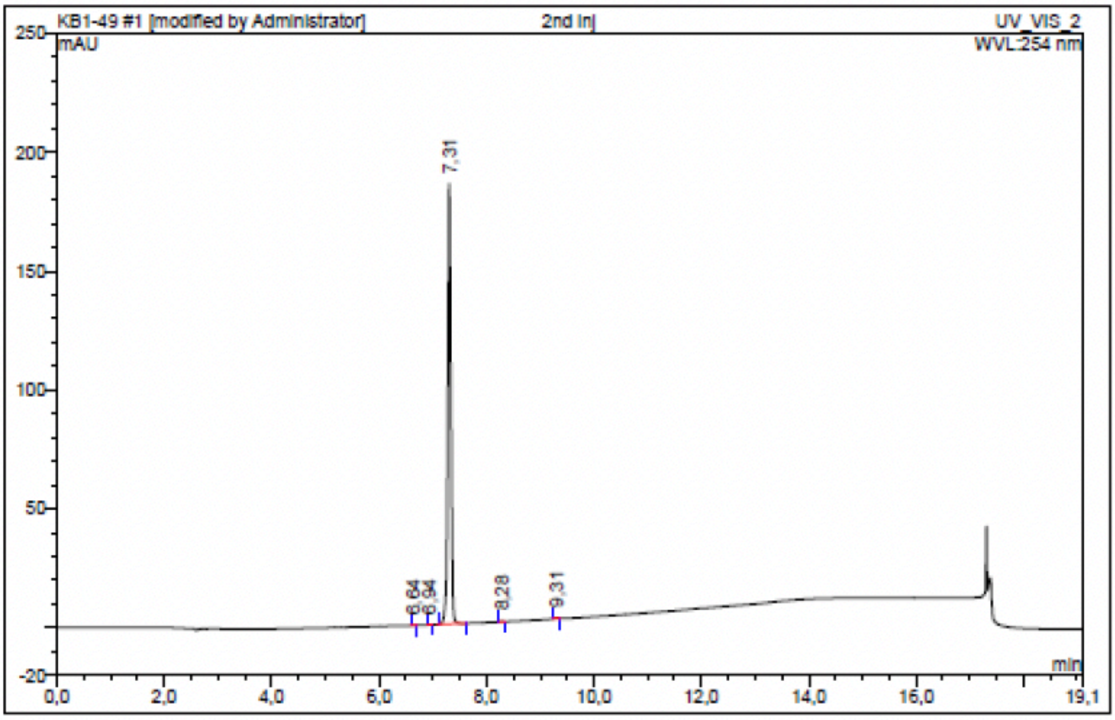

| No. | Ret.Time min | Peak Name | Height mAU | Area mAU*min | Rel.Area % | Amount | Resolution(EP) |
|---|---|---|---|---|---|---|---|
| 1 | 6,64 | n.a. | 0,038 | 0,002 | 0,02 | n.a. | 3,26 |
| 2 | 6,94 | n.a. | 0,128 | 0,007 | 0,05 | n.a. | 3,97 |
| 3 | 7,31 | n.a. | 184,968 | 12,410 | 99,64 | n.a. | 10,50 |
| 4 | 8,28 | n.a. | 0,190 | 0,010 | 0,08 | n.a. | 11,34 |
| 5 | 9,31 | n.a. | 0,445 | 0,027 | 0,21 | n.a. | n.a. |
| Total: | | | 185,769 | 12,455 | 100,00 | 0,000 | |

Compound 27 - TUG-2294

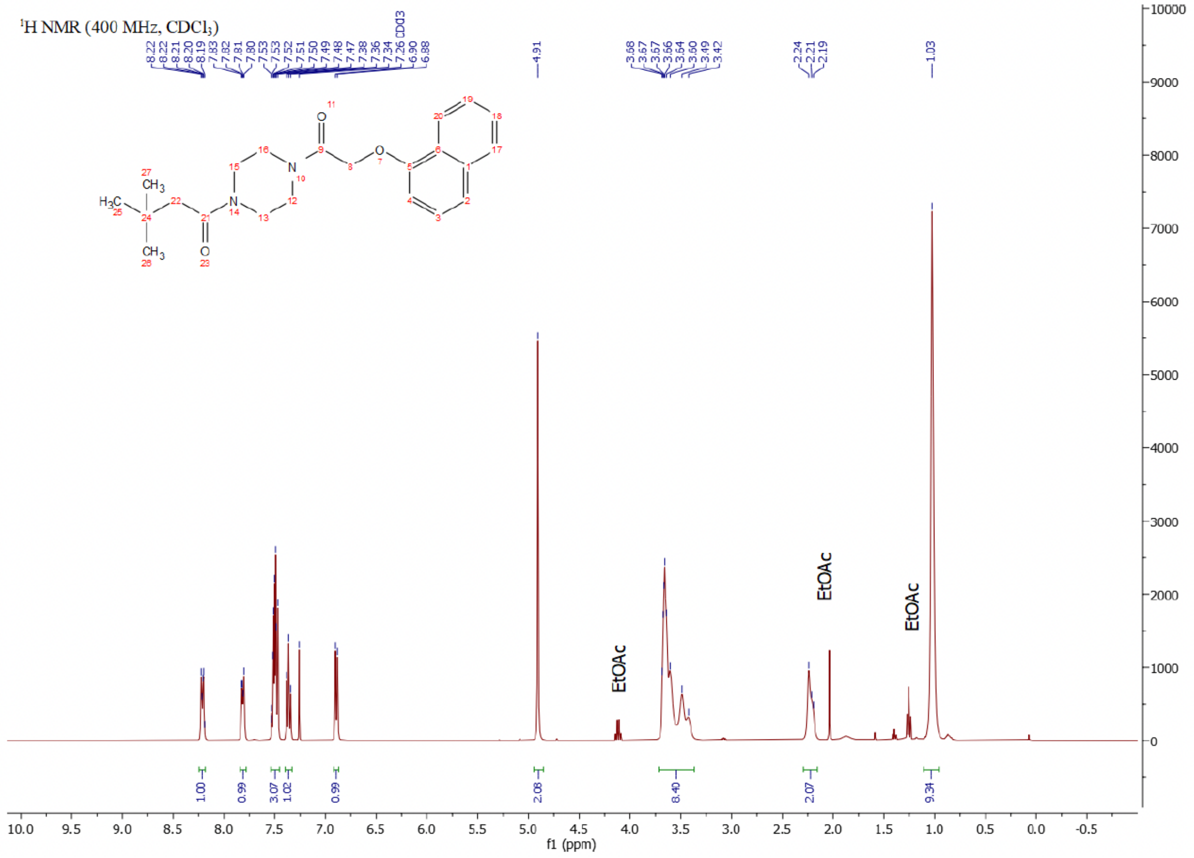

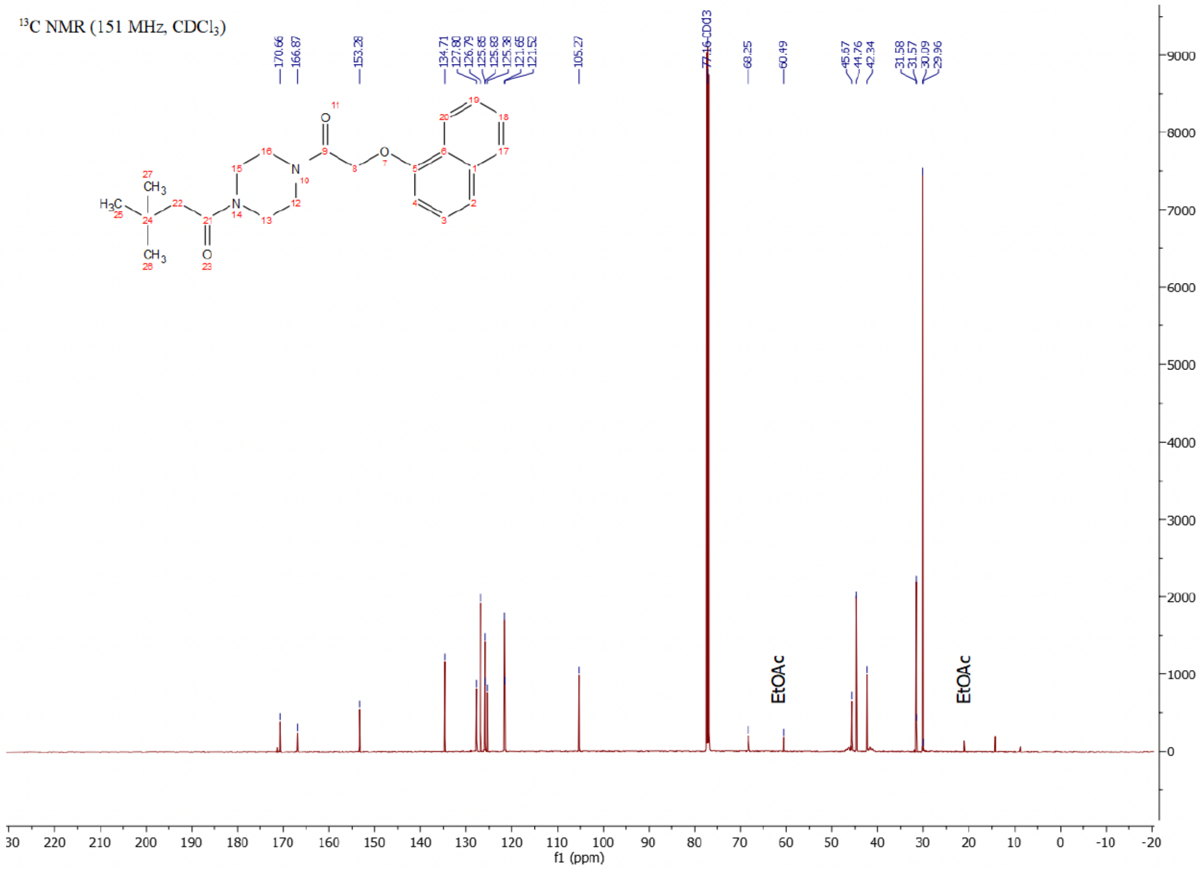

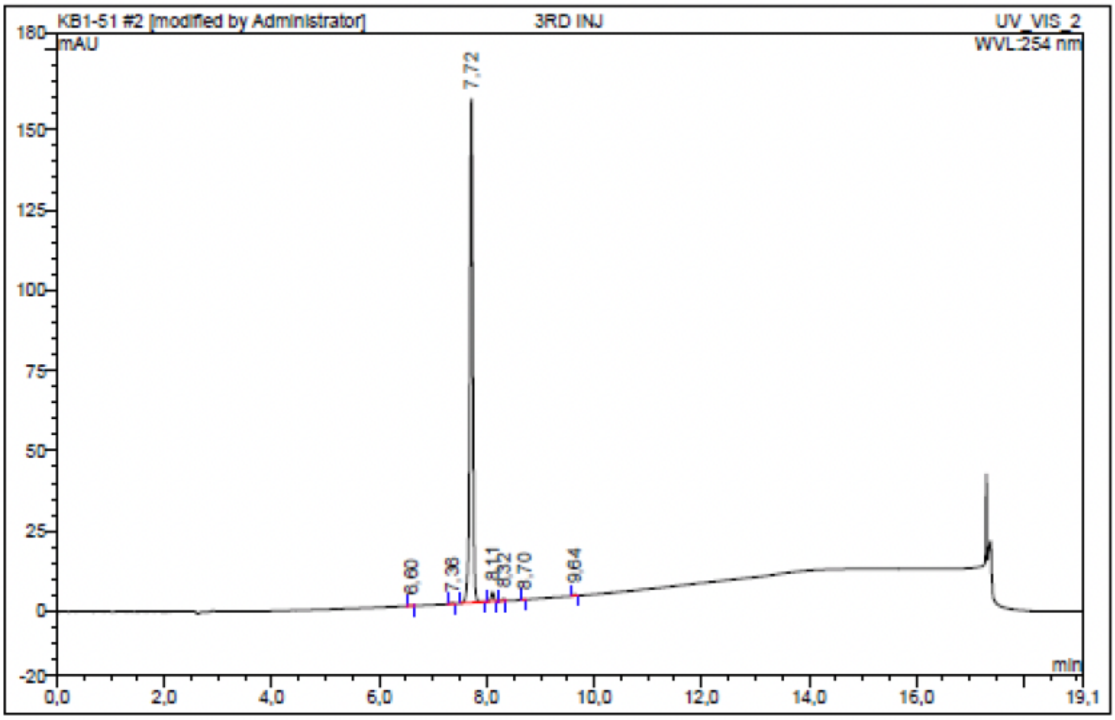

| No. | Ret.Time min | Peak Name | Height mAU | Area mAU*min | Rel.Area % | Amount | Resolution(EP) |
|---|---|---|---|---|---|---|---|
| 1 | 6,60 | n.a. | 0,091 | 0,010 | 0,09 | n.a. | 5,88 |
| 2 | 7,36 | n.a. | 0,339 | 0,020 | 0,18 | n.a. | 3,74 |
| 3 | 7,72 | n.a. | 156,781 | 10,343 | 97,56 | n.a. | 4,22 |
| 4 | 8,11 | n.a. | 2,941 | 0,171 | 1,61 | n.a. | 2,30 |
| 5 | 8,32 | n.a. | 0,635 | 0,036 | 0,34 | n.a. | 4,35 |
| 6 | 8,70 | n.a. | 0,038 | 0,002 | 0,02 | n.a. | 10,70 |
| 7 | 9,64 | n.a. | 0,361 | 0,020 | 0,19 | n.a. | n.a. |
| Total: | | | 161,186 | 10,601 | 100,00 | 0,000 | |

Compound 28 - TUG-2297

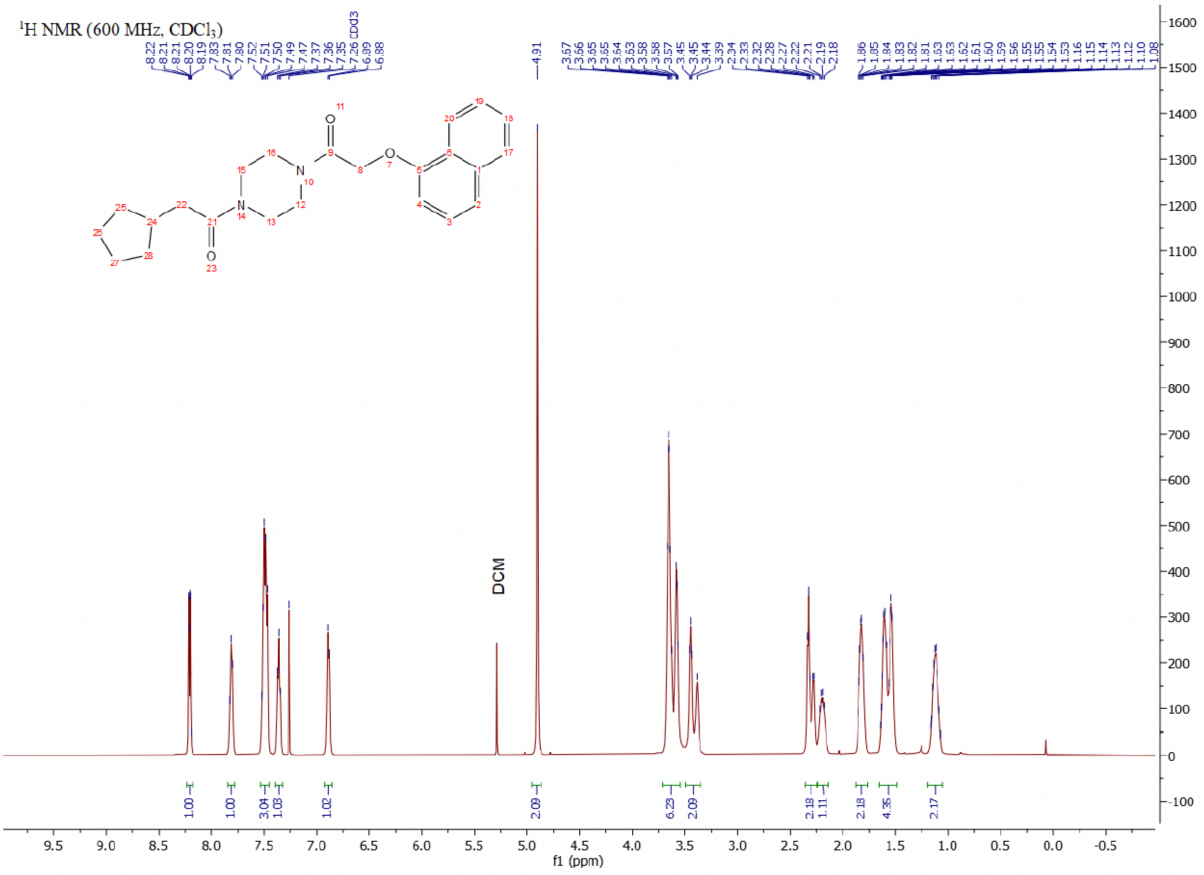

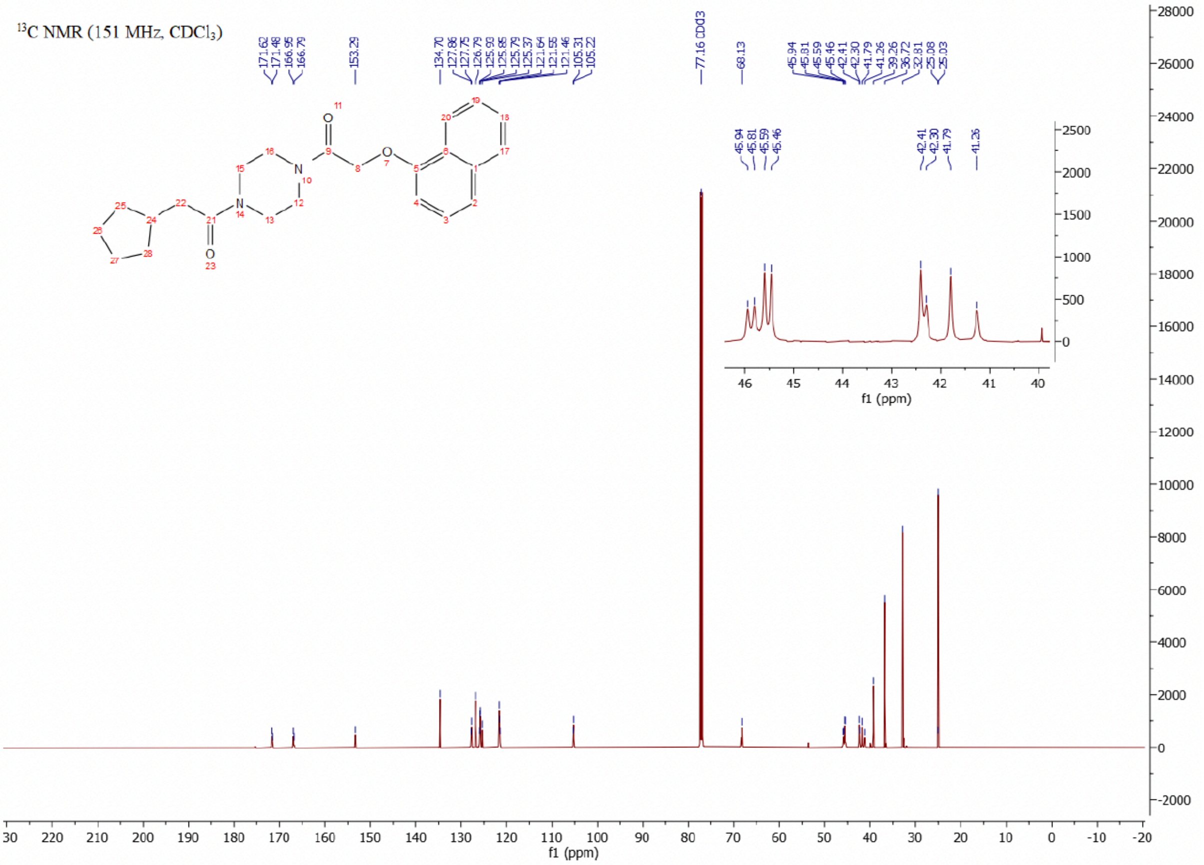

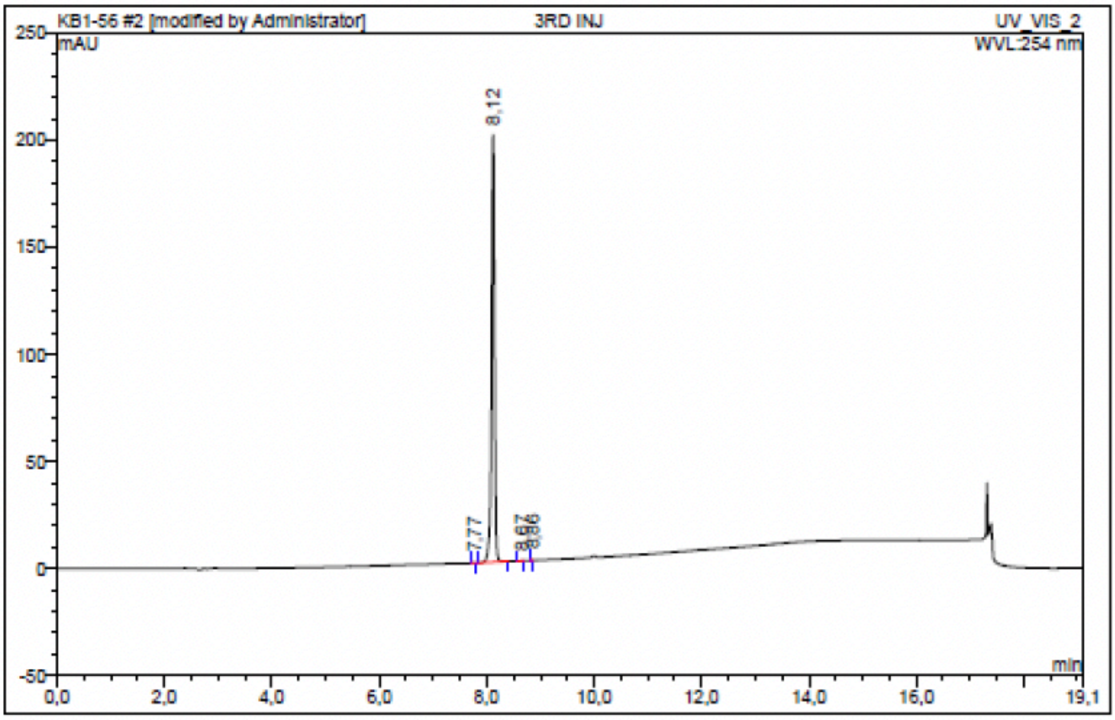

| No. | Ret.Time min | Peak Name | Height mAU | Area mAU*min | Rel.Area % | Amount | Resolution(EP) |
|---|---|---|---|---|---|---|---|
| 1 | 7,77 | n.a. | 0,126 | 0,005 | 0,04 | n.a. | 4,07 |
| 2 | 8,12 | n.a. | 199,399 | 13,373 | 99,93 | n.a. | 6,32 |
| 3 | 8,67 | n.a. | 0,057 | 0,002 | 0,02 | n.a. | 2,88 |
| 4 | 8,86 | n.a. | 0,041 | 0,002 | 0,01 | n.a. | n.a. |
| Total: | | | 199,623 | 13,383 | 100,00 | 0,000 | |

Compound 29 - TUG-2258

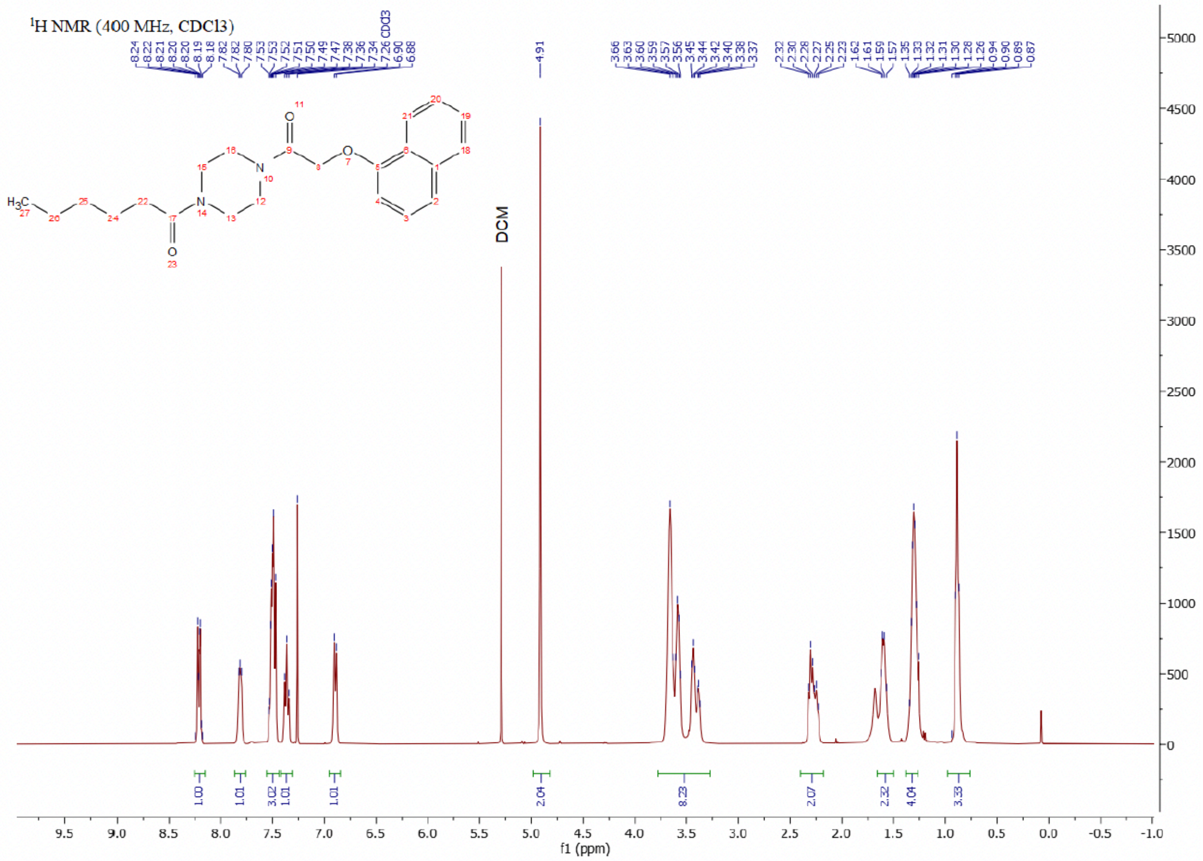

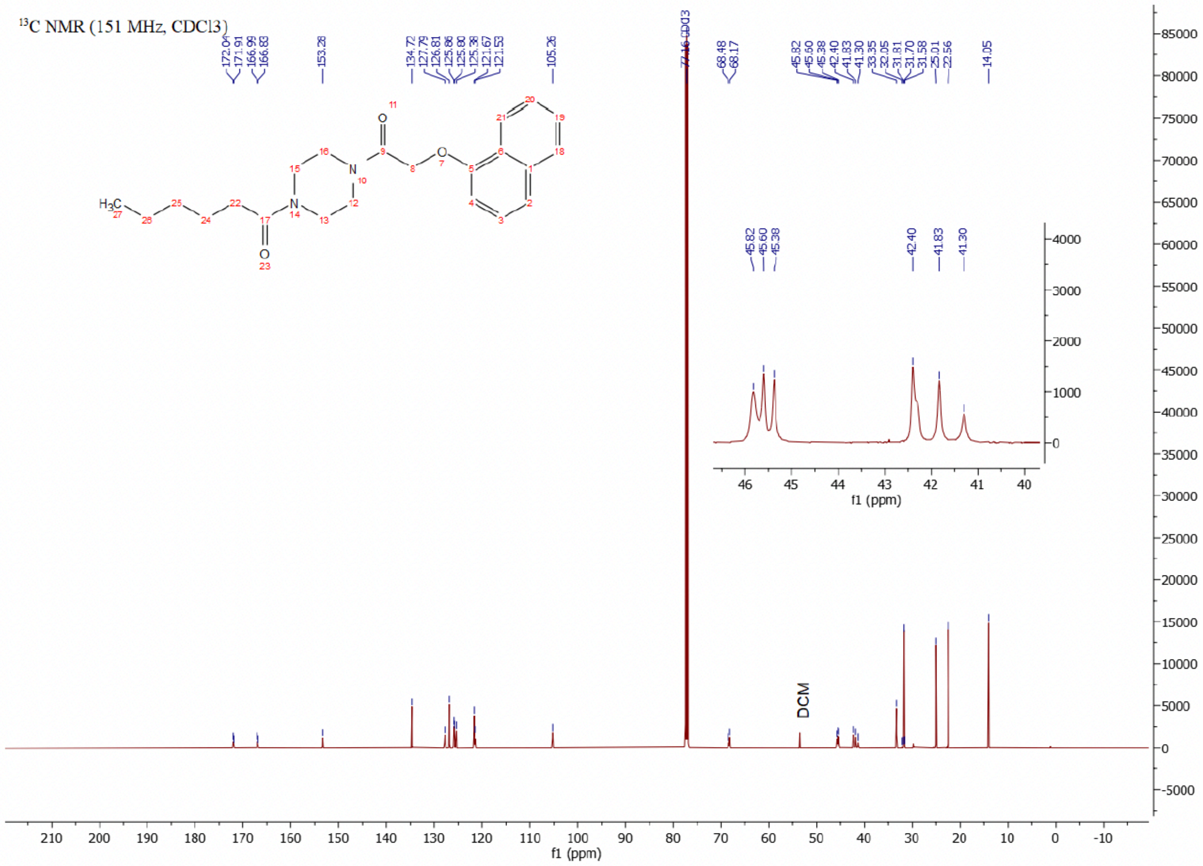

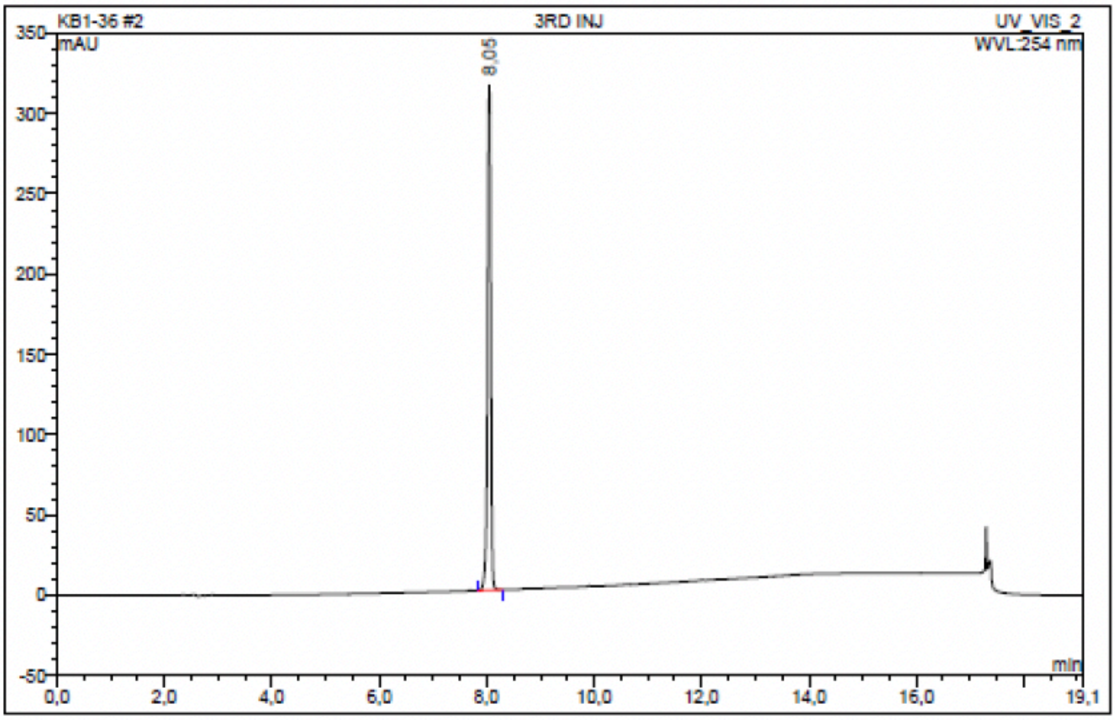

| No. | Ret.Time min | Peak Name | Height mAU | Area mAU*min | Rel.Area % | Amount | Resolution(EP) |
|---|---|---|---|---|---|---|---|
| 1 | 8,05 | n.a. | 314,315 | 20,498 | 100,00 | n.a. | n.a. |
| Total: | | | 314,315 | 20,498 | 100,00 | 0,000 | |

### Compound 30 - TUG-2709

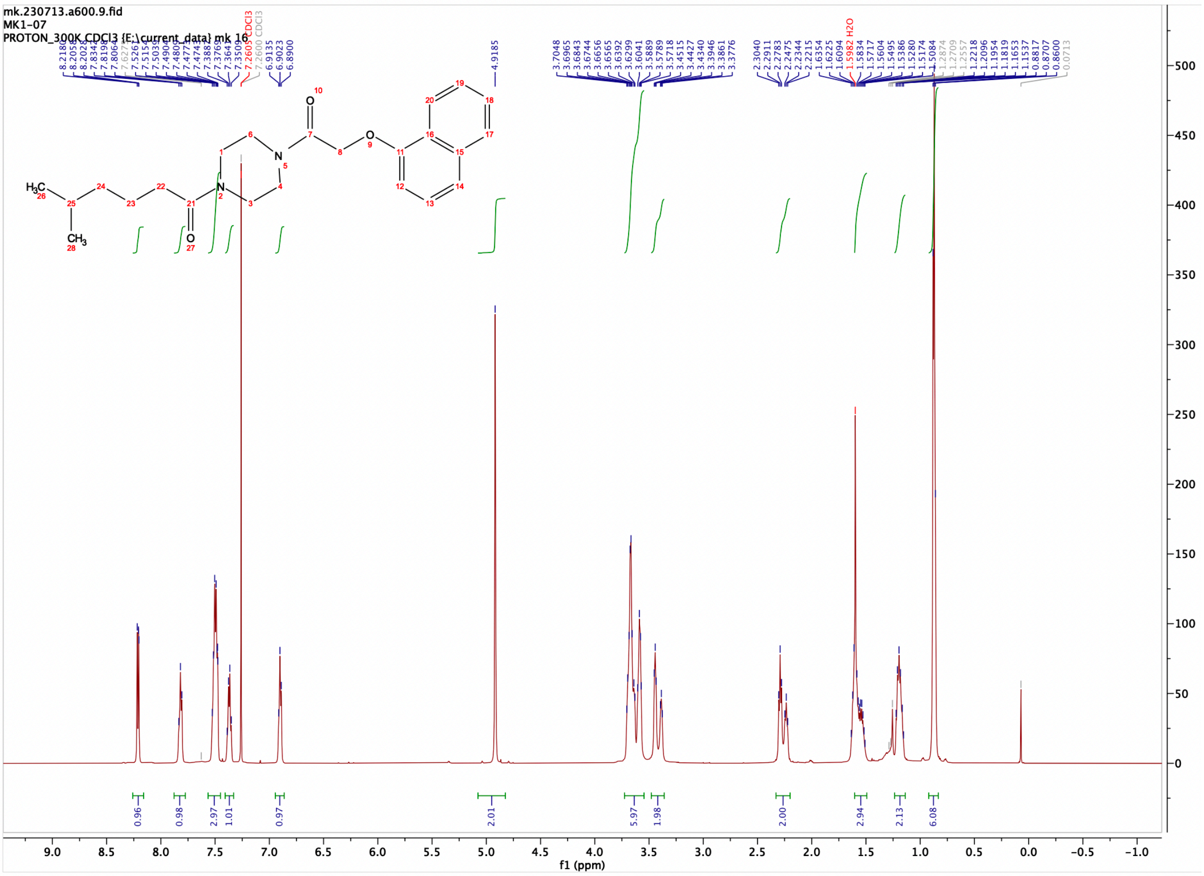

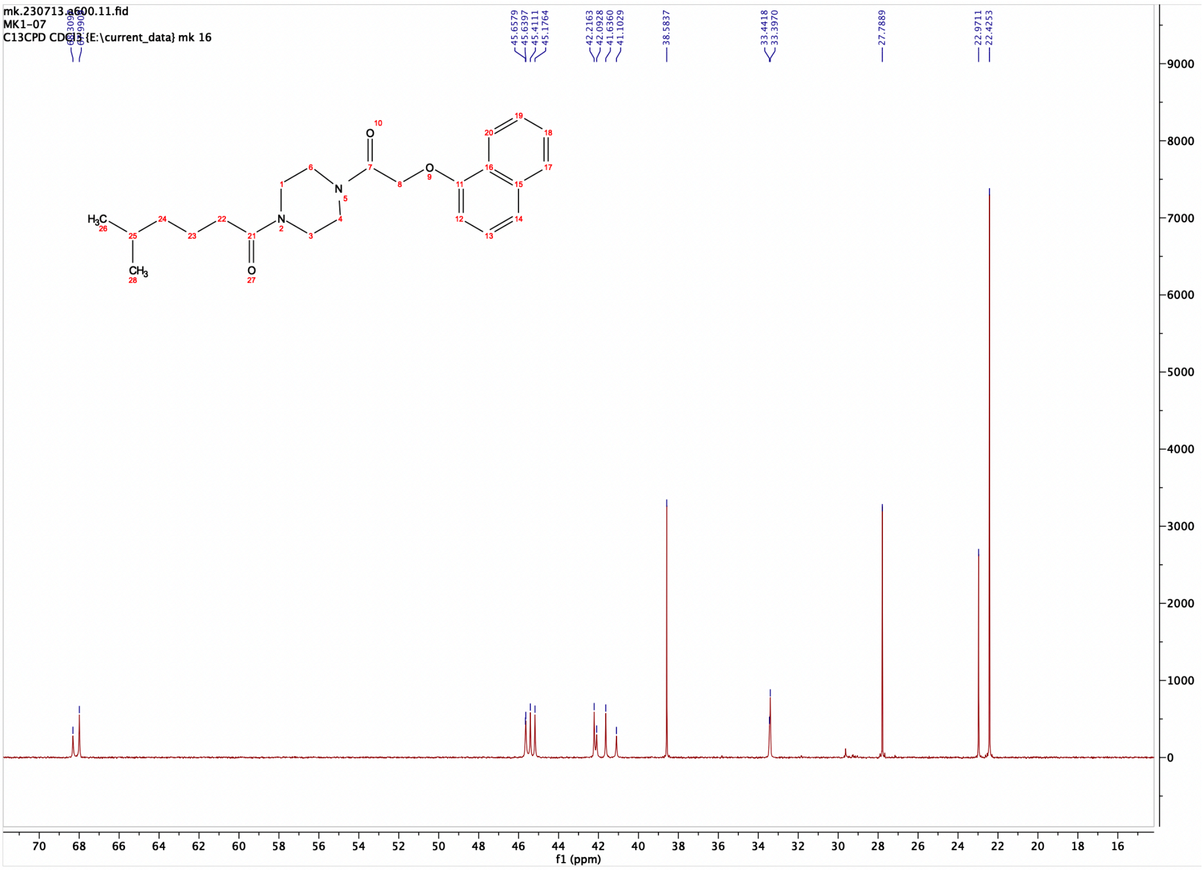

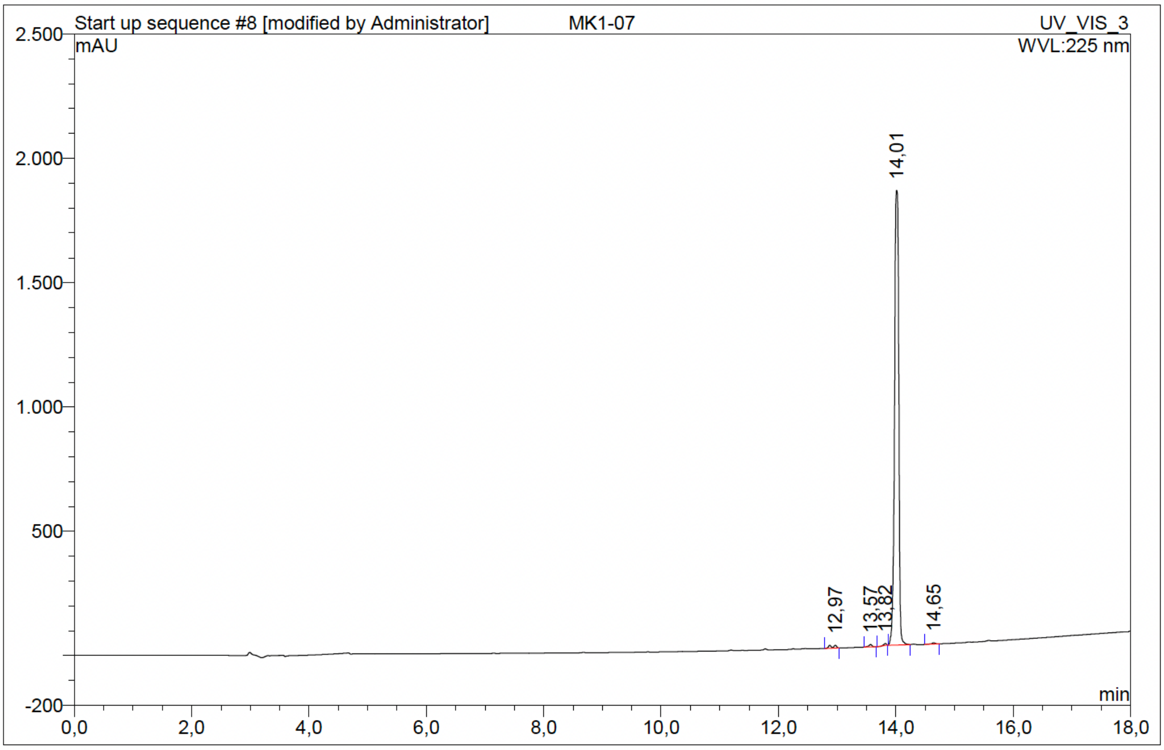

| No. | Ret.Time min | Peak Name | Height mAU | Area mAU*min | Rel.Area % | Amount | Resolution(EP) |
|---|---|---|---|---|---|---|---|
| 1 | 12,97 | n.a. | 11,765 | 1,279 | 0,81 | n.a. | 3,50 |
| 2 | 13,57 | n.a. | 9,449 | 0,626 | 0,39 | n.a. | 2,79 |
| 3 | 13,82 | n.a. | 7,400 | 0,328 | 0,21 | n.a. | 1,72 |
| 4 | 14,01 | n.a. | 1827,790 | 155,916 | 98,31 | n.a. | 5,28 |
| 5 | 14,65 | n.a. | 5,412 | 0,445 | 0,28 | n.a. | n.a. |
| Total: | | | 1861,816 | 158,594 | 100,00 | 0,000 | |

Compound 31 - TUG-2374

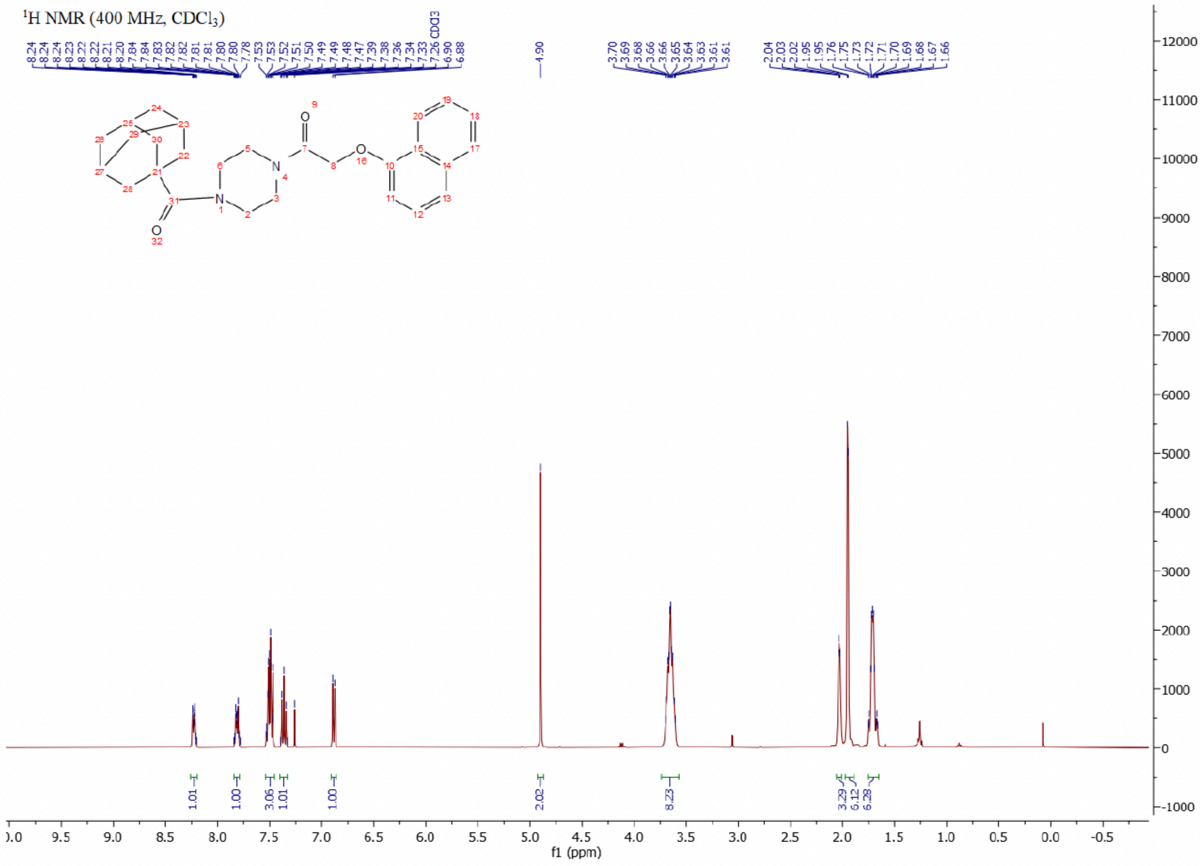

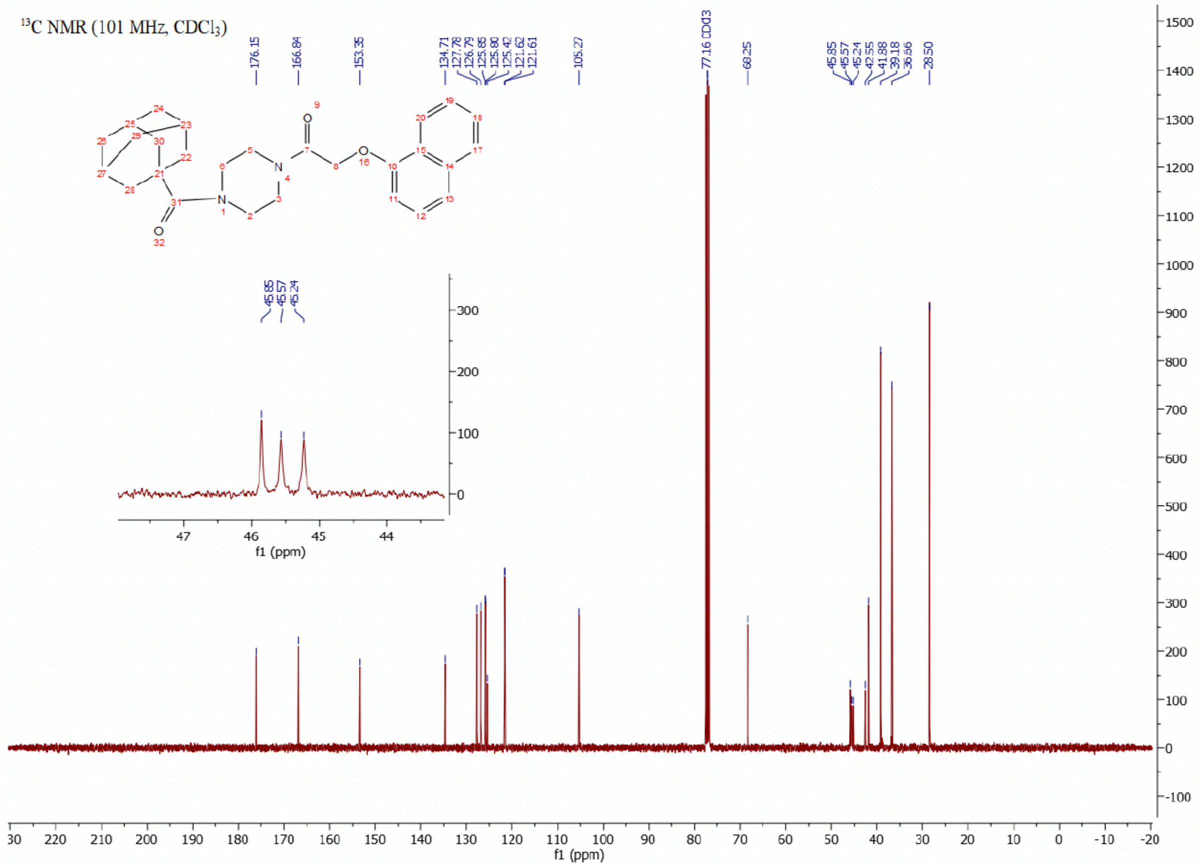

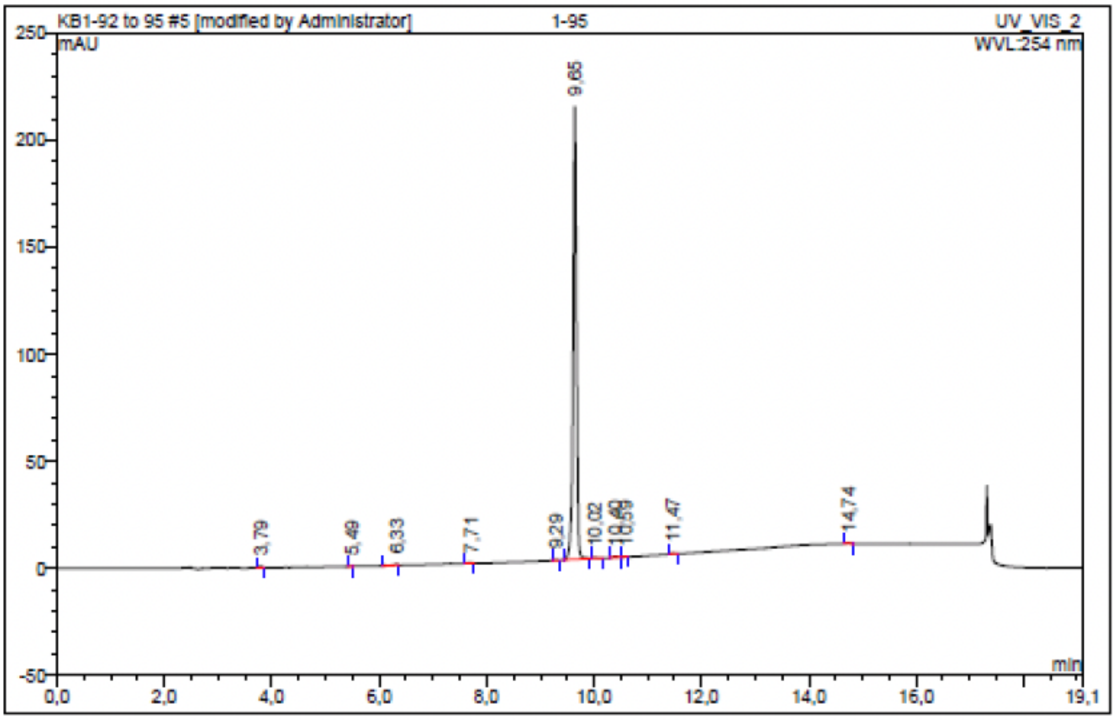

| No. | Ret.Time min | Peak Name | Height mAU | Area mAU*min | Rel.Area % | Amount | Resolution(EP) |
|---|---|---|---|---|---|---|---|
| 1 | 3,79 | n.a. | 0,763 | 0,044 | 0,28 | n.a. | 20,02 |
| 2 | 5,49 | n.a. | 0,128 | 0,006 | 0,04 | n.a. | 3,36 |
| 3 | 6,33 | n.a. | 0,460 | 0,042 | 0,27 | n.a. | 4,50 |
| 4 | 7,71 | n.a. | 0,409 | 0,042 | 0,27 | n.a. | 11,09 |
| 5 | 9,29 | n.a. | 0,212 | 0,013 | 0,08 | n.a. | 3,51 |
| 6 | 9,65 | n.a. | 211,479 | 15,272 | 98,27 | n.a. | 3,81 |
| 7 | 10,02 | n.a. | 0,266 | 0,014 | 0,09 | n.a. | 3,77 |
| 8 | 10,40 | n.a. | 0,641 | 0,048 | 0,31 | n.a. | 1,98 |
| 9 | 10,59 | n.a. | 0,186 | 0,010 | 0,06 | n.a. | 9,51 |
| 10 | 11,47 | n.a. | 0,319 | 0,018 | 0,12 | n.a. | 28,09 |
| 11 | 14,74 | n.a. | 0,394 | 0,033 | 0,22 | n.a. | n.a. |
| Total: | | | 215,256 | 15,541 | 100,00 | 0,000 | |

Compound 32 - TUG-2608

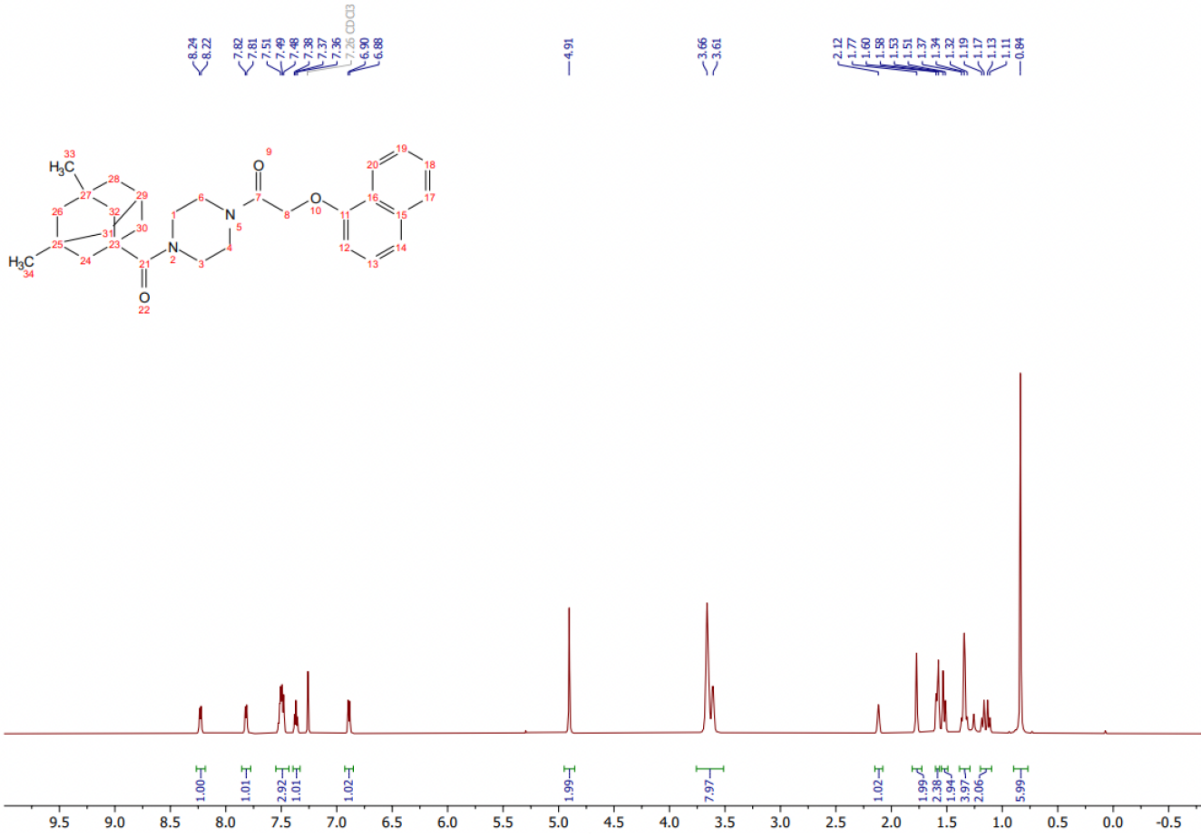

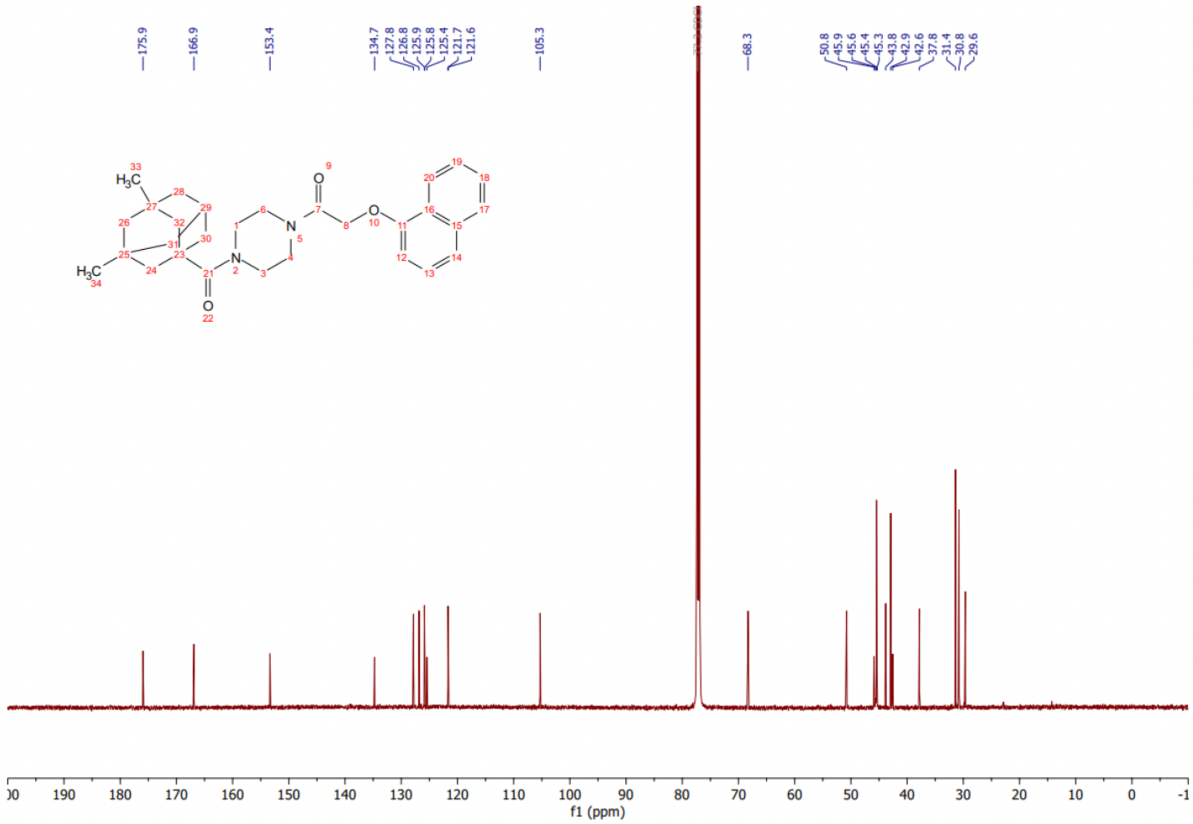

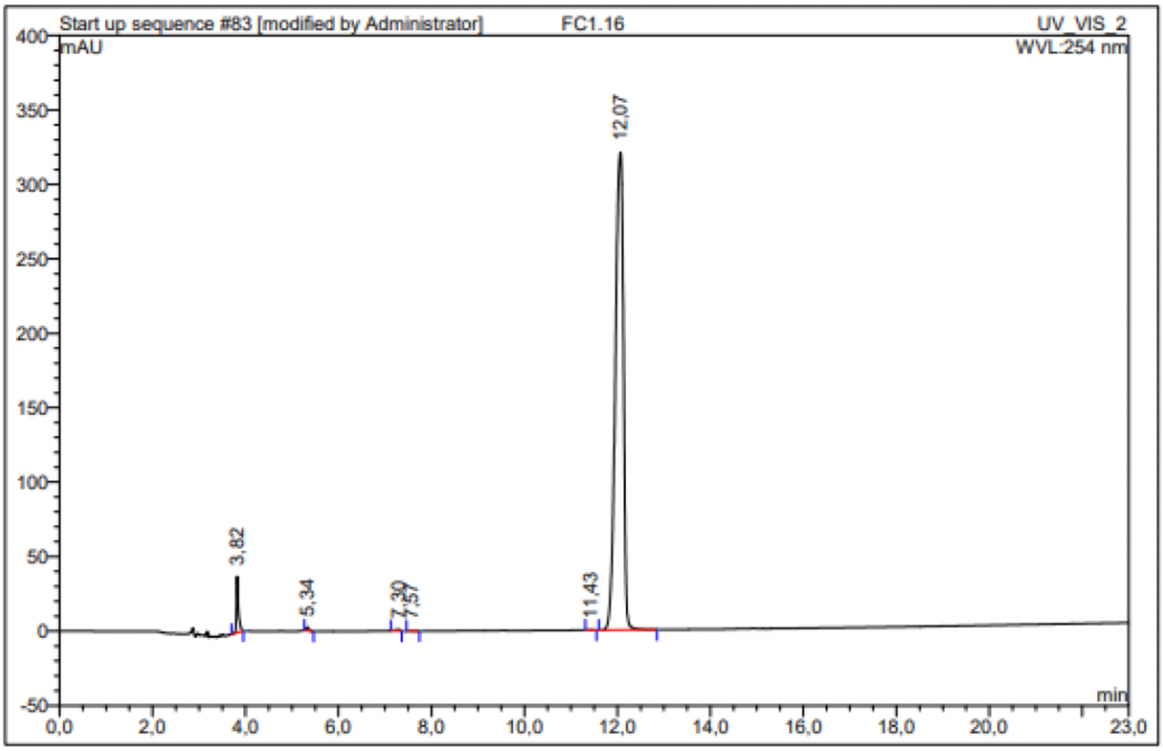

| No. | Ret.Time min | Peak Name | Height mAU | Area mAU*min | Rel.Area % | Amount | Resolution(EP) |
|---|---|---|---|---|---|---|---|
| 1 | 3,82 | n.a. | 37,765 | 1,940 | 2,87 | n.a. | 16,79 |
| 2 | 5,34 | n.a. | 2,346 | 0,159 | 0,24 | n.a. | 11,13 |
| 3 | 7,30 | n.a. | 0,909 | 0,118 | 0,17 | n.a. | 1,28 |
| 4 | 7,57 | n.a. | 0,153 | 0,011 | 0,02 | n.a. | 17,96 |
| 5 | 11,43 | n.a. | 0,158 | 0,022 | 0,03 | n.a. | 2,21 |
| 6 | 12,07 | n.a. | 320,739 | 65,365 | 96,67 | n.a. | n.a. |
| **Total:** | | | 362,070 | 67,616 | 100,00 | 0,000 | |

Compound 33 - TUG-2840

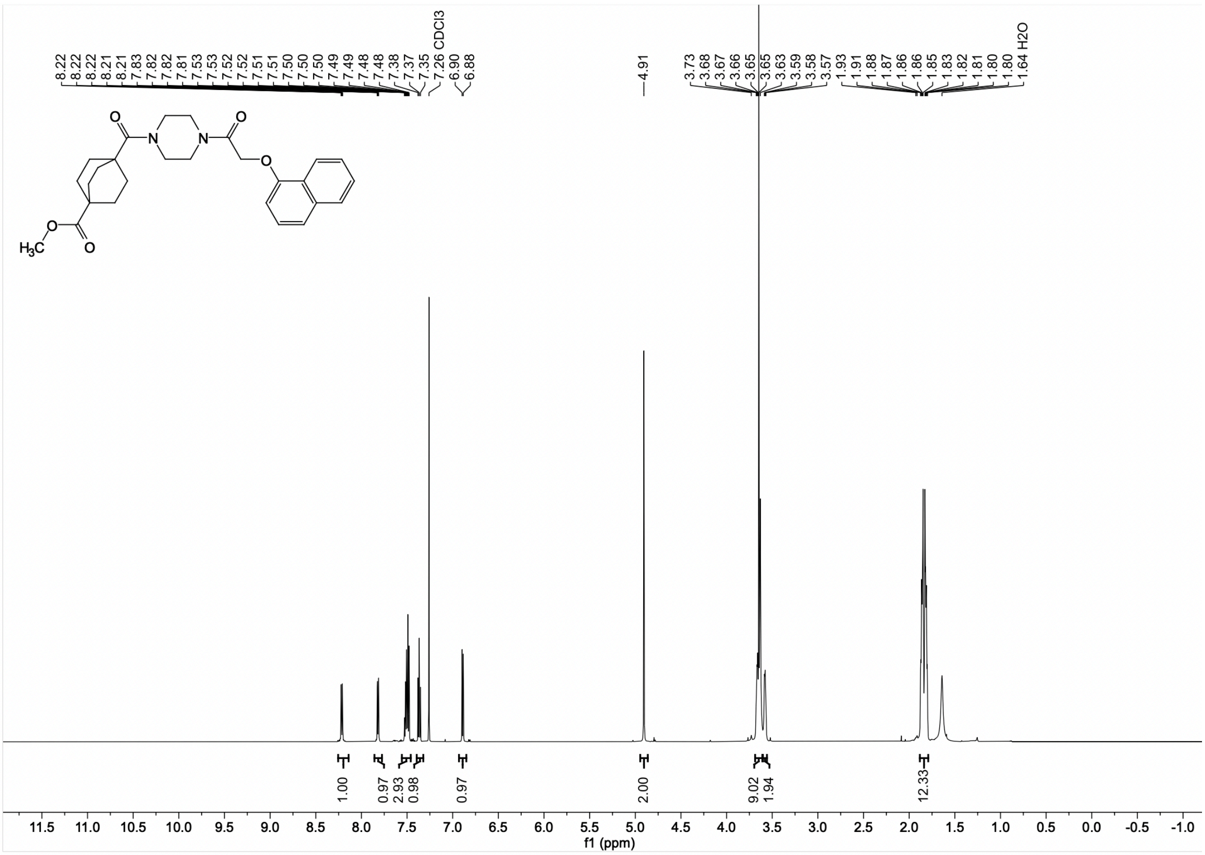

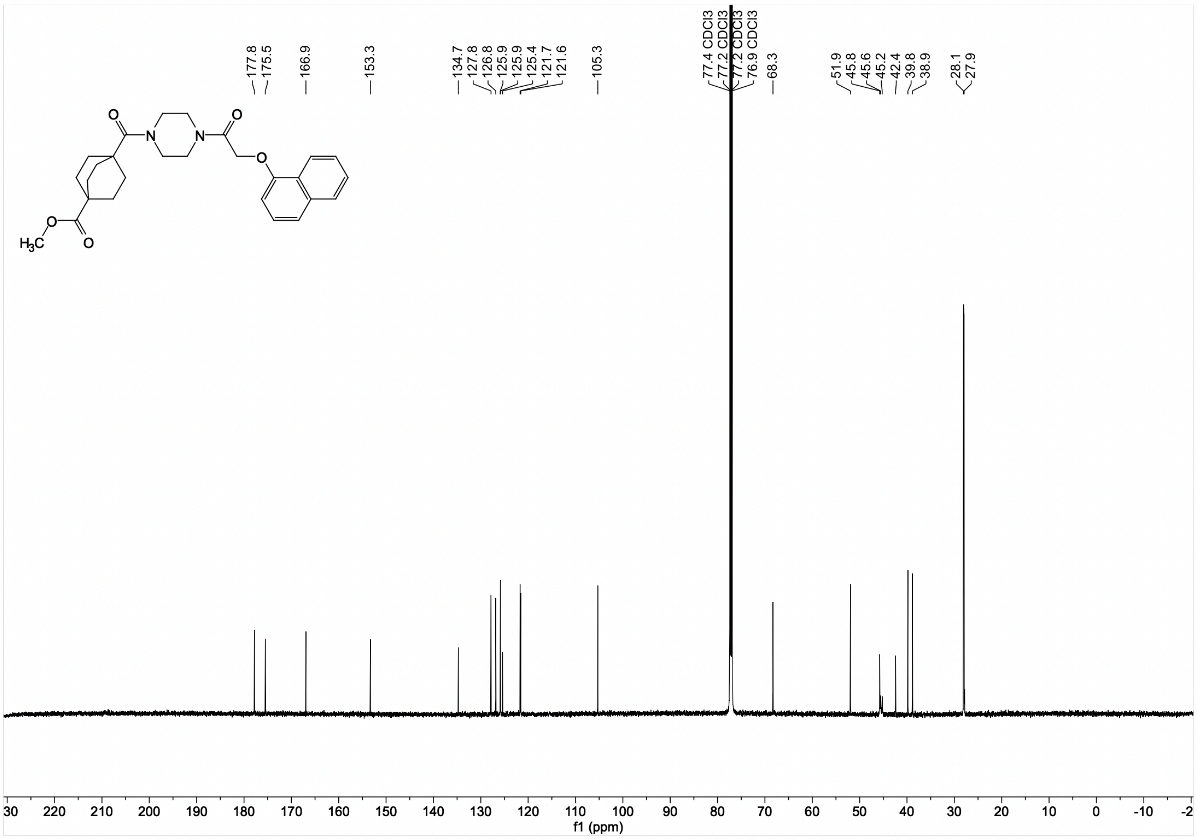

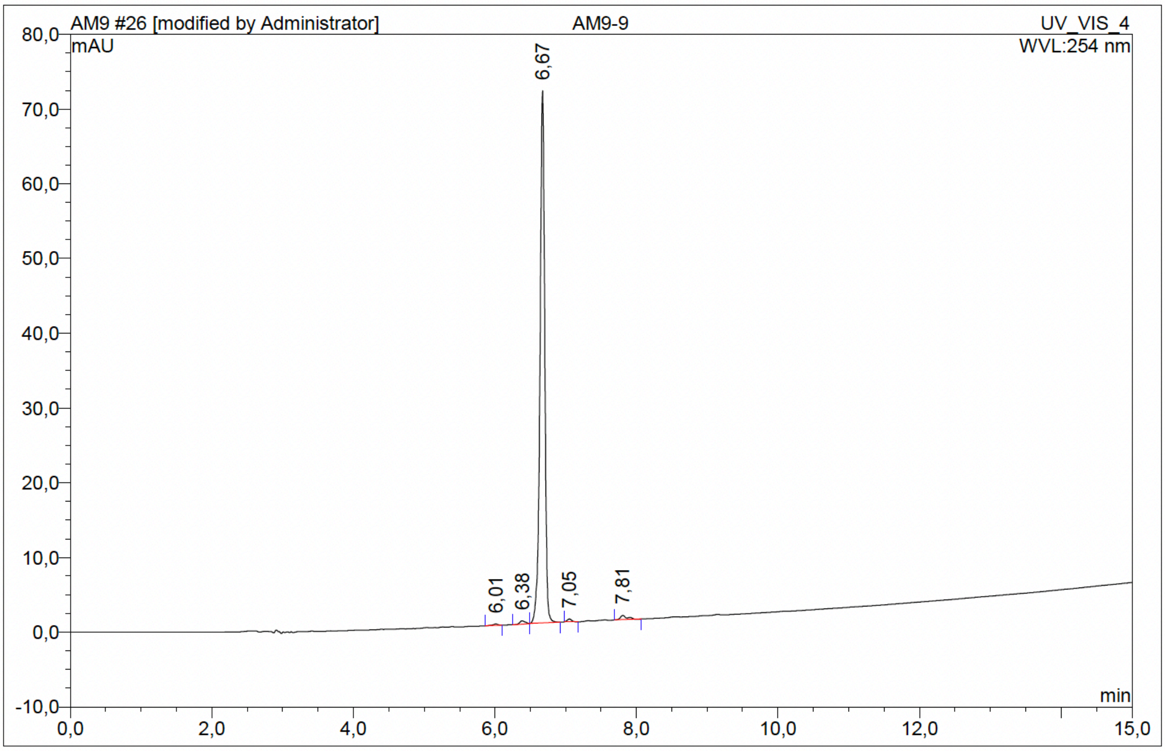

| No. | Ret.Time min | Peak Name | Height mAU | Area mAU*min | Rel.Area % | Amount | Resolution(EP) |
|---|---|---|---|---|---|---|---|
| 1 | 6,01 | n.a. | 0,186 | 0,015 | 0,25 | n.a. | 3,02 |
| 2 | 6,38 | n.a. | 0,432 | 0,036 | 0,63 | n.a. | 2,25 |
| 3 | 6,67 | n.a. | 71,230 | 5,615 | 97,50 | n.a. | 3,17 |
| 4 | 7,05 | n.a. | 0,385 | 0,029 | 0,50 | n.a. | 6,13 |
| 5 | 7,81 | n.a. | 0,570 | 0,064 | 1,11 | n.a. | n.a. |
| Total: | | | 72,802 | 5,759 | 100,00 | 0,000 | |

Compound 34 - TUG-2842

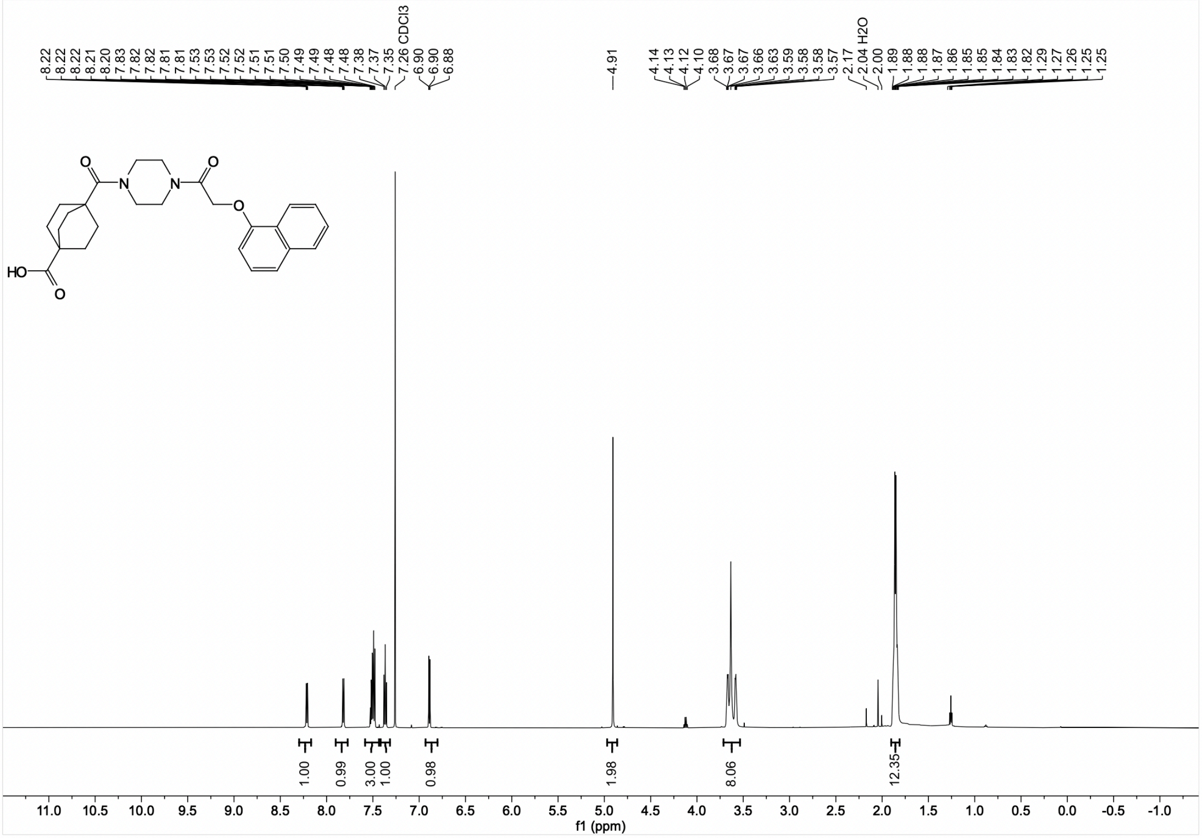

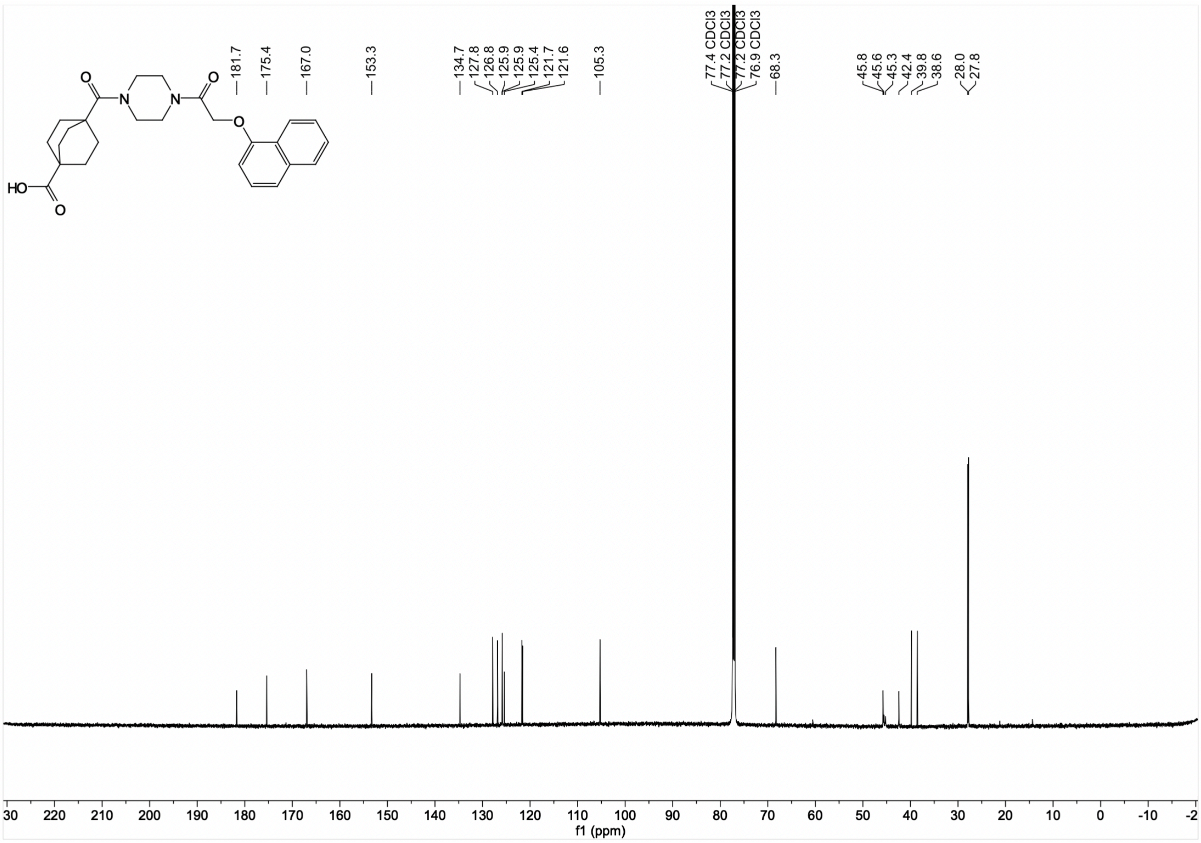

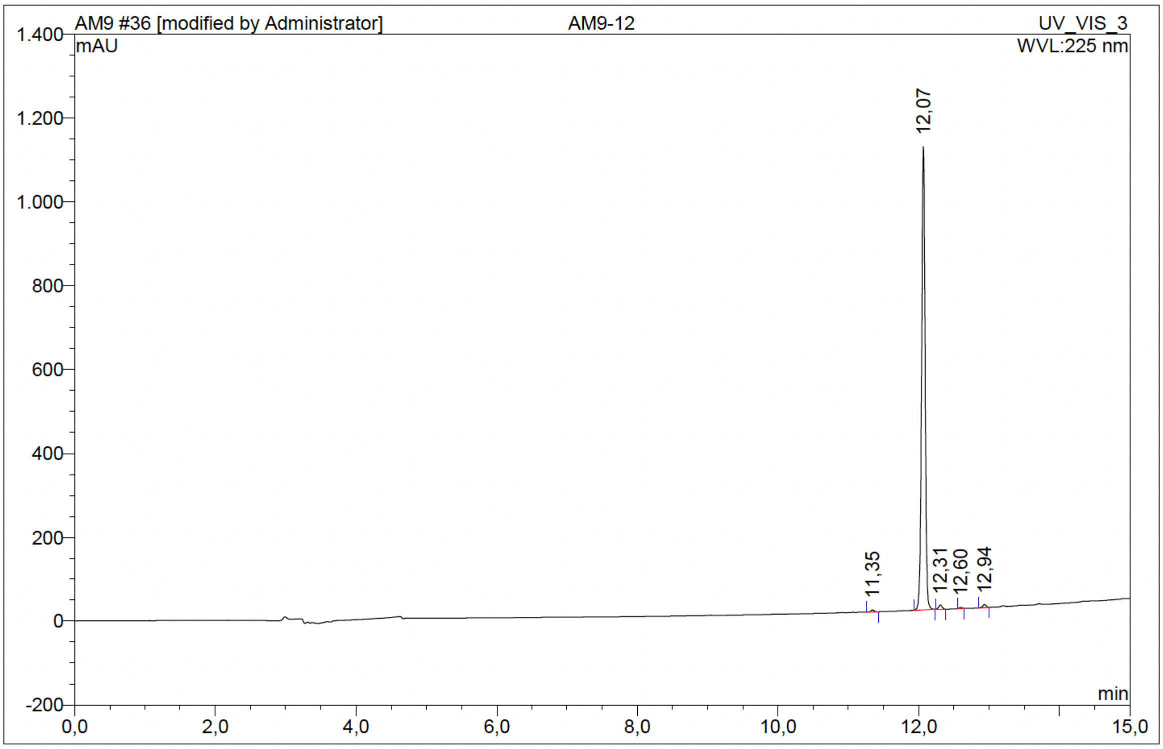

| No. | Ret.Time min | Peak Name | Height mAU | Area mAU*min | Rel.Area % | Amount | Resolution(EP) |
|---|---|---|---|---|---|---|---|
| 1 | 11,35 | n.a. | 4,670 | 0,251 | 0,38 | n.a. | 8,25 |
| 2 | 12,07 | n.a. | 1105,297 | 65,405 | 97,98 | n.a. | 2,66 |
| 3 | 12,31 | n.a. | 9,627 | 0,539 | 0,81 | n.a. | 3,40 |
| 4 | 12,60 | n.a. | 2,693 | 0,133 | 0,20 | n.a. | 3,93 |
| 5 | 12,94 | n.a. | 7,406 | 0,423 | 0,63 | n.a. | n.a. |
| Total: | | | 1129,693 | 66,752 | 100,00 | 0,000 | |

Compound 35 - TUG-2327

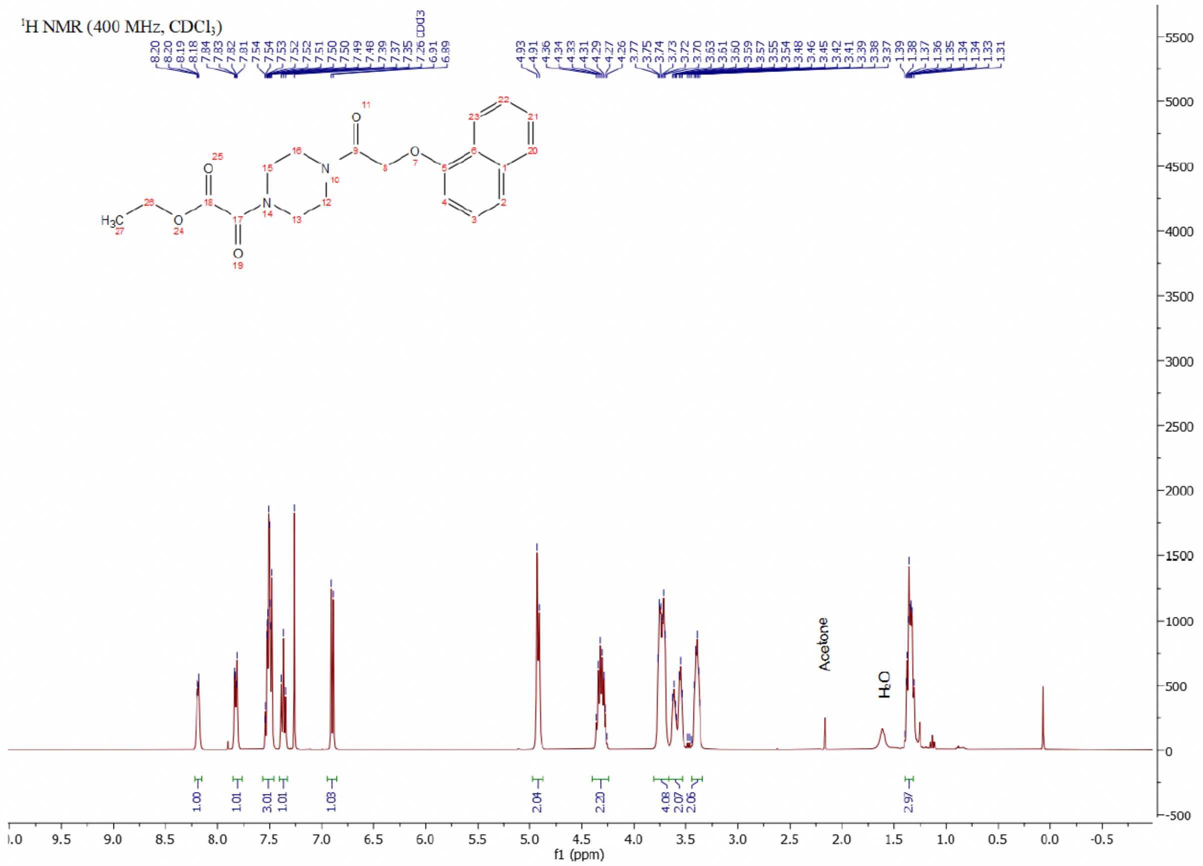

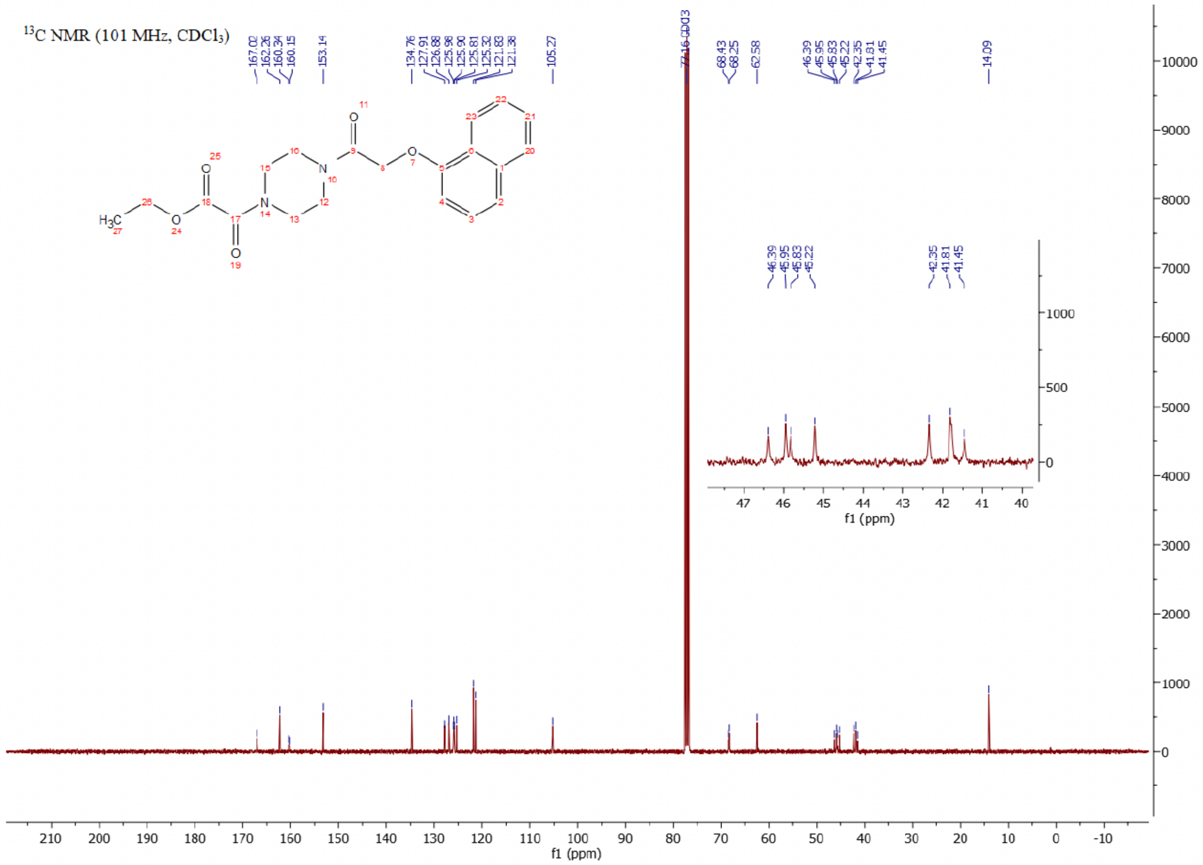

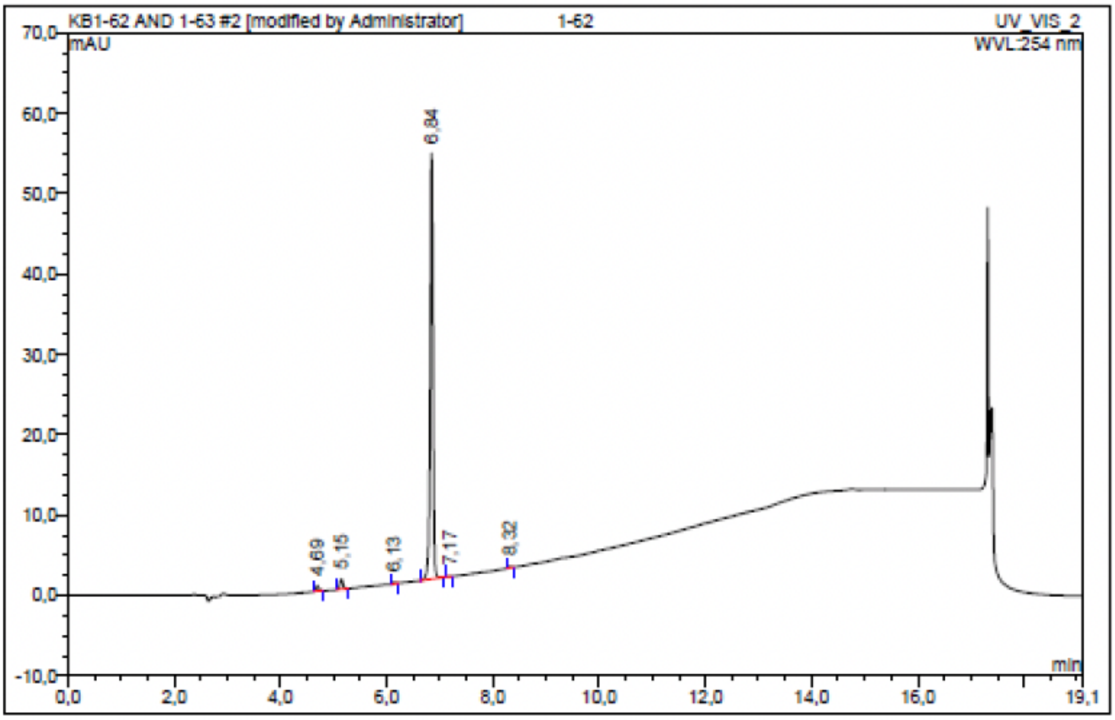

| No. | Ret.Time min | Peak Name | Height mAU | Area mAU*min | Rel.Area % | Amount | Resolution(EP) |
|---|---|---|---|---|---|---|---|
| 1 | 4,69 | n.a. | 0,645 | 0,043 | 1,25 | n.a. | 4,50 |
| 2 | 5,15 | n.a. | 1,242 | 0,080 | 2,31 | n.a. | 10,43 |
| 3 | 6,13 | n.a. | 0,183 | 0,010 | 0,29 | n.a. | 7,77 |
| 4 | 6,84 | n.a. | 52,964 | 3,320 | 95,86 | n.a. | 3,89 |
| 5 | 7,17 | n.a. | 0,037 | 0,002 | 0,05 | n.a. | 14,22 |
| 6 | 8,32 | n.a. | 0,162 | 0,008 | 0,24 | n.a. | n.a. |
| Total: | | | 55,231 | 3,464 | 100,00 | 0,000 | |

Compound 36 - TUG-2328

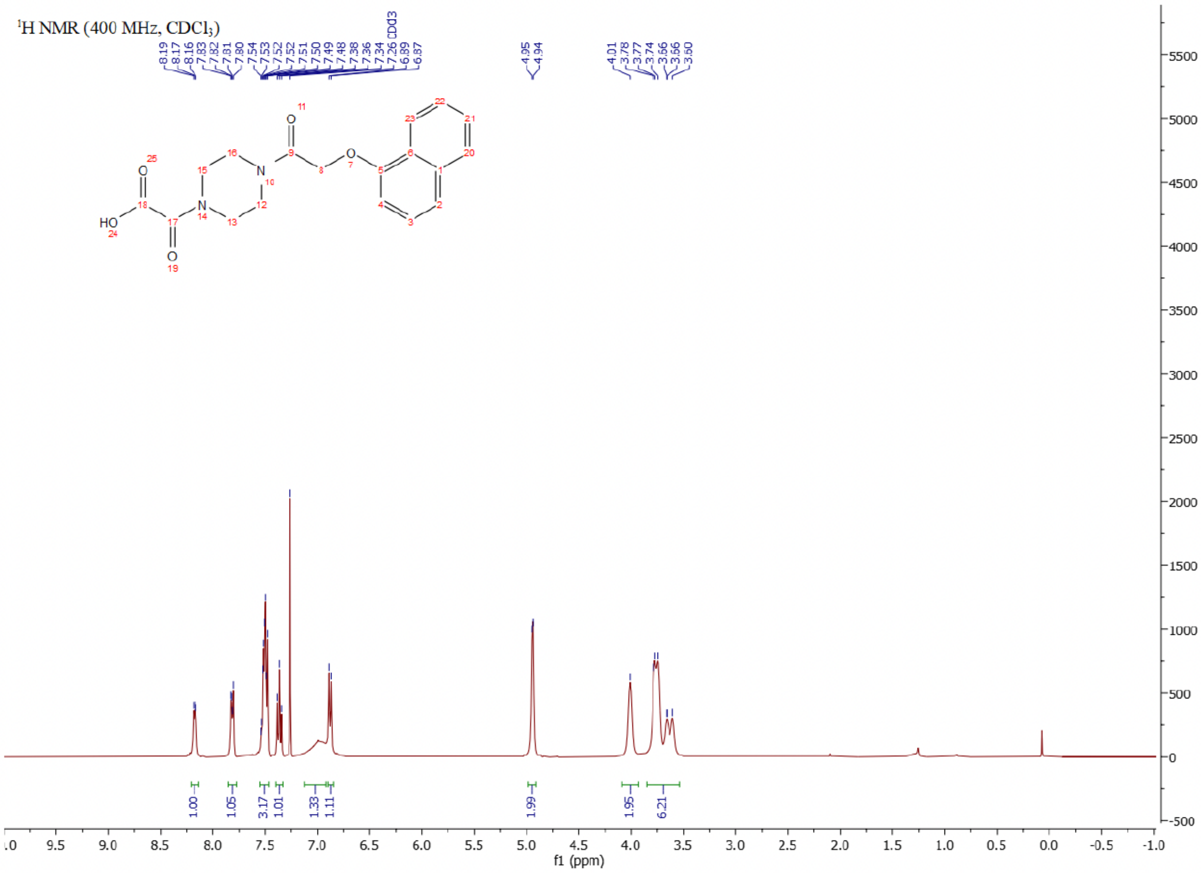

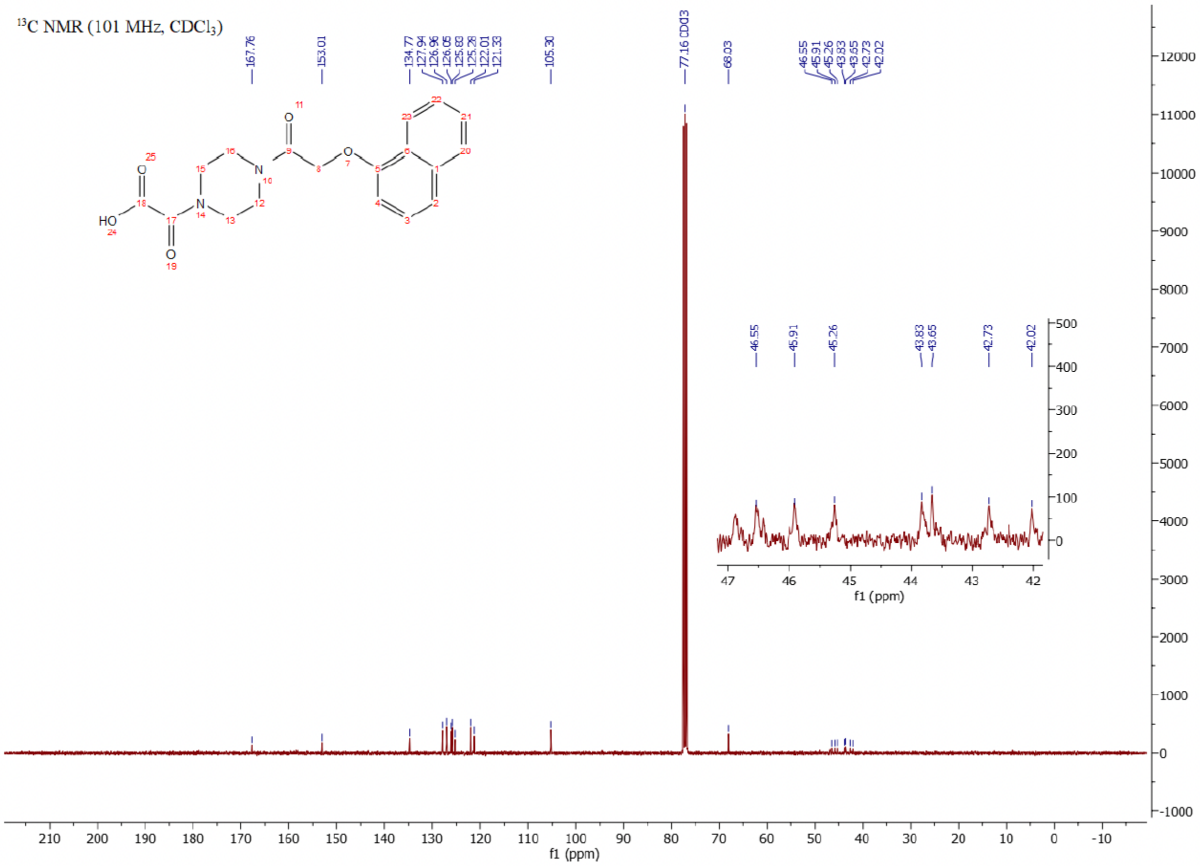

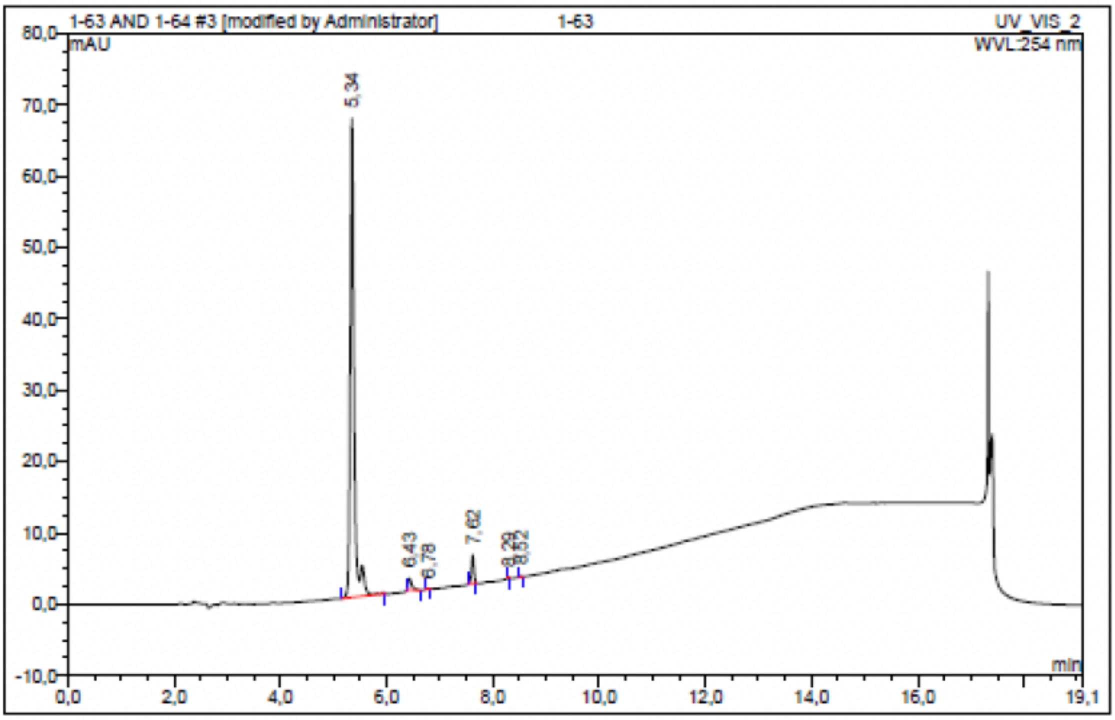

| No. | Ret.Time min | Peak Name | Height mAU | Area mAU*min | Rel.Area % | Amount | Resolution(EP) |
|---|---|---|---|---|---|---|---|
| 1 | 5,34 | n.a. | 67,096 | 5,669 | 94,02 | n.a. | 9,08 |
| 2 | 6,43 | n.a. | 1,577 | 0,133 | 2,21 | n.a. | 3,14 |
| 3 | 6,78 | n.a. | 0,096 | 0,005 | 0,09 | n.a. | 9,29 |
| 4 | 7,62 | n.a. | 3,941 | 0,215 | 3,56 | n.a. | 8,83 |
| 5 | 8,29 | n.a. | 0,085 | 0,003 | 0,05 | n.a. | 3,36 |
| 6 | 8,52 | n.a. | 0,108 | 0,004 | 0,07 | n.a. | n.a. |
| Total: | | | 72,903 | 6,030 | 100,00 | 0,000 | |

Compound 37 - TUG-2257

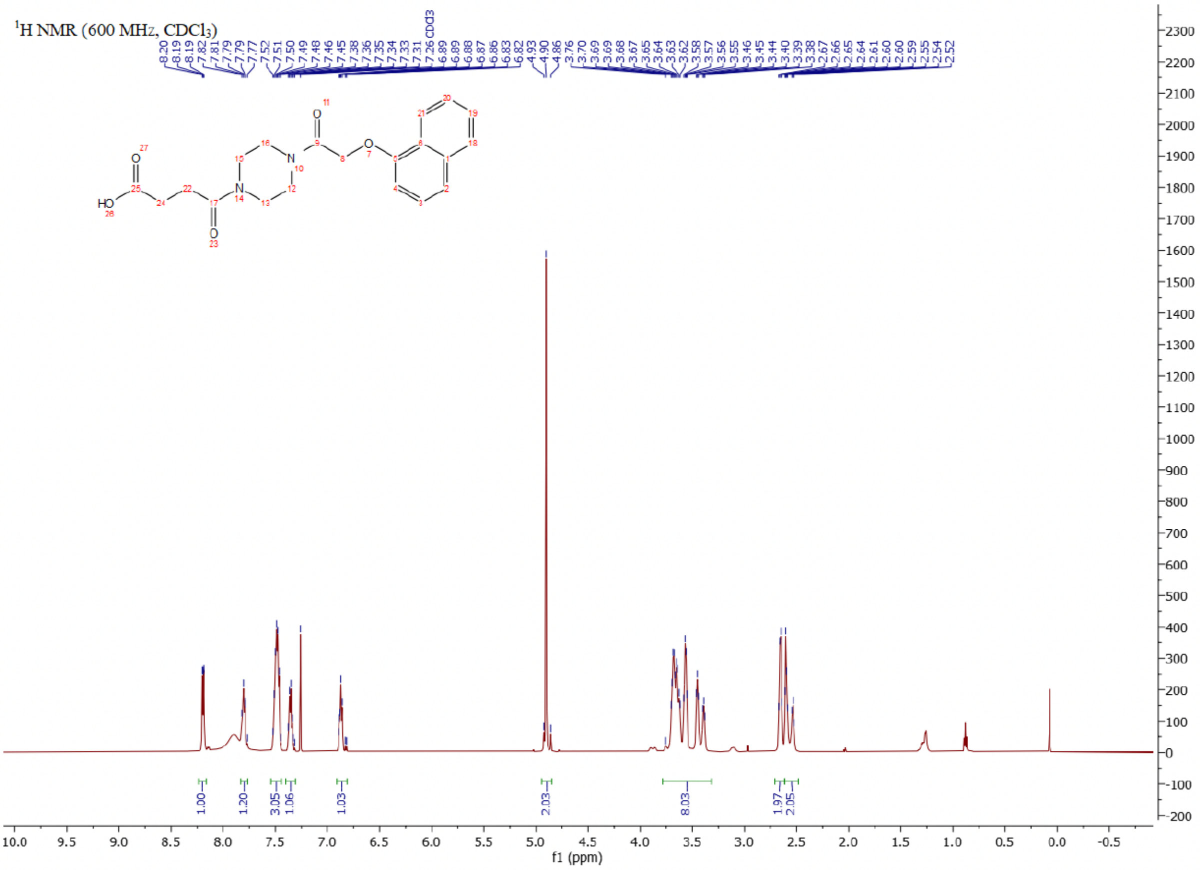

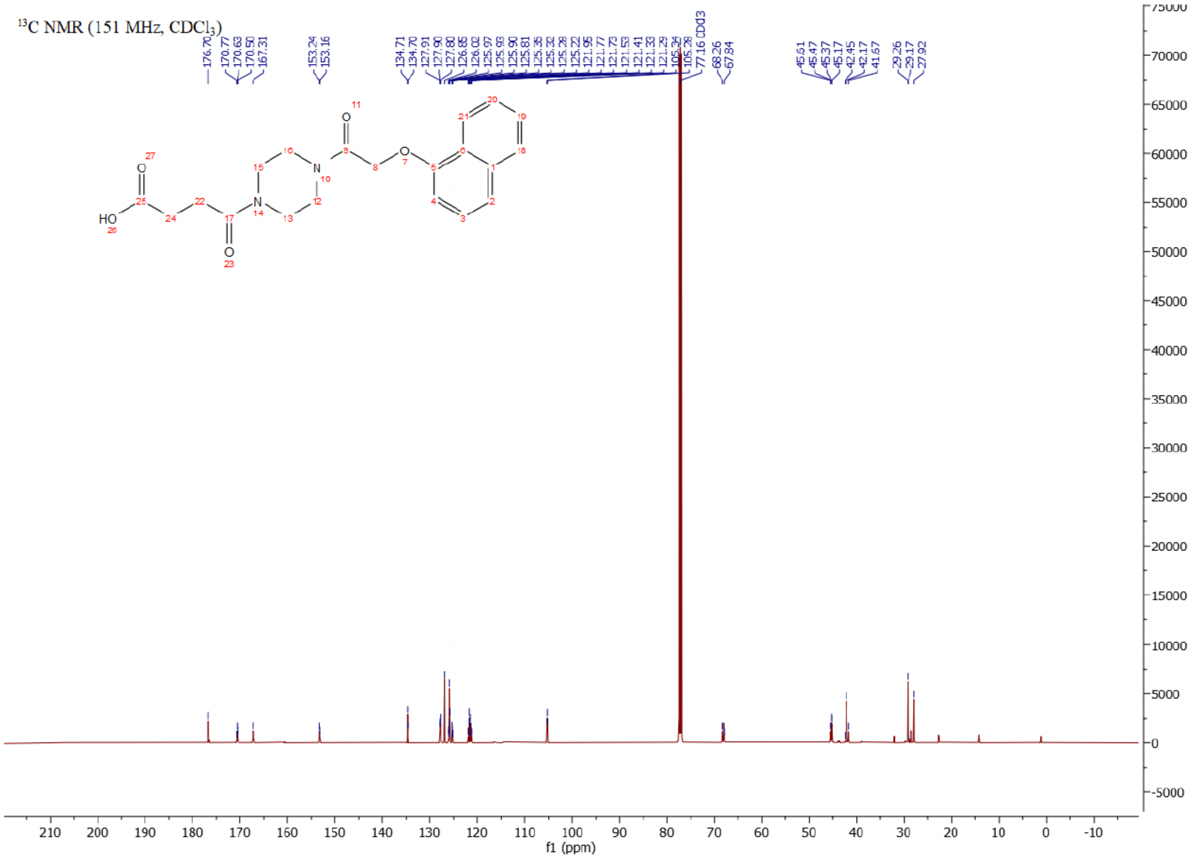

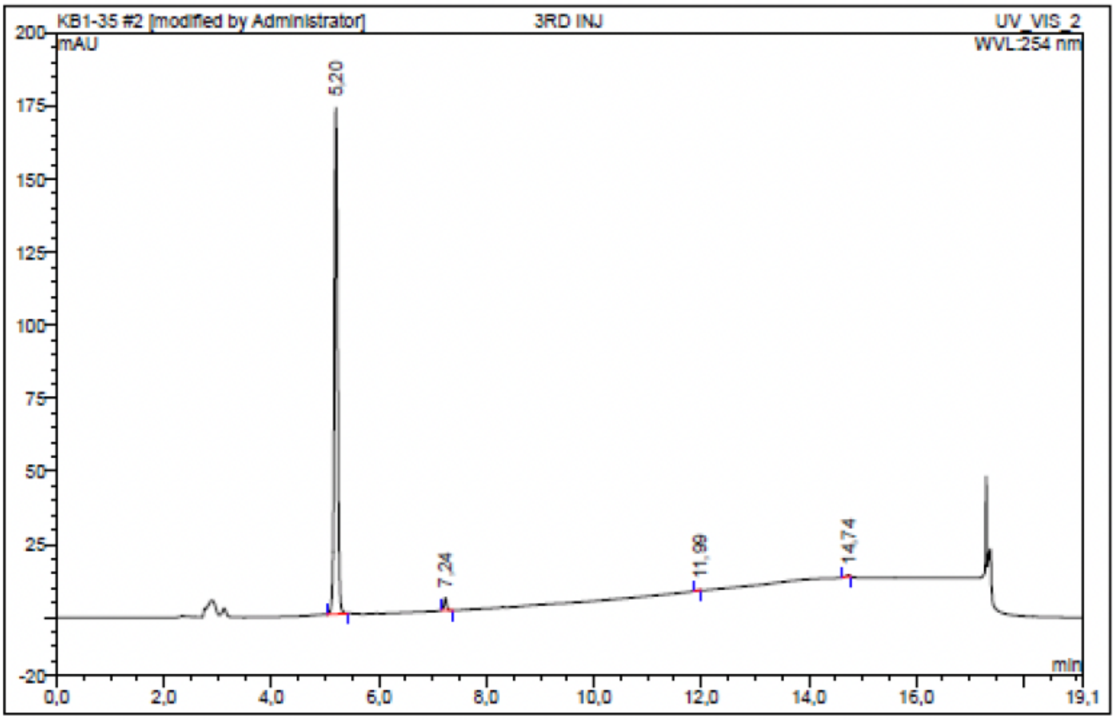

| No. | Ret.Time min | Peak Name | Height mAU | Area mAU*min | Rel.Area % | Amount | Resolution(EP) |
|---|---|---|---|---|---|---|---|
| 1 | 5,20 | n.a. | 173,342 | 11,734 | 97,68 | n.a. | 21,17 |
| 2 | 7,24 | n.a. | 4,192 | 0,239 | 1,99 | n.a. | 34,85 |
| 3 | 11,99 | n.a. | 0,023 | 0,001 | 0,01 | n.a. | 18,22 |
| 4 | 14,74 | n.a. | 0,526 | 0,039 | 0,32 | n.a. | n.a. |
| Total: | | | 178,083 | 12,012 | 100,00 | 0,000 | |

Compound 38 - TUG-2606

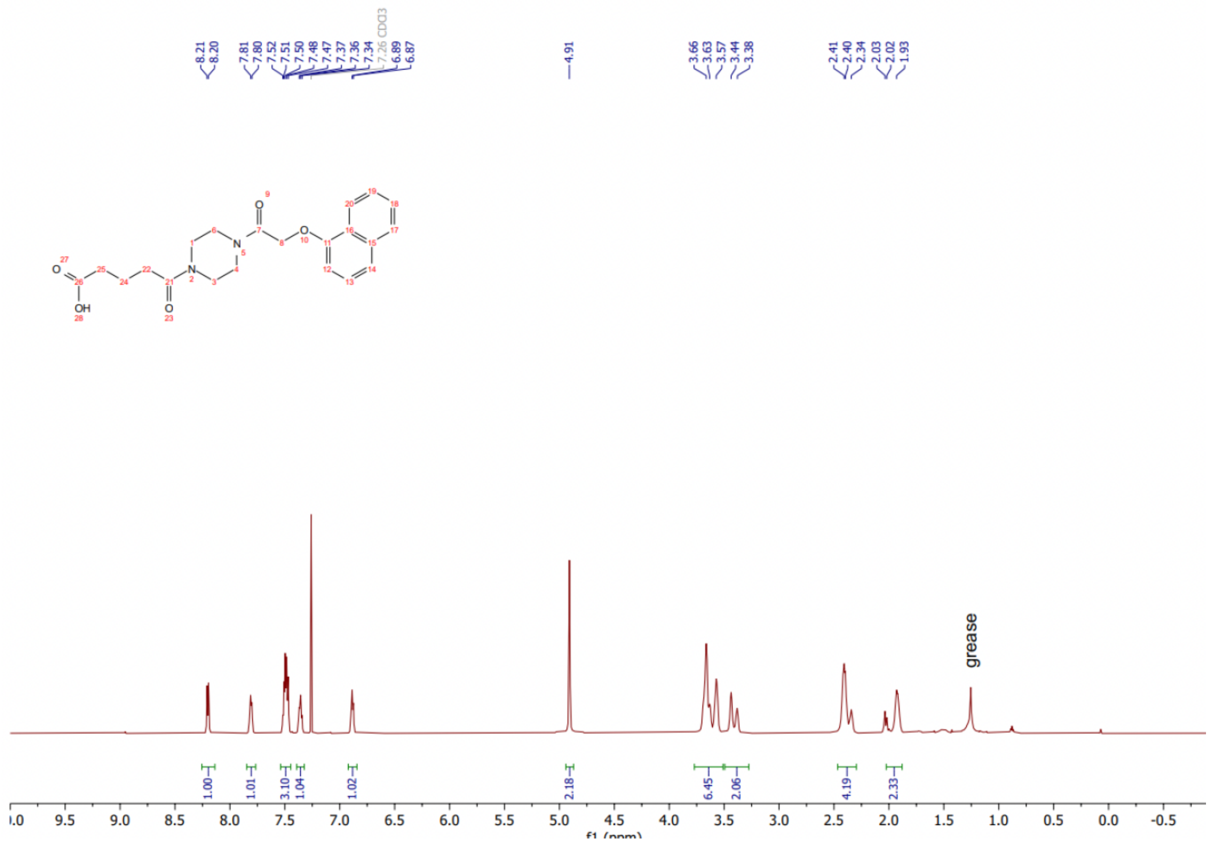

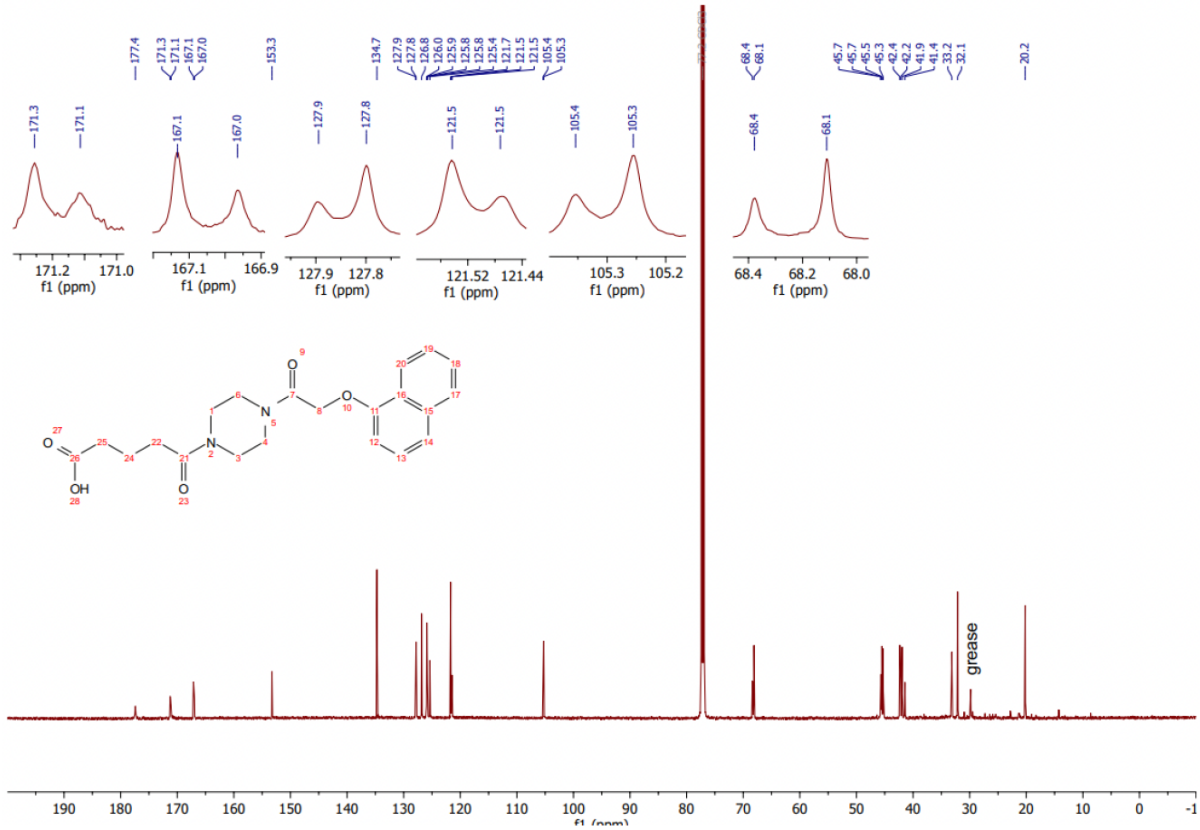

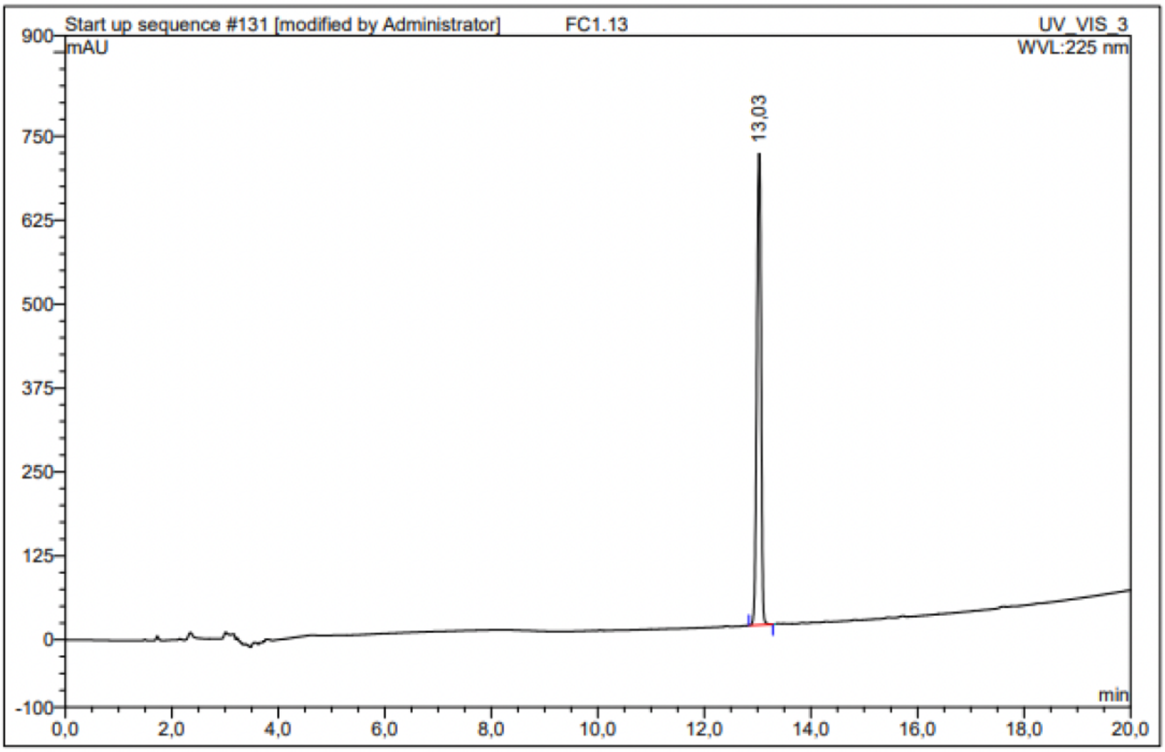

| No. | Ret.Time min | Peak Name | Height mAU | Area mAU*min | Rel.Area % | Amount | Resolution(EP) |
|---|---|---|---|---|---|---|---|
| 1 | 13,03 | n.a. | 702,004 | 62,889 | 100,00 | n.a. | n.a. |
| Total: | | | 702,004 | 62,889 | 100,00 | 0,000 | |

Compound 39 - TUG-2607

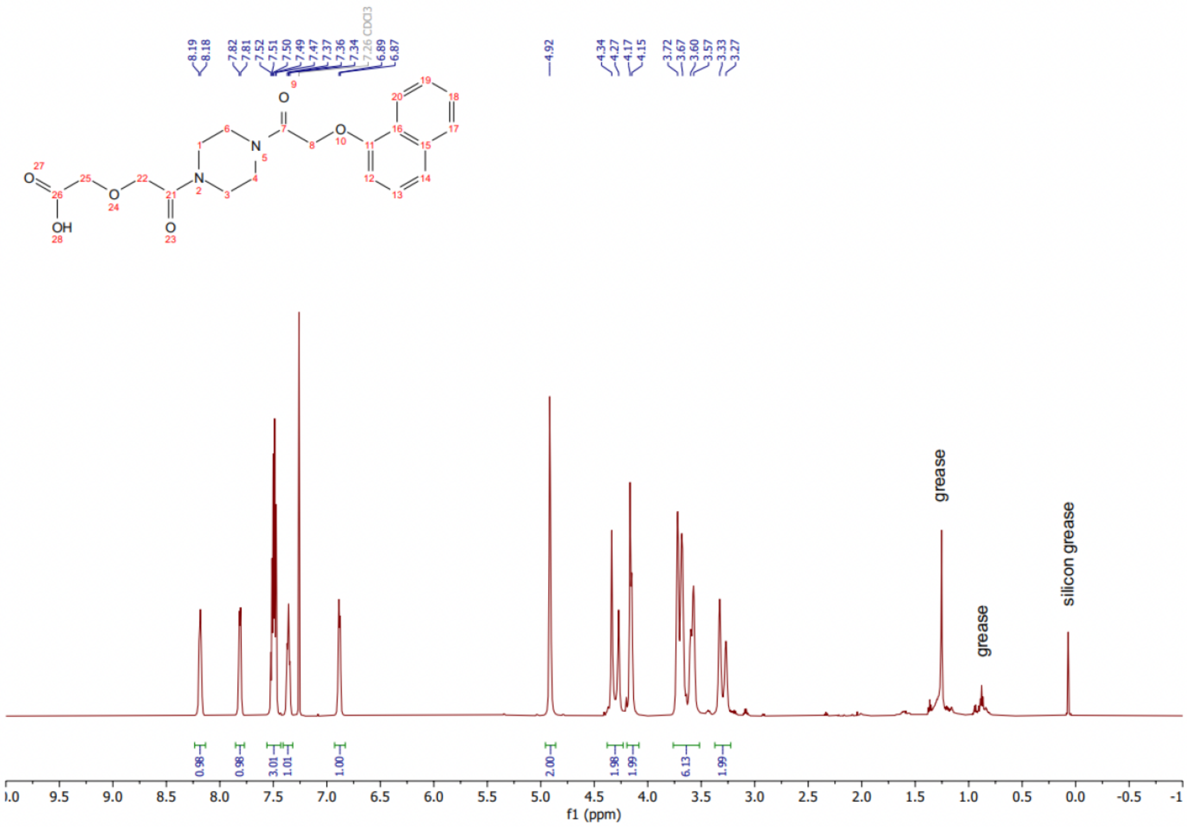

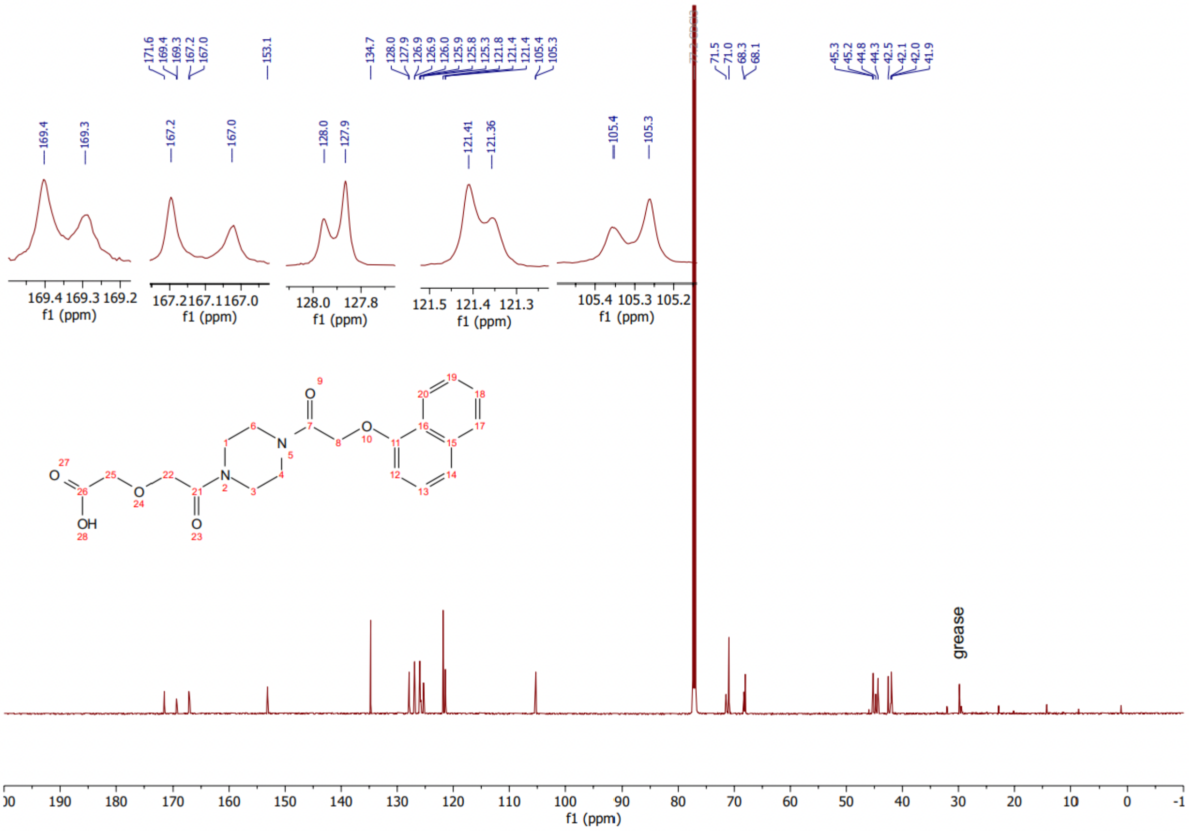

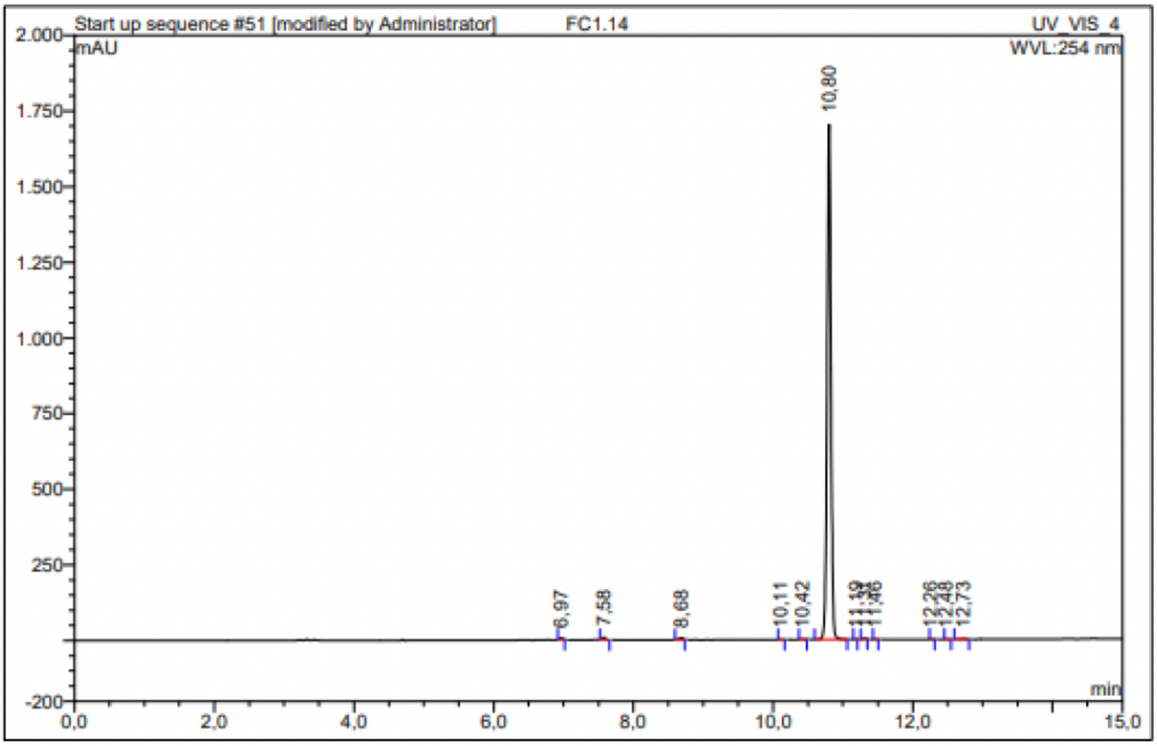

| No. | Ret.Time min | Peak Name | Height mAU | Area mAU*min | Rel.Area % | Amount | Resolution(EP) |
|---|---|---|---|---|---|---|---|
| 1 | 6,97 | n.a. | 5,451 | 0,296 | 0,27 | n.a. | 7,35 |
| 2 | 7,58 | n.a. | 8,127 | 0,375 | 0,34 | n.a. | 12,40 |
| 3 | 8,68 | n.a. | 5,550 | 0,354 | 0,33 | n.a. | 15,02 |
| 4 | 10,11 | n.a. | 0,738 | 0,037 | 0,03 | n.a. | 3,36 |
| 5 | 10,42 | n.a. | 1,629 | 0,093 | 0,09 | n.a. | 3,97 |
| 6 | 10,80 | n.a. | 1700,930 | 107,194 | 98,59 | n.a. | 4,89 |
| 7 | 11,19 | n.a. | 0,396 | 0,014 | 0,01 | n.a. | 1,82 |
| 8 | 11,31 | n.a. | 3,540 | 0,169 | 0,16 | n.a. | 1,93 |
| 9 | 11,46 | n.a. | 0,595 | 0,024 | 0,02 | n.a. | 11,87 |
| 10 | 12,26 | n.a. | 0,780 | 0,031 | 0,03 | n.a. | 3,35 |
| 11 | 12,48 | n.a. | 0,268 | 0,009 | 0,01 | n.a. | 1,76 |
| 12 | 12,73 | n.a. | 1,504 | 0,129 | 0,12 | n.a. | n.a. |
| Total: | | | 1729,508 | 108,723 | 100,00 | 0,000 | |

Compound 40 - TUG-2329

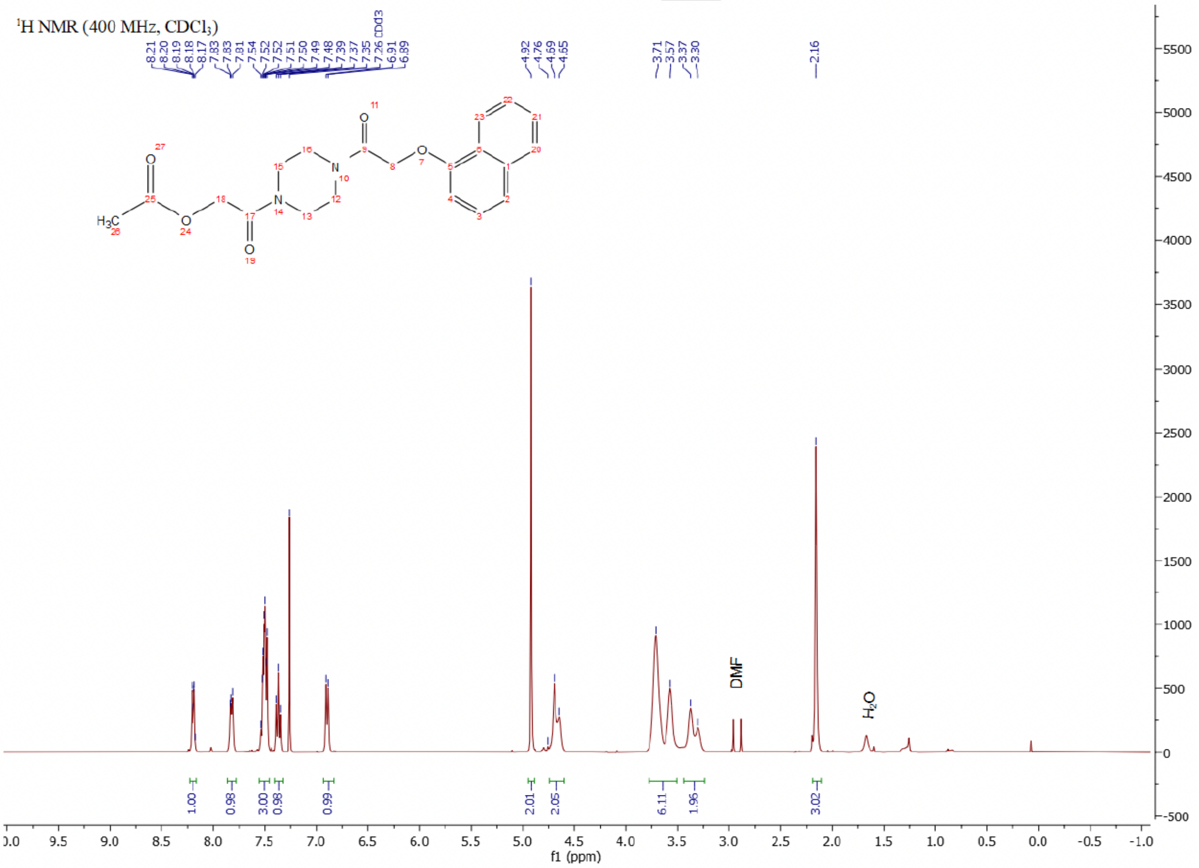

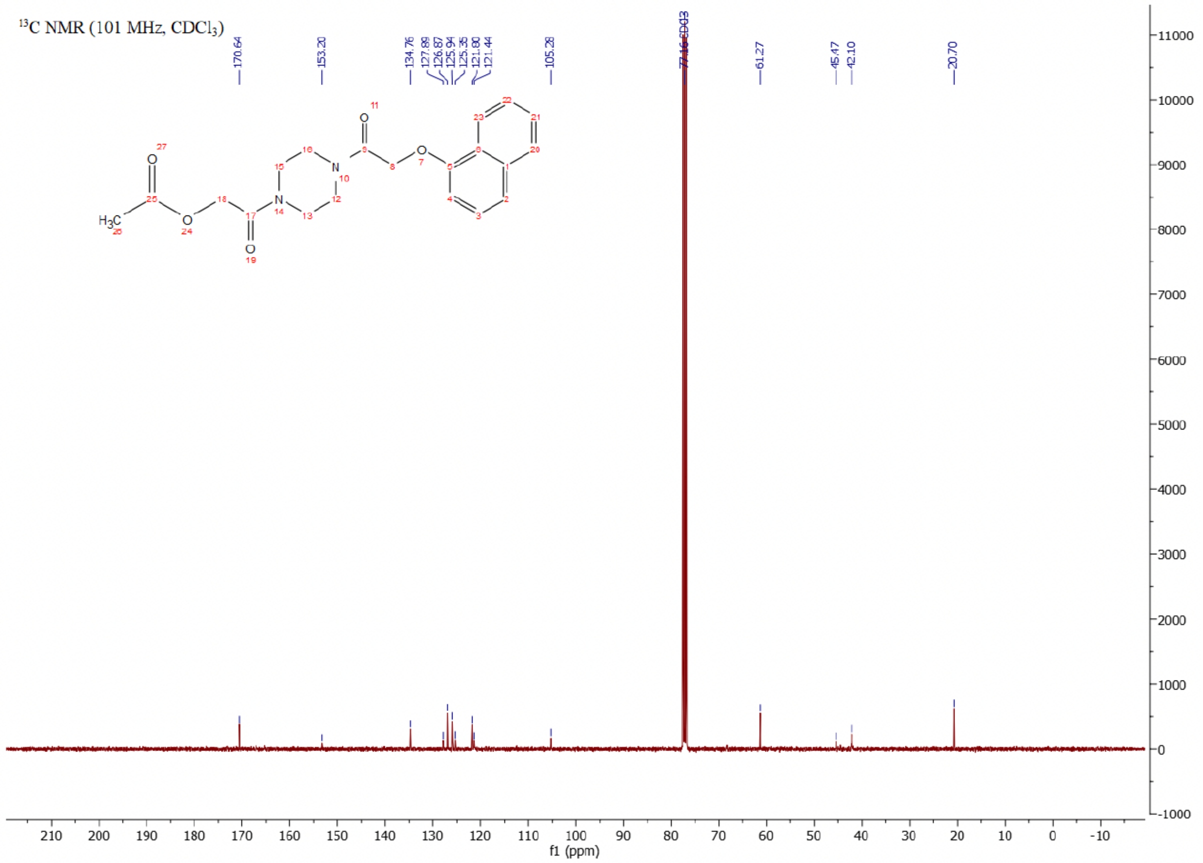

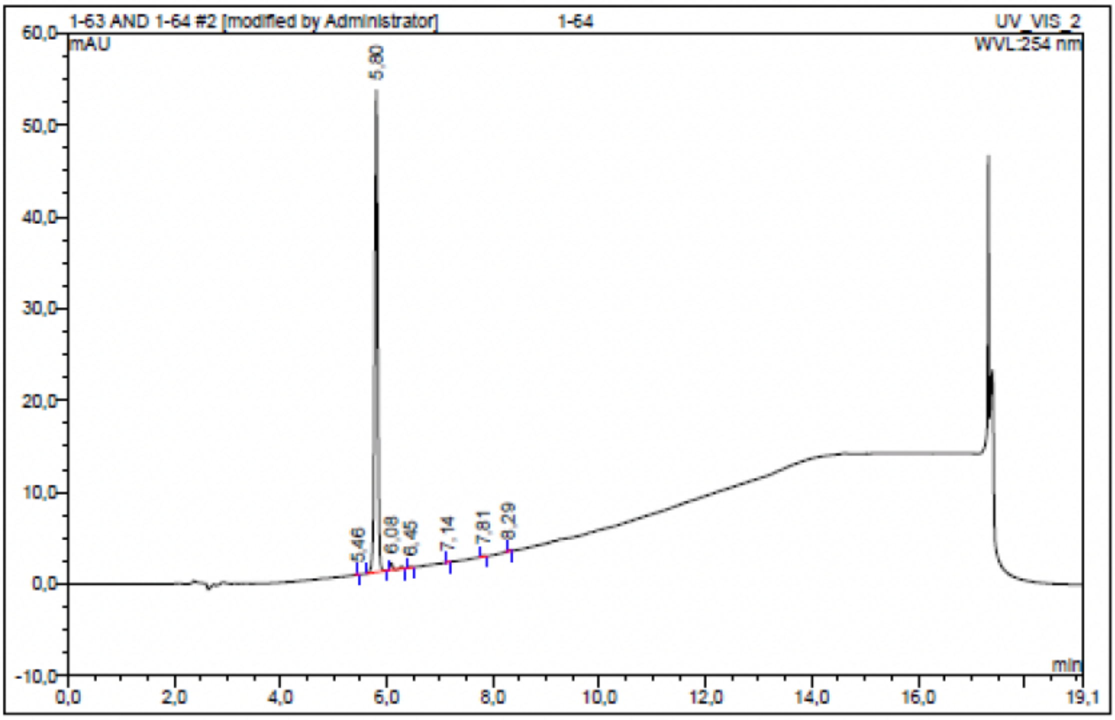

| No. | Ret.Time min | Peak Name | Height mAU | Area mAU*min | Rel.Area % | Amount | Resolution(EP) |
|---|---|---|---|---|---|---|---|
| 1 | 5,46 | n.a. | 0,026 | 0,001 | 0,03 | n.a. | 4,36 |
| 2 | 5,80 | n.a. | 52,494 | 3,443 | 97,80 | n.a. | 2,87 |
| 3 | 6,08 | n.a. | 0,767 | 0,057 | 1,61 | n.a. | 4,00 |
| 4 | 6,45 | n.a. | 0,098 | 0,005 | 0,15 | n.a. | 8,61 |
| 5 | 7,14 | n.a. | 0,028 | 0,001 | 0,03 | n.a. | 8,46 |
| 6 | 7,81 | n.a. | 0,114 | 0,006 | 0,17 | n.a. | 5,76 |
| 7 | 8,29 | n.a. | 0,161 | 0,008 | 0,22 | n.a. | n.a. |
| Total: | | | 53,687 | 3,521 | 100,00 | 0,000 | |

Compound 41 - TUG-2261

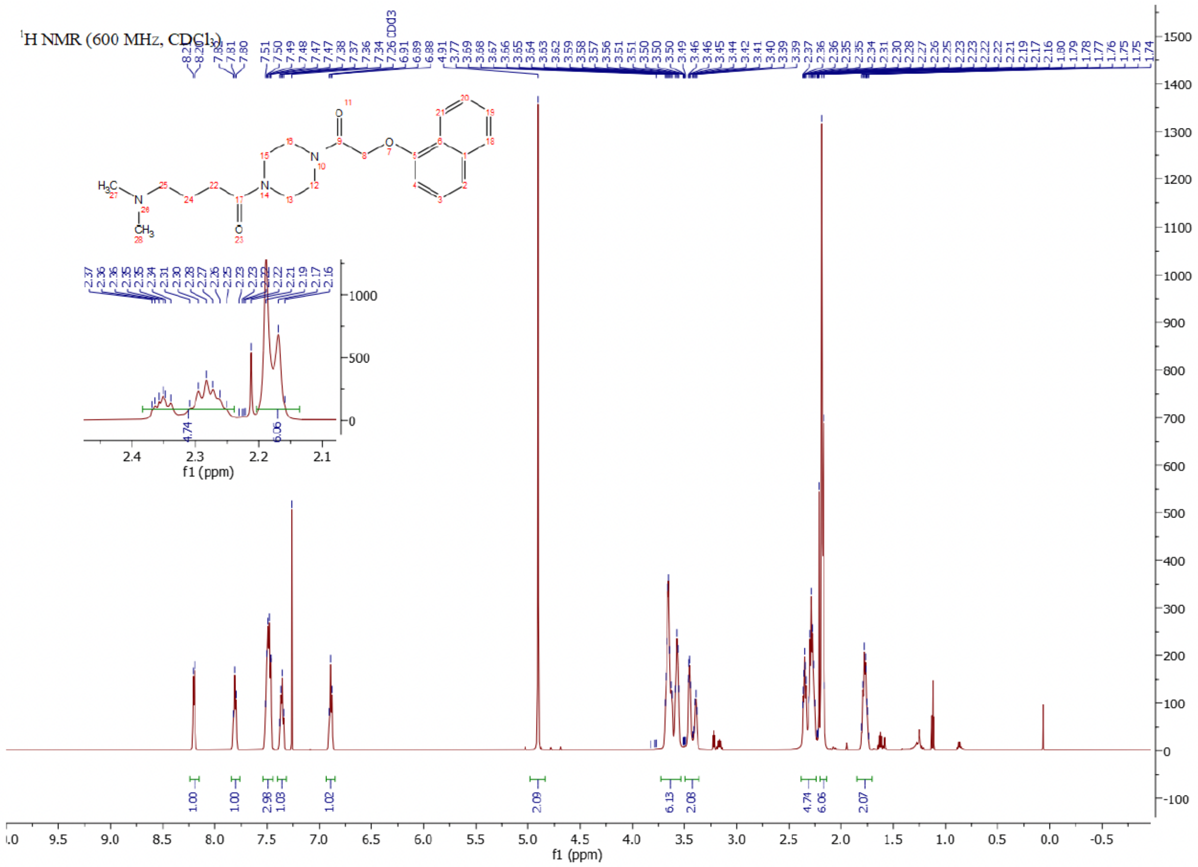

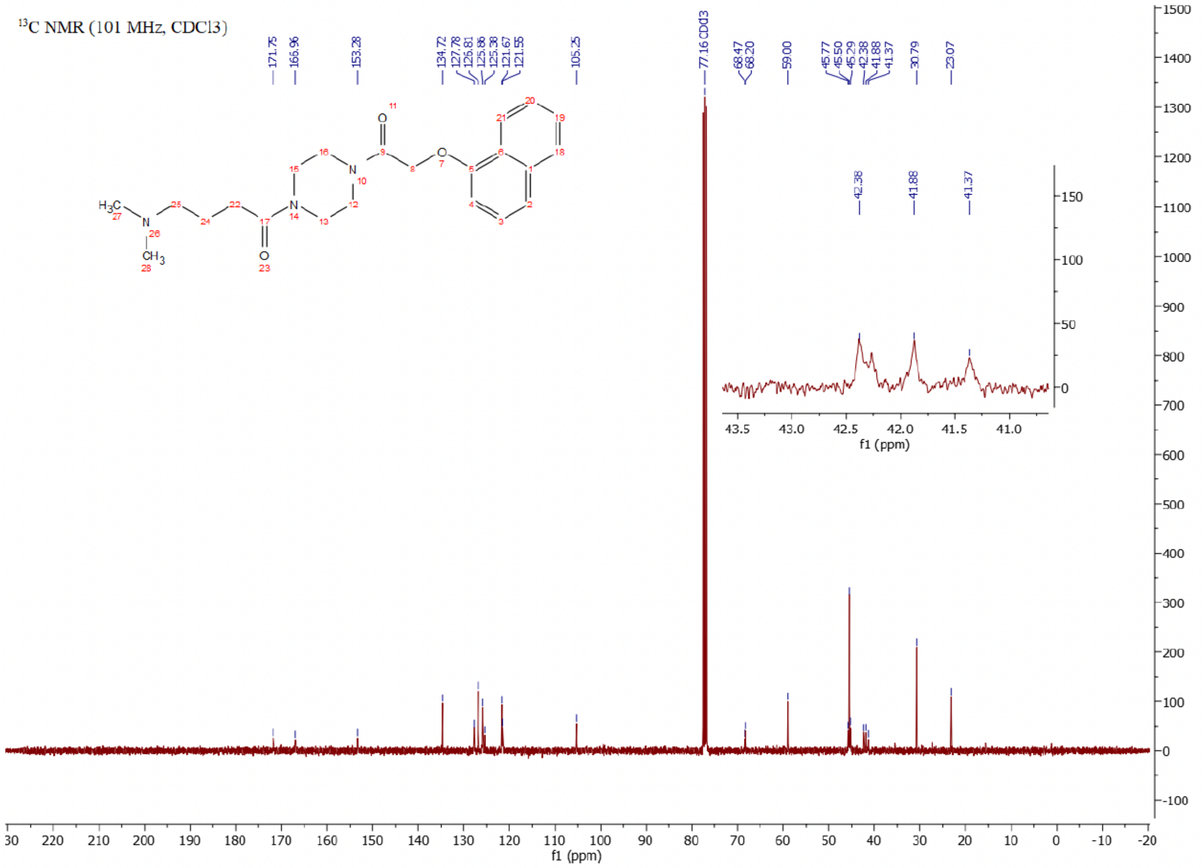

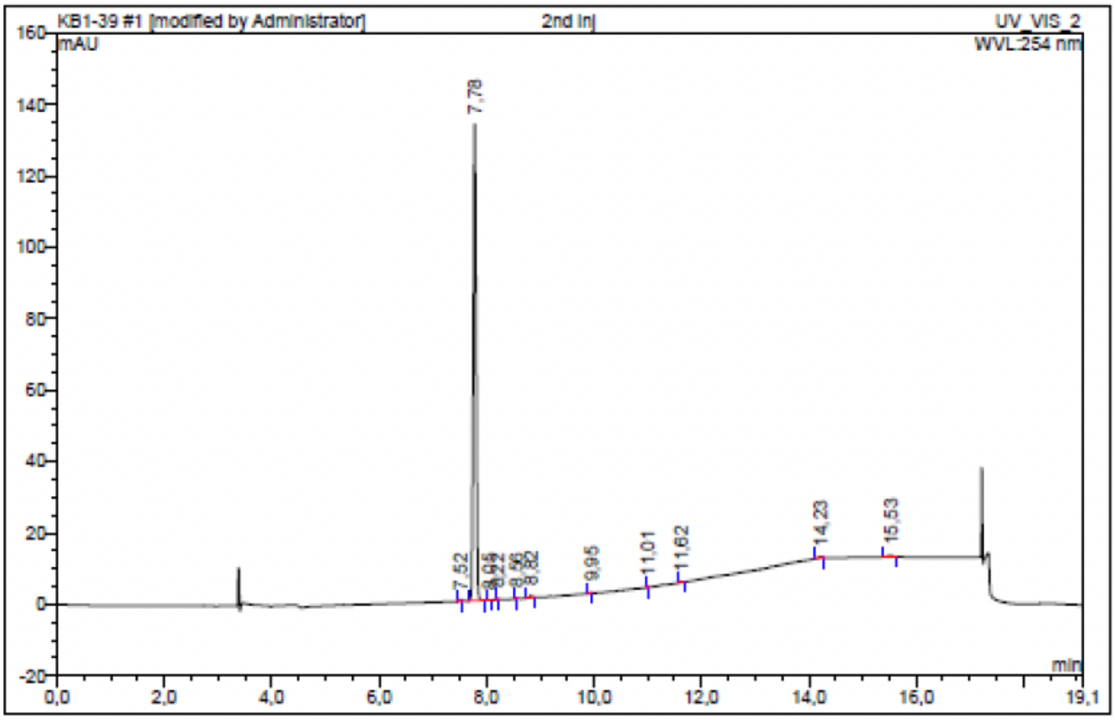

| No. | Ret.Time min | Peak Name | Height mAU | Area mAU*min | Rel.Area % | Amount | Resolution(EP) |
|---|---|---|---|---|---|---|---|
| 1 | 7,52 | n.a. | 0,405 | 0,025 | 0,30 | n.a. | 2,64 |
| 2 | 7,78 | n.a. | 133,412 | 7,947 | 98,00 | n.a. | 3,37 |
| 3 | 8,05 | n.a. | 0,067 | 0,002 | 0,03 | n.a. | 2,07 |
| 4 | 8,22 | n.a. | 0,061 | 0,004 | 0,05 | n.a. | 4,36 |
| 5 | 8,56 | n.a. | 0,032 | 0,001 | 0,02 | n.a. | 3,86 |
| 6 | 8,82 | n.a. | 0,554 | 0,028 | 0,35 | n.a. | 13,29 |
| 7 | 9,95 | n.a. | 0,026 | 0,000 | 0,00 | n.a. | 13,53 |
| 8 | 11,01 | n.a. | 0,185 | 0,007 | 0,09 | n.a. | 7,61 |
| 9 | 11,62 | n.a. | 0,225 | 0,013 | 0,16 | n.a. | 22,18 |
| 10 | 14,23 | n.a. | 0,461 | 0,040 | 0,49 | n.a. | 8,76 |
| 11 | 15,53 | n.a. | 0,401 | 0,041 | 0,51 | n.a. | n.a. |
| Total: | | | 135,829 | 8,110 | 100,00 | 0,000 | |

### Compound 42 - TUG-2711

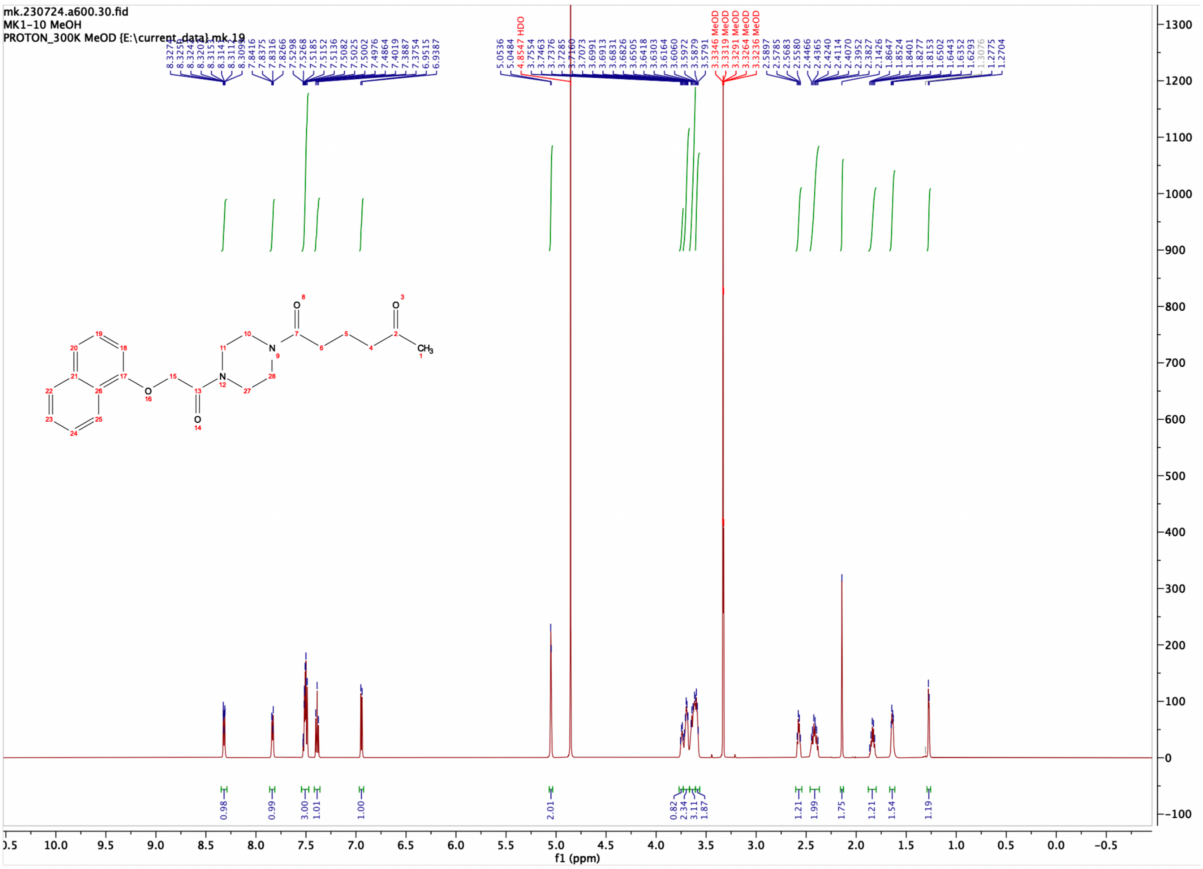

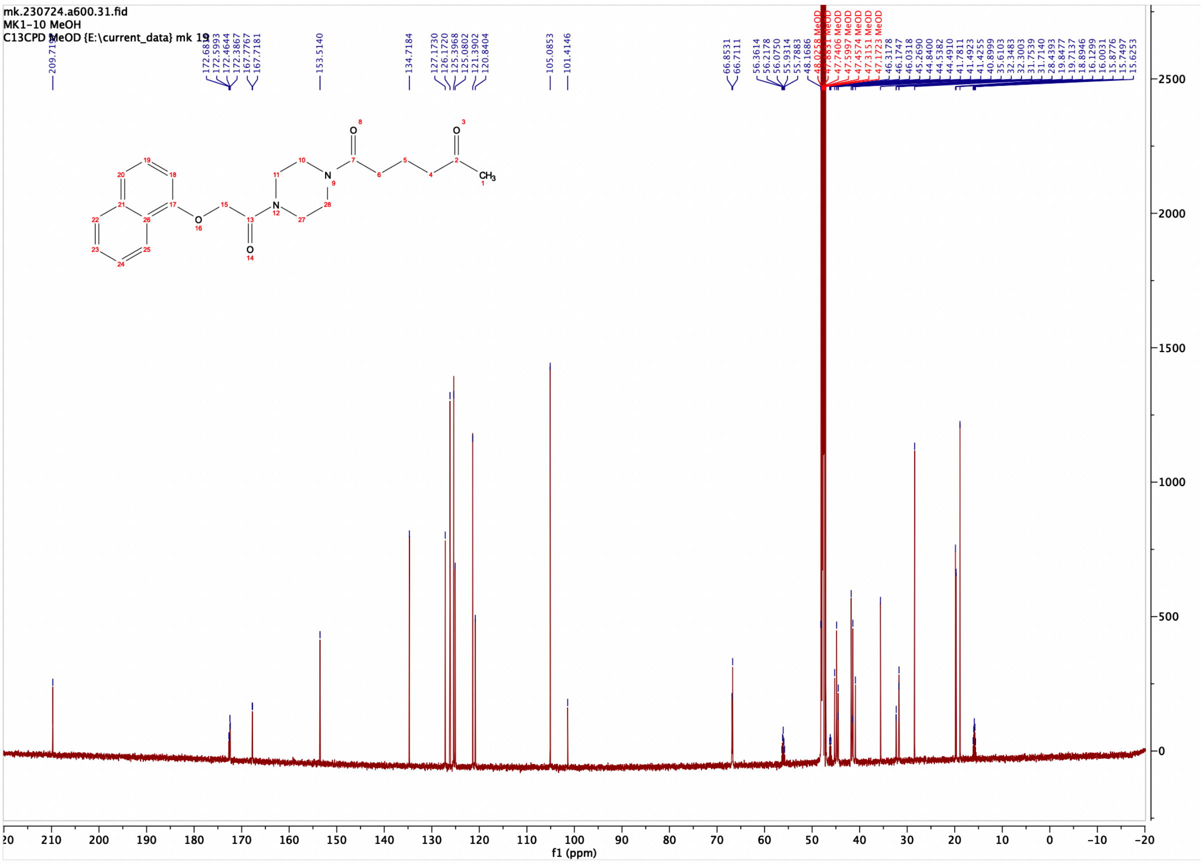

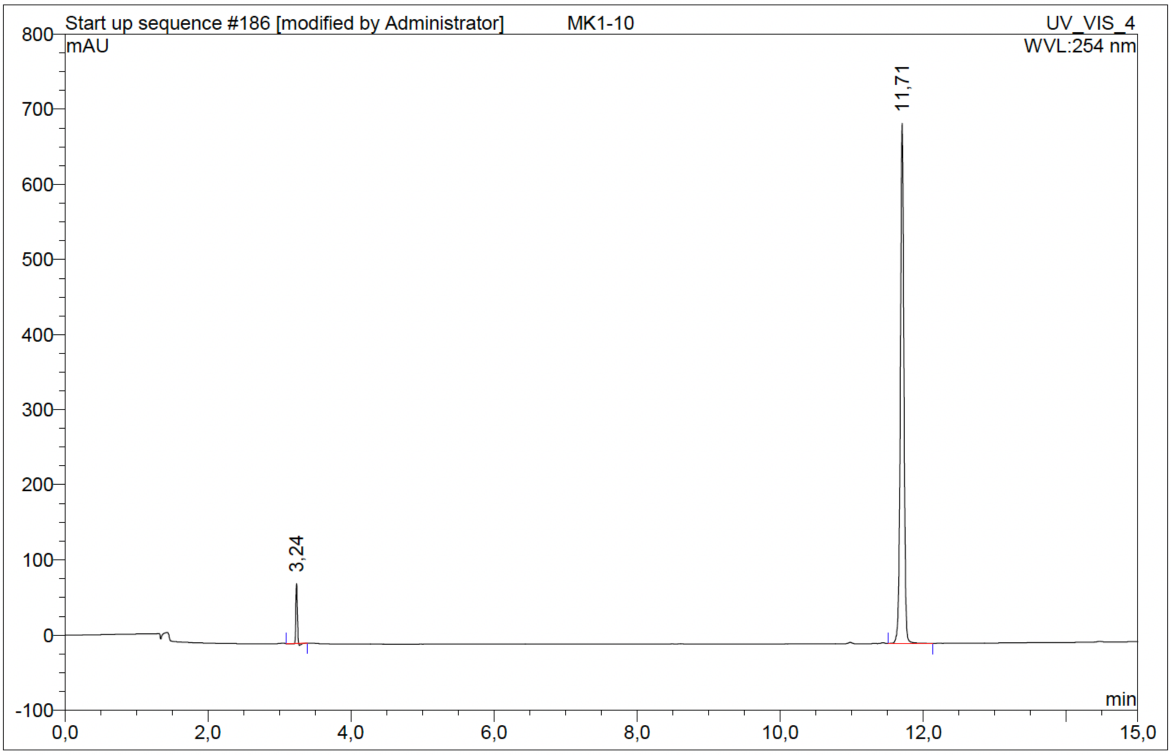

| No. | Ret.Time min | Peak Name | Height mAU | Area mAU*min | Rel.Area % | Amount | Resolution(EP) |
|---|---|---|---|---|---|---|---|
| 1 | 3,24 | n.a. | 79,651 | 1,837 | 4,30 | n.a. | 129,47 |
| 2 | 11,71 | n.a. | 692,211 | 40,874 | 95,70 | n.a. | n.a. |
| Total: | | | 771,862 | 42,711 | 100,00 | 0,000 | |

Compound 43 - TUG-2710

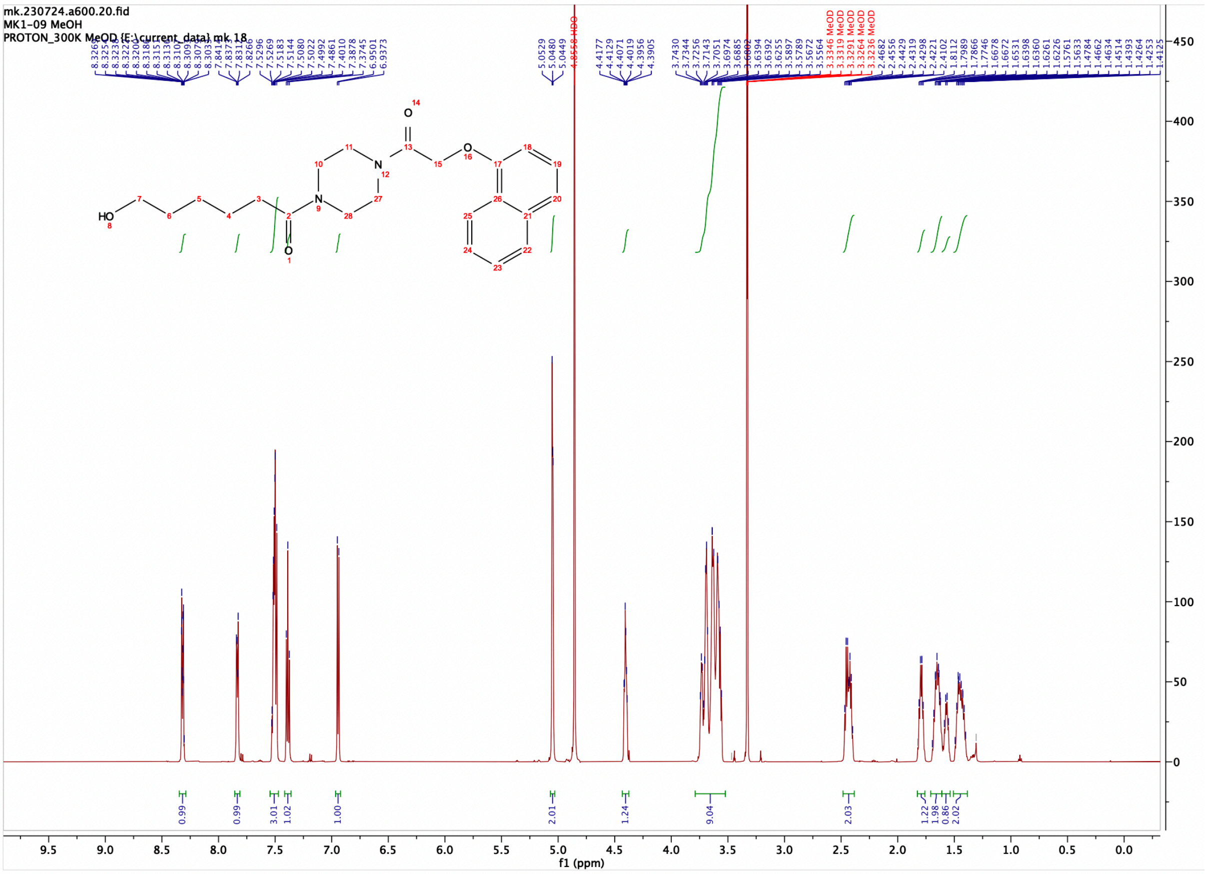

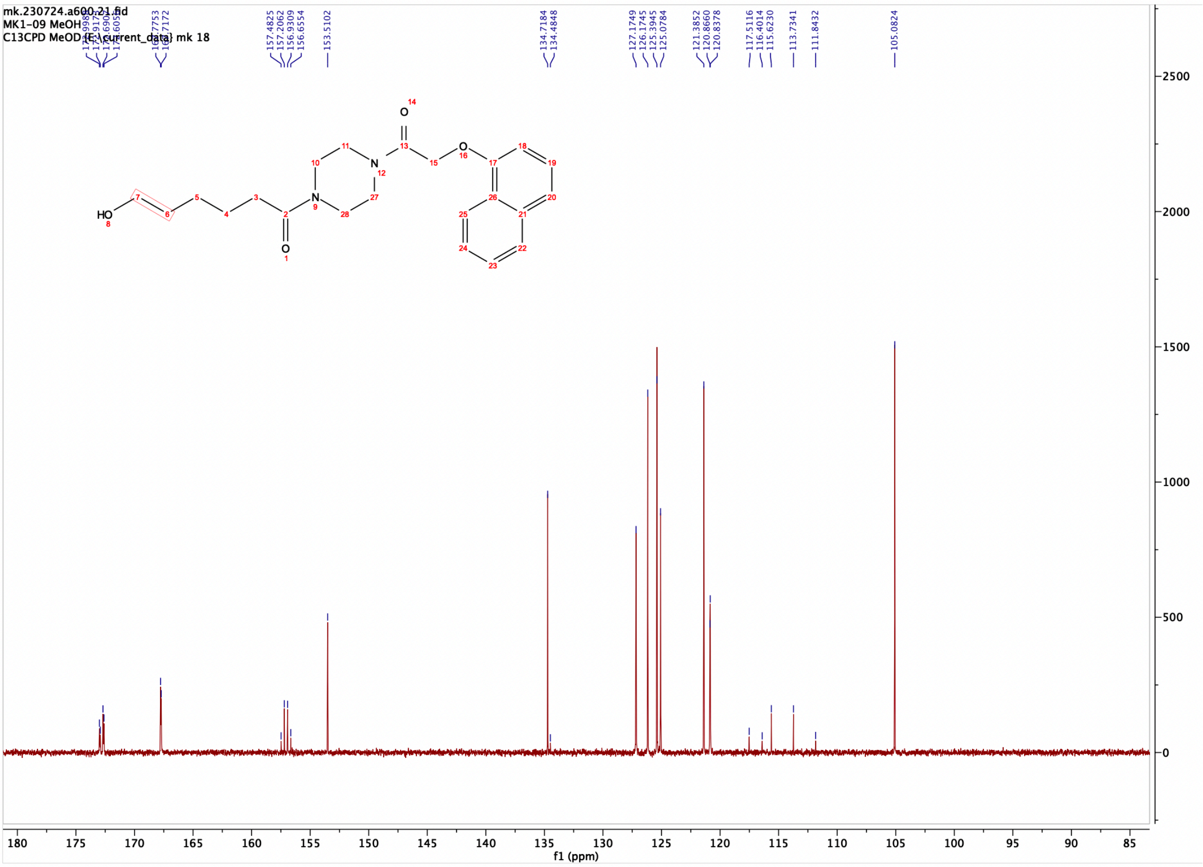

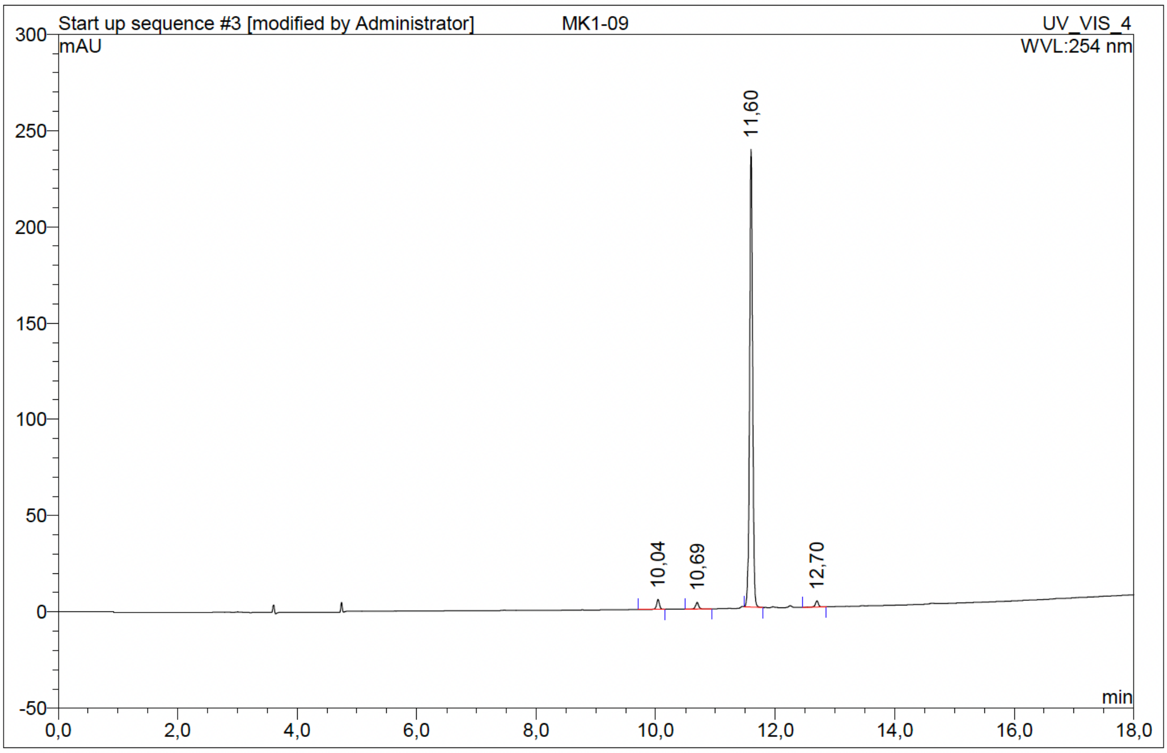

| No. | Ret.Time min | Peak Name | Height mAU | Area mAU*min | Rel.Area % | Amount | Resolution(EP) |
|---|---|---|---|---|---|---|---|
| 1 | 10,04 | n.a. | 5,178 | 0,290 | 1,95 | n.a. | 7,55 |
| 2 | 10,69 | n.a. | 3,448 | 0,202 | 1,36 | n.a. | 10,10 |
| 3 | 11,60 | n.a. | 237,812 | 14,174 | 95,36 | n.a. | 11,93 |
| 4 | 12,70 | n.a. | 3,186 | 0,198 | 1,33 | n.a. | n.a. |
| Total: | | | 249,623 | 14,864 | 100,00 | 0,000 | |

### Compound 44 - TUG-2759

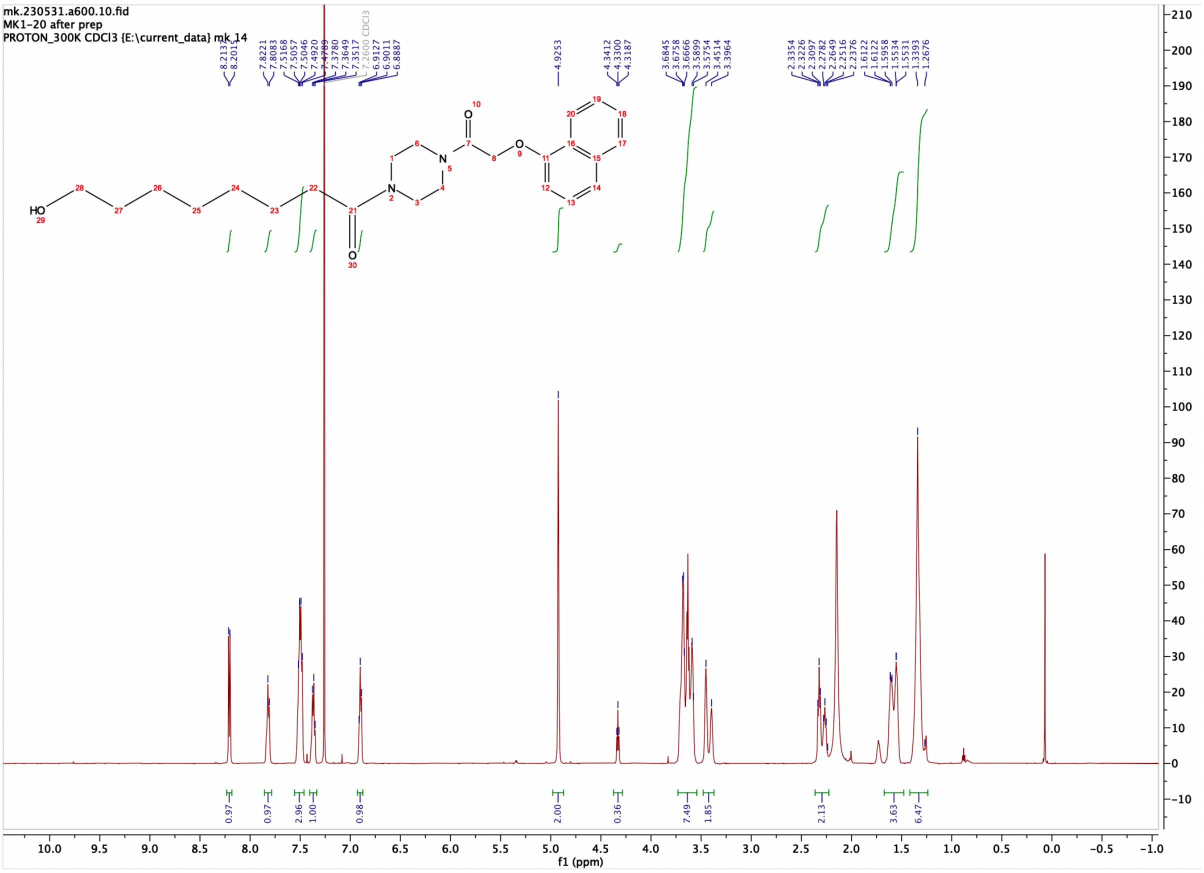

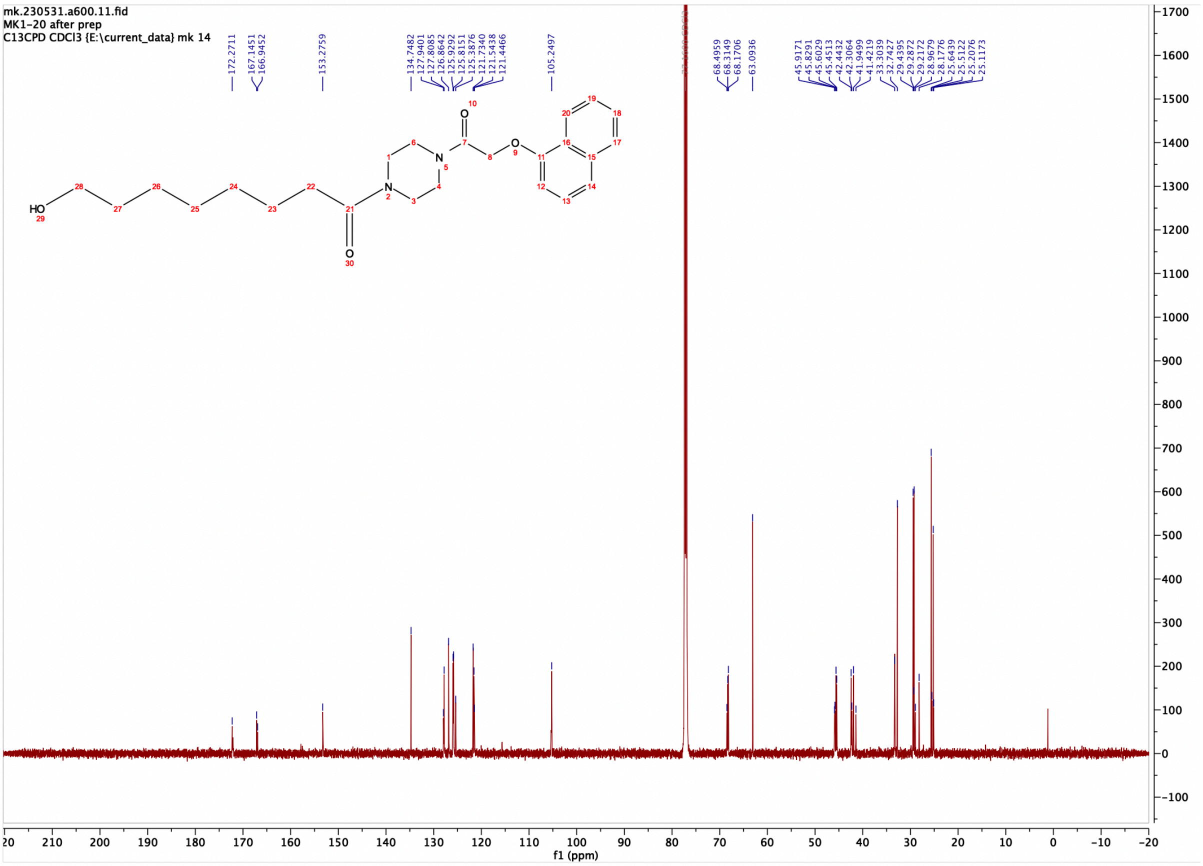

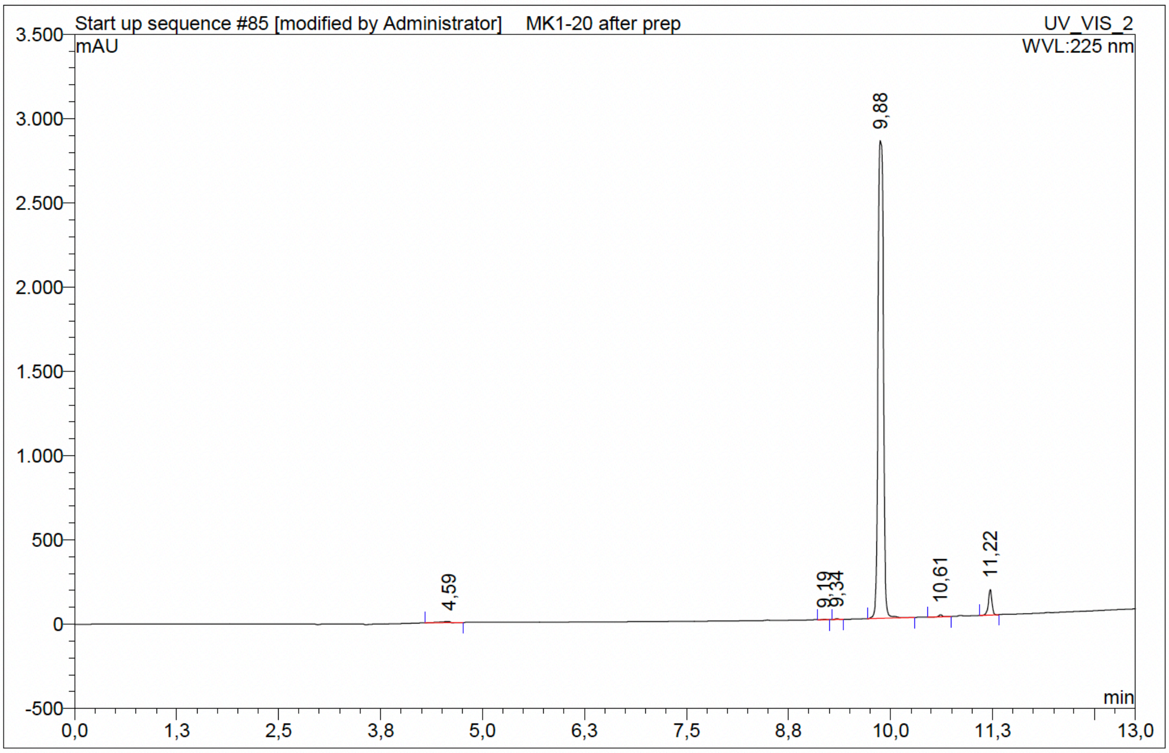

| No. | Ret.Time min | Peak Name | Height mAU | Area mAU*min | Rel.Area % | Amount | Resolution(EP) |
|---|---|---|---|---|---|---|---|
| 1 | 4,59 | n.a. | 5,962 | 0,821 | 0,37 | n.a. | 29,23 |
| 2 | 9,19 | n.a. | 2,249 | 0,091 | 0,04 | n.a. | 2,12 |
| 3 | 9,34 | n.a. | 5,537 | 0,239 | 0,11 | n.a. | 5,70 |
| 4 | 9,88 | n.a. | 2835,933 | 210,189 | 95,75 | n.a. | 7,81 |
| 5 | 10,61 | n.a. | 12,251 | 0,531 | 0,24 | n.a. | 8,24 |
| 6 | 11,22 | n.a. | 151,048 | 7,651 | 3,49 | n.a. | n.a. |
| Total: | | | 3012,980 | 219,523 | 100,00 | 0,000 | |

### Compound 45 - TUG-2760

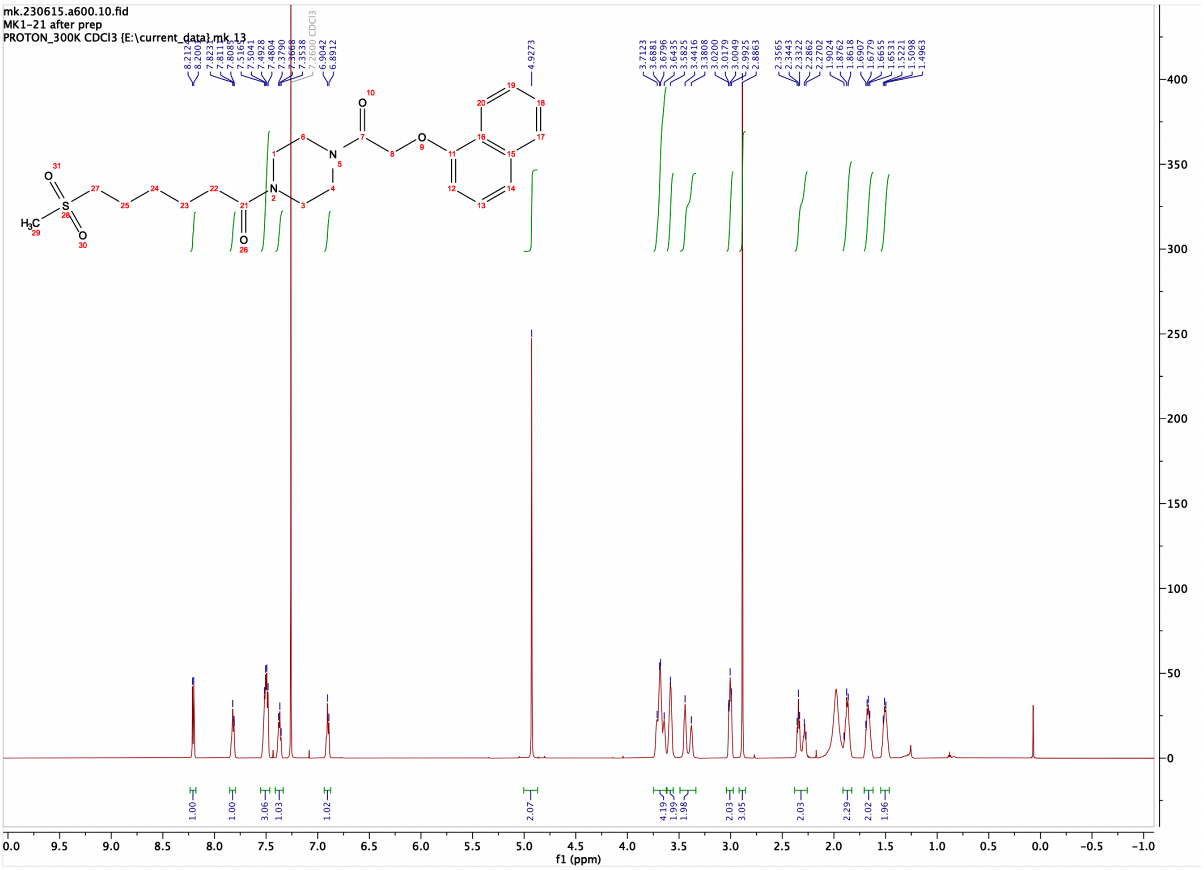

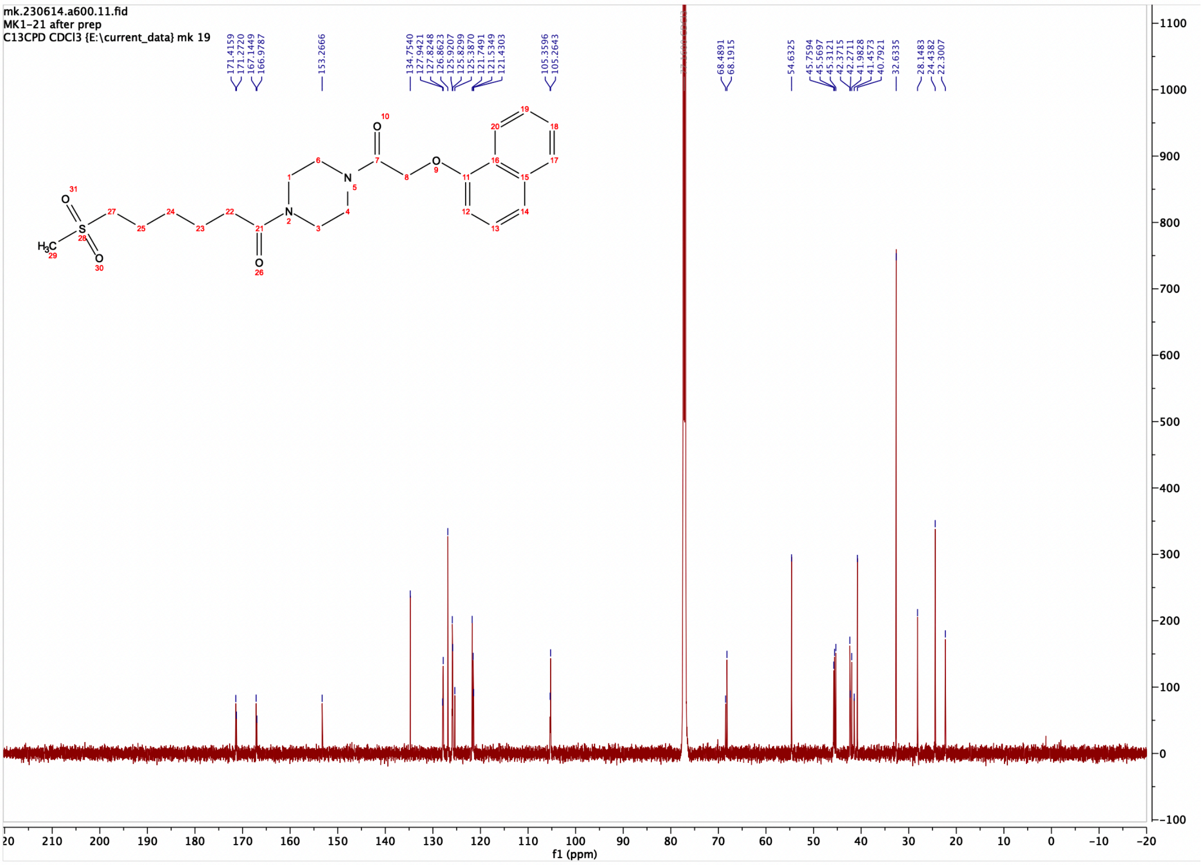

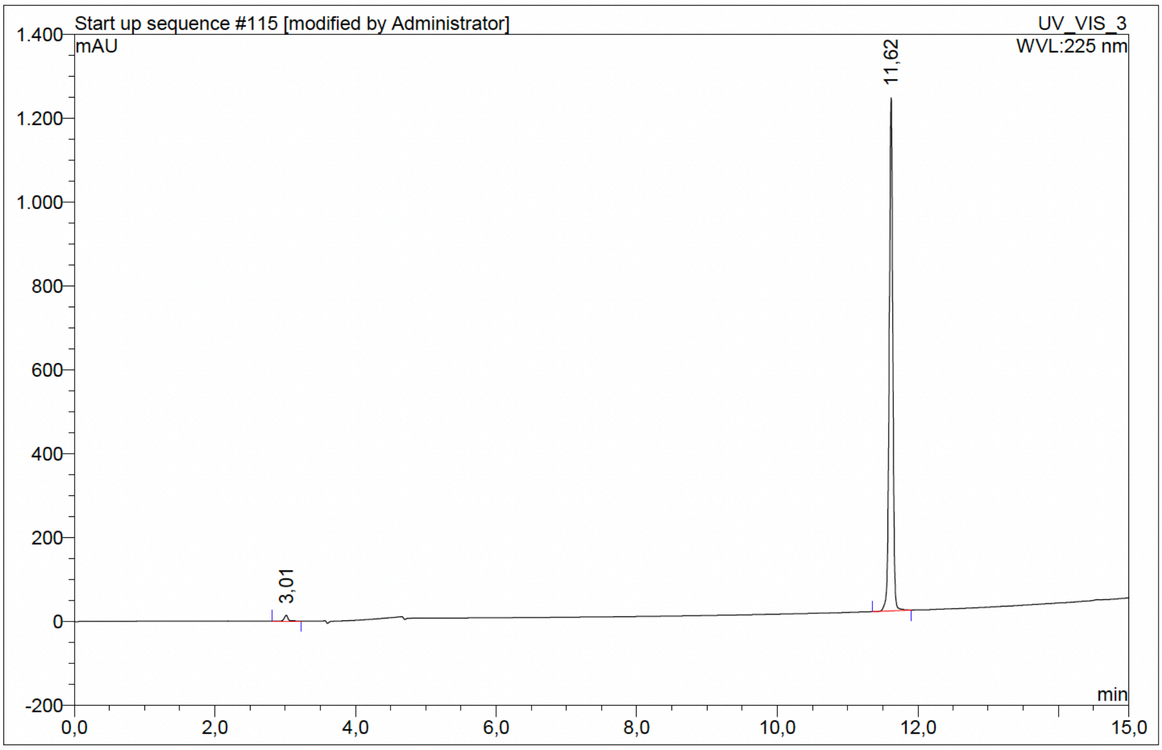

| No. | Ret.Time min | Peak Name | Height mAU | Area mAU*min | Rel.Area % | Amount | Resolution(EP) |
|---|---|---|---|---|---|---|---|
| 1 | 3,01 | n.a. | 13,615 | 0,748 | 1,05 | n.a. | 101,69 |
| 2 | 11,62 | n.a. | 1223,273 | 70,627 | 98,95 | n.a. | n.a. |
| Total: | | | 1236,887 | 71,375 | 100,00 | 0,000 | |

Compound 46 - TUG-2761

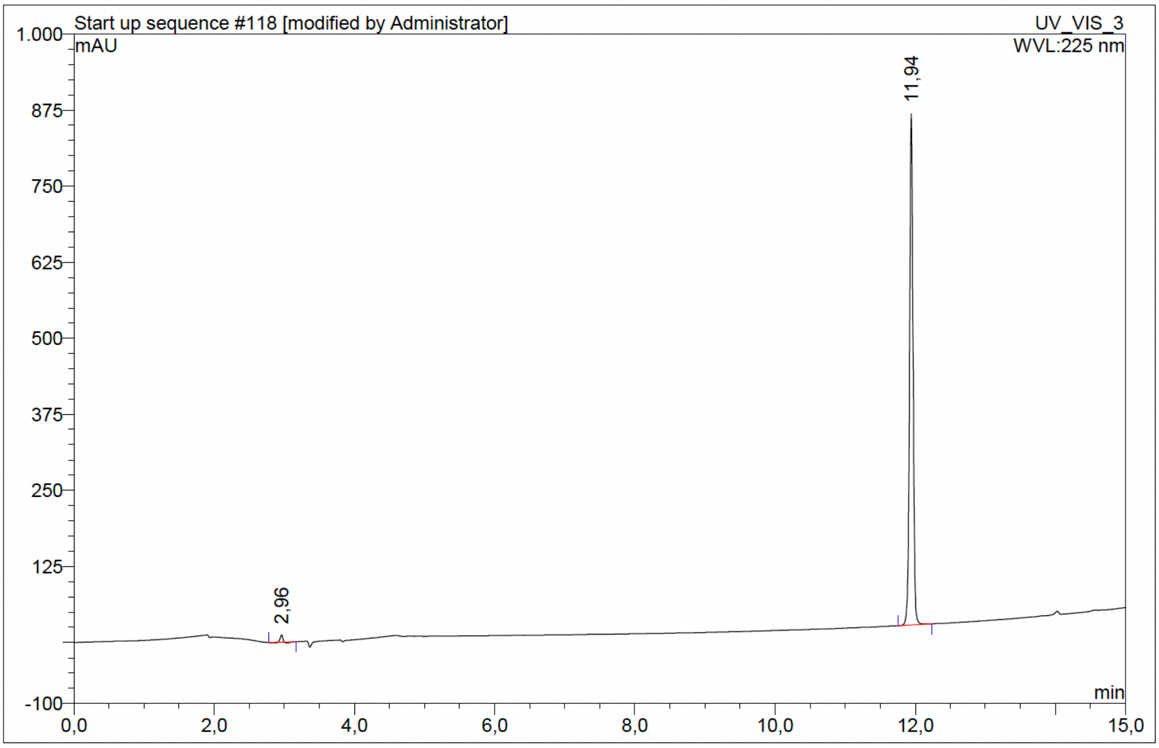

| No. | Ret.Time min | Peak Name | Height mAU | Area mAU*min | Rel.Area % | Amount | Resolution(EP) |
|---|---|---|---|---|---|---|---|
| 1 | 2,96 | n.a. | 12,041 | 0,362 | 0,75 | n.a. | 111,93 |
| 2 | 11,94 | n.a. | 839,871 | 47,920 | 99,25 | n.a. | n.a. |
| Total: | | | 851,912 | 48,282 | 100,00 | 0,000 | |

Compound 47 - TUG-2725

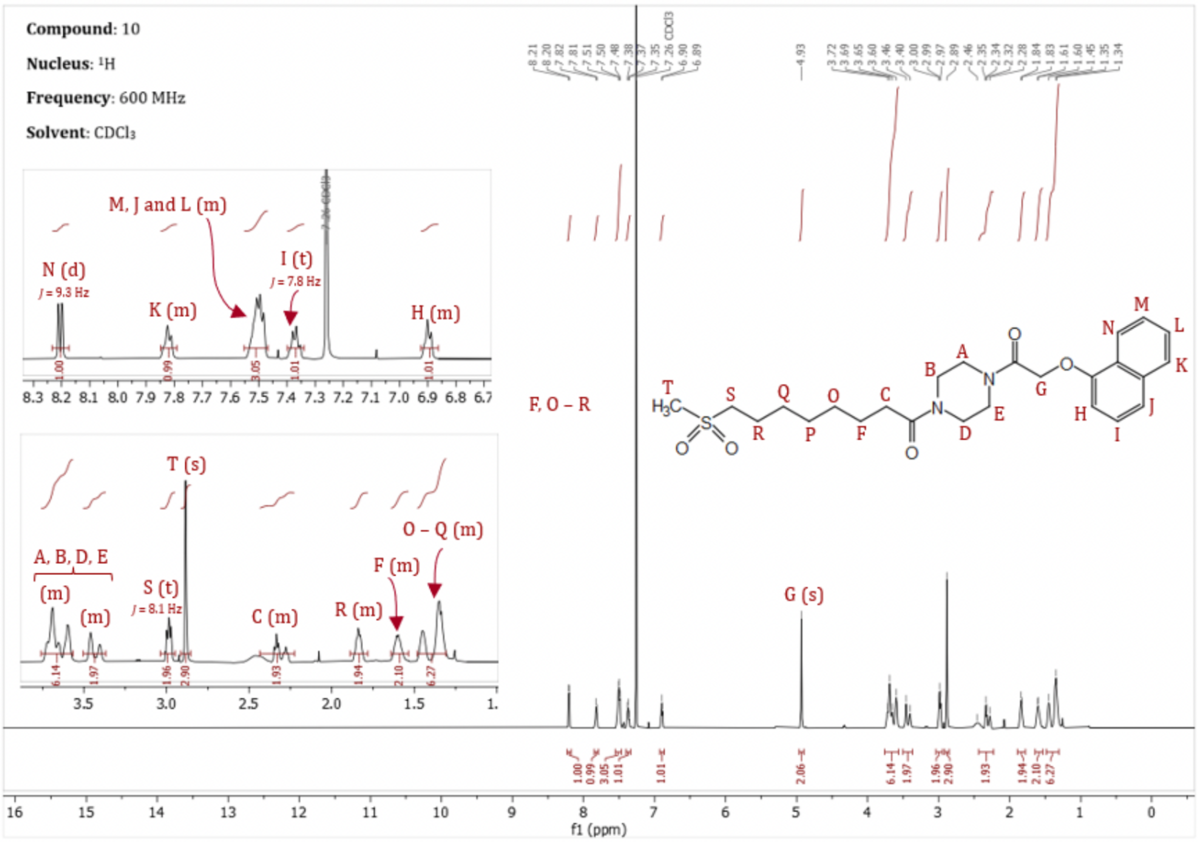

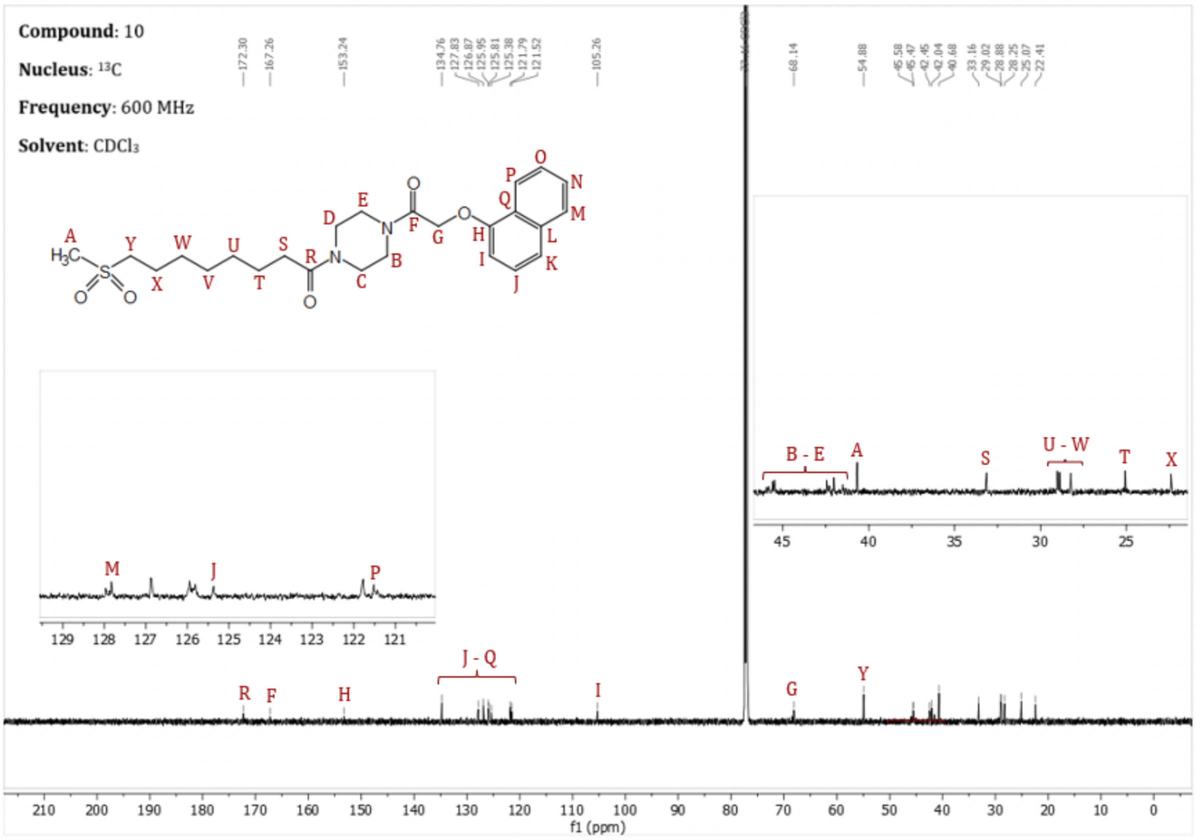

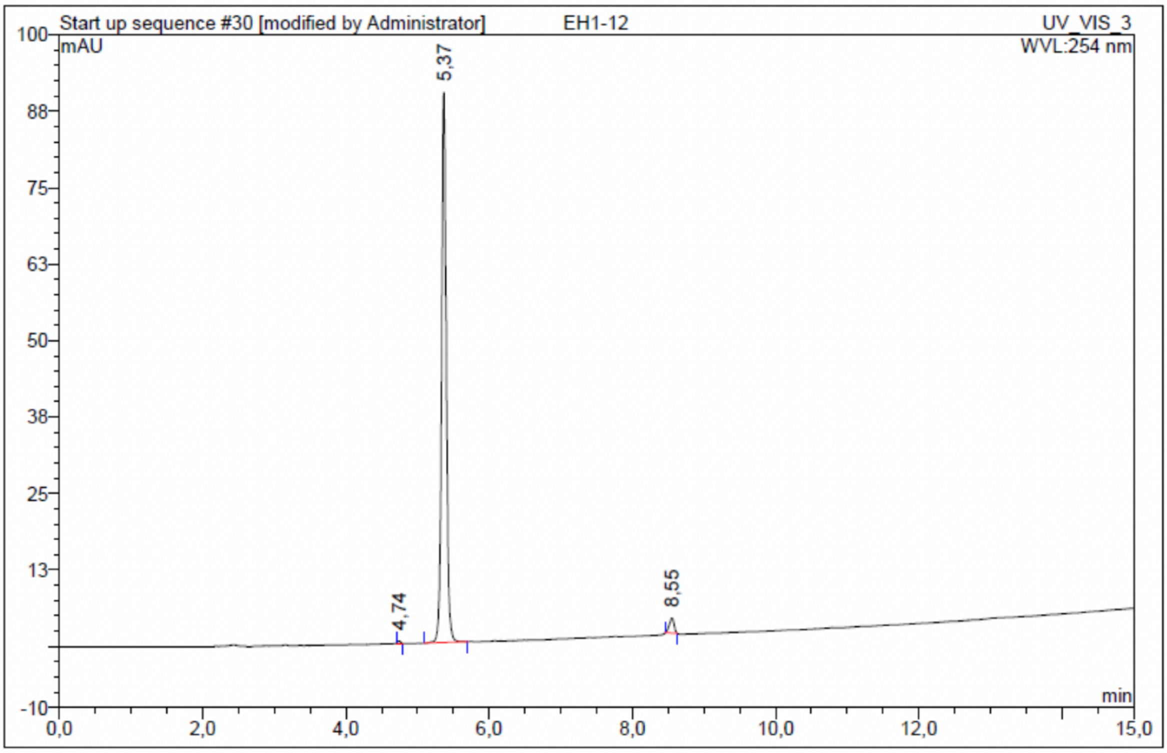

| No. | Ret.Time min | Peak Name | Height mAU | Area mAU*min | Rel.Area % | Amount | Resolution(EP) |
|---|---|---|---|---|---|---|---|
| 1 | 4,74 | n.a. | 0,388 | 0,019 | 0,27 | n.a. | 6,52 |
| 2 | 5,37 | n.a. | 89,957 | 6,681 | 97,02 | n.a. | 27,39 |
| 3 | 8,55 | n.a. | 2,483 | 0,187 | 2,71 | n.a. | n.a. |
| Total: | | | 92,828 | 6,887 | 100,00 | 0,000 | |

#### Compound 48 - TUG-2762

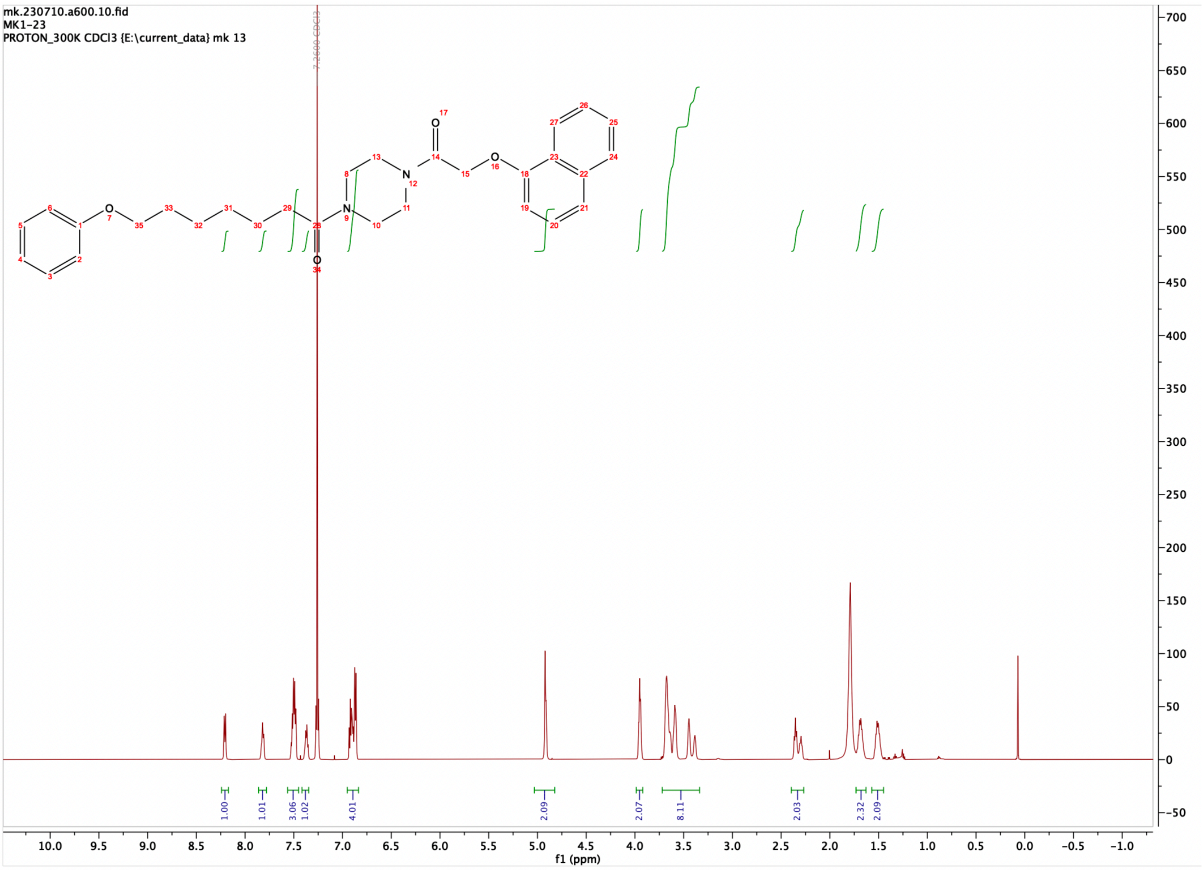

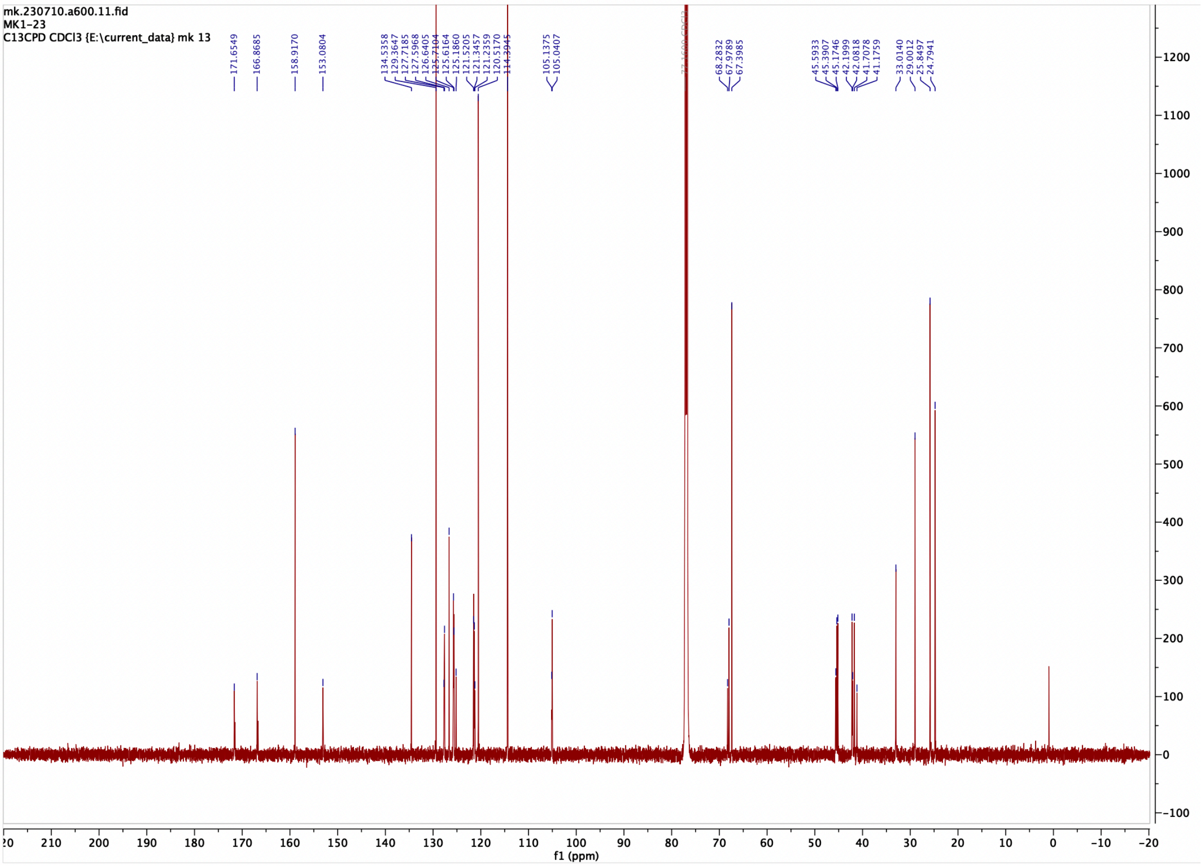

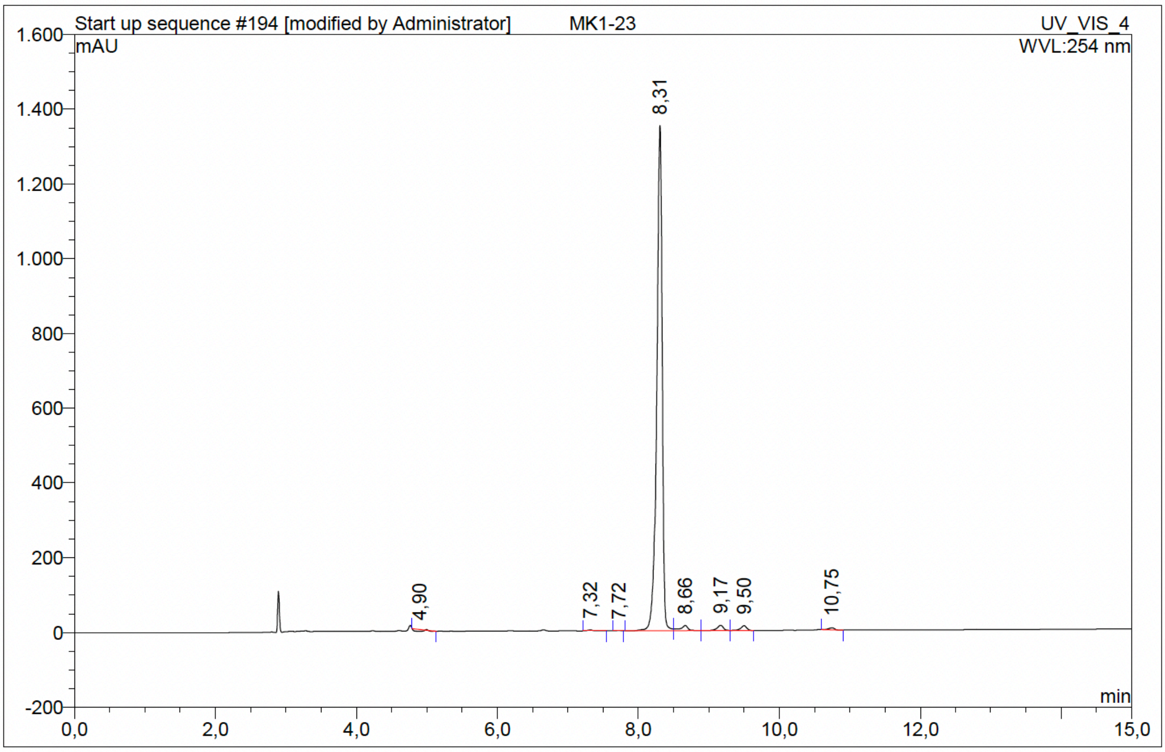

| No. | Ret.Time min | Peak Name | Height mAU | Area mAU*min | Rel.Area % | Amount | Resolution(EP) |
|---|---|---|---|---|---|---|---|
| 1 | 4,90 | n.a. | 5,035 | 0,873 | 0,68 | n.a. | 12,26 |
| 2 | 7,32 | n.a. | 2,620 | 0,235 | 0,18 | n.a. | 3,44 |
| 3 | 7,72 | n.a. | 0,811 | 0,058 | 0,05 | n.a. | 4,69 |
| 4 | 8,31 | n.a. | 1350,972 | 121,699 | 95,25 | n.a. | 2,50 |
| 5 | 8,66 | n.a. | 13,639 | 1,720 | 1,35 | n.a. | 3,38 |
| 6 | 9,17 | n.a. | 13,856 | 1,432 | 1,12 | n.a. | 2,31 |
| 7 | 9,50 | n.a. | 12,857 | 1,256 | 0,98 | n.a. | 8,58 |
| 8 | 10,75 | n.a. | 5,744 | 0,495 | 0,39 | n.a. | n.a. |
| Total: | | | 1405,534 | 127,768 | 100,00 | 0,000 | |

### Compound 49 - TUG-2763

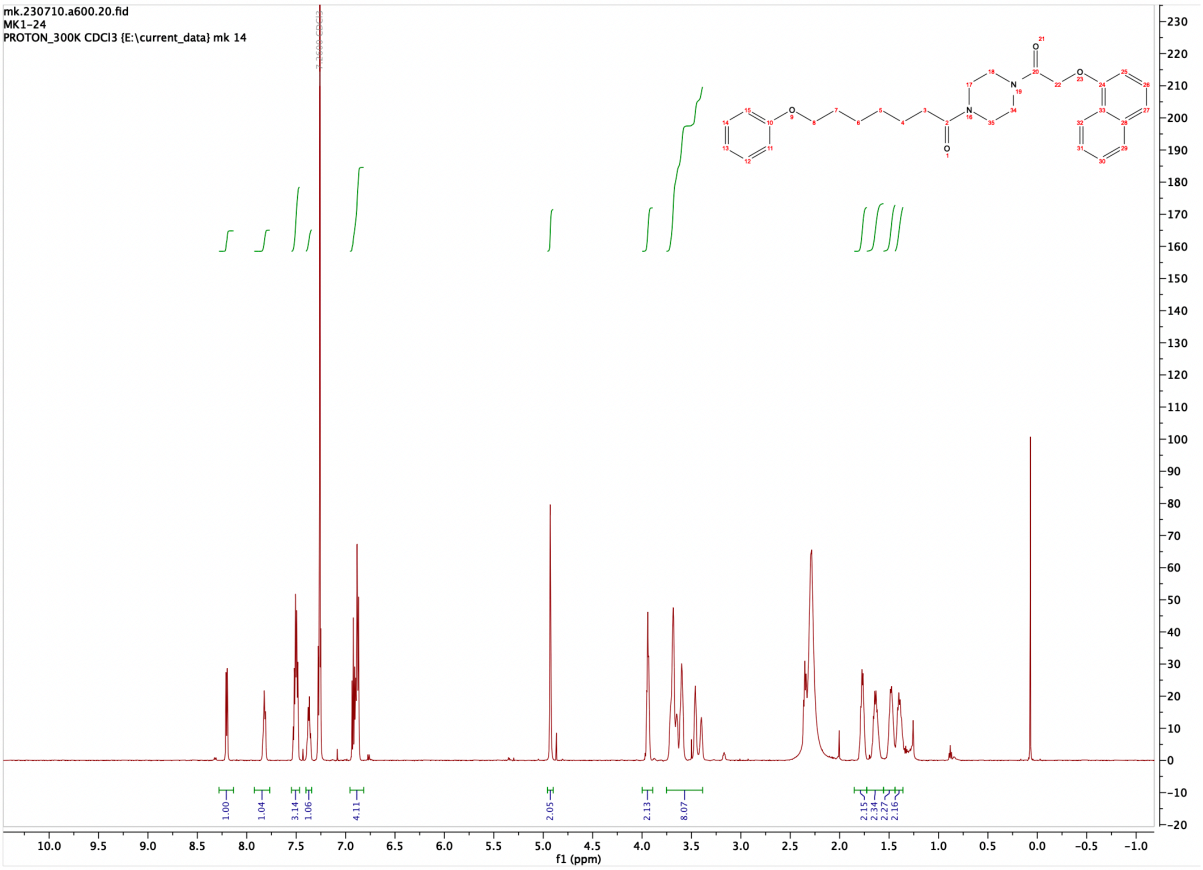

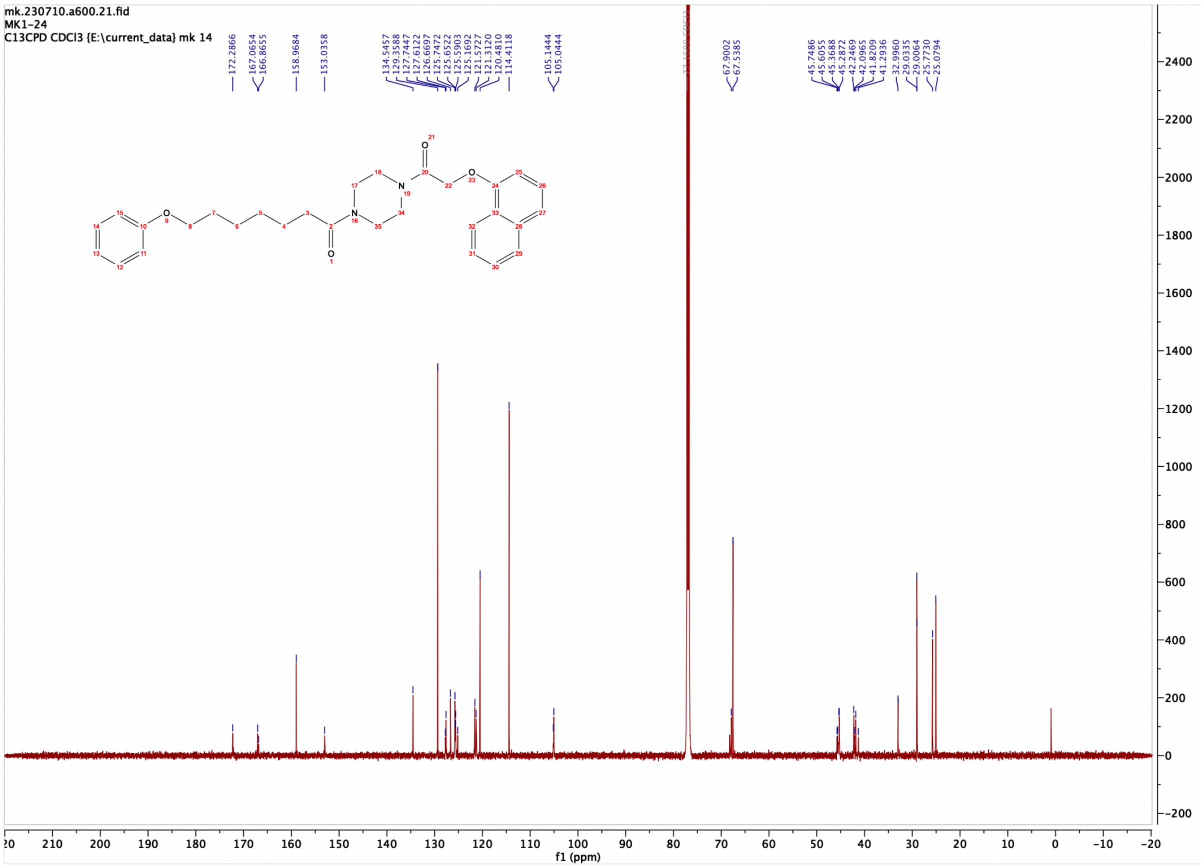

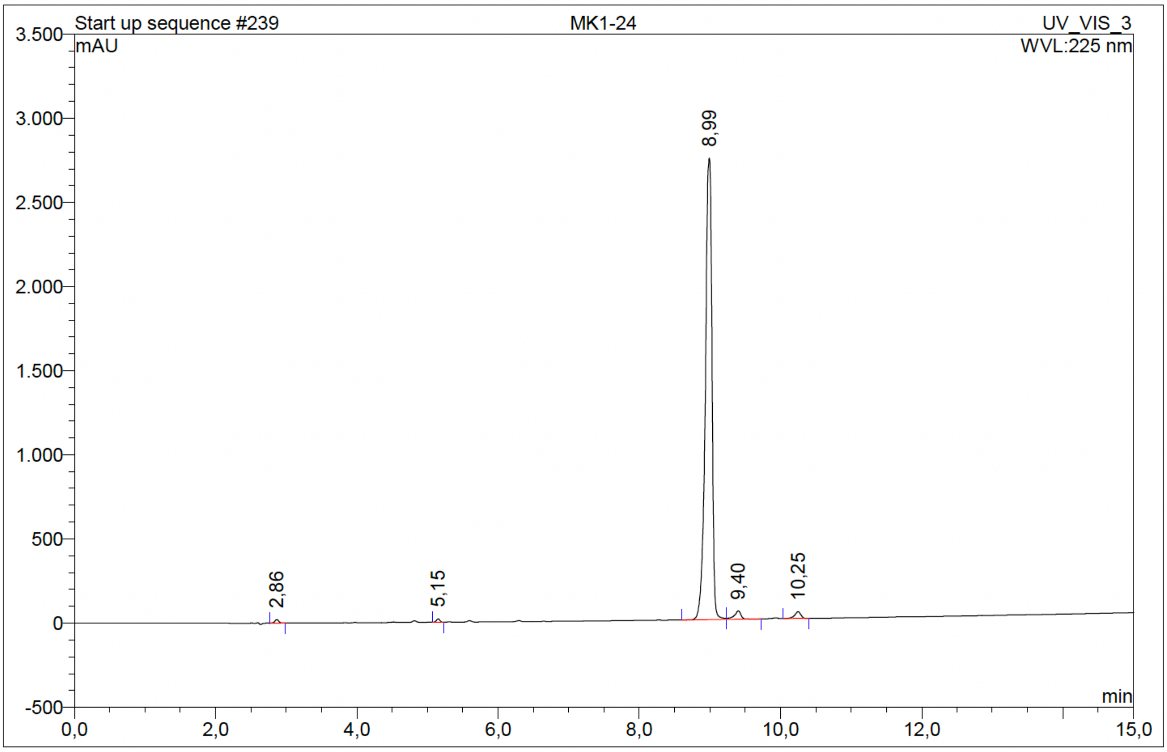

| No. | Ret.Time min | Peak Name | Height mAU | Area mAU*min | Rel.Area % | Amount | Resolution(EP) |
|---|---|---|---|---|---|---|---|
| 1 | 2,86 | n.a. | 21,007 | 1,332 | 0,44 | n.a. | 23,84 |
| 2 | 5,15 | n.a. | 20,185 | 1,175 | 0,38 | n.a. | 30,31 |
| 3 | 8,99 | n.a. | 2741,331 | 293,475 | 95,99 | n.a. | 2,61 |
| 4 | 9,40 | n.a. | 49,900 | 5,619 | 1,84 | n.a. | 5,61 |
| 5 | 10,25 | n.a. | 40,852 | 4,131 | 1,35 | n.a. | n.a. |
| Total: | | | 2873,274 | 305,732 | 100,00 | 0,000 | |

Compound 50 - TUG-2616

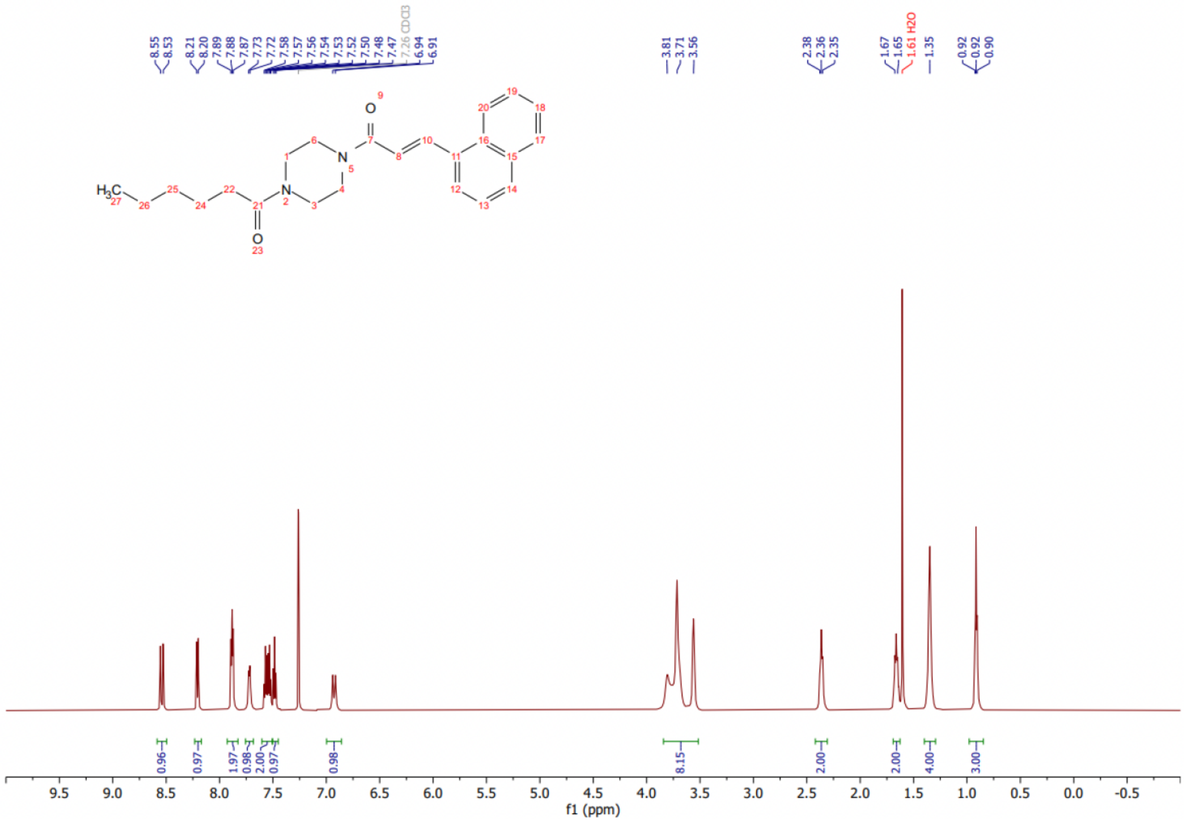

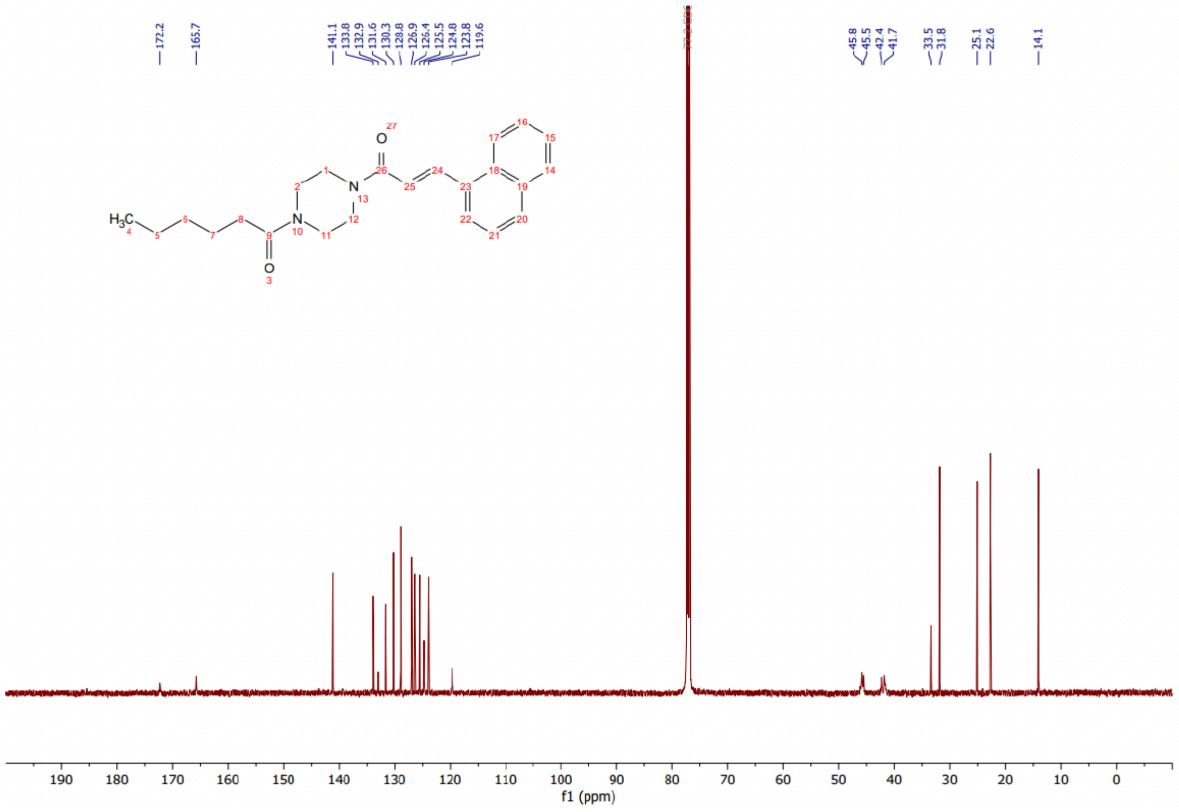

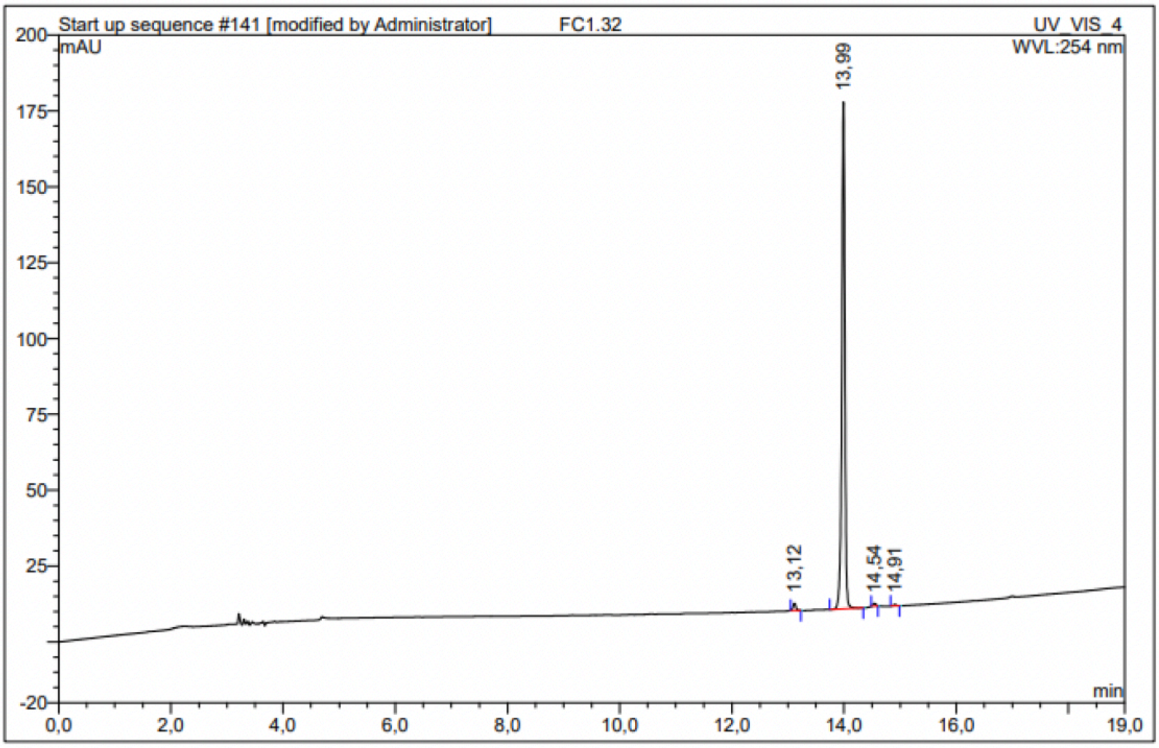

| No. | Ret.Time min | Peak Name | Height mAU | Area mAU*min | Rel.Area % | Amount | Resolution(EP) |
|---|---|---|---|---|---|---|---|
| 1 | 13,12 | n.a. | 2,355 | 0,133 | 1,23 | n.a. | 9,74 |
| 2 | 13,99 | n.a. | 167,007 | 10,632 | 98,06 | n.a. | 5,84 |
| 3 | 14,54 | n.a. | 0,872 | 0,050 | 0,46 | n.a. | 3,89 |
| 4 | 14,91 | n.a. | 0,451 | 0,027 | 0,25 | n.a. | n.a. |
| Total: | | | 170,685 | 10,842 | 100,00 | 0,000 | |

Compound 51 - TUG-2778

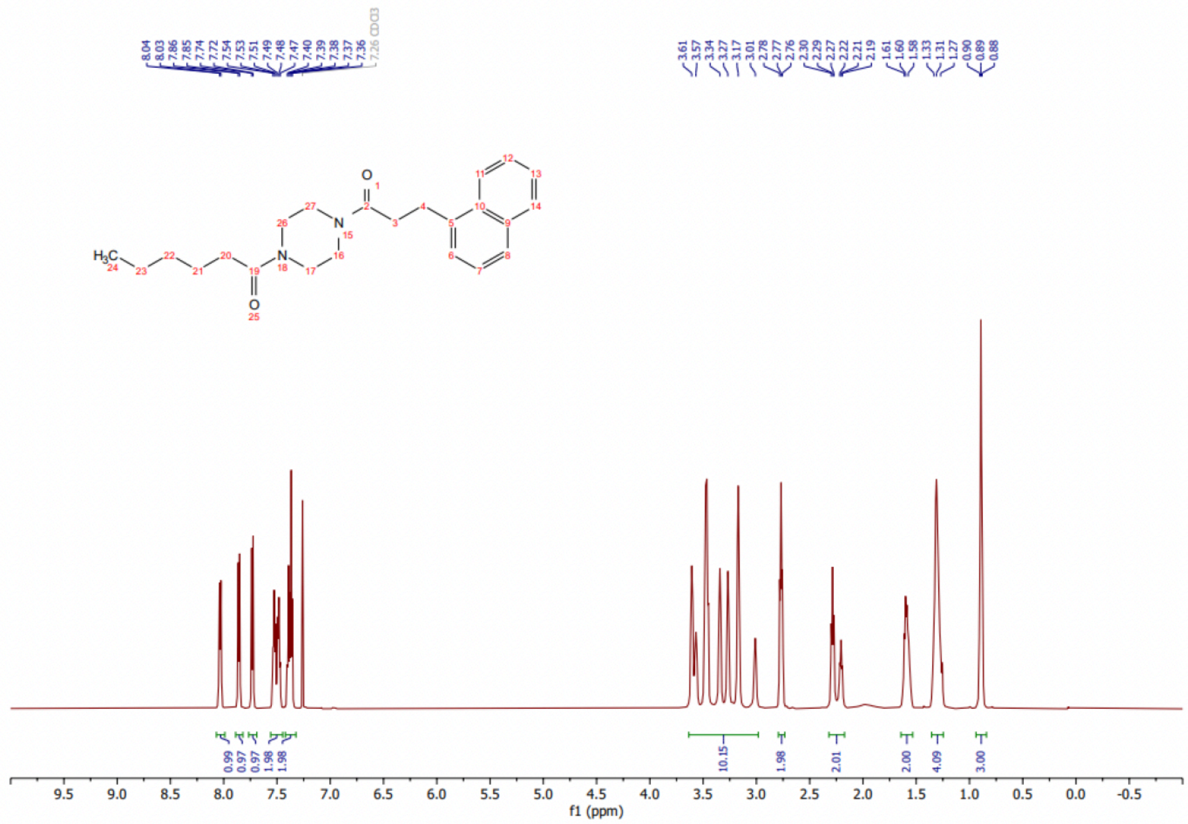

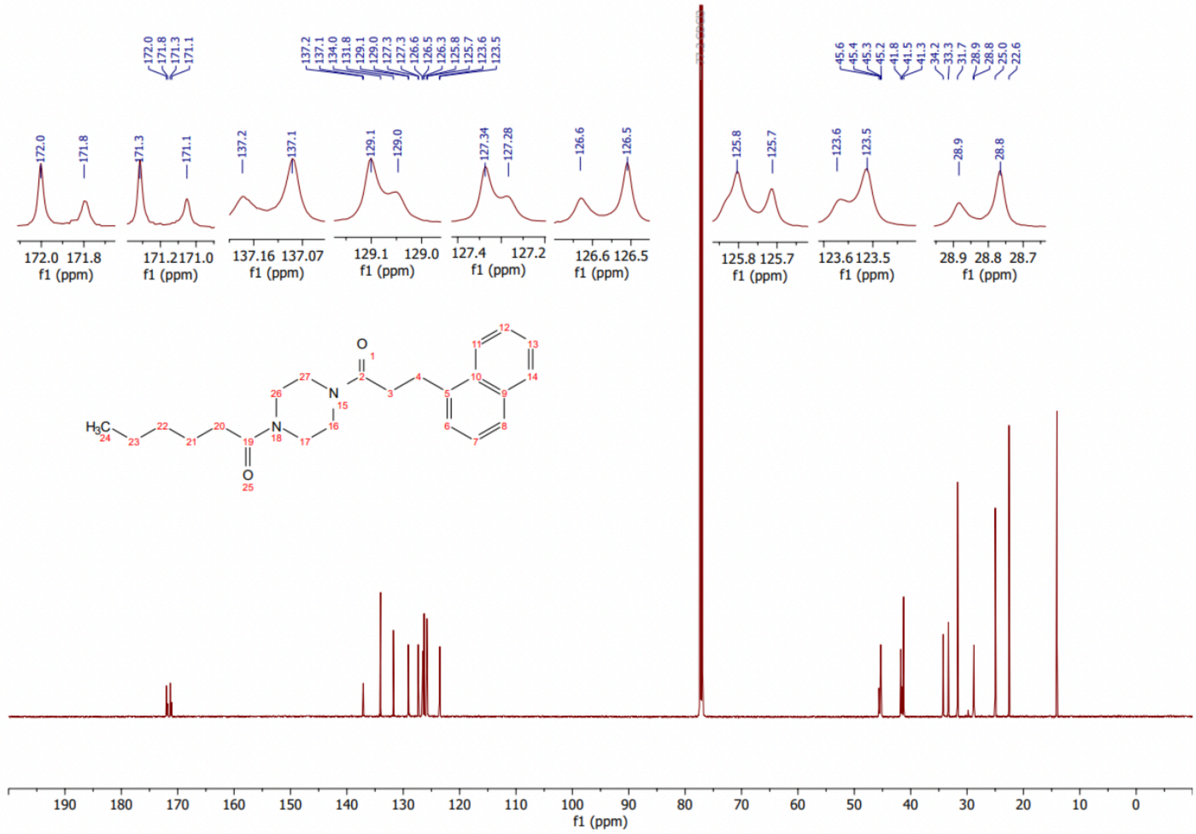

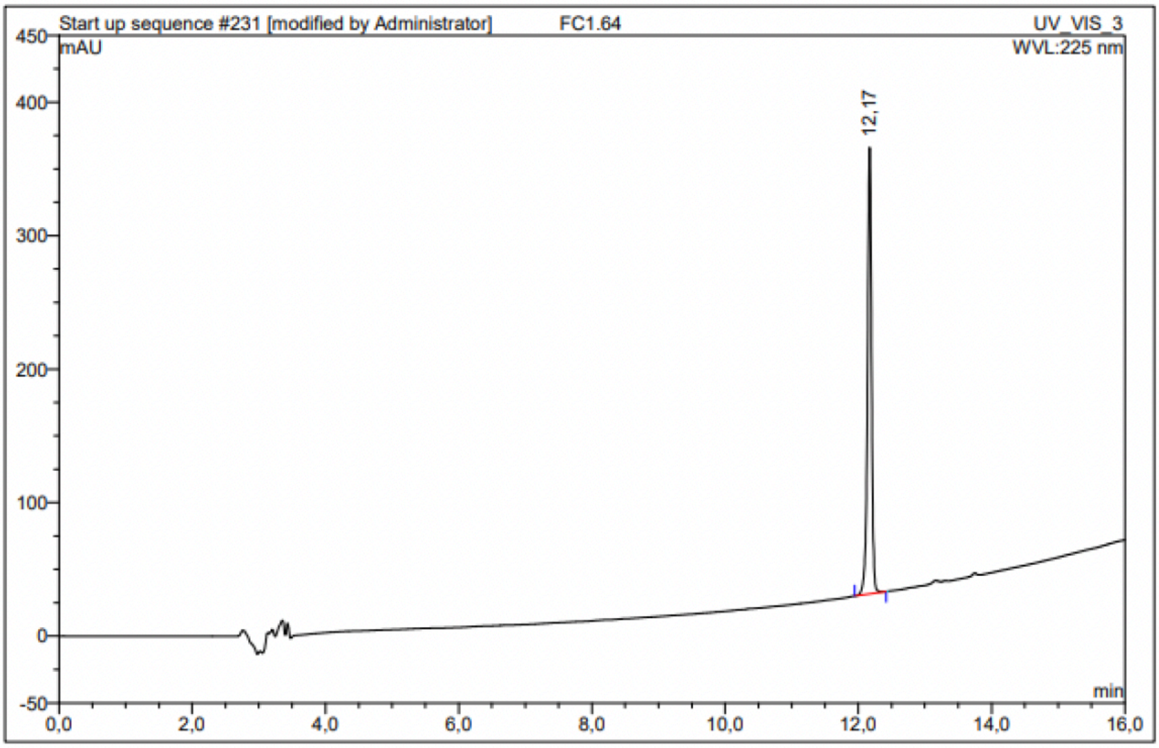

| No. | Ret.Time min | Peak Name | Height mAU | Area mAU*min | Rel.Area % | Amount | Resolution(EP) |
|---|---|---|---|---|---|---|---|
| 1 | 12,17 | n.a. | 334,373 | 23,998 | 100,00 | n.a. | n.a. |
| Total: | | | 334,373 | 23,998 | 100,00 | 0,000 | |

Compound 52 - TUG-2370

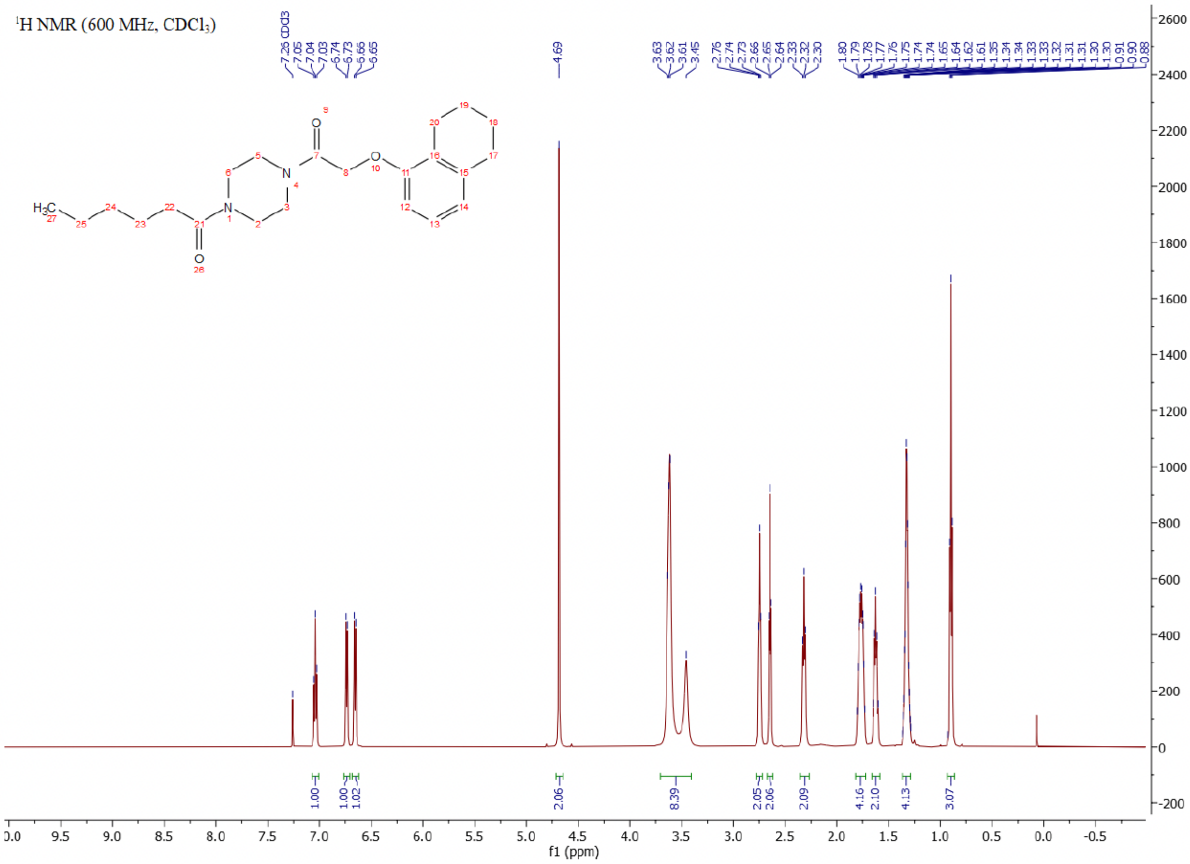

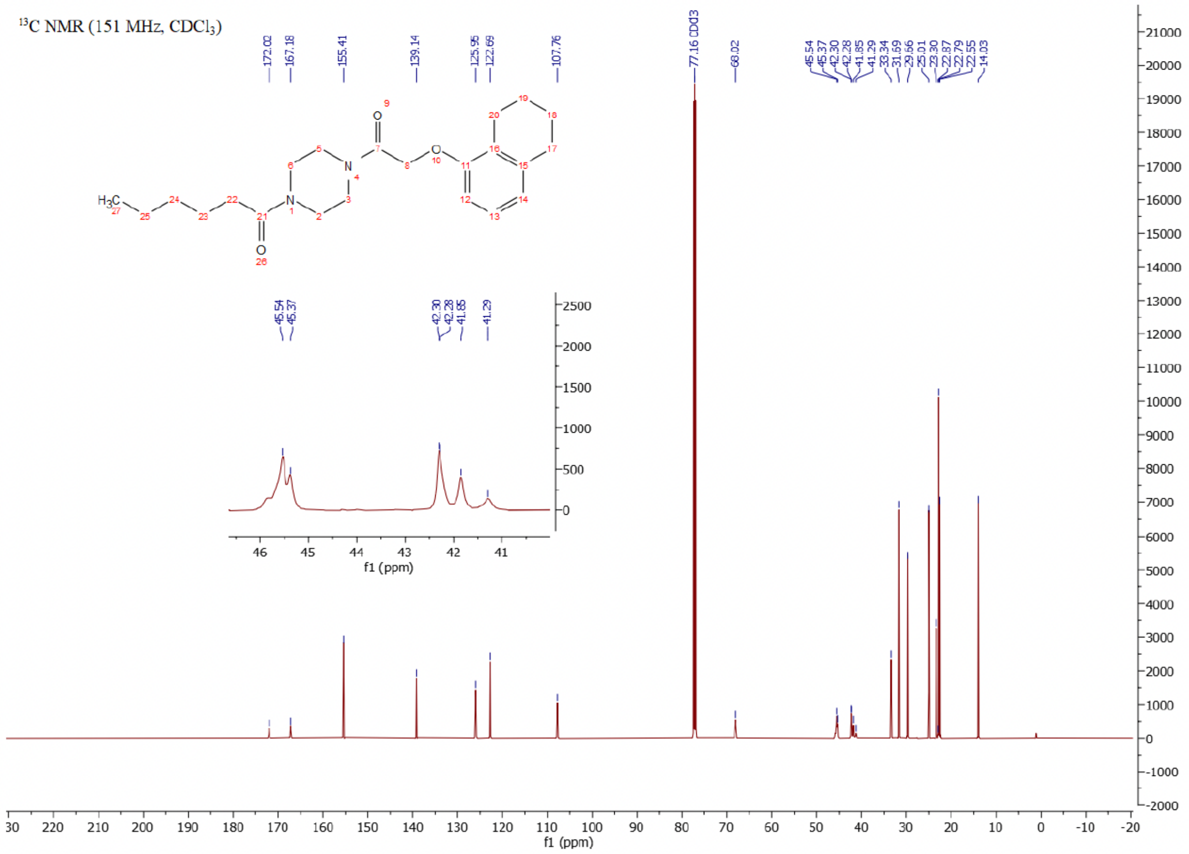

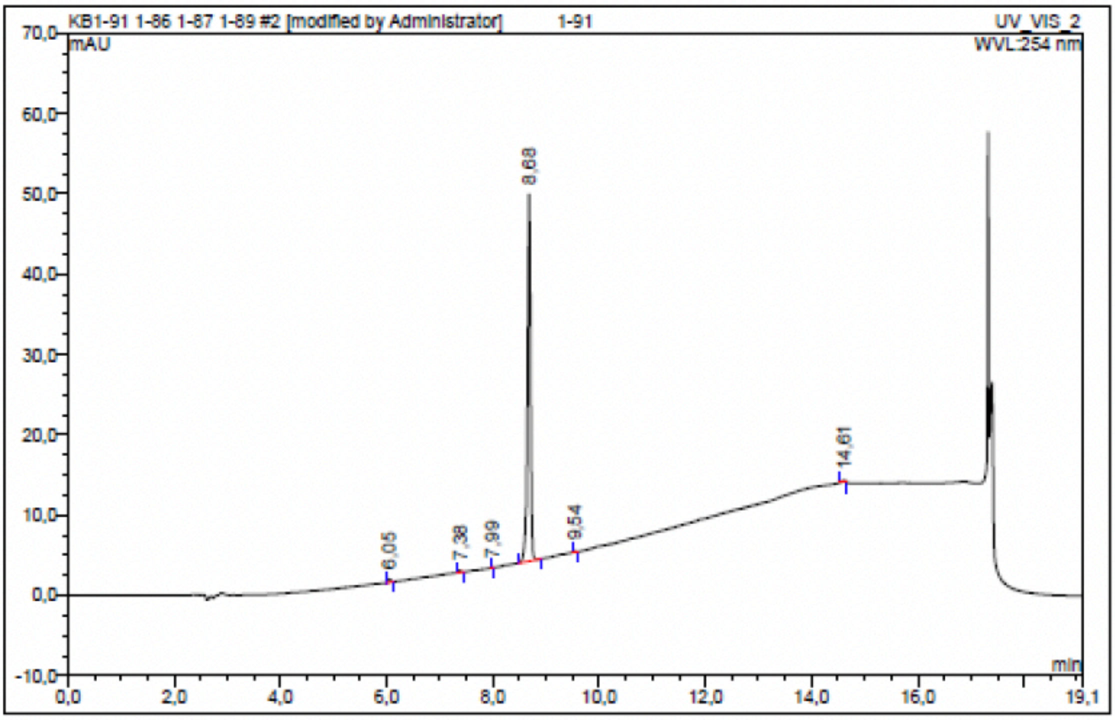

| No. | Ret.Time min | Peak Name | Height mAU | Area mAU*min | Rel.Area % | Amount | Resolution(EP) |
|---|---|---|---|---|---|---|---|
| 1 | 6,05 | n.a. | 0,433 | 0,022 | 0,68 | n.a. | 16,42 |
| 2 | 7,38 | n.a. | 0,252 | 0,012 | 0,39 | n.a. | 9,02 |
| 3 | 7,99 | n.a. | 0,033 | 0,001 | 0,03 | n.a. | 8,94 |
| 4 | 8,68 | n.a. | 45,700 | 3,104 | 98,09 | n.a. | 9,86 |
| 5 | 9,54 | n.a. | 0,124 | 0,005 | 0,17 | n.a. | 53,01 |
| 6 | 14,61 | n.a. | 0,286 | 0,020 | 0,64 | n.a. | n.a. |
| Total: | | | 46,827 | 3,164 | 100,00 | 0,000 | |

Compound 53 - TUG-2365

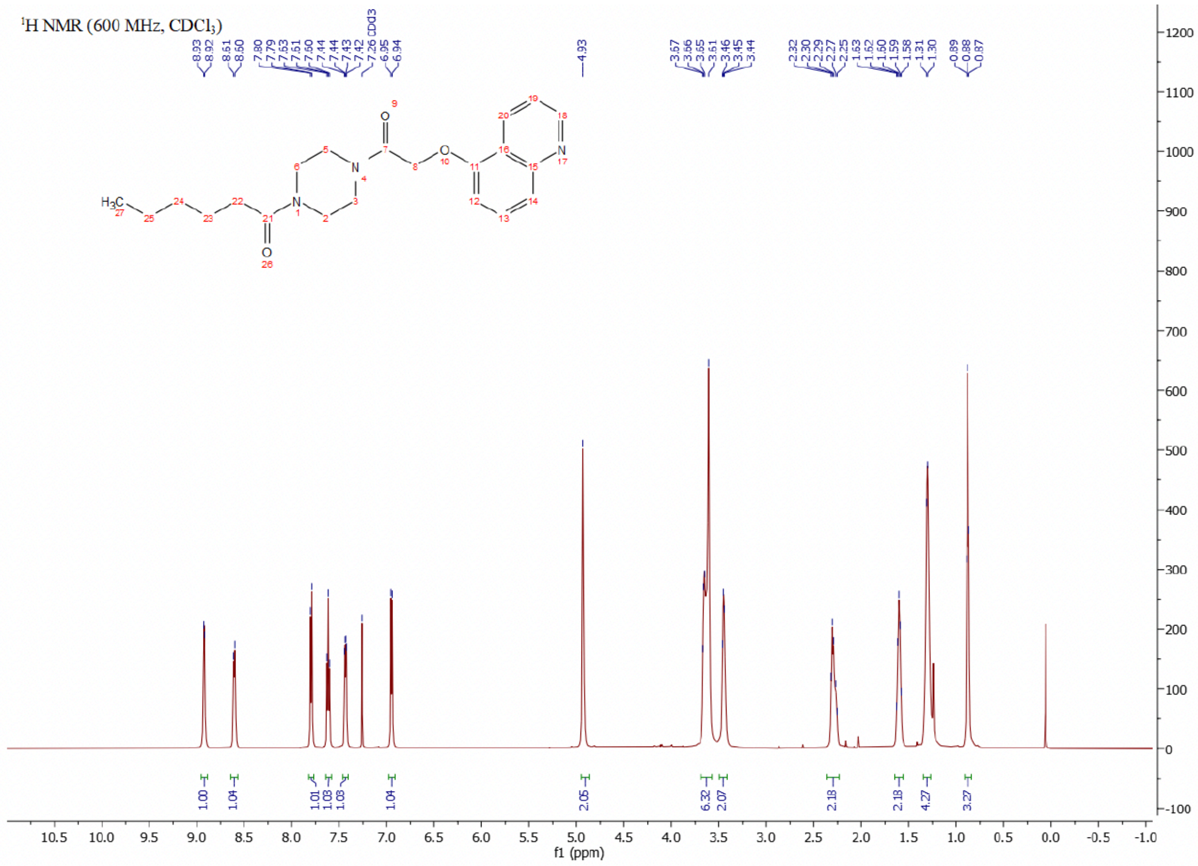

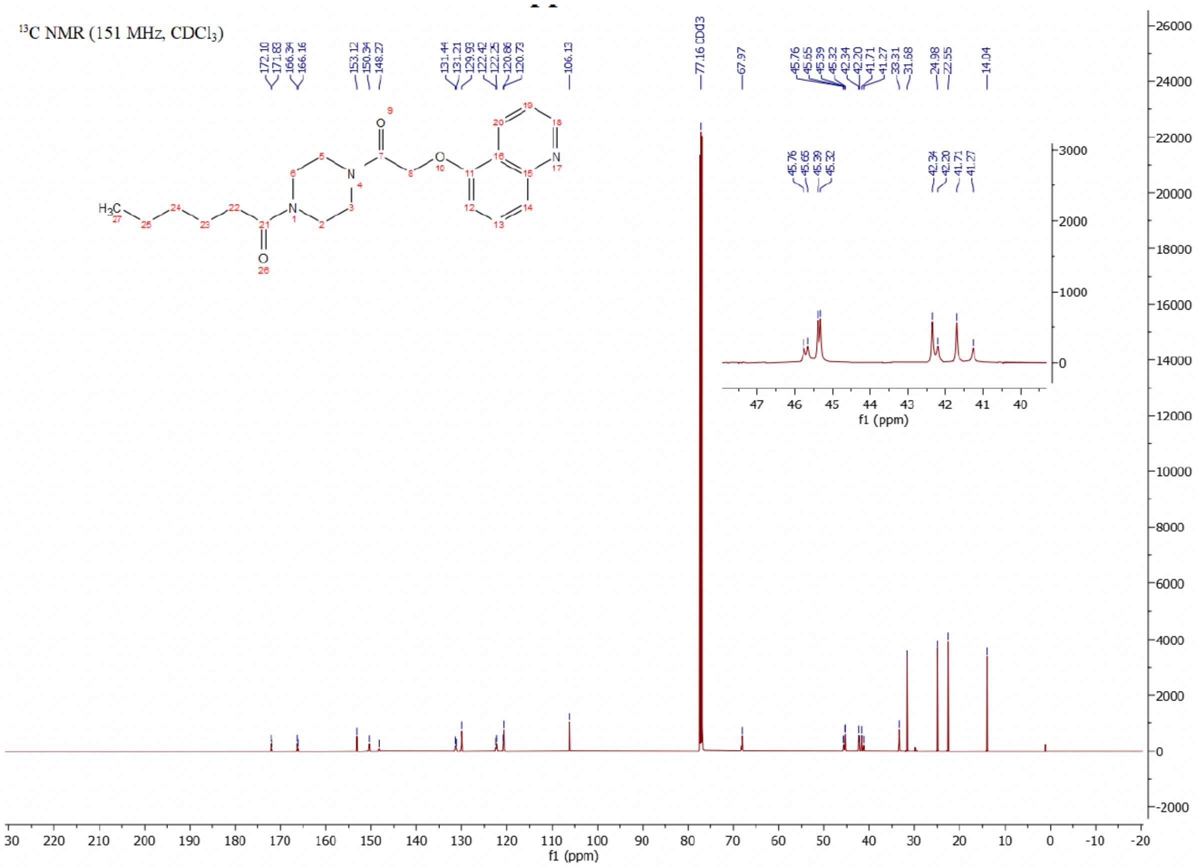

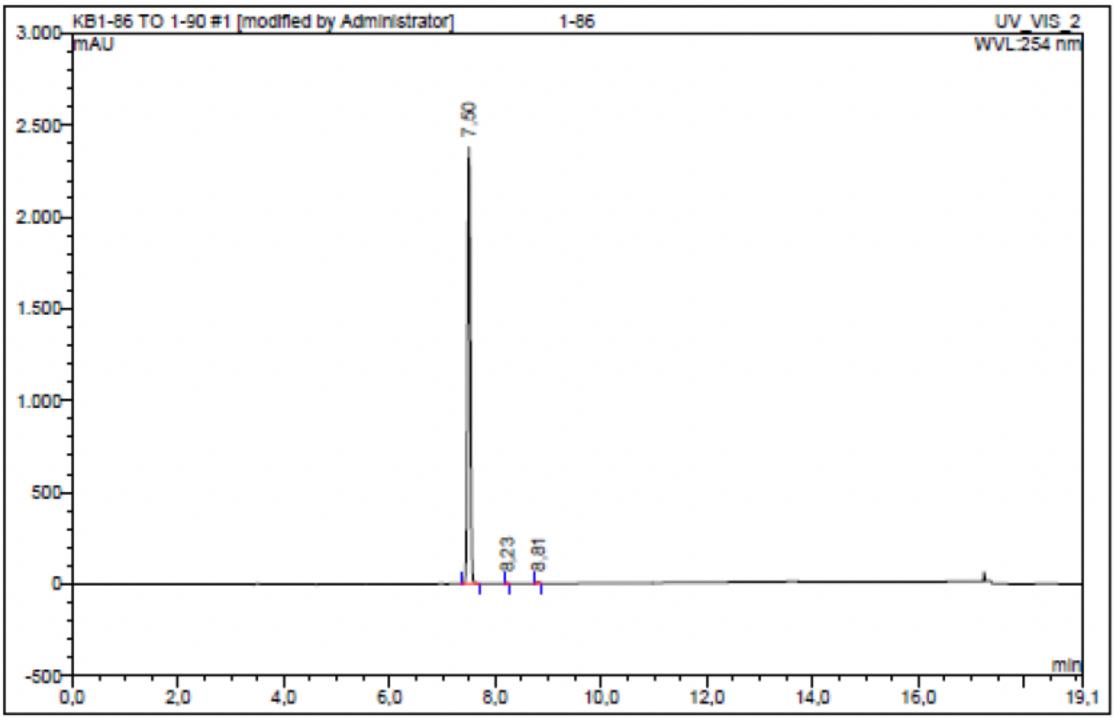

| No. | Ret.Time min | Peak Name | Height mAU | Area mAU*min | Rel.Area % | Amount | Resolution(EP) |
|---|---|---|---|---|---|---|---|
| 1 | 7,50 | n.a. | 2377,503 | 150,020 | 99,65 | n.a. | 8,32 |
| 2 | 8,23 | n.a. | 2,573 | 0,114 | 0,08 | n.a. | 7,55 |
| 3 | 8,81 | n.a. | 8,423 | 0,419 | 0,28 | n.a. | n.a. |
| Total: | | | 2388,498 | 150,553 | 100,00 | 0,000 | |

Compound 54 - TUG-2367

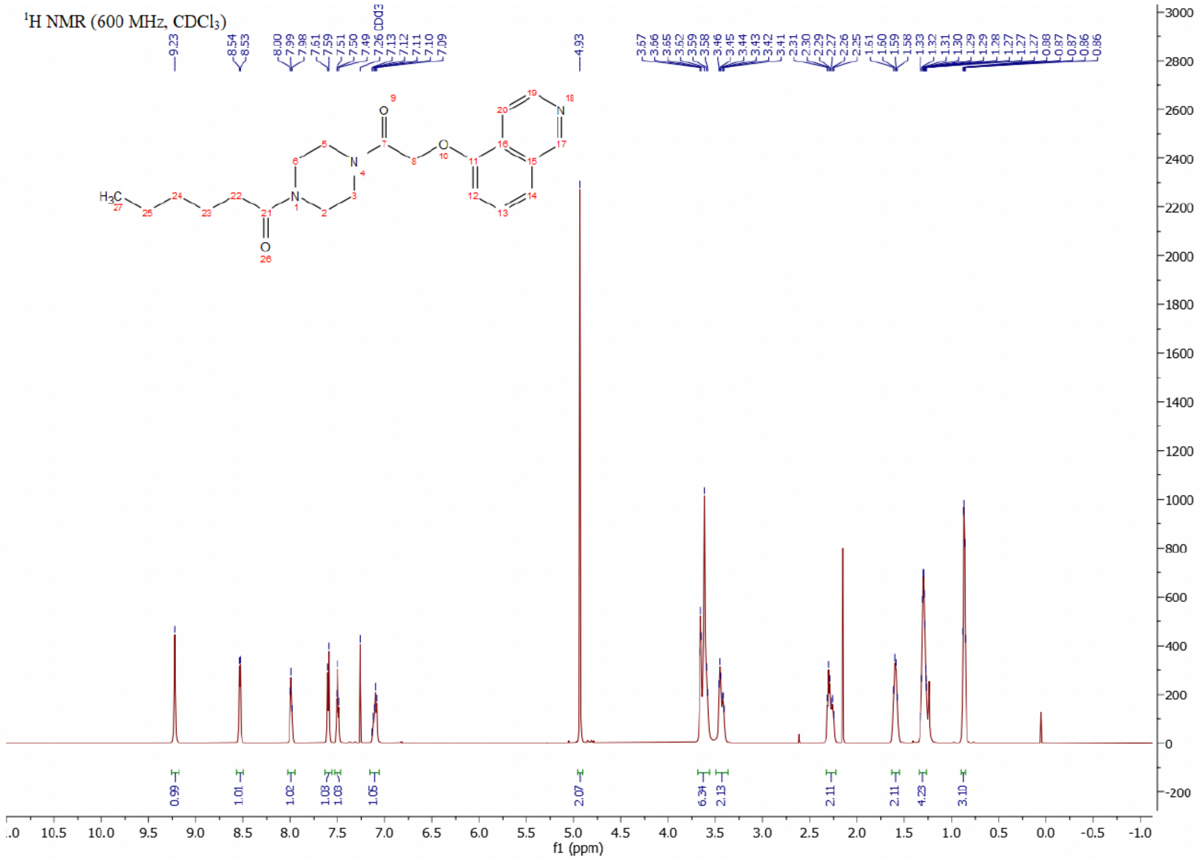

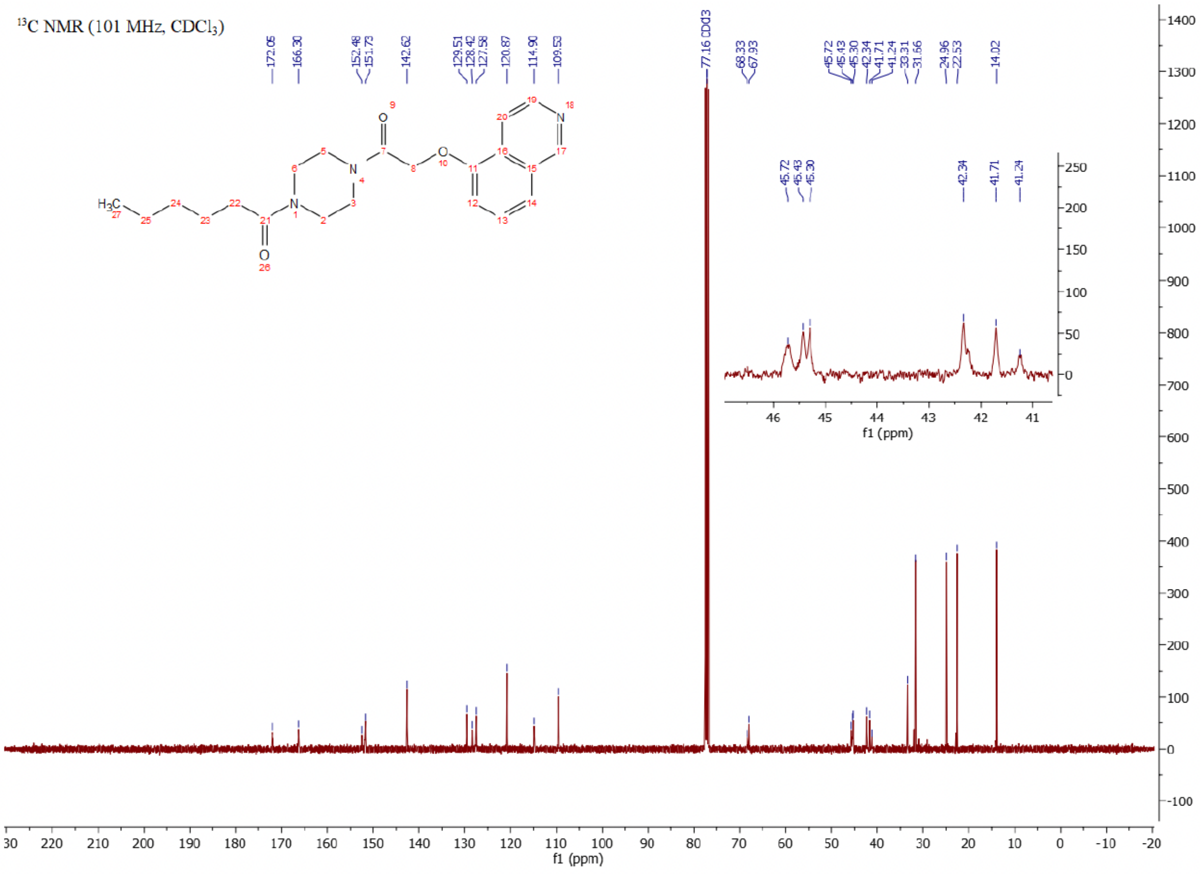

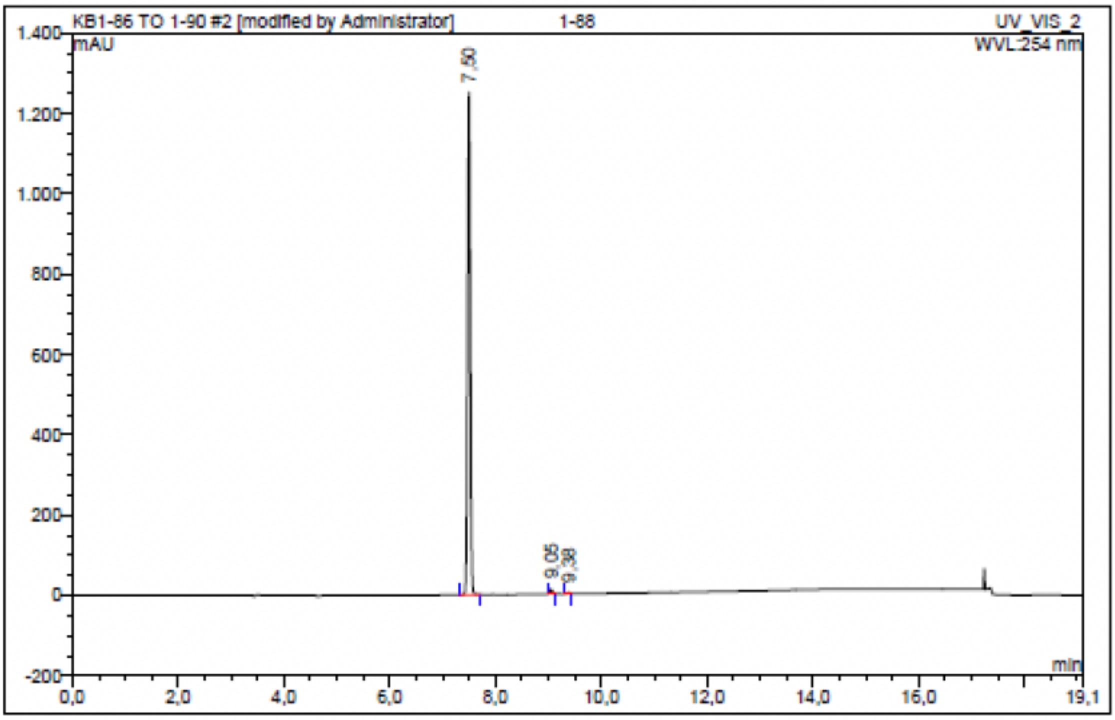

| No. | Ret.Time min | Peak Name | Height mAU | Area mAU*min | Rel.Area % | Amount | Resolution(EP) |
|---|---|---|---|---|---|---|---|
| 1 | 7,50 | n.a. | 1251,363 | 75,966 | 98,95 | n.a. | 16,76 |
| 2 | 9,05 | n.a. | 11,652 | 0,648 | 0,84 | n.a. | 3,78 |
| 3 | 9,38 | n.a. | 2,932 | 0,159 | 0,21 | n.a. | n.a. |
| Total: | | | 1265,947 | 76,773 | 100,00 | 0,000 | |

Compound 55 - TUG-2369

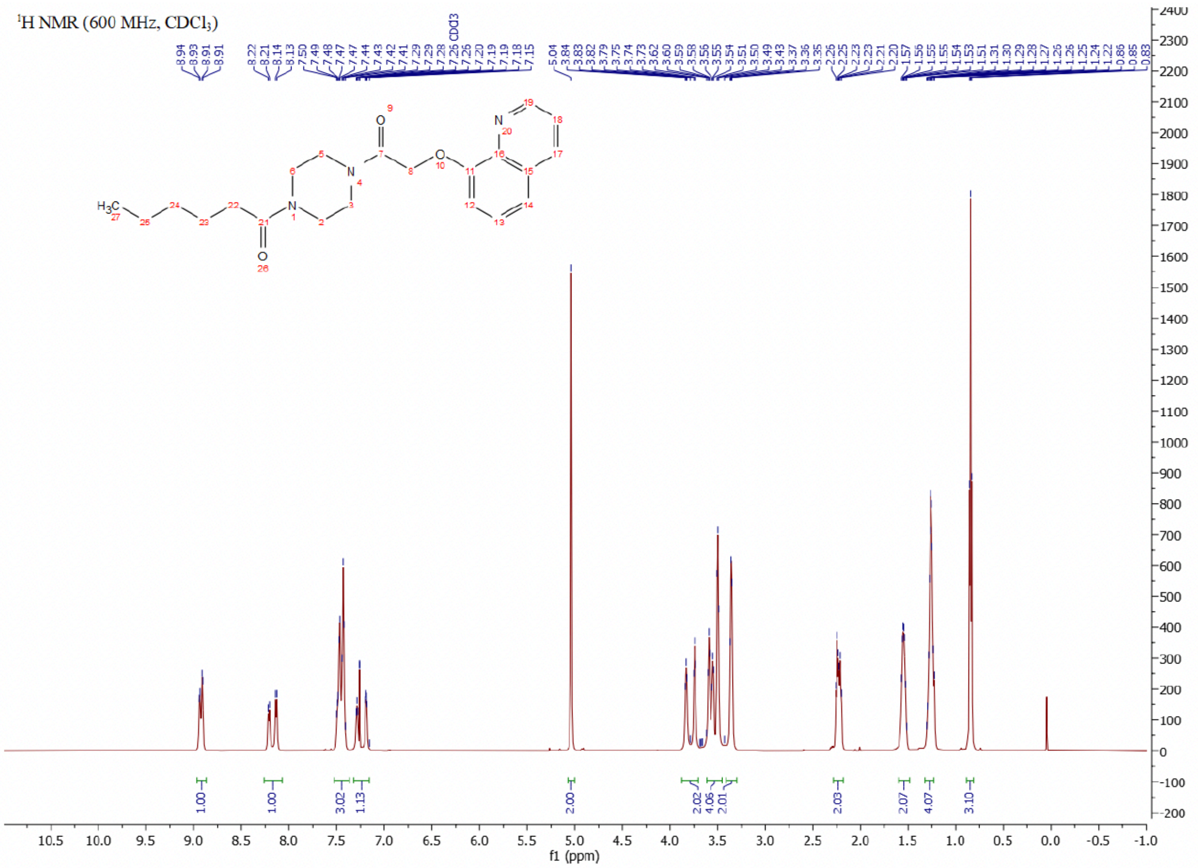

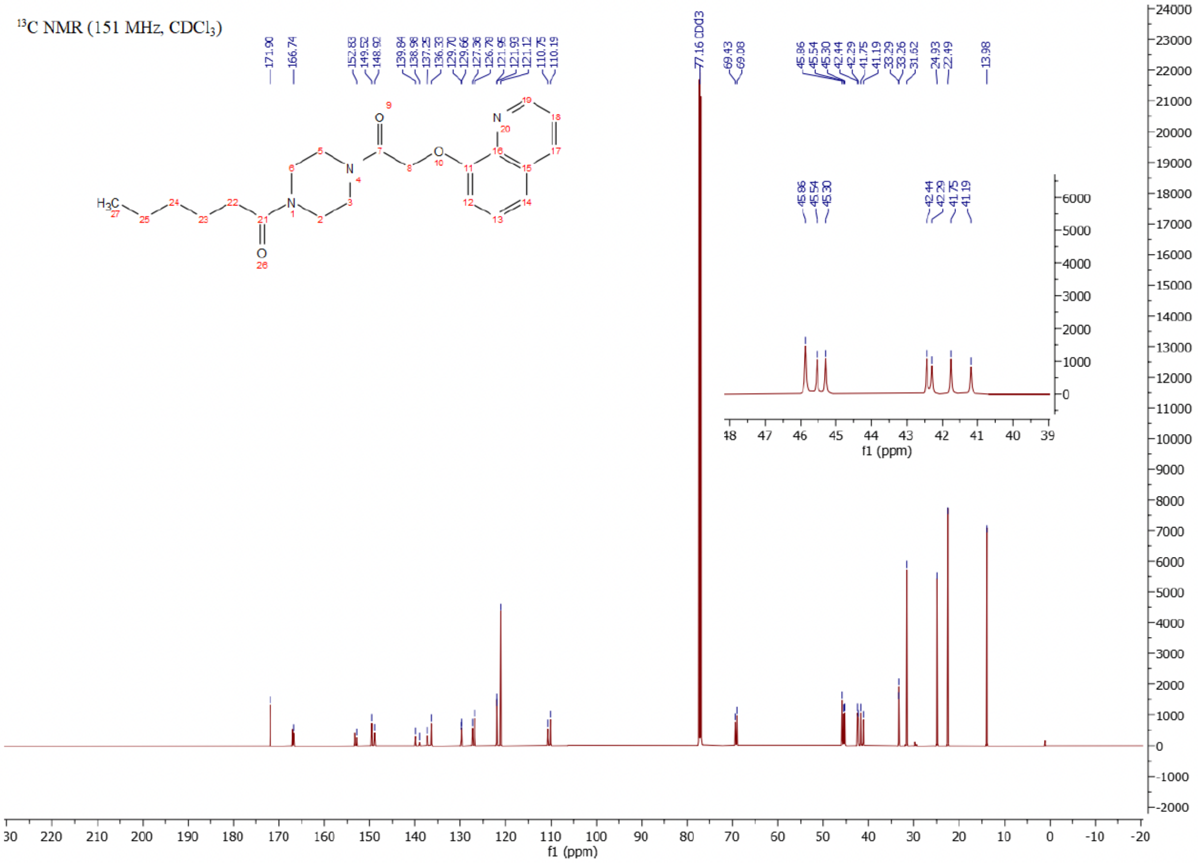

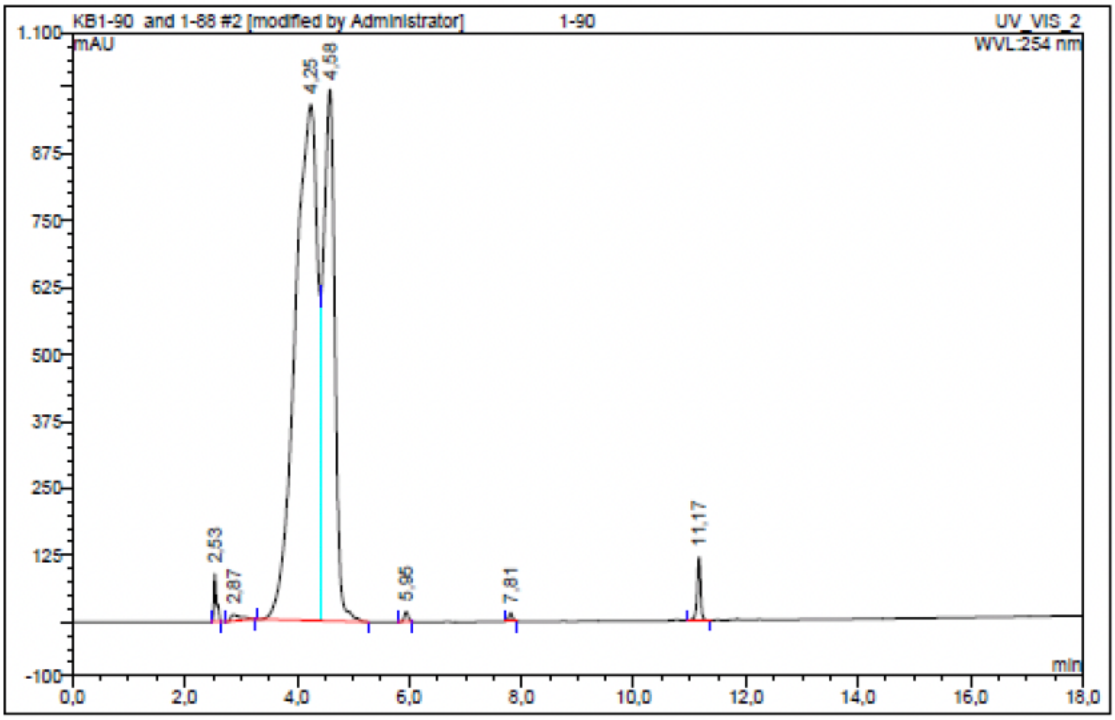

| No. | Ret.Time min | Peak Name | Height mAU | Area mAU*min | Rel.Area % | Amount | Resolution(EP) |
|---|---|---|---|---|---|---|---|
| 1 | 2,53 | n.a. | 88,674 | 5,236 | 0,75 | n.a. | 1,36 |
| 2 | 2,87 | n.a. | 11,199 | 2,507 | 0,36 | n.a. | n.a. |
| 3 | 4,25 | n.a. | 963,860 | 440,716 | 62,84 | n.a. | n.a. |
| 4 | 4,58 | n.a. | 992,073 | 241,811 | 34,48 | n.a. | n.a. |
| 5 | 5,95 | n.a. | 18,594 | 1,503 | 0,21 | n.a. | 15,11 |
| 6 | 7,81 | n.a. | 14,623 | 1,105 | 0,16 | n.a. | 30,12 |
| 7 | 11,17 | n.a. | 118,165 | 8,402 | 1,20 | n.a. | n.a. |
| Total: | | | 2207,188 | 701,280 | 100,00 | 0,000 | |

Compound 56 - TUG-2368

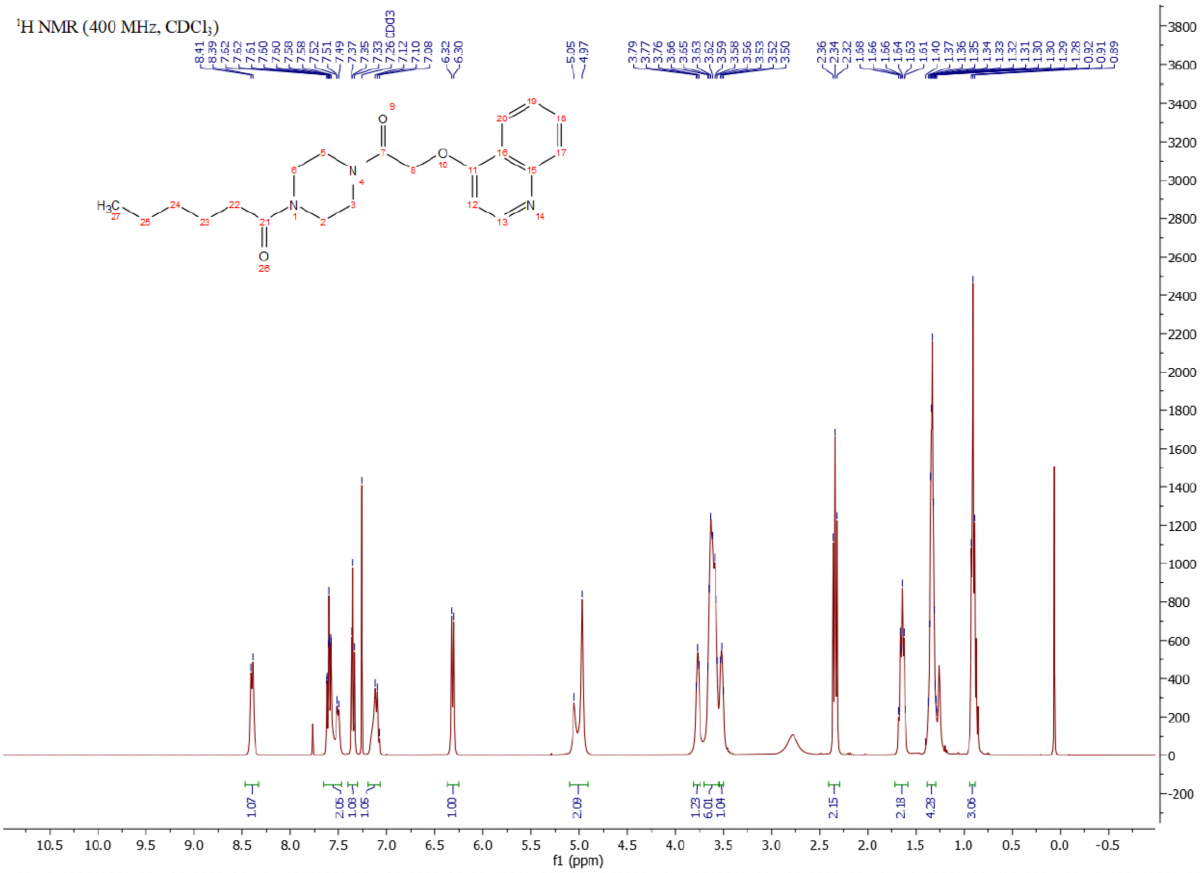

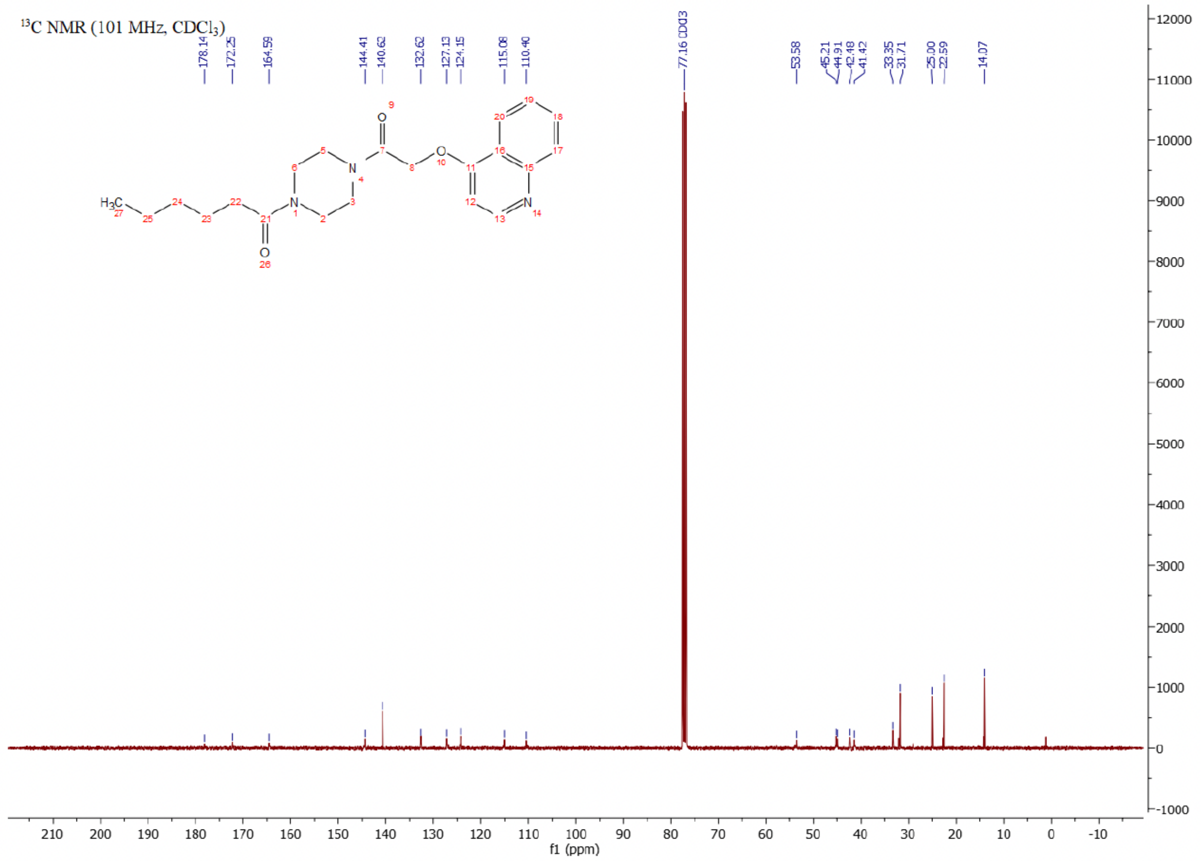

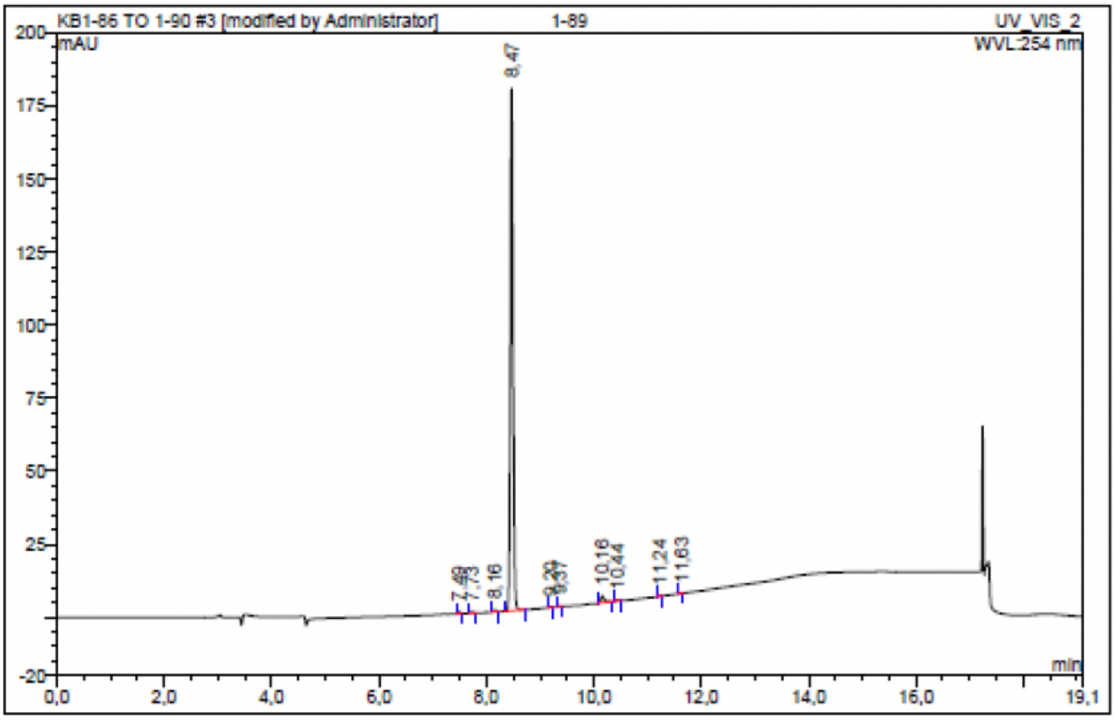

| No. | Ret.Time min | Peak Name | Height mAU | Area mAU*min | Rel.Area % | Amount | Resolution(EP) |
|---|---|---|---|---|---|---|---|
| 1 | 7,49 | n.a. | 0,701 | 0,036 | 0,33 | n.a. | 2,69 |
| 2 | 7,73 | n.a. | 0,616 | 0,034 | 0,31 | n.a. | 4,67 |
| 3 | 8,16 | n.a. | 0,644 | 0,039 | 0,36 | n.a. | 3,35 |
| 4 | 8,47 | n.a. | 178,710 | 10,520 | 96,83 | n.a. | 7,76 |
| 5 | 9,20 | n.a. | 0,206 | 0,011 | 0,11 | n.a. | 2,18 |
| 6 | 9,37 | n.a. | 0,202 | 0,008 | 0,07 | n.a. | 10,04 |
| 7 | 10,16 | n.a. | 2,402 | 0,181 | 1,66 | n.a. | 3,14 |
| 8 | 10,44 | n.a. | 0,346 | 0,018 | 0,17 | n.a. | 9,19 |
| 9 | 11,24 | n.a. | 0,196 | 0,011 | 0,10 | n.a. | 4,10 |
| 10 | 11,63 | n.a. | 0,099 | 0,006 | 0,05 | n.a. | n.a. |
| Total: | | | 184,121 | 10,864 | 100,00 | 0,000 | |

Compound 57 - TUG-2366

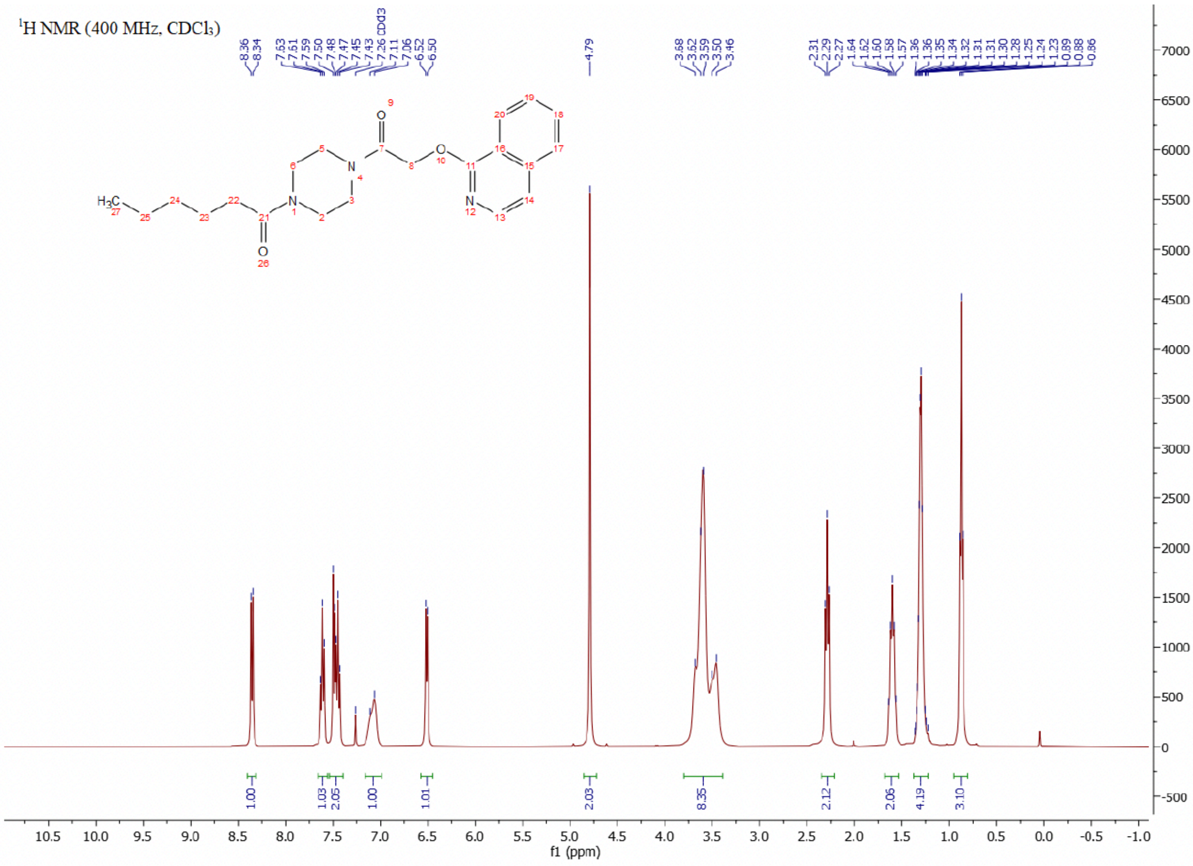

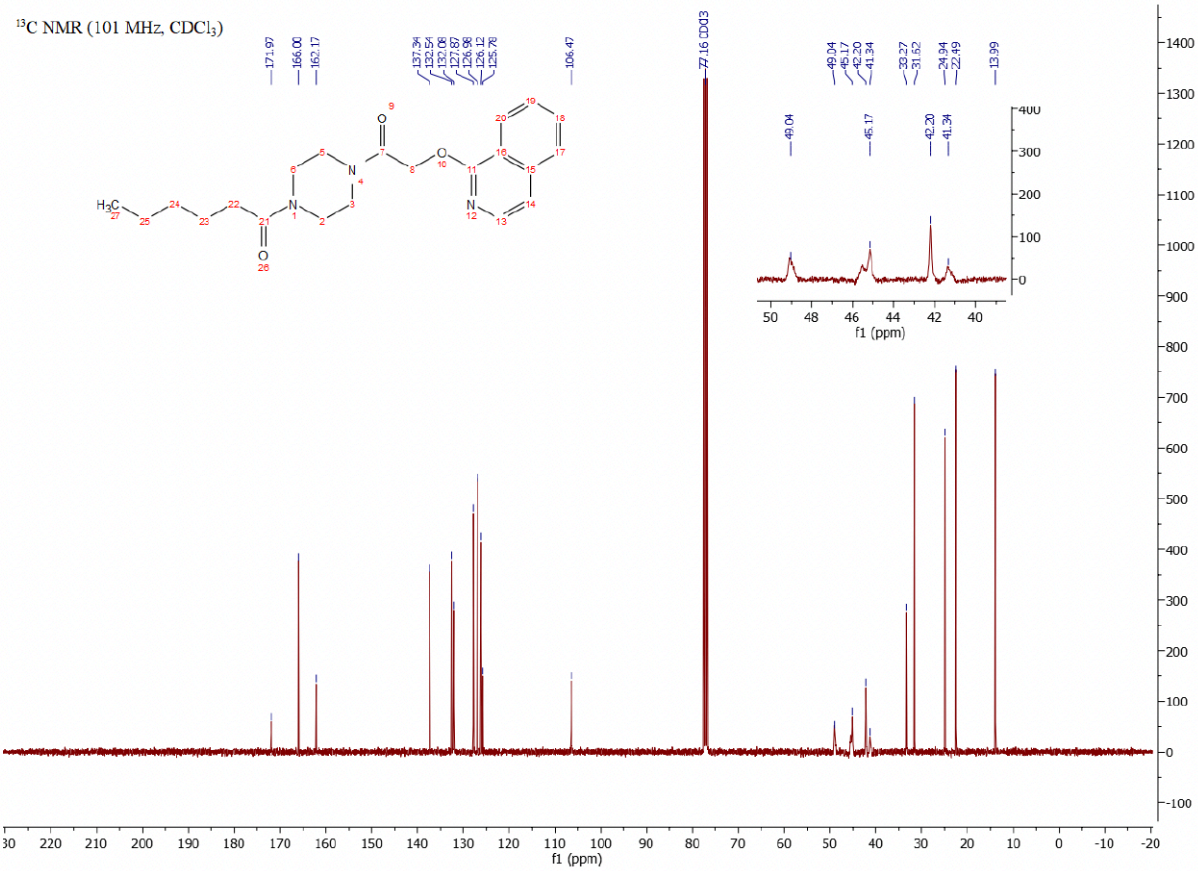

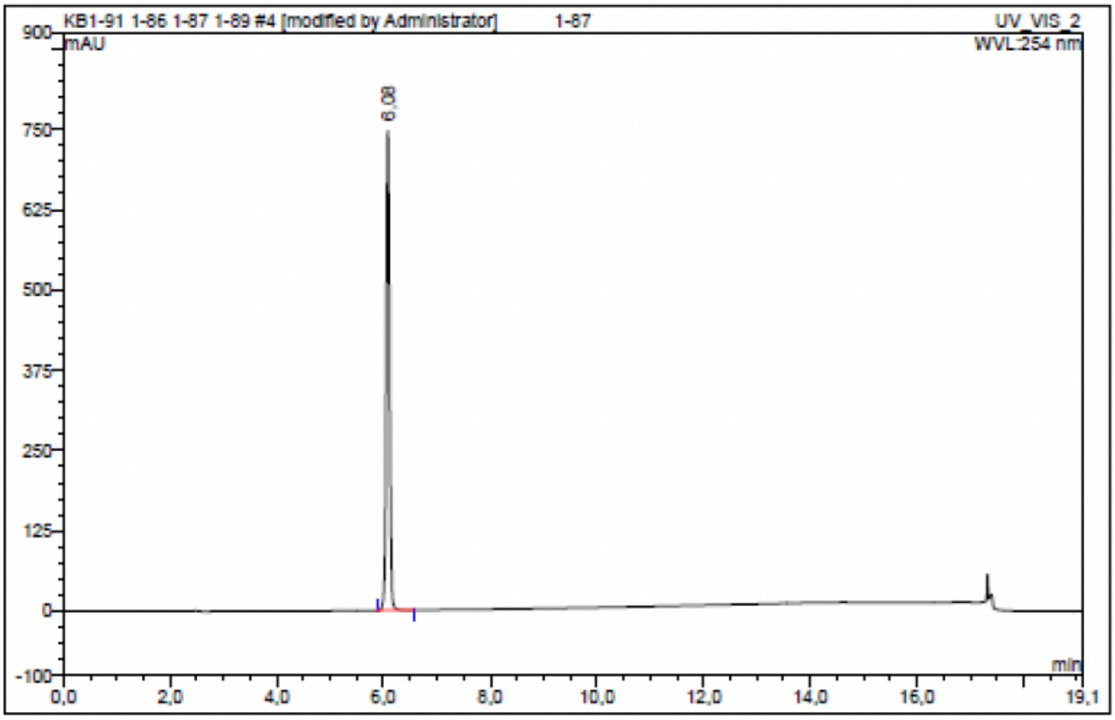

| No. | Ret.Time min | Peak Name | Height mAU | Area mAU*min | Rel.Area % | Amount | Resolution(EP) |
|---|---|---|---|---|---|---|---|
| 1 | 6,08 | n.a. | 745,591 | 52,672 | 100,00 | n.a. | n.a. |
| Total: | | | 745,591 | 52,672 | 100,00 | 0,000 | |

### Compound 58 - TUG-2757

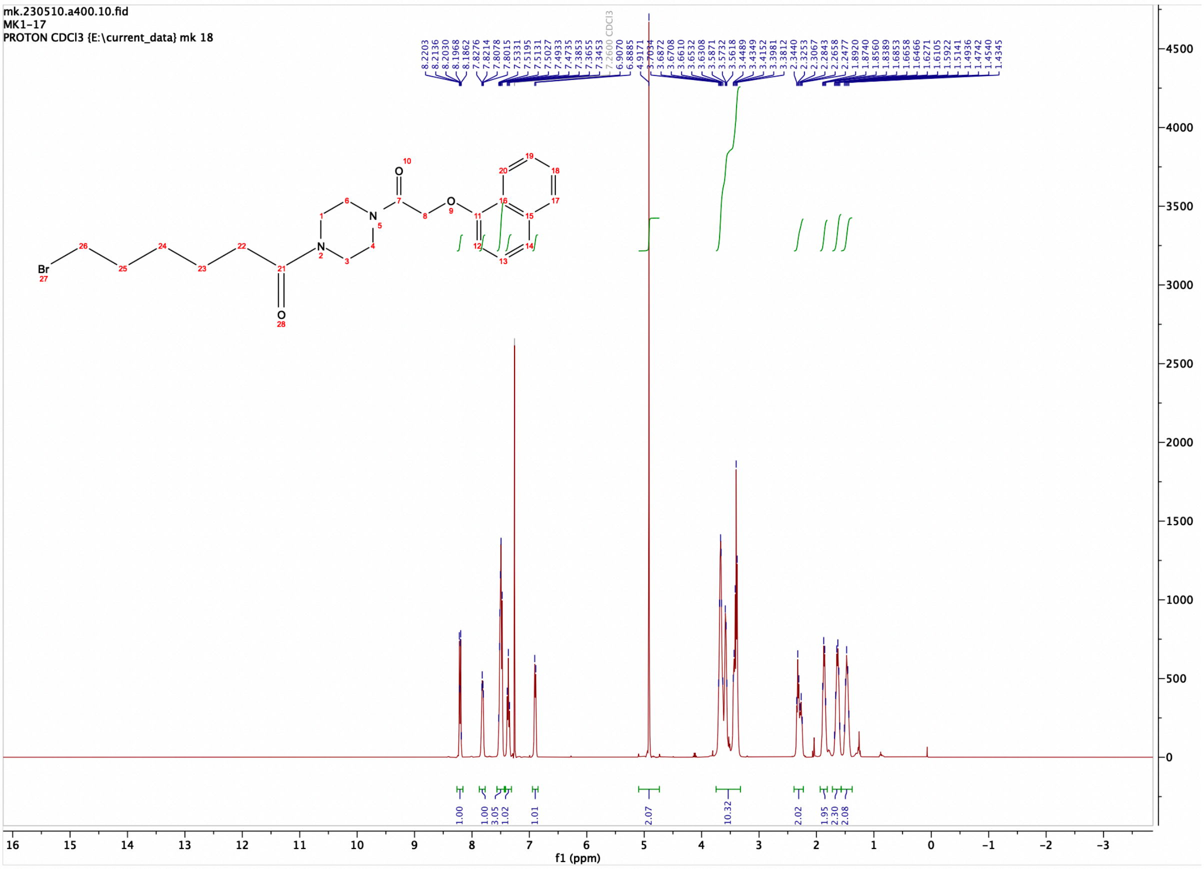

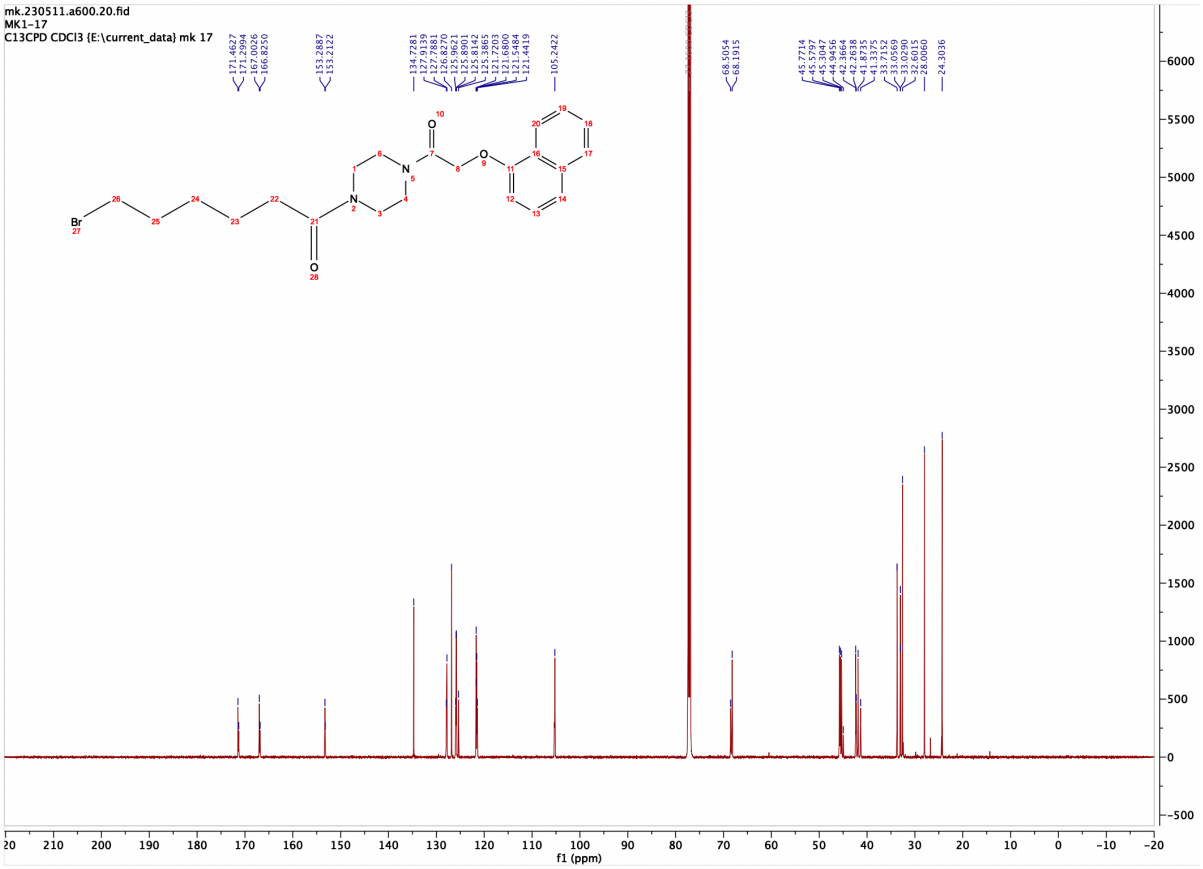

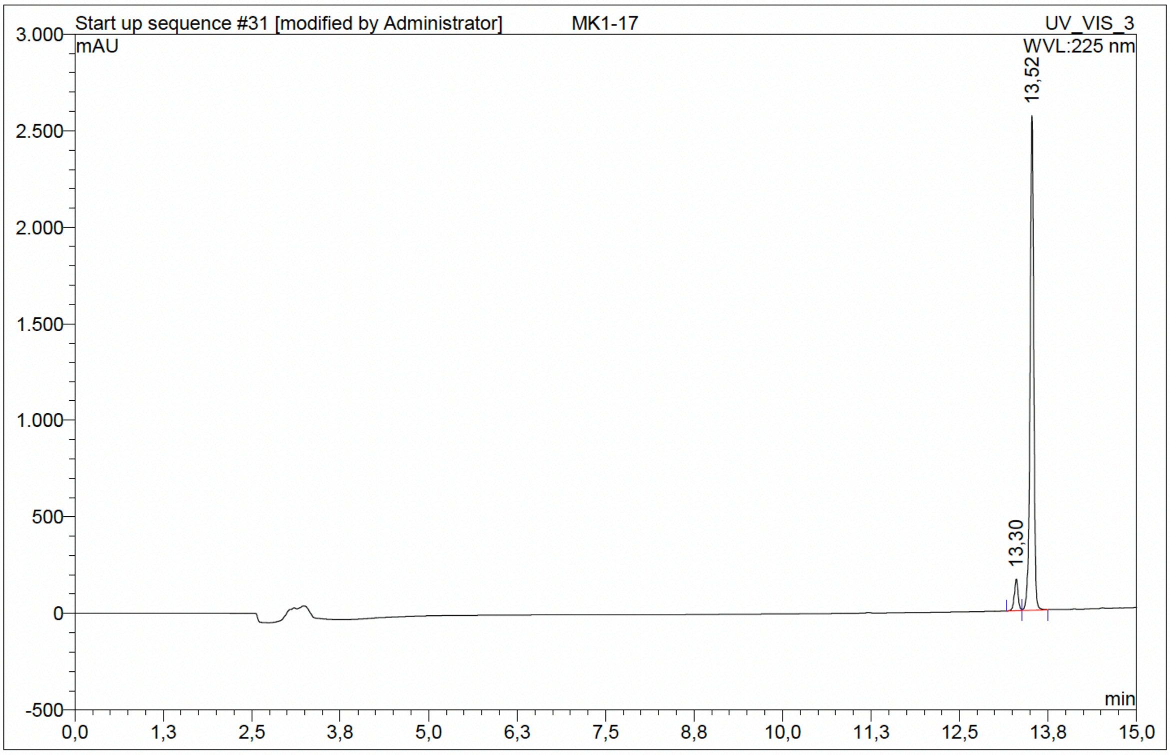

| No. | Ret.Time min | Peak Name | Height mAU | Area mAU*min | Rel.Area % | Amount | Resolution(EP) |
|---|---|---|---|---|---|---|---|
| 1 | 13,30 | n.a. | 163,974 | 9,996 | 6,02 | n.a. | 2,43 |
| 2 | 13,52 | n.a. | 2561,713 | 156,127 | 93,98 | n.a. | n.a. |
| Total: | | | 2725,687 | 166,124 | 100,00 | 0,000 | |

### Compound 59 - TUG-2758

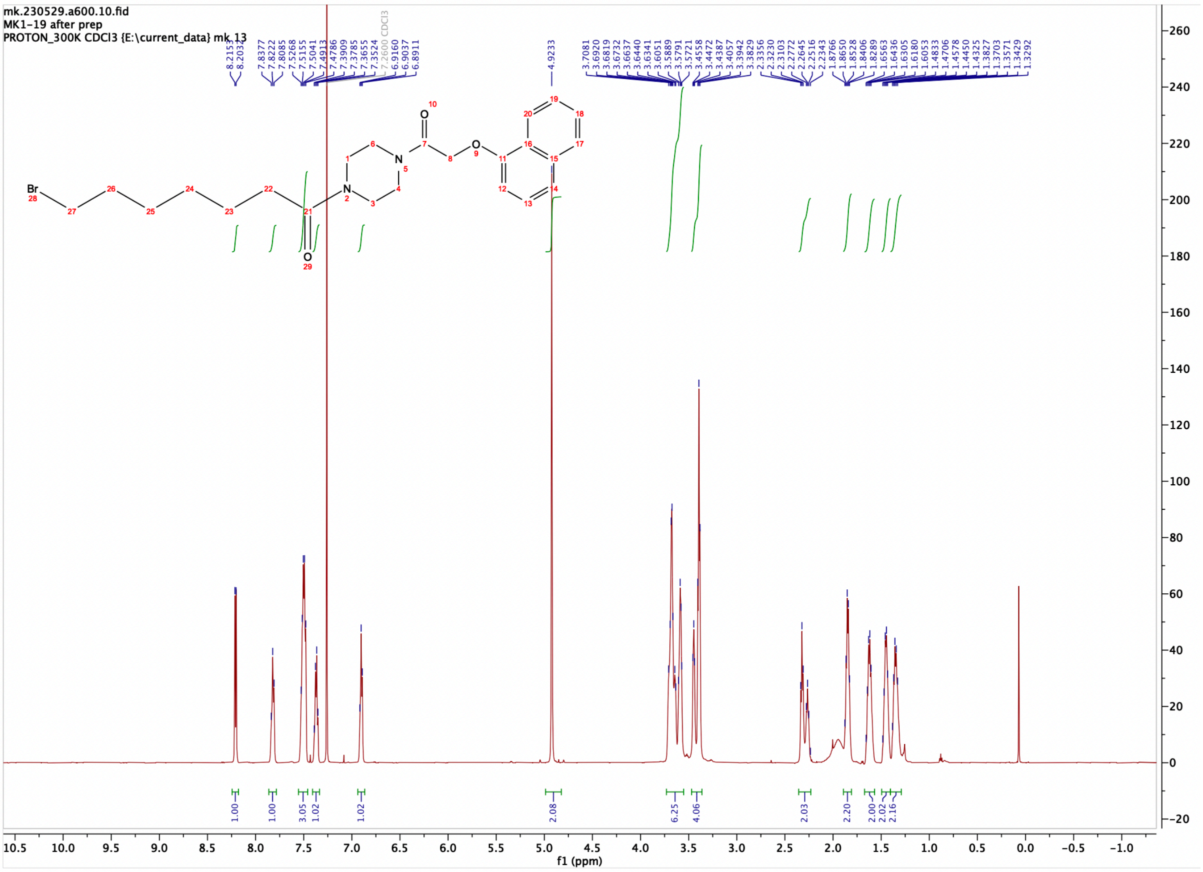

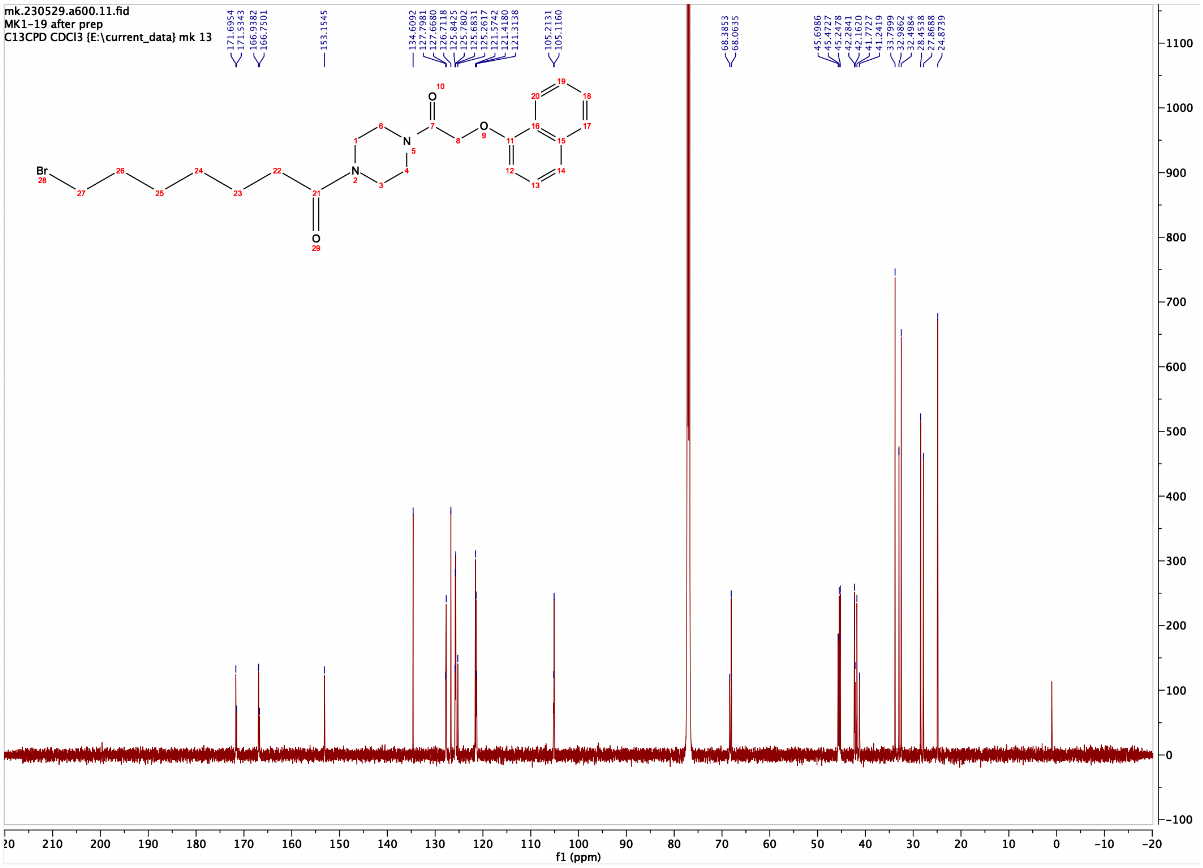

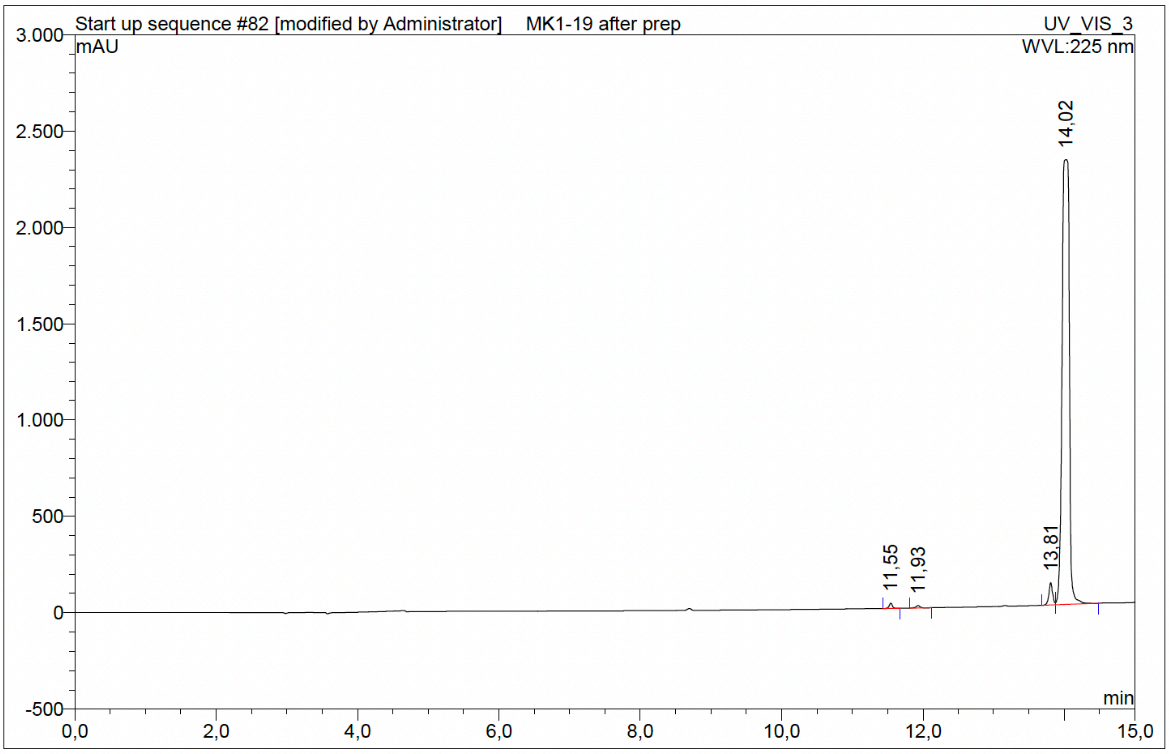

| No. | Ret.Time min | Peak Name | Height mAU | Area mAU*min | Rel.Area % | Amount | Resolution(EP) |
|---|---|---|---|---|---|---|---|
| 1 | 11,55 | n.a. | 27,059 | 1,440 | 0,52 | n.a. | 4,43 |
| 2 | 11,93 | n.a. | 13,082 | 0,836 | 0,30 | n.a. | 20,57 |
| 3 | 13,81 | n.a. | 115,373 | 6,912 | 2,49 | n.a. | 1,56 |
| 4 | 14,02 | n.a. | 2309,553 | 268,426 | 96,69 | n.a. | n.a. |
| Total: | | | 2465,068 | 277,613 | 100,00 | 0,000 | |

Compound 60 - TUG-2724

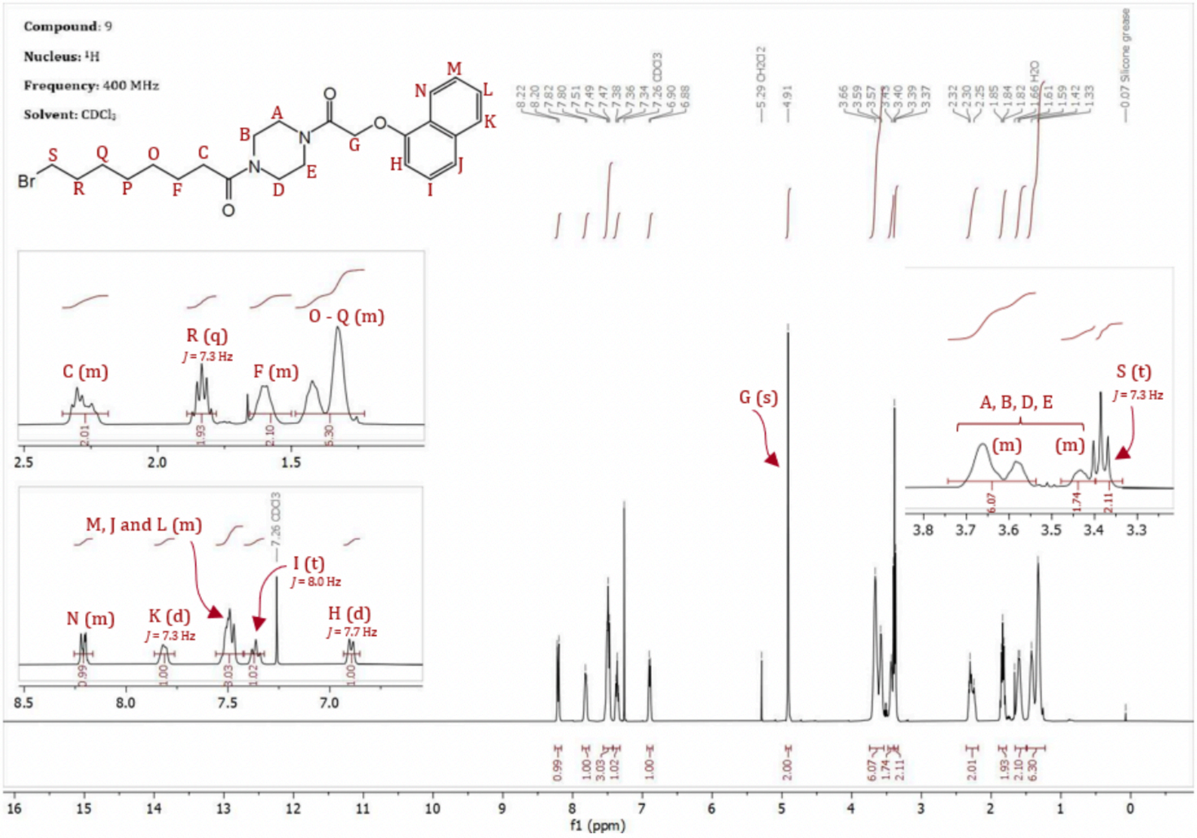

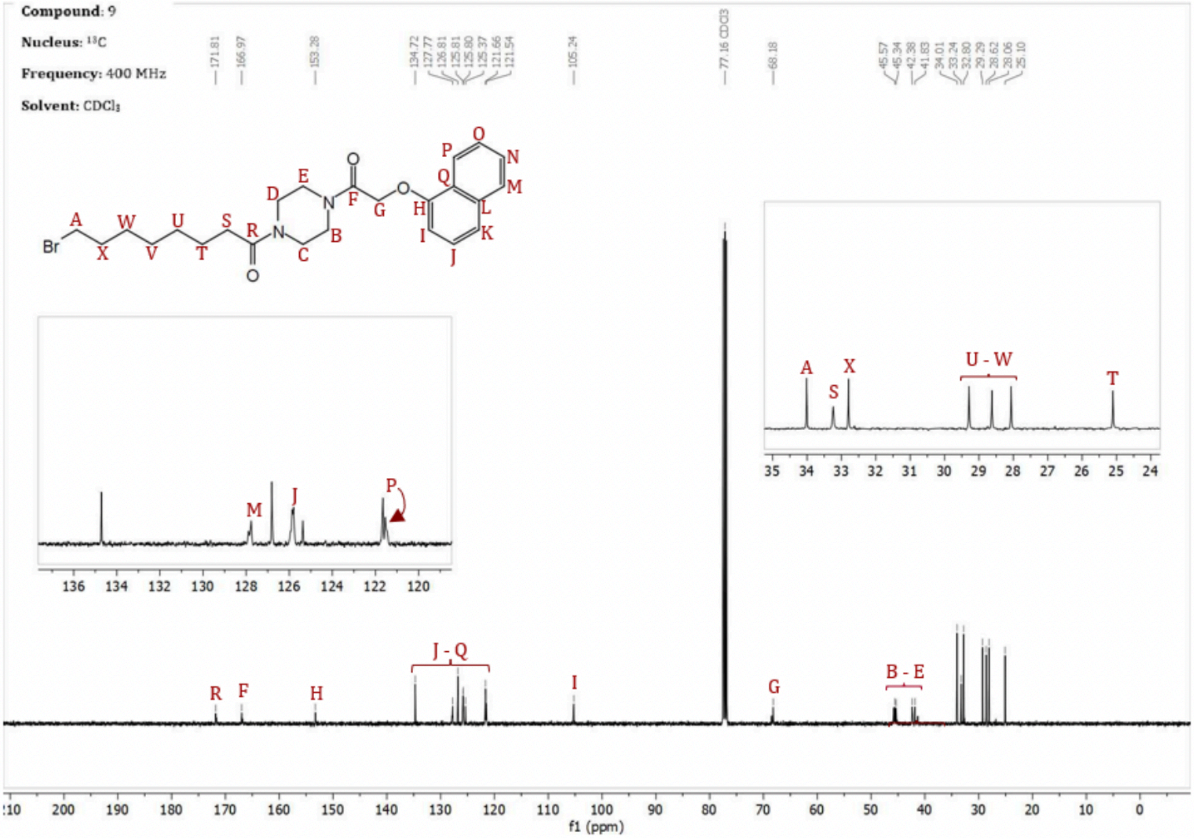

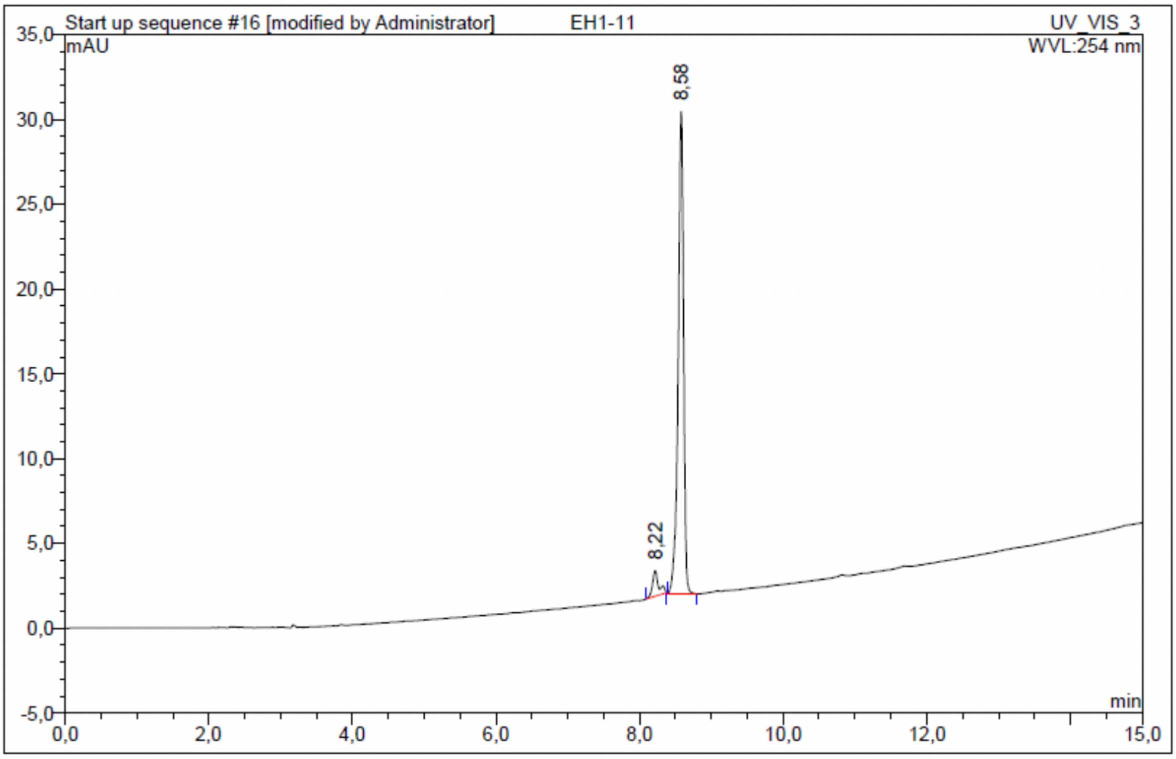

| No. | Ret.Time min | Peak Name | Height mAU | Area mAU*min | Rel.Area % | Amount | Resolution(EP) |
|---|---|---|---|---|---|---|---|
| 1 | 8,22 | n.a. | 1,501 | 0,151 | 5,64 | n.a. | 2,82 |
| 2 | 8,58 | n.a. | 28,424 | 2,521 | 94,36 | n.a. | n.a. |
| Total: | | | 29,925 | 2,671 | 100,00 | 0,000 | |

Compound 61 - TUG-2643

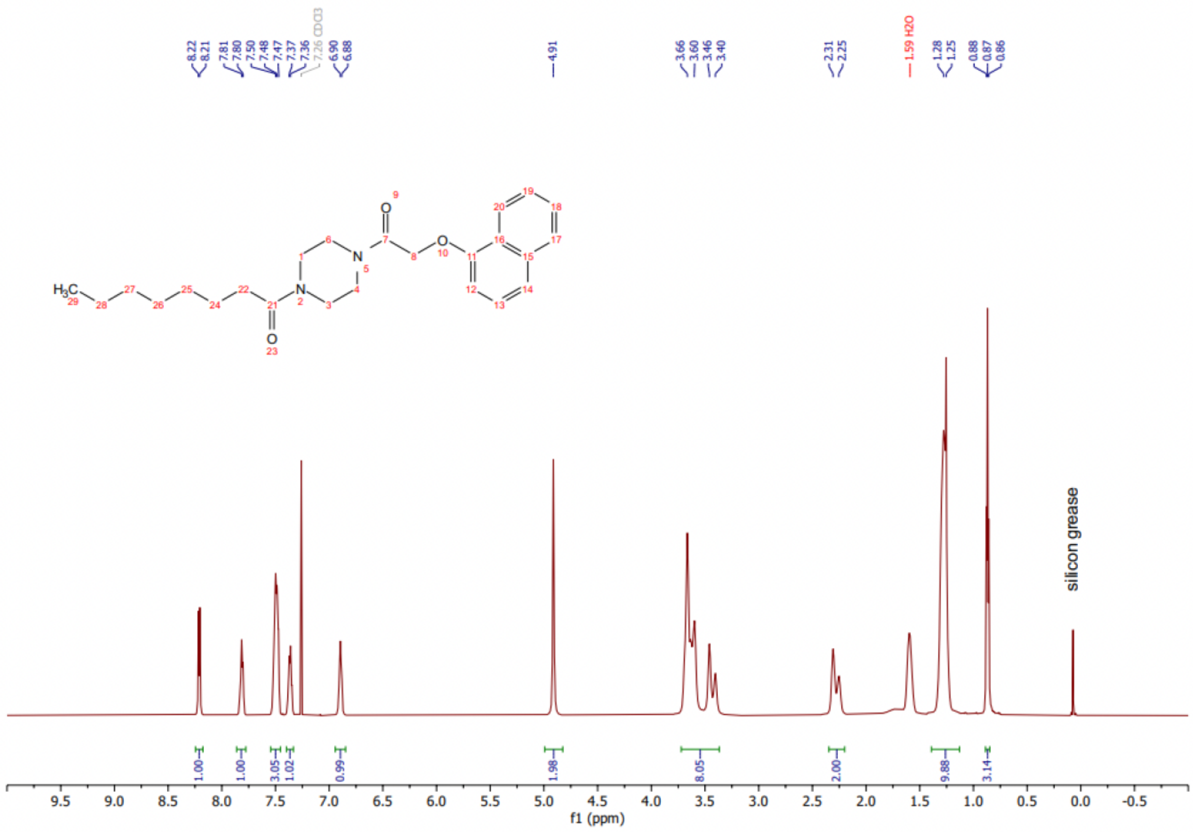

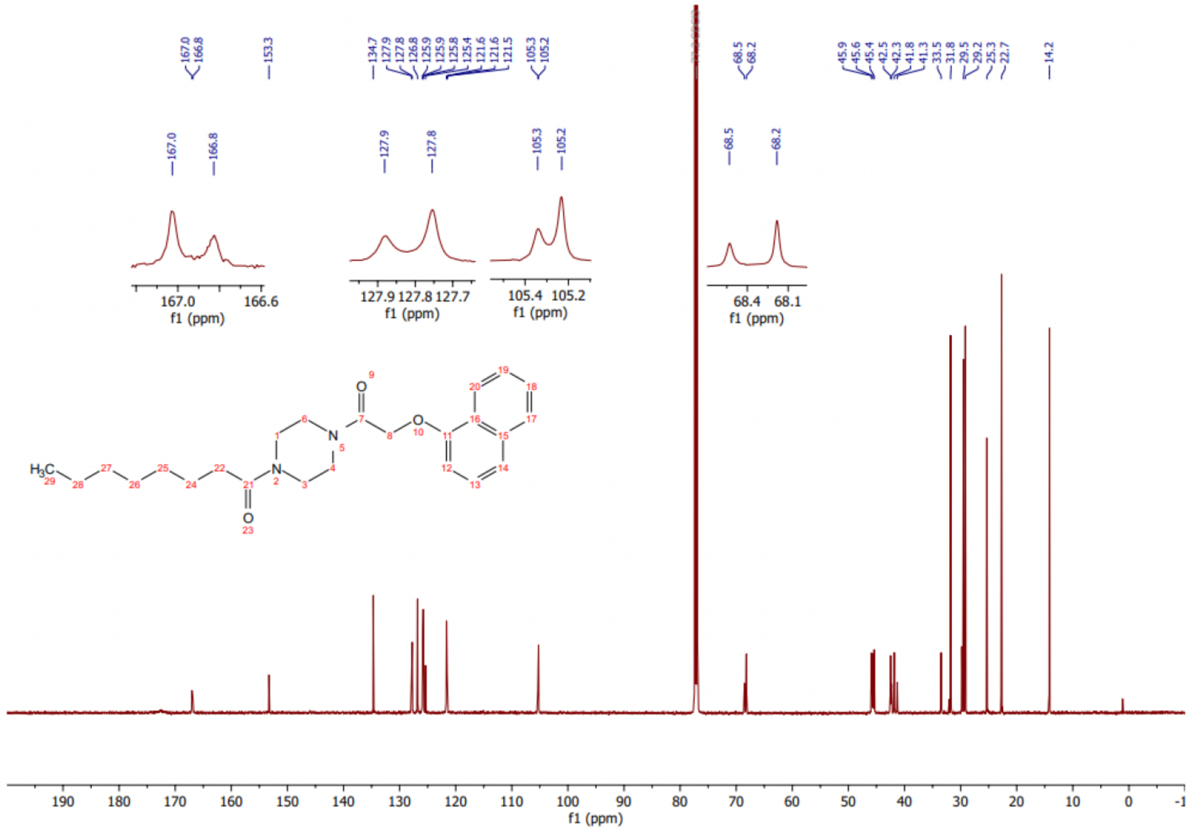

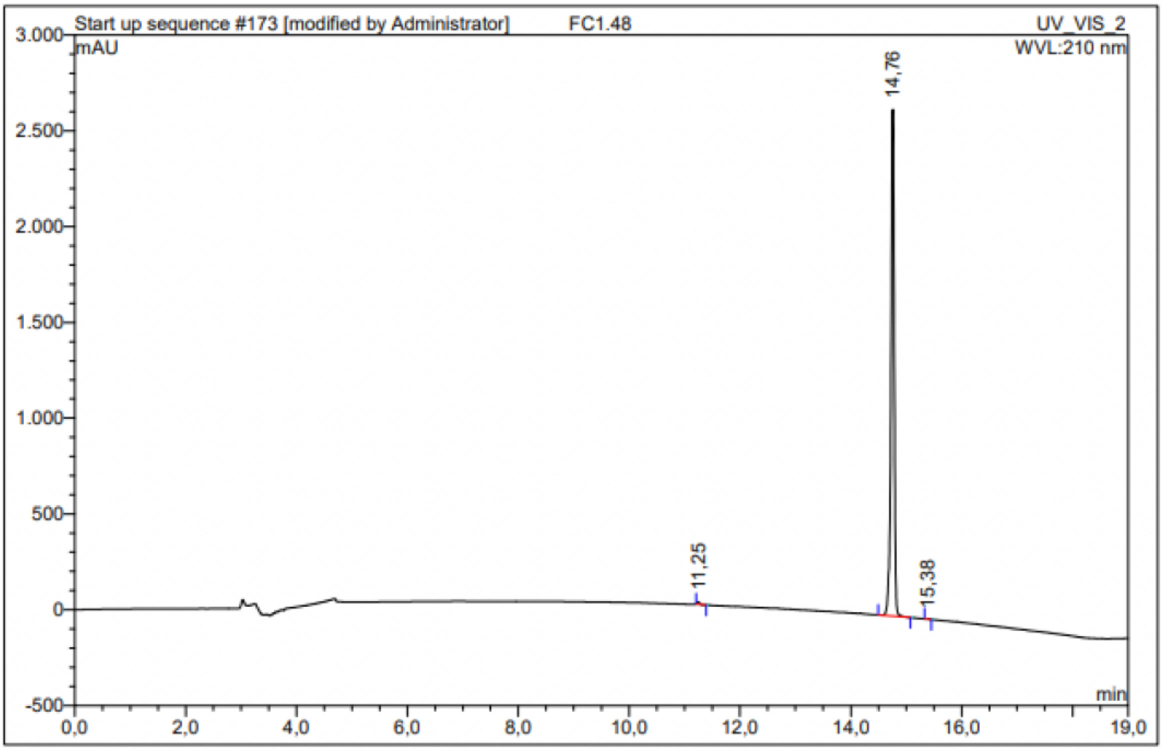

| No. | Ret.Time min | Peak Name | Height mAU | Area mAU*min | Rel.Area % | Amount | Resolution(EP) |
|---|---|---|---|---|---|---|---|
| 1 | 11,25 | n.a. | 12,731 | 0,569 | 0,31 | n.a. | 37,85 |
| 2 | 14,76 | n.a. | 2646,044 | 181,063 | 99,62 | n.a. | 6,38 |
| 3 | 15,38 | n.a. | 2,170 | 0,127 | 0,07 | n.a. | n.a. |
| Total: | | | 2660,944 | 181,759 | 100,00 | 0,000 | |

Compound 62 - TUG-2644

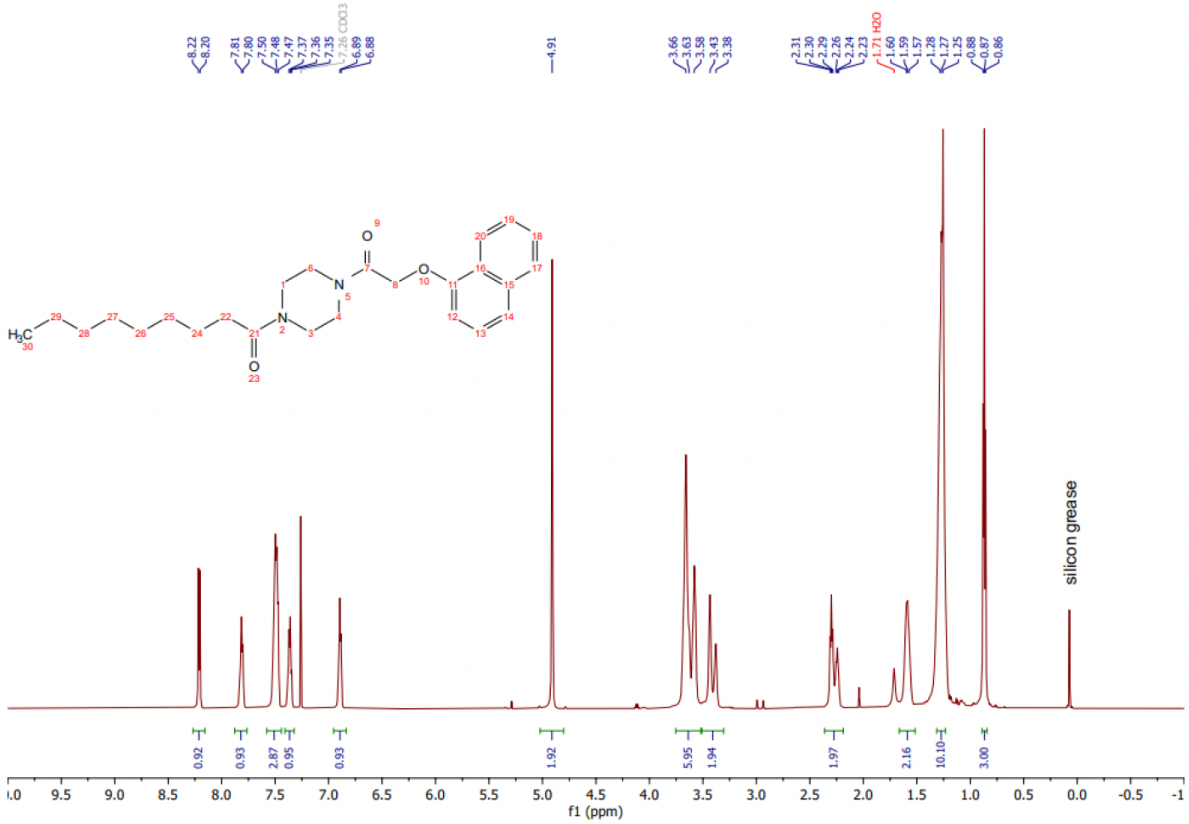

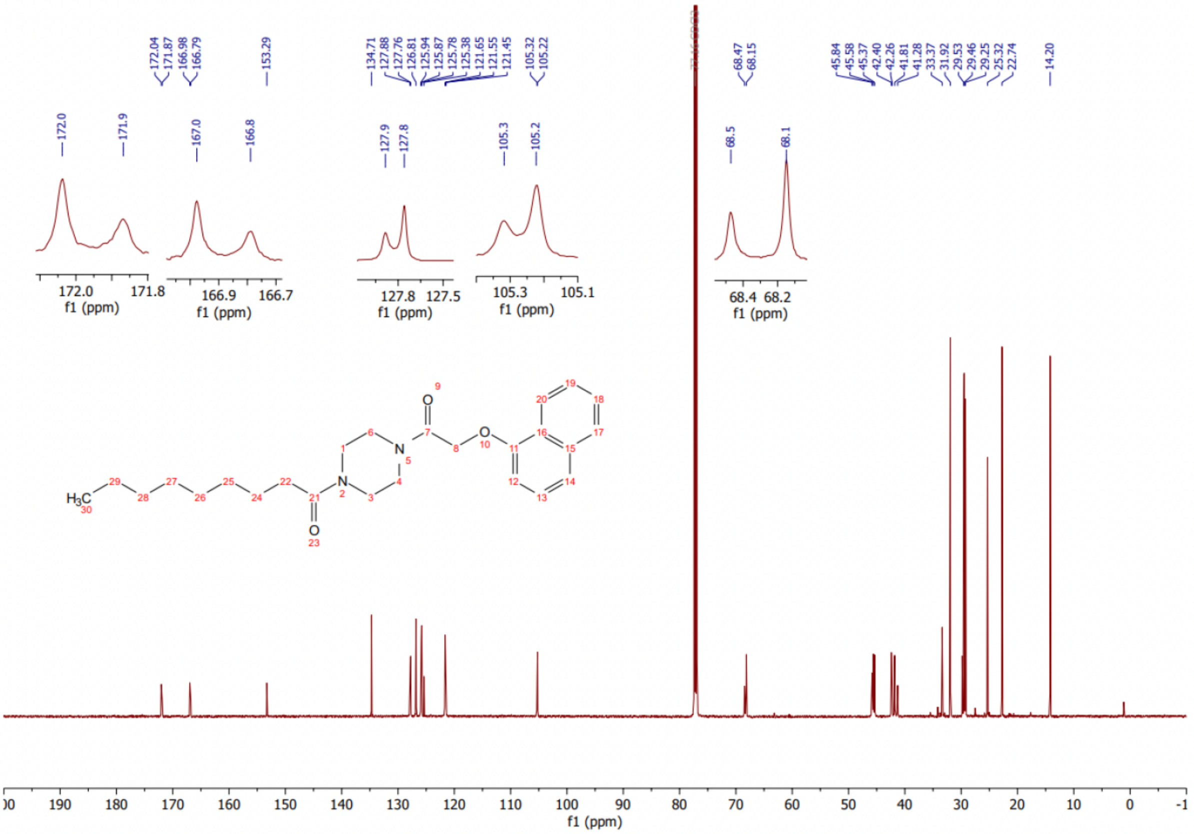

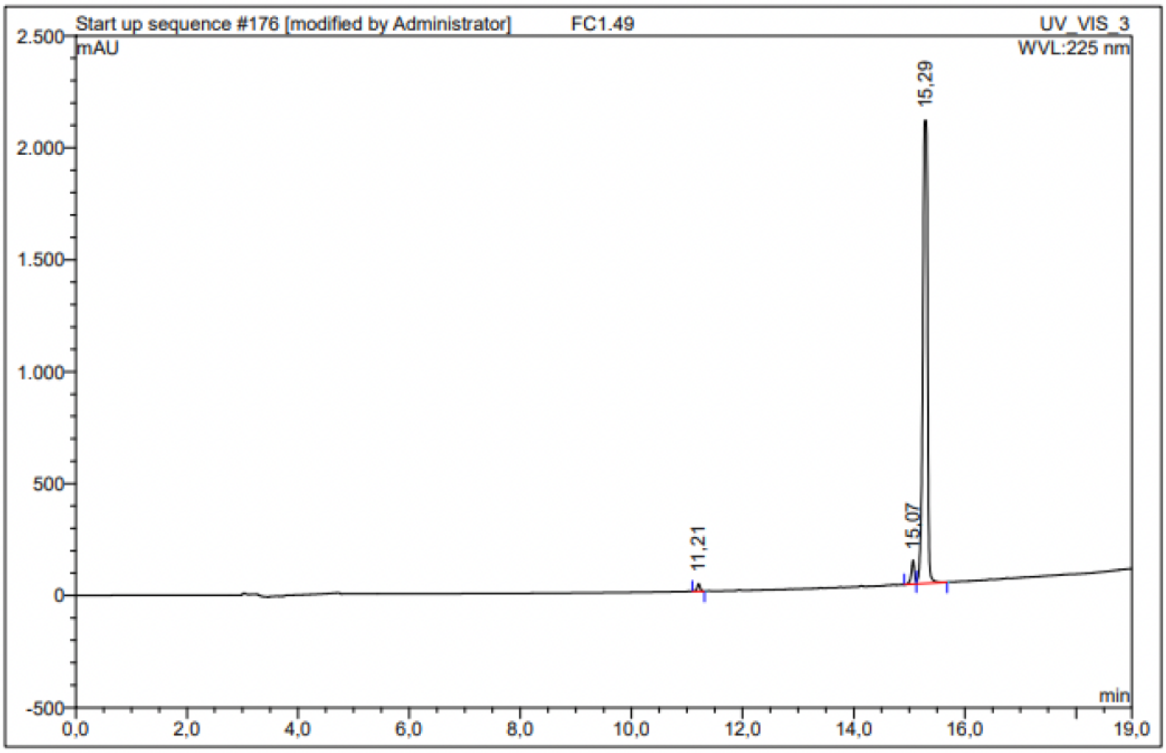

| No. | Ret.Time min | Peak Name | Height mAU | Area mAU*min | Rel.Area % | Amount | Resolution(EP) |
|---|---|---|---|---|---|---|---|
| 1 | 11,21 | n.a. | 33,784 | 2,037 | 0,93 | n.a. | 39,90 |
| 2 | 15,07 | n.a. | 106,091 | 7,162 | 3,28 | n.a. | 1,69 |
| 3 | 15,29 | n.a. | 2069,064 | 209,373 | 95,79 | n.a. | n.a. |
| Total: | | | 2208,940 | 218,572 | 100,00 | 0,000 | |

Compound 63 - TUG-2604

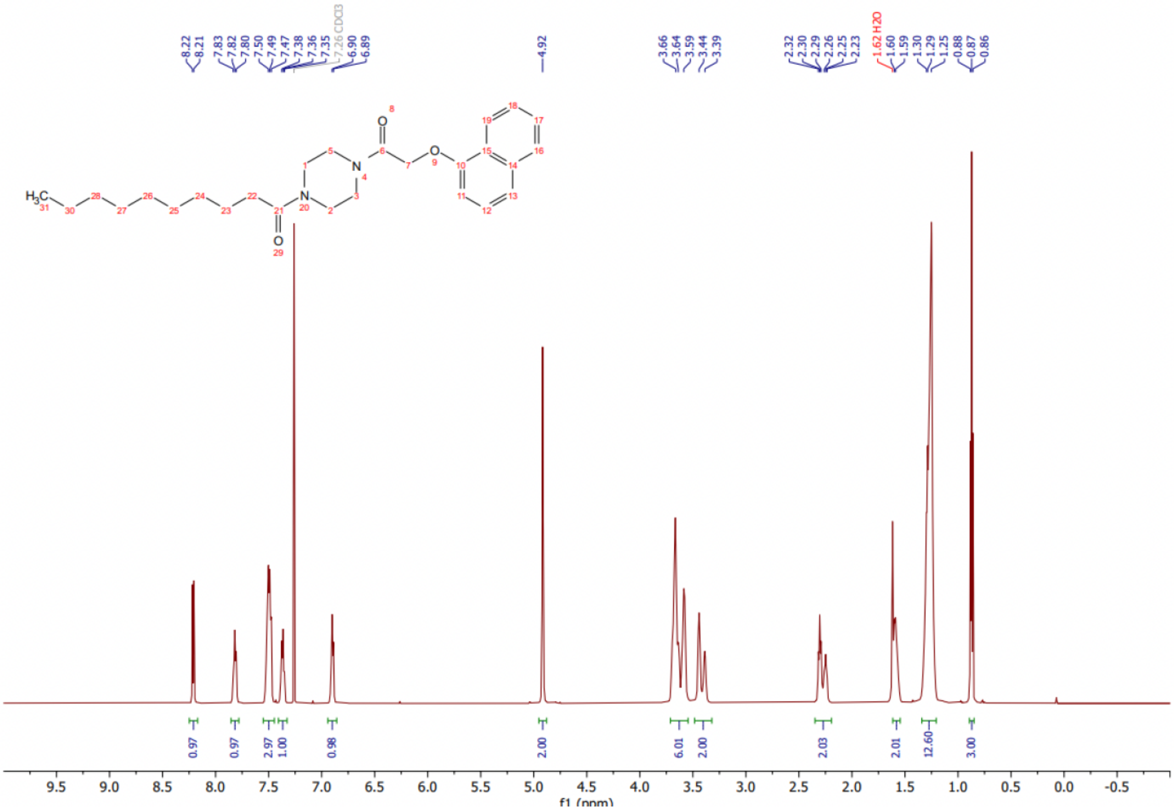

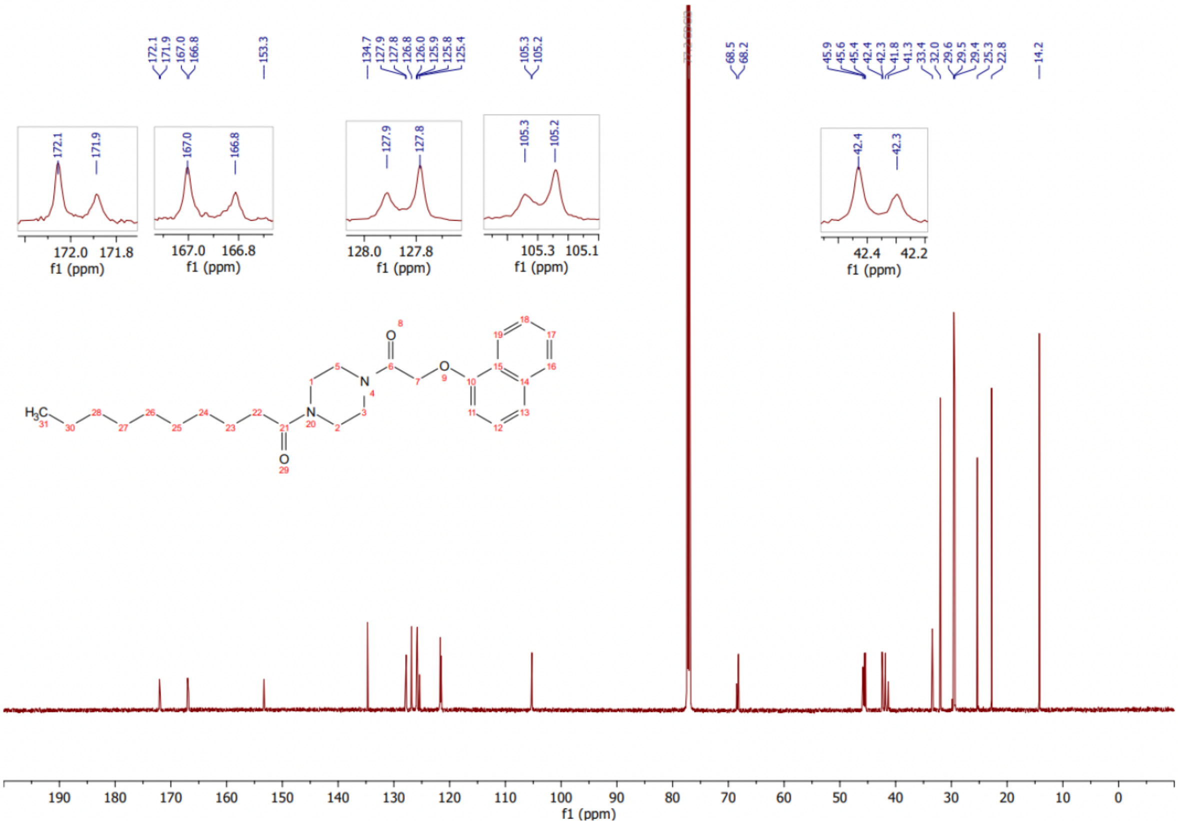

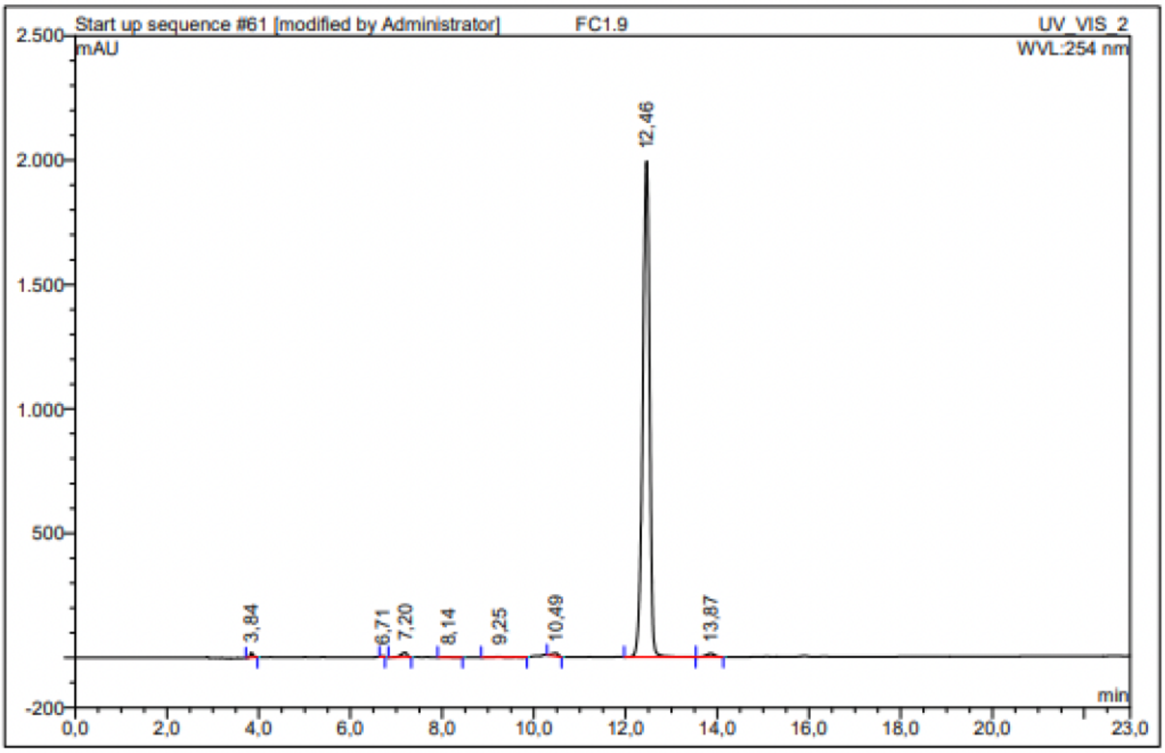

| No. | Ret.Time min | Peak Name | Height mAU | Area mAU*min | Rel.Area % | Amount | Resolution(EP) |
|---|---|---|---|---|---|---|---|
| 1 | 3,84 | n.a. | 19,631 | 1,438 | 0,40 | n.a. | 23,93 |
| 2 | 6,71 | n.a. | 2,115 | 0,161 | 0,05 | n.a. | 2,65 |
| 3 | 7,20 | n.a. | 18,464 | 2,739 | 0,77 | n.a. | 4,04 |
| 4 | 8,14 | n.a. | 1,244 | 0,178 | 0,05 | n.a. | 3,09 |
| 5 | 9,25 | n.a. | 2,254 | 0,408 | 0,11 | n.a. | 3,30 |
| 6 | 10,49 | n.a. | 12,555 | 1,919 | 0,54 | n.a. | 7,26 |
| 7 | 12,46 | n.a. | 1991,939 | 346,221 | 97,29 | n.a. | 4,84 |
| 8 | 13,87 | n.a. | 14,021 | 2,787 | 0,78 | n.a. | n.a. |
| Total: | | | 2062,223 | 355,851 | 100,00 | 0,000 | |

Compound 64 - TUG-2331

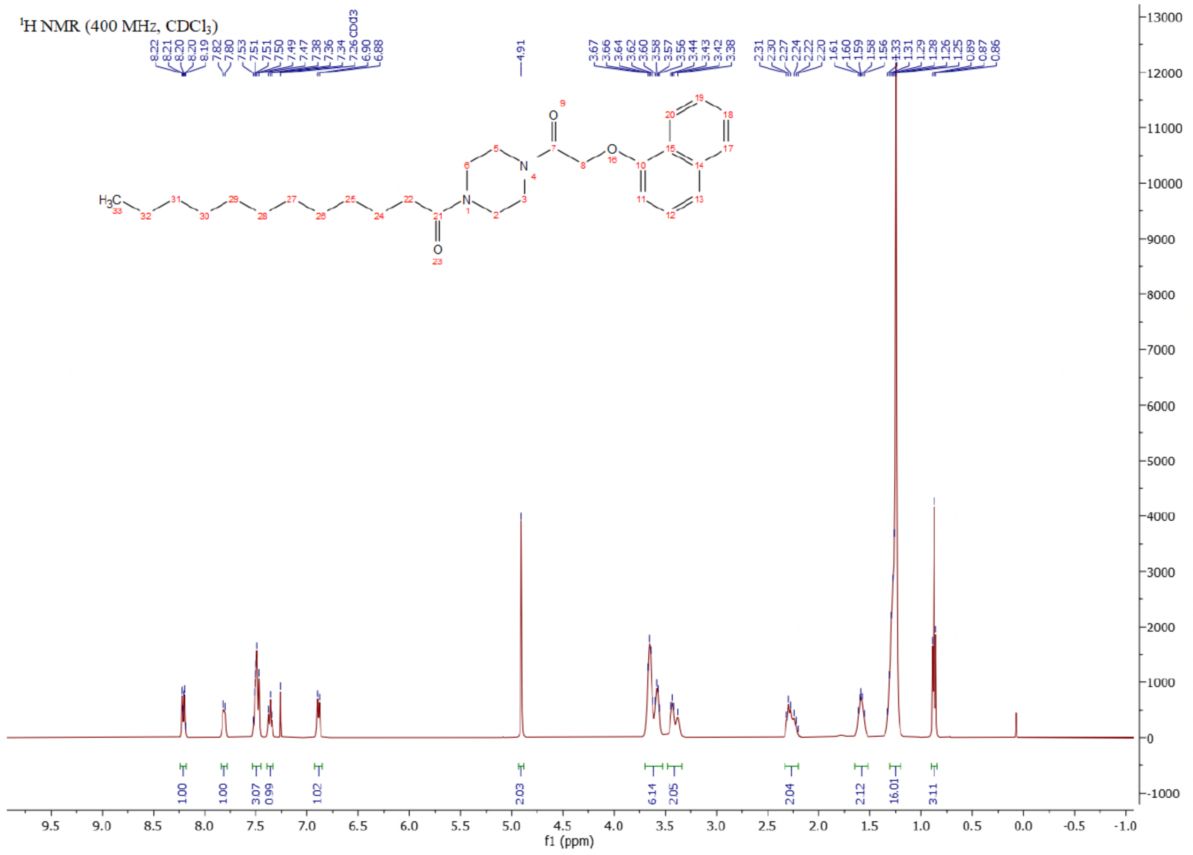

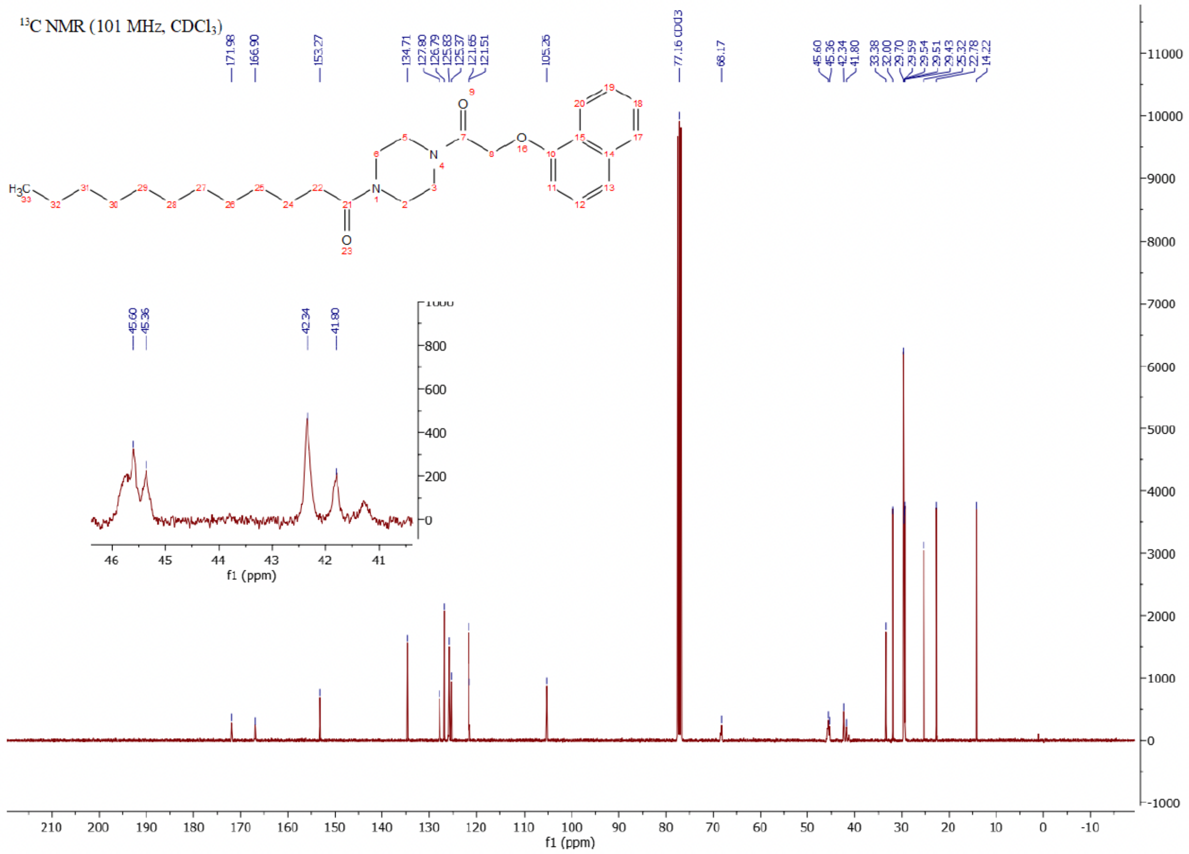

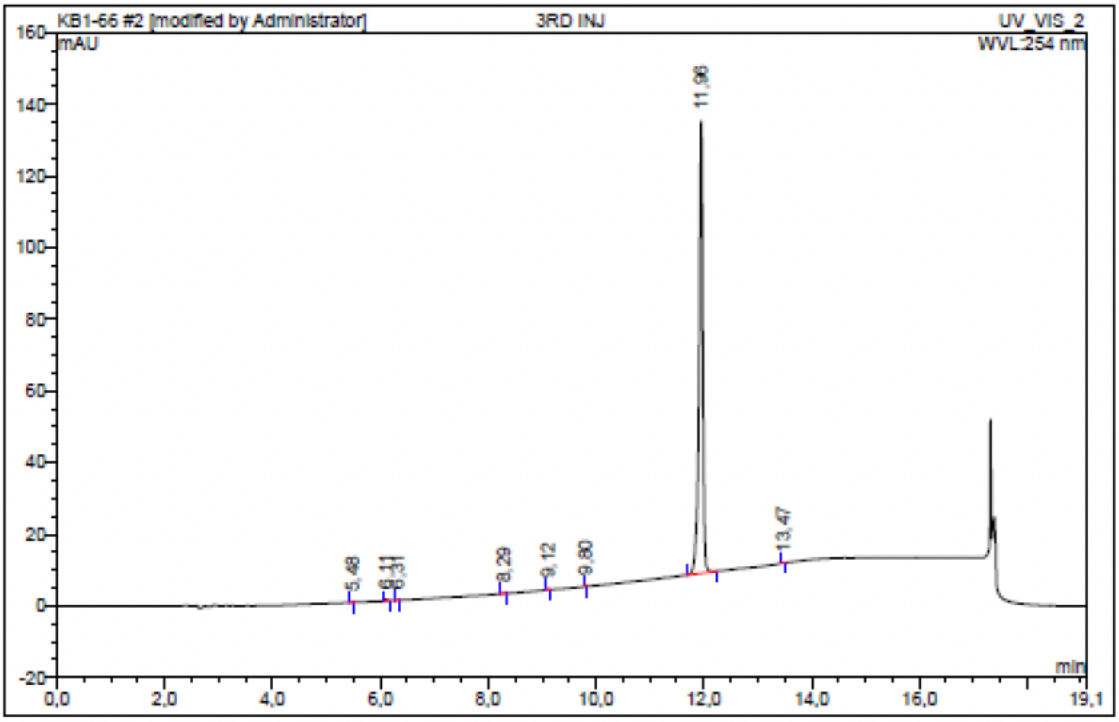

| No. | Ret.Time min | Peak Name | Height mAU | Area mAU*min | Rel.Area % | Amount | Resolution(EP) |
|---|---|---|---|---|---|---|---|
| 1 | 5,48 | n.a. | 0,108 | 0,005 | 0,05 | n.a. | 7,31 |
| 2 | 6,11 | n.a. | 0,496 | 0,027 | 0,26 | n.a. | 2,33 |
| 3 | 6,31 | n.a. | 0,198 | 0,011 | 0,10 | n.a. | 22,32 |
| 4 | 8,29 | n.a. | 0,175 | 0,010 | 0,09 | n.a. | 10,02 |
| 5 | 9,12 | n.a. | 0,218 | 0,010 | 0,09 | n.a. | 10,17 |
| 6 | 9,80 | n.a. | 0,088 | 0,003 | 0,03 | n.a. | 24,47 |
| 7 | 11,96 | n.a. | 126,263 | 10,349 | 99,32 | n.a. | 15,56 |
| 8 | 13,47 | n.a. | 0,126 | 0,006 | 0,06 | n.a. | n.a. |
| Total: | | | 127,673 | 10,419 | 100,00 | 0,000 | |

Compound 65 - TUG-2391

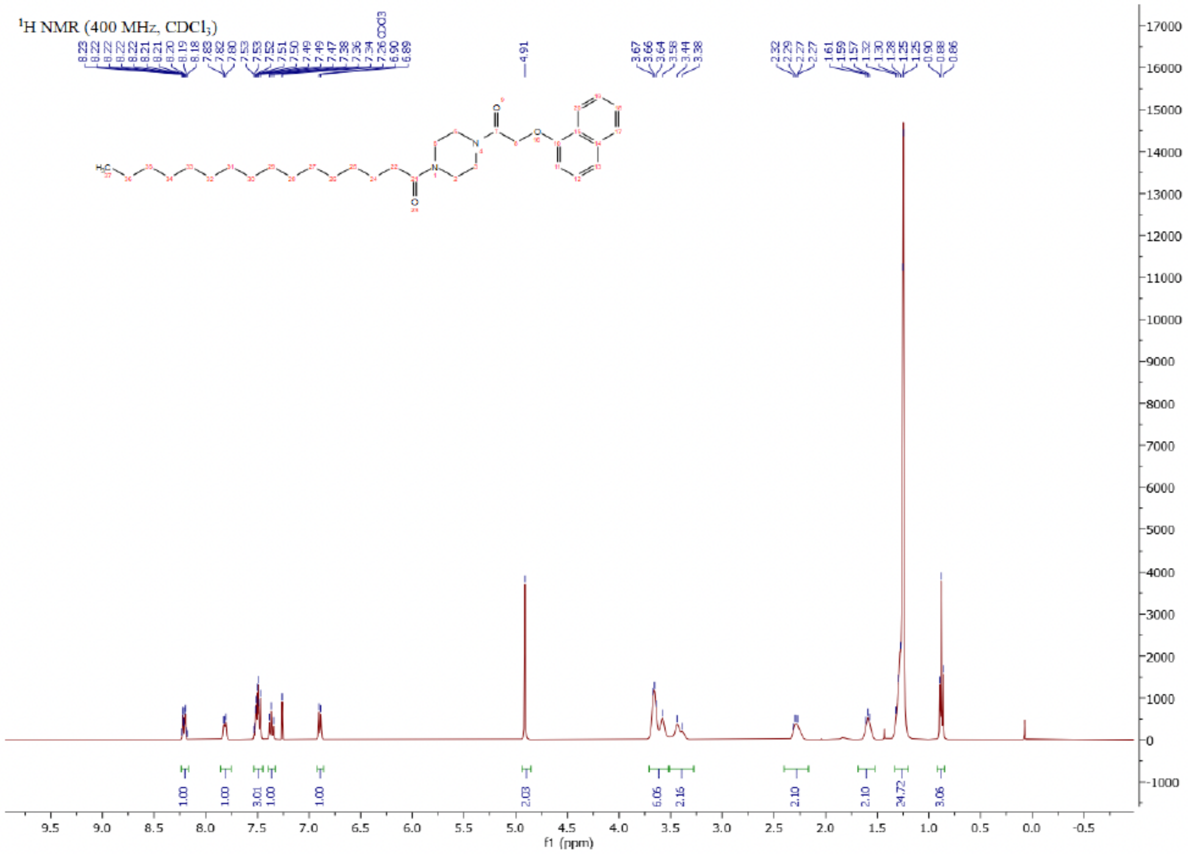

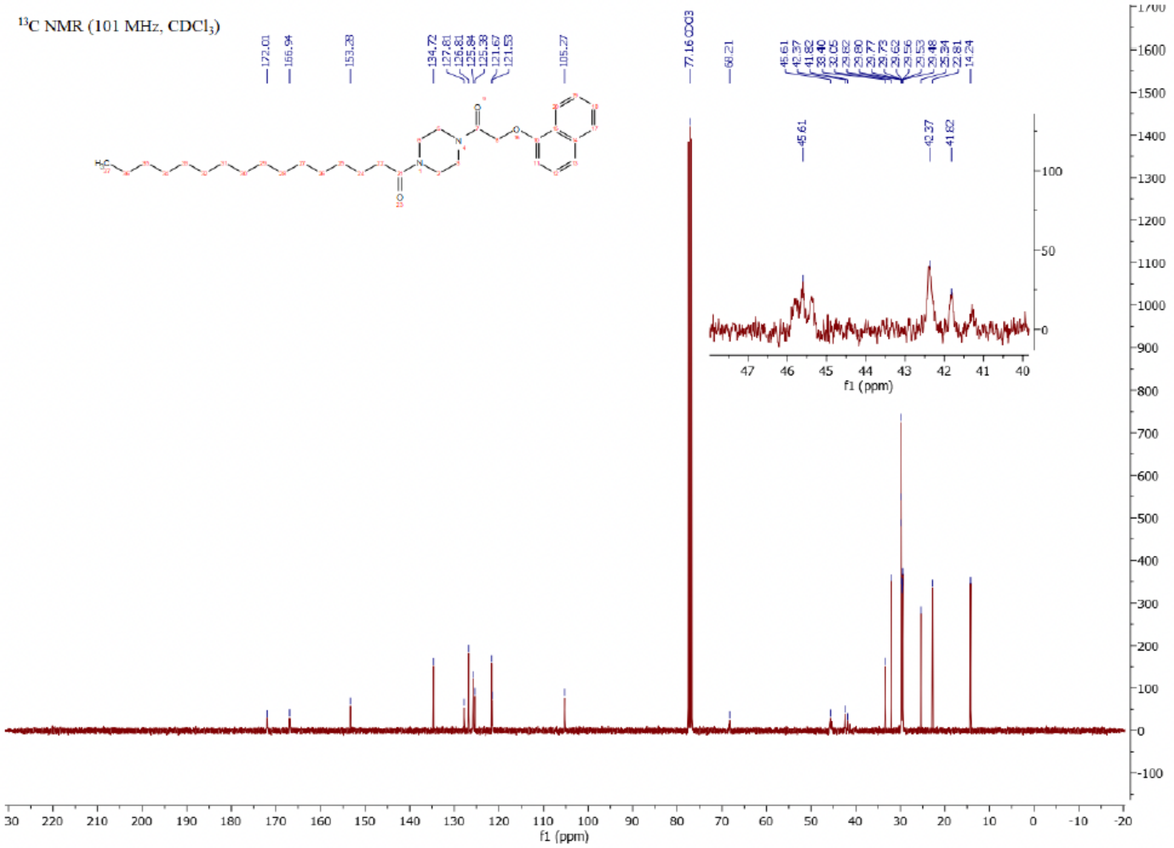

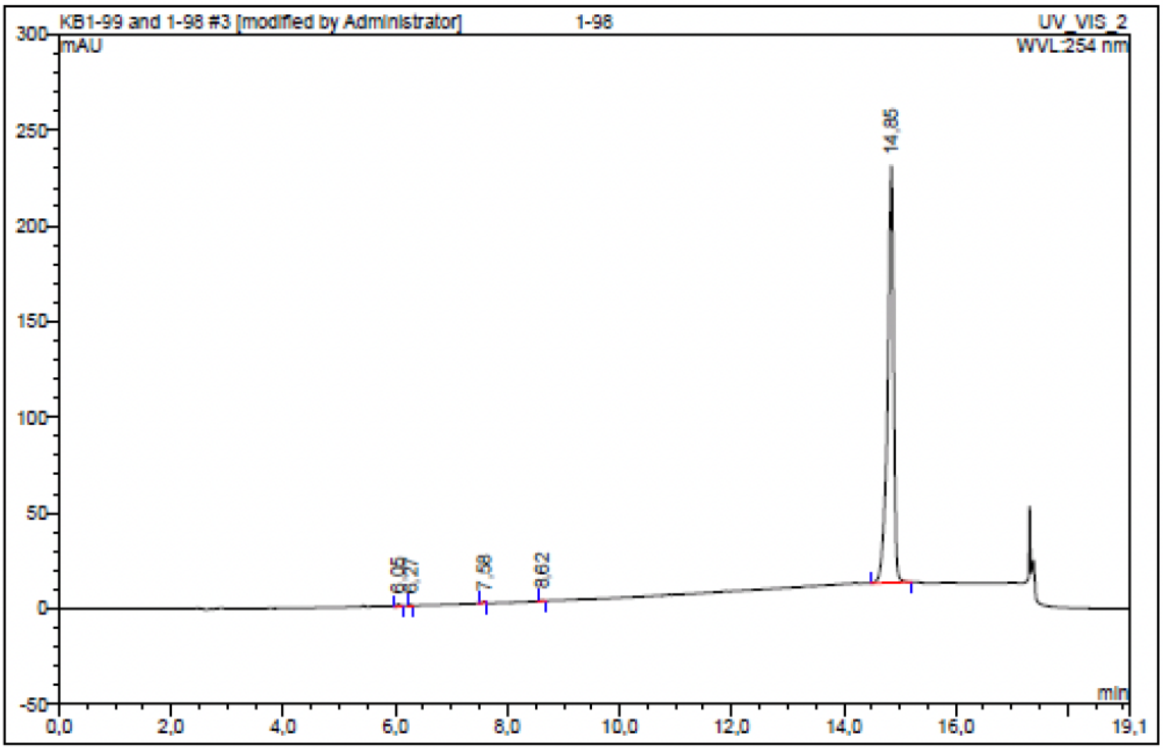

| No. | Ret.Time min | Peak Name | Height mAU | Area mAU*min | Rel.Area % | Amount | Resolution(EP) |
|---|---|---|---|---|---|---|---|
| 1 | 6,05 | n.a. | 1,392 | 0,077 | 0,28 | n.a. | 2,72 |
| 2 | 6,27 | n.a. | 0,479 | 0,020 | 0,08 | n.a. | 18,87 |
| 3 | 7,58 | n.a. | 0,721 | 0,025 | 0,09 | n.a. | 12,99 |
| 4 | 8,62 | n.a. | 0,497 | 0,029 | 0,11 | n.a. | 46,68 |
| 5 | 14,85 | n.a. | 217,685 | 26,962 | 99,44 | n.a. | n.a. |
| Total: | | | 220,774 | 27,114 | 100,00 | 0,000 | |

Compound 66 - TUG-2650

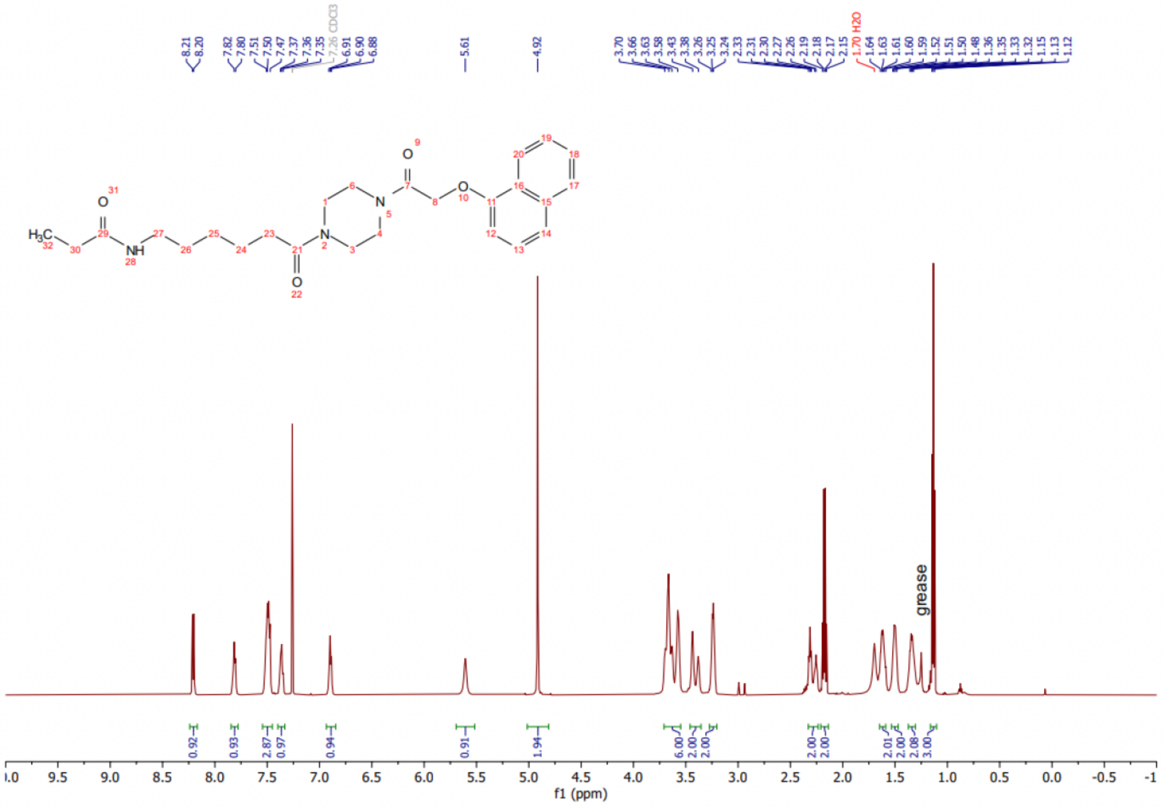

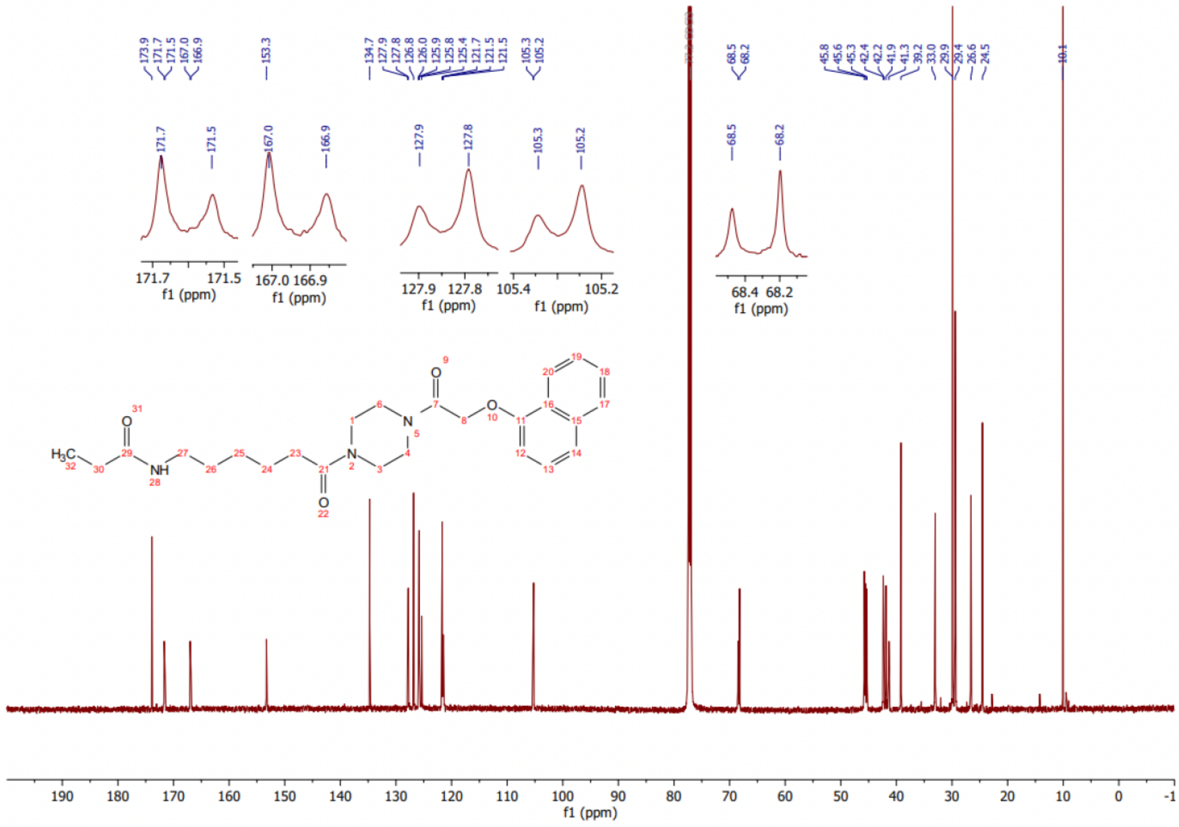

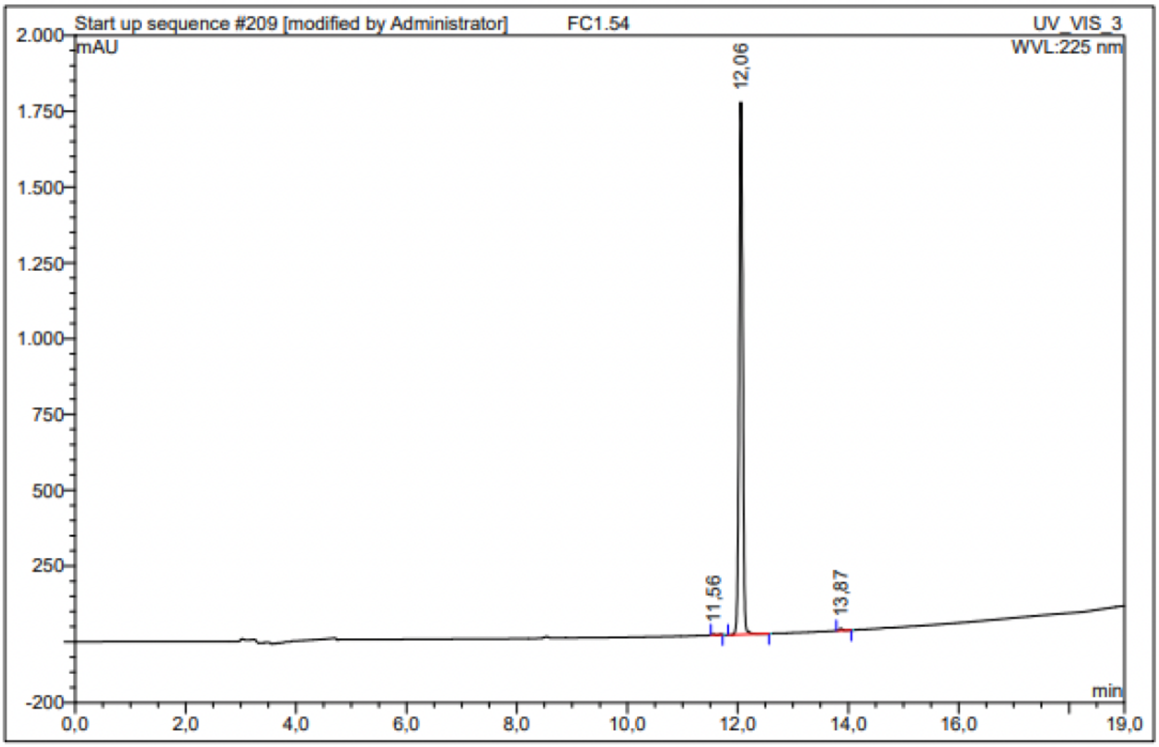

| No. | Ret.Time min | Peak Name | Height mAU | Area mAU*min | Rel.Area % | Amount | Resolution(EP) |
|---|---|---|---|---|---|---|---|
| 1 | 11,56 | n.a. | 2,494 | 0,120 | 0,09 | n.a. | 5,06 |
| 2 | 12,06 | n.a. | 1755,719 | 128,304 | 99,53 | n.a. | 15,85 |
| 3 | 13,87 | n.a. | 6,704 | 0,489 | 0,38 | n.a. | n.a. |
| Total: | | | 1764,917 | 128,913 | 100,00 | 0,000 | |

### Compound 67 - TUG-2756

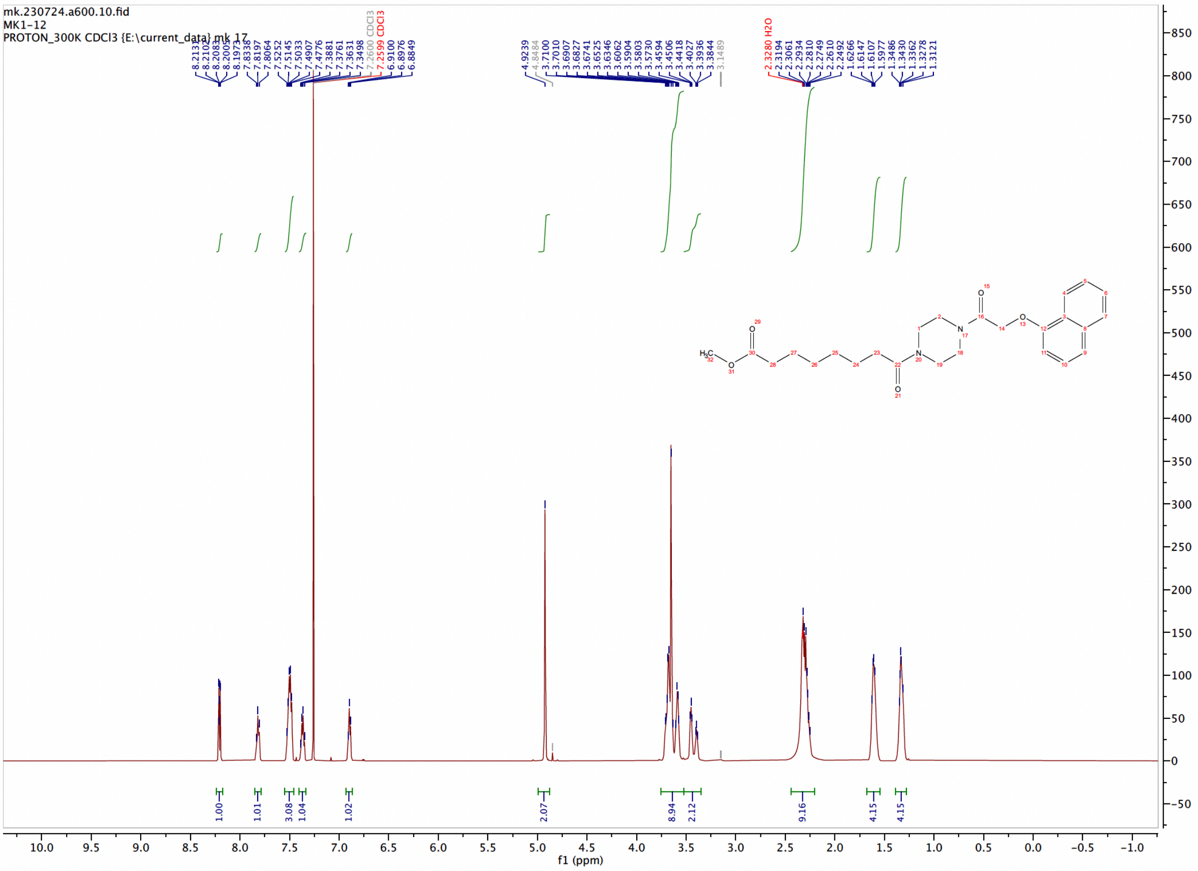

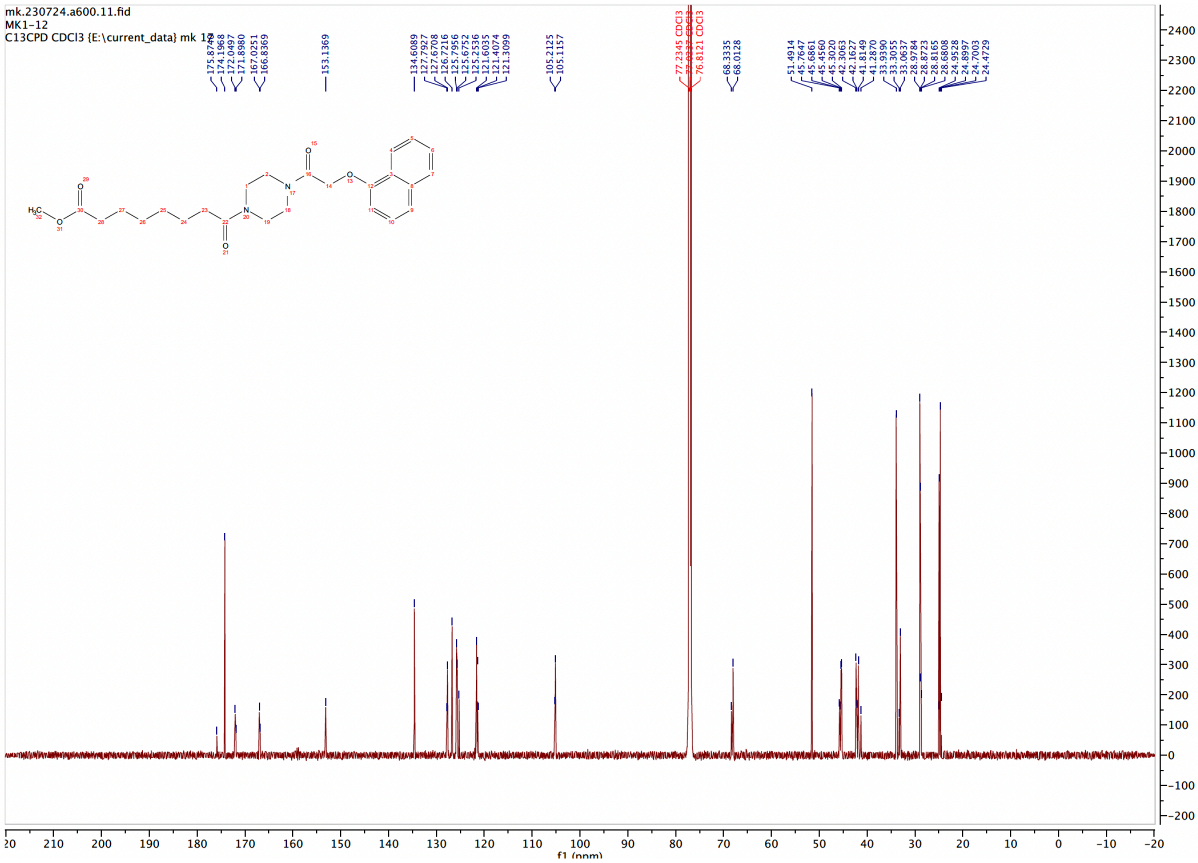

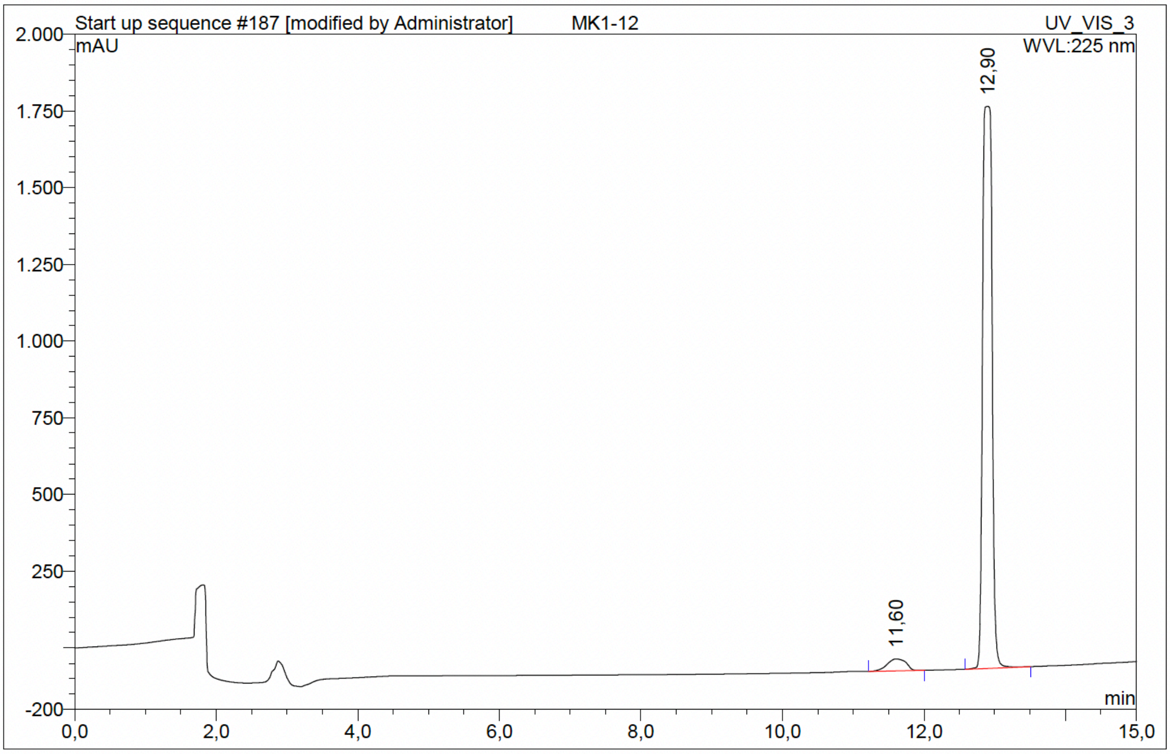

| No. | Ret.Time min | Peak Name | Height mAU | Area mAU*min | Rel.Area % | Amount | Resolution(EP) |
|---|---|---|---|---|---|---|---|
| 1 | 11,60 | n.a. | 39,182 | 12,084 | 4,12 | n.a. | 3,40 |
| 2 | 12,90 | n.a. | 1830,676 | 281,242 | 95,88 | n.a. | n.a. |
| Total: | | | 1869,859 | 293,326 | 100,00 | 0,000 | |

Compound 68 - TUG-2841

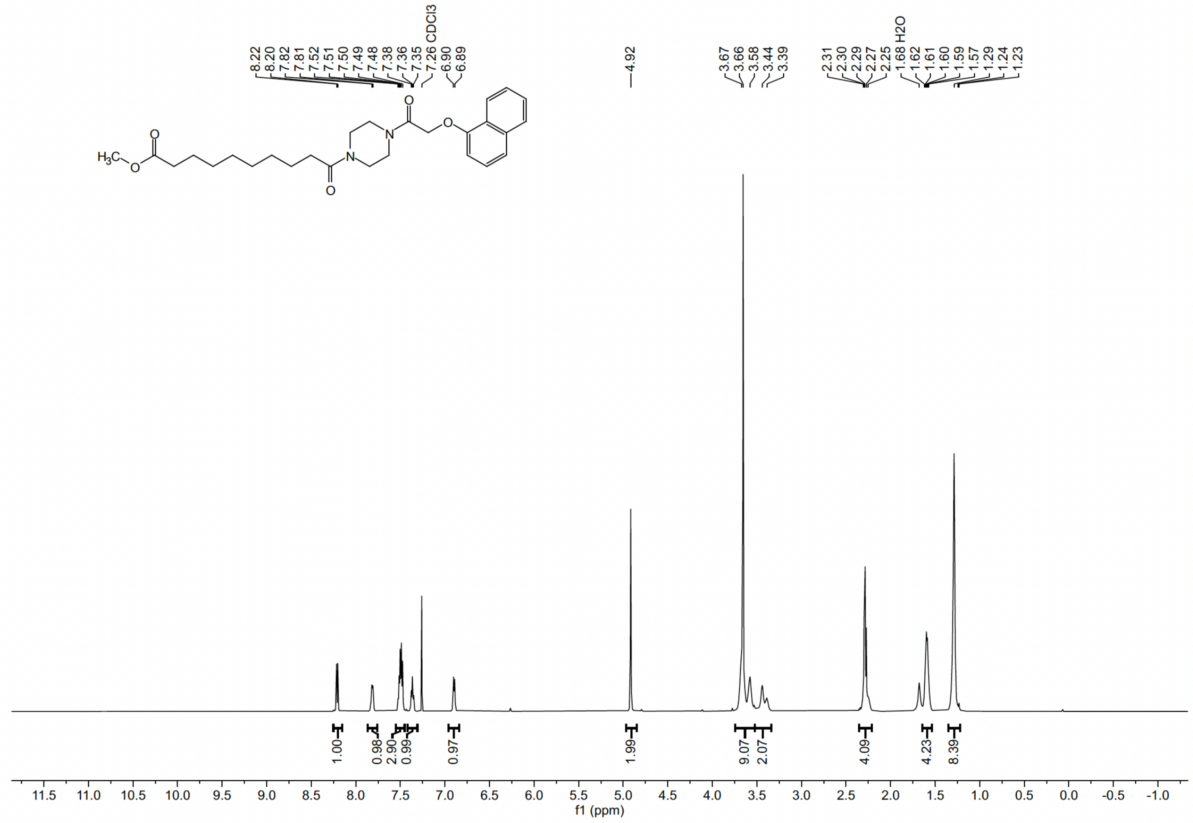

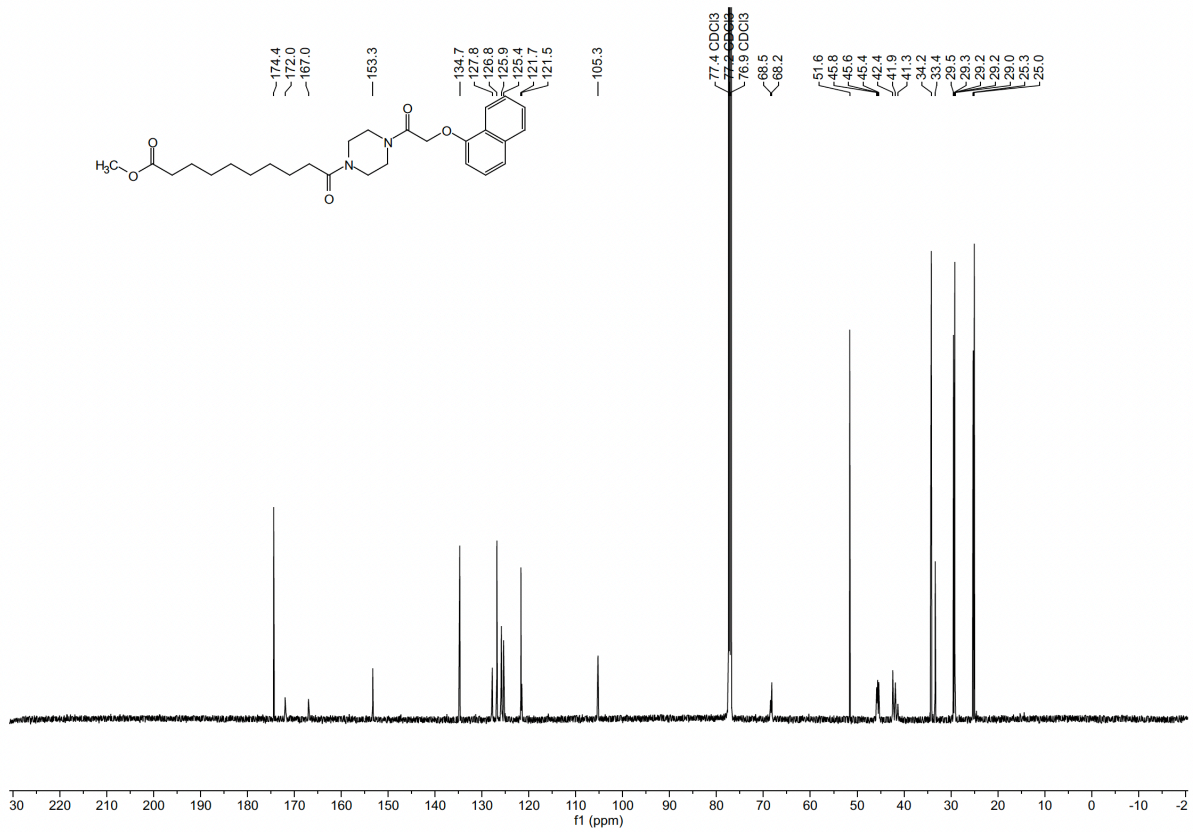

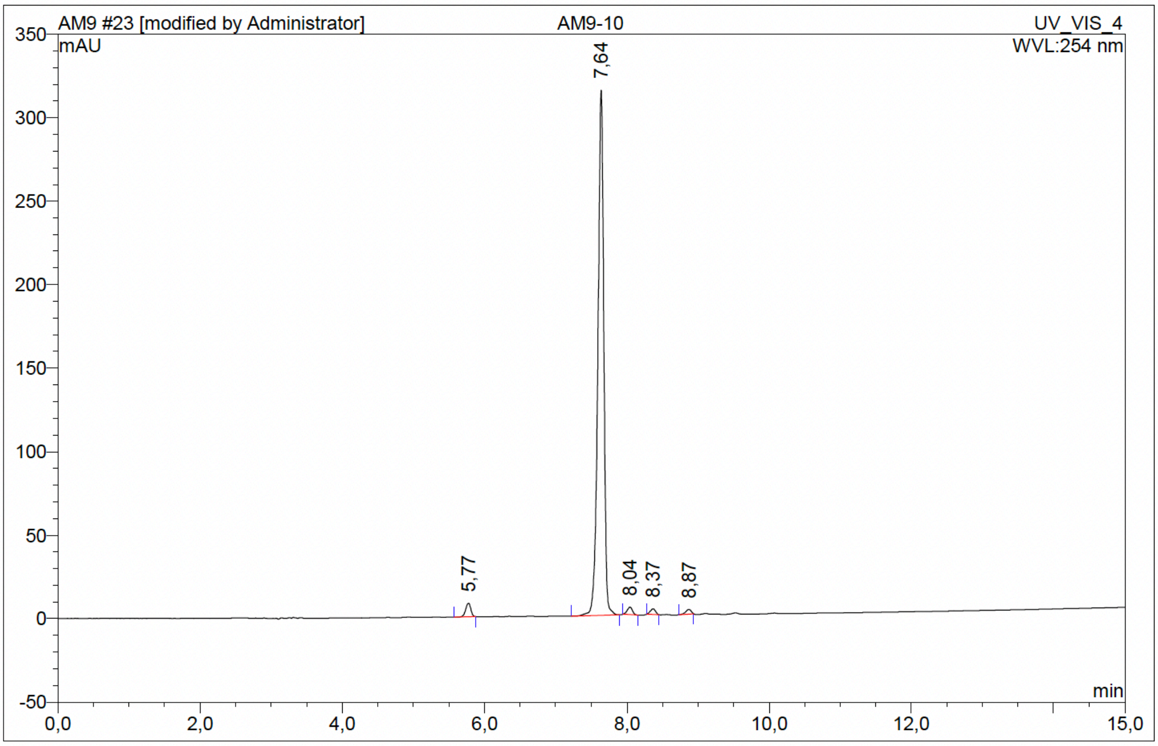

| No. | Ret.Time min | Peak Name | Height mAU | Area mAU*min | Rel.Area % | Amount | Resolution(EP) |
|---|---|---|---|---|---|---|---|
| 1 | 5,77 | n.a. | 8,096 | 0,661 | 2,09 | n.a. | 13,63 |
| 2 | 7,64 | n.a. | 314,611 | 30,010 | 95,00 | n.a. | 2,87 |
| 3 | 8,04 | n.a. | 4,513 | 0,389 | 1,23 | n.a. | 2,34 |
| 4 | 8,37 | n.a. | 3,363 | 0,281 | 0,89 | n.a. | 3,67 |
| 5 | 8,87 | n.a. | 2,825 | 0,247 | 0,78 | n.a. | n.a. |
| Total: | | | 333,407 | 31,588 | 100,00 | 0,000 | |

Compound 69 - TUG-2843

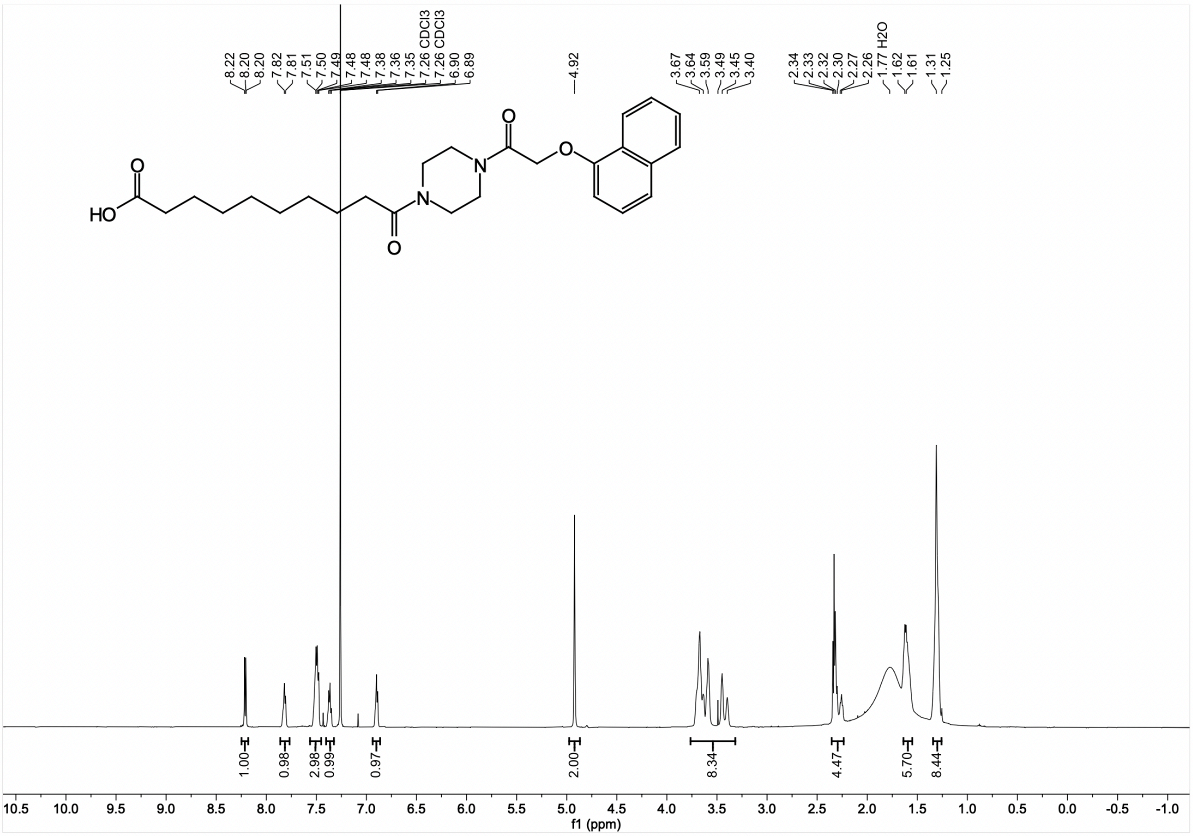

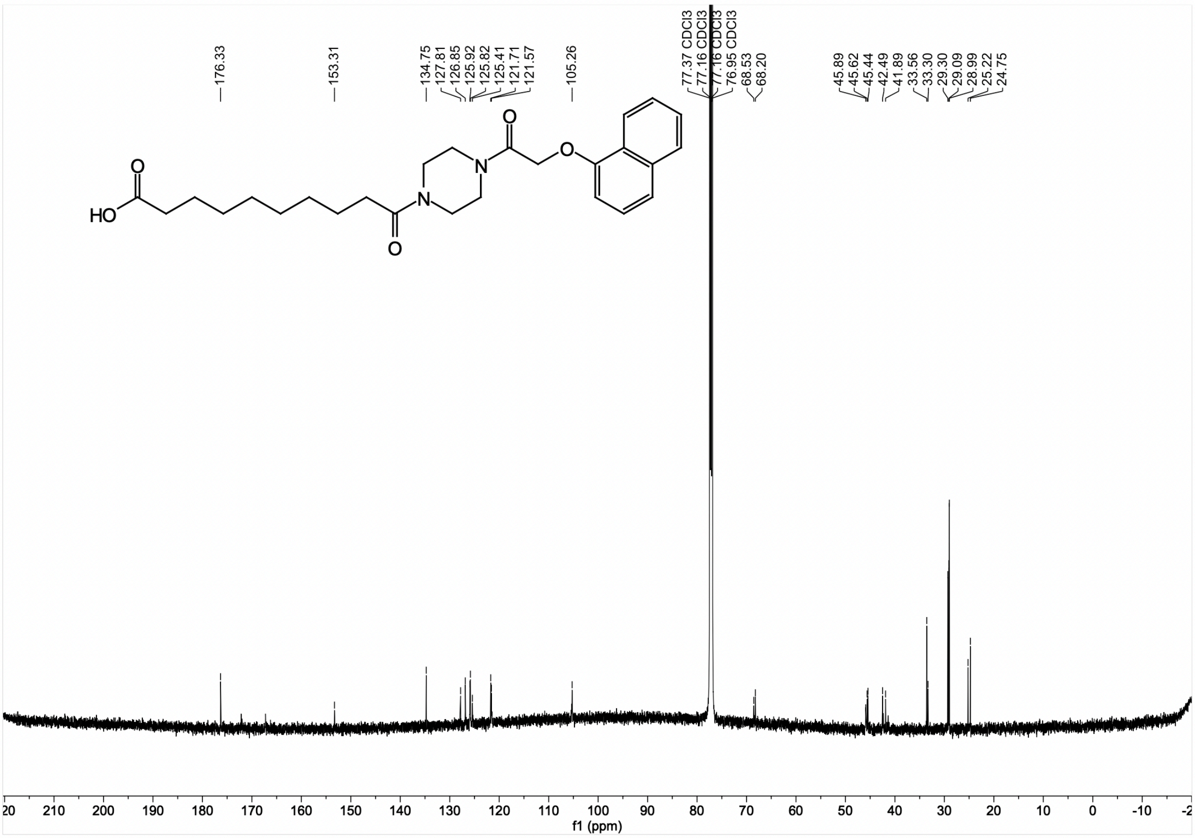

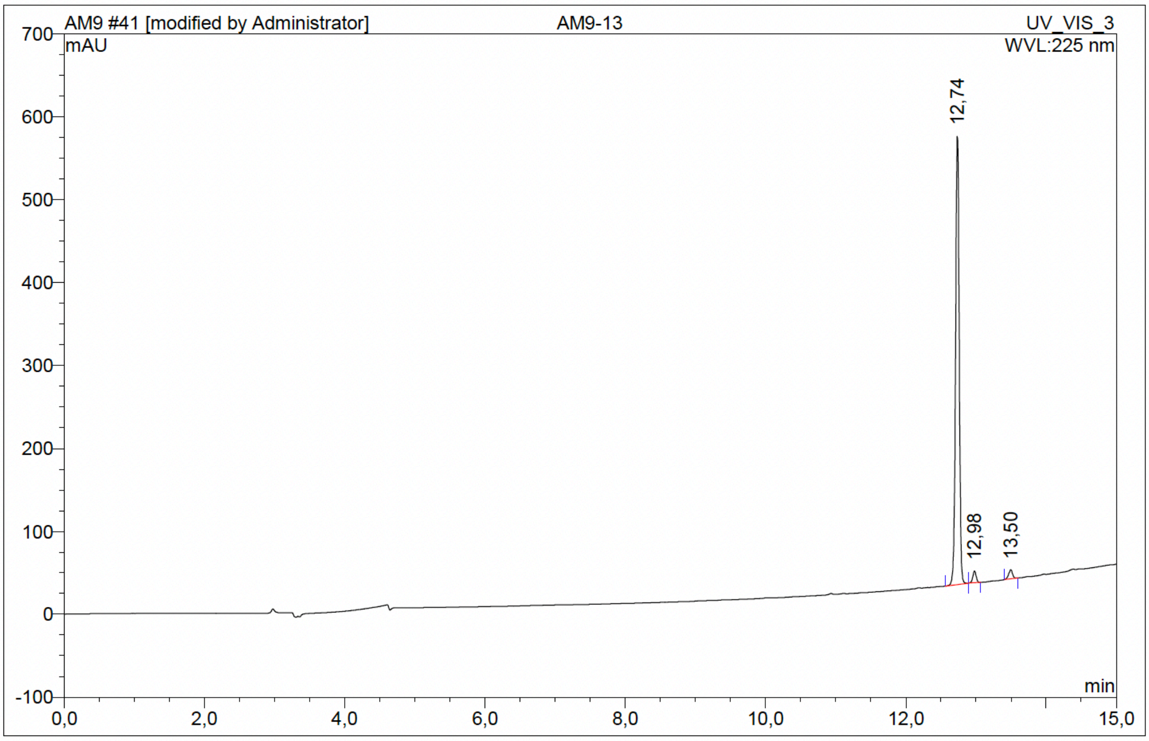

| No. | Ret.Time min | Peak Name | Height mAU | Area mAU*min | Rel.Area % | Amount | Resolution(EP) |
|---|---|---|---|---|---|---|---|
| 1 | 12,74 | n.a. | 541,069 | 30,998 | 95,49 | n.a. | 2,78 |
| 2 | 12,98 | n.a. | 14,108 | 0,785 | 2,42 | n.a. | 5,64 |
| 3 | 13,50 | n.a. | 10,955 | 0,679 | 2,09 | n.a. | n.a. |
| Total: | | | 566,133 | 32,462 | 100,00 | 0,000 | |

Compound 70 - TUG-2392

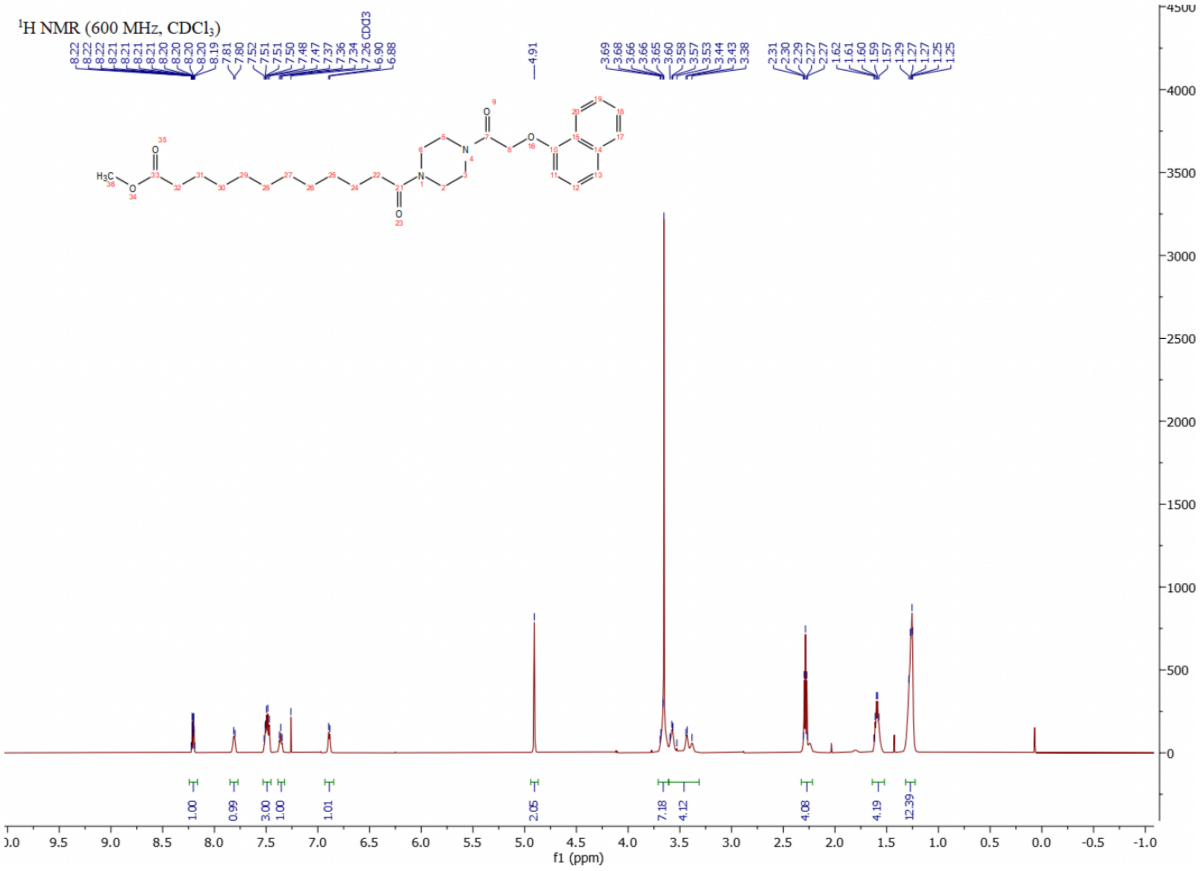

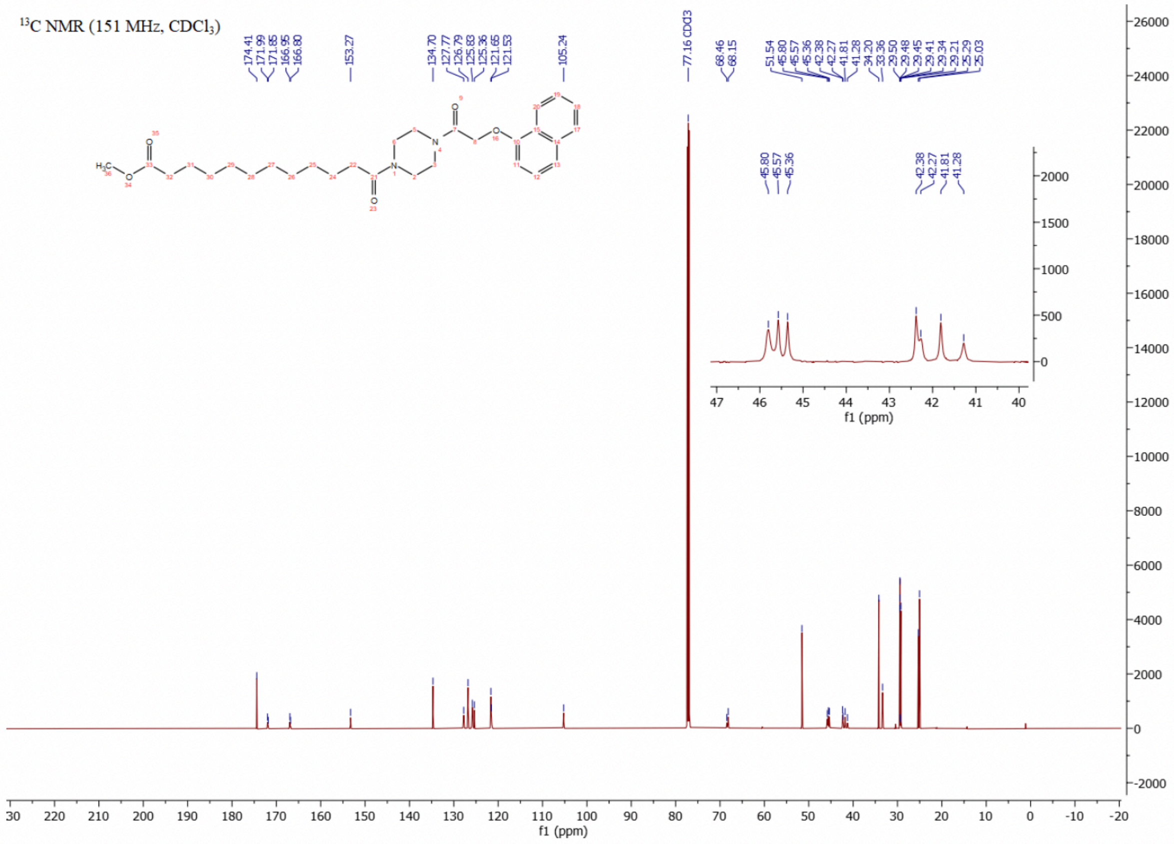

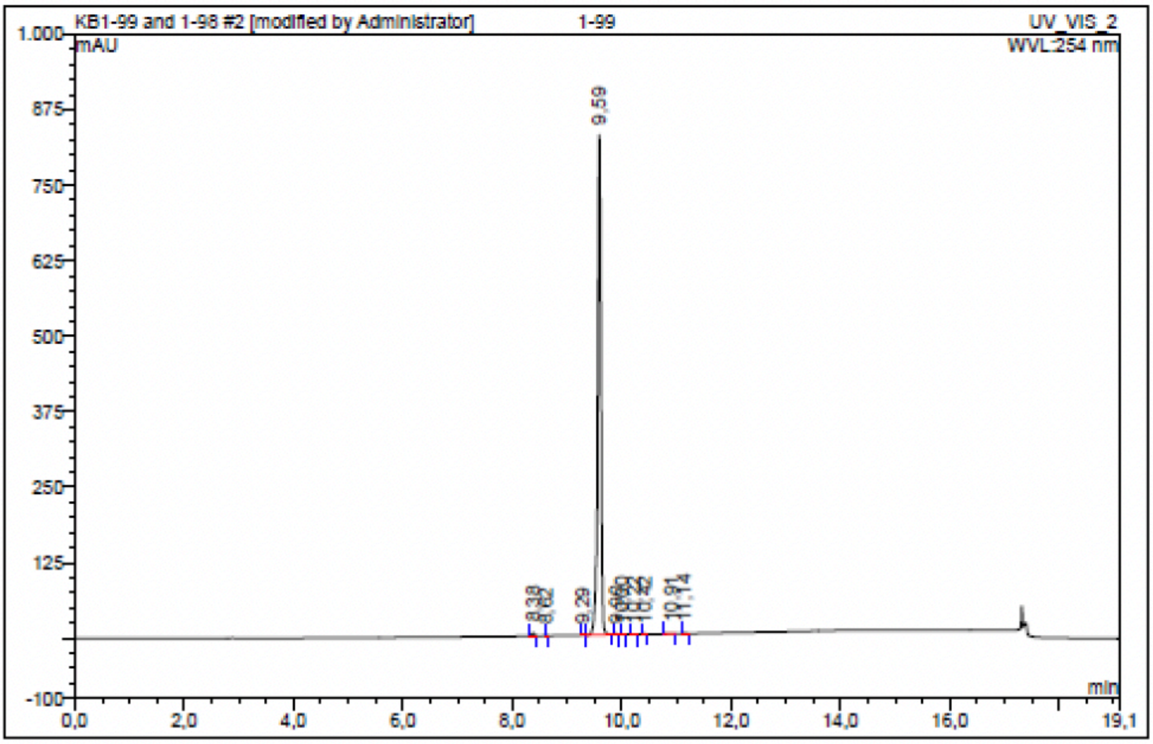

| No. | Ret.Time min | Peak Name | Height mAU | Area mAU*min | Rel.Area % | Amount | Resolution(EP) |
|---|---|---|---|---|---|---|---|
| 1 | 8,38 | n.a. | 4,331 | 0,242 | 0,42 | n.a. | 2,98 |
| 2 | 8,62 | n.a. | 0,641 | 0,025 | 0,04 | n.a. | 8,70 |
| 3 | 9,29 | n.a. | 0,808 | 0,043 | 0,07 | n.a. | 3,17 |
| 4 | 9,59 | n.a. | 827,603 | 57,146 | 99,06 | n.a. | 3,23 |
| 5 | 9,90 | n.a. | 0,833 | 0,044 | 0,08 | n.a. | 1,31 |
| 6 | 10,00 | n.a. | 0,224 | 0,007 | 0,01 | n.a. | 2,43 |
| 7 | 10,22 | n.a. | 0,668 | 0,040 | 0,07 | n.a. | 2,28 |
| 8 | 10,42 | n.a. | 0,645 | 0,028 | 0,05 | n.a. | 4,74 |
| 9 | 10,91 | n.a. | 0,580 | 0,054 | 0,09 | n.a. | 2,03 |
| 10 | 11,14 | n.a. | 0,986 | 0,062 | 0,11 | n.a. | n.a. |
| Total: | | | 837,319 | 57,691 | 100,00 | 0,000 | |

Compound 71 - TUG-2393

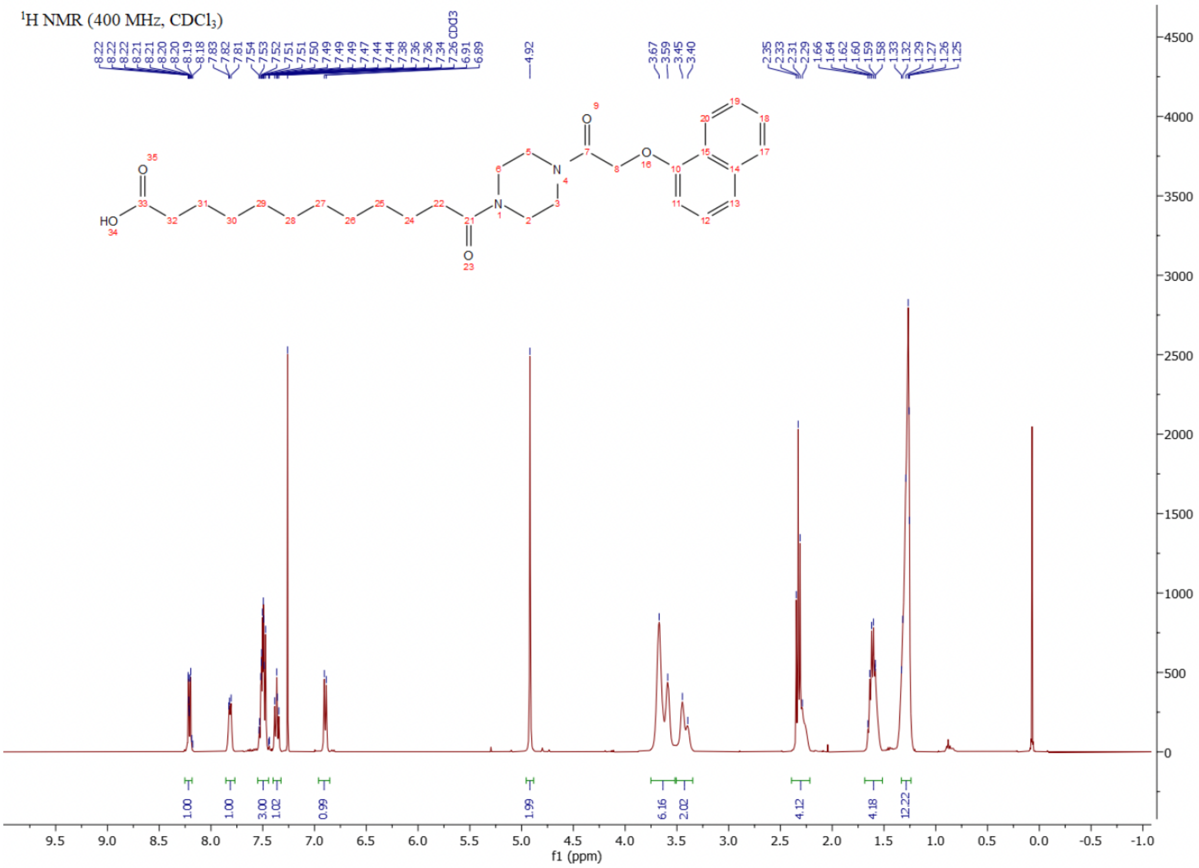

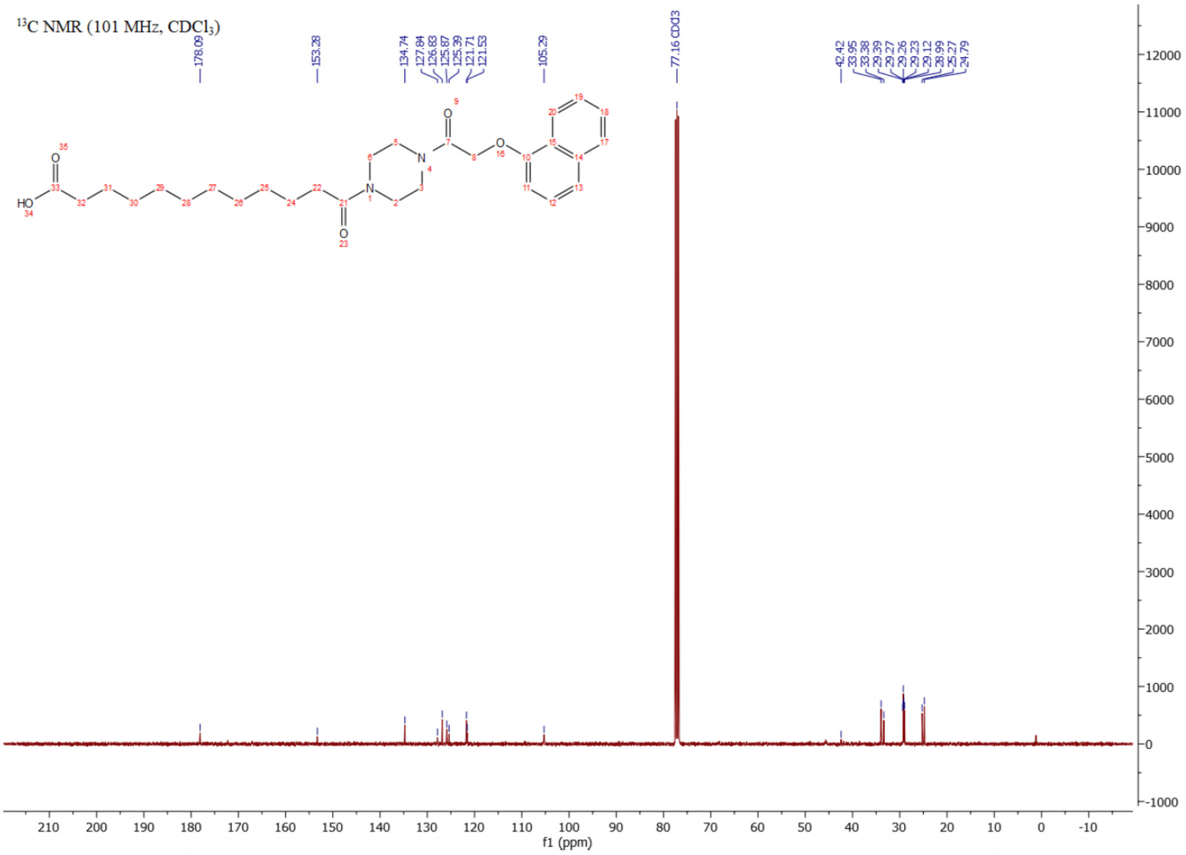

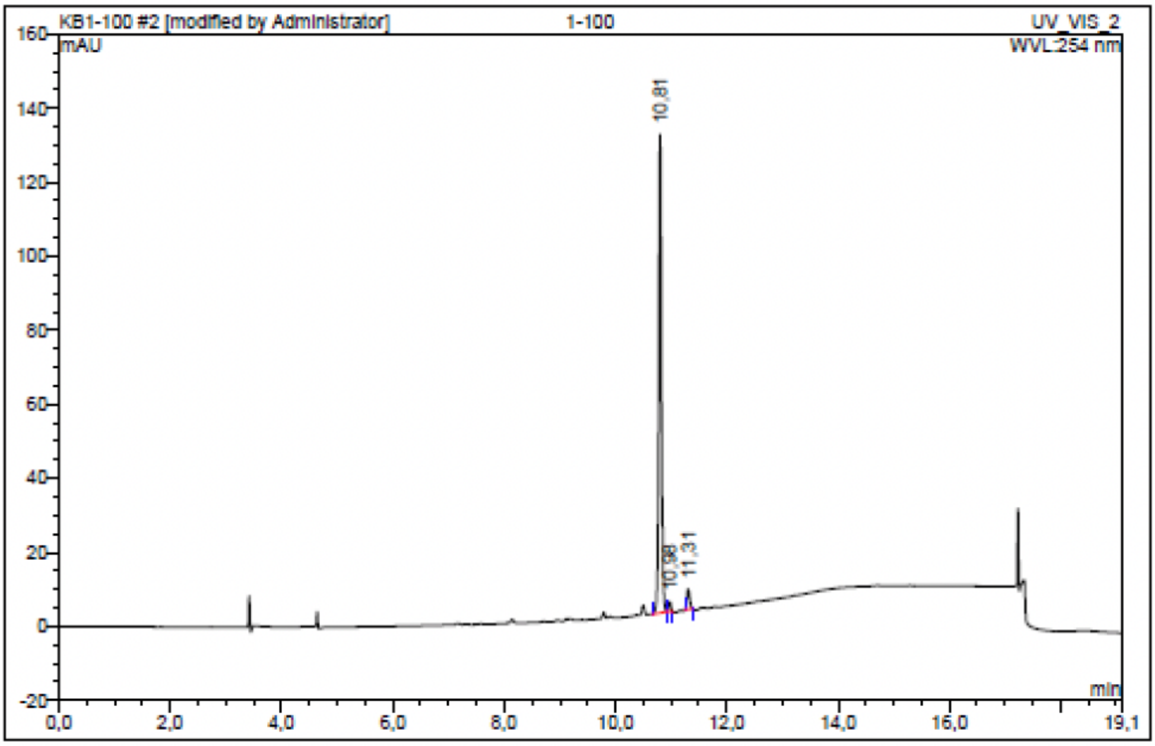

| No. | Ret.Time min | Peak Name | Height mAU | Area mAU*min | Rel.Area % | Amount | Resolution(EP) |
|---|---|---|---|---|---|---|---|
| 1 | 10,81 | n.a. | 129,228 | 7,189 | 94,51 | n.a. | 2,15 |
| 2 | 10,98 | n.a. | 2,577 | 0,121 | 1,59 | n.a. | 3,99 |
| 3 | 11,31 | n.a. | 5,432 | 0,297 | 3,90 | n.a. | n.a. |
| Total: | | | 137,236 | 7,606 | 100,00 | 0,000 | |
